# A high-coverage genome from a 200,000-year-old Denisovan

**DOI:** 10.1101/2025.10.20.683404

**Authors:** Stéphane Peyrégne, Diyendo Massilani, Yaniv Swiel, Michael James Boyle, Leonardo N. M. Iasi, Arev P. Sümer, Alba Bossoms Mesa, Cesare de Filippo, Bence Viola, Elena Essel, Sarah Nagel, Julia Richter, Antje Weihmann, Barbara Schellbach, Hugo Zeberg, Johann Visagie, Maxim B. Kozlikin, Michael V. Shunkov, Anatoly P. Derevianko, Kay Prüfer, Benjamin M. Peter, Matthias Meyer, Svante Pääbo, Janet Kelso

**Author notes:** Corresponding authors Correspondence to Stéphane Peyrégne or Janet Kelso.

## Abstract

Denisovans, an extinct sister group of Neandertals who lived in Eastern Eurasia during the Middle and Late Pleistocene, are known only from a handful of skeletal remains and limited genetic data, including the high-coverage genome of a woman who lived ∼65,000 years ago. Here, we present a second high-quality Denisovan genome, reconstructed from a molar found at Denisova Cave. It belonged to a man who lived ∼200,000 years ago in a small Denisovan group. This group mixed with early Neandertals and was then replaced by Denisovans who had mixed with later Neandertals. We show that in addition Denisovans received gene flow from hominins that diverged before the split of the ancestors of Denisovans and modern humans. The two Denisovan genomes allow us to disentangle Denisovan ancestry in present-day humans revealing contributions from at least three distinct Denisovan groups. In particular, Oceanians and South Asians independently inherited DNA from a deeply diverged Denisovan population which was likely isolated in South Asia. This supports an early migration of the ancestors of Oceanians through South Asia followed by the later arrival of the ancestors of present-day South Asians. East Asians do not share this Denisovan component in their genomes, suggesting that their ancestors arrived independently, perhaps by a northerly route. Finally, the two high-quality Denisovan genomes allow us to refine the catalogue of genetic changes that arose on the Denisovan lineage, some of which were contributed to present-day humans.

## Main text

Denisovans, an extinct human group, were first identified based on ancient DNA extracted from *Denisova 3*, a finger phalanx discovered at Denisova Cave in the Altai Mountains of Siberia in 2008^1,2^. Analysis of the nuclear genome from this individual revealed that Denisovans were a sister group to Neandertals^2^, another group of now extinct humans who lived in Western Eurasia in the middle and late Pleistocene. While twelve fragmentary remains and one cranium have since been attributed to Denisovans based on DNA or protein analysis^1-12^, only *Denisova 3* has yielded a high-quality genome (∼30-fold genomic coverage)^13^.

The Denisovan genomic data available from Denisova Cave, which also includes three partial nuclear genomes, seven mitochondrial genomes, and Denisovan DNA recovered from sediment samples, indicate recurrent Denisovan presence in the Altai region between 200 and 50 thousand years ago (ka)^8,14,15^, interspersed with periods of Neandertal occupation. The identification of a child of a Neandertal mother and a Denisovan father shows that these groups sometimes overlapped and interbred^4^. However, Denisovans remained genetically distinct from Neandertals, perhaps because these events were limited to a few contact zones. Mitochondrial DNA analyses at Denisova Cave and Baishiya Karst Cave on the Tibetan Plateau^16^ suggest that different Denisovan populations at these locations at times replaced each other.

Comparisons of the genomes of *Denisova 3* and modern humans revealed that interbreeding between Denisovans and the ancestors of present-day people took place in Asia and possibly Oceania^2,13,17-20^. Evidence of multiple Denisovan components with different relationships to the sequenced Denisovan in the genomes of present-day people indicates mixture with genetically distinct Denisovan populations^20-25^, about whom we know very little. Additional Denisovan genomes would be helpful to better understand Denisovan population history, the relationships among Denisovan ancestry components in present-day people, and the biology of this now-extinct human group.

### A new Denisovan genome sequence

In 2020, a complete left upper molar was found in layer 17, one of the lowest cultural layers of the South Chamber of Denisova Cave, dated to 200-170ka by optically stimulated luminescence^15^. Designated as *Denisova 25*, this molar is similar in size to the other molars discovered at Denisova Cave, *Denisova 4* and *Denisova 8*^2,3^, and larger than those of Neandertals as well as most other Middle Pleistocene and later hominins (**Supplementary Note 1, Figures S3** and **S4**), suggesting that it potentially belonged to a Denisovan. Yet, *Denisova 25* shows a simpler morphology than *Denisova 4* and *Denisova 8*, with only three main cusps and lacking the very robust and strongly diverging roots seen in *Denisova 4*. Its outline is most similar to that of an upper molar preserved in the Harbin cranium from northeastern China, dated to at least 146ka^26^. Two samples of 2.7 mg and 8.9 mg were removed by drilling one hole at the cemento-enamel junction of the tooth, and twelve subsamples, ranging from 4.5 mg to 20.2 mg, were obtained by gently scratching the outer layer of one of the roots with a dentistry drill (**Supplementary Note 2**). After DNA extraction and library preparation, we identified two samples with exceptional ancient DNA preservation. Notably, 20 to 47% of DNA sequences generated mapped to the human genome and less than ∼3% of these sequences stemmed from present-day human DNA contamination (**Figure 1**). We sequenced these libraries more deeply and reconstructed a 23.6-fold coverage of the genome of this individual (**Supplementary Note 3**). As the coverage of the X and Y chromosomes is half that of the autosomes, we concluded that the tooth belonged to a male individual (**Figure 1**; **Supplementary Note 3**).

**Figure 1.**
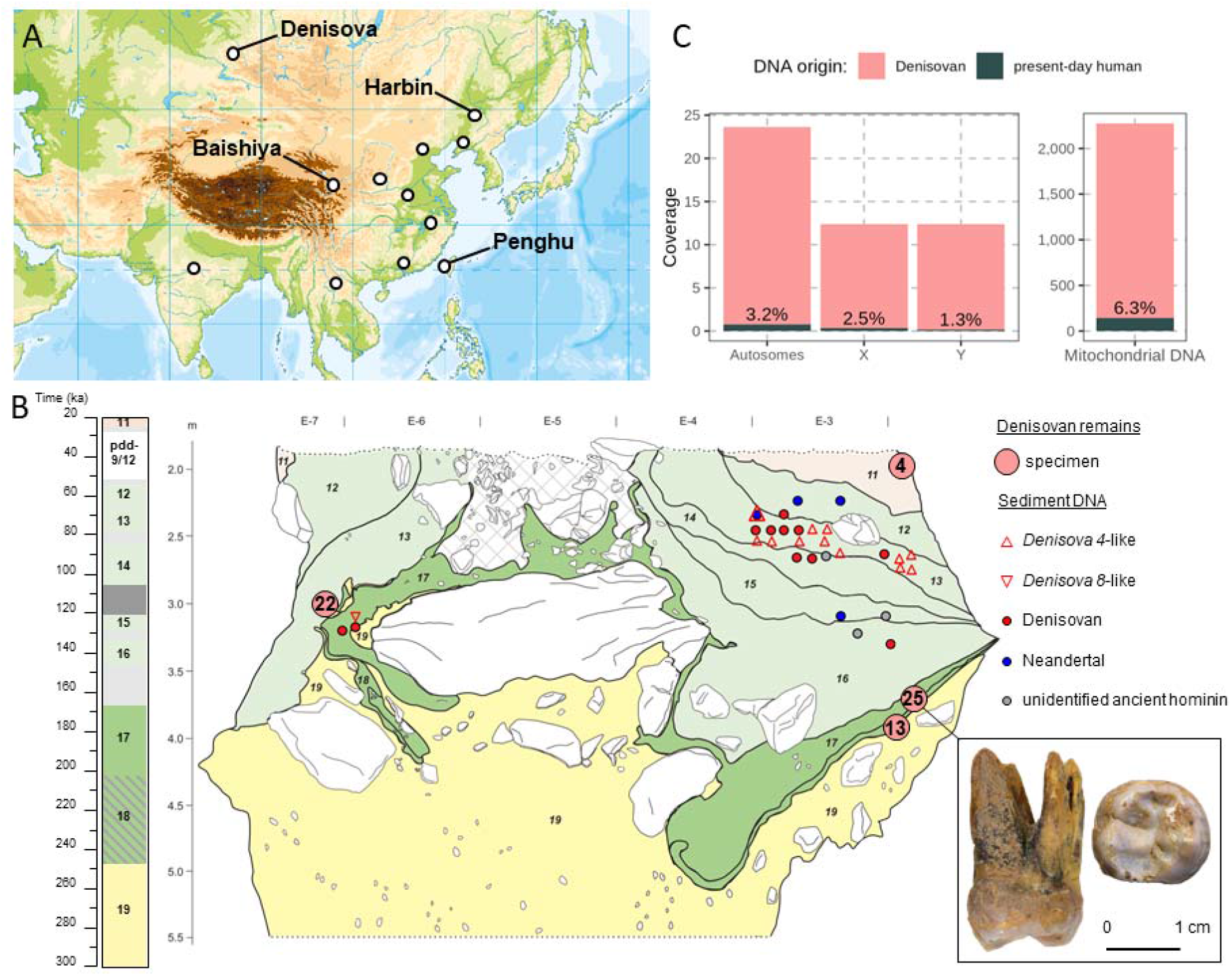
Origin of *Denisova 25* and available genetic data. (**A**) Map of Asia showing the locations of Denisova Cave, Baishiya Cave, the Harbin cranium and Penghu mandible discoveries, and other sites with potential Denisovan remains^27-29^. (**B**) Stratigraphic profile of the South Chamber in Denisova Cave, illustrating the approximate origins of *Denisova 25* and other Denisovan remains (specimen numbers circled in the corresponding layers). A composite section (modified from^15^) to the left of the profile displays modelled start and end ages (in ka) for sedimentary layers (95% highest posterior probability). Circles denote previously described sediment sample locations, coloured by detected hominin mitochondrial DNA: red (Denisovan), blue (Neandertal), grey (unidentified ancient hominin), and white (no ancient hominins detected). Triangles indicate sediment samples assigned to a specific Denisovan mitochondrial lineage. Distal and occlusal views of *Denisova 25* are shown. (**C**) Genomic coverage and proportion of present-day human DNA contamination for *Denisova 25*.

### Molecular dating

The mitochondrial genome and Y chromosome of *Denisova 25* are of the Denisovan type (**Figures 2B** and **2C**; **Supplementary Notes 4** and **5**) and are more closely related to the mitochondrial genome and Y chromosome of *Denisova 8*, a Denisovan previously dated to 136-106ka, than those of *Denisova 4* dated to 84-55ka^30^. Molecular dating using a Bayesian analysis^31^ of the mitochondrial genome and the Y chromosome (restricted to regions that facilitate accurate dating^32^) results in age estimates for the *Denisova 25* individual of 219ka (95% Highest Posterior Density Interval, HPDI: 275-159ka) and 232ka (95% HPDI: 288-177ka), respectively. Given that the mitochondrial lineage of *Denisova 8* is longer than that of *Denisova 25* by at most two mutations, we estimate that the *Denisova 8* individual lived approximately 215ka (95% HPDI: 273-158ka), which is consistent with the assignment of *Denisova 8* to a deeper layer (dated to ∼190ka) in a recent re-evaluation of the stratigraphy of Denisova Cave^15^. The number of missing mutations in the nuclear genome of *Denisova 25* relative to present-day humans yields a molecular date of 205ka (215ka-195ka, 2 s.d.; **Supplementary Note 6**). The mitochondrial, Y chromosome and nuclear estimates are thus consistent with the age of layer 17 where the molar was found. *Denisova 25* is therefore about 140,000 years older than the previously sequenced *Denisova 3* (75-55ka, 2 s.d.; **Supplementary Note 6**).

**Figure 2.**
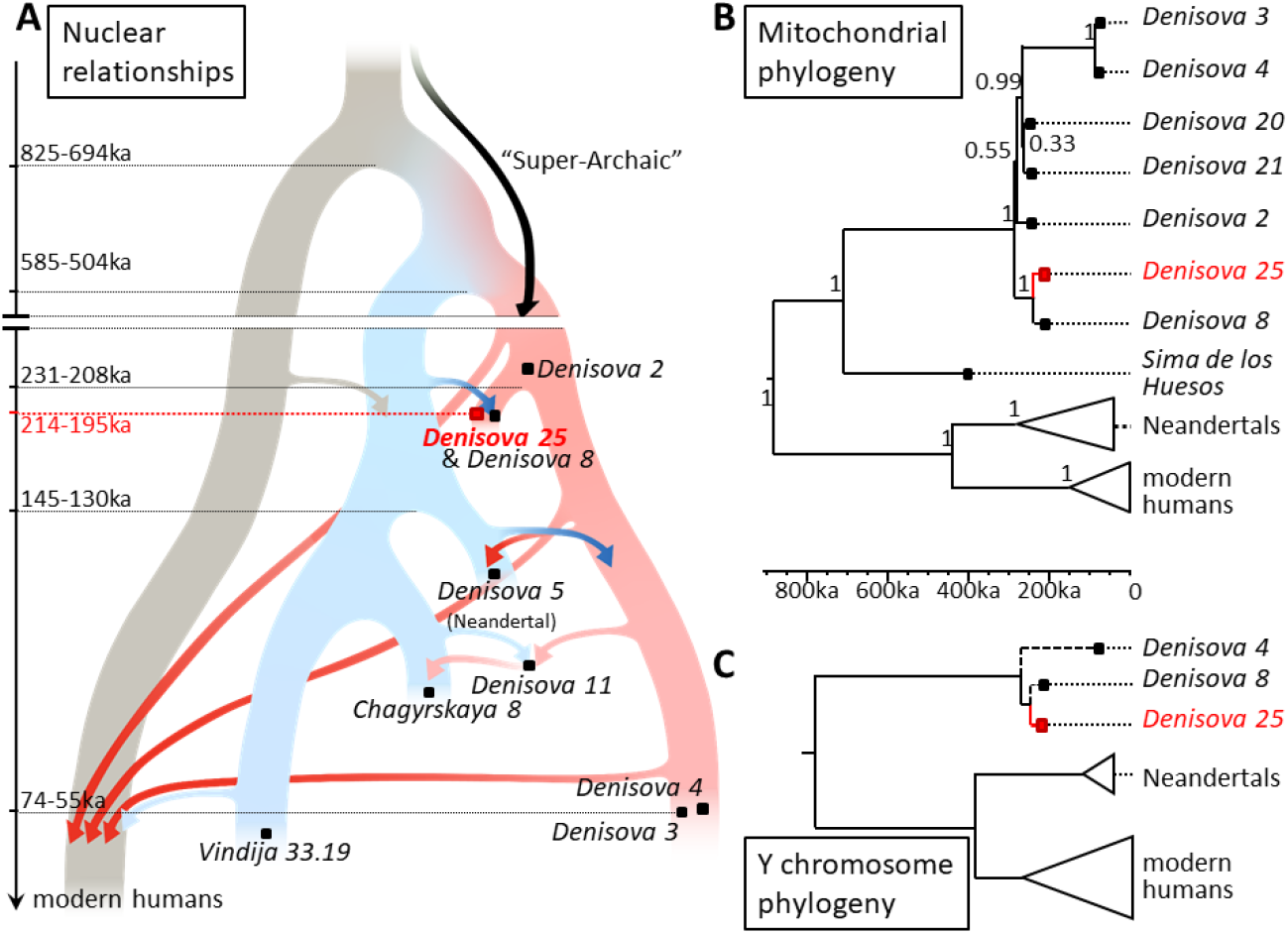
Relationships of *Denisova 25* to archaic and modern humans. (**A**) Tree representing average population relationships inferred from nuclear genomes. Arrows indicate gene flow between populations, with darker arrows denoting those identified or examined in this study. Age estimates for high-coverage genomes are inferred from branch shortening (i.e., missing mutations in archaic genomes), with intervals corresponding to two standard deviations from the estimate as computed with a block jackknife. Population split time estimates are indicated by dotted lines. The split between the populations of *Denisova 3* and *Denisova 25* is estimated at 15,000 years (±2,000 years) before *Denisova 25* lived, adjusted for the uncertainty in the age of *Denisova 25*. Neandertal-Denisovan and modern-archaic split times correspond to the range of estimates obtained with calibrations of the divergence in genomic regions that presumably evolve neutrally. The split between the population of the *Denisova 5* Neandertal and other Neandertals was previously estimated^33^. The tree is not to scale. (**B**) Bayesian phylogenetic tree of mitochondrial genomes of 7 Denisovans, 27 Neandertals, 64 modern humans and a hominin from Sima de los Huesos. The numbers at the nodes correspond to the posterior probabilities of the depicted branching orders. The Denisovan mitochondrial sequence of *Denisova 19* is not included because it is identical to that of *Denisova 21*^8^. (**C**) Bayesian phylogenetic tree of the Y chromosomes of 3 Denisovans, 2 Neandertals, and 32 modern humans. The Y chromosomes of *Denisova 4* and *Denisova 8* were excluded from the analysis due to the lower coverage available, but positioned on the tree based on the sharing of derived alleles with the Y chromosome of *Denisova 25*. The length of the branches for *Denisova 4* and *Denisova 8* is inferred from the age of these individuals on the mitochondrial tree. The scale for the mitochondrial and Y chromosome phylogenies is indicated. Throughout the figure, Denisovans, Neandertals and modern humans are highlighted in light red, light blue, and light yellow, respectively.

### Relationships to other hominins

We next compared the derived allele sharing between the high-quality genomes of *Denisova 25*, the *Denisova 3* Denisovan and three Neandertals, as well as the lower-quality genomes from four additional Denisovans (**Supplementary Note 8**). These comparisons show that *Denisova 25* is more closely related to *Denisova 3* than to Neandertals, indicating that he belonged to an early Denisovan population (**Figure 2A**). Furthermore, *Denisova 8* shares more derived alleles with *Denisova 25* than with *Denisova 3*, whereas *Denisova 4* and *Denisova 11* share more alleles with *Denisova 3* (**Supplementary Note 8**). This is consistent with the mitochondrial and Y chromosome relationships and supports a temporal transition between two distinct Denisovan populations at Denisova Cave. The partial genome of *Denisova 2*, a deciduous molar found in the deepest layer of the Main Chamber, is equally related to *Denisova 3* and *Denisova 25*, which could suggest the existence of a third population, but the high level of present-day human DNA contamination in the *Denisova 2* genome may obscure the relationships.

*Denisova 25* shares more alleles with Neandertals than *Denisova 3* does (**Supplementary Note 8**), suggesting that *Denisova 25* received more Neandertal gene flow than *Denisova 3*, who has been estimated to carry approximately 0.5% DNA from Neandertals more closely related to the 120,000-year-old Neandertal *Denisova 5* than later Neandertals^33,34^. In addition, the Neandertal *Denisova 5* shares more alleles with both *Denisova 3* and *Denisova 25* than the *Vindija 33*.*19* and *Chagyrskaya 8* Neandertals do, who appear equally related to Denisovans (**Supplementary Note 8**). This difference between *Denisova 5* and the other Neandertals in their relationship to *Denisova 25* cannot be due to gene flow from the Neandertal *Denisova 5* population to the *Denisova 25* population as *Denisova 25* predates the existence of the *Denisova 5* population, which splits from the other Neandertal populations around 140ka^33,34^. The explanation must therefore be gene flow from Denisovans to the ancestors of the Neandertal population to which *Denisova 5* belonged.

### Recurrent Neandertal gene flow into Denisovans

Using a method^35^ that leverages reference genomes to identify local ancestry along the Denisovan genomes, we estimated that *Denisova 25* and *Denisova 3* carry 3.6-5.2% and 1.8-2.5% Neandertal ancestry, respectively, and identified between 56Mb and 166Mb of Neandertal segments in *Denisova 25*, and between 34Mb and 87Mb in *Denisova 3*, depending on the parameters used for calling segments (**Figure 3**; **Supplementary Note 9**). While the Neandertal ancestry of *Denisova 3* is more closely related to the *Denisova 5* Neandertal than to the two other Neandertals, the Neandertal ancestry of *Denisova 25* is equally related to all three Neandertal genomes. Given the age of *Denisova 25*, this suggests that his Neandertal ancestry comes from a yet undescribed Neandertal population that split over 200,000 years ago from the Neandertal populations for which we have genomes. Moreover, the length distribution of their Neandertal segments fits better to a mixture of exponential distributions than a single exponential decay (Figure 3), suggesting that both Denisovans inherited their Neandertal ancestry from multiple gene flow events. We estimated that the most recent Neandertal ancestors of *Denisova 25* predate him by an average of 13,000-7,000 years, and that his ancestors may also have mixed with even earlier Neandertals (**Supplementary Note 9**). Similarly, the ancestors of *Denisova 3* mixed with Neandertals on multiple occasions, with both recent gene flow 16,000 to 12,000 years before she lived, as well as older gene flow. This indicates repeated contacts between Neandertals and Denisovans throughout their history (**Supplementary Note 9**). Despite this recurrent mixing, distinct Neandertal and Denisovan groups can be identified at Denisova Cave from around 200,000 years ago until around 70,000 years ago. This suggests that mixed individuals did not contribute much to the gene pool of later individuals, which is consistent with archaic population turnovers in the Denisova Cave region. It is possible that mixing between Neandertals and Denisovans occurred locally in the region of Denisova Cave at the edges of their respective geographical ranges without spreading further. Recurrent migrations from elsewhere then replaced these mixed groups with Denisovans and Neandertals with less mixed ancestry.

**Figure 3.**
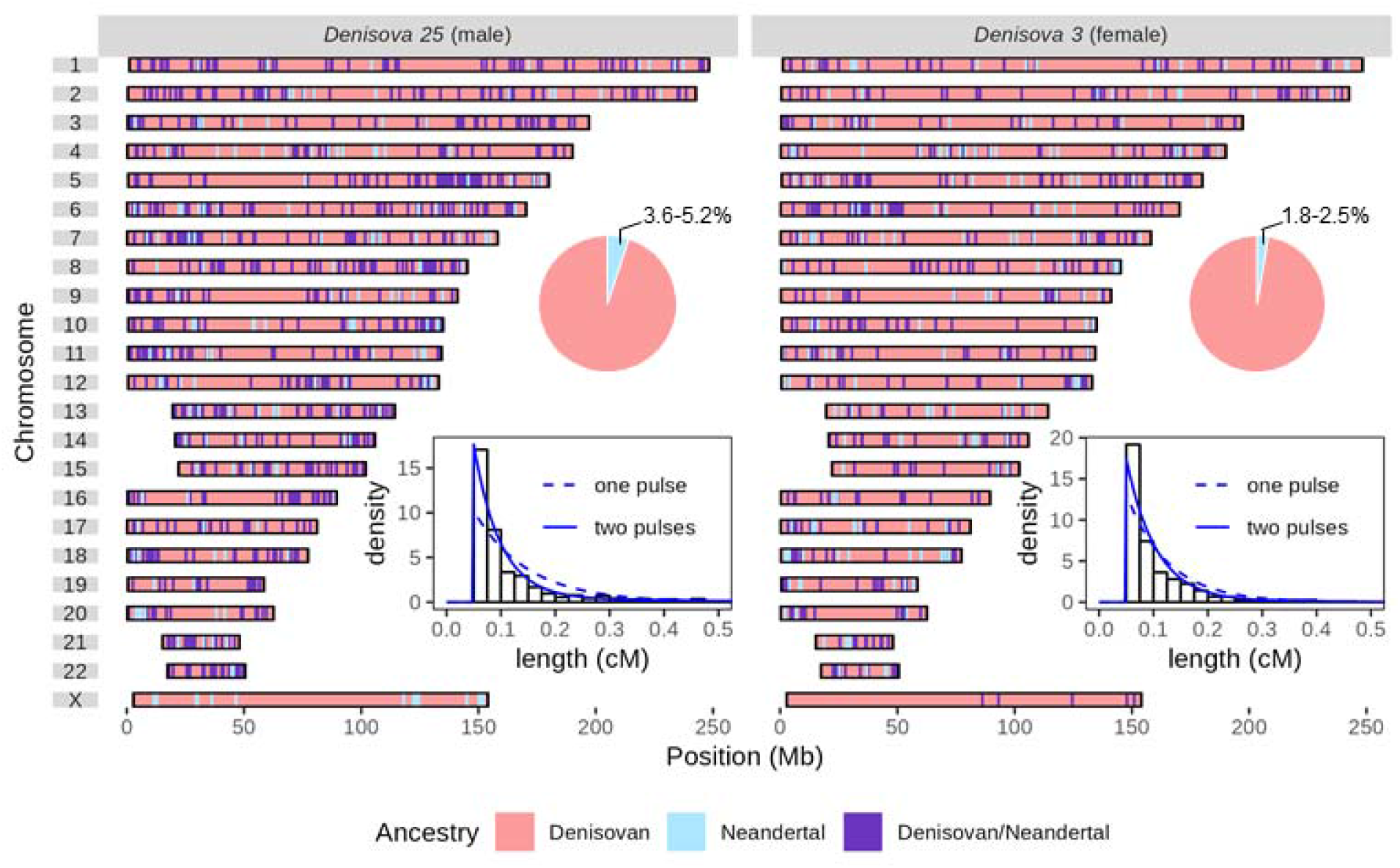
Distribution of Neandertal ancestry in Denisovan genomes. Colours indicate the inferred local ancestry along the two copies of each chromosome (or the single X chromosome of *Denisova 25*). Neandertal DNA segments of at least 0.05cM are shown. Pie charts depict the total proportion of Neandertal DNA identified in each genome. The length distribution of Neandertal DNA segments (in centimorgans, cM) is presented for each genome as histograms, alongside fitted distributions modeling one or two pulses of admixture with Neandertals. The two-pulse model provides a significantly better fit than the one-pulse model for both genomes (likelihood ratio test p-values: <10^-6^ for *Denisova 3* and <10^-23^ for *Denisova 25*).

### Revised population split times

The amount of Neandertal DNA in these Denisovan genomes could affect population split time estimates. We show with simulations that a previously used statistic (*F(A*|*B)*, which measures the extent to which individuals share genetic polymorphisms^13,34,36^, is highly sensitive to gene flow, and likely under-estimates the true split times, while split time estimates based on sequence divergence are less biased (**Supplementary Note 8**). In addition, background selection has a large effect on demographic inferences^37-39^. To account for this, we masked regions exhibiting a reduced genetic diversity in present-day humans, a signal indicative of selective constraints^40^. When restricting the analysis to neutrally evolving regions and using sequence divergence, a statistic less sensitive to gene flow, we estimate that modern and archaic humans separated 825-694ka, while Neandertals and Denisovans separated 585-504ka. These dates are substantially older than previous estimates (630-520ka and 440-390ka, respectively^33^) and consistent with the time to the most recent common ancestor between Denisovan and modern human mitochondrial genome and Y chromosome (**Figure 2B and 2C**). The revised dates are also consistent with estimates based on dental evolutionary rates^41^ and the presence of derived Neandertal dental and mandibular traits in the early Neandertals from Sima de los Huesos^42-44^. Finally, we estimate that the ancestors of *Denisova 25* and *Denisova 3* separated 15,000 (±2,000) years before *Denisova 25* lived. Considering the uncertainty in the age of *Denisova 25*, this results in a split time of 231-204ka (**Supplementary Note 8**). We caution that the split time estimates, particularly those between archaic and modern humans, remain approximate; in particular, gene flow from more divergent groups into modern humans^45,46^ or Denisovans^33,34^ could inflate our estimates by tens, but not hundreds, of thousands of years (**Supplementary Figure S45**; **Supplementary Note 8**).

### Genetic diversity and population sizes of early and late Denisovans

As *Denisova 25* lived about fifteen thousand years after the divergence from the ancestral population with *Denisova 3*, we investigated the demographic history of the *Denisova 25* population (Supplementary Note S7). Using the Pairwise Sequential Markovian Coalescent model, we detected a decline in population size approximately 220,000 years ago, following the split with the ancestors of *Denisova 3* (**Figure 4A**). Additionally, about 15.8% of the *Denisova 25* genome are present in homozygous regions longer than 2.5cM, indicating a history of consanguinity among the ancestors of *Denisova 25* (**Figure 4B**). In particular, the fraction of the *Denisova 25* genome covered by homozygous regions longer than 10cM (8.9%) is similar to that in the *Denisova 5* (13%) and *Chagyrskaya 8* (8.3%) Neandertal genomes, compatible with various scenarios of recent consanguinity, including parents that were double first cousins to half-siblings^34^. Despite these long runs of homozygosity, *Denisova 25* exhibits higher heterozygosity than other archaic genomes. However, if the regions derived from Neandertals are excluded in the *Denisova 3* and *Denisova 25* genomes, their heterozygosity becomes similar to the Neandertal genomes, showing that the apparent differences in heterozygosity stem from the presence of Neandertal ancestry in the Denisovans (**Figure 4C**).

**Figure 4.**
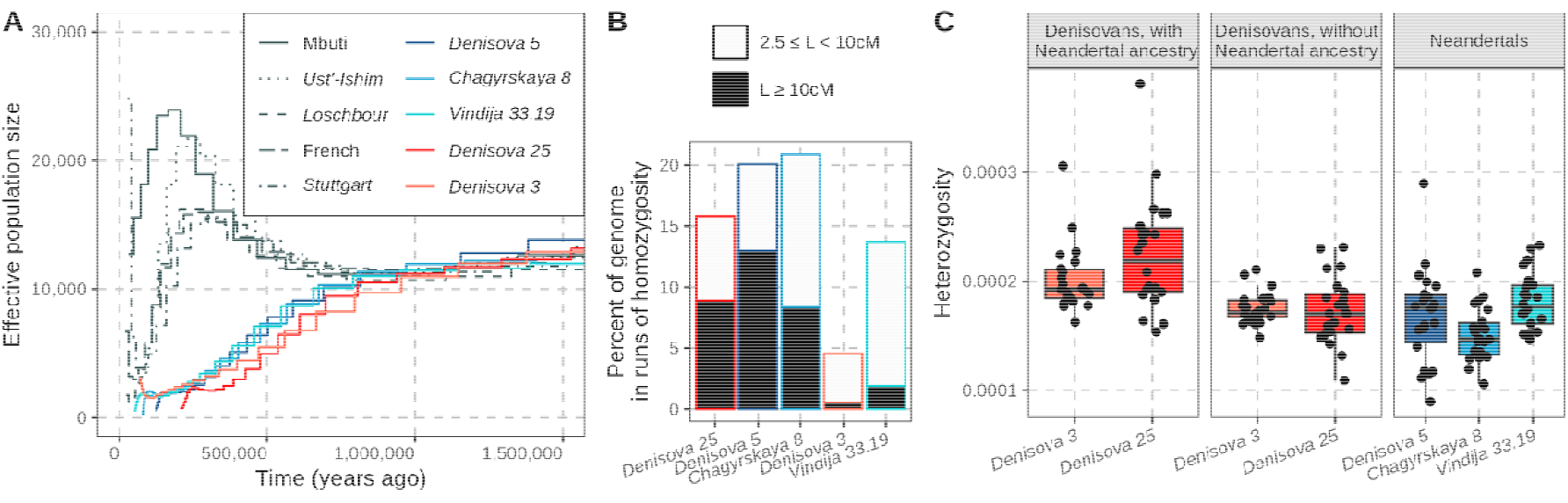
Population size, runs of homozygosity, and heterozygosity of *Denisova 25*. (**A**) Population size estimates over time based on the Pairwise Sequential Markovian Coalescent model, calculated for presumably neutrally evolving genomic regions (B-score^40^≥0.9). (**B**) Percentage of the genome in runs of homozygosity (ROH) of at least 2.5cM in high-coverage archaic genomes, assuming a uniform recombination rate of 1.3cM/Mb. Black bars indicate the proportion of the genome in ROH of at least 10cM, suggesting close parental relatedness. (**C**) Heterozygosity estimates in high-coverage archaic human genomes. For comparison, Neandertal segments (identified with *admixfrog*, **Supplementary Note 9**) were masked in the Denisovan genomes. Each dot represents the heterozygosity for a chromosome, while diamonds denote the average across all autosomes.

Using an approach which estimates group size from the length and number of runs of homozygosity^47^, we estimated that the *Denisova 25* lived in a group of 50-60 individuals, with approximately one migrant per generation, similar to the estimates for the 80,000-year-old *Chagyrskaya 8* Neandertal genome and about twice that of the 120,000-year-old *Denisova 5* Neandertal genome (**Supplementary Note 7**). In contrast, the 60,000-year-old *Denisova 3* Denisovan and the 50,000-year-old *Vindija 33*.*19* Neandertal likely lived in larger groups (>100 individuals), suggesting that group sizes were not inherently different between Neandertals and Denisovans. The small size of the early groups and the population turnovers suggest that conditions may not have allowed large groups to persist during early periods in Southern Siberia, potentially contributing to the disappearance of the *Denisova 5 Neandertal* and *Denisova 25* populations.

### Evidence of “Super-Archaic” ancestry in Denisovans

The availability of a second Denisovan genome provides an opportunity to investigate a previous hypothesis that Denisovans have ancestry from archaic humans who diverged from the modern human lineage earlier than Denisovan ancestors did (“Super-archaic” humans)^33,34,48^. This was suggested because Neandertals share more derived alleles with present-day Africans than Denisovans do, particularly at positions with derived alleles fixed in present-day Africans^34^, a result we replicate with *Denisova 25* (**Figure 5A**; **Supplementary Note 10**). Given more recent evidence for early gene flow from the ancestors or relatives of modern humans to Neandertals^48-54^, the presence of any such gene flow into Denisovans may not be necessary anymore to account for this observation.

**Figure 5.**
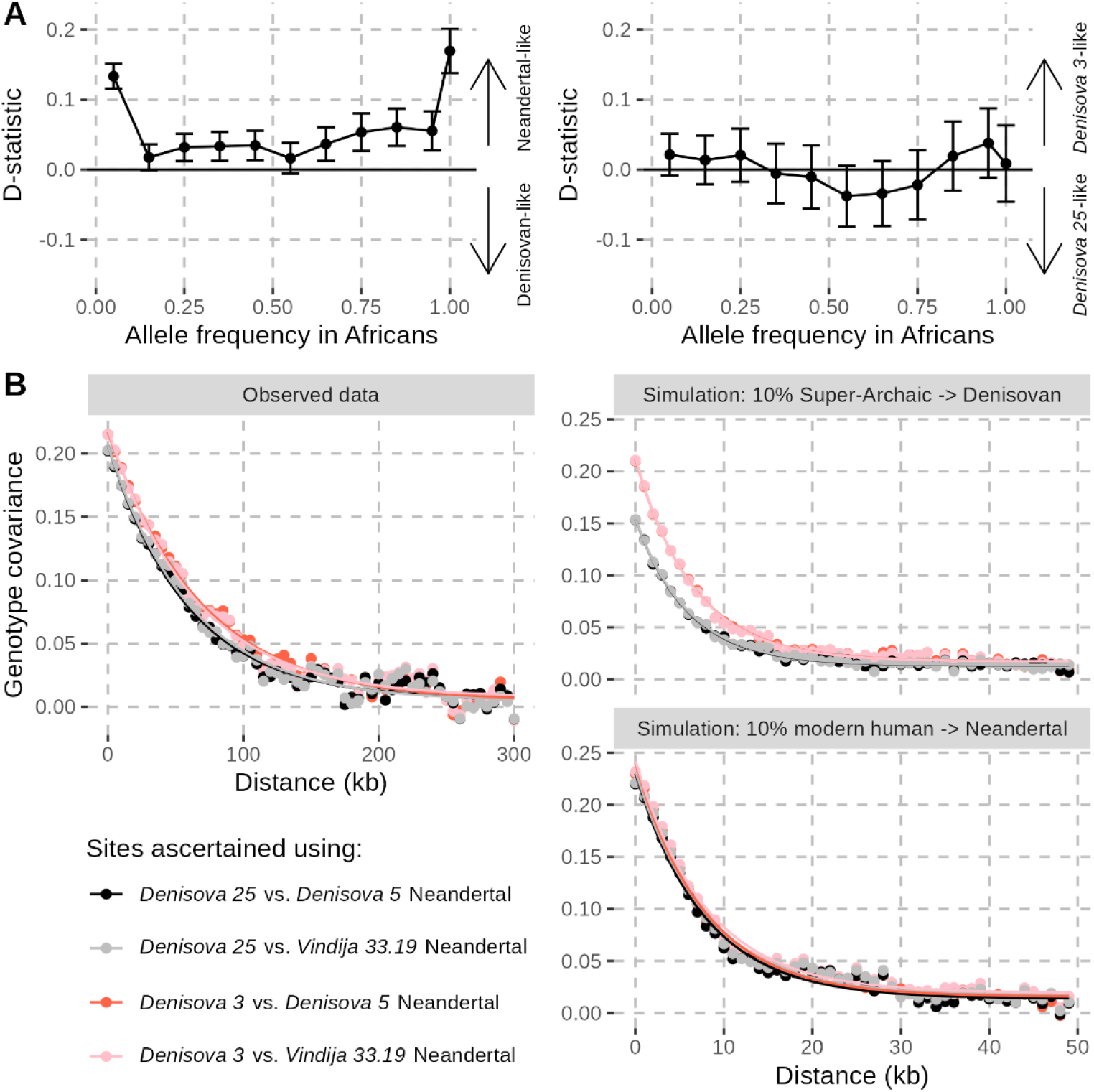
Evidence of “Super-Archaic” ancestry in Denisovans. (**A**) Stratified D-statistics of the form D(Neandertal, Denisovan, African, outgroup) (left) and D(*Denisova 3, Denisova 25*, African, outgroup) (right), binned by allele frequency i SGDP Sub-Saharan Africans (in intervals of 10%, and for fixed derived alleles). The outgroup allele was determined from four great ape reference genomes (**Supplementary Note 8**). Only transversion polymorphisms were used to avoid biases observe with transitions (**Supplementary Note 8**), likely due to recurrent mutations. Whiskers indicate ±2 standard errors estimated with a block jackknife in 5Mb windows. (**B**) Decay of genotype covariance between sites carrying alleles potentially inherited from a “Super-Archaic” hominin (fixed derived in Africans, ancestral in one pseudohaploid archaic genome, and shared with Africans in the other). Points show mean covariance for site pairs binned by distance (kb); curves are exponential fits. Colours indicate genome pairs. Observed data (top left) are compared to coalescent simulations with 10% gene flow from a “Super-Archaic” hominin to Denisovans (top) or from modern humans to Neandertals (bottom) (see **Supplementary Note 10** for more details).

We first analysed the relationship of *Denisova 3* and *Denisova 25* to present-day Africans by comparing their derived allele sharing stratified by the allele frequency in present-day Africans and found no significant differences across the allele frequency spectrum (**Figure 5A**; **Figure S65**). This suggest that if Denisovans received any gene flow from a more divergent archaic group, it is likely to have occurred over 200,000 years ago, i.e., before the *Denisova 3* and *Denisova 25* populations split.

As *Denisova 25* is much older than *Denisova 3* and therefore closer in time to any potential contact with a divergent archaic group, any DNA segments contributed by such a group should be longer in the *Denisova 25* genome than in the *Denisova 3* genome. By examining the decay of linkage disequilibrium between alleles potentially inherited from such a hominin (i.e. at positions where derived alleles are fixed in present-day Africans but where a Neandertal or Denisovan genome carries the ancestral allele), we found a stronger decay in *Denisova 3* than in *Denisova 25*, but no differences among Neandertals (**Figure 5B**). Simulations exploring the effect of deeply divergent and modern human DNA contributions shows that the difference in decay among Denisovans can be explained by gene flow from a deeply diverged hominin in Denisovans but not by only early modern human gene flow in Neandertals. This shows that, in addition to modern human ancestry in Neandertals, an unknown “Super-Archaic” hominin contributed DNA to the ancestors of Denisovans.

Using a method that infers introgressed DNA segments from a divergent population based on an elevated density of single nucleotide variants^55^, we identified 46 and 70 candidate segments for “Super-Archaic” ancestry in *Denisova 3* and *Denisova 25*, respectively, with 33% of *Denisova 3* segments overlapping those reported in a previous study^48^. The higher number of candidate segments in *Denisova 25* may reflect the closer temporal proximity of this individual to the admixture event, facilitating the detection of “Super-Archaic” DNA segments. Notably, such DNA segments were detected also in the Neandertal genomes (44, 45, and 37 candidate segments in *Denisova 5, Chagyrskaya 8*, and *Vindija 33*.*19*, respectively). Since Neandertals should have little or no “Super-Archaic” ancestry, this suggests a high rate of false positives, currently making it difficult to identify “Super-Archaic” DNA segments in Denisovans, perhaps because admixture was ancient enough for segments to be short. However, in Denisovans, unlike in Neandertals, several candidate regions exhibit extreme heterozygosity as well as unusually high divergence both from Neandertals and between the two Denisovan genomes, consistent with retention of ancestry from a deeply divergent lineage (**Extended Figure 1**).

**Extended Figure 1.**
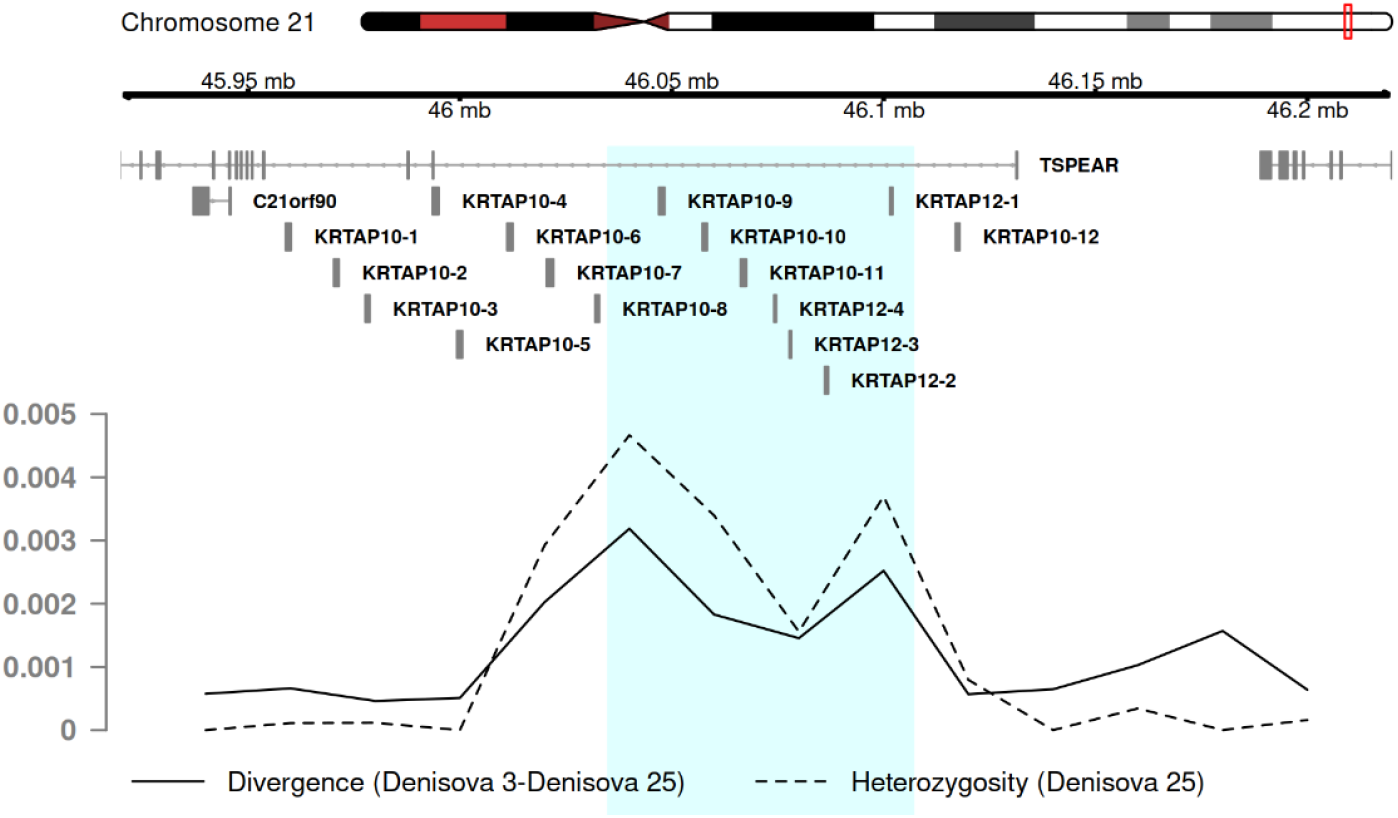
Example of a genomic region in *Denisova 25* with a candidate “Super-Archaic” segment (shaded region), showing high heterozygosity (dashed line) and high divergence from *Denisova 3* (solid line). The region overlaps *TSPEAR* and keratin genes (shown on top) and lies near the end of chromosome 21 (red frame on the ideogram).

### Denisovan ancestry in present-day humans

As genetic contributions from distinct Denisovan populations have been detected in the genomes of present-day humans^20-25^, we next compared how these components relate to the *Denisova 3* and *Denisova 25* genomes. We used a published map of Denisovan ancestry in the 1,000 Genomes Project dataset^21^ as well as new maps we reconstructed for genomes from the SGDP dataset^56^ using a method that relies on the high quality archaic genomes to detect introgressed archaic segments (“reference-based” method)^35^ and a method that infers archaic introgressed segments based on an increase in the density of single nucleotide variants (“reference-free” method)^55^(**Supplementary Note 11**). We present the results using the reference-based method, which are qualitatively consistent with the results from the reference-free approaches (**Supplementary Note 11**). On average, 1.8-fold more Denisovan-like segments are identified in non-Africans when adding the *Denisova 25* genome (**Figure 6A**; **Figures S83** and **S85**). The lowest levels are found in Europeans (average 0.4 Mb) and the highest levels in Oceanians (average 31 Mb) (**Extended Data Table 1**; **Table S66**).

**Figure 6.**
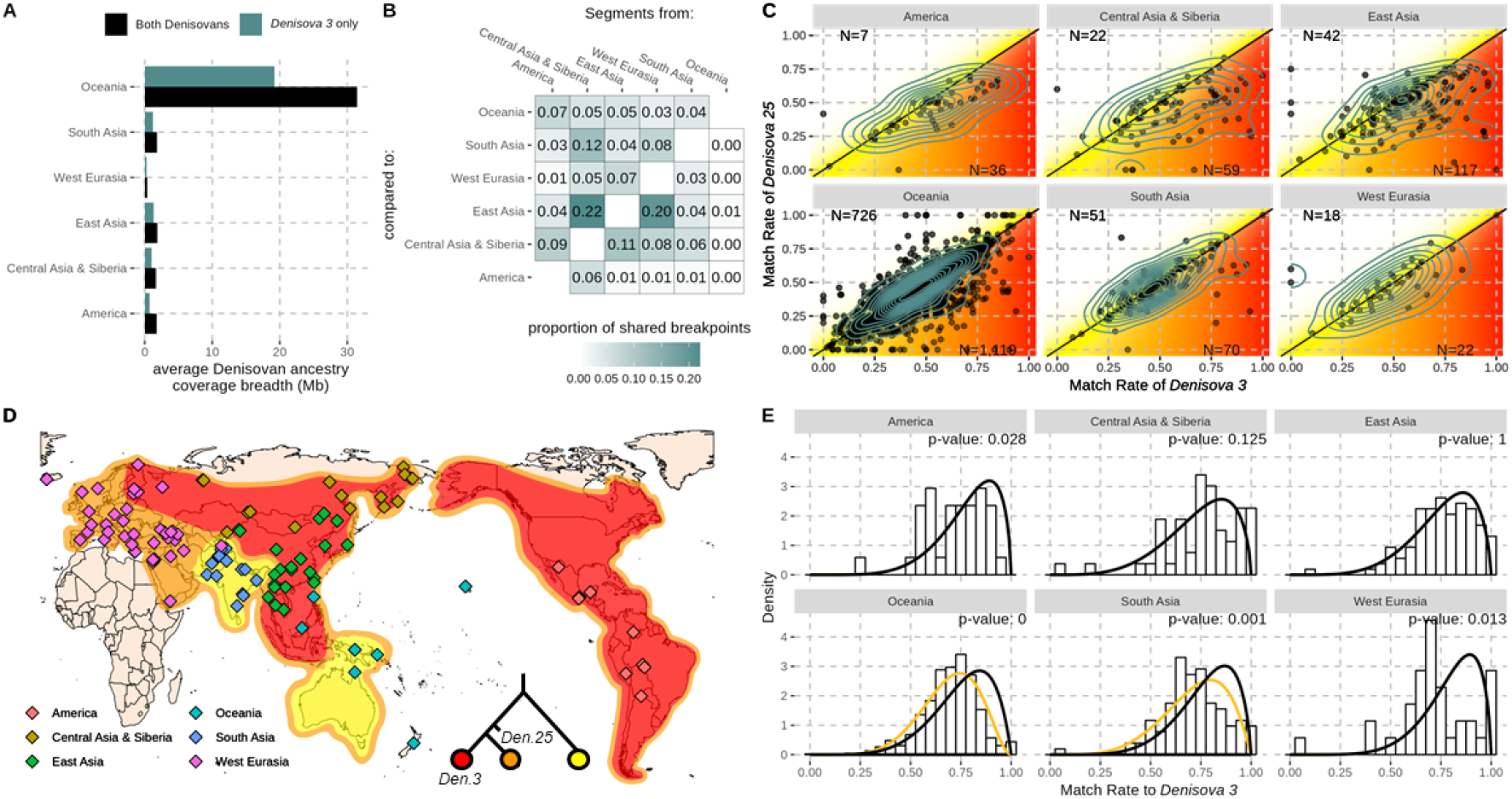
Characterisation of Denisovan ancestry in present-day human genomes. (**A**) Average Denisovan ancestry coverage across genomes from different geographical groups, using one or both Denisovan genomes as references. More details are provided in **Extended Data Table 1. (B**) Proportion of Denisovan-like segments, grouped by population (columns), sharing a breakpoint with those from other populations (rows). (**C**) Match rate of Denisovan-like segments in present-day human genomes to *Denisova 3* (x-axis) and *Denisova 25* (y-axis), by counting the proportion of derived alleles in each segment that are share with the Denisovan genome, after excluding alleles shared with Africans. Each dot represents a different segment and contours show their density. Counts of segments above and below the diagonal are indicated. Roughly expected match rates for each of the three identified Denisovan ancestry components are shown with different colours (as shown in D). (**D**) Tentative geographical distribution of the three Denisovan ancestry components detected in present-day populations, with their relationship depicted in the tree. Diamonds represent the geographical origin of analysed genomes. (**E**) Distributions of match rates to *Denisova 3* at sites where *Denisova 3* has a derived allele absent in an African genome. The fit of a beta-binomial distribution to the match rate of *Denisova 25* to *Denisova 3* within the same genomic segments identified as Denisovan-like in present-day humans is shown (black line). A mixture of two beta-binomial distributions including a more distant component is shown (yellow line) when this mixture provides a better fit to the match rate of Denisovan-like segments in present-day humans (histogram). The distributions were fitted using an Expectation-Maximization algorithm and the reported p-values were computed using a likelihood ratio test. (**A-E**) Segments used in all panels were identified with *admixfrog* using both Denisovan genomes as references. For overlapping segments within a geographical group, one segment is randomly chosen.

The Denisovan-like segments match *Denisova 3* more frequently than *Denisova 25* (**Figure 6C**), a pattern that persists even after masking Neandertal-like segments in both Denisovan genomes, indicating that potential mis-assignment of the Neandertal segments in modern humans, which is more closely related to the Neandertal ancestry in *Denisova 3*, does not explain the observation. Thus, all non-Africans inherited at least part of their Denisovan ancestry from Denisovans who were more closely related to *Denisova 3* than to *Denisova 25*. Consistent with previous findings^21^, we observed an additional Denisovan component that is even closer to *Denisova 3* in the genomes of present-day East Asians, Central Asians, Siberians, Native Americans, and some Europeans (e.g. Finnish). In South Asians and Oceanians, the match rate between Denisovan-like segments and the Denisovan genomes is lower than the *Denisova 3*-*Denisova 25* match rate in the same genomic regions, providing evidence for a different Denisovan component that diverged earlier from the common ancestral population of *Denisova 3* and *Denisova 25* (**Figure 6D** and **6E**). This divergent Denisovan component is not found in populations other than South Asians and Oceanians. Therefore, we detect at least three distinct Denisovan components in present-day humans that are differently distributed across populations today.

To test whether South Asians and Oceanians inherited the most divergent Denisovan DNA component through separate events we compared the breakpoints of Denisovan-like segments among non-Africans, assuming that segments inherited from a common introgression would share breakpoints from past recombination events (**Figure 6B**). However, we found that segments identified in Oceanians almost never share breakpoints with segments in other populations, including South Asians, while the segments identified in other populations share breakpoints broadly across all non-Africans. This suggests that ancestors of Oceanians inherited their Denisovan ancestry independently from other modern humans. It further implies that although the deeply diverged Denisovan component is seen in both South Asians and Oceanians, South Asians must have inherited it in a separate event from Oceanians. These observations support the hypothesis that Oceanians descend from an early wave of modern humans that reached Oceania via a route through South Asia^57^. The observation that East Asians do not carry the deeply diverged Denisovan component, but do have a component most closely related to *Denisova 3*, suggests that their ancestors migrated into East Asia via a more Northern route. Notably, Neandertal-like DNA segments are shared among all populations^58^ (**Supplementary Note 11, Figure S114**), including Oceanians, consistent with a single out-of-Africa event, but independent Denisovan gene flows suggest multiple migrations into Asia (**Extended Figure 2**).

**Extended Figure 2.**
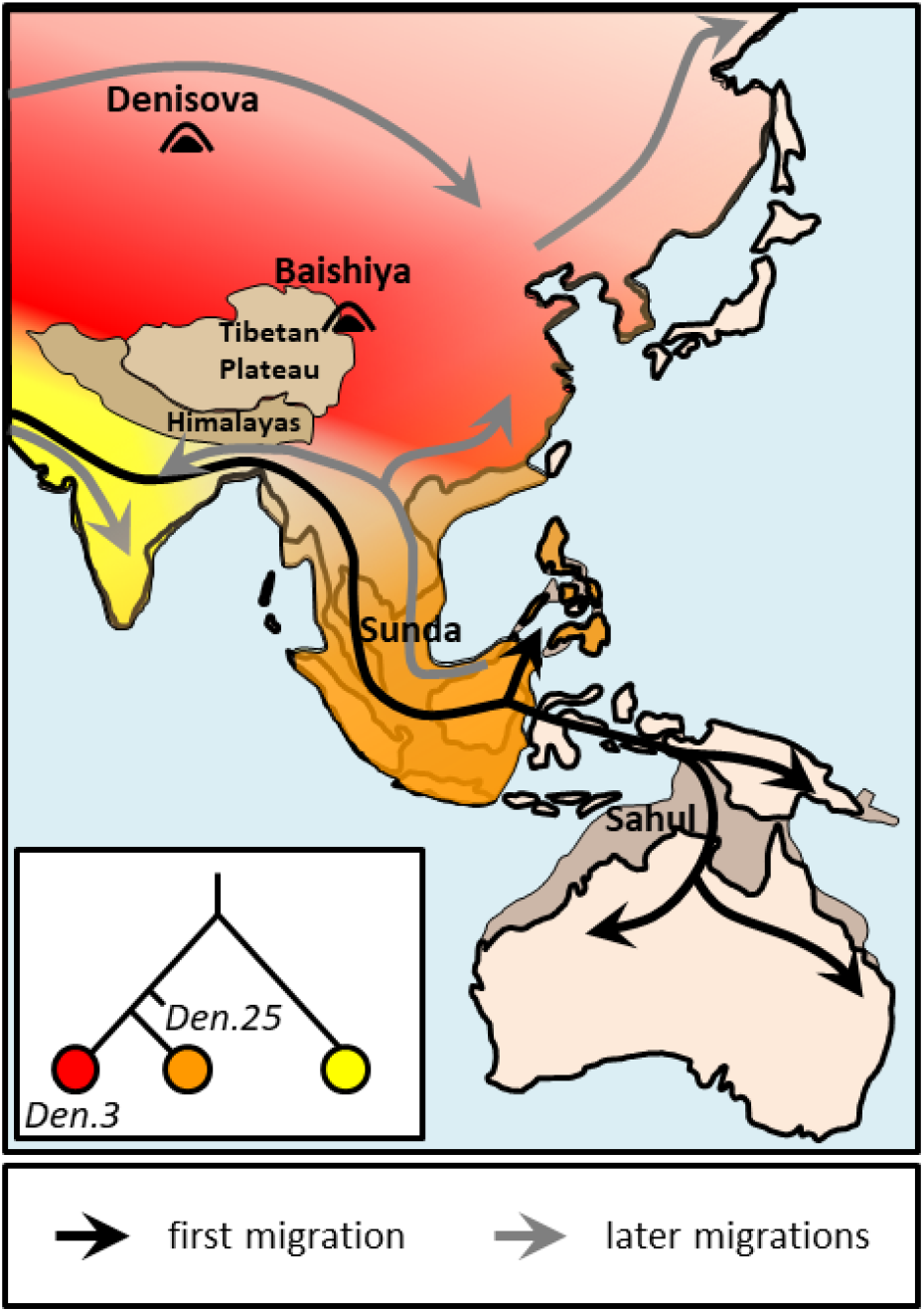
Potential admixture history between Denisovans and modern humans. The phylogeny illustrates the relationship between Denisovans, with colours corresponding to their hypothesised geographical distribution shown on the map. Molecular evidence from Denisova and Baishiya caves (locations indicated) suggests a northern distribution of *Denisova 3*-like Denisovans^16^. Denisovans most divergent from *Denisova 3* were likely in South Asia, given the distribution of this ancestry today. The hypothesised distribution of other Denisovans (orange) is based on morphological evidence^59^ but could have been instead further west, e.g. in the Middle East. Arrows represent the minimum number of migrations required (for this geographical distribution of Denisovans), with colours illustrating their order. Assuming this distribution of Denisovans, the later migration from Southeast Asia into East and South Asia (grey arrows) is necessary to explain the wide distribution of the orange Denisovan ancestry. However, this migration would not be necessary if these Denisovans were found further to the West.

**Extended Table 1.** Average Denisovan ancestry coverage breadth (in Mb). Statistics were computed from putative Denisovan segments of at least 0.05cM. Segments identified with *hmmix* exhibit a posterior probability of at least 0.8 and are assigned as Denisovan based on a majority of alleles shared with a Denisovan genome compared to Neandertal genomes. For the segments identified with *admixfrog*, we report the results using the reference that only includes *Denisova 3* and the reference that includes both *Denisova 3* and *Denisova 25*.

| Region | Reference-based method (*admixfrog*) |  |  |  | Reference-free method (*hmmix*) |  |
| --- | --- | --- | --- | --- | --- | --- |
|  | using *Denisova 3* only |  | using both *Denisova 3* and 25 |  |  |  |
|  | per genome | cumulative | per genome | cumulative | per genome | cumulative |
| America | 0.70 | 7.44 | 1.74 | 17.71 | 3.74 | 27.95 |
| Central Asia/Siberia | 1.02 | 16.41 | 1.69 | 25.05 | 4.38 | 53.67 |
| West Eurasia | 0.26 | 8.60 | 0.37 | 14.53 | 1.28 | 42.36 |
| Oceania | 19.25 | 203.46 | 31.38 | 317.35 | 61.31 | 511.49 |
| Oceania (Australia, Papua New Guinea, Bougainville ) | 24.87 | 194.98 | 40.65 | 305.31 | 79.16 | 489.20 |
| South Asia | 1.18 | 31.56 | 1.76 | 46.88 | 4.63 | 108.41 |
| East Asia | 1.33 | 30.94 | 1.85 | 42.76 | 4.98 | 97.61 |
| non-Africans | 2.73 | 256.27 | 4.41 | 392.08 | 9.37 | 667.64 |

### Dating Denisovan gene flows into modern humans

Assuming that the same Denisovan population mixed with the ancestors of present-day Oceanians and South Asians, the geographic distribution of this ancestry makes South Asia a likely contact zone where Denisovans different from those identified in Denisova Cave mixed first with the ancestors of Oceanians, and later with those of South Asians.

To test whether we can distinguish these admixture events, we estimated Denisovan admixture times using the exponential decay of the linkage disequilibrium between alleles inherited from Denisovans in present-day individuals. We replicated a previous estimate that admixture between the ancestors of Oceanians and Denisovans occurred about 10% more recently than Neandertal admixture^60^ (**Extended Figure 3**; **Supplementary Note 11**). Although uncertainties in recombination rates prevent an absolute time estimate for Oceanians^60^, population-specific recombination maps available for other populations^61^ allowed us to estimate Denisovan admixture times of 50-39ka in South Asians and 44-35ka in East Asians, approximately 20% and 29% more recent than Neandertal admixture time estimates in these populations (62-49ka – the range of estimates obtained for Eurasians), respectively (**Extended Figure 3**; **Supplementary Note 11**). Therefore, the difference in Denisovan admixture times between Oceanians and other populations supports that Oceanians descend from an early wave of modern humans that encountered Denisovans before other non-Africans did.

Europeans are likely to have acquired Denisovan ancestry indirectly through contact with Asian populations as there is no evidence of Denisovan DNA contributions in early modern humans in Europe^62^. East Asians are the most likely contributors of this ancestry, as Europeans share more Denisovan-like segment breakpoints with East Asians than with other Asians (**Figure 6B**). The date of the Denisovan DNA fragments in Europeans (46-34ka) is consistent with that estimated for East Asians.

We could not further deconvolute the timing of introgression for the different Denisovan ancestry components due to limitations in accurately determining the length of Denisovan segments, which we show is sensitive to the choice of methods and parameters (**Supplementary Note 11**). It may in future be possible to get more precise dates for the Denisovan admixtures using ancient genomes, as has been recently shown for Neandertal admixture^58,62^.

**Extended Figure 3.**
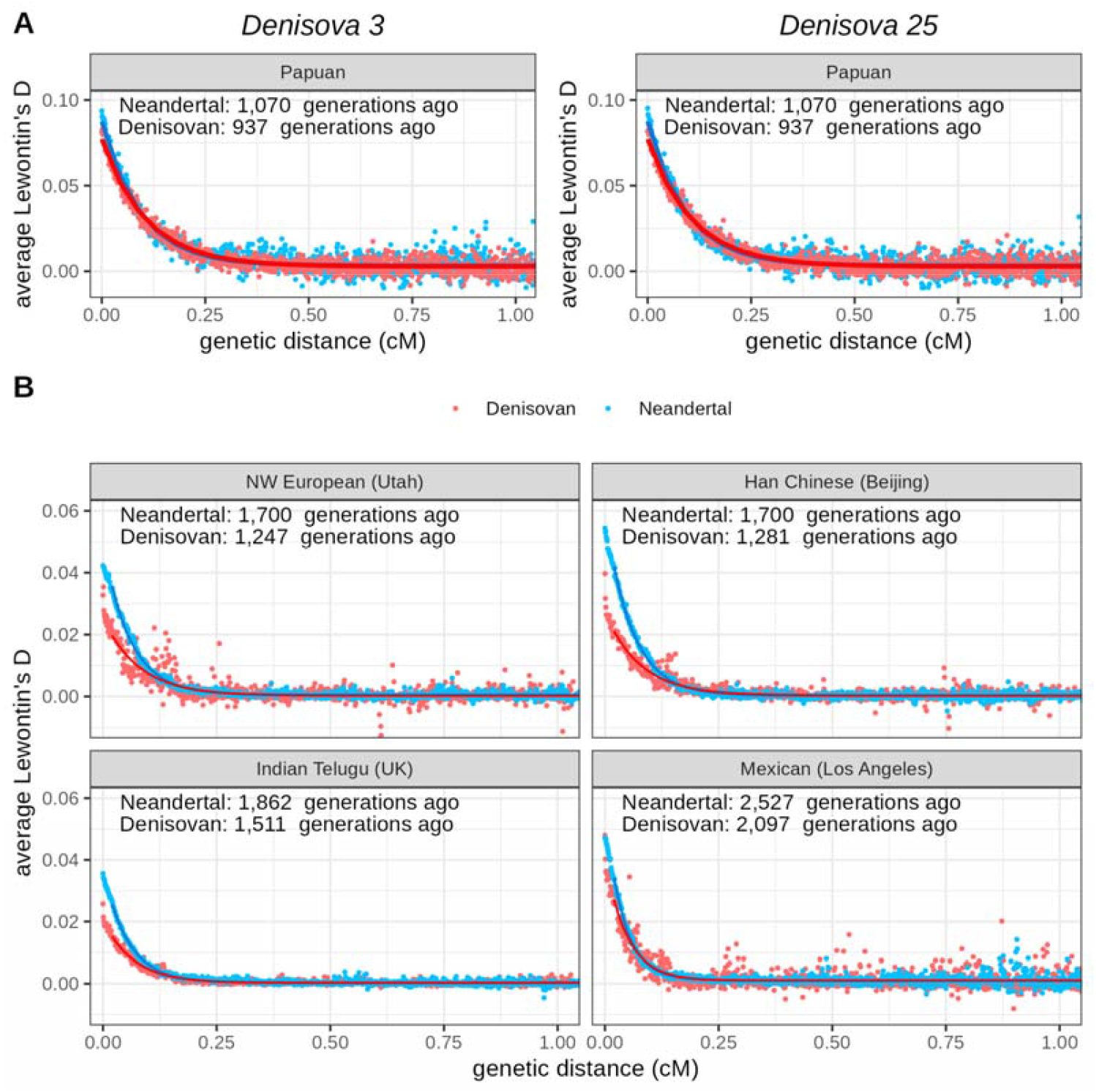
Estimation of Denisovan and Neandertal admixture dates. (**A**) Admixture dates in present-day Oceanians (SGDP dataset^56^), inferred from linkage disequilibrium (Lewontin’s D) decay between putative Neandertal (reference: *Denisova 5*) or Denisovan introgressed alleles (references: *Denisova 3*, left; *Denisova 25*, right). Genetic distances were calculated using the African-American recombination map^63^. (**B**) Admixture dates in other present-day populations (1,00 Genomes dataset^64^, inferred from the linkage disequilibrium decay between putative Neandertal (reference: *Denisova 5*) or Denisovan introgressed alleles (reference: *Denisova 3*, closest to most Denisovan ancestry). Genetic distances were calculated using population-specific recombination maps^61^. For both panels, average linkage disequilibrium in bins of 0.001 cM is shown with fitted exponential decay curves. Estimates in other populations are provided in **Supplementary Note 11**). The estimate of the Neandertal admixture in Native Americans is inconsistent with estimates for all other groups. This seems to be driven by over-estimates of the recombination rates in the population-specific recombination maps perhaps due to the complex admixture history of Native Americans.

### Local adaptation through Denisovan introgression

Denisovans inhabited much of Asia for hundreds of thousands of years and likely adapted to diverse local environments. These adaptations may in turn have benefited incoming modern humans when genetic variants were passed on to them from Denisovans. Previous studies identified Denisovan-like DNA segments at high frequency in modern humans^20,22,65-71^, potentially due to adaptive introgression, but had limited ability to identify the contributing Denisovan population when only one Denisovan genome was available. Leveraging the increased power to detect Denisovan DNA segments using both *Denisova 3* and *Denisova 25*, we scanned the genomes of 236 present-day non-Africans from six geographic regions for candidates of adaptive introgression and compared each segment’s relationship to the two Denisovan genomes (**Supplementary Note 12**).

We defined candidates as segments in the extreme upper tail of the frequency distribution (Z-score ≥ 4.5) in the genomic regions where at least one individual carries Denisovan ancestry. We identified a total of 38 candidate regions, with the highest number (27) identified in Oceanians, who have the highest levels of Denisovan ancestry in our study (**Supplementary Note 11**), the majority of which were not identified in a previous scan of the same genomes^60^. Interestingly, most candidates were unique to single geographic groups, with only two candidates shared by more than one group, suggesting that selection on introgressed Denisovan DNA differed between geographic regions and contributed primarily to local adaptation.

In Oceania and South Asia, segments that are more divergent from *Denisova 3* than *Denisova 25* are significantly more frequent among candidates than expected from the genome-wide composition of Denisovan-like DNA segments. By contrast, in Native Americans, East Asians, and West Eurasians, most candidate segments are closer to *Denisova 3* than *Denisova 25* is, significantly more frequently than expected from Denisovan-like segments genome-wide. These patterns are consistent with local adaptation with northern, *Denisova 3*-like Denisovans contributing most to adaptation in northern populations, and the more divergent, likely more southern Denisovans contributing most to adaptation in Oceanians and South Asians.

Finally, we investigated the potential functional roles of candidates. Functional enrichment analysis of the set of adaptive introgression candidates yields only a single significant association: in South Asia two of the four candidate regions overlap genes, *ATP6V0A4* and *COL4A3*, and both genes are associated with sensory perception of sound and mechanical stimulus (**Table S69**). The introgressed Denisovan segments do not modify the protein coding sequences of either gene, suggesting that any functional impacts could be related to differences in gene expression. Notably, there are four potentially regulatory variants on the introgressed haplotype in *ATP6V0A4*, including a variant in the 5’ UTR of the canonical transcript. The adaptive advantage of these candidates is unclear; as mutations in both *ATP6V0A4* and *COL4A3* also cause renal diseases^72-74^ any advantage may also be related to kidney function.

### An updated catalogue of Denisovan-derived variants

Little phenotypic information is available for Denisovans, as they are known from only a few skeletal remains. However, their genomes have the potential to provide insights into their biology and its similarities and differences with modern humans^75-78^. A previous analysis of the *Denisova 3* genome identified 692,818 single nucleotide changes that may be Denisovan-derived^13^, though many of these likely represent variants specific to that individual.

Using the newly available high-coverage Denisovan genome and three high-coverage Neandertal genomes, we identified 237,112 single nucleotide variants shared between both Denisovan genomes (allowing a missing genotype in one), thereby refining the set of variants that likely arose on the Denisovan lineage (**Supplementary Note 13**). Of these, 1,307 result in amino acid substitutions in 1,142 protein-coding genes. Variants at evolutionary conserved sites are likely to have the largest functional effects. Using a deep learning approach that integrates nucleotide conservation across 236 primate species with protein 3D structure information^79^, we identified ten missense changes predicted to have the strongest effects (PrimateAI-3D score between 0.8 and 1). Five of these genes, *COPB1, KIF14, PIGG, PLCE1* and *VWA8* have been linked to morphological traits^80^, several of which show changes in Denisovan fossils. In particular, *COPB1, KIF14*, and *PIGG* influence cranial size, head circumference, and orbital size, all of which have been described to be large in the Denisovan cranium from Harbin^11,12,26^. Furthermore, *COPB1* and *PIGG* are also associated with finger breadth, which is potentially relevant given that the *Denisova 3* phalanx is more gracile than that of Neandertals^81^, while *KIF14* and *PIGG* are associated with projection of the lower jaw, and *KIF14* with a sloping forehead, which are also described in the Harbin cranium^26^. These genes are therefore promising candidates for understanding the distinct cranio-facial morphology of Denisovans.

Changes that influence gene regulation may also have an important impact on phenotypes. Although many regulatory variants occur within non-coding regions where their target genes are difficult to determine, those located in promoter-proximal regions typically allow for a more direct link to gene expression. We identified 1,126 changes in promoter-proximal regions potentially impacting transcription initiation of 1,052 genes. Those with changes at highly conserved positions, inferred from 447 mammalian genomes^82^, are significantly enriched in the gene ontology terms central nervous system development (GO:0007417), brain development (GO:0007420), and neuron projection membrane (GO:0032589) (family-wise error rates ≤ 0.05) (**Table S67**). The change at the most conserved position (phyloP score 8.097) lies 395 base pairs upstream the *FOXP2* transcription start site, within an “ultra-conserved” 22-base-pair region identical across 235 mammalian genomes^83^. The Denisovan allele likely disrupted binding of the transcription factor ISL1 that is predicted to bind strongly to the ancestral allele (**Supplementary Note 13**), and thereby potentially modified the expression of *FOXP2*. This gene encodes forkhead box protein P2, a transcription factor expressed in many tissues. In the brain, it is involved in cerebral cortex development (GO:0021987), as well as speech and language development^84,85^.

The effects of introgressed Denisovan alleles on modern human phenotypes may also provide some hints into Denisovan biology. Using alleles that have been associated with phenotypes in modern humans^86^, we identified 16 associations with 11 Denisovan alleles (**Table S75**), including height, blood pressure, monocyte count, and levels of cholesterol, haemoglobin and C-reactive protein. We also identified 305 expression quantitative trait loci (QTL) and 117 alternative splicing QTL affecting gene expression in modern humans across nineteen tissues, the strongest effects include eQTLs in thyroid, arterial tibial, testis and muscle (**Figure S125**). These molecular effects can be leveraged to explore further phenotypes not preserved in the fossil record, and this updated catalogue provides a more reliable basis for exploring Denisovan traits, adaptations, and disease susceptibilities, some of which may have been contributed to present-day humans through admixture.

### Future perspectives

Using this second Denisovan genome has shown that there was recurrent mixing between Neandertals and Denisovans in the Altai region, but that these mixed populations were replaced by Denisovans from elsewhere, supporting the idea that Denisovans were widespread and that the Altai may have been at the edge of their geographic range.

The Denisovan ancestry in modern humans provides a glimpse into the diversity of Denisovans, and suggests that Denisovans may have lived in populations that were geographically isolated from one another over an extended time. This isolation may simply reflect the large distances over which Denisovans were distributed or be due to more substantial geographic barriers. The presence of deeply divergent Denisovan ancestry in both Oceanians and South Asians makes it plausible that the Denisovans that contributed ancestry to these groups were isolated in South Asia by the Himalayas.

Retrieving genetic data from these deeply diverged Denisovans will be critical to fully understand the origins and diversity of Denisovans, including their relationship to Neandertals and to the more deeply diverged hominin that contributed DNA to Denisovans.

## Data availability

The sequences and genotypes from this study will be deposited in the European Nucleotide Archive. The FASTA file of the reconstructed mitochondrial genome will be deposited in GenBank.

## Acknowledgment

This project was funded by the Max Planck Society and the European Research Council (grant agreement no. 694707). Archaeological research at Denisova Cave by M.V.S. and M.B.K. was supported by the Russian Science Foundation, grant number 24-18-00069. We acknowledge support from the National Genomics Infrastructure in Stockholm funded by Science for Life Laboratory, the Knut and Alice Wallenberg Foundation and the Swedish Research Council, and SNIC/Uppsala Multidisciplinary Center for Advanced Computational Science for assistance with massively parallel sequencing and access to the UPPMAX computational infrastructure. We thank Isabel Pineda Yupanqui for helping prepare Figures 2 and 6.

## Author contributions

S.Pe. and J.K. designed the study. D.M., E.E., S.N. and J.R. performed ancient DNA lab work. A.W., B.S., H.Z., J.V., S.Pe. and J.K. performed DNA sequencing and raw data processing. M.B.K., M.V.S. and A.P.D. provided and analysed archaeological material. B.V. performed the morphological analyses. S.Pe., Y.S., M.J.B., L.N.M.I., A.P.S., A.B.M., C.d.F. and K.P. performed or aided the genetic analyses. J.K., S.Pä., B.M.P. and M.M. supervised the study. J.K. S.Pe and J.K. wrote the manuscript with input from all co-authors.

## Ethics declarations Competing interests

The authors declare no competing interests.

## Supplementary information

This file contains Supplementary Notes 1-13 including figures, tables and references. See contents page of the Supplementary Information file for more details.

