## Supplementary Notes for "A high-coverage genome from a 200,000-year-old Denisovan"

### Supplementary Note 1: Archaeological context and morphology of *Denisova 25*

#### Archaeological context

The *Denisova 25* molar was found in the central part of the South Chamber of Denisova Cave in the upper part of layer 17 (square D-2). The sediments of layer 17 are light silty loam, brownish-gray and dark gray with a brown tint, with the inclusion of fine smoothed weathered limestone debris covered with a whitish reaction rim. According to the results of OSL dating and biostratigraphic indicators, the lithological and stratigraphic analogue of layer 17 is layer 20 in the Main Chamber of the cave and layer 14 in the East Chamber, which correspond to MIS 7^1,2^.

The diverse large mammal fauna from layer 17 and the underlying layer 18, the lowest cultural unit in the South Chamber of the cave, includes numerous remains of the dhole Cuon alpinus, wild horse Equus ferus, Siberian roe deer Capreolus pygargus, giant deer Megaloceros giganteus, dzeren Procapra gutturosa, and Merck's rhinoceros Stephanorhinus kirchbergensis^3^. The small vertebrates include the Altai mole Asioscalops, wood mouse Apodemus (Alsomys), voles Clethrionomys and Siberian zokor Myospalax myospalax, and the tree squirrels include chipmunks Eutamias, a representative of taiga biotopes, and Siberian flying squirrel Pteromys, which lives in tall-trunk birch forests^4^. In general, the remains of the theriofauna indicate that these deposits were formed in relatively warm climatic conditions of the interglacial period.

The taphocoenosis from layers 18 and 17 is characterized by active human impact on bone remains. There are more bone fragments with numerous traces of cuts and cutting of meat by stone tools, split and chipped bones, bone flakes and burnt bones than in the overlying layers.

The highest density of archaeological finds in the Pleistocene strata is noted in the deposits of layers 18 and 17. The collection of lithic products includes more than 10,000 artifacts that have similar technical and typological characteristics and belong to the same complex of the initial stage of the Middle Paleolithic.

The creators of this stone industry were well acquainted with the physical properties of the lithic raw materials of local alluvial sources. For the manufacture of tools, they selected homogeneous pieces with a hardness of 5–6.5 on the Mohs scale, mainly sedimentary rocks – siltstones and sandstones, less often – volcanic aphyric or porphyritic effusives.

The lithic industry from layers 18 and 17 is based mainly on radial technology (**Figure S1, 3**) of pebbles or secondary cores from large massive flakes. In accordance with this technology, the main products of primary flaking are shortened and short blanks with a residual striking platform beveled towards the ventral side. Numerous débordantes of radial cores (**Figure S1, 14**) also served as blanks. At the same time, the industry contains samples of parallel cores (**Figure S1, 15, 16**), while the proportion of blades (**Figure S1, 13**) among the blanks is insignificant.

Among the artefacts with secondary processing, large flakes with ventral thinning (**Figure S1, 5, 6**) predominate, formed as a result of wide removals from the distal or one or two lateral edges. Another characteristic type of products, technologically close to the previous one, are basally-truncated flakes (**Figure S1, 2, 8, 9**), the proximal edge of which is truncated by ventral removals, often thinning the base of the blank. The next most important category is a variety of side-scrapers (**Figure S1, 1, 11, 12**), a significant part of which are Quina-type products (**Figure S1, 4, 10**), including those that have undergone several cycles of re-shaping. These three main categories of inventory are represented by stable series and are considered as regional cultural-identifying markers of the initial stage of the Middle Paleolithic. The tool set is complemented by a variety of denticulate (**Figure S1, 7**) and notched forms.

In general, the lithic industries of the early Middle Paleolithic from different areas of Denisova Cave number about 50,000 artifacts^5^. At present, in Altai Mountains, as well as throughout the entire territory of Northern and Central Asia, there are no reliable cultural and chronological analogies to this the most ancient Middle Paleolithic complex.

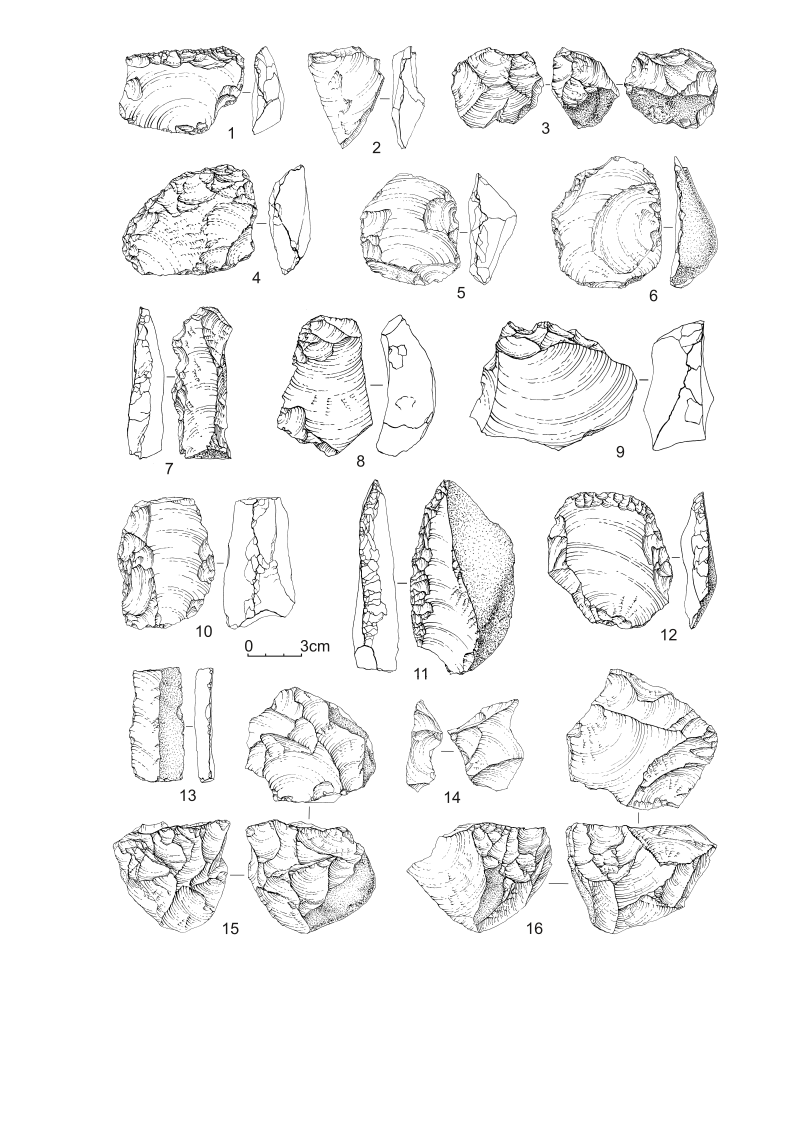

**Figure S1: Lithic artifacts from layer 17 in the South Chamber of Denisova Cave.**

1, 4, 10–12 – side-scrapers; 2, 8, 9 – basally truncated flakes; 3, 15, 16 – cores; 5, 6 – ventrally thinned flakes; 7 – denticulate tool; 13 – blade; 14 – radial core débordantes.

#### *Denisova 25* description

*Denisova 25* is a complete left upper molar. The preservation is in general good, with most of the surface well preserved. The lingual root has been sampled for DNA, consisting of the debriding of some of the lingual surface along the whole length of the root, and the drilling of a 1.5x2mm whole that reached the pulp cavity. Dental calculus is present on the crown, especially on the mesial aspect around the interproximal facet (IPF), with smaller amounts on the lingual and distal surfaces.

There are clear wear facets on the occlusal surface, but without any dentine exposures. There is a large, suboval (5x 2.6 mm) interproximal facet on the mesial aspect of the crown. The IPF shows several subvertical striations on the lingual part of the facet. Distally, there is no interproximal facet. A small chip of enamel (1.2x0.6 mm) is missing on the distobuccal corner of the crown adjacent to the cervix.

The crown outline is rather rounded, and tapers distally, with a mesiodistal diameter of 12 mm, and a buccolingual diameter of 13.7 mm. The lingual half of the crown is dominated by the large protocone. On the occlusal surface, the hypocone is only represented by a very slight elevation on the distolingual margin of the tooth (ASUDAS Grade 0). Wear might have obscured its presence, but it is not separated from the protocone by any kind of fissure, and is only faintly visible on the enamel-dentin junction (EDJ).

On the buccal half, both paracone and metacone (ASUDAS Grade 5) are large, and are separated by a sharply delimited buccal groove. The buccal groove intersects the marked central groove at a right angle. The central groove is deeply incised, and interrupts both the transverse crest connecting the protocone and paracone and the crista obliqua.

The mesial marginal ridge is obscured by wear, but it probably carried several accessory tubercles, as three discrete ones are visible on the EDJ.

Two roots are present, with the larger buccal root (length 16.2 mm, as measured from the cemento-enamel junction) the result of a fusion of a thinner mesiobuccal and somewhat more robust distobuccal root. The cross section of the this fused buccal root is C-shaped, but with a marked developmental groove on the buccal surface. The lingual root is 16.8 mm long (measured from the enamel-cementum junction) lacks the pronounced flare seen in *Denisova 4*, and is close to vertical. In its present state, the lingual root appears relatively slender, but this is probably in part due to debriding of the lingual surface. The root apices are closed.

We visualized the enamel dentine junction via µCT scanning (BIR Actis 300 µCT, resolution 0.0128mm isovoxel). The enamel and dentine were then segmented slice by slice in Avizo 8.0, using a combination of thereshold based segmentation and the magic wand tool.

The dentine horns are placed on the marginal ridges (centrally placed dentine horns absent, according to Martin *et al*., 2017’s scoring scheme^6^). The tallest dentine horn is the protocone and metacone. Just like on the occlusal surface, the crista obliqua is absent. The dentine horn of the hypocone is present as a small raised area on the marginal ridge running from the protocone towards the distal margin of the crown. There is a slight triangular depression on the lingual aspect of the crown next to the hypocone, and a slight essential crest that descends from the hypocone dentine horn towards the central basin. A minor post-paracone tubercle^6^ is visible. Three accessory tubercles are present on the mesial marginal ridge, descending in size from lingual to buccal, with the lingual and central one showing clear essential ridges descending into the mesial fossa. The distal marginal ridge does not show any clear accessory tubercles, but is sinuous in distal view, with two slight elevations.

#### Identification and comparisons

The assignment of isolated molars to position can be difficult, especially in populations like the Denisovans for which there is a very limited comparative sample. Both previously described molars from Denisova, *Denisova 4*^7^ and *Denisova 8*^8^ were described as an M^2^ or M^3^.

Both lack distal interproximal facets, which in the case of *Denisova 4* could also be due to the M^3^ not yet erupting, but in the case of *Denisova 8* this is less likely due to the more advanced wear on the tooth. A possibility is also that these are both second molars, without corresponding third molars. M3 agenesis or strongly reduced M3s are quite frequent in the Asian fossil record, and is also seen in the probably Denisovan Baishiya mandible^9^, the likely contemporaneous Penghu mandible^10^ and probably in the Harbin cranium^11^.

For *Denisova 25*, we have a problem similar to the two previously published molars. In this case as well, there is no distal interproximal facet, but due to the relatively limited wear, it is possible that a potential tooth behind this would not have erupted yet. The strongly reduced distal part of the crown, with a lack of the hypocone fits with an M^2^ or M^3^.

In Neandertals, M2s usually have a hypocone (11.6% ASUDAS 0-1)^12^, while it is more often missing in M^3^s (35.2% ASUDAS 0-1)^12^. In the Atapuerca Sima de los Huesos sample, which is close to the likely split between the Denisovan and Neandertal lineages, hypocone reduction is much more frequent in M^2^s (33.4% ASUDAS 0-1)^12^, and is pretty ubiquitous in M^3^s (95.2% ASUDAS 0-1)^12^.

The relatively large metacone in *Denisova 25* contrasts with the usually reduced metacone in SH M^3^s (81% ASUDAS 0-2)^12^, while in Neandertals this is more frequent (50% ASUDAS 0-2)^12^.

The last trait that can be compared to these samples is the absence of the crista obliqua in *Denisova 25*. All 20 M^3^s from Atapuerca Sima de los Huesos lack a crista oblique, while it is present in 83.3% of the 18 M^2^s^12^. In Neandertals, it is missing in 28.6% of M^2^s, and 70.6% of M^3^s.

Taking into account the strong reduction of the hypocone and the lack of the crista obliqua, it seems more likely that *Denisova 25* is an M^3^. As for the other two molars from Denisova, *Denisova 4*, with its large hypocone is in our opinion most likely an M^2^, while the unusual, five-cusped morphology of *Denisova 8* fits better with an M^3^.

*Denisova 25* expands the morphological pattern seen in the other two upper molars from Denisova. While *Denisova 4* and *8* are characterized by extremely large crowns with large main cusps and numerous accessory cusps, *Denisova 25* shows a simpler morphology, with only three main cusps, and accessory cusps only visible on the EDJ. It also lacks the very large and strongly diverging roots seen in *Denisova 4*. Despite the slightly smaller mesiodistal and buccolingual diameters than the other two teeth from Denisova, it is still above the range of variation seen in Neandertals (with the exception of the likely pathological *Obi Rakhmat 1* molar), and most other Middle Pleistocene and later *Homo* specimens. Interestingly, the Harbin cranium^11^ preserves an upper M2 that is similar to *Denisova 25* in its outline and general shape, though detailed comparisons are difficult due to the strong wear on Harbin. It is striking though that the Harbin molar is one of very few specimens that are close in size to the Denisova teeth.

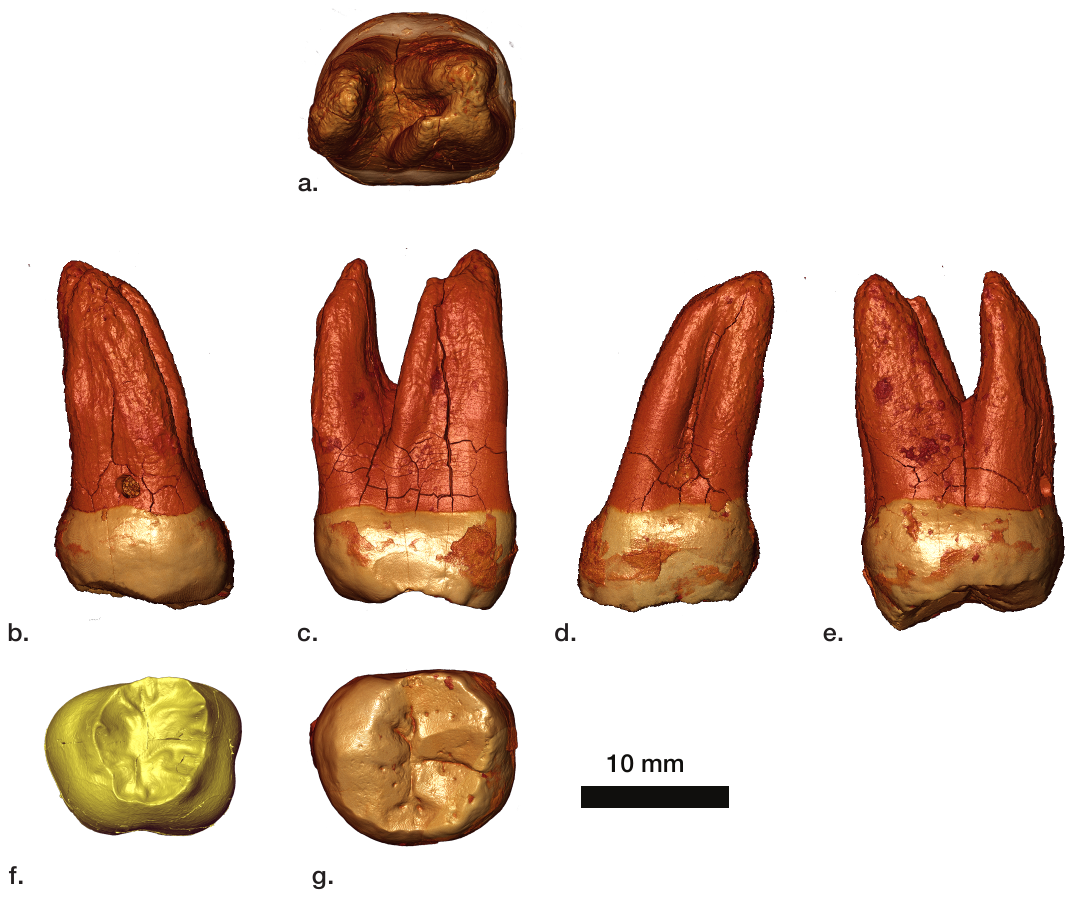

**Figure S2: µCT based renderings of Denisova 25.** a. apical view, b. lingual view, c. mesial view, d. buccal view, e. distal view, f. Enamel-dentine junction in occlusal view, g. crown in occlusal view.

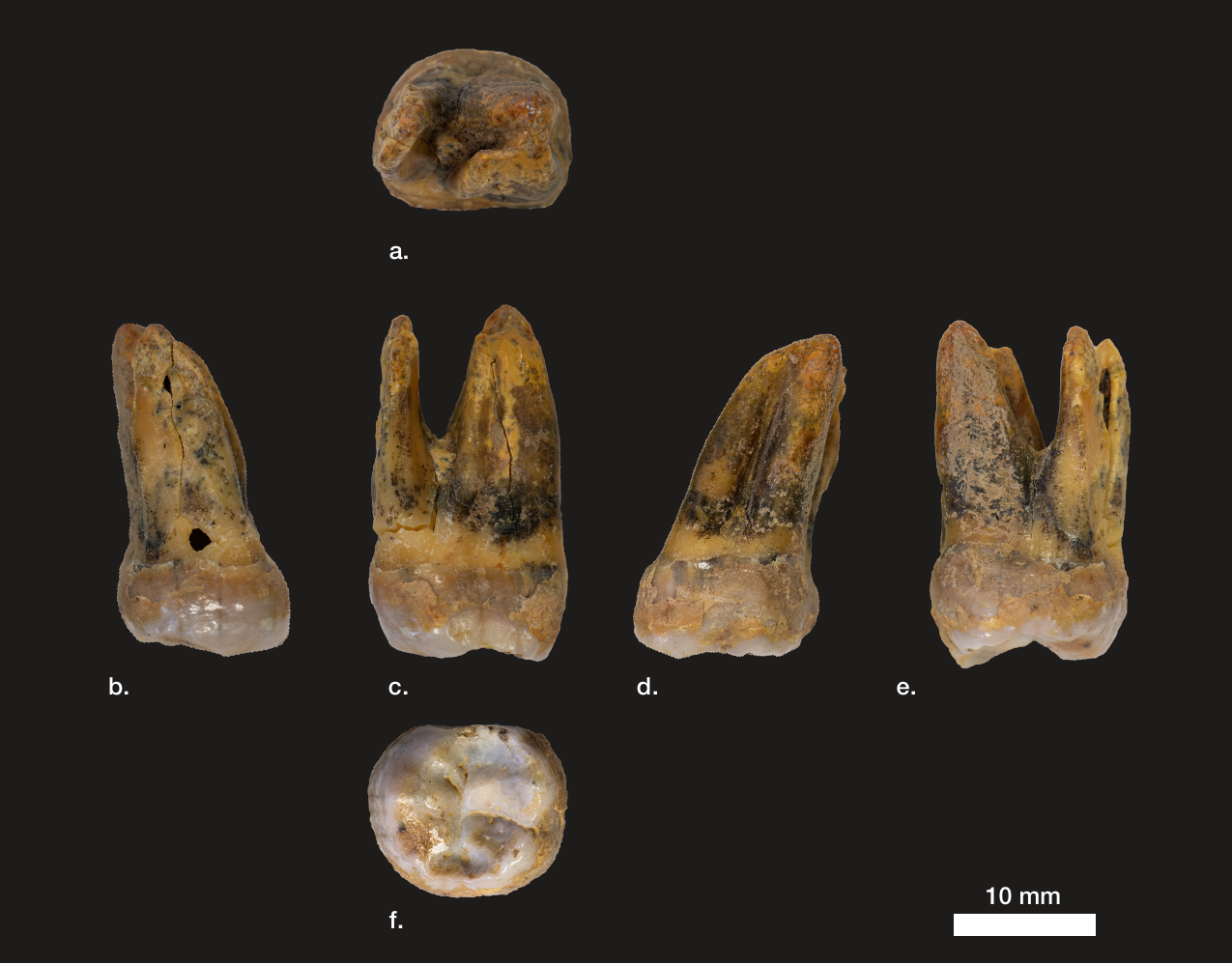

**Figure S3: Denisova 25.** a. apical view, b. lingual view, c. mesial view, d. buccal view, e. distal view, f. crown in occlusal view.

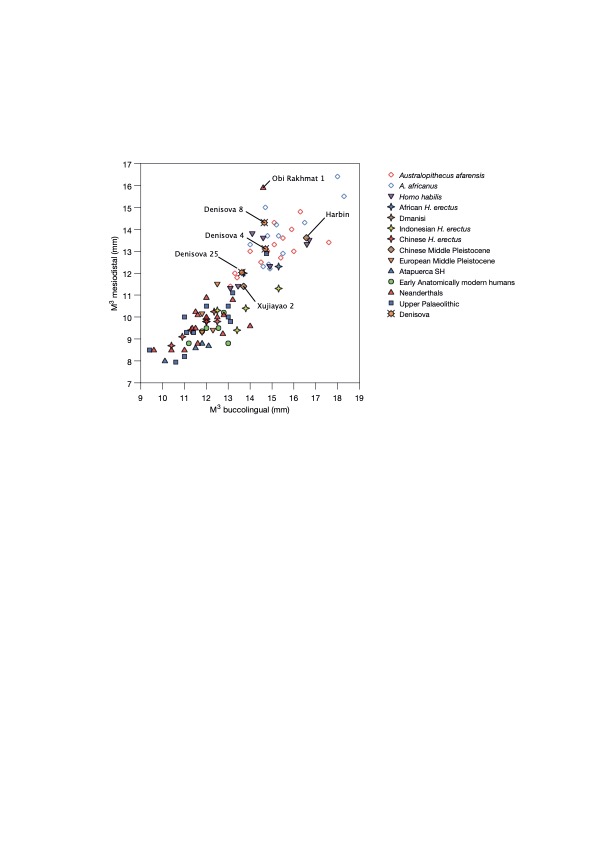

**Figure S4: Biplot of mesiodistal and buccolingual measurements of the Denisova molars and other hominin M^3^**s. Denisova 4 and Harbin are plotted here for comparison, despite clearly (Harbin) and more likely being an M^2^ (Denisova).

**Table S1: Metric comparisons of hominin third upper molars.** (Mean+-standard deviation (N) and mean (range; N), if N <5)

|  | **M^3^ md** | **M^3^ bl** |
| --- | --- | --- |
| ***A. afarensis*** | 13.1±1 (14) | 15±1.3 (14) |
| ***A. africanus*** | 13.8±1.3 (12) | 15.6±1.4 (12) |
| ***Homo habilis*** | 12.7±1.1 (7) | 14.8±1.4 (7) |
| **Dmanisi** | 9.8 (1) | 12 (1) |
| ***H. erectus* (Africa)** | 12.2 (12-12.3; 2) | 14.5 (13.7-15.3; 2) |
| ***H. erectus* (Indonesia)** | 10.4 (9.4-11.3; 4) | 13.8 (12.5-15.3; 4) |
| ***H. erectus* (China)** | 9.6±0.5 (7) | 11.6±0.8 (7) |
| **Atapuerca SH** | 8.5±0.4 (4) | 11.4±0.9 (4) |
| ***H. heidelbergensis* (Europe)** | 10.1 (9.3-11.5; 4) | 12.1 (11.8-12.5; 4) |
| **Neandertals** | 10.1±1.8 (17) | 12±1.3 (17) |
| **Early AMH** | 9.4±0.5 (6) | 12.2±0.7 (6) |
| **Upper Palaeolithic** | 9.8±1.4 (12) | 12±1.5 (12) |
| **Harbin M^2^** | 13.6 | 16.6 |
| ***Denisova 4* M^2^** | 13.1 | 14.7 |
| ***Denisova 8*** | 14.3 | 14.65 |
| ***Denisova 25*** | 12 | 13.7 |

*A. afarensis*: Hadar, Omo (own measurements)

*A. africanus*: Stekfontein, Makapansgat^13^

*Homo habilis*: Olduvai^14^, East Turkana^13^

Dmanisi^15^

*H. erectus* (Africa): East Turkana^13^, Nariokotome^16^, Konso^17^, Swartkrans^13^

*H. erectus* (China): Zhoukoudian^18^, Hexian^19^

*H. erectus* (Indonesia): Trinil^13^, Sangiran^20^ (own measurements)

Atapuerca SH^12^

*H. heidelbergensis* (Europe): La Chaise^21^, Biache^22^, Arago^23^, Petralona^21^

Neandertals: Amud^24^, Châteauneuf^25^, St. Brelade^26^, Krapina^27^, La Croze de Dua^26^, La Quina^26^, Le Moustier^26^, Obi-Rakhmat (own measurements), Saccopastore^26^, Shanidar^28^, Spy^26^, Tabun^26^, Vergisson la Falaise^26^

Early AMH: Skhul^29^, Qafzeh^30^, Temara^31^

Upper Palaeolithic: Brno^26^, Changwu^19^, Cro-Magnon^26^, Dolni Vestonice^32^, Grotte des Enfants^26^, Kostenki (own measurements), La Rochette^26^, Leuca^26^, Mladec^33^, Oase^34^, Predmosti^26^, Sungir (own measurements)

Harbin^11^

### Supplementary Note 2: Generation of preliminary sequencing data for the assessment of ancient DNA preservation

#### Specimen sampling

The *Denisova 25* specimen was sampled in a clean room facility dedicated to the processing of ancient DNA at the Max Planck Institute to Evolutionary Anthropology in Leipzig (Germany). Using a dentistry drill, two small samples of 2.7 mg and 8.9 mg were obtained by drilling one hole at the cemento-enamel junction of the tooth. Additionally, to improve retrieval of ancient endogenous DNA, twelve small subsamples ranging from 4.5 mg to 20.2 mg were obtained by gently scratching the outer layer of one of the tooth root with a dentistry drill (**Table S2**)^35,36^.

#### DNA extraction

DNA was extracted from the two cemento-enamel junction subsamples and nine subsamples from the outer layer of the root using silica-coated magnetic beads on an automated liquid handling platform^37^. The remaining three subsamples from the outer layer of the root were incubated with 500 µL of 0.5% sodium hypochlorite (bleach) at room temperature for 15 minutes under rotation (**Table S2**). After centrifugation and removal of the bleach supernatant, the pellet was washed three times with 500 µL of water for 3 minutes to remove residual bleach^38,39^. DNA was extracted from the bleached treated pellet following the same silica-coated magnetic beads automated protocol as for the non-bleached extracts.

Due to the low quantities of input material, the volume of lysis buffer was reduced to 300 µL, of which 150 µL were used to produce 30 µL of DNA extract. Extracts E24830, E24831 and E24832 were made using the remaining 150 µL of the lysates used to make extracts E23935, E23937 and E23939, respectively (**Table S2**). Extraction negative controls were carried through each step of DNA extraction for both the bleach treated and the non-treated DNA extracts.

#### DNA library preparation

The entire 30 µL of DNA extracts were converted into single stranded DNA libraries following a protocol developed for highly degraded ancient DNA, as previously described^40^ using an automated liquid handling platform^41^. The libraries were amplified according to a double indexing scheme^42^. Library negative controls were carried through all steps of the experiments. No uracil-DNA glycosylase treatment was performed to preserve the C to T substitution patterns to authenticate DNA sequences.

A total of 17 single-stranded DNA libraries were made including three for which extracts were pre-treated with bleach (**Table S2**). Using quantitative PCR, we estimated the number of DNA molecules incorporated in each library to be between 3.09x10^8^ and 4.70x10^9^ for bleach treated extracts and between 1.51x10^9^ and 9.46x10^10^ for non-bleached treated extracts, which is higher than for libraries prepared from negative controls (**Table S2**). Unfortunately, we found that the bleach treatment greatly reduced the amount of endogenous DNA extracted, making it unsuitable for the subsample of the outer layer of the tooth root of *Denisova 25*.

**Table S2: List of DNA extracts and libraries prepared from Denisova 25.** The extraction and library preparation negative controls are marked in grey. The four libraries selected for further sequencing are marked in bold and highlighted with a grey shading.

|  | **extract ID** | **tooth powder used (mg)** | **pre-treatment** | **number of molecules in library** | **number of control oligos in library** | **indexed library ID** | **captured library ID (MT)** |
| --- | --- | --- | --- | --- | --- | --- | --- |
| *Denisova 25* | E22998 | 5,5 | no treatment | 8.30E+09 | 7.52E+05 | A41435 | B48024 |
|  | E22999 | 7,2 | no treatment | 1.11E+10 | 6.40E+05 | A41436 | B48025 |
|  | E23000 | 2,7 | no treatment | 3.44E+09 | 1.00E+06 | A41437 | B48026 |
|  | E23001 | 8,9 | no treatment | 9.46E+10 | 9.46E+05 | A41438 | B48027 |
|  | E23934 | 9,1 | Sodium hypochlorite | 3.09E+08 | 1.06E+06 | A43614 | B49759 |
|  | **E23935** | **6,3** | **no treatment** | **1.96E+09** | **1.02E+06** | **A43615** | **B49760** |
|  | **E24830** |  |  | **1.51E+09** | **7.28E+05** | **A45393** | **B50617** |
|  | E23936 | 20,2 | Sodium hypochlorite | 1.61E+09 | 9.89E+05 | A43616 | B49761 |
|  | **E23937** | **11,3** | **no treatment** | **3.91E+09** | **1.10E+06** | **A43617** | **B49762** |
|  | **E24831** |  |  | **3.07E+09** | **8.56E+05** | **A45394** | **B50618** |
|  | E23938 | 13,5 | Sodium hypochlorite | 4.70E+09 | 7.27E+05 | A43618 | B49763 |
|  | E23939 | 4,5 | no treatment | 1.19E+10 | 9.76E+05 | A43619 | B49764 |
|  | E24832 |  |  | 1.10E+10 | 8.40E+05 | A45395 | B50619 |
|  | E24834 | 8,3 | no treatment | 6.97E+09 | 8.29E+05 | A45399 | B50621 |
|  | E24835 | 14,1 | no treatment | 1.57E+10 | 8.83E+05 | A45400 | B50622 |
|  | E24836 | 6,1 | no treatment | 7.86E+09 | 7.70E+05 | A45401 | B50623 |
|  | E24837 | 13,1 | no treatment | 1.62E+10 | 8.34E+05 | A45402 | B50624 |
| Extraction negative controls | E23006 | - | no treatment | 8.29E+07 | 9.04E+05 | A41445 | B48111 |
|  | E23940 | - | Sodium hypochlorite | 5.96E+07 | 1.01E+06 | A43620 | B49765 |
|  | E23941 | - | no treatment | 5.99E+07 | 1.04E+06 | A43621 | B49766 |
|  | E24838 | - | no treatment | 8.20E+07 | 8.34E+05 | A45403 | B50625 |
| Library preparation negative controls | - | - | - | 6.06E+07 | 4.92E+05 | A41443 | B48110 |
|  | - | - | - | 5.54E+07 | 1.07E+06 | A43644 | B49784 |
|  | - | - | - | 7.76E+07 | 8.84E+05 | A45397 | B50620 |

#### Sequencing and raw data processing

For a preliminary assessment of ancient DNA preservation, the DNA libraries were pooled with libraries from other projects and paired-end shotgun sequencing was carried out on a MiSeq platform in 76 cycles^43^. Each run also included a spiked-in ΦX 174 library. Bases were called using *Bustard* (*Illumina*). Adapters were removed and overlapping forward and reverse reads were merged with *leeHom*^44^. Demultiplexing was performed by requiring perfect index matching with the expected combination for each library. Sequences were mapped to the human reference genome (version hg19 with additional decoy sequences as described in Prüfer *et al*., 2017^45^) using *BWA*^46^ with parameters adapted to ancient DNA^47^. For each library, sequences with identical start and end coordinates were considered PCR duplicates and collapsed into single sequences using *bam-rmdup* (https://github.com/mpieva/biohazard-tools). Sequences shorter than 35 base pairs (bp) were removed as well as those with a mapping quality lower than 25. Sequences that mapped outside the Map35_100% mappability track from Prüfer *et al*. (2014)^48^ were also removed.

#### Assessment of ancient DNA preservation

Sequences from all libraries are short, on average 52 base pairs long (excluding sequences shorter than 35bp), and show evidence of ancient DNA damage, with C-to-T substitutions reaching on average 46.7% and 41.6% on the 5’ and 3’ termini, respectively (***Table S4***). Two libraries (A41435 and A41436) stand out with fewer C-to-T substitutions (less than 40% and 35% at the 5’ and 3’ termini, respectively) and slightly different average sequence lengths. Estimates of present-day DNA contamination with *AuthentiCT*^49^, an approach based on patterns of ancient DNA damage, range from 0.1 to 6.2%, except in the two libraries noted above, as well as another library (A41438, with only 0.025% of the sequences mapped to the human genome, **Table S3**), that exhibit more than 14% present-day DNA contamination (**Table S4**). These estimates are similar to those from another method that relies on the proportion of shared derived alleles with a modern human genome to estimate present-day human DNA contamination^50^. Among the libraries with low levels of DNA contamination, we picked four libraries with exceptional DNA preservation for further sequencing (highlighted in bold in **Table S3** and **Table S4**). The qPCR estimates of the number of DNA fragments in these four libraries suggest that they would yield 3.5 to 6 copies of the nuclear genome each, if sequenced to exhaustion. In addition, 20 to 47% of the sequences in these libraries align to the human genome, so that an unusually large proportion of generated sequences is usable.

**Table S3: Sequencing summary statistics in the screened DNA libraries of Denisova 25.** The extraction and library preparation negative controls are marked in grey. Summary statistics for the four libraries selected for further sequencing are marked in bold and highlighted with a grey shading. L=length; MQ=mapping quality

|  | **indexed library ID** | **raw number of sequences** | **L≥35** | **mapped**  **(L≥35, MQ≥25)** | **unique** | **mapped fragments (%)** | **duplication rate** | **depth (screening)** | **estimated depth in library** |
| --- | --- | --- | --- | --- | --- | --- | --- | --- | --- |
| *Denisova 25* | A41435 | 687,449 | 298,855 | 21,493 | 19,348 | 7.19 | 1.00 | 0.0005 | 4.24 |
|  | A41436 | 897,594 | 420,346 | 29,171 | 26,397 | 6.94 | 1.00 | 0.0006 | 5.94 |
|  | A41437 | 1,014,605 | 521,768 | 1,122 | 1,014 | 0.22 | 1.00 | <10^-4^ | 0.05 |
|  | A41438 | 913,437 | 520,683 | 131 | 115 | 0.025 | 1.00 | <10^-4^ | 0.19 |
|  | A43614 | 1,092,517 | 340,226 | 154,872 | 140,566 | 45.52 | 1.00 | 0.0031 | 0.67 |
|  | **A43615** | **967,208** | **399,865** | **186,382** | **168,662** | **46.61** | **1.00** | **0.0039** | **5.89** |
|  | **A45393** | **725,279** | **299,016** | **136,250** | **123,686** | **45.57** | **1.00** | **0.0028** | **4.44** |
|  | A43616 | 952,572 | 455,290 | 54,741 | 49,734 | 12.02 | 1.00 | 0.0011 | 1.41 |
|  | **A43617** | **1,068,535** | **359,847** | **73,786** | **66,721** | **20.51** | **1.00** | **0.0015** | **4.20** |
|  | **A45394** | **1,234,133** | **441,937** | **89,729** | **81,079** | **20.30** | **1.00** | **0.0019** | **3.56** |
|  | A43618 | 1,058,688 | 429,138 | 7,923 | 7,210 | 1.85 | 1.00 | 0.0002 | 0.55 |
|  | A43619 | 1,224,756 | 538,985 | 32,481 | 29,405 | 6.03 | 1.00 | 0.0007 | 4.96 |
|  | A45395 | 952,515 | 424,931 | 24,526 | 22,178 | 5.77 | 1.00 | 0.0005 | 4.45 |
|  | A45399 | 937,027 | 383,303 | 26,590 | 24,096 | 6.94 | 1.00 | 0.0005 | 3.05 |
|  | A45400 | 1,104,089 | 489,537 | 15,250 | 13,742 | 3.12 | 1.00 | 0.0003 | 3.28 |
|  | A45401 | 888,192 | 359,993 | 32,256 | 29,276 | 8.96 | 1.00 | 0.0007 | 4.40 |
|  | A45402 | 1,109,396 | 506,954 | 11,095 | 9,969 | 2.19 | 1.00 | 0.0002 | 2.44 |
| Extraction negative controls | A41445 | 55,328 | 1,847 | 21 | 21 | 1.14 | 1.00 | - | - |
|  | A43620 | 60,736 | 3,001 | 22 | 21 | 0.73 | 1.00 | - | - |
|  | A43621 | 62,832 | 2,489 | 18 | 15 | 0.72 | 1.00 | - | - |
|  | A45403 | 50,894 | 2,184 | 23 | 20 | 1.05 | 1.00 | - | - |
| Library preparation negative controls | A41443 | 105,189 | 2,701 | 1 | 1 | 0.04 | 1.00 | - | - |
|  | A43644 | 80,434 | 2,957 | 2 | 2 | 0.07 | 1.00 | - | - |
|  | A45397 | 44,553 | 1,980 | 1 | 1 | 0.05 | 1.00 | - | - |

**Table S4: Summary statistics related to DNA preservation in the screened DNA libraries of Denisova 25.** Summary statistics for the four libraries selected for further sequencing are marked in bold and highlighted with a grey shading. Two present-day human DNA contamination estimates are provided, one based on the proportion of derived alleles shared with a modern human genome (“lineage assignment”)^50^, and another based on patterns of ancient DNA damage (“AuthentiCT”)^49^. Note that the reported average length was computed after excluding sequences shorter than 35bp.

| **indexed library ID** | **5' C->T (%)** | **3' C->T (%)** | **average length (bp)** | **Contamination estimates**  **(%, [95% CI])** | |
| --- | --- | --- | --- | --- | --- |
|  |  |  |  | **lineage assignment** | ***AuthentiCT*** |
| A41435 | 37.8 | 34.8 | 54.5 | 15.9 [7.4-28.4] | 14.8 [12.3-17.3] |
| A41436 | 39.7 | 33.5 | 54.4 | 18.3 [10.3-29.4] | 17.4 [15.4-19.5] |
| A41437 | 45.6 | 43.6 | 47.8 | 0.0 [0.0-75.0] | 0.1 [0.0-10.9] |
| A41438 | 61.1 | 42.3 | 47.4 | 60.4 [0.0-100.0] | 25.8 [8.8-42.8] |
| A43614 | 49.9 | 45.3 | 50.4 | 1.0 [0.0-3.1] | 0.4 [0.0-1.3] |
| **A43615** | **47.3** | **43.3** | **51.7** | **0.3 [0.0-1.9]** | **0.1 [0.0-1.1]** |
| **A45393** | **47.8** | **43.6** | **51.8** | **3.1 [1.2-5.9]** | **0.3 [0.0-1.3]** |
| A43616 | 46.9 | 43.9 | 50.5 | 3.1 [0.3-7.7] | 2.6 [1.3-3.8] |
| **A43617** | **46.3** | **42.8** | **51.6** | **4.5 [1.7-8.8]** | **4.1 [2.9-5.3]** |
| **A45394** | **45.1** | **42.0** | **53.0** | **4.6 [1.9-8.6]** | **3.5 [2.4-4.6]** |
| A43618 | 49.6 | 43.5 | 51.1 | 0.0 [0.0-15.3] | 0.1 [0.0-3.7] |
| A43619 | 47.7 | 41.9 | 52.1 | 4.7 [0.6-12.2] | 1.5 [0.0-3.4] |
| A45395 | 48.5 | 42.4 | 51.9 | 0.6 [0.0-6.9] | 0.1 [0.0-2.3] |
| A45399 | 45.9 | 42.3 | 51.1 | 3.8 [0.0-12.4] | 1.3 [0.0-3.5] |
| A45400 | 41.8 | 39.2 | 50.3 | 3.6 [0.0-17.1] | 6.2 [3.3-9.0] |
| A45401 | 48.2 | 41.9 | 51.0 | 4.1 [0.7-13.0] | 2.0 [0.1-4.0] |
| A45402 | 45.5 | 41.4 | 50.3 | 10.8 [1.6-29.4] | 4.6 [1.4-7.7] |

### Supplementary Note 3: Generation and processing of sequencing data, genotyping and filtering

#### Generation and processing of NovaSeq sequencing data

The DNA libraries selected for further sequencing were pooled. One pool contained the libraries A43615 and A45393, and another pool contained A43617 and A45394, which seem to have slightly more present-day DNA contamination than the former pool of libraries according to our preliminary assessment of the DNA preservation (**Supplementary Note 2**). Sequencing was performed on a NovaSeq platform and about 12.8 billion sequences were generated (**Table S5**). Bases were called with the Real-Time Analysis (RTA) software on the instrument. Demultiplexing was performed with *bcl2fastq* (Illumina) allowing one mismatch in the indices to recover more sequences (those with a potential sequencing error in the indices) without much risk of cross contamination with our pooling strategy. The resulting FASTQ files were converted into the BAM format with *samtools import*^51^. Adapters were removed, overlapping reads were merged and the resulting sequences were aligned as described in **Supplementary Note 2**. The data produced over multiple runs were merged per library with *samtools* v1.3.1^51^, including the data produced on the MiSeq platform for the preliminary assessment of DNA preservation in these libraries. PCR duplicates were removed and filters on mappability, mapping quality and sequence length were applied as described in **Supplementary Note 2**. Indel realignment was then performed with *GATK* v1.3-14^52^ on this merged and filtered dataset.

The sequences from all four libraries exhibit patterns of ancient DNA damage (**Table S6**, **Figure S5**) in concordance with the screening data produced on the MiSeq platform (**Supplementary Note 2**). The average length of sequenced DNA fragments is 52bp (see **Figure S6** for the length distribution). The average sequence depth per base is 23.6x on the autosomes (only counting bases with base quality ≥30; see **Table S5** for each library), in regions of the human genome to which fragments of length 35 can be aligned uniquely (map35_100 track). The sequence depth on the X chromosome (and the Y chromosome) is half that of the autosomes, indicating that *Denisova 25* was a male. Estimates of present-day human DNA contamination using the lineage assignment method and *AuthentiCT* (**Table S6**) are consistent with those obtained in the screening. Taking advantage that *Denisova 25* was a male, we also estimated present-day human DNA contamination on the X chromosome using the two-consensus method from *contaminationX*^53^. Assuming that the contaminant has likely European ancestry, we used the allele frequencies of the CEU reference panel (provided with *contaminationX* and downloaded from <https://github.com/sapfo/contaminationX/tree/master/HapMapFreqs>). Beyond the filters described above, we excluded the pseudo-autosomal regions, variants with a minor allele frequency below 0.05, bases with a base quality below 20, and focused on positions with a sequence depth between 2 and 20x. This method provided similar estimates of present-day human DNA contamination (**Table S6**).

**Table S5: Sequencing summary statistics for the deeply sequenced DNA libraries of Denisova 25.** “On target” refers to the number of sequences that align to regions of the human genome to which sequences of length 35bp can be aligned uniquely (map35_100 track). L=length; MQ=mapping quality

| **indexed library ID** | **raw number of sequences** | **L≥35** | **mapped**  **(L≥35, MQ≥25)** | **mapped fragments (%)** | **on target** | **unique** | **duplication rate** | **depth** | | |
| --- | --- | --- | --- | --- | --- | --- | --- | --- | --- | --- |
|  |  |  |  |  |  |  |  | **autosomes** | **X** | **Y** |
| A43615 | 3,440,256,904 | 1,400,292,404 | 646,178,185 | 46.15 | 584,188,809 | 391,150,665 | 1.49 | 9.12 | 4.78 | 4.80 |
| A45393 | 3,711,212,665 | 1,539,329,762 | 689,457,578 | 44.79 | 623,444,009 | 360,217,727 | 1.73 | 8.49 | 4.44 | 4.46 |
| A43617 | 3,791,000,525 | 1,271,720,839 | 241,051,950 | 18.96 | 217,749,232 | 160,888,561 | 1.35 | 3.75 | 1.97 | 1.95 |
| A45394 | 1,904,131,260 | 690,051,274 | 131,812,462 | 19.10 | 118,755,635 | 94,648,258 | 1.25 | 2.27 | 1.19 | 1.18 |
| **Total** | **12,846,601,354** | **4,901,394,279** | **1,708,500,175** | **34.86** | **1,544,137,685** | **921,721,778** | **-** | **23.63** | **12.38** | **12.39** |

**Table S6: Summary statistics related to DNA preservation in the deeply sequenced DNA libraries of Denisova 25.** Three present-day (human) DNA contamination estimates are provided, one based on the proportion of derived alleles shared with a modern human genome (“lineage assignment”)^50^, another based on the observation of alternative alleles on the haploid X chromosome (“contaminationX”)^53^ and another based on patterns of ancient DNA damage (“AuthentiCT”)^49^. Note that the reported average length was computed after excluding sequences shorter than 35bp. The screening data produced on the MiSeq platform were included.

| **indexed library ID** | **5' C->T (%)** | **3' C->T (%)** | **average length (bp)** | **Contamination estimates**  **(%, [95% CI])** | | |
| --- | --- | --- | --- | --- | --- | --- |
|  |  |  |  | **lineage assignment** | **contaminationX** | **AuthentiCT** |
| A43615 | 47.8 | 42.6 | 51.6 | 2.80 [2.76-2.84] | 1.91 [1.89-1.93] | 0.1 [0.0-1.1] |
| A45393 | 47.9 | 44.0 | 52.1 | 2.83 [2.80-2.88] | 2.03 [2.01-2.05] | 0.7 [0.0-1.7] |
| A43617 | 45.8 | 41.9 | 51.7 | 4.41 [4.35-4.48] | 3.55 [3.50-3.60] | 3.4 [2.4-4.4] |
| A45394 | 45.9 | 42.1 | 53.2 | 4.63 [4.54-4.72] | 3.27 [3.16-3.38] | 3.6 [2.6-4.6] |
| ***Denisova 25*** | **47.3** | **42.9** | **51.9** | **3.24 [3.22-3.27]** | **2.55 [2.53-2.56]** | **1.3 [0.2-2.3]** |

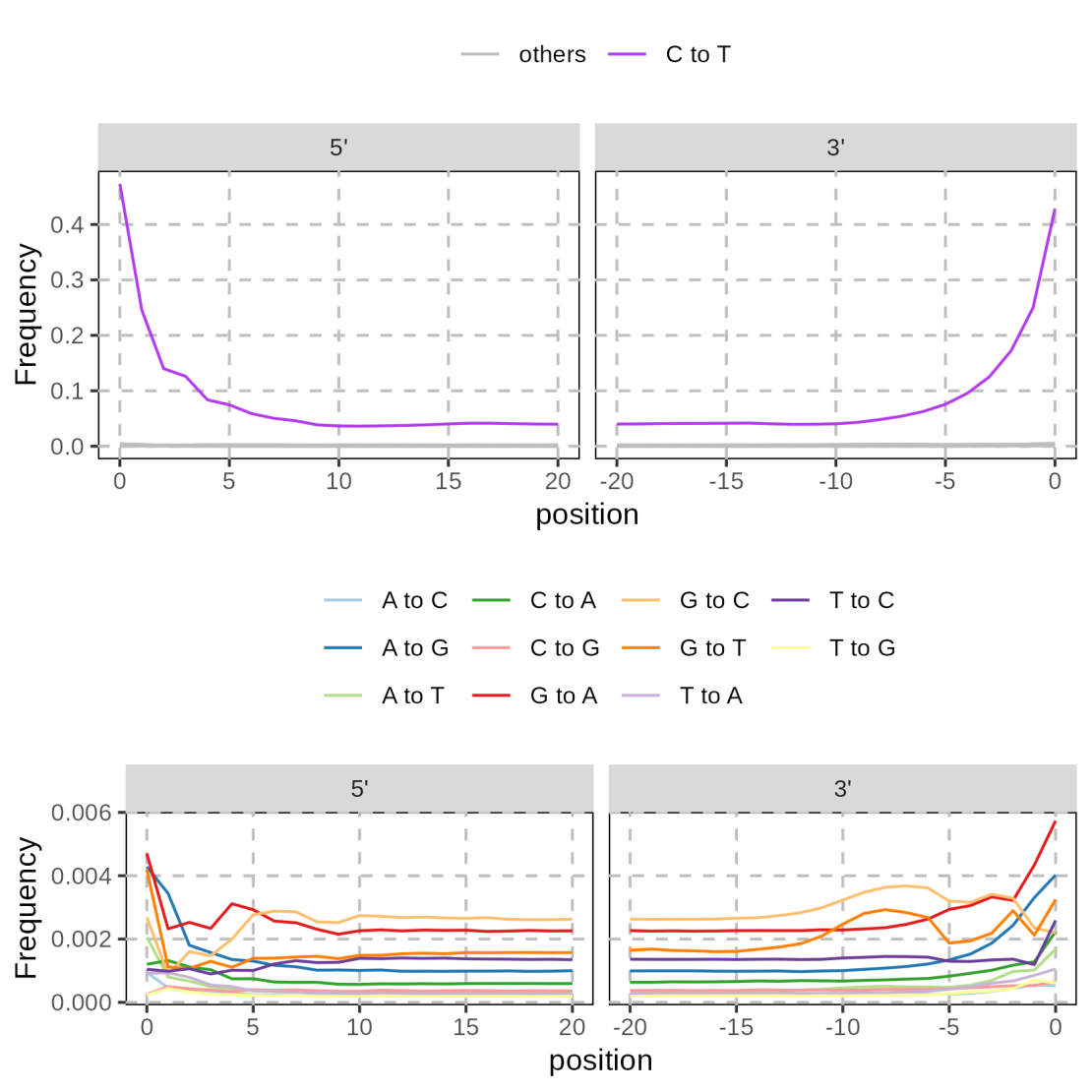

**Figure S5: Base substitution frequencies at the end of Denisova 25 sequences relative to the bases in the human reference genome, hg19.** C-to-T substitutions are shown in the top panels, whereas other substitutions, which occur at lower rates, are shown below at a finer scale. All sequences of at least 35 bases aligning within the Map35_100% mappability track with a mapping quality of at least 25 were used (from the A43615, A43617, A45393 and A45394 libraries).

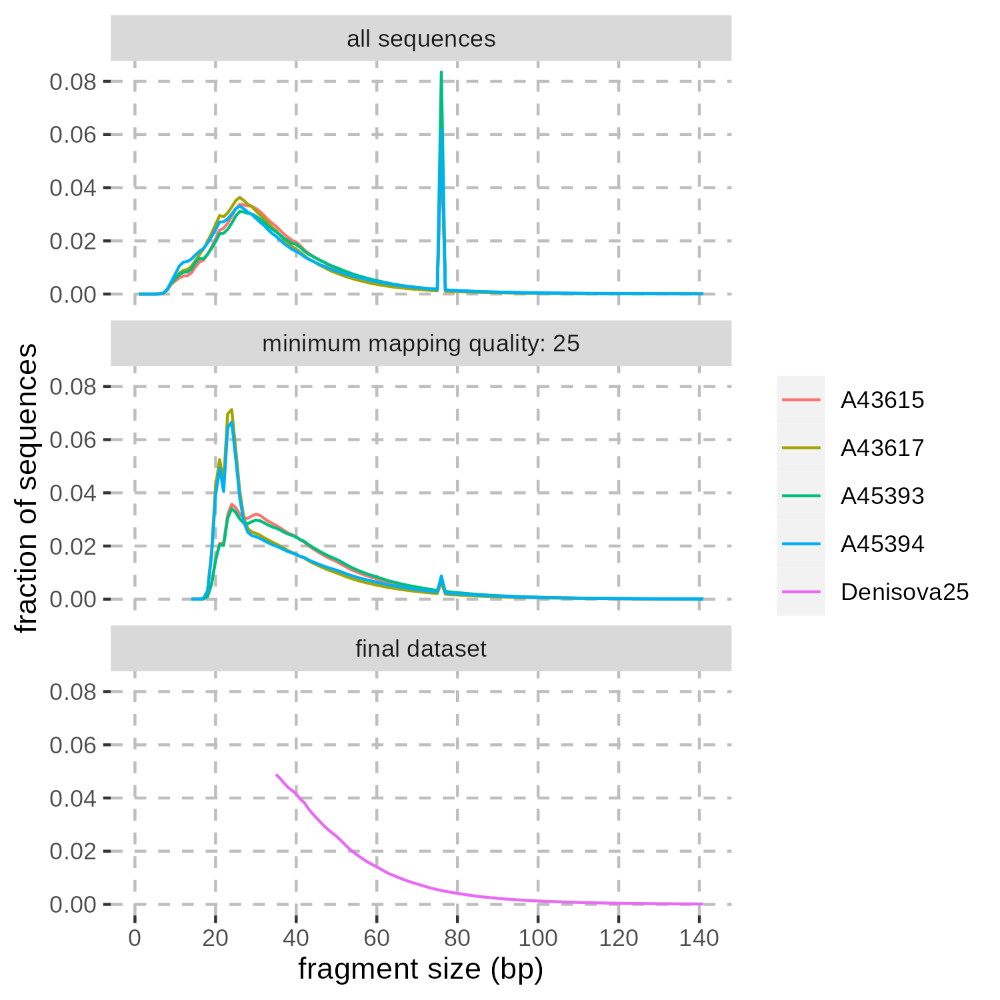

**Figure S6: Fragment size distribution** **in the deeply sequenced DNA libraries of Denisova 25**. The top panel corresponds to all sequences after removing duplicates, the middle panel corresponds to those sequences aligning to the human reference genome, hg19, within the Map35_100% mappability track with a mapping quality of at least 25, and the bottom panel corresponds to only those that are also merged from overlapping paired-end reads and are at least 35bp long. The peak at 76bp in the upper two panels corresponds to paired-end reads that could not be merged. Peaks between 20 and 30bp in the middle panel are due to spurious alignments of short reads, which were filtered out in the final dataset.

#### Genotyping

Input files for the ancient DNA genotype caller *snpAD*^54^ (version 0.2.1) were generated for each chromosome using bases with a quality of at least 30 (from sequences of at least 35 bases with a mapping quality of at least 25, as described above). Positions outside the Map35_100% mappability track from Prüfer *et al*. (2014)^48^ were removed. Genotype frequencies and error rates were then estimated independently for each chromosome (**Table S7**) and used to call the most likely genotype at each position (**Table S8** and **Table S9**).

**Table S7: Estimated prior probabilities for each genotype per chromosome**. Prior probabilities for heterozygous genotypes on the haploid X and Y chromosomes were included for completeness. The frequencies of each genotype after genotype calling are presented in **Table S8**.

| **Chr.** | **AA** | **CC** | **GG** | **TT** | **AC** | **AG** | **AT** | **CG** | **CT** | **GT** |
| --- | --- | --- | --- | --- | --- | --- | --- | --- | --- | --- |
| 1 | 0.29 | 0.21 | 0.21 | 0.29 | 1.9E-05 | 6.2E-05 | 1.9E-05 | 1.7E-05 | 6.1E-05 | 1.8E-05 |
| 2 | 0.30 | 0.20 | 0.20 | 0.30 | 2.2E-05 | 7.4E-05 | 2.3E-05 | 2.1E-05 | 7.3E-05 | 2.2E-05 |
| 3 | 0.30 | 0.20 | 0.20 | 0.30 | 2.2E-05 | 6.8E-05 | 2.3E-05 | 2.0E-05 | 6.8E-05 | 2.2E-05 |
| 4 | 0.31 | 0.19 | 0.19 | 0.31 | 1.9E-05 | 5.3E-05 | 2.1E-05 | 1.4E-05 | 5.5E-05 | 1.8E-05 |
| 5 | 0.30 | 0.19 | 0.20 | 0.30 | 2.6E-05 | 8.7E-05 | 2.7E-05 | 2.4E-05 | 8.8E-05 | 2.6E-05 |
| 6 | 0.31 | 0.19 | 0.19 | 0.31 | 2.5E-05 | 8.4E-05 | 2.5E-05 | 2.2E-05 | 8.5E-05 | 2.6E-05 |
| 7 | 0.30 | 0.20 | 0.20 | 0.30 | 2.5E-05 | 8.3E-05 | 2.6E-05 | 2.2E-05 | 8.0E-05 | 2.4E-05 |
| 8 | 0.30 | 0.20 | 0.20 | 0.30 | 2.7E-05 | 8.5E-05 | 2.6E-05 | 2.3E-05 | 8.6E-05 | 2.5E-05 |
| 9 | 0.29 | 0.21 | 0.21 | 0.29 | 2.6E-05 | 8.3E-05 | 2.6E-05 | 2.6E-05 | 8.6E-05 | 2.7E-05 |
| 10 | 0.29 | 0.21 | 0.21 | 0.29 | 1.8E-05 | 6.1E-05 | 1.9E-05 | 1.6E-05 | 6.2E-05 | 1.8E-05 |
| 11 | 0.29 | 0.21 | 0.21 | 0.29 | 2.8E-05 | 9.3E-05 | 2.7E-05 | 2.5E-05 | 9.4E-05 | 2.7E-05 |
| 12 | 0.30 | 0.20 | 0.20 | 0.30 | 2.1E-05 | 7.0E-05 | 2.3E-05 | 1.8E-05 | 6.6E-05 | 2.1E-05 |
| 13 | 0.31 | 0.19 | 0.19 | 0.31 | 2.9E-05 | 9.2E-05 | 3.1E-05 | 2.3E-05 | 9.2E-05 | 2.9E-05 |
| 14 | 0.30 | 0.20 | 0.20 | 0.30 | 2.0E-05 | 7.4E-05 | 2.0E-05 | 1.9E-05 | 7.0E-05 | 2.2E-05 |
| 15 | 0.29 | 0.21 | 0.21 | 0.29 | 2.6E-05 | 9.1E-05 | 2.5E-05 | 2.7E-05 | 9.2E-05 | 2.7E-05 |
| 16 | 0.28 | 0.22 | 0.22 | 0.28 | 2.1E-05 | 6.8E-05 | 1.9E-05 | 2.7E-05 | 7.2E-05 | 2.1E-05 |
| 17 | 0.27 | 0.23 | 0.23 | 0.27 | 2.5E-05 | 9.3E-05 | 2.2E-05 | 2.6E-05 | 9.4E-05 | 2.4E-05 |
| 18 | 0.30 | 0.20 | 0.20 | 0.30 | 2.1E-05 | 6.9E-05 | 2.4E-05 | 1.8E-05 | 6.8E-05 | 2.1E-05 |
| 19 | 0.25 | 0.25 | 0.25 | 0.25 | 2.0E-05 | 7.3E-05 | 2.0E-05 | 2.3E-05 | 7.1E-05 | 2.1E-05 |
| 20 | 0.28 | 0.22 | 0.22 | 0.28 | 1.8E-05 | 6.9E-05 | 2.0E-05 | 2.1E-05 | 7.1E-05 | 2.0E-05 |
| 21 | 0.30 | 0.20 | 0.20 | 0.30 | 3.2E-05 | 1.1E-04 | 3.6E-05 | 2.9E-05 | 1.1E-04 | 3.3E-05 |
| 22 | 0.26 | 0.24 | 0.24 | 0.26 | 3.4E-05 | 1.4E-04 | 2.4E-05 | 4.6E-05 | 1.5E-04 | 3.4E-05 |
| X | 0.31 | 0.19 | 0.19 | 0.31 | 3.6E-06 | 6.5E-06 | 6.3E-06 | 2.8E-06 | 5.1E-06 | 3.7E-06 |
| Y | 0.31 | 0.19 | 0.19 | 0.31 | 2.2E-05 | 3.5E-05 | 2.3E-05 | 1.5E-05 | 3.7E-05 | 1.4E-05 |

**Table S8: Frequencies of each genotype called per chromosome**. The frequencies of heterozygous genotypes on the haploid X and Y chromosomes were included for completeness (including the pseudoautosomal regions of the X chromosome). These frequencies were computed after applying the standard filters described in this supplementary note.

| **Chr.** | **AA** | **CC** | **GG** | **TT** | **AC** | **AG** | **AT** | **CG** | **CT** | **GT** |
| --- | --- | --- | --- | --- | --- | --- | --- | --- | --- | --- |
| 1 | 0.29 | 0.21 | 0.21 | 0.29 | 1.5E-05 | 5.4E-05 | 1.2E-05 | 1.4E-05 | 5.4E-05 | 1.4E-05 |
| 2 | 0.30 | 0.20 | 0.20 | 0.30 | 1.9E-05 | 6.8E-05 | 1.7E-05 | 1.9E-05 | 6.8E-05 | 1.8E-05 |
| 3 | 0.30 | 0.20 | 0.20 | 0.30 | 1.7E-05 | 6.4E-05 | 1.6E-05 | 1.7E-05 | 6.3E-05 | 1.8E-05 |
| 4 | 0.31 | 0.19 | 0.19 | 0.31 | 1.5E-05 | 4.9E-05 | 1.3E-05 | 1.2E-05 | 5.0E-05 | 1.4E-05 |
| 5 | 0.30 | 0.19 | 0.20 | 0.30 | 2.2E-05 | 8.2E-05 | 2.0E-05 | 2.2E-05 | 8.3E-05 | 2.2E-05 |
| 6 | 0.31 | 0.19 | 0.19 | 0.30 | 2.1E-05 | 7.8E-05 | 1.8E-05 | 1.9E-05 | 7.8E-05 | 2.2E-05 |
| 7 | 0.30 | 0.20 | 0.20 | 0.30 | 2.0E-05 | 7.5E-05 | 1.9E-05 | 2.0E-05 | 7.4E-05 | 2.0E-05 |
| 8 | 0.30 | 0.20 | 0.20 | 0.30 | 2.2E-05 | 7.9E-05 | 1.9E-05 | 2.1E-05 | 8.1E-05 | 2.1E-05 |
| 9 | 0.29 | 0.21 | 0.21 | 0.29 | 2.2E-05 | 7.6E-05 | 1.9E-05 | 2.4E-05 | 8.1E-05 | 2.3E-05 |
| 10 | 0.29 | 0.21 | 0.21 | 0.29 | 1.4E-05 | 5.3E-05 | 1.2E-05 | 1.4E-05 | 5.5E-05 | 1.4E-05 |
| 11 | 0.29 | 0.21 | 0.21 | 0.29 | 2.3E-05 | 8.6E-05 | 1.9E-05 | 2.3E-05 | 8.7E-05 | 2.4E-05 |
| 12 | 0.30 | 0.20 | 0.20 | 0.30 | 1.8E-05 | 6.4E-05 | 1.6E-05 | 1.6E-05 | 6.2E-05 | 1.8E-05 |
| 13 | 0.31 | 0.19 | 0.19 | 0.31 | 2.4E-05 | 8.7E-05 | 2.2E-05 | 2.2E-05 | 8.7E-05 | 2.5E-05 |
| 14 | 0.30 | 0.20 | 0.20 | 0.30 | 1.6E-05 | 6.7E-05 | 1.3E-05 | 1.7E-05 | 6.4E-05 | 1.7E-05 |
| 15 | 0.29 | 0.21 | 0.21 | 0.29 | 2.3E-05 | 8.6E-05 | 1.9E-05 | 2.6E-05 | 8.7E-05 | 2.2E-05 |
| 16 | 0.28 | 0.22 | 0.22 | 0.28 | 1.6E-05 | 5.9E-05 | 1.3E-05 | 2.3E-05 | 6.2E-05 | 1.6E-05 |
| 17 | 0.27 | 0.23 | 0.23 | 0.27 | 2.1E-05 | 8.1E-05 | 1.6E-05 | 2.3E-05 | 8.2E-05 | 2.0E-05 |
| 18 | 0.30 | 0.20 | 0.20 | 0.30 | 1.7E-05 | 6.4E-05 | 1.6E-05 | 1.5E-05 | 6.3E-05 | 1.7E-05 |
| 19 | 0.25 | 0.25 | 0.25 | 0.25 | 1.6E-05 | 6.1E-05 | 1.1E-05 | 1.9E-05 | 6.1E-05 | 1.7E-05 |
| 20 | 0.28 | 0.22 | 0.22 | 0.28 | 1.5E-05 | 6.2E-05 | 1.3E-05 | 1.8E-05 | 6.4E-05 | 1.7E-05 |
| 21 | 0.30 | 0.20 | 0.20 | 0.30 | 2.6E-05 | 1.0E-04 | 2.5E-05 | 2.5E-05 | 9.8E-05 | 2.4E-05 |
| 22 | 0.26 | 0.24 | 0.24 | 0.26 | 2.8E-05 | 1.3E-04 | 1.9E-05 | 4.2E-05 | 1.3E-04 | 2.9E-05 |
| X | 0.31 | 0.19 | 0.19 | 0.31 | 1.8E-06 | 2.1E-06 | 2.7E-06 | 1.0E-06 | 2.1E-06 | 2.1E-06 |
| Y | 0.31 | 0.19 | 0.19 | 0.31 | 8.1E-06 | 9.2E-06 | 4.9E-06 | 4.5E-06 | 1.0E-05 | 4.0E-06 |

**Table S9: Number of each genotype called per chromosome.** The number of heterozygous genotypes on the haploid X and Y chromosomes were included for completeness and include those in the pseudo autosomal regions of both chromosomes. Only those genotypes that pass the standard filters described in this supplementary note were considered.

| **Chr.** | **AA** | **CC** | **GG** | **TT** | **AC** | **AG** | **AT** | **CG** | **CT** | **GT** |
| --- | --- | --- | --- | --- | --- | --- | --- | --- | --- | --- |
| 1 | 40043771 | 28145327 | 28145077 | 40060704 | 2040 | 7411 | 1663 | 1911 | 7381 | 1959 |
| 2 | 45263030 | 29894275 | 29917788 | 45319049 | 2791 | 10266 | 2495 | 2900 | 10183 | 2765 |
| 3 | 37812307 | 24321145 | 24326554 | 37782943 | 2169 | 7903 | 1981 | 2132 | 7863 | 2232 |
| 4 | 36760910 | 21981615 | 21972187 | 36745082 | 1733 | 5753 | 1507 | 1459 | 5923 | 1690 |
| 5 | 33917883 | 21704538 | 21727903 | 33935831 | 2432 | 9123 | 2181 | 2396 | 9271 | 2477 |
| 6 | 32154903 | 20533589 | 20550811 | 32144438 | 2178 | 8204 | 1879 | 1995 | 8248 | 2285 |
| 7 | 27246639 | 18096110 | 18074621 | 27229012 | 1841 | 6823 | 1708 | 1811 | 6730 | 1802 |
| 8 | 27383887 | 17959905 | 17918147 | 27284753 | 1967 | 7137 | 1704 | 1916 | 7320 | 1925 |
| 9 | 20288483 | 14144804 | 14126598 | 20232453 | 1507 | 5225 | 1302 | 1622 | 5584 | 1563 |
| 10 | 23492286 | 16532421 | 16519446 | 23483233 | 1119 | 4261 | 924 | 1125 | 4370 | 1090 |
| 11 | 23684462 | 16810941 | 16820214 | 23695083 | 1902 | 6967 | 1576 | 1846 | 7047 | 1935 |
| 12 | 23937231 | 16013101 | 16007681 | 23953811 | 1401 | 5139 | 1286 | 1286 | 4939 | 1413 |
| 13 | 19183696 | 11700373 | 11700803 | 19243207 | 1506 | 5366 | 1373 | 1333 | 5377 | 1516 |
| 14 | 16264336 | 11000320 | 11036761 | 16341498 | 852 | 3646 | 723 | 932 | 3514 | 939 |
| 15 | 13941080 | 9997642 | 9983509 | 13905663 | 1100 | 4094 | 900 | 1223 | 4168 | 1069 |
| 16 | 12336589 | 9885864 | 9932608 | 12383294 | 729 | 2609 | 596 | 1029 | 2770 | 735 |
| 17 | 11942628 | 9913766 | 9880499 | 11936147 | 896 | 3554 | 708 | 987 | 3600 | 875 |
| 18 | 14872979 | 9649496 | 9659740 | 14879926 | 846 | 3122 | 767 | 757 | 3090 | 855 |
| 19 | 6554333 | 6369583 | 6391406 | 6528514 | 404 | 1565 | 288 | 489 | 1567 | 427 |
| 20 | 10671941 | 8450196 | 8484165 | 10762976 | 575 | 2391 | 489 | 673 | 2442 | 634 |
| 21 | 6300804 | 4282393 | 4284441 | 6281018 | 542 | 2150 | 520 | 521 | 2068 | 515 |
| 22 | 5014804 | 4707596 | 4712154 | 4994244 | 544 | 2518 | 364 | 811 | 2594 | 571 |
| X | 24402946 | 15449606 | 15459111 | 24396785 | 146 | 170 | 215 | 80 | 170 | 165 |
| Y | 1370802 | 859589 | 864637 | 1372337 | 36 | 41 | 22 | 20 | 46 | 18 |

#### General filters and quality controls

A set of standard filters were applied to remove potential alignment errors. We excluded simple repeats as identified by TandemRepeatFinder^55^, positions overlapping indels called with GATK v1.3-14^52^, as well as positions with a coverage that falls outside the central 95% of the coverage distribution per GC category (**Figure S7**). Positions covered by less than 10 sequences were further excluded for the autosomes, but not for the haploid X- and Y-chromosomes given their lower sequence coverage.

After filtering, 1,670,458,986 genotypes remained, which represents 59Mb (3.4%) less compared to the 30-fold coverage *Denisova 3* genome (the same genotype caller, snpAD, and standard filters were previously applied to this genome^45^; see **Table S10** for a comparison with other high-coverage archaic human genomes). These include 351,710 heterozygous sites on the autosomes, compared to 329,997 for the *Denisova 3* genome. After removing pseudo-autosomal regions (PAR1: X:60001-2699520 and PAR2: X:154931044-155260560), there are 815 heterozygous calls out of 79,285,026 genotypes on the X chromosome. As *Denisova 25* is male, this represents one error every 100,000bp for this chromosome. Note that the error rate should be lower for the autosomes as the autosomal sequence coverage is two-fold higher and we applied an additional coverage cutoff of minimum 10 sequences per position. Heterozygous sites on the X and Y chromosome were filtered out in subsequent analyses, unless stated otherwise (**Supplementary Note 5**).

Genotype qualities are lower for *Denisova 25* than for *Denisova 3*, as expected given the lower sequence coverage, particularly for the heterozygous transition calls (**Figure S8**). Yet, the proportions of each substitution type at heterozygous sites in *Denisova 25* are similar to those in the high-coverage genomes of other archaic humans and one present-day Mbuti individual (**Figure S9**), even for heterozygous transitions. Therefore, there is no evidence of more errors due to deamination-derived substitutions in *Denisova 25* compared to previously published archaic human genomes. The allelic imbalance at heterozygous positions, where we expect half of the sequences to carry the alternative allele, is also not different from that of *Denisova 3* (**Figure S10**).

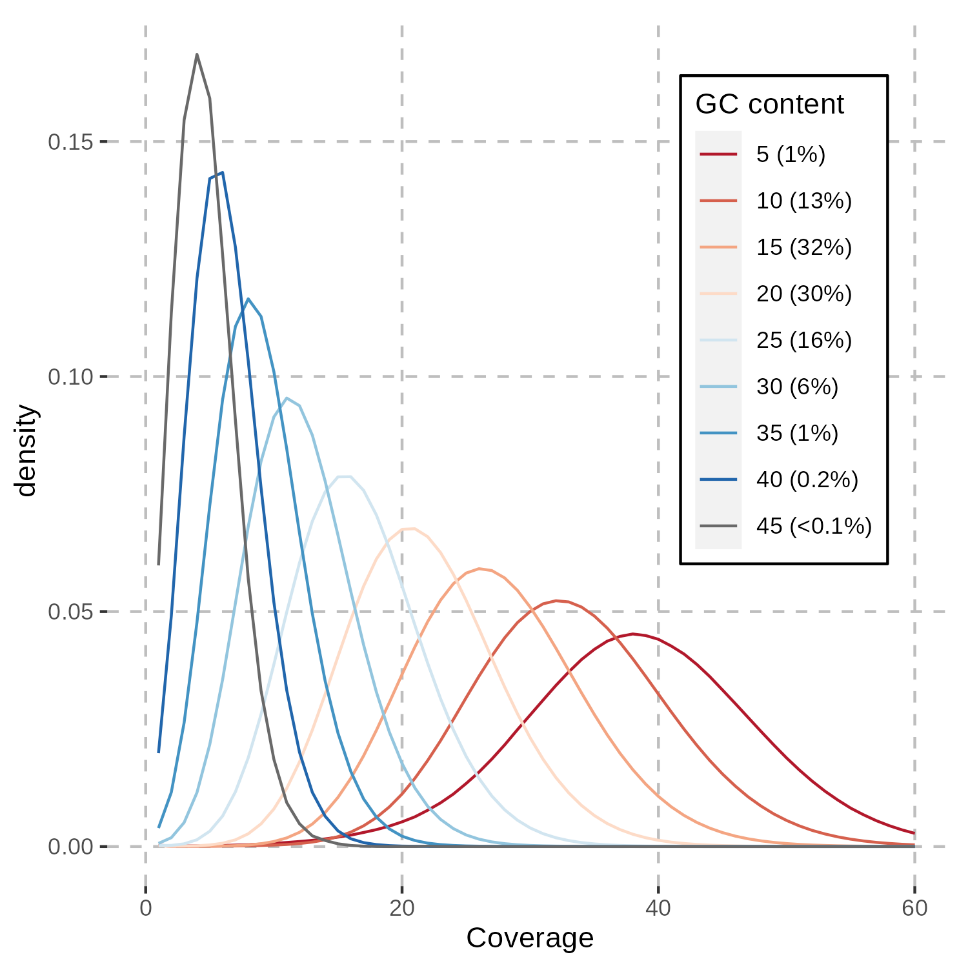

**Figure S7: Coverage conditional on the local GC content** (25bp upstream and downstream of each site, i.e., in 51bp sliding windows along the genome). The numbers in the legend correspond to the number of G or C bases per 51bp window (5 indicates 5-9 G or C bases, 10 indicates 10-14, etc.) and the percentage of all mappable bases belonging to each GC content category.

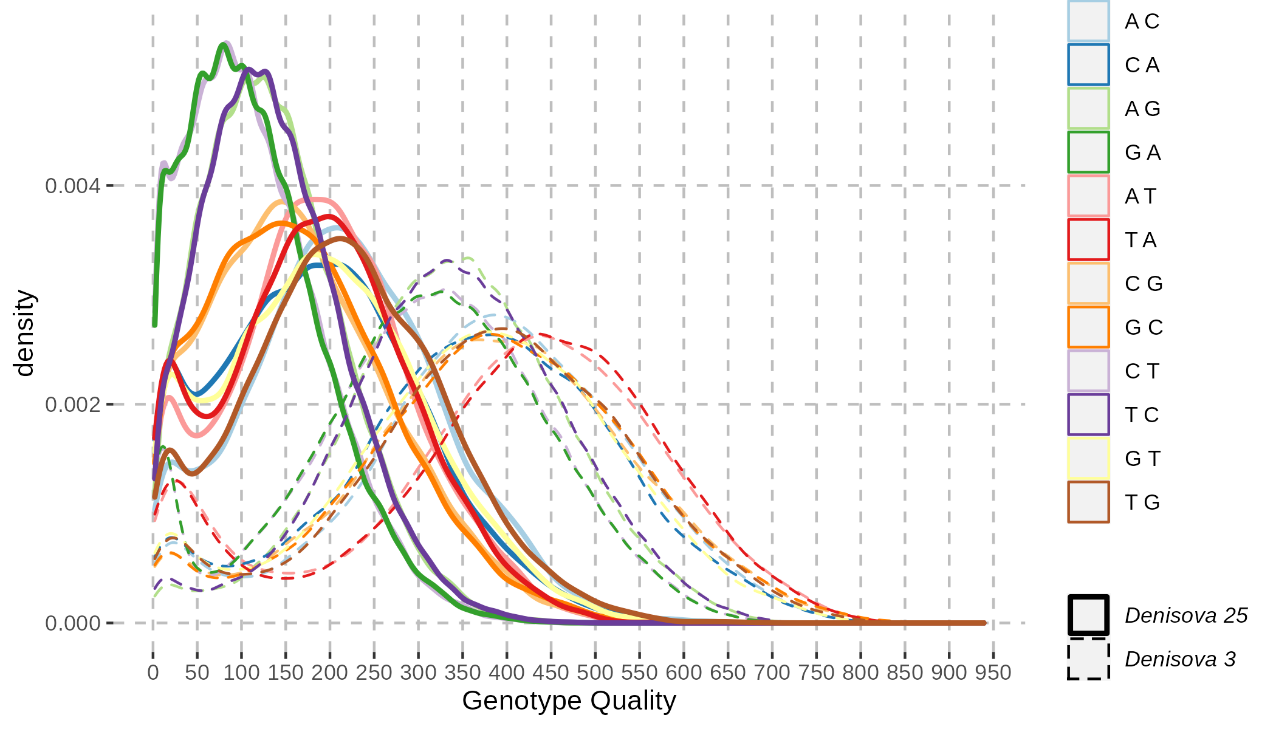

**Figure S8: Distribution of genotype qualities per substitution type (different colours) at heterozygous positions in the genomes of Denisova 25 and Denisova 3.** The first base in the legend corresponds to the reference allele, whereas the second base corresponds to the alternative allele.

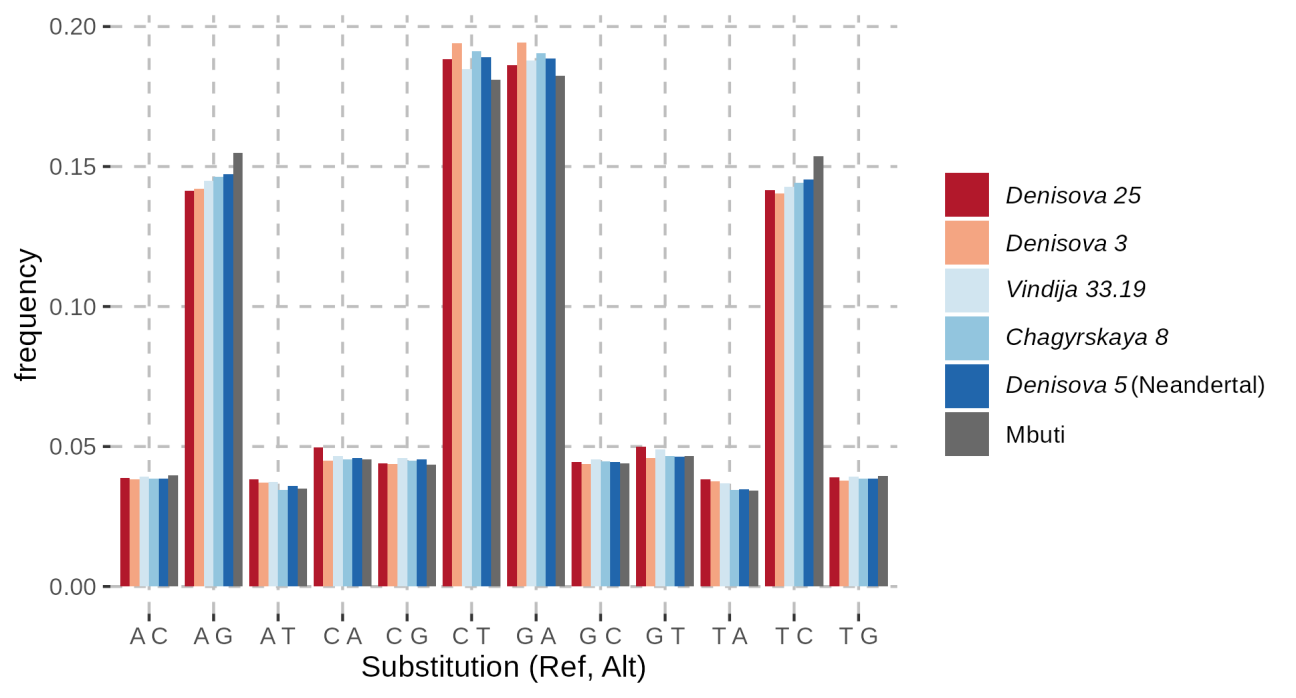

**Figure S9: Frequency of heterozygous genotypes in high coverage genomes.** A high coverage genome from a present-day Mbuti individual (SS6004471 or HGDP00982)^48^ was included for comparison with Denisova 25 and other archaic human genomes.

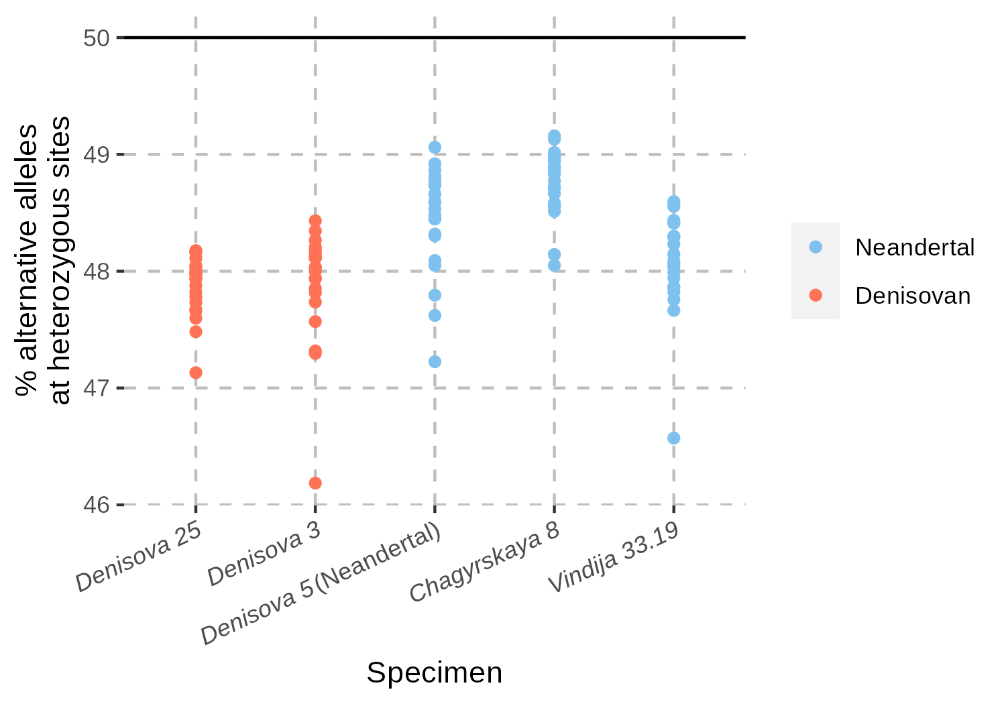

**Figure S10: Allelic imbalance at heterozygous positions called with snpAD in high-coverage archaic human genomes.** We computed the proportion of sequences that carry the alternative allele (with a base quality of at least 30) among those sequences that are at least 35bp and align to the human genome with a mapping quality of at least 25. We only considered sequences overlapping heterozygous positions that pass the minimal set of filters described in this Supplementary Note. Each dot represents the computed proportion for an autosome.

**Table S10: Number of genotypes called per chromosome for each high-coverage archaic human genome.** Only those genotypes that pass the standard filters described in this supplementary note were considered.

| **Chromosome** | **Altai Neandertal (*Denisova 5*)** | ***Vindija 33.19*** | ***Chagyrskaya 8*** | ***Denisova 3*** | ***Denisova 25*** |
| --- | --- | --- | --- | --- | --- |
| 1 | 137,765,719 | 137,406,985 | 135,859,530 | 136,957,354 | 136,417,244 |
| 2 | 151,729,000 | 151,547,577 | 149,862,249 | 151,138,410 | 150,425,542 |
| 3 | 125,404,994 | 125,225,876 | 123,873,415 | 124,799,608 | 124,267,229 |
| 4 | 118,251,176 | 118,185,270 | 116,953,088 | 117,914,451 | 117,477,859 |
| 5 | 112,211,096 | 112,132,343 | 110,834,393 | 111,795,737 | 111,314,035 |
| 6 | 106,148,559 | 106,105,861 | 104,860,113 | 105,852,040 | 105,408,530 |
| 7 | 91,459,558 | 91,362,676 | 90,264,428 | 91,008,365 | 90,667,097 |
| 8 | 91,290,377 | 91,188,319 | 90,233,110 | 90,907,937 | 90,568,661 |
| 9 | 69,417,366 | 69,252,030 | 68,431,704 | 69,009,598 | 68,809,141 |
| 10 | 80,826,273 | 80,660,966 | 79,753,747 | 80,421,672 | 80,040,275 |
| 11 | 81,916,633 | 81,655,502 | 80,755,694 | 81,293,513 | 81,031,973 |
| 12 | 80,503,183 | 80,419,065 | 79,441,580 | 80,189,571 | 79,927,288 |
| 13 | 62,252,773 | 62,217,521 | 61,579,328 | 62,039,994 | 61,844,550 |
| 14 | 55,101,494 | 55,056,780 | 54,432,506 | 54,877,615 | 54,653,521 |
| 15 | 48,299,692 | 48,164,970 | 47,594,656 | 48,062,914 | 47,840,448 |
| 16 | 45,007,869 | 44,801,619 | 44,207,224 | 44,529,265 | 44,546,823 |
| 17 | 44,174,028 | 43,931,205 | 43,330,088 | 43,731,584 | 43,683,660 |
| 18 | 49,464,460 | 49,427,725 | 48,900,547 | 49,268,539 | 49,071,578 |
| 19 | 26,229,900 | 25,944,323 | 25,411,631 | 25,680,804 | 25,848,576 |
| 20 | 38,827,750 | 38,672,275 | 38,252,717 | 38,549,727 | 38,376,482 |
| 21 | 21,322,661 | 21,304,193 | 21,058,290 | 21,213,340 | 21,154,972 |
| 22 | 19,797,946 | 19,636,250 | 19,429,710 | 19,512,261 | 19,436,200 |
| X | 80,962,031 | 81,025,732 | 80,040,723 | 80,703,574 | 79,709,394 |
| Y | - | - | - | - | 4,467,548 |

### Supplementary Note 4: The mitochondrial genome of *Denisova 25*

#### Mitochondrial DNA capture, sequencing and processing

An aliquot of each of the DNA libraries previously prepared (**Supplementary Note 2**) was enriched for human mitochondrial DNA fragments using a bead-based hybridization capture method^56^. Capture probes (described in Fu, *et al*., 2013^57^) covering the complete mitochondrial genome of the revised Cambridge Reference Sequence^58^ (rCRS, Genbank acc. no. NC_012920.1) were used as baits. Two successive rounds of automated hybridization capture were performed on the automated liquid handling platform (Agilent Bravo NGS Workstation) as previously described^59^. The enriched DNA libraries, along with negative controls, were pooled with DNA libraries from other projects, sequenced on a MiSeq platform, and processed as described for the shotgun sequencing in **Supplementary Note 2**. However, the sequences were only aligned to the mitochondrial genome reference, rCRS, instead of the whole genome. Summary statistics of the sequencing of these enriched DNA libraries are presented in **Table S11**.

The level of present-day human DNA contamination was estimated from the proportion of sequences that carry the modern human allele at positions where the mitochondrial genomes of 312 present-day humans^60^ (including the rCRS) differ from that of 4 Denisovans^7,8,61,62^ and a hominin from Sima de los Huesos^63^. To avoid errors due to cytosine deamination, sequences that align in the forward or reverse orientation were excluded if one of the two possible states was C or G, respectively. The estimates of present-day human DNA contamination varied between 1.6 to 63.0% across libraries (**Table S12**). For further analysis, we selected the 9 libraries that showed less than 10% contamination.

**Table S11: Sequencing summary statistics for the DNA libraries of Denisova 25 enriched for mitochondrial DNA.** The libraries highlighted in grey are those with minimal present-day human DNA contamination that were used to reconstruct the consensus mitochondrial sequence of Denisova 25. Sequences were aligned to the rCRS. L=length; MQ=mapping quality

| **captured library ID** | **raw number of sequences** | **L≥35** | **mapped**  **(L≥35, MQ≥25)** | **unique** | **mapped fragments (%)** | **duplication rate** | **average depth** |
| --- | --- | --- | --- | --- | --- | --- | --- |
| B48024 | 224,004 | 171,344 | 115,724 | 79,301 | 67.54 | 1.46 | 299 |
| B48025 | 206,621 | 163,080 | 105,330 | 77,106 | 64.59 | 1.37 | 302 |
| B48026 | 329,330 | 251,462 | 154,051 | 26,895 | 61.26 | 5.73 | 81 |
| **B48027** | **238,358** | **191,074** | **103,883** | **43,703** | **54.37** | **2.38** | **139** |
| **B49759** | **187,510** | **122,659** | **88,732** | **30,939** | **72.34** | **2.87** | **102** |
| **B49760** | **288,028** | **214,221** | **142,341** | **85,820** | **66.45** | **1.66** | **298** |
| **B49761** | **252,265** | **180,239** | **120,810** | **56,095** | **67.03** | **3.15** | **184** |
| **B49762** | **498,090** | **359,015** | **253,871** | **122,158** | **70.71** | **2.08** | **418** |
| **B49763** | **249,694** | **173,427** | **106,842** | **31,756** | **61.61** | **3.36** | **103** |
| B49764 | 193,646 | 153,691 | 87,573 | 60,677 | 56.98 | 1.44 | 218 |
| **B50617** | **446,154** | **335,536** | **215,501** | **103,045** | **64.23** | **2.09** | **352** |
| **B50618** | **633,818** | **463,083** | **307,389** | **121,793** | **66.38** | **2.52** | **431** |
| **B50619** | **418,611** | **310,607** | **170,754** | **72,862** | **54.97** | **2.34** | **249** |
| B50621 | 509,830 | 370,552 | 225,504 | 82,020 | 60.86 | 2.75 | 279 |
| B50622 | 353,306 | 270,797 | 156,960 | 60,742 | 57.96 | 2.58 | 210 |
| B50623 | 383,818 | 295,513 | 178,254 | 86,298 | 60.32 | 2.07 | 299 |
| B50624 | 504,332 | 406,385 | 219,438 | 61,035 | 54.00 | 3.60 | 218 |
| **analysed**  **dataset** | **3,212,528** | **2,349,861** | **1,510,123** | **668,171** | **64.26** | **2.26** | **2,275** |

**Table S12: Present-day human DNA contamination estimates in the DNA libraries of Denisova 25 enriched for mitochondrial DNA.** The number of sequences with a modern human or a Denisovan allele are reported in the second and third column, respectively. The diagnostic positions used in this analysis correspond to a set of 107 sites where the mitochondrial genome sequences of 4 Denisovans^7,8,61,62^ and a hominin from Sima de los Huesos^63^ differ from that of 312 modern humans^60^ (including the rCRS). Sequences are considered conflicting when overlapping multiple diagnostic positions and carrying both the modern and Denisovan alleles. We further discarded sequences obtained in the forward (reverse) orientation at C/T (G/A) polymorphisms or sequences carrying a third allele. Numbers in brackets correspond to DNA sequences with C-to-T substitutions within the three last positions of either end. The libraries highlighted in grey are those with minimal present-day human DNA contamination that were used to reconstruct the consensus mitochondrial sequence of Denisova 25.

| **Library ID** | **modern human state** | **Denisovan state** | **conflicting** | **Contamination estimate (%)** | **Binomial 95% confidence intervals** |
| --- | --- | --- | --- | --- | --- |
| B48024 | 6,419  [44] | 4,661  [991] | 2  [0] | 57.9  [4.3] | 57.0-58.8  [3.1-5.7] |
| B48025 | 7,120  [42] | 4,176  [869] | 2  [0] | 63.0  [4.6] | 62.1-63.9  [3.3-6.2] |
| B48026 | 411  [2] | 2,139  [547] | 0  [0] | 16.1  [0.4] | 14.7-17.6  [0.0-1.3] |
| **B48027** | **306**  **[8]** | **3,864**  **[945]** | **0**  **[0]** | **7.3**  **[0.8]** | **6.6-8.2**  **[0.3-1.6]** |
| **B49759** | **60**  **[5]** | **3,139**  **[724]** | **2**  **[1]** | **1.9**  **[0.7]** | **1.4-2.4**  **[0.2-1.6]** |
| **B49760** | **441**  **[13]** | **9,493**  **[1,948]** | **3**  **[0]** | **4.4**  **[0.7]** | **4.0-4.9**  **[0.4-1.1]** |
| **B49761** | **102**  **[2]** | **5,896**  **[1,266]** | **1**  **[1]** | **1.7**  **[0.2]** | **1.4-2.1**  **[0.0-0.6]** |
| **B49762** | **1,167**  **[11]** | **12,011**  **[2,729]** | **0**  **[0]** | **8.9**  **[0.4]** | **8.4-9.4**  **[0.2-0.7]** |
| **B49763** | **51**  **[3]** | **3,248**  **[718]** | **1**  **[0]** | **1.6**  **[0.4]** | **1.2-2.0**  **[0.1-1.2]** |
| B49764 | 792  [12] | 5,779  [1,336] | 3  [1] | 12.1  [0.9] | 11.3-12.9  [0.5-1.5] |
| **B50617** | **463**  **[18]** | **10,421**  **[2,376]** | **5**  **[1]** | **4.3**  **[0.8]** | **3.9-4.7**  **[0.4-1.2]** |
| **B50618** | **1,236**  **[20]** | **12,828**  **[2,804]** | **8**  **[2]** | **8.8**  **[0.7]** | **8.3-9.3**  **[0.4-1.1]** |
| **B50619** | **722**  **[10]** | **6,754**  **[1,530]** | **0**  **[0]** | **9.7**  **[0.7]** | **9.0-10.3**  **[0.3-1.2]** |
| B50621 | 953  [16] | 7,679  [1,699] | 3  [0] | 11.0  [0.9] | 10.4-11.7  [0.5-1.5] |
| B50622 | 1,709  [6] | 4,857  [1,062] | 0  [0] | 26.0  [0.6] | 25.0-27.1  [0.2-1.2] |
| B50623 | 2,236  [21] | 8,100  [1,762] | 5  [2] | 21.6  [1.2] | 20.8-22.4  [0.7-1.8] |
| B50624 | 3,272  [15] | 4,057  [964] | 2  [1] | 44.6  [1.5] | 43.5-45.8  [0.9-2.5] |
| **analysed**  **dataset** | **4,548**  **[90]** | **67,654**  **[15,040]** | **20**  **[5]** | **6.3**  **[0.6]** | **6.1-6.5**  **[0.5-0.7]** |

#### Reconstruction of the mitochondrial genome

To reconstruct the mitochondrial genome sequence of *Denisova 25*, the raw sequences from the 9 libraries selected above were realigned to the mitochondrial genome sequence of *Denisova 3* (reconstructed with high-coverage data)^62^. Using a Denisovan as reference limits the loss of ancient DNA sequences during alignment due to divergence to the reference genome. The same processing steps and filters were applied as described above. After merging the data from these libraries, a total of 703,678 sequences were retained, yielding a 2,422-fold average coverage of the mitochondrial genome. A consensus base was then called by a majority vote at every position with at least five overlapping sequences and at least 80% of the sequences carrying the same base (as done previously^63^). Any terminal T in sequences aligned in the orientation as sequenced, and terminal A in sequences aligned in the reverse orientation, were masked (converted into N) at the three last positions of either end to avoid introduction of deamination-induced substitutions into the consensus. At 16 positions, the majority base was supported by less than 80% of sequences, which may be due to present-day human DNA contamination. Using only sequences with signs of ancient DNA damage (at least one C-to-T substitution within the first three and last three positions), the majority base at nine of these positions was then supported by at least 80% of sequences. The other positions were in or adjacent to poly-C stretches and were resolved by inspection of the alignments in these regions (focusing on the fragments sequenced in the reverse orientation that span the poly-C stretches, as sequences in the forward orientation may exhibit C-to-T substitutions due to deamination that complicate their alignment). Comparison of the reconstructed mitochondrial sequence of *Denisova 25*, 6 Denisovans^7,8,61,62,64^, and a hominin from Sima de los Huesos^63^, shows that it is closest to the mitochondrial sequence of *Denisova 8* and most distant to that of *Denisova 3*, at the exception of that of the Sima de los Huesos hominin (**Table S13**).

**Table S13: Number of pairwise nucleotide differences between the full mitochondrial genome sequences of 7 Denisovans and a hominin from Sima de los Huesos (SH).** Ambiguous positions (gaps or missing data) were removed for each pair.

|  | SH | *Denisova3* | *Denisova4* | *Denisova2* | *Denisova20* | *Denisova21* | *Denisova8* |
| --- | --- | --- | --- | --- | --- | --- | --- |
| SH |  |  |  |  |  |  |  |
| *Denisova3* | 212 |  |  |  |  |  |  |
| *Denisova4* | 214 | 2 |  |  |  |  |  |
| *Denisova2* | 185 | 72 | 70 |  |  |  |  |
| *Denisova20* | 184 | 69 | 67 | 20 |  |  |  |
| *Denisova21* | 184 | 69 | 67 | 20 | 4 |  |  |
| *Denisova8* | 192 | 85 | 83 | 29 | 30 | 30 |  |
| *Denisova25* | 192 | 86 | 84 | 29 | 27 | 27 | 11 |

**Table S14: Bases in the mitochondrial genome where Denisova 8 and Denisova 25 differ.** The likely derived alleles are shaded in grey. Positions correspond to rCRS coordinates^58^. Excluding the positions in the hypervariable region (i.e., before position 577 and after position 16,023), there are 5 mutations on the Denisova 8 lineage (perhaps 6 if the A at position 11314 is a back mutation to the ancestral state) compared to 4 mutations on the Denisova 25 lineage (perhaps 5 if the G at position 11314 is a recurrent mutation). SH stands for Sima de los Huesos. Missing bases and indels are indicated with “N” and “-“, respectively.

| position | hypervariable? | *Den.8* | *Den.25* | rCRS | SH | *Den.3* | *Den.4* | *Den.2* | *Den.20* | *Den.21* |
| --- | --- | --- | --- | --- | --- | --- | --- | --- | --- | --- |
| 301 | yes | C | - | A | N | - | - | - | C | C |
| 2366 | no | G | A | G | G | G | G | G | G | G |
| 2649 | no | C | T | T | T | T | T | T | T | T |
| 3221 | no | G | A | A | A | A | A | A | A | A |
| 5809 | no | A | - | - | - | - | - | - | - | - |
| 6293 | no | T | C | T | T | T | T | T | T | T |
| 6617 | no | T | C | C | C | C | C | C | C | C |
| 8696 | no | T | C | T | T | T | T | T | T | T |
| 11314 | no | A | G | A | A | G | G | G | G | G |
| 14040 | no | A | G | G | G | G | G | G | G | G |
| 15119 | no | G | A | G | G | G | G | G | G | G |
| 16086 | yes | T | C | T | C | T | T | N | C | C |
| 16188 | yes | C | T | C | N | - | - | - | T | T |

#### Phylogenetic analysis

We estimated how long ago the *Denisova 25* individual lived and resolved his mitochondrial relationship with other archaic and modern humans using BEAST2 (Bayesian Evolutionary Analysis Sampling Trees, program version 2.6.1)^65^. The reconstructed mitochondrial genome sequence from the *Denisova 25* individual was first aligned with MAFFT (version 7.271)^66^ to the mitochondrial genome sequences of 6 Denisovans, 27 Neandertals, 54 present-day humans, 10 early modern humans, a hominin from Sima de los Huesos and a chimpanzee (see **Table S15** for a list of individuals and the corresponding accession numbers in the NCBI database). For consistency with previous analyses^57^, the hypervariable region was excluded, *i.e.*, only the coding parts between positions 577 and 16,023 (in rCRS coordinates)^57^ were used in subsequent analyses.

Time constraints were set with uniform priors for the age of ancient individuals and for the time to the most recent common ancestor of relevant groups (**Table S16**). The best fitting substitution model was identified with jModelTest 2.1.10^67^ to be the Tamura-Nei model^68^ with a faction of invariable sites and gamma distributed rate variation among sites (**Table S17**). Two possible tree priors (constant population size or Bayesian skyline) and two possible clock models (strict or relaxed) were evaluated with the path sampling approach implemented in the MODEL_SELECTION package^65^. The marginal likelihood of each model combination was computed from 40 path steps, comprising each 50,000,000 iterations, and sampling values and trees every 10,000 iterations after a pre-burn-in of 500,000 iterations, followed by a burn-in of 80%. Path sampling steps were spaced out using a beta distribution with parameter alpha equal to 0.3^69^. For the strict clock model, the mutation rate was set to 1.57 x 10^-8^ substitutions per site per year^57^. For the relaxed clock model, the mutation rate was allowed to vary between branches following a lognormal distribution with the mean set to 1.57 x 10^-8^ substitutions per site per year and the standard deviation following a gamma distribution estimated from the data. The strict clock with a Bayesian Skyline tree model showed a significantly better fit to the data than the next best fit model (log10 Bayes Factor>2, **Table S18**) following the scale provided by Kass and Raftery^70^.

We performed three Markov Chain Monte Carlo runs of 75,000,000 iterations with this model, sampling parameters every 5,000 iterations, with a burn-in of 10,000,000 iterations. We merged the results of these chains with the program *logcombiner*^71^ and report the estimates and their 95% High Posterior Density Intervals (HPDI) in **Table S19**. We estimated that the *Denisova 25* individual lived around 219 thousand years ago (ka; 95% HPDI: 159-275 ka) and shared a common mitochondrial ancestor with the *Denisova 8* individual around 238 ka (95% HPDI: 181-293 ka) and with later Denisovans around 285 ka (95% HPDI: 236-332 ka) (**Figure S11**).

The reconstructed tree differs from the published maximum parsimony trees^61,64^ in placing *Denisova 8* and *Denisova 25*, rather than *Denisova 3* and *Denisova 4*, as outgroups to the other Denisovans. Unlike maximum parsimony, BEAST uses a substitution model that better accounts for recurrent mutations and reversions (mutations back to the ancestral allele). The substantial age difference between early and late Denisovans may lead to the erroneous placement of the late Denisovans (*Denisova 3* and *Denisova 4*) as outgroups under maximum parsimony, owing to more reversions occurring on their long common branch (the longest among Denisovans).

Because the more divergent lineage of the Sima de los Huesos hominin is clearly an outgroup to all Denisovans, cases where it shares derived alleles with some Denisovans suggest either reversions in other Denisovans (with the ancestral allele represented by the modern human reference) or recurrent mutations occurring independently in the Sima de los Huesos lineage and in Denisovans with the derived allele. We identified six such positions consistent with reversions in the lineage leading to *Denisova 3* and *Denisova 4*, including two that are private to either *Denisova 3* or *Denisova 4* (**Table S20**). We also found five positions where *Denisova 3* and *Denisova 4* share a derived allele exclusively with the Sima de los Huesos mitochondrial genome, indicating recurrent mutations (**Table S21**). In contrast, only two positions suggest reversions in the lineage leading to the common ancestor of *Denisova 8* and *Denisova 25* (or recurrent mutations that other Denisovans share with the Sima de los Huesos lineage; **Table S22**).

Taken together, the multiple instances of recurrent mutations, and in particular reversions, support the *BEAST* tree as the more reliable reconstruction. The placement of *Denisova 3* and *Denisova 4* as outgroups in maximum parsimony trees is likely an artifact of reversions on their longer branch.

**Table S15: Dataset of mtDNA genomes used in the phylogenetic analysis.** “Accession” refers to the mtDNA genome identifier in the NCBI database.

| **Individual** | **Accession** | **Individual** | **Accession** | **Individual** | **Accession** |
| --- | --- | --- | --- | --- | --- |
| **Neandertal mtDNA** | | **Present-day modern human mtDNA** | | **Present-day modern human mtDNA** | |
| *Broion 1* | - | rCRS | NC_012920 | Mbuti | AF346998 |
| *Denisova 11* | KU131206 | Australian | AF346964 | Mbuti | AF346999 |
| *Denisova 15* | MK033602 | Australian | AF346963 | Mkamba | AF347000 |
| *Denisova 5 ("Altai")* | KC879692 | Australian | AF346965 | Piman | AF347001 |
| *El Sidrón 1253* | FM865409 | Bamileke | AF346967 | Papuan (coast) | AF347003 |
| *Feldhofer 1* | FM865407 | Biaka | AF346968 | Papuan (coast) | AF347002 |
| *Feldhofer 2* | FM865408 | Biaka | AF346969 | Papuan (Highlands) | AF347005 |
| *Goyet Q305-4* | KX198087 | Buriat | AF346970 | Papuan (Highlands) | AF347004 |
| *Goyet Q305-7* | KX198086 | Chinese | AF346973 | Saami | AF347006 |
| *Goyet Q374a-1* | KX198085 | Chinese | AF346972 | Samoa | AF347007 |
| *Goyet Q56-1* | KX198084 | Chukchi | AF346971 | San | AF347008 |
| *Goyet Q57-1* | KX198082 | Dutch | AF346975 | San | AF347009 |
| *Goyet Q57-2* | KX198088 | Effik | AF346976 | Siberian Inuit | AF347010 |
| *Goyet Q57-3* | KX198083 | Effik | AF346977 | Tatar | AF346974 |
| *Hohlenstein-Stadel* | KY751400.2 | English | AF346978 | Uzbek | AF347011 |
| *Les Cottés Z4-1514* | MG025536 | Evenki | AF346979 | Warao | AF347012 |
| *Mezmaiskaya 2* | MG025537 | Ewondo | AF346980 | Warao | AF347013 |
| *Mezmaiskaya 1* | FM865411 | French | AF346981 | Yoruba | AF347015 |
| *Okladnikov 2* | KF982693 | Georgian | AF346982 | Yoruba | AF347014 |
| *Scladina I-4A* | MK123269 | German | AF346983 | **Ancient modern human mtDNA** | |
| *Spy 94a* | MG025538 | Guarani | AF346984 | *Boshan 11* | KC521454 |
| *Stajnia S5000* | MT795654 | Hausa | AF346985 | *Dolní Věstonice 13* | KC521459 |
| *Strashnaya 2* | MT921957 | Ibo | AF346986 | *Dolní Věstonice 14* | KC521458 |
| *Vindija 33.16* | AM948965 | Ibo | AF346987 | *Iceman* | EU810403 |
| *Vindija 33.17* | KJ533544 | India | AF346966 | *Kostenki 14* | FN600416 |
| *Vindija 33.19* | KJ533545 | Italian | AF346988 | *Loschbour* | KC521455 |
| *Vindija 33.25* | FM865410 | Japanese | AF346989 | *Oberkassel 998* | KC521457 |
| **Denisovan mtDNA** | | Japanese | AF346990 | *Saqqaq Eskimo* | EU725621 |
| *Denisova 2* | KX663333 | Khirgiz | AF346991 | *Tianyuan* | KC417443 |
| *Denisova 20* | MT576652 | Kikuyu | AF346992 | *Ust-Ishim* | PRJEB6622 |
| *Denisova 21* | MT576653 | Korean | AF346993 | **Sima de los Huesos mtDNA** | |
| *Denisova 25* | This study | Lisongo | AF346994 | *Femur VIII* | NC_023100 |
| *Denisova 3* | NC_013993 | Mandenka | AF346995 | **Chimpanzee mtDNA** | |
| *Denisova 4* | FR695060 | Mbenzele | AF346997 | Jenny | X93335 |
| *Denisova 8* | KT780370 | Mbenzele | AF346996 |  |  |

**Table S16: Constraints set for the age of individuals and for the time to the most recent common ancestor (TMRCA) in the phylogenetic analysis performed with BEAST.** Uniform priors for the age of samples with direct radiocarbon dates were set within the 95% confidence intervals of the calibrated dates (calibrated using OxCal 4.3^72^ with IntCal13^73^ as reported in Douka et al. (2019)^74^, Fewlass, et al. (2019)^75^ and Picin, et al. (2020)^76^. The uniform priors for the age of Denisova 3 and the Sima de los Huesos hominin reflect prior knowledge from the analysis of the nuclear genome^45^ or the archaeological context^77^, respectively. Groups with constraints on their TMRCA were enforced to be monophyletic in the sampled trees.

| **Individual** | **Uniform prior:**  **initial value (range)** | **Individual** | **Uniform prior:**  **initial value (range)** |
| --- | --- | --- | --- |
| **Neandertal mtDNA** | | **Denisovan mtDNA** | |
| *Broion 1* | 50,000 (30,000-200,000) | *Denisova 3* | 50,000 (30,000-100,000) |
| *Denisova 11* |  | *Denisova 20* | 50,000 (30,000-300,000) |
| *Denisova 15* |  | *Denisova 21* |  |
| *Denisova 5 ("Altai")* |  | *Denisova 25* |  |
| *El Sidrón 1253* |  | *Denisova 2* |  |
| *Goyet Q305-7* |  | *Denisova 4* |  |
| *Goyet Q374a-1* |  | *Denisova 8* |  |
| *Hohlenstein-Stadel* |  | **Ancient modern human mtDNA** | |
| *Mezmaiskaya 1* |  | *Boshan 11* | 8,234 (8,152-8,316) |
| *Okladnikov 2* |  | *Dolní Věstonice 13* | 31,071 (30,884-31,249) |
| *Scladina I-4A* |  | *Dolní Věstonice 14* | 30,934 (30,741-31,120) |
| *Stajnia S5000* |  | *Iceman* | 5,300 (5,275-5,325) |
| *Strashnaya 2* |  | *Kostenki 14* | 37,473 (36,262-38,684) |
| *Vindija 33.17* |  | *Loschbour* | 8,054 (7,948-8,160) |
| *Vindija 33.19* |  | *Oberkassel 998* | 14,077 (13,755-14,105) |
| *Vindija 33.25* |  | *Saqqaq Eskimo* | 4,504 (4,423-4,585) |
| *Feldhofer 1* | 43,707 (42,670-44,744) | *Tianyuan* | 39,008 (37,761-40,254) |
| *Feldhofer 2* | 43,268 (42,193-44,342) | *Ust-Ishim* | 45,045 (43,212-46,878) |
| *Goyet Q305-4* | 44,236 (43,386-45,085) | **Sima de los Huesos mtDNA** | |
| *Goyet Q56-1* | 42,515 (42,063-42,967) | Femur VIII | 300,000 (260,000-780,000) |
| *Goyet Q57-1* | 44,696 (43,834-45,558) | **TMRCA** | |
| *Goyet Q57-2* | 41,185 (40,595-41,775) | Denisovans (including Sima) | (260,000 - Infinity) |
| *Goyet Q57-3* | 42,407 (41,946-42,867) | Denisovans (excluding Sima) | (50,000 - Infinity) |
| *Les Cottés Z4-1514* | 43,230 (42,720-43,740) | Neandertals | (100,000 - Infinity) |
| *Mezmaiskaya 2* | 43,834 (42,038-45,630) | modern humans | (50,000 - Infinity) |
| *Spy 94a* | 40,463 (39,840-41,085) | Neandertals – modern humans | (100,000 - Infinity) |
| *Vindija 33.16* | 43,707 (39,234-48,179) | All humans | (400,000 - Infinity) |

**Table S17: Best substitution models according to the three model selection measures computed by jModelTest 2.1.10.** These measures are the Bayesian Information Criterion (BIC), a measure based on Decision Theory (DT) and the corrected Akaike Information Criterion (AICc).

| **Measure** | **Model** | **Rank** | **Log-likelihood** | **Measure value** |
| --- | --- | --- | --- | --- |
| **AICc** | **GTR+I+G** | **1** | **-34,622.65** | **69,666.99** |
| AICc | TrN+I+G | 2 | -34,626.54 | 69,668.63 |
| AICc | GTR+I | 3 | -34,635.76 | 69,691.16 |
| **BIC** | **TrN+I+G** | **1** | **-34,626.54** | **71,230.51** |
| BIC | TrN+I | 2 | -34,639.37 | 71,246.52 |
| BIC | GTR+I+G | 3 | -34,622.65 | 71,251.65 |
| **DT** | **TrN+I+G** | **1** | **-34,626.54** | **0.000005** |
| DT | GTR+I+G | 2 | -34,622.65 | 0.002268 |
| DT | GTR+I | 3 | -34,635.76 | 0.014015 |

**Table S18: Marginal likelihoods of the different tested clock and tree models obtained from a path sampling analysis.** The best model is marked in bold.

| **Clock model** | **Tree model** | **Marginal log-likelihood** |
| --- | --- | --- |
| Strict | Coalescent with constant  population size | -35,207.2200 |
| Relaxed (uncorrelated lognormal) | Coalescent with constant  population size | -35,208.2570 |
| **Strict** | **Bayesian Skyline** | **-35,177.4179** |
| Relaxed (uncorrelated lognormal) | Bayesian Skyline | -35,182.2935 |

**Table S19: Estimates of molecular age and divergence times (years before present).** Estimates were obtained from three Markov Chain Monte Carlo runs of 75,000,000 iterations each (sampling parameter values and trees every 5,000 iterations, with a burn-in of 10,000,000 iterations). Dates were rounded to the nearest 10 years. TMRCA: Time to the Most Recent Common Ancestor. HPDI: Highest Posterior Density Interval. ESS: effective sample size

| **Parameter** | **Mean** | **95% HPDI** | **ESS** |
| --- | --- | --- | --- |
| Human-Chimpanzee TMRCA | 5,882,100 | 6,715,500 – 5,084,400 | 2,438 |
| Archaic and modern human TMRCA | 884,240 | 967,320 – 799,760 | 1,006 |
| Neandertal and modern human TMRCA | 439,070 | 495,120 – 387,410 | 3,585 |
| Modern Human TMRCA | 151,020 | 174,250 – 128,070 | 7,906 |
| Denisovan - Sima TMRCA | 709,530 | 811,240 – 611,910 | 657 |
| Neandertal TMRCA | 287,180 | 332,150 – 241,150 | 1,986 |
| *Denisova 25* and *Denisova 8* TMRCA | 238,060 | 292,850 – 181,460 | 243 |
| *Denisova 3*, *4*, *8* and *25* TMRCA | 285,150 | 331,950 – 235,730 | 218 |
| *Denisova 2*, *3*, *4*, *8* and *25* TMRCA | 286,000 | 332,230 – 236,430 | 218 |
| *Strashnaya 2* age | 135,270 | 172,250 – 98,770 | 1,143 |
| *Scladina I-4A* age | 128,600 | 169,960 – 87,960 | 1,079 |
| *Stajnia S5000* age | 119,690 | 155,360 – 84,100 | 1,225 |
| *Hohlenstein-Stadel* Neandertal age | 139,170 | 197,140 – 85,520 | 3,464 |
| Altai Neandertal (*Denisova 5*) age | 139,740 | 179,650 – 99,860 | 798 |
| *Broion 1* age | 41,190 | 49,200 – 31,520 | 16,569 |
| *Denisova 15* Neandertal age | 135,950 | 175,670 – 95,110 | 775 |
| *Vindija 33.17* age | 51,070 | 60,470 – 42,200 | 6,635 |
| *Vindija 33.19* age | 44,990 | 53,220 – 35,610 | 9,485 |
| *Vindija 33.25* age | 44,150 | 54,940 – 33,180 | 6,087 |
| *Mezmaiskaya 1* age | 101,390 | 138,320 – 64,190 | 1,437 |
| *Okladnikov 2* age | 95,210 | 121,480 – 71,250 | 1,651 |
| *Goyet Q305-7* age | 40,730 | 46,960 – 32,670 | 14,931 |
| *Goyet Q374a-1* age | 40,774 | 47,150 – 32,770 | 15,820 |
| *Denisova 11* age | 102,460 | 125,690 – 79,660 | 1,248 |
| *Denisova 3* age | 80,460 | 100,000 – 54,080 | 182 |
| *Denisova 4* age | 84,754 | 110,190 – 54,070 | 175 |
| *Denisova 8* age | 214,740 | 273,480 – 157,630 | 269 |
| ***Denisova 25* age** | **218,560** | **275,210** – **159,430** | **258** |
| *Denisova 2* age | 249,120 | 300,000 – 200,840 | 261 |
| *Denisova 20* age | 253,930 | 299,820 – 209,590 | 235 |
| *Denisova 21* age | 250,130 | 295,980 – 202,530 | 240 |
| Sima de los Huesos hominin age | 408,840 | 527,860 – 281,790 | 778 |

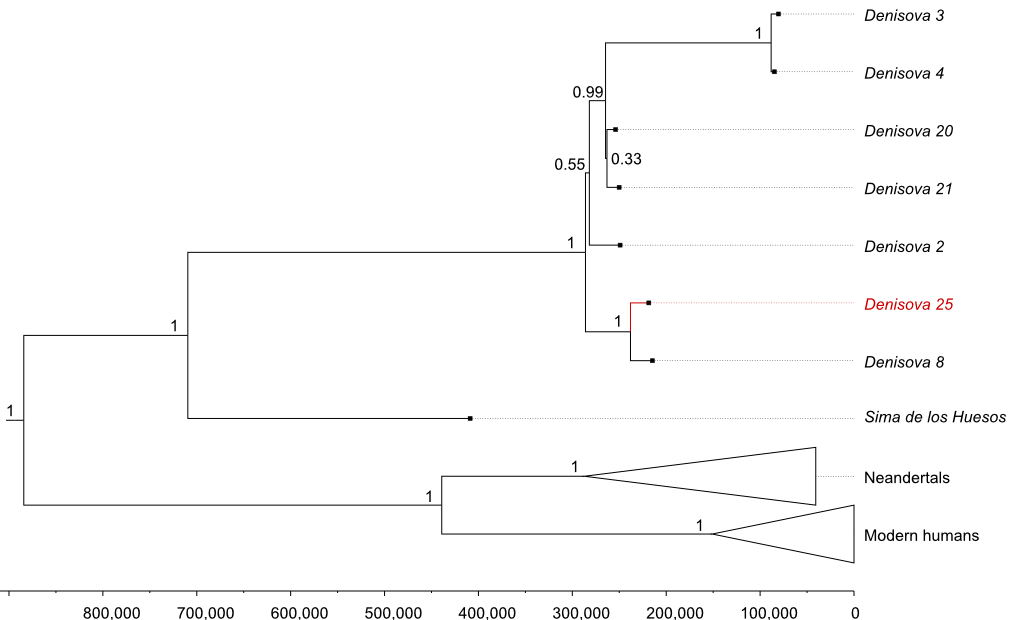

**Figure S11: Phylogenetic relationships among the mitochondrial genomes of the Denisova 25 individual and other humans.** The tree was reconstructed using the program BEAST 2^65^ and plotted using FigTree (v1.4.4)^78^. The mitochondrial lineage of the Denisova 25 individual is highlighted in red. The branch corresponding to the mtDNA of a chimpanzee used to root the tree is not shown. Mitochondrial lineages from modern humans and Neandertals were collapsed. The x-axis represents time in years before present. The numbers at the nodes correspond to the posterior probabilities of the depicted branching orders.

**Table S20: Positions in the mitochondrial genome with evidence that Denisova 3 and Denisova 4 may represent an outgroup to other Denisovans.** Note that at six positions the sequence of the Sima de los Huesos hominin^63^, which is an out-group to all Denisovans, carries the putative derived allele seen in the other Denisovans. Therefore, the ancestral allele (shaded in grey) seen in Denisova 3 and/or Denisova 4 at these six positions may result from mutations back to this state (i.e., reversions). Positions correspond to rCRS coordinates^58^.

| **position** | **Den. 3** | **Den. 4** | **Sima** | **rCRS** | **Other Denisovans** | **Comment** |
| --- | --- | --- | --- | --- | --- | --- |
| 94 | G | G | A | G | A | likely reversion |
| 146 | T | T | C | T | C | likely reversion |
| 1,040 | T | T | C | T | C | likely reversion |
| 3,600 | C | T | C | C | T | likely reversion in Den. 3 |
| 3,603 | C | C | N | C | T |  |
| 5,320 | C | C | C | C | T |  |
| 8,659 | A | A | A | A | G |  |
| 11,623 | C | C | C | C | T |  |
| 11,977 | C | C | C | C | T |  |
| 13,722 | A | A | G | A | G | likely reversion |
| 15,924 | A | A | A | A | G |  |
| 16,111 | C | C | C | C | T  (Den. 2: N) |  |
| 16,399 | G | A | G | A | A | likely reversion in Den. 4 |

**Table S21: Positions in the mitochondrial genome where Denisova 3 and Denisova 4 share an allele exclusively with a Sima de los Huesos hominin.** The 6 positions where the shared allele is ancestral (shaded in grey) are also reported in **Table S20** and may be evidence that Denisova 3 and Denisova 4 were an outgroup to other Denisovans. The 5 positions where the shared allele is derived are not compatible with a single mutation event and may represent recurrent mutations in the mitochondrial sequences of the Sima de los Huesos hominin, Denisova 3 and Denisova 4. Positions correspond to rCRS coordinates^58^.

| **position** | **Den. 3** | **Den. 4** | **Sima** | **rCRS** | **Other Denisovans** | **Comment** |
| --- | --- | --- | --- | --- | --- | --- |
| 151 | T | T | T | C | C | potential recurrent mutation |
| 189 | G | G | G | A | A | potential recurrent mutation |
| 1,393 | A | A | A | G | G | potential recurrent mutation |
| 5,320 | C | C | C | C | T |  |
| 8,659 | A | A | A | A | G |  |
| 9,380 | A | A | A | G | G | potential recurrent mutation |
| 11,623 | C | C | C | C | T |  |
| 11,977 | C | C | C | C | T |  |
| 15,924 | A | A | A | A | G |  |
| 16,111 | C | C | C | C | T  (Den. 2: N) |  |
| 16,320 | T | T | T | C | C | potential recurrent mutation |

**Table S22: Positions in the mitochondrial genome with evidence that Denisova 8 and Denisova 25 may represent an out-group to other Denisovans.** Note that at two positions the sequence of the Sima de los Huesos hominin^63^, which is an out-group to all Denisovans, carries the putative derived allele seen in the other Denisovans. Therefore, the ancestral allele (shaded in grey) seen in Denisova 8 and Denisova 25 at these two positions may result from mutations back to the ancestral state (i.e., reversions). Positions correspond to rCRS coordinates^58^.

| **position** | **Den. 8** | **Den. 25** | **Sima** | **rCRS** | **Other Denisovans** | **Comment** |
| --- | --- | --- | --- | --- | --- | --- |
| 5,471 | G | G | A | G | A | likely reversion |
| 10,930 | C | C | C | C | T |  |
| 11,314 | A | A | A | A | G |  |
| 12,654 | A | A | A | A | G |  |
| 13,437 | T | T | T | T | C  (Den. 2: T) |  |
| 15,358 | A | A | G | A | G | likely reversion |
| 16,134 | C | C | C | C | T |  |
| 16,294 | C | C | C | C | T |  |

### Supplementary Note 5: The Y-chromosome of *Denisova 25*

#### Data Processing

The Y chromosome sequences of *Denisova 25* were processed as described in Supplementary Note 3. However, we observed 154 heterozygous sites after genotyping (***Table S23***). As heterozygous sites are unexpected outside the pseudoautosomal regions of the Y chromosome, we interpreted these heterozygous positions as evidence of errors. As mapping errors can happen around indels, we removed sequences containing indels following indel realignment. After applying the *map35_100* filter from Prüfer *et al*. (2014)^48^, 1,859,721 confidently aligned sequences remained for analysis (***Table S24***). Read length distribution (***Figure S12***) and substitution patterns (***Figure S13***) of the retained sequences match that described for the rest of the nuclear genome (Supplementary Note 3).

Genotypes were also called with *snpAD*^54^ using bases with a minimum base quality of 30. As *snpAD* is designed to call diploid genotypes, we converted the remaining 145 heterozygous genotype calls to the most likely homozygous genotype based on the genotype posterior probabilities computed by *snpAD*. Genotypes were then filtered based on GC-dependent coverage cutoffs to remove positions below the 2.5% percentile and above the 97.5% percentile of each GC-dependent coverage distribution, as described in Supplementary Note 3. Finally, we applied the *capture_full* filter from Petr *et al*. (2020)^79^ to the final genotype calls to remove tandem repeats identified by the Tandem Repeat Finder^55^ and other repetitive regions missed by the *map35_100* filter. Due to the repetitive nature of the human Y chromosome, out of the 59.37 megabases (Mb) comprising the Y chromosome of the human genome assembly (hg19), *Denisova 25* genotypes could be confidently called at 4.27 million (or 7.19 %) sites. The mean coverage of the genotyped sites is 12.15 and the mean genotype quality is 73.32.

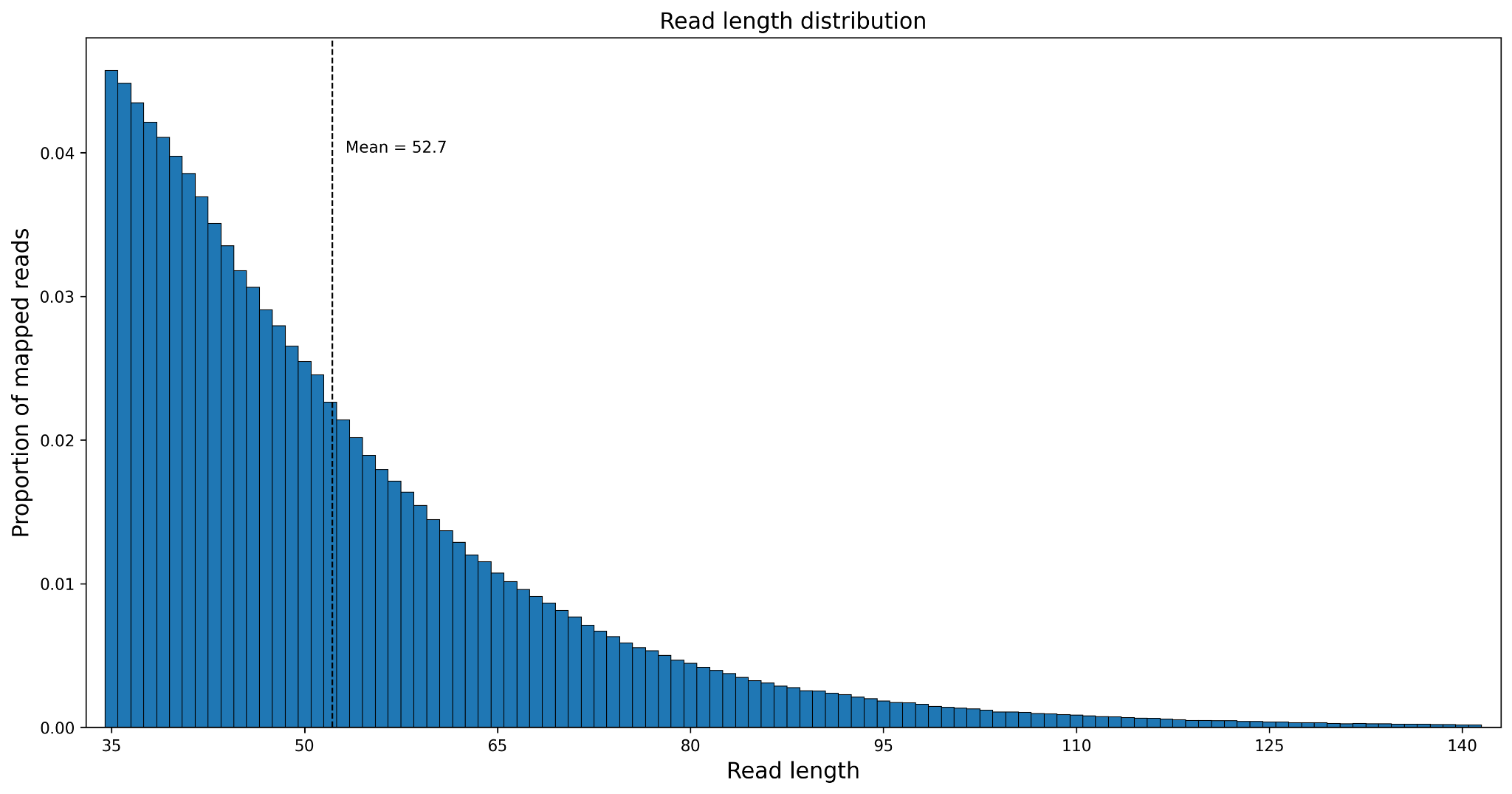

**Figure S12: Length distribution of Y chromosome sequences passing the quality and mappability filters.**

**Table S23: Effect of filtering out sequences with indels on the number of heterozygous genotype calls in the Y chromosome.**

|  | **all sequences** | **excluding sequences with indels** |
| --- | --- | --- |
| **Number of sequences** | 1,891,017 | 1,859,721 |
| **Number of genotypes** | 4,276,408 | 4,270,465 |
| **Number of heterozygotes** | 154 | 145 |

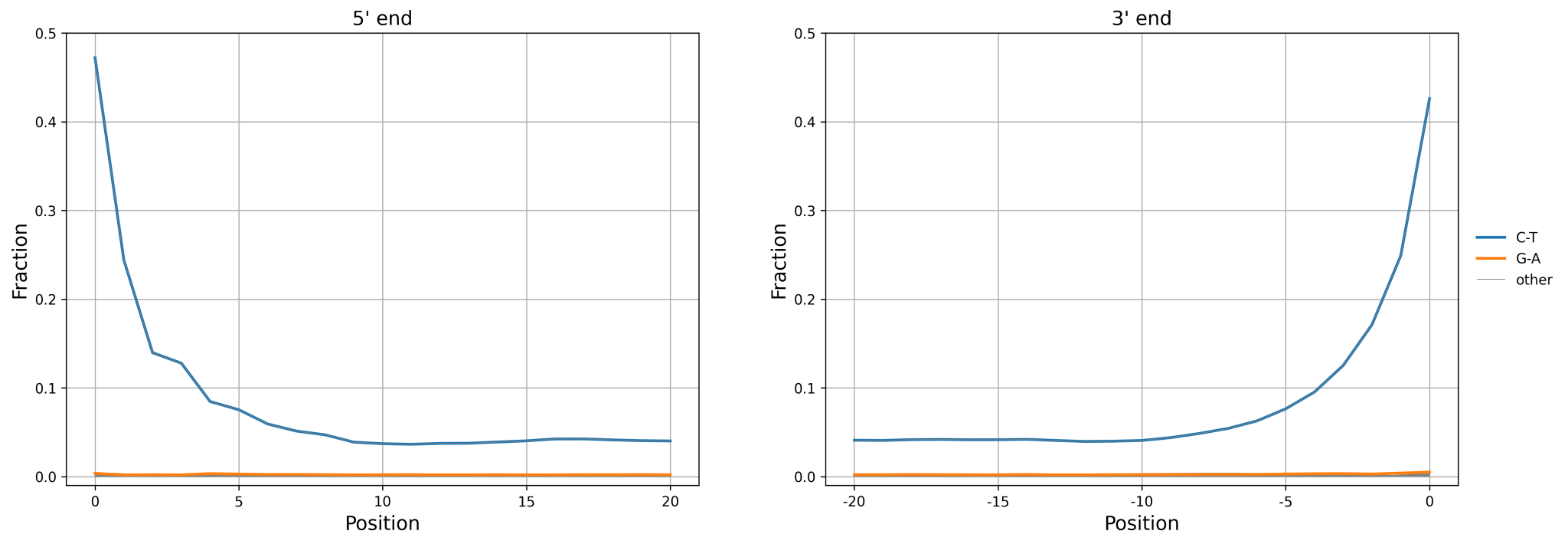

**Figure S13: Ancient DNA damage patterns from uniquely mapped Y chromosome sequences.**

**Table S24: Number of uniquely mapped sequences and proportion of sequences with terminal C-to-T substitutions per library.** Terminal C-to-T substitutions are those observed at the three first or last positions of the sequences.

| **Library** | **Mapped sequences** | **Mean sequence length** | **Sequences with terminal C-to-T substitutions** | **% sequences with terminal C-to-T substitutions** |
| --- | --- | --- | --- | --- |
| **A43615** | 724,218 | 52.3 | 203,309 | 28.1 |
| **A43617** | 294,260 | 52.4 | 80,056 | 27.2 |
| **A45393** | 666,998 | 52.8 | 188,285 | 28.2 |
| **A45394** | 174,245 | 54.0 | 46,808 | 26.9 |
| **Combined** | 1,859,721 | 52.7 | 518,458 | 27.9 |

#### Processing of a comparative Y chromosome dataset

For the analysis of the *Denisova 25* Y chromosome, we used a dataset comprising Y chromosomes from present-day humans, ancient humans and archaic humans (**Table S25**). For the present-day humans, we selected 10 African and 15 non-African Y chromosomes from the Human Genome Diversity Project (HGDP)^80^ to capture the variation in the Y chromosome phylogeny. To represent the most deeply divergent modern human lineages in our dataset, we also included two African Y chromosomes from the 1000 Genomes Project (1kGP)^81^ as well as Y chromosomes from two closely related individuals carrying the A00 haplogroup^82^. Due to the relatively low coverage of the two A00 Y chromosomes, we merged the A00 sequence data into a single BAM file, as done in the original study, because the two chromosomes differ at only nine positions. The dataset also includes shotgun-sequenced Y chromosomes from the following radiocarbon-dated ancient humans: *Ust’-Ishim*^83^, an individual from Shum Laka with the A00 haplogroup^84^, Loschbour^85^, and *Yana 1*^86^. Finally, the dataset includes Y chromosomes from two Neandertals (*Chagyrskaya 2*^87^ and *Mezmaiskaya 2*^79^) and two Denisovans (*Denisova 4* and *Denisova 8*)^79^ that were obtained by capture sequencing.

As we obtained raw, unaligned reads for the HGDP and 1KGP individuals, we used *MarkIlluminaAdapters* from Picard tools (v2.18.29) to flag adapter sequences and aligned the reads to hg19 using BWA-MEM (v0.7.17)^46^*.* For the ancient individual from Shum Laka, we aligned the sequences to hg19 using BWA-ALN (v0.4.9) with ancient parameters (-n 0.01 -o 2 -l 16500). We then applied the processing described for the *Denisova 25* Y chromosome to the Y chromosome sequences of all individuals. For the Y chromosome sequences of *Denisova 4* and *Denisova 8,* however, we only removed sites with coverage higher than the 99% percentile of the respective GC-dependent coverage distribution due to the low coverage available for both individuals.

We used the Y chromosome of the *panTro6* chimpanzee genome assembly to identify ancestral alleles. From the UCSC hg19-panTro6 multiple alignment file (hg19.panTro6.synNet.maf), we extracted the panTro6 Y chromosome sequences that aligned to the hg19 Y chromosome to call panTro6 genotypes.

**Table S25: Individuals whose Y chromosomes were included in the comparative dataset.**

| **Individual** | **Data Source** | **Sample accession** | **Haplogroup** |
| --- | --- | --- | --- |
| Chagyrskaya2 | ^87^ | SAMEA110640443 | Neandertal |
| Mezmaiskaya2 | ^79^ | SAMEA7062760 | Neandertal |
| Denisova 4 |  | SAMEA7062758 | Denisovan |
| Denisova 8 |  | SAMEA7062759 | Denisovan |
| Shum Laka (2/SE II) | ^84^ | SAMEA6104836 | A00 |
| Loschbour | ^85^ | SAMEA2697124 | I2 |
| Ust'Ishim | ^83^ | SAMEA2656891 | K |
| Yana1 | ^86^ | SAMEA5684442 | P1 |
| A00 | ^82^ | GRC13292545 | A00 |
| HG02982 | ^81^ | GRC13292546 | A0a1 |
| HG02666 |  | SAMN01761225 | A1a |
| HGDP01029 | ^80^ | SAMN00779975 | A1b1a1a1 |
| HGDP01406 |  | SAMEA2580841 | A1b1b2b |
| HGDP00931 |  | SAMEA2580853 | B-M181 |
| cHGDP00478 |  | SAMEA2580834 | B2a |
| HGDP00453 |  | SAMEA2580803 | B2b2 |
| HGDP00475 |  | SAMEA2580818 | B2b1a1c |
| HGDP00984 |  | SAMEA2580836 | B2b1a1a |
| HGDP00757 |  | SAMEA2581095 | D1b2a2 |
| HGDP01214 |  | SAMEA2581057 | D1a1a |
| HGDP01031 |  | SAMEA2580848 | E2b1a1 |
| HGDP01200 |  | SAMEA2580900 | E1a2a1b1 |
| HGDP00908 |  | SAMEA2580889 | E1b1a1a1c2c3b |
| HGDP00103 |  | SAMEA2581152 | C2b1c |
| HGDP00213 |  | SAMEA2581196 | G2b1 |
| HGDP00127 |  | SAMEA2581169 | I2a2a1a2a2 |
| HGDP00057 |  | SAMEA2581295 | J1a2b |
| HGDP00056 |  | SAMEA2581294 | J2a1 |
| HGDP00549 |  | SAMEA2581681 | M1 |
| HGDP01009 |  | SAMEA2581745 | Q1a2a1a1 |
| HGDP00136 |  | SAMEA2581321 | R1a1a1b2 |
| HGDP00218 |  | SAMEA2581199 | R2a |
| HGDP01298 |  | SAMEA2581140 | L1a2 |
| HGDP01192 |  | SAMEA2581001 | N1 |
| HGDP01225 |  | SAMEA2581048 | O2a2b1a1a |
| HGDP01190 |  | SAMEA2580999 | O1b1a1a1a |

#### Present-day Human DNA Contamination

To estimate present-day human DNA contamination among the sequences obtained from the *Denisova 25* Y chromosome, we measured the proportion of modern human-derived alleles at positions where Denisovans carry an ancestral allele. We defined a set of 960 informative positions (consisting of 362 transversions and 598 transitions) where we observe the modern human reference allele in all present-day human Y chromosomes whereas *Denisova 4* and *Denisova 8* carry the chimpanzee allele.

We first used the aligned sequences of the *Denisova 25* Y chromosome to estimate contamination. We provide sequence-based contamination estimates for all of the aligned sequences as well as for a subset of sequences that exhibit signs of ancient DNA damage, a C-to-T substitution within the first three or last three bases. To avoid any error due to DNA damage on our sequence-based contamination estimates, we disregard a sequence in the forward or reverse orientation if one of the informative alleles is a C or G, respectively. We estimated that 1.32% and 0.78% of all sequences and sequences from putatively deaminated DNA fragments stem from modern human contamination, respectively (**Table S26**). Using *AuthentiCT*^49^, which uses patterns of ancient DNA damage to estimate present-day DNA contamination, we obtained a similar contamination estimate of 1.98% ± 0.518 for all sequences (**Table S27**).

At the informative sites, we were able to confidently call 874 genotypes for *Denisova 25*, with 868 positions matching the ancestral genotype resulting in a genotype-based contamination estimate of 0.69% (95% CI 0.14 - 1.23). If only transversions are considered, the modern human allele is not observed in any of the 332 genotype calls for *Denisova 25*.

We then used the high coverage *Denisova 25* Y chromosome to estimate present-day human contamination among the sequences of the *Denisova 4* and *Denisova 8* Y chromosomes. We defined another set of informative sites where modern humans are derived but the *Denisova 25* genotype matches the ancestral state of the chimpanzee Y chromosome. We then estimated contamination in the low-coverage Denisovan genomes using all aligned sequences or those with terminal C-to-T substitutions at the last three positions of either end. As in the *Denisova 25* contamination estimates, we disregard a sequence if one of the informative alleles is a C when the sequence is in the forward orientation or if one of the informative alleles is a G when the sequence is in the reverse orientation. Contamination estimates are reported in **Table S28**.

**Table S26: Present-day human DNA contamination of the Denisova 25 Y chromosome dataset.**

|  | **Library** | **Sequences with the modern human allele** | **Number of sequences overlapping informative sites** | **Contamination %** | **95% binomial CI** |
| --- | --- | --- | --- | --- | --- |
| **All sequences** | **A43615** | 12 | 1,797 | 0.66 | (0.29, 1.04) |
|  | **A43617** | 23 | 767 | 2.91 | (1.74, 4.08) |
|  | **A45393** | 18 | 1,731 | 1.03 | (0.56, 1.50) |
|  | **A45394** | 10 | 423 | 2.31 | (0.89, 3.72) |
|  | **Combined** | 63 | 4,718 | 1.32 | (0.99, 1.64) |
| **Sequences with terminal C-to-T substitutions** | **A43615** | 2 | 463 | 0.43 | (0.00, 1.02) |
|  | **A43617** | 2 | 214 | 0.93 | (0.00, 2.20) |
|  | **A45393** | 6 | 472 | 1.26 | (0.26, 2.25) |
|  | **A45394** | 0 | 128 | 0 | (0.00, 0.00) |
|  | **Combined** | 10 | 1,277 | 0.78 | (0.30, 1.26) |

**Table S27: Present-day human DNA contamination among the Denisova 25 Y chromosome sequences as estimated by AuthentiCT.**

| **Library** | **Contamination %** | **1 std. error** |
| --- | --- | --- |
| **A43615** | 0.7853 | 0.5434 |
| **A43617** | 4.0675 | 0.5041 |
| **A45393** | 0.1475 | 0.5265 |
| **A45394** | 4.0134 | 0.5032 |
| **Combined** | 1.9849 | 0.5181 |

**Table S28: Present-day human DNA contamination in the Denisova 4 and Denisova 8 Y chromosome sequences.**

|  |  | **Sequences with the modern human allele** | **Number of sequences overlapping informative sites** | **Contamination %** | **95% binomial CI** |
| --- | --- | --- | --- | --- | --- |
| ***Denisova 4*** | **All sequences** | 63 | 837 | 7.00 | (5.33, 8.67) |
|  | **Sequences with terminal C-to-T substitutions** | 1 | 109 | 0.91 | (0.00, 2.68) |
| ***Denisova 8*** | **All sequences** | 44 | 2,182 | 1.98 | (1.40, 2.55) |
|  | **Sequences with terminal C-to-T substitutions** | 4 | 743 | 0.54 | (0.01, 1.06) |

#### Missense variants in the Y chromosome of *Denisova 25*

To identify missense variants in the *Denisova 25* Y chromosome, we applied the Ensembl Variant Effect Predictor (VEP, tool version 107, database version 107)^88^ using the GRCh37 assembly database. This revealed 7 missense variants in 4 genes (**Table S29**). The same variants are also found in the Y chromosomes of *Denisova 4* and *Denisova 8* suggesting that they occurred early in the Denisovan Y chromosome phylogeny (**Table S30**). All detected variants are reported by VEP as having MODERATE impact. No clinically significant phenotypes are reported in ClinVar for the three variants found in present-day populations. In the gnomAD dataset (v4.0.0), rs7067496 is commonly found in present-day populations (allele frequency = 7.8%), while rs866266594 and rs1034100630 are rare variants (allele frequency < 0.1%).

**Table S29: Missense variants on the Denisova 25 Y chromosome.**

| **Missense Variant** | **SNP** | **Gene** | **Gene Function** | **Phenotypes in modern humans associated with other variants in these genes** |
| --- | --- | --- | --- | --- |
| Y_6932181_C/G | unknown SNP | *TBL1Y* | Transcription activator | Deafness, Y-Linked 2 |
| Y_6955466_A/G | unknown SNP |  |  |  |
| Y_9384122_A/C | rs866266594 | *FAM197Y1* | RNA Gene | Not available |
| Y_14832620_G/T | rs7067496 | *USP9Y* | Peptidase | Spermatogenic failure, Y-linked, 2  Partial Chromosome Y Deletion |
| Y_14850182_G/A | unknown SNP |  |  |  |
| Y_14952462_T/A | unknown SNP |  |  |  |
| Y_21905071_C/T | rs1034100630 | *KDM5D* | Histone demethylase | Spermatogenic failure, Y-linked, 2 |

**Table S30: Sharing of missense variants amongst Denisovans.** The numbers in brackets represent the coverage at these positions.

| **Missense Variant** | **Ancestral allele** | ***Denisova 25*** | ***Denisova 4*** | ***Denisova 8*** |
| --- | --- | --- | --- | --- |
| Y_6932181_C/G | C | G (5) | G (5) | G (3) |
| Y_6955466_A/G | A | G (9) | G (4) | G (1) |
| Y_9384122_A/C | G | C (7) | C (3) | C (3) |
| Y_14832620_G/T | T | T (9) | - | T (5) |
| Y_14850182_G/A | G | A (15) | - | - |
| Y_14952462_T/A | A | A (19) | A (2) | A (7) |
| Y_21905071_C/T | T | T (13) | T (1) | T (5) |

#### Relationship to other Denisovans

To determine the relationship between the high coverage Y chromosome of *Denisova 25* and the low coverage Y chromosomes of the other two male Denisovans, we defined a set of diagnostic sites where the *Denisova 25* allele (genotype call) differs from both the chimpanzee and the human reference allele. We then used the genotypes as well as the aligned sequences of *Denisova 4* and *Denisova 8* to determine whether *Denisova 25* is more closely related to either individual.

Using genotypes, we counted the number of diagnostic sites where one of the low-coverage Denisovan Y chromosomes has the derived allele (i.e. the *Denisova 25* allele) while the other has the ancestral allele (the allele that matches both the chimpanzee and the modern human reference) (**Table S31**). We also tested whether the minimum coverage cutoff used for the low-coverage Denisovan genotypes affects the results. We find that *Denisova 25* shares more alleles with *Denisova 8* irrespective of the minimum coverage cutoff.

Using sequences, we counted the proportion of alleles matching the allele of *Denisova 25* for all sequences or those with at least one C-to-T substitution within the last three positions of either end (**Table S32**). To avoid errors stemming from deamination we disregard a sequence if one of the informative alleles is a C or G when the sequence is in the forward or reverse orientation, respectively. Using the sequences, we find that *Denisova 25* is more closely related to *Denisova 8* than to *Denisova 4*, supporting the results obtained with the genotypes.

**Table S31:** **Y chromosome relationship between Denisova 25 and the other two male Denisovans based on genotype calls.** Minimum genotype coverage refers to the minimum cutoff applied to the low-coverage Denisovan genotype calls. Only positions with genotype information for all three Denisovans were counted among those where Denisova 25 carries a derived allele that is absent from modern humans.

|  | **Minimum Genotype Coverage** | **Number of positions** | ***Denisova 4*** | ***Denisova 8*** |
| --- | --- | --- | --- | --- |
| **All positions** | **1** | 913 | 10 | 51 |
|  | **2** | 381 | 0 | 15 |
|  | **3** | 128 | 0 | 4 |
| **Transversions only** | **1** | 309 | 4 | 14 |
|  | **2** | 133 | 0 | 7 |
|  | **3** | 47 | 0 | 2 |

**Table S32: Y chromosome relationship between Denisova 25 and the other two male Denisovans based on sequences.**

|  | **All sequences** | | | | **Sequences with terminal C-to-T substitutions** | | | |
| --- | --- | --- | --- | --- | --- | --- | --- | --- |
|  | **Ancestral allele count** | **Derived allele count** | **% matching** | **95% binomial CI** | **Ancestral allele count** | **Derived allele count** | **% matching** | **95% binomial CI** |
| ***Denisova 4*** | 106 | 821 | 88.57 | (86.52, 90.61) | 10 | 98 | 90.74 | (85.27, 96.21) |
| ***Denisova 8*** | 119 | 2,024 | 94.40 | (93.43, 95.38) | 29 | 722 | 96.14 | (94.76, 97.52) |

#### Time to the most recent common ancestor with modern humans

We next estimated the time to the most recent common ancestor (TMRCA) between the *Denisova 25* Y chromosome and the Y chromosomes of modern humans. As reference bias results in missing mutations in deeply divergent human Y chromosomes, we used the method described in Swiel, Kelso, Peyrégne (2025)^89^, which filters out Y chromosome regions based on modern human-chimpanzee sequence divergence to minimise the effect of this bias on TMRCA estimates. For comparison, we also estimated the TMRCA between the Y chromosomes of modern humans and the two other Denisovans or two Neandertals, as well as between the African A00 Y chromosome^82^ and other modern human Y chromosomes.

We first computed the human Y chromosome mutation rate using the high-coverage genome of *Ust’-Ishim*^83^ -- a 45,000-year-old modern human from Siberia -- by determining the number of mutations missing on the *Ust’-Ishim* Y chromosome compared to present-day non-African Y chromosomes (**Figure S14**). The Y chromosome mutation rate was estimated to be 7.16x10^-10^ (95% CI 4.80x10^-10^ to 9.72x10^-10^) mutations/bp/year, while the mutation rate estimate for the regions of the Y chromosome that remain after filtering based on modern human-chimpanzee sequence divergence was found to be 6.61x10^-10^ (95% CI 4.12x10^-10^ to 9.57x10^-10^) mutations/bp/year. We then computed the number of mutations that happened on the modern human lineage since the most recent common ancestor with A00 or the archaic human Y chromosomes, using the chimpanzee Y chromosome as an outgroup to identify the derived and ancestral alleles. Finally, we used the estimated mutation rate to convert this number of mutations into a TMRCA estimate. To calculate confidence intervals for the TMRCA estimates, we used a bootstrapping approach whereby we randomly sampled branch lengths (i.e., number of mutations) from a Poisson distribution with the λ parameter set to the observed branch lengths. We then calculated the TMRCA for each iteration and used this simulated TMRCA distribution to compute the 95% confidence intervals. Results are reported in **Figure S15**, **Figure S16**, and **Figure S17** for the modern human TMRCA, Neandertal-modern human TMRCA, and Denisovan-modern human TMRCA, respectively.

If the Y chromosome is not filtered to minimise the effect of reference bias, the modern human-Denisovan Y chromosome TMRCA estimates for the low coverage Denisovans are older than the TMRCA calculated using the high coverage *Denisova 25* Y chromosome. Furthermore, these estimates differ depending on whether an African or a non-African Y chromosome is used. By contrast, the modern human-Denisovan Y chromosome TMRCA estimate is around 800ka using either the African or the non-African branch after correcting for the effect of reference bias.

We then estimated the age of *Denisova 25* based on the number of missing mutations on his Y chromosome compared to those of modern humans. The difference was then calibrated into years before the present based on our estimate of the mutation rate. While the age estimate of 180ka [105ka-255ka 95% CI] for the filtered Y chromosome dataset is similar to the estimated age of layer 17 where *Denisova 25* was found^2^, and that obtained with the autosomes and the mitochondrial genome, the estimate obtained with the unfiltered dataset is unreasonably old (over 300ka), consistent with the effect of missing mutations due to reference bias.

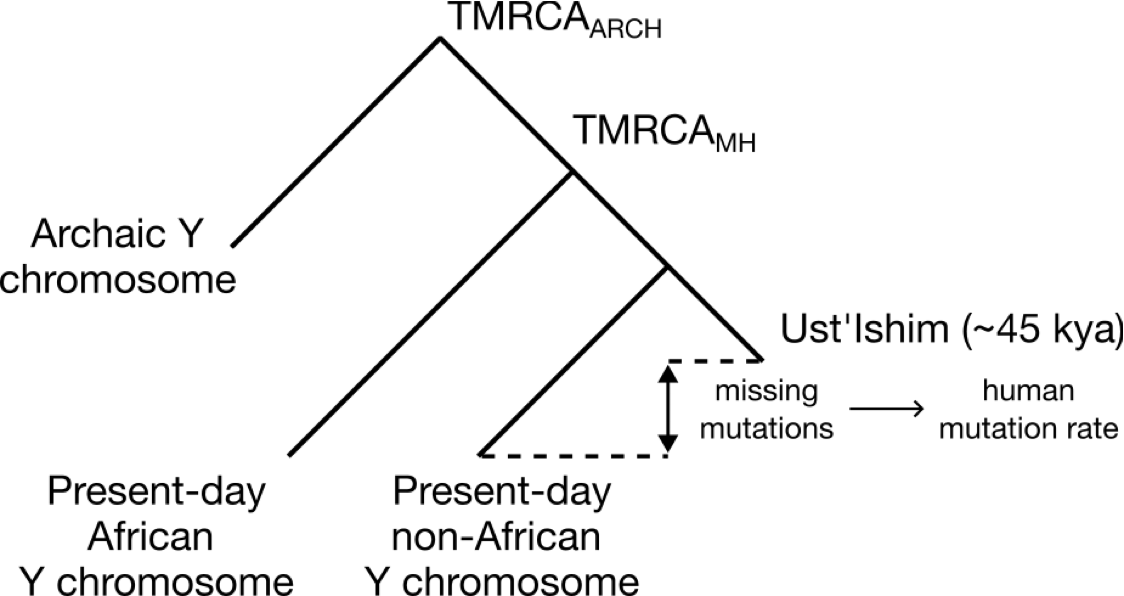

**Figure S14: Illustration of the method used to compute the Y chromosome TMRCA between modern and archaic humans.** We used the Y chromosome of a present-day African, a present-day non-African and Ust’-Ishim (an ancient radiocarbon-dated non-African).

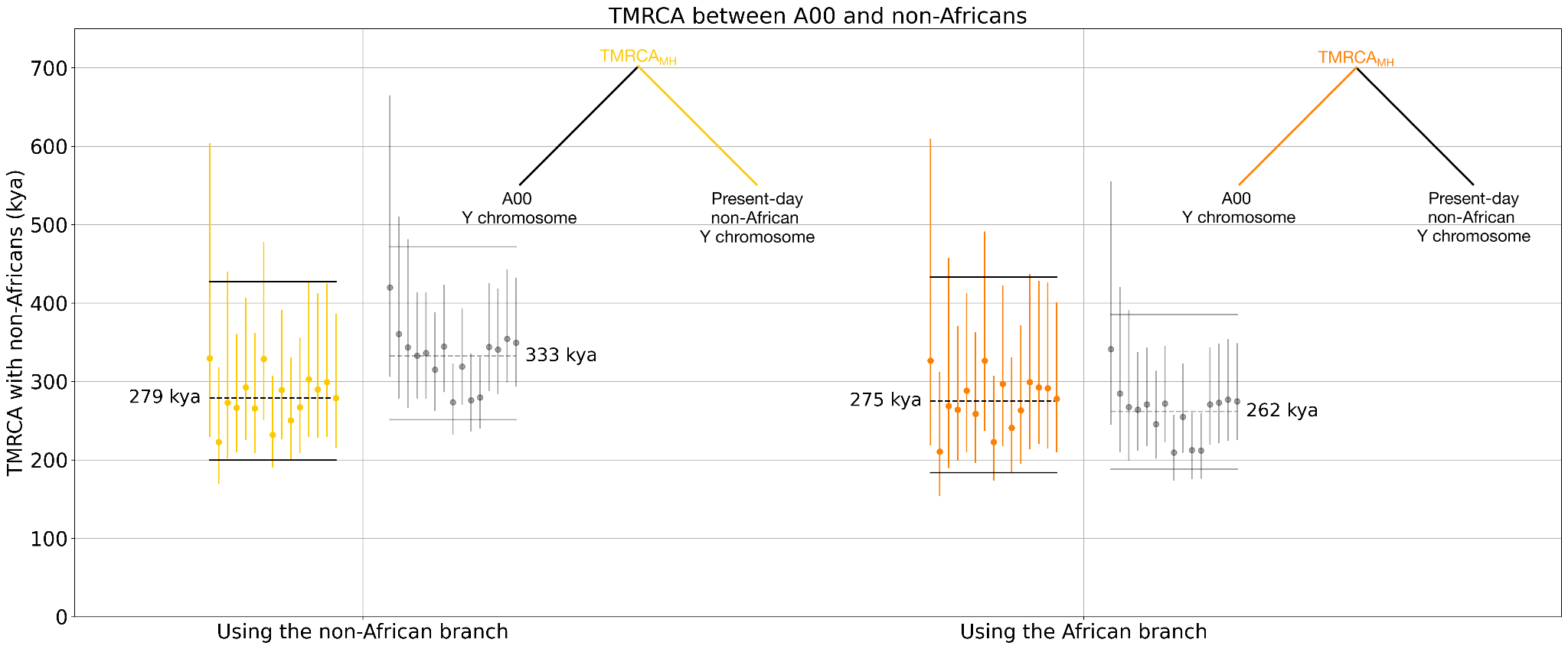

**Figure S15: Y chromosome TMRCA estimates between a present-day A00 Y chromosome and 15 present-day non-African Y chromosomes.** The points in colour represent TMRCAs computed with the filtered Y chromosome with minimal mapping bias, while the dark grey points show the TMRCAs calculated using the full Y chromosome. Each dot represents the TMRCA with a single present-day individual Y chromosome while error bars denote 95% CIs based on the resampling approach described in the text. The horizontal lines represent the mean TMRCA together with 95% CIs computed over all non-Africans.

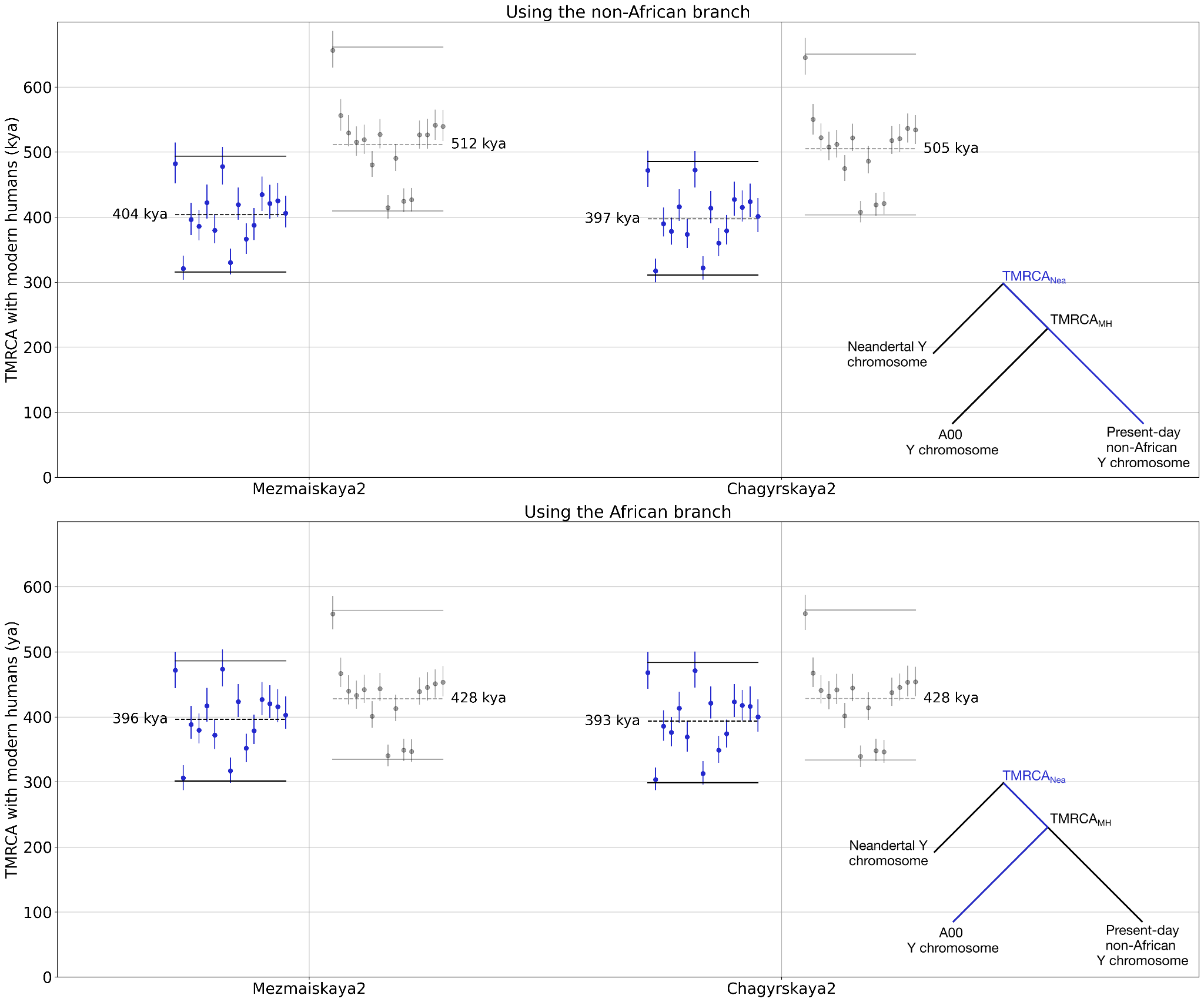

**Figure S16: Y chromosome TMRCA estimates between Neandertal Y chromosomes and 15 present-day non-African Y chromosomes.** See the caption of **Figure S15** for more information.

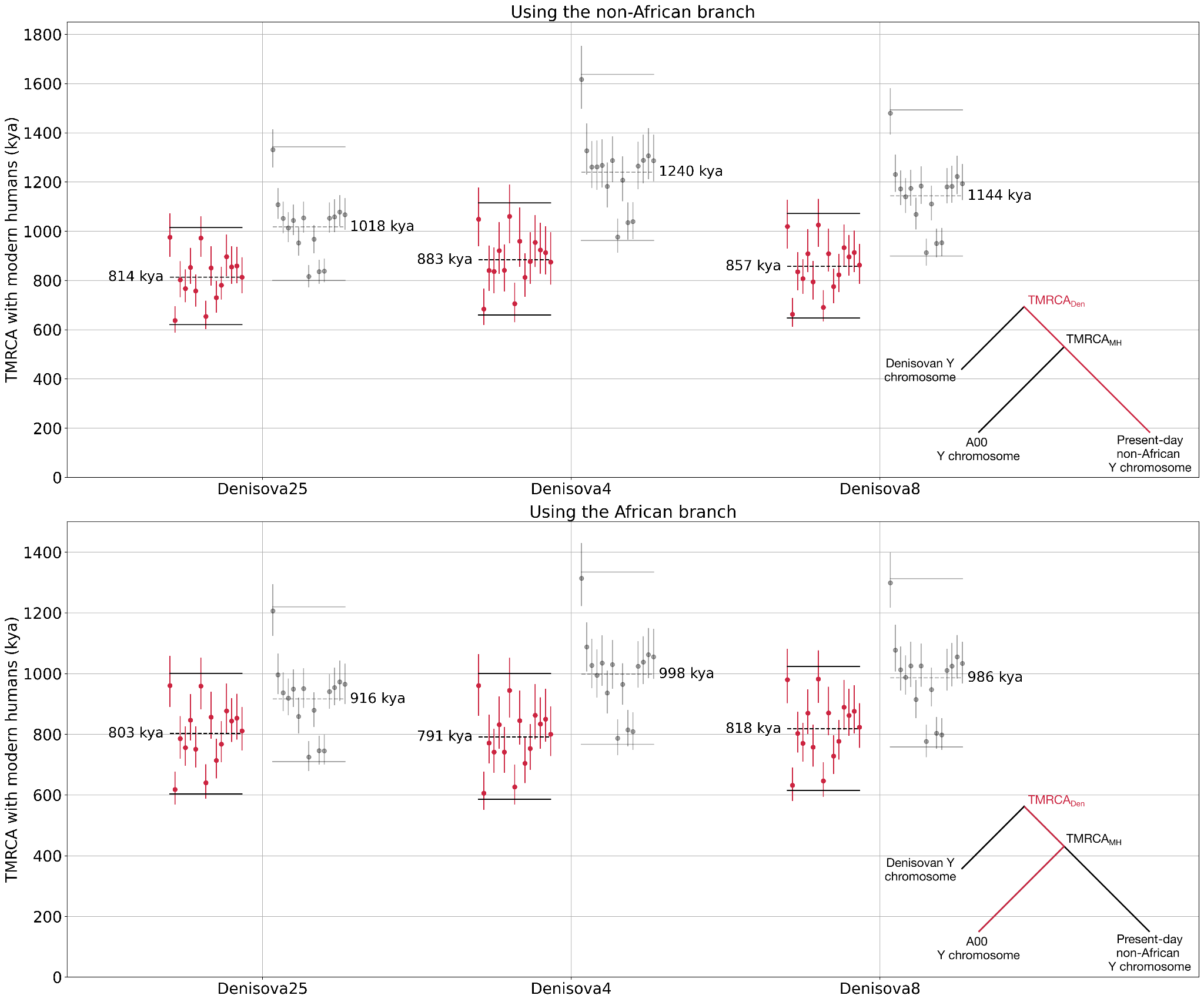

**Figure S17:** **Y chromosome TMRCA estimates between Denisovan Y chromosomes and 15 present-day non-African Y chromosomes.** See the caption of **Figure S15** for more information.

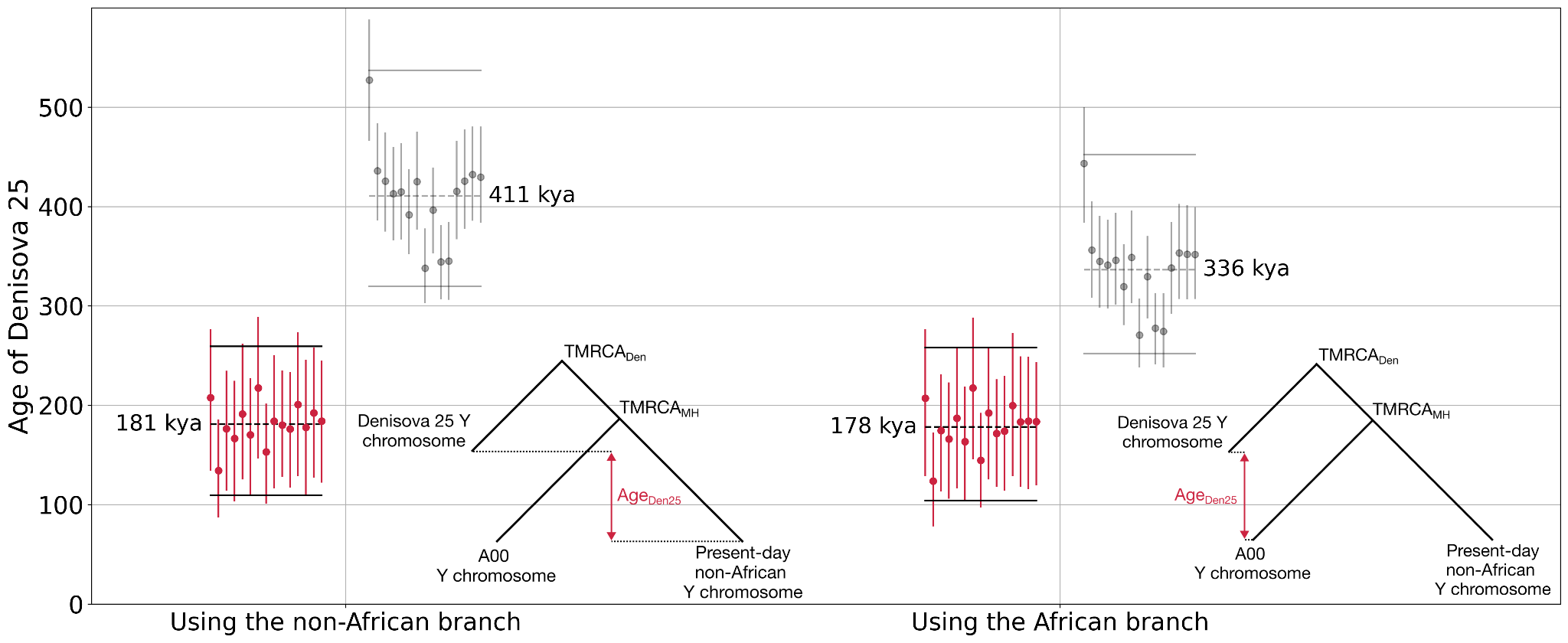

**Figure S18: Estimate of the age of Denisova 25.** This age estimate is based on the number of missing mutations on his Y chromosome compared to those of modern humans, which is then calibrated in years before the present based on our estimate of the TMRCA with modern humans.

#### Bayesian Phylogenetic Analysis

We used *BEAST2* (v2.6.7)^65^ to reconstruct the Y chromosome phylogeny and estimate both the modern human-*Denisova 25* TMRCA and the age of *Denisova 25*. Due to the low coverage available for *Denisova 4* and *Denisova 8*, we did not include their Y chromosome in this analysis.

*BEAST* requires a FASTA file as input, and we used a custom Python script to generate a FASTA file from an input VCF file. To reduce the computational complexity of the Bayesian analysis, only variable sites were included in the input FASTA file. We informed *BEAST* about the nucleotide composition of the invariant sites by editing the *BEAST* .xml file as in Karmin *et al*. (2015)^82^.

The *bModelTest* *BEAST2* package^90^ was used to estimate the best-fitting nucleotide substitution model for the Y chromosome dataset. *bModelTest* averages over all substitution models while simultaneously reconstructing the phylogeny which allowed us to omit the additional step of selecting a substitution model.

The tip dates of the present-day individuals were set to zero and uniform priors were used to represent the ages of the ancient individuals (**Table S33**). For the radiocarbon-dated ancient individuals, we set the range of the priors as the 95% confidence interval of the respective IntCal20 calibration curves. The starting tip date of the *Denisova 25* branch was set to 200,000 years ago and the range of the prior was set as 40,000 to 300,000 years ago. For *Chagyrskaya 2* we set the starting tip date to 64,000 years ago^87^ and we set a uniform prior ranging from 40,000 to 120,000 years ago. We set the initial Y chromosome mutation rate to 7.34x10^-10^ mutations/bp/year as estimated in Petr *et al.* (2020)^79^. To allow *BEAST* to effectively estimate the Y chromosome mutation rate, we set a wide uniform prior of 4x10^-10^ to 10x10^-10^ mutations/bp/year.

We used a path sampling approach (as implemented in the *BEAST2* MODEL_SELECTION package) to select the tree model and the clock model that best fit our data. The following model combinations were tested:

- Strict clock, constant population size
- Relaxed log-normal clock, constant population size
- Strict clock, Coalescent Bayesian Skyline
- Relaxed log-normal clock, Coalescent Bayesian Skyline

For each model combination the marginal likelihood was estimated using 50 path steps, an alpha parameter of 0.3 for the Beta distribution used to space out the steps, and a chain length of 10 million MCMC iterations for each step. The pre-burn-in stage was set at 5 million iterations and an 80% burn-in was used for each chain.

The two model combinations with the highest support based on the marginal likelihood were those with the Coalescent Bayesian Skyline tree model with slightly higher support for the relaxed clock model (**Table S34**). As the support for the relaxed clock model over the strict clock model was not significant, (Bayes factor = 0.67), we used the simpler model of the fixed clock together with the Coalescent Bayesian Skyline tree model to estimate the Y chromosome phylogeny.

For reconstructing the Y chromosome phylogeny, four independent MCMC runs of 55 million iterations were performed. A 10 million iteration pre-burnin stage was used for each run and each MCMC chain was sampled every 5,000 steps. The resulting log files were combined with *logcombiner* (v2.6.7) using a 10% burnin for each run and a single best-supported tree (**Figure S19**) was generated with *TreeAnnotator* (v2.6.7).

The Y chromosome mutation rate was estimated to be 6.78x10^-10^ (95% HPD 6.05x10^-10^ to 7.51x10^-10^) mutations/bp/year. Based on *bModelTest*, the nucleotide substitution model with the highest support is the unnamed 123421 model. The nucleotide substitution rates are summarised in **Table S35**. We estimated that the modern human-*Denisova 25* TMRCA is 784ka (95% Highest Posterior Density, HPD, Interval: 697ka to 876ka) and the age of *Denisova 25* is 232ka (95% HPD Interval: 177ka-288ka). These estimates are consistent with those obtained using the branch length method described above considering the associated large HPD intervals (around 800ka and 180ka for the TMRCA and age, respectively).

**Table S33: Uniform tip date priors for the ancient individuals included in the BEAST analysis.**

| **Individual** | **Starting Tip Date (years before present)** | **Tip Date Prior (years before present)** |
| --- | --- | --- |
| *A00_I10871* | 7,913 | 7,839 - 8,008 |
| *Loschbour* | 8,025 | 7,936 - 8,171 |
| *Yana 1* | 31,850 | 31,532 - 32,200 |
| *Mezmaiskaya 2* | 43,449 | 42,156 - 44,965 |
| *Ust’-Ishim* | 44,366 | 42,904 - 45,930 |
| *Chagyrskaya 2* | 64,000 | 40,000 - 120,000 |
| *Denisova 25* | 200,000 | 40,000 - 300,000 |

**Table S34: Marginal log-likelihood of the model combinations tested using a path sampling approach.** The two best-supported model combinations are marked in bold.

| **Clock Model** | **Tree Model** | **Marginal log-likelihood** |
| --- | --- | --- |
| Strict | Coalescent Constant Population | -3,213,972.91 |
| **Strict** | **Coalescent Bayesian Skyline** | **-3,213,960.59** |
| Relaxed Log Normal | Coalescent Constant Population | -3,213,972.66 |
| **Relaxed Log Normal** | **Coalescent Bayesian Skyline** | **-3,213,959.05** |

**Table S35: Nucleotide substitution rates estimated by bModelTest^90^.**

| **Substitution rates from *bModelTest*** | |
| --- | --- |
| AC | 0.548 |
| AG | 1.852 |
| AT | 0.267 |
| CG | 0.933 |
| CT | 1.852 |
| GT | 0.548 |

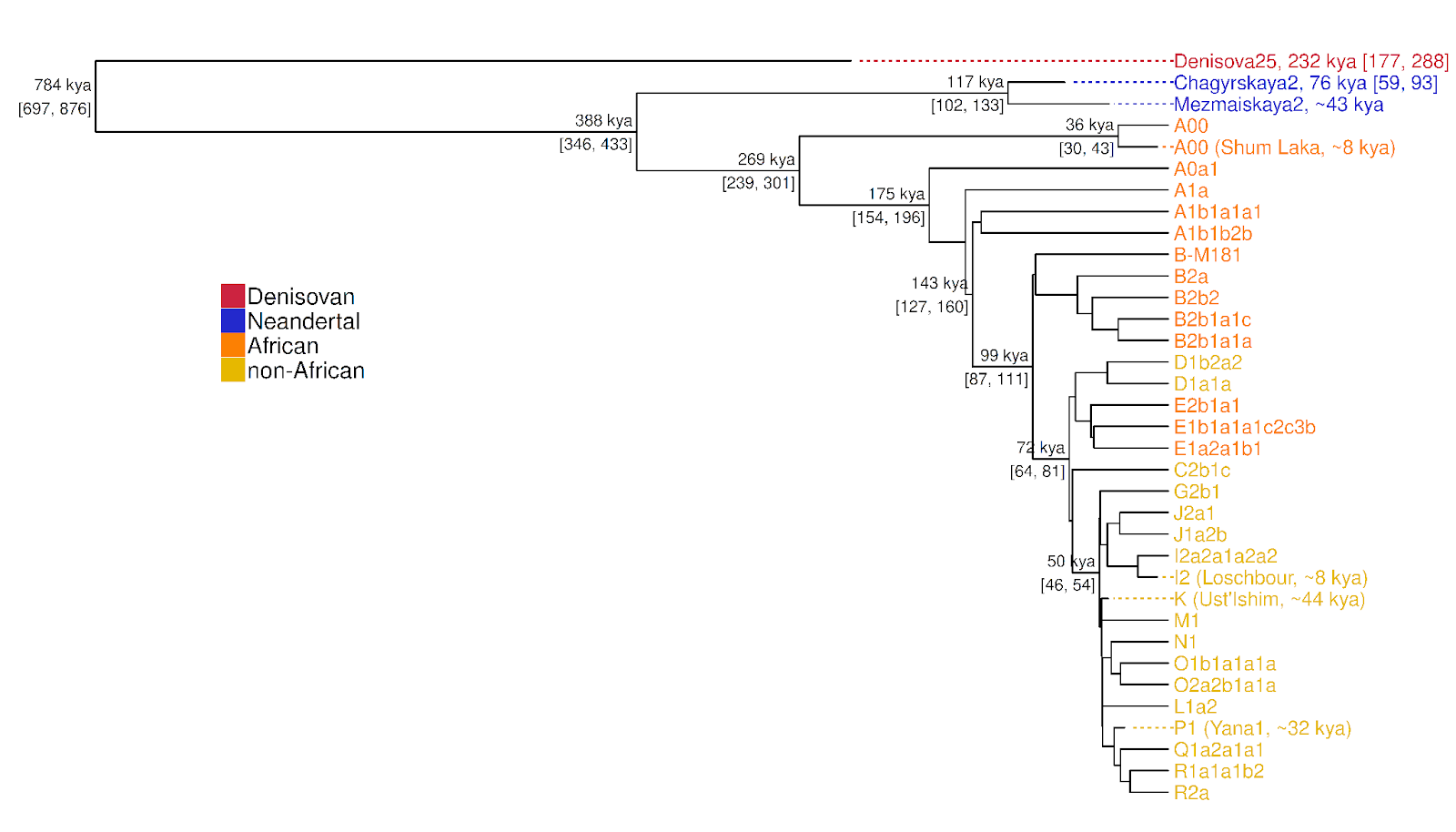

**Figure S19: Y chromosome phylogeny based on a Bayesian analysis.** Calibrated (IntCal20) radiocarbon dates are provided for ancient individuals. Age estimates for Denisova 25 and Chagyrskaya 2 are based on the mean of the respective posterior probability distributions. The numbers in brackets represent the 95% Highest Posterior Density intervals.

### Supplementary Note 6: Nuclear genetic dating of *Denisova 25*

#### Branch shortening compared to other present-day and ancient genomes

To estimate when the *Denisova 25* individual lived, we counted the number of derived alleles on his lineage since the last common ancestor with the great apes and compared it to that of a present-day human, as done previously^45^ and described here for completeness. A smaller number of derived alleles indicates a shorter lineage, so that the difference in the number of derived alleles (*i.e.*, branch shortening) between the present-day human and *Denisova 25* lineages indicates how long ago the *Denisova 25* individual lived. The present-day human genome we used for calibration was from an Mbuti individual (HGDP00982, SS6004471, or B_Mbuti-4)^48^. For comparison, we also counted the number of derived alleles in four high-coverage archaic human genomes^45,47,48,91^ and three ancient modern human genomes, Ust’-Ishim^83^, Loschbour and Stuttgart^85^. For each genome, we randomly sampled one allele at heterozygous sites so that we are effectively measuring the length of a single lineage. We only included bases in the autosomes that align to the reference genomes of four great apes (chimpanzee panTro4, bonobo panpan1.1, gorilla gorGor3, and orangutan ponAbe2) and where all four great apes carry the same allele (different from a gap “-“ or a missing base “N”), which was then considered to be the ancestral allele. Any difference to this ancestral state was counted as a derived allele. We calculated the relative age of each ancient individual as (𝑛_𝑀_ − 𝑛_𝐴_)/𝑛_𝑀_, with 𝑛_𝑀_ and 𝑛_𝐴_ denoting the number of derived alleles in the Mbuti genome and ancient genome, respectively. To arrive at an estimate in years, we assumed a divergence time to the common ancestor of humans and chimpanzee of 13 million years, which corresponds to a mutation rate of 0.5 × 10^−9^ per base pair per year^92^. We computed the variance in the dating estimates with a block-jackknife approach using a window size of 5Mb^93^.

This branch shortening analysis is sensitive to the quality and filtering of the data. To minimize biases due to the different quality of the genomes, we excluded positions that are outside the intersection of the basic filter of *Denisova 25* (**Supplementary Note 3**) and that of the other genomes^45,91^. We also tested different filters on the genotype quality (from no additional filter, **Figure S20** and **Table S36**, to minimum genotype qualities of 10 to 60, **Figure S21**) and excluded transitions. We estimated that the *Denisova 25* individual lived 205ka (195-215ka, 2 standard deviations) and the estimates for the other genomes (**Table S36**) are consistent with previous estimates^45^. In particular, the Stuttgart^85^, Loschbour^85^, Ust’-Ishim^83^ and *Vindija 33.19*^45^ individuals, whose remains were radiocarbon dated to ca. 7ka, 8ka, 45ka and a minimum of 46ka, respectively, lived 5ka (0-13ka), 9ka (0-18ka), 42ka (33-51ka) and 47ka (37-57ka) according to our estimates (intervals correspond to 2 standard deviations around the estimates). The branch shortening estimate for *Denisova 25* is consistent with those obtained from the mitochondrial genome 219ka (95% High Posterior Density Interval: 159-275ka; **Supplementary Note 4**) and Y chromosome 232ka (95% High Posterior Density Interval: 177-288ka; **Supplementary Note 5**).

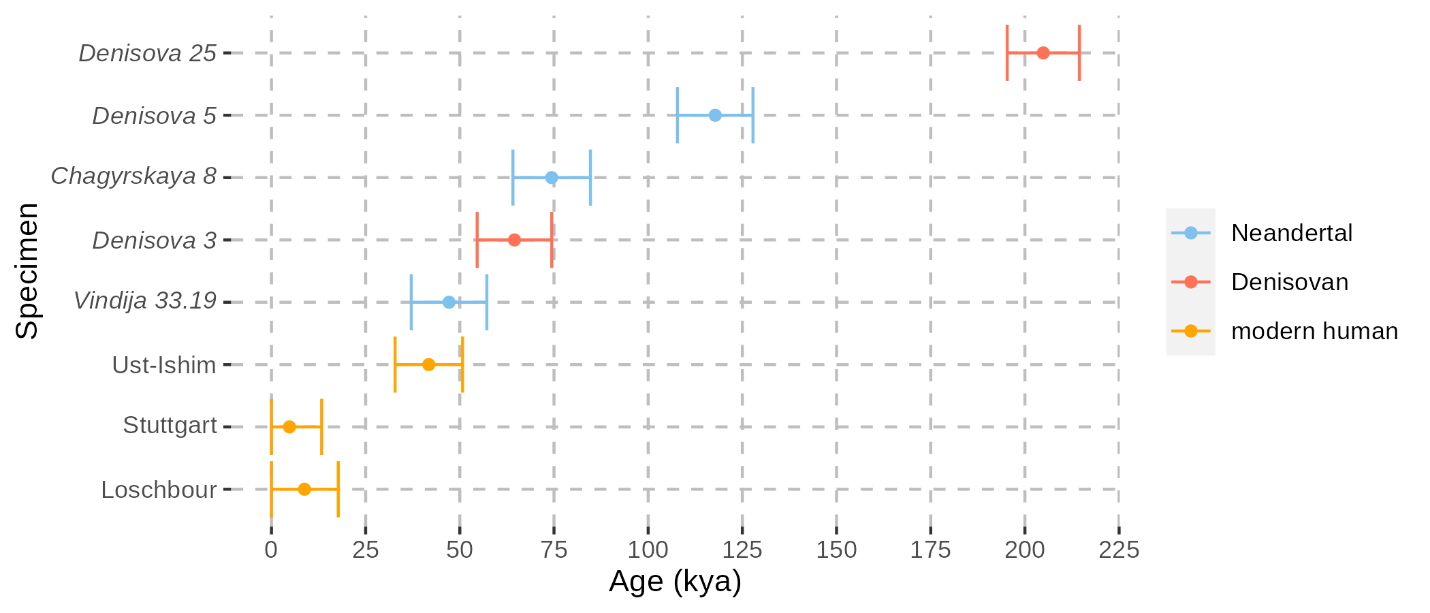

**Figure S20: Autosomal branch shortening estimates.** Bars correspond to two standard deviations computed with a block-Jackknife using a window size of 5Mb. We applied the minimal set of filters described in **Supplementary Note 3** (no genotype quality cutoff was applied; see **Figure S21** for the effect of genotype quality).

**Table S36: Autosomal branch shortening estimates.** These estimates correspond to the data presented in **Figure S20**.

| **Individual** | **Age estimate (years ago)** | **Standard deviation (years)** |
| --- | --- | --- |
| Denisova 25 | 204,880 | 4,791 |
| Denisova 5 (Neandertal) | 117,798 | 5,012 |
| Chagyrskaya 8 | 74,384 | 5,142 |
| Denisova 3 | 64,506 | 4,952 |
| Vindija 33.19 | 47,144 | 5,015 |
| Ust’-Ishim | 41,774 | 4,478 |
| Loschbour | 8,771 | 4,485 |
| Stuttgart | 4,796 | 4,263 |

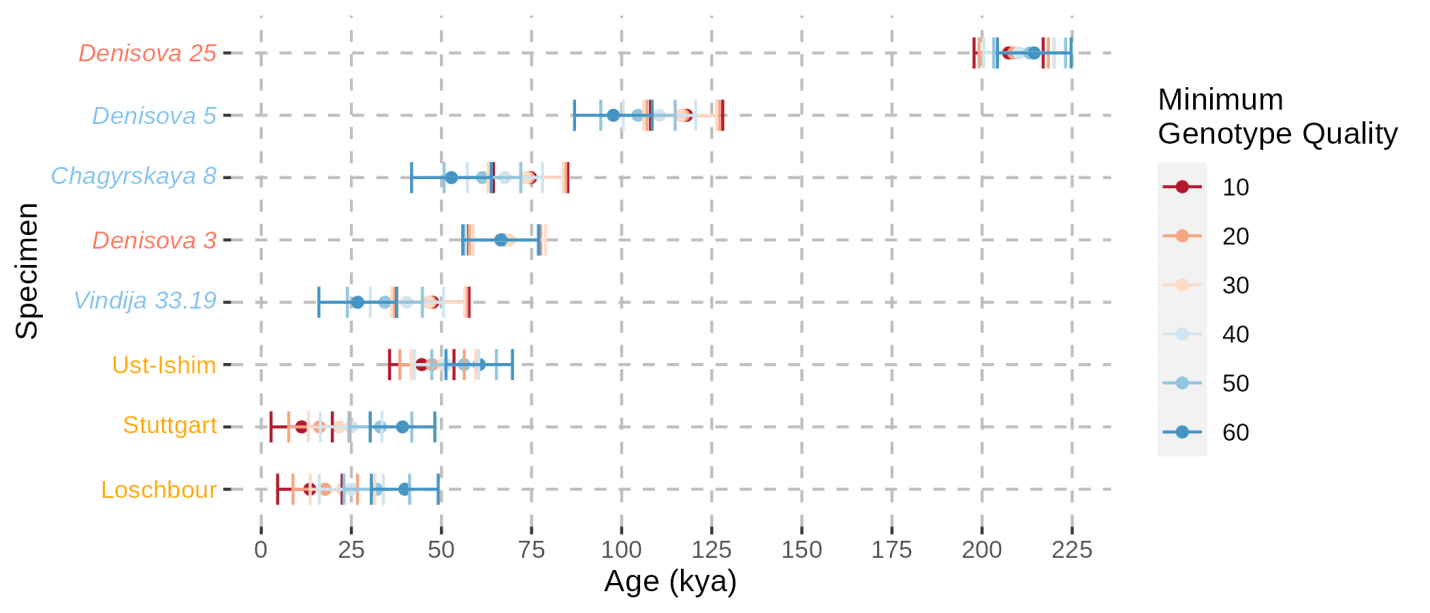

**Figure S21: Effect of filtering based on genotype quality on the autosomal branch shortening estimates.** Bars correspond to two standard deviations computed with a block-Jackknife using a window size of 5Mb. We applied the minimal set of filters described in **Supplementary Note 3** and different minimum genotype quality cutoffs (different colours). Specimen names in red, blue and orange correspond to Denisovans, Neandertals and modern humans, respectively.

#### Relative genetic dating from the demographic histories of *Denisova 3* and *Denisova 25*

To get another estimate of the age of *Denisova 25* from the nuclear genome, we compared the history of population size changes for the population of *Denisova 3* and that of *Denisova 25*, as estimated with the *Pairwise Sequential Markovian Coalescent* (*PSMC*^94^; see **Supplementary Note 7**). Both demographic histories should match up to the time that the ancestors of *Denisova 3* and *Denisova 25* separated. By truncating the demographic history of the more recent individual at a time T in the past and testing the fit of this truncated demography to the genome of the older individual, Fu *et al*. (2014)^83^ have shown that it is possible to estimate the time difference between when the two individuals lived. This method has the advantage of not relying exclusively on private mutations, which can be affected by sequencing or genotyping errors^83^.

We followed the procedure described in Mafessoni *et al*. (2020)^91^, which takes into account the effects that different filters have on the demographic estimates for different genomes, with a slight modification in the first step:

1) Simulate whole genomes with *scrm*^95^ using the corrected demographic history of *Denisova 3* (see **Supplementary Note 7**), by sampling the genomes in the past with increasing ages (instead of using the script psmc_trunc.pl from the *PSMC* suite of tools as reported in Mafessoni *et al.* (2020)^91^. We varied age differences between 0 and 5,600 generations in steps of 175 generations (approximately 0 to 160,000 years, in steps of 5,000 years, assuming a generation time of 29 years).
2) Apply the filters of the *Denisova 25* genome to those simulated genomes and convert these genomes into input files for *PSMC,* using a custom python script scrm2psmc.py available at https://github.com/StephanePeyregne/calibratePSMC together with an example of a snakemake pipeline to run the whole procedure.
3) Run *PSMC* on the filtered version of the simulated data to get a truncated demographic history of *Denisova 3* as inferred from a genome filtered as *Denisova 25.*
4) Run *PSMC* on the *Denisova 25* genome while setting the demographic history to that estimated with the simulated data, with the following command: psmc -N1 -i simulated_demography -o output Denisova25.psmcfa. This computes the likelihood of the *Denisova 25* data given that demographic history.
5) Identify the time difference that maximizes the likelihood, thus representing an estimate of the time difference between when the two individuals lived.

We evaluated uncertainty in the estimate by simulating ten genomes for each age difference, and retained the 5% top simulations (among 330) with the highest likelihood. The retained simulations correspond to an age difference of 3,500 to 4,725 generations between *Denisova 3* and *Denisova 25*. Assuming that *Denisova 3* lived 72,000 years ago^45^ and a generation time of 29 years, *Denisova 25* most likely lived between 174,000 and 209,000 years ago (**Figure S22**). This is again consistent with the other branch shortening estimates described above and in previous sections.

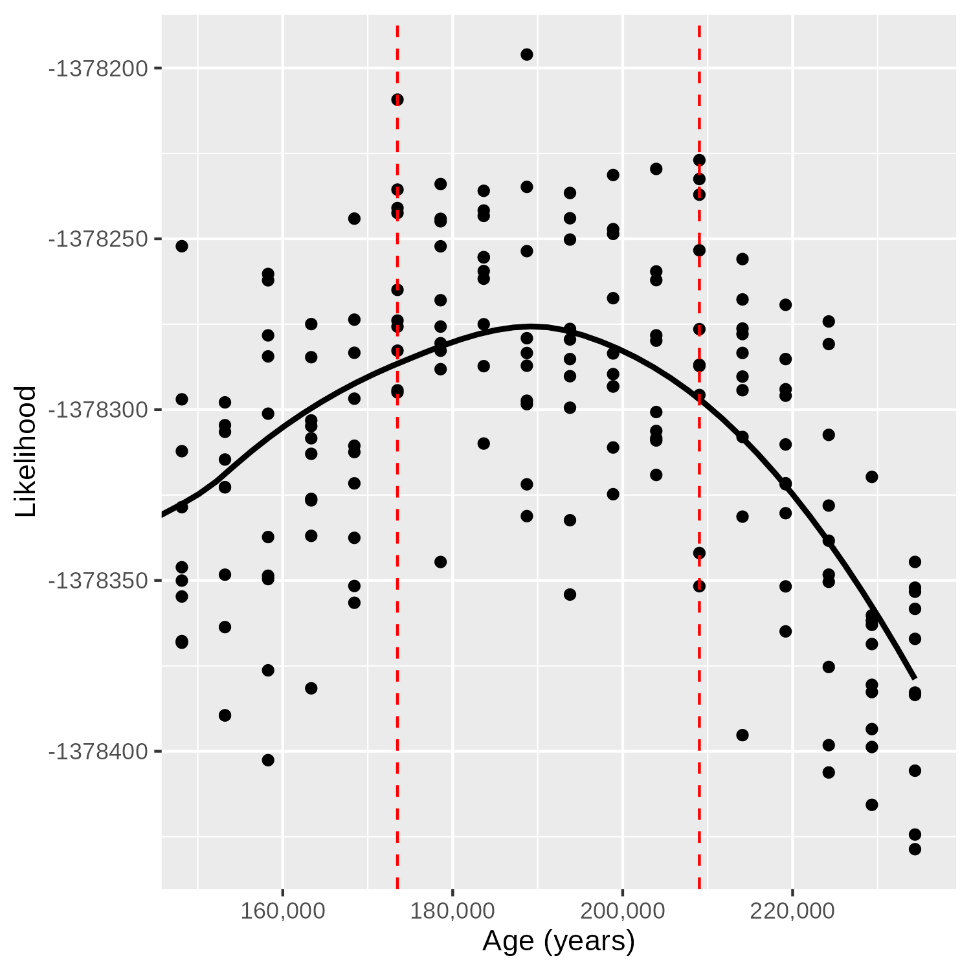

**Figure S22: Dating of Denisova 25 using the inferred demographic history of Denisovans.** Each point represents the likelihood of a particular truncation of the demographic history of Denisova 3 given the Denisova 25 genome. The curve corresponds to a LOESS approximation of the likelihood surface. The vertical red dashed lines highlight the minimum and maximum age difference among the 5% top simulations with the highest likelihood. The difference in the number of generations between Denisova 3 and Denisova 25 was converted into an age before present by assuming that Denisova 3 lived 72,000 years ago^45^ and a generation time of 29 years. The plot is limited to between 150,000 and 235,000 years ago, but the likelihood decreases further for both more recent and older ages.

### Supplementary Note 7: Population size

#### Heterozygosity

Previous studies have shown that the heterozygosity in *Denisova 3* is slightly higher than in Neandertal genomes^45,48,96^. To test whether this holds true, we computed the heterozygosity of *Denisova 25* and compared it with those of *Denisova 3* and three high-coverage Neandertal genomes (*Denisova 5*^48^), *Chagyrskaya 8*^91^, and *Vindija 33.19*^45^; using snpAD genotype calls from Prüfer *et al.* (2017)^45^ and Mafessoni *et al.* (2020)^91^. We calculated heterozygosity as the number of heterozygous sites over the total number of sites that passed the minimal set of filters for each genome. We found that heterozygosity is similar in both Denisovan genomes and slightly higher than in Neandertal genomes. As Neandertal ancestry that is in the heterozygous state in the Denisovan genomes would inflate heterozygosity, we next masked the Neandertal segments with a minimum length of 0.05cM that could be confidently identified with *admixfrog* (version 0.7.2^97^; using a penalty of 0.4 and a window size of 0.005cM, which includes more putative Neandertal segments than other parameter values; see **Supplementary Note 9**). This resulted in lower values of heterozygosity in both Denisovan genomes that are comparable to those observed in the Neandertal genomes (**Figure S23**). After masking regions that are homozygous by descent due to consanguinity (described in the next section), the heterozygosity of *Denisova 3* is even lower than that of Neandertals, whereas the heterozygosity of *Denisova 25* and Neandertals is similar (**Figure S24**). We conclude that the observed difference in heterozygosity between Denisovans and Neandertals does not reflect differences in population size, but rather is a consequence of the presence of Neandertal ancestry in the Denisovan genomes.

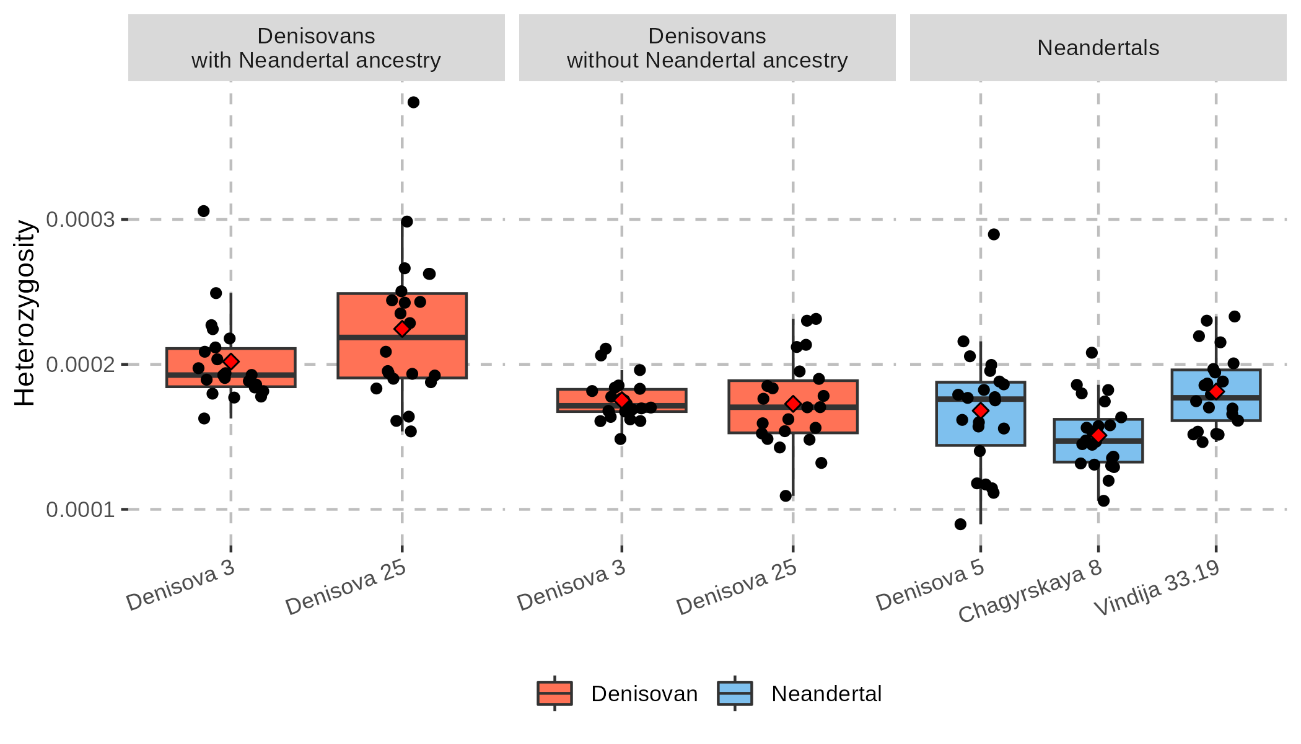

**Figure S23: Estimates of heterozygosity in high-coverage archaic human genomes.** Each box plot represents the heterozygosity distribution across the 22 autosomes, with different colours for Neandertals and Denisovans. The black dots represent the heterozygosity for each chromosome, and the red diamonds correspond to the average over all autosomes (for the overall genome-wide heterozygosity, see **Table S37**). The two panels on the left correspond to the results for the Denisovan genomes with and without masking Neandertal segments of at least 0.05cM identified with admixfrog using a penalty of 0.4 and a window size of 0.005cM (see **Supplementary Note 9**).

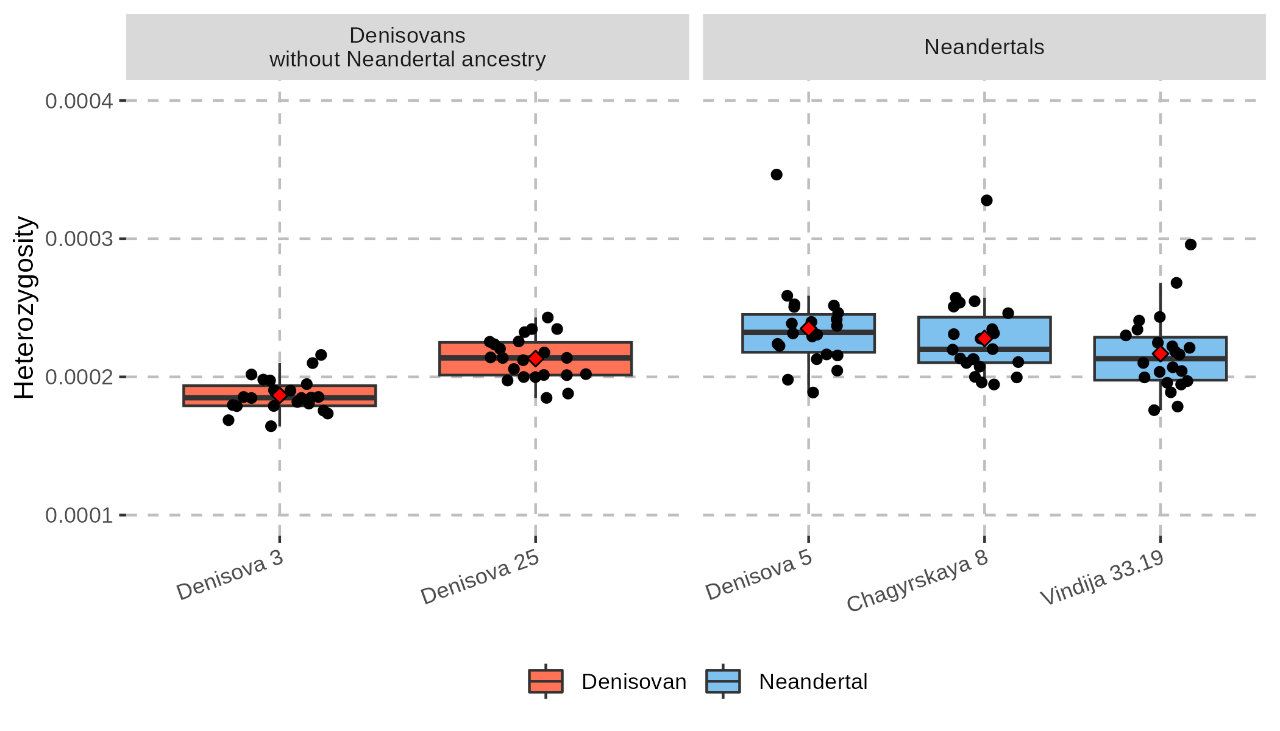

**Figure S24: Estimates of heterozygosity in high-coverage archaic human genomes after excluding homozygous-by-descent (HBD) segments.** HBD segments longer than 1Mb, identified using the best π parameter value for each genome (see next section below; **Figure S25**), were excluded. Neandertal segments of at least 0.05cM identified with admixfrog using a penalty of 0.4 and a window size of 0.005cM were masked in the Denisovan genomes (see **Supplementary Note 9**). See **Figure S23** for more details on the figure layout.

**Table S37: Average autosomal heterozygosity in high-coverage archaic human genomes.** The estimates in the regions of the Denisovan genomes “without Neandertal ancestry” were computed after masking the Neandertal segments of at least 0.05cM identified with admixfrog using a penalty of 0.4 and a window size of 0.005cM (see **Supplementary Note 9**).

| **Individual** | **Population** | **Heterozygosity** | |
| --- | --- | --- | --- |
|  |  | **Whole genome** | **Without Neandertal ancestry** |
| *Denisova 3* | Denisovan | 0.00020 | 0.00017 |
| *Denisova 25* |  | 0.00021 | 0.00017 |
| *Chagyrskaya 8* | Neandertal | 0.00015 | - |
| *Denisova 5* |  | 0.00017 | - |
| *Vindija 33.19* |  | 0.00018 | - |

#### Runs of homozygosity

We detected regions that are homozygous by descent (HBD) due to consanguinity in the genome of *Denisova 25* using a previously described method^45,48,91,98^. Briefly, the method uses genotype data to identify continuous runs of homozygosity (ROH) of at least 50kb, and then merges these windows into larger HBD tracts of at least 1Mb if a minimum proportion π of positions in that tract are in ROH. This parameter π is measured in overlapping, sliding windows of 1Mb with an offset of 100kb and allows some 10kb windows with heterozygous sites in each 1Mb window to account for genotyping errors and rare *de novo* mutations. To determine the adequate π parameter for each genome, we tested a set of values (π = 0.8, 0.825, …, 0.975, 0.99), and plotted the total length of regions of at least 2.5cM using a uniform recombination rate of 1.3cM/Mb, the African American recombination map^99^ or the deCode recombination map^100^ (**Figure S25**, **Figure S26**, and **Figure S27**). Following Sümer *et al.* (2025), the most adequate value of π was chosen as the one that yields the highest proportion of the genome in long HBD segments (≥10cM) relative to the total proportion in segments of at least 2.5cM. We then quantified inbreeding by dividing the total number of sites within the HBD segments by the total number of sites that passed the minimal set of filters for each genome (**Table S38**).

We found that 15.5% of the *Denisova 25* genome is in HBD segments compared to 4.6% for *Denisova 3*. *Denisova 25* carries particularly more segments of at least 10cM, covering 8.9% of the genome compared to 0.5% for *Denisova 3*. The fraction of the *Denisova 25* genome in long HBD segments is in fact similar to that observed for the *Denisova 5* (13%) and *Chagyrskaya 8* (8.3%) Neandertal genomes, with values that are within the variation expected for various scenarios of recent consanguinity, including parents that were double first cousins to half siblings^48^. The length distribution and number of HBD segments in *Denisova 25* are most similar to those observed for the *Denisova 5* and *Chagyrskaya 8* Neandertal (**Figure S28**). Altogether, these results suggest that these levels of consanguinity might not represent a difference between Neandertals and Denisovans, but rather differ over time, with higher levels of consanguinity among the earlier individuals. More Neandertal and Denisovan genomes across space and time will be necessary to explore the cultural, demographic and environmental factors that may have influenced this variation.

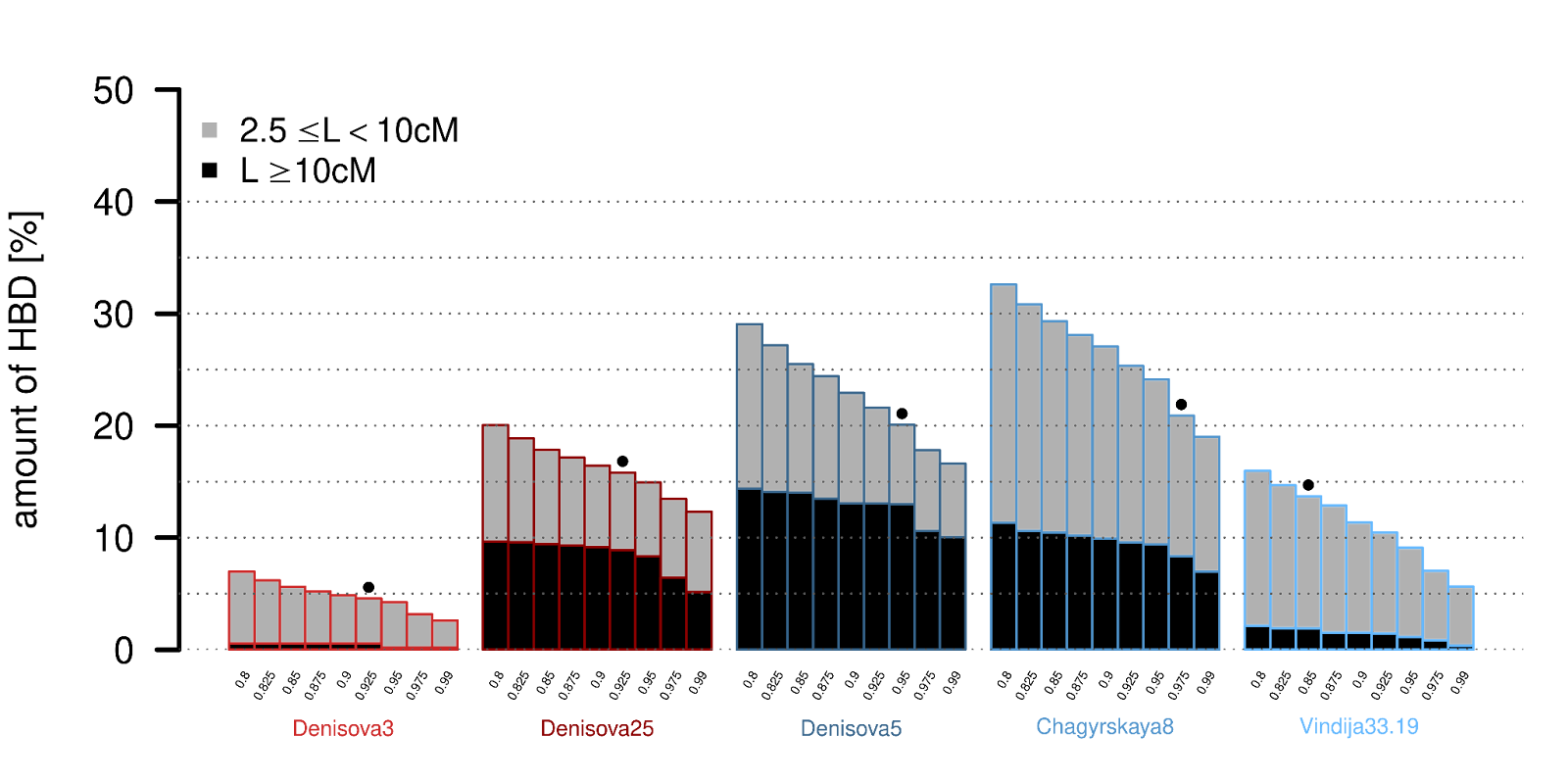

**Figure S25: Amount of HBD with different π parameters in high-coverage archaic human genomes, using a uniform recombination rate of 1.3cM/Mb.** Best value of π is indicated for each genome.

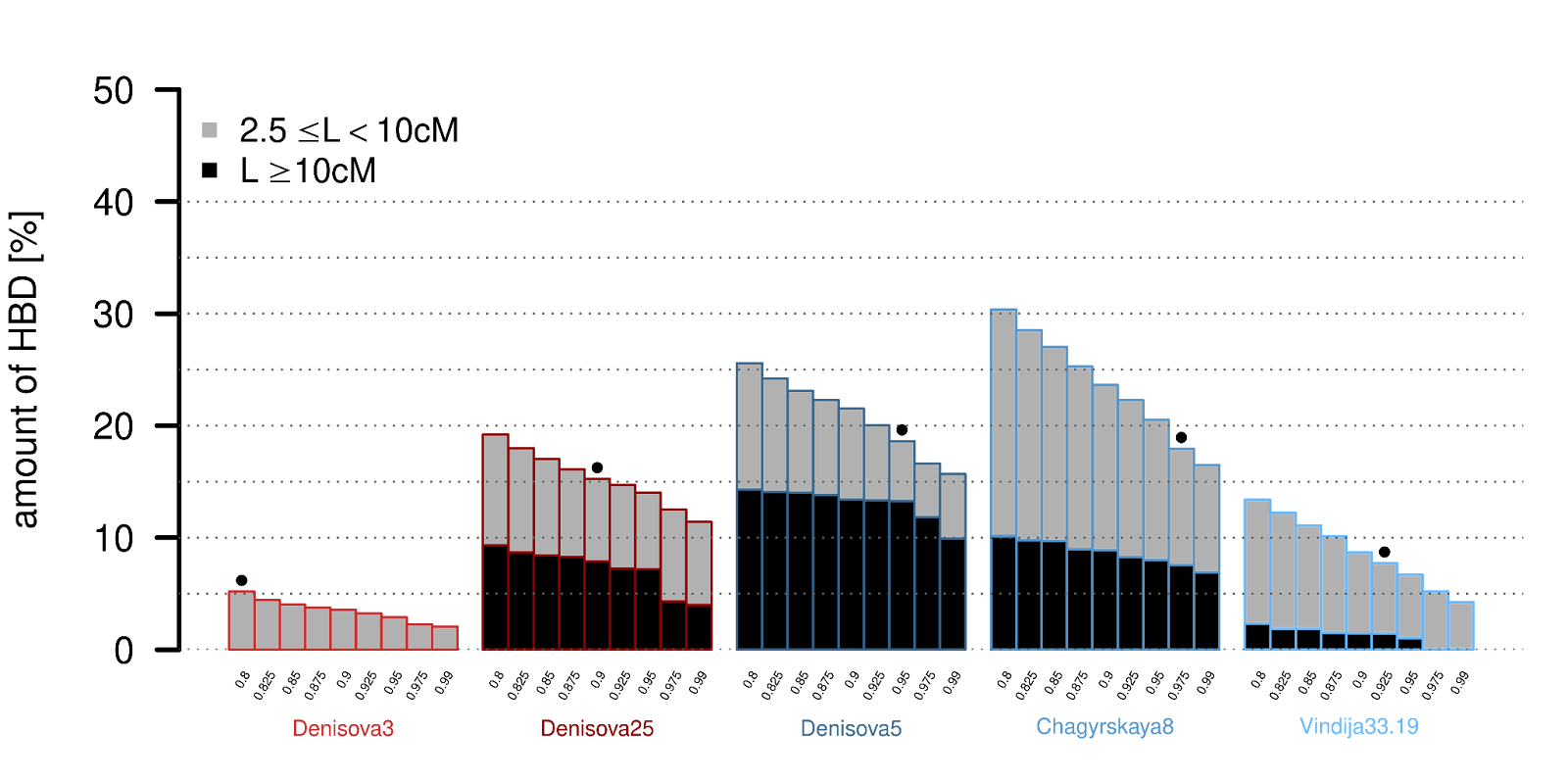

**Figure S26: Amount of HBD with different π parameters in high-coverage archaic human genomes, using the African American recombination map.** Best value of π is indicated for each genome.

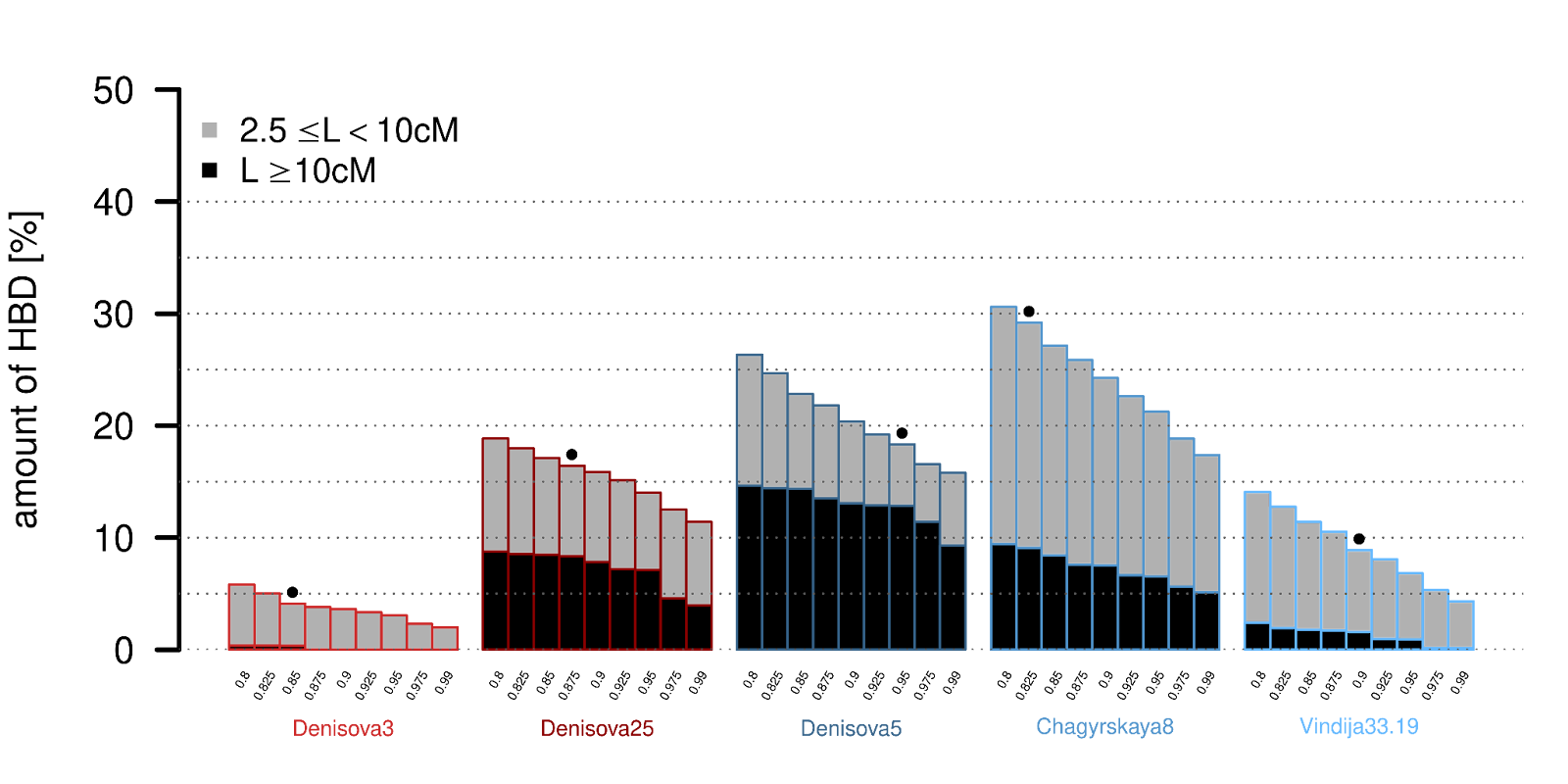

**Figure S27: Amount of HBD with different π parameters in high-coverage archaic human genomes, using the deCode recombination map.** Best value of π is indicated for each genome.

**Table S38: Amount of HBD in different high-coverage archaic human genomes with their best π values, using a uniform recombination rate and the two recombination maps.** The table shows the amount of HBD in percentage, with lengths higher or equal to 2.5cM (all) and 10cM (long). These values are equivalent to those in **Figure S25**-**Figure S27**.

| **Samples** | **Proportion of HBD [%]** | | | | | |
| --- | --- | --- | --- | --- | --- | --- |
|  | **Uniform recombination rate** | | **African American map** | | **deCode map** | |
|  | **long** | **all** | **long** | **all** | **long** | **all** |
| *Denisova 3* | 0.5 | 4.6 | 0 | 5.2 | 0.4 | 8.5 |
| *Denisova 25* | 8.9 | 15.8 | 7.9 | 15.3 | 8.3 | 16.4 |
| *Denisova 5* | 13.0 | 20.1 | 13.3 | 18.6 | 12.9 | 19.2 |
| *Chagyrskaya 8* | 8.3 | 20.9 | 7.6 | 17.9 | 9.1 | 29.2 |
| *Vindija 33.19* | 1.9 | 13.7 | 1.5 | 7.7 | 1.6 | 8.9 |

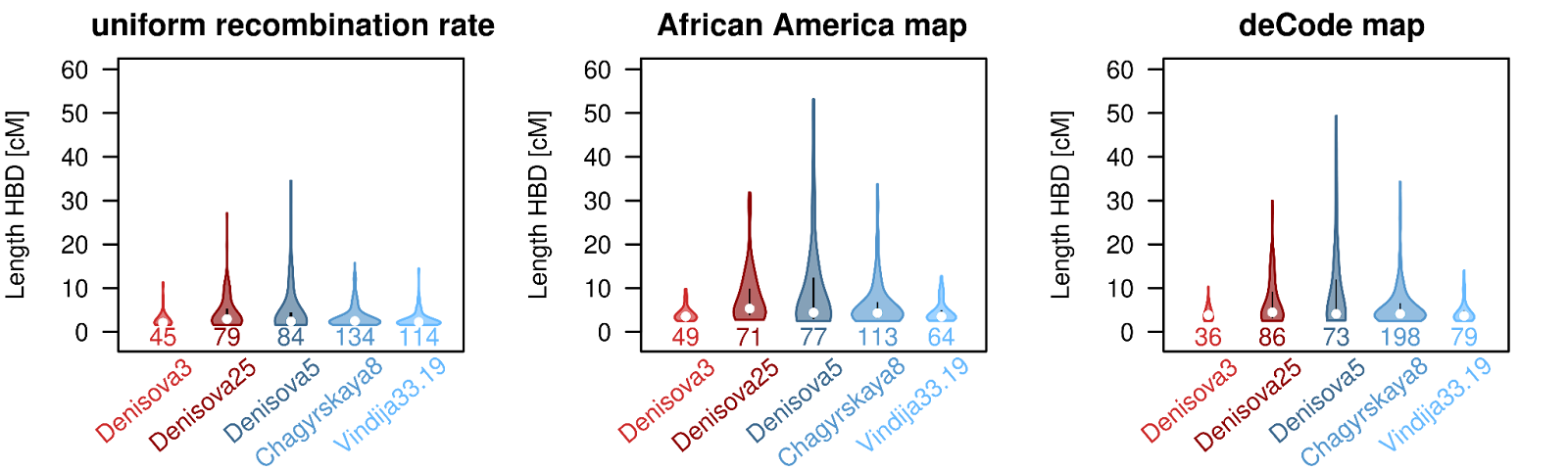

**Figure S28:  HBD length distribution and the number of segments, using different recombination rates**. The width of the violins is proportional to the number of HBD segments reported under the violins and the points represent the median of each distribution.

#### Inference of local population structure

To infer the local structure of the Denisova 25 population, we applied an analytical approach previously used for archaic genomes^91^. This method leverages the fact that most HBD segments result from recent coalescence events, providing insights into the size of the deme or subpopulation in which an individual lived, assuming a model where the population is subdivided into multiple demes with migrations between them. For completeness, we briefly outline this approach below.

##### Proportion of HBD segments in an island model

To estimate $H\left( L \right)$, the probability that a DNA segment of length $L$ is homozygous by descent (HBD) in the genome of an individual, we assume that HBD segments arise from a common ancestor within a deme. Coalescent theory asserts that the two chromosomal segments an individual carries find a common ancestor with probability $\frac{1}{2N}$ per generation, where $N$ is the effective population size. Migration into the deme occurs with probability $2m$, where $m$ is the migration rate per generation. For a segment to remain HBD, it must avoid mutation and recombination, which we assume always disrupt homozygosity. The probability of recombination affecting a segment of length $L$ is$2rL$ , where $r$ is the recombination rate per Morgan ($r=1$), and the probability of mutation affecting it is $2\mu L$, where $\mu$ is the mutation rate per Morgan per generation. The probability of HBD for a segment of length $L$ is given by	$H\left( L \right)=\frac{\frac{1}{2N}}{\frac{1}{2N}+2m+2rL+2\mu L}$ .	(1)

Because our approach to identifying HBD segments permits a few differences between the two haplotypes of an individual to account for sequencing errors, new mutations since the last common ancestor do not disrupt an HBD segment. Therefore, we set the mutation rate to zero, following Mafessoni *et al*., (2020).

##### Parameter estimation

To estimate the population structure parameters for archaic humans, we calculate the probability that a randomly chosen tract of length $L$ is HBD. This probability is determined by counting the possible positions where a random tract may fall. We consider the total genome length $L_{tot}$ and the number of autosomes, all longer than $L$. Since the last $L$ of each chromosome cannot serve as a starting point without extending beyond the chromosome's end, the number of valid starting points for a tract of length L is $L_{tot}-L$. Similarly, for each observed HBD segment, with $L$ _i_ corresponding to the length of this i-th HBD segment, a randomly chosen tract of length $L$ is HBD if its starting point falls within the first $L_{i}-L$ bases of that HBD segment. Since HBD segments do not overlap, the total number of possible starting points for a random tract to be HBD is thus $\sum_{i|L_{i}>L} (L_{i}-L)$ and the overall probability $H\left( L \right)$is the sum of the probabilities for each HBD segment: $H\left( L \right)=\sum_{i|L_{i}>L} \frac{L_{i}-L}{L_{tot}-22L}$. We computed $H\left( L \right)$ from the observed data for tract lengths (L) between 1 and 10cM in steps of 1cM. The total genetic length ($L_{tot}$) was measured as the sum of genomic regions passing the standard filters (about 3,540cM for all genomes), including gaps shorter than 100kb since HBD segments are at least 1Mb and include missing data. Including larger gaps had a minimal effect, producing similar values for $L_{tot}$. Finally, we estimated parameters by fitting numerically $H\left( L \right)$ using equation (1) with the non-linear least squares method implemented in the *nlsLM* function of the *minpack.lm* library in R. Standard deviations were computed from the covariance matrix extracted with the *vcov* function.

##### Observed HBD distributions

The observed distribution of HBD sequence lengths is well-captured for the *Denisova 25* genome (**Figure S29**), and the *Denisova 5* and *Chagyrskaya 8* Neandertal genomes, but less so for the *Denisova 3* and *Vindija 33.19* genomes, where the observed HBD-region lengths are inconsistent with a low effective deme size. Our analysis suggests that the *Denisova 25* and *Chagyrskaya 8* genomes come from populations with effective deme sizes between 50 and 60 individuals, with approximately one migrant per generation (**Table S39**). In contrast, the *Denisova 5* Neandertal genome originates from a population with an effective deme size of around 20 individuals, also with about one migrant per generation. It is important to note that we estimate the effective size of the demes, which may significantly differ from the census size due to high variance in reproductive success among individuals. Moreover, our estimates are sensitive to the value of the π parameter for identifying HBD segments, and may be twice as large using the strictest value of 0.99 (***Figure S30***; ***Table S40***).

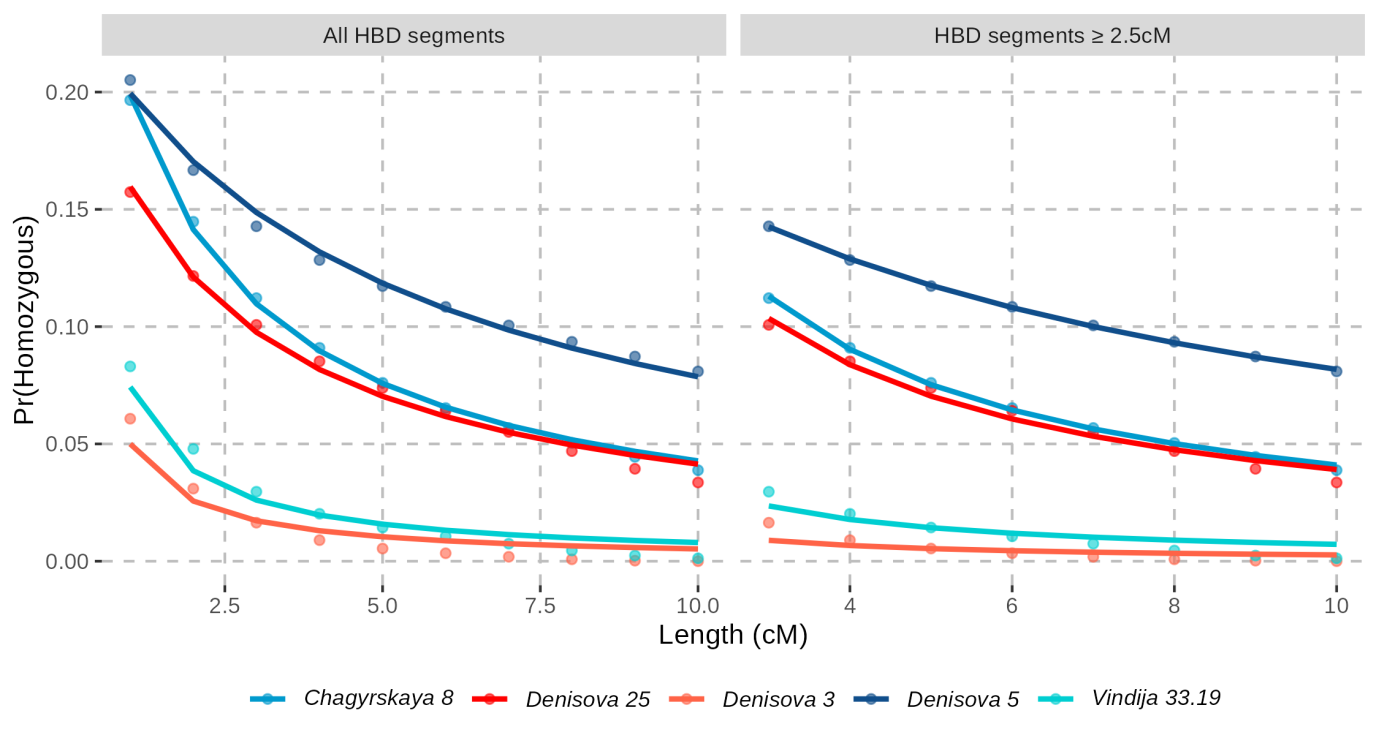

**Figure S29: Observed and expected HBD probabilities in archaic humans**. We show the probability that a randomly chosen genomic segment of length L cM is HBD. Data points represent observed values for π values determined with the African-American map, while the lines correspond to the best-fit estimates. Each individual is represented by a distinct colour, with red shades for Denisovans and blue shades for Neandertals. Estimated parameter values are provided in **Table S39**. The left panel presents results from all HBD segments (of at least 1cM for the first data point), while the right panel excludes segments shorter than 2.5cM.

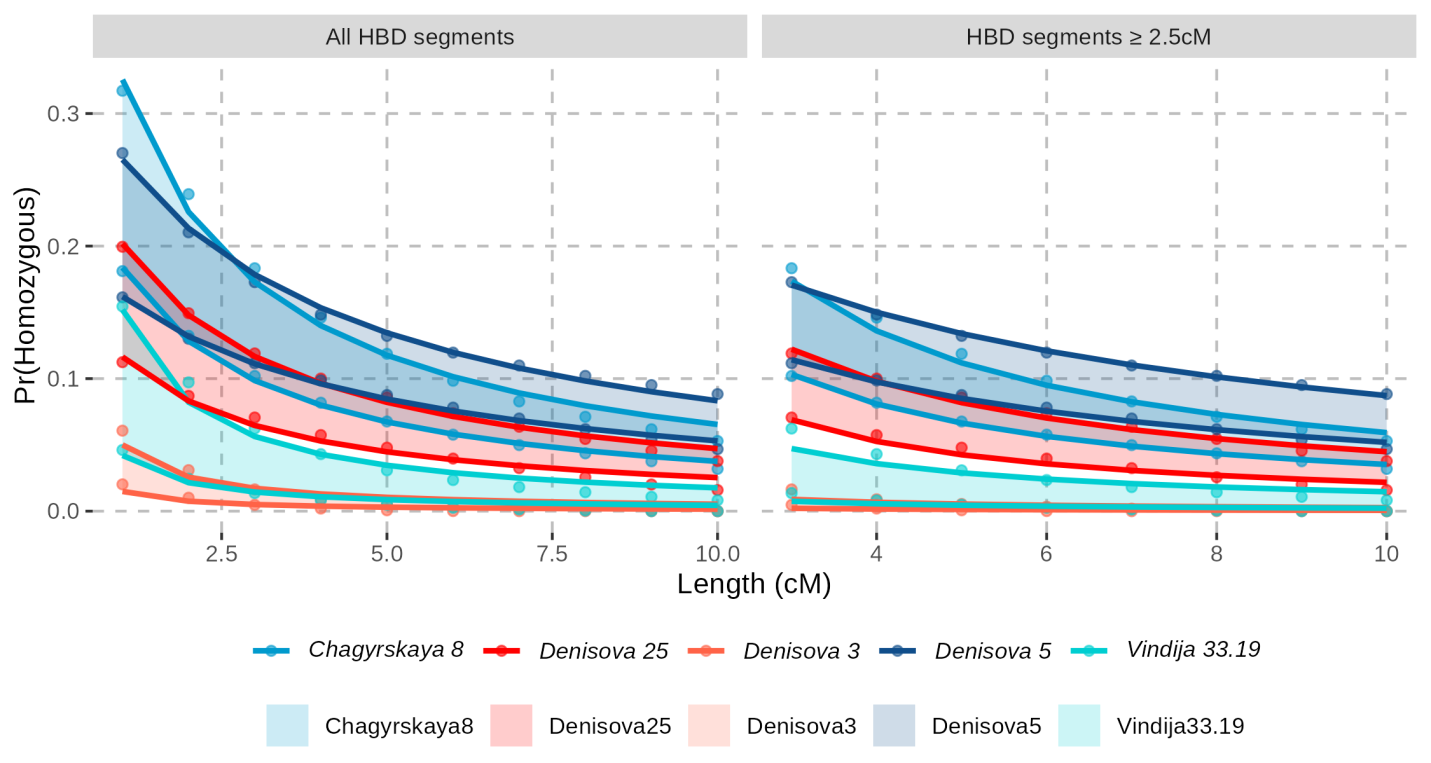

**Figure S30: Range of observed and expected HBD probabilities in archaic humans depending on the value of π.** For each archaic genome, the top and bottom lines correspond to the best-fit estimates for π=0.8 and π=0.99, respectively. Estimated parameter values are provided in **Table S40**. See **Figure S29**for more details.

**Table S39: Estimates of deme sizes (N) and migration rates (m).** Values correspond to the mean and standard deviation for the parameters N and m.

| **Individual** | **HBD segments** | **N** | **m** | **sd(N)** | **sd(m)** |
| --- | --- | --- | --- | --- | --- |
| *Denisova 25* | all | 49.7 | 0.016 | 2.3 | 0.002 |
|  | ≥2.5cM | 56.9 | 0.008 | 4.0 | 0.004 |
| *Denisova 3* | all | 475.1 | 0.0 | 127.8 | 0.003 |
|  | ≥2.5cM | 927.7 | 0.0 | 762.0 | 0.032 |
| *Denisova 5* (Neandertal) | all | 21.4 | 0.037 | 0.8 | 0.002 |
|  | ≥2.5cM | 18.6 | 0.051 | 0.2 | 0.001 |
| *Vindija 33.19* | all | 311.6 | 0.0 | 54.2 | 0.002 |
|  | ≥2.5cM | 345.4 | 0.0 | 134.6 | 0.015 |
| *Chagyrskaya 8* | all | 51.0 | 0.010 | 1.1 | 0.001 |
|  | ≥2.5cM | 55.5 | 0.005 | 1.1 | 0.001 |

**Table S40: Range of estimates for deme sizes (N) and migration rates (m) depending on the value of π.** Values correspond to the mean and standard deviation for the parameters N and m.

| **Individual** | **HBD segments** | ***π*** | **N** | **m** | **sd(N)** | **sd(m)** |
| --- | --- | --- | --- | --- | --- | --- |
| *Denisova 25* | all | 0.8 | 45.2 | 0.012 | 1.8 | 0.001 |
|  |  | 0.99 | 85.9 | 0.012 | 7.5 | 0.003 |
|  | ≥2.5cM | 0.8 | 50.4 | 0.006 | 3.5 | 0.003 |
|  |  | 0.99 | 112.6 | 0.0 | 14.3 | 0.005 |
| *Denisova 3* | all | 0.8 | 475.1 | 0.0 | 127.8 | 0.003 |
|  |  | 0.99 | 1,654.8 | 0.0 | 657.6 | 0.005 |
|  | ≥2.5cM | 0.8 | 927.7 | 0.0 | 762.0 | 0.032 |
|  |  | 0.99 | 3,846.9 | 0.0 | 4,522.6 | 0.046 |
| *Denisova 5* (Neandertal) | all | 0.8 | 22.9 | 0.020 | 0.7 | 0.001 |
|  |  | 0.99 | 35.3 | 0.027 | 1.2 | 0.002 |
|  | ≥2.5cM | 0.8 | 20.0 | 0.031 | 0.5 | 0.002 |
|  |  | 0.99 | 37.6 | 0.022 | 2.2 | 0.004 |
| *Vindija 33.19* | all | 0.8 | 139.5 | 0.0 | 15.6 | 0.001 |
|  |  | 0.99 | 569.3 | 0.0 | 126.7 | 0.003 |
|  | ≥2.5cM | 0.8 | 168.1 | 0.0 | 54.1 | 0.013 |
|  |  | 0.99 | 1,110.8 | 0.0 | 947.9 | 0.033 |
| *Chagyrskaya 8* | all | 0.8 | 33.9 | 0.005 | 1.8 | 0.001 |
|  |  | 0.99 | 58.9 | 0.009 | 1.9 | 0.001 |
|  | ≥2.5cM | 0.8 | 39.7 | 0.0 | 3.2 | 0.003 |
|  |  | 0.99 | 66.9 | 0.003 | 2.2 | 0.001 |

#### Population size changes over time

We used the *Pairwise Sequentially Markovian Coalescent* (*PSMC*, version 0.6.5^94^) to reconstruct the demographic history of the population of *Denisova 25*. For comparison, we also analyzed the previously published high-coverage genomes from *Denisova 3*^47^, three Neandertals (*Denisova 5*^48^, *Chagyrskaya 8*^91^, and *Vindija 33.19*^45^), three ancient modern humans (Loschbour^85^, Stuttgart^85^, and Ust’-Ishim^83^), and two present-day humans (Mbuti HGDP00982 and French HGDP00533^48^).

Input files were prepared by converting the genotypes (*snpAD* calls from Prüfer *et al.* (2017)^45^ and Mafessoni *et al*. (2020)^91^, or GATK calls from Prüfer *et al*. (2014)^48^ for the two present-day human genomes; all filtered as described in **Supplementary Note 3**) into the psmcfa format using a custom python script scrm2psmc.py available at https://github.com/StephanePeyregne/calibratePSMC. This conversion involved splitting each genome into windows of 100bp, and assigning a state to each window based on the presence of at least one heterozygous site (state K), all called sites being homozygous (state T), or more than 10% missing data (state N). After excluding the sex chromosomes, the psmcfa files, one for each chromosome, were concatenated.

We ran *PSMC* using the following command: psmc -N25 -t15 -r5 -p "4+25*2+4+6" -o output.psmc input.psmcfa, where N is the maximum number of iterations, r is the ratio between θ/ρ (where θ and ρ are the scaled mutation and recombination rates, respectively), and -p is a string that specifies time intervals (see https://github.com/lh3/psmc). To prevent overfitting, we used the results from the 20th iteration from a total of 25 iterations, following the default setting of psmc2history.pl^94^, which extracts the demographic history from the output of *PSMC*. As *PSMC* is sensitive to the missing data caused by filtering^45,91^, we followed the approach described in Mafessoni *et al*. (2020)^91^, which involved calibrating the parameters θ and ρ with coalescent simulations (using *scrm*^95^).

We performed the following steps:

1) Generate the *scrm* simulation code from the output (.psmc) file using two scripts provided with *PSMC*: psmc2history.pl output.psmc | history2ms.pl -L [GENOME_LENGTH] -u 1.45e-08 -g 29 | sed 's/msHOT-lite/scrm/g' | sed 's/-l//g' > scrm_code.txt
2) Simulate whole genomes, each of size 2,881,033,286bp, while correcting θ and ρ by varying factors, fθ and fρ, with values from 1 to 2 in steps of 0.05. We simulated one diploid genome for each factor combination.
3) Apply the same filters used for the real data to the simulated genomes, convert the simulated genotypes into input files for *PSMC* (using again scrm2psmc.py), and apply *PSMC* on that simulated data (with the same parameters described above for the real data).
4) Run *PSMC* on the real data while setting the demographic history to that estimated using *PSMC* on the simulated data to compute the likelihood of the real data given that demographic history: psmc -N1 -i simulated_demography -o output input.psmcfa.
5) Identify the combination of correction-factors (fθ and fρ) that maximizes the likelihood (**Figure S31**), thus obtaining the corrected demographic history that best fits the real data (**Figure S32**, **Table S41**).

Note that the estimate of effective population size for *Denisova 3* for the most recent time interval in the uncalibrated demographic history was extremely high (in the order of 160 million individuals). This is likely an artifact due to the large uncertainty typically associated with the estimates for the most recent and oldest time intervals. As the Denisovan effective population size is unlikely to have increased to this extent, we set the effective population size of this time interval to the estimate of the next time interval before calibrating the demographic history.

Estimates of the past effective size (Ne) of the *Denisova 3* and *Denisova 25* populations are similar for most of their history until about 240ka when the estimates for the *Denisova 25* population decrease abruptly (**Figure S32**). This may represent the time when these populations split (see **Supplementary Note 8**). However, there is also a slight difference in Ne as early as 400ka, when the estimates of Ne for the population of *Denisova 25* get slightly smaller (as low as 2,000 individuals) than those for the population of *Denisova 3* (about 2,700 individuals throughout). This could reflect population structure between these Denisovan populations or could be due to different gene flows (e.g. from Neandertals). Differences in the estimates of Ne for Neandertals and Denisovans (**Figure S32**) are also apparent beyond the population split time of 392-438ka previously estimated for these groups^45^, as well as between modern and archaic humans. The difference between Neandertals and Densiovans is visible as early as 900ka, with a larger Ne for Neandertals than Denisovans, whereas the Ne of modern and archaic humans never really match. Gene flow from modern humans to Neandertals could potentially inflate the estimates of Ne for Neandertals, whereas the differences between modern and archaic humans could be due to selection, in particular background selection^101,102^, which may have been less efficient in the small archaic populations.

We repeated the procedure after applying different filters to evaluate the effect of Neandertal ancestry and runs of homozygosity in the Denisovan genomes, modern human ancestry in the Neandertal genomes, and background selection. Masking the runs of homozygosity in the *Denisova 25* genome had no effect on the demographic history estimated with *PSMC* (**Figure S33**). Masking Neandertal segments of at least 0.05cM identified with *admixfrog* (using a penalty of 0.4 and a window size of 0.005cM) decreased the estimates of Ne for both Denisovan populations (**Figure S34**), but the difference between ~400ka and ~250ka remained, suggesting that this is not due to Neandertal ancestry. Masking the modern human segments previously identified in two Neandertal genomes (identified in the M1A model from Hubisz *et al*. (2020)^103^) did not much affect the estimates of Ne for these Neandertals (**Figure S35**). The modern human ancestry in Neandertals may not be the cause for the early differences in Ne between Neandertals and Denisovans, but it is also likely that modern human segments were missed as the detection power of the method used to detect these segments is below 50%^103^. Therefore, it remains unclear whether these early differences in the estimates of Ne between Neandertals and Denisovans are due to the modern human ancestry of Neandertals.

Finally, to assess the effect of background selection, we focused on genomic regions presumed to evolve neutrally (B-score ≥ 0.9^104^). As the mutation rate may differ in these regions, we repeated an analysis originally done to estimate the mutation rate by comparing the reduced number of mutations in the genome of the directly radiocarbon-dated Ust’-Ishim individual with the number of mutations observed in a present-day human genome, since the common ancestor with chimpanzees. We used four high-quality present-day genomes for comparison: HGDP00982 Mbuti, HGDP00775 Han, HGDP00533 French, and HGDP00546 Papuan. At each autosomal site passing standard filters, we randomly sampled an allele from the diploid genotype calls for both the Ust’-Ishim genome (snpAD genotype calls^45^) and the comparison genome (GATK genotype calls^48^). To remain consistent with the published analysis, we excluded CpG sites and corrected mutation rate estimates accordingly^83^. The ancestral state was determined using the inferred human-chimpanzee ancestor (Ensembl Compara v71; https://ftp.ensembl.org/pub/release-75/fasta/ancestral_alleles/), excluding bases with low confidence ancestral calls (i.e. supported by a single sequence). Assuming the Ust’-Ishim individual is 45,000 years old^83^ and using a generation time of 29 years, we estimated the mutation rate as follows: $\frac{\# missing mutations}{\# sites that pass the filters}\times\frac{1}{45000}\times29$. Based on the derived alleles that are found in the lineage leading to the comparison individual, we estimated that CpG dinucleotides constitute on average 14% of analyzed mutations. Thus, we adjusted our estimates by dividing the mutation rate by (1 - 0.14) to correct for the filtering of CpG sites. We found that the mutation rate in presumed neutrally evolving regions is about 9% higher than the genome-wide mutation rate. For subsequent analyses, we used a mutation rate of 1.58 x10^-8^ mutations per base pair per generation in these regions. Applying the PSMC model and the calibration steps described above to these regions increased the estimates of Ne for all modern and archaic human populations, particularly for times beyond one million years ago (**Figure S36**, **Figure S37**). Interestingly, the estimates of effective population size beyond one million years ago are similar between modern and archaic humans when focusing on these regions (**Figure S37**), as expected given their ancestors belonged then to a common ancestral population. Therefore, the differences observed for the whole genome suggest differences in how much background selection affects the estimates of Ne for modern and archaic humans, and perhaps differences in the efficacy of selection. The larger increase in the estimates of Ne for modern humans using regions presumably evolving neutrally is consistent with a stronger effect of selection in modern humans compared to archaic humans^105^.

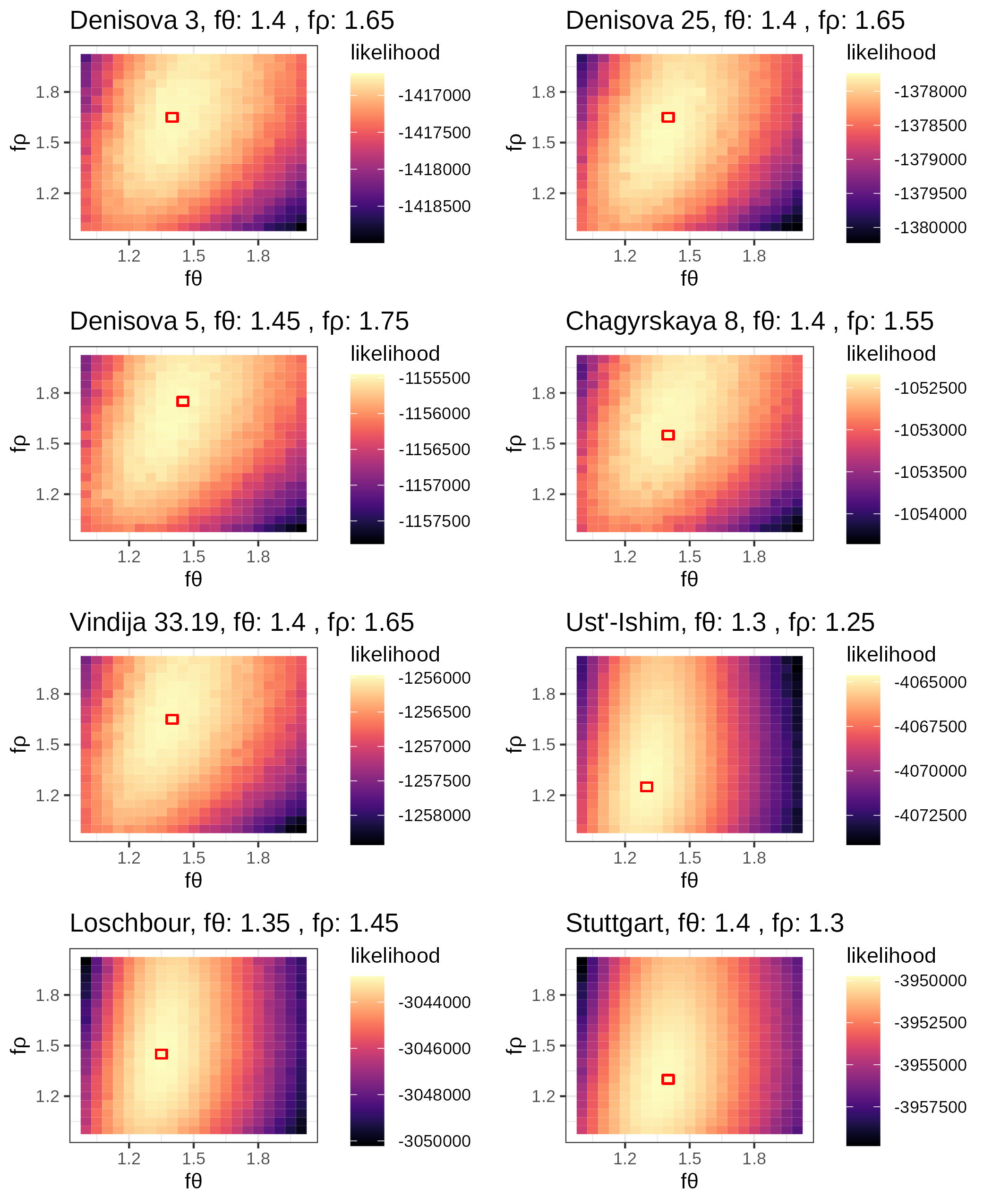

**Figure S31: Estimated corrections for the demographic histories of Denisova 25, four archaic humans, and three ancient modern humans.** The colours represent the likelihood of the real data given different correction factors of the theta (θ) and rho (ρ) parameters (fθ and fρ, respectively). The squares with red borders represent the combination of correction factors for which PSMC returns the highest likelihood (corresponding values are reported on top).

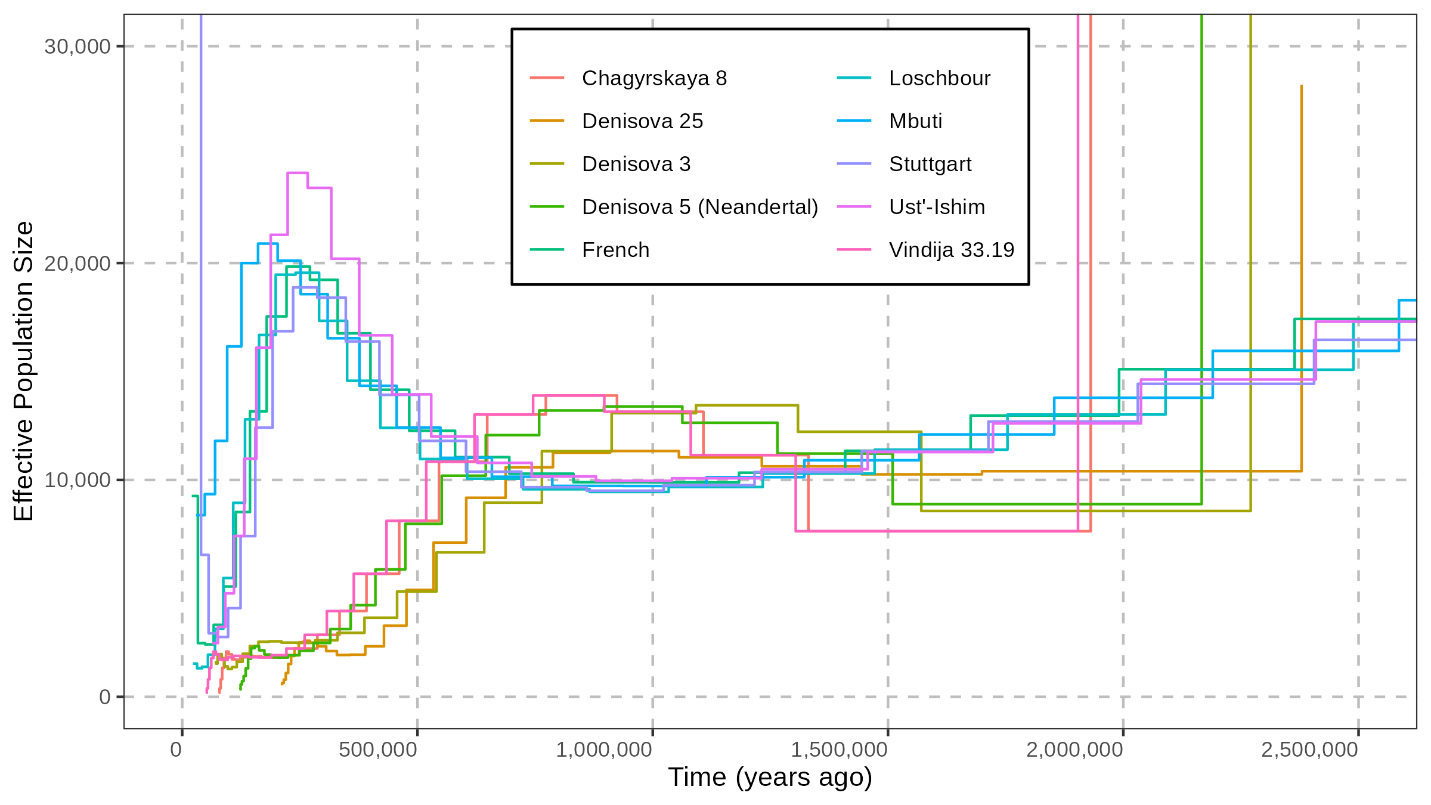

**Figure S32: Corrected demographic histories estimated with PSMC.** Time is calibrated using a mutation rate of 0.5x10^-9^ per base pair per year (corresponding to 1.45x10^-8^ per base pair per generation, for a generation time of 29 years).

**Table S41:** scrm codes to simulate a diploid genome following the corrected demographic histories (**Figure S32**).

| **Genome** | **Simulation code** |
| --- | --- |
| *Denisova 25* | scrm 2 1 -SC abs -p 10 -t 8.4663e+06 -r 1.76109e+06 2881033286 -en 0.0010 1 0.0127 -en 0.0017 1 0.0237 -en 0.0025 1 0.0226 -en 0.0035 1 0.0327 -en 0.0047 1 0.0428 -en 0.0062 1 0.0471 -en 0.0079 1 0.0529 -en 0.0099 1 0.0635 -en 0.0124 1 0.0741 -en 0.0154 1 0.0765 -en 0.0190 1 0.0699 -en 0.0233 1 0.0604 -en 0.0285 1 0.0526 -en 0.0348 1 0.0505 -en 0.0423 1 0.0587 -en 0.0513 1 0.0838 -en 0.0622 1 0.1322 -en 0.0753 1 0.2001 -en 0.0910 1 0.2627 -en 0.1099 1 0.2972 -en 0.1327 1 0.3076 -en 0.1600 1 0.3068 -en 0.1929 1 0.3012 -en 0.2325 1 0.2883 -en 0.2801 1 0.2730 -en 0.3373 1 0.2854 -en 0.4889 1 0.7113 |
| *Denisova 3* | scrm 2 1 -SC abs -p 10 -t 522943 -r 113663 2881033286 -en 0.0157 1 1.0000 -en 0.0260 1 0.6164 -en 0.0385 1 1.0993 -en 0.0537 1 1.0188 -en 0.0719 1 0.7008 -en 0.0940 1 0.5259 -en 0.1206 1 0.5275 -en 0.1528 1 0.6309 -en 0.1916 1 0.7533 -en 0.2385 1 0.8513 -en 0.2952 1 0.9506 -en 0.3637 1 1.0587 -en 0.4463 1 1.1309 -en 0.5462 1 1.1247 -en 0.6668 1 1.0705 -en 0.8125 1 1.0542 -en 0.9885 1 1.1706 -en 1.2011 1 1.5379 -en 1.4578 1 2.2723 -en 1.7679 1 3.3175 -en 2.1424 1 4.3586 -en 2.5948 1 5.0827 -en 3.1412 1 5.4878 -en 3.8012 1 5.5792 -en 4.5983 1 5.1841 -en 5.5612 1 3.7828 -en 8.1287 1 17.3710 |
| *Denisova 5* (Neandertal) | scrm 2 1 -SC abs -p 10 -t 108678 -r 25321.5 2881033286 -en 0.0631 1 0.7872 -en 0.1050 1 1.0666 -en 0.1558 1 1.5586 -en 0.2174 1 2.3170 -en 0.2920 1 3.1639 -en 0.3825 1 3.9307 -en 0.4922 1 4.4744 -en 0.6252 1 4.5890 -en 0.7864 1 4.3349 -en 0.9818 1 4.0577 -en 1.2188 1 3.8987 -en 1.5059 1 3.8966 -en 1.8541 1 4.0490 -en 2.2761 1 4.3184 -en 2.7877 1 4.7591 -en 3.4079 1 5.7627 -en 4.1597 1 8.0772 -en 5.0710 1 12.2885 -en 6.1758 1 17.7456 -en 7.5151 1 22.3110 -en 9.1386 1 24.7240 -en 11.1067 1 25.6403 -en 13.4924 1 25.7977 -en 16.3846 1 25.2012 -en 19.8906 1 23.5697 -en 24.1406 1 19.3198 -en 35.5383 1 65.3383 |
| *Chagyrskaya 8* | scrm 2 1 -SC abs -p 10 -t 833051 -r 173462 2881033286 -en 0.0068 1 0.0412 -en 0.0113 1 0.1138 -en 0.0168 1 0.2079 -en 0.0235 1 0.3076 -en 0.0316 1 0.4253 -en 0.0415 1 0.5332 -en 0.0535 1 0.5381 -en 0.0682 1 0.4930 -en 0.0860 1 0.4819 -en 0.1076 1 0.4937 -en 0.1339 1 0.4900 -en 0.1659 1 0.4774 -en 0.2048 1 0.4782 -en 0.2521 1 0.4943 -en 0.3096 1 0.5230 -en 0.3796 1 0.5778 -en 0.4647 1 0.6921 -en 0.5681 1 0.9413 -en 0.6939 1 1.4409 -en 0.8469 1 2.2048 -en 1.0330 1 2.9482 -en 1.2592 1 3.3335 -en 1.5343 1 3.3950 -en 1.8689 1 3.3206 -en 2.2757 1 3.0814 -en 2.7705 1 2.1683 -en 4.1038 1 8.3474 |
| *Vindija 33.19* | scrm 2 1 -SC abs -p 10 -t 1.78171e+07 -r 3.96304e+06 2881033286 -en 0.0004 1 0.0058 -en 0.0007 1 0.0078 -en 0.0010 1 0.0127 -en 0.0014 1 0.0174 -en 0.0019 1 0.0227 -en 0.0024 1 0.0257 -en 0.0031 1 0.0259 -en 0.0039 1 0.0246 -en 0.0049 1 0.0236 -en 0.0062 1 0.0234 -en 0.0076 1 0.0226 -en 0.0094 1 0.0214 -en 0.0115 1 0.0213 -en 0.0141 1 0.0227 -en 0.0172 1 0.0256 -en 0.0210 1 0.0301 -en 0.0255 1 0.0379 -en 0.0310 1 0.0533 -en 0.0377 1 0.0795 -en 0.0457 1 0.1118 -en 0.0554 1 0.1382 -en 0.0671 1 0.1529 -en 0.0812 1 0.1585 -en 0.0983 1 0.1561 -en 0.1189 1 0.1426 -en 0.1438 1 0.1097 -en 0.2102 1 0.4048 |
| *Ust’-Ishim* | scrm 2 1 -SC abs -p 10 -t 389608 -r 61600.1 2881033286 -en 0.1029 1 1.6713 -en 0.1686 1 1.8597 -en 0.2464 1 2.8044 -en 0.3382 1 4.9495 -en 0.4466 1 8.1397 -en 0.5747 1 10.9428 -en 0.7260 1 12.3559 -en 0.9047 1 13.0335 -en 1.1158 1 12.6460 -en 1.3651 1 10.9750 -en 1.6597 1 9.1046 -en 2.0076 1 7.6185 -en 2.4186 1 6.6143 -en 2.9040 1 6.0101 -en 3.4775 1 5.6722 -en 4.1548 1 5.5308 -en 4.9549 1 5.5755 -en 5.9000 1 5.8332 -en 7.0163 1 6.3723 -en 8.3350 1 7.3060 -en 9.8926 1 8.6948 -en 11.7325 1 10.3341 -en 13.9058 1 11.6892 -en 16.4730 1 12.2752 -en 19.5053 1 12.0909 -en 23.0873 1 11.5810 -en 32.3161 1 14.9450 |
| *Loschbour* | scrm 2 1 -SC abs -p 10 -t 1.11598e+06 -r 225929 2881033286 -en 0.0245 1 0.2395 -en 0.0404 1 0.2893 -en 0.0592 1 0.3252 -en 0.0816 1 0.4459 -en 0.1082 1 0.6964 -en 0.1398 1 1.1829 -en 0.1774 1 2.0122 -en 0.2222 1 3.0057 -en 0.2753 1 3.7760 -en 0.3385 1 4.0545 -en 0.4136 1 3.8697 -en 0.5029 1 3.4645 -en 0.6091 1 3.0140 -en 0.7353 1 2.5937 -en 0.8854 1 2.2684 -en 1.0638 1 2.0510 -en 1.2759 1 1.9290 -en 1.5280 1 1.8901 -en 1.8277 1 1.9180 -en 2.1840 1 2.0200 -en 2.6076 1 2.2486 -en 3.1112 1 2.6723 -en 3.7098 1 3.2937 -en 4.4215 1 3.9198 -en 5.2676 1 4.1980 -en 6.2735 1 3.8993 -en 8.8908 1 4.8223 |
| *Stuttgart* | scrm 2 1 -SC abs -p 10 -t 1.52402e+07 -r 2.23187e+06 2881033286 -en 0.0027 1 1.0000 -en 0.0045 1 0.1023 -en 0.0065 1 0.0426 -en 0.0090 1 0.0412 -en 0.0118 1 0.0661 -en 0.0152 1 0.1211 -en 0.0192 1 0.1970 -en 0.0239 1 0.2604 -en 0.0295 1 0.2880 -en 0.0360 1 0.2816 -en 0.0438 1 0.2541 -en 0.0529 1 0.2184 -en 0.0637 1 0.1858 -en 0.0765 1 0.1623 -en 0.0915 1 0.1483 -en 0.1093 1 0.1422 -en 0.1303 1 0.1430 -en 0.1550 1 0.1504 -en 0.1842 1 0.1647 -en 0.2187 1 0.1877 -en 0.2594 1 0.2198 -en 0.3074 1 0.2569 -en 0.3642 1 0.2888 -en 0.4311 1 0.3020 -en 0.5101 1 0.2922 -en 0.6034 1 0.2680 -en 0.8435 1 0.7945 |
| Mbuti HGDP00982 | scrm 2 1 -SC abs -p 10 -t 3.26666e+06 -r 474753 2881033286 -en 0.0172 1 0.5575 -en 0.0280 1 0.8056 -en 0.0408 1 0.9256 -en 0.0557 1 1.1389 -en 0.0733 1 1.3670 -en 0.0938 1 1.4694 -en 0.1180 1 1.4534 -en 0.1463 1 1.3397 -en 0.1795 1 1.1608 -en 0.2184 1 0.9714 -en 0.2641 1 0.8267 -en 0.3177 1 0.7323 -en 0.3806 1 0.6739 -en 0.4544 1 0.6430 -en 0.5409 1 0.6375 -en 0.6424 1 0.6570 -en 0.7615 1 0.7043 -en 0.9012 1 0.7875 -en 1.0650 1 0.9179 -en 1.2572 1 1.0974 -en 1.4827 1 1.2915 -en 1.7472 1 1.4320 -en 2.0575 1 1.4771 -en 2.4214 1 1.4476 -en 2.8483 1 1.4089 -en 3.3491 1 1.5570 -en 4.6258 1 5.2190 |
| French HGDP00533 | scrm 2 1 -SC abs -p 10 -t 6.5964e+07 -r 1.15208e+07 2881033286 -en 0.0006 1 0.0231 -en 0.0010 1 0.0082 -en 0.0014 1 0.0079 -en 0.0019 1 0.0113 -en 0.0025 1 0.0196 -en 0.0033 1 0.0337 -en 0.0041 1 0.0502 -en 0.0052 1 0.0622 -en 0.0064 1 0.0668 -en 0.0078 1 0.0637 -en 0.0095 1 0.0554 -en 0.0115 1 0.0470 -en 0.0139 1 0.0409 -en 0.0167 1 0.0368 -en 0.0200 1 0.0341 -en 0.0240 1 0.0327 -en 0.0287 1 0.0325 -en 0.0342 1 0.0338 -en 0.0407 1 0.0368 -en 0.0484 1 0.0424 -en 0.0576 1 0.0509 -en 0.0684 1 0.0614 -en 0.0812 1 0.0698 -en 0.0964 1 0.0722 -en 0.1143 1 0.0697 -en 0.1355 1 0.0674 -en 0.1903 1 0.1994 |

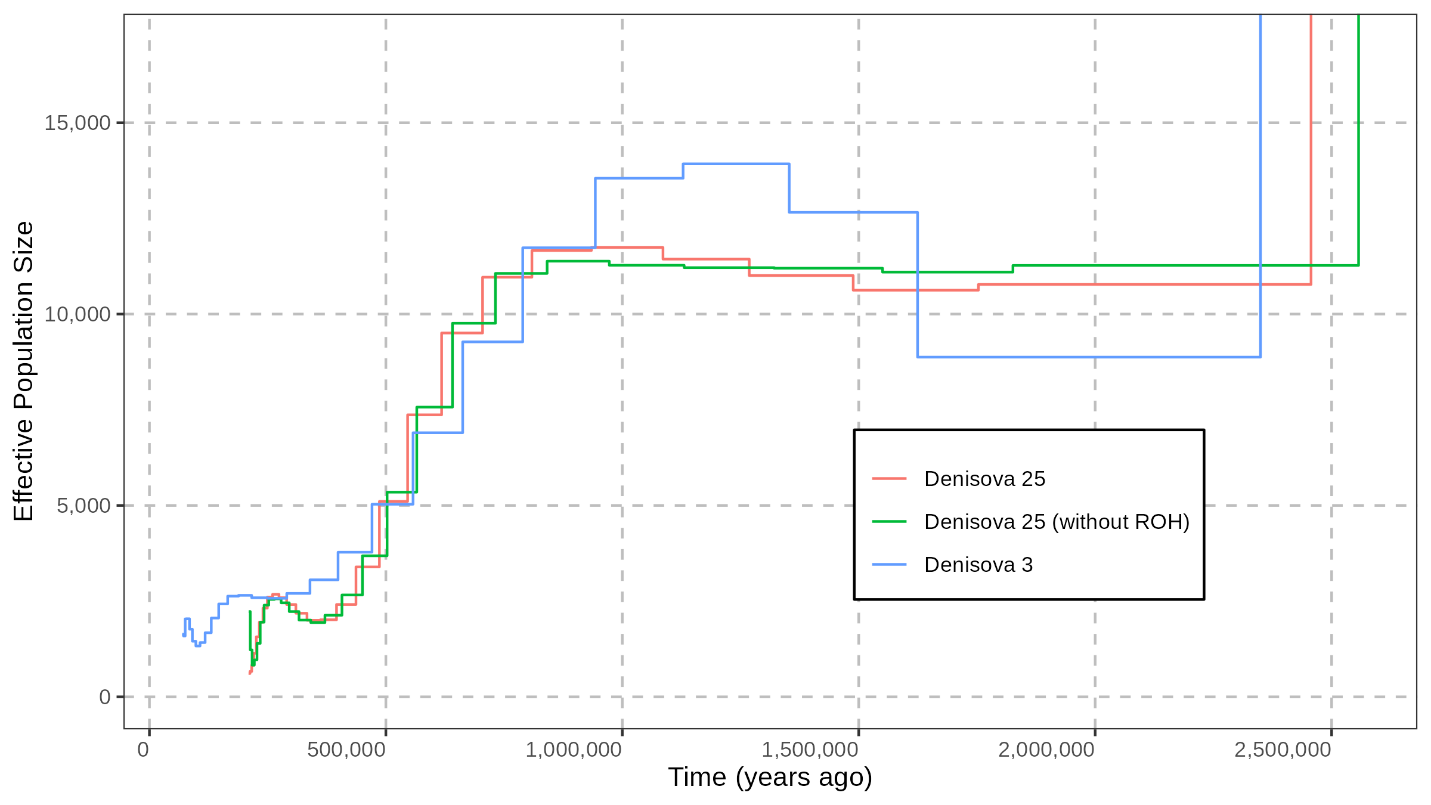

**Figure S33: No effect of HBD segments in the Denisova 25 genome on the demographic history estimated with PSMC.** We removed runs of homozygosity as described in **Supplementary Note 7** but did not apply a length cutoff. Time is calibrated using a mutation rate of 0.5x10^-9^ per base pair per year (corresponding to 1.45x10^-8^ per base pair per generation, for a generation time of 29 years).

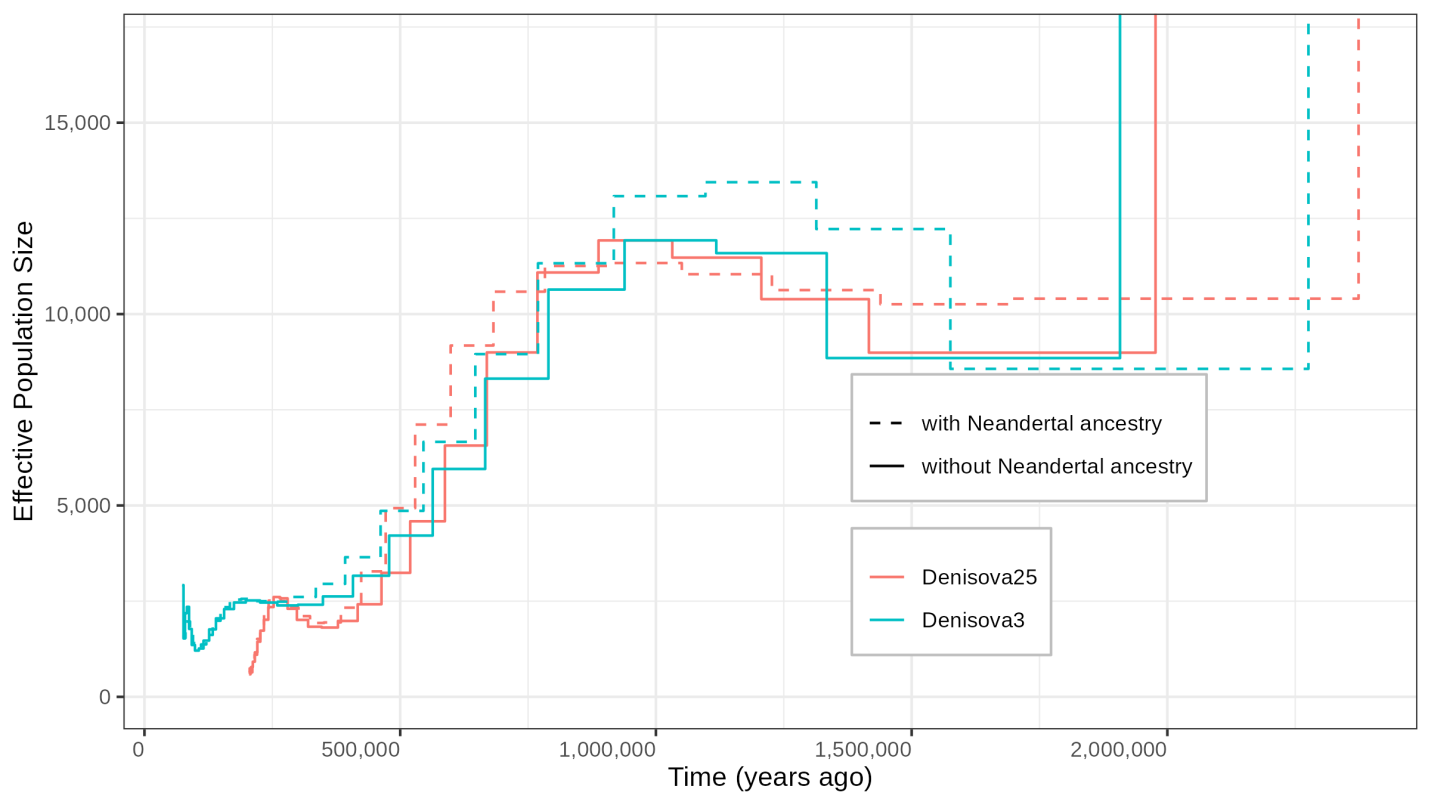

**Figure S34: Effect of masking the Neandertal ancestry on the Denisovan** **demographic histories estimated with PSMC**. Only the Neandertal segments of at least 0.05cM (as identified with admixfrog with a penalty of 0.4 and a window size of 0.005cM) were masked in the corresponding Denisovan genome. Time is calibrated using a mutation rate of 0.5x10^-9^ per base pair per year (corresponding to 1.45x10^-8^ per base pair per generation, for a generation time of 29 years).

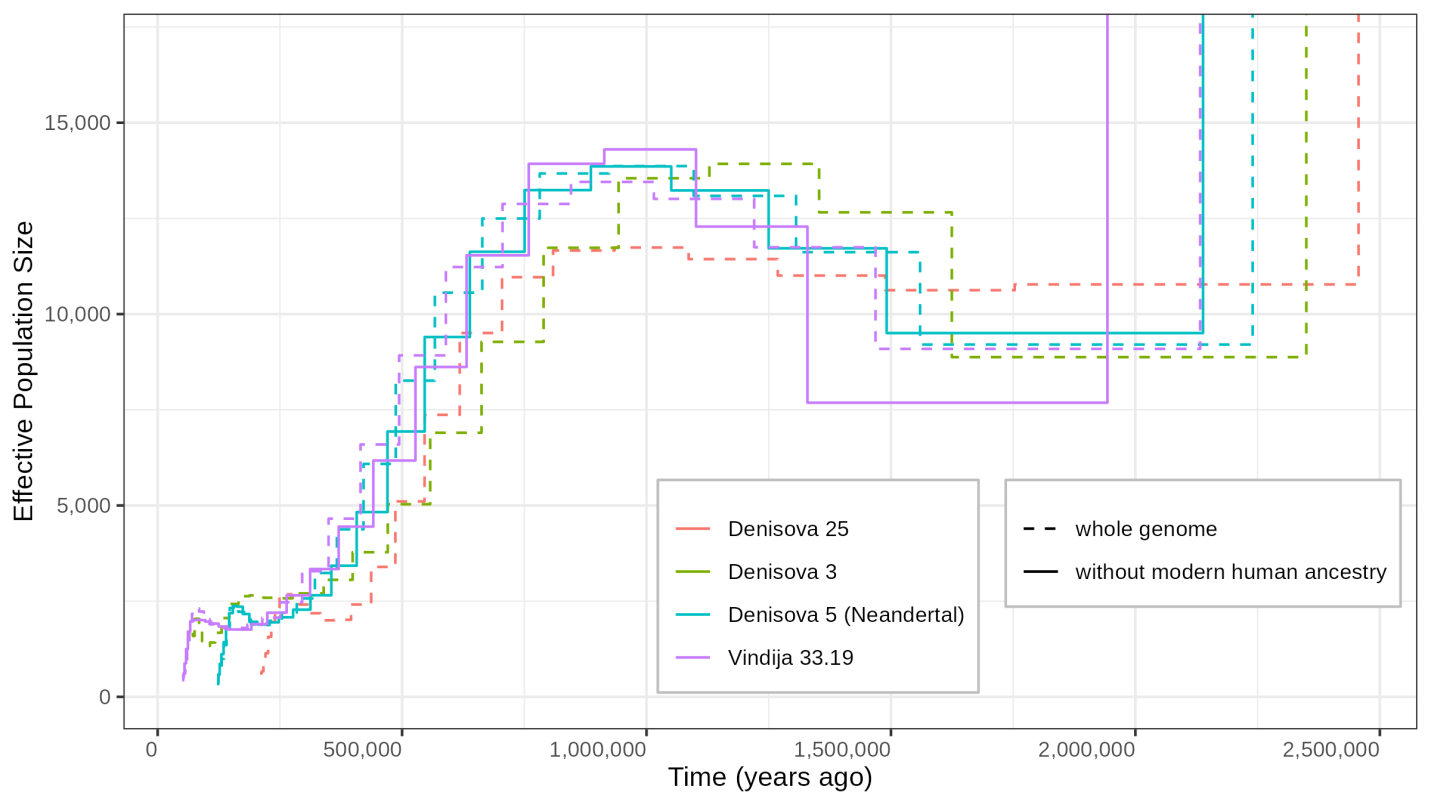

**Figure S35: Effect of masking modern human ancestry on the Neandertal** **demographic histories estimated with PSMC**. Modern human segments identified in the M1A model from Hubisz et al. (2020)^103^ were masked in the corresponding Neandertal genome. For comparison, the calibrated demographic histories of the two Denisovans (inferred from the whole genome) are also plotted. Time is calibrated using a mutation rate of 0.5x10^-9^ per base pair per year (corresponding to 1.45x10^-8^ per base pair per generation, for a generation time of 29 years).

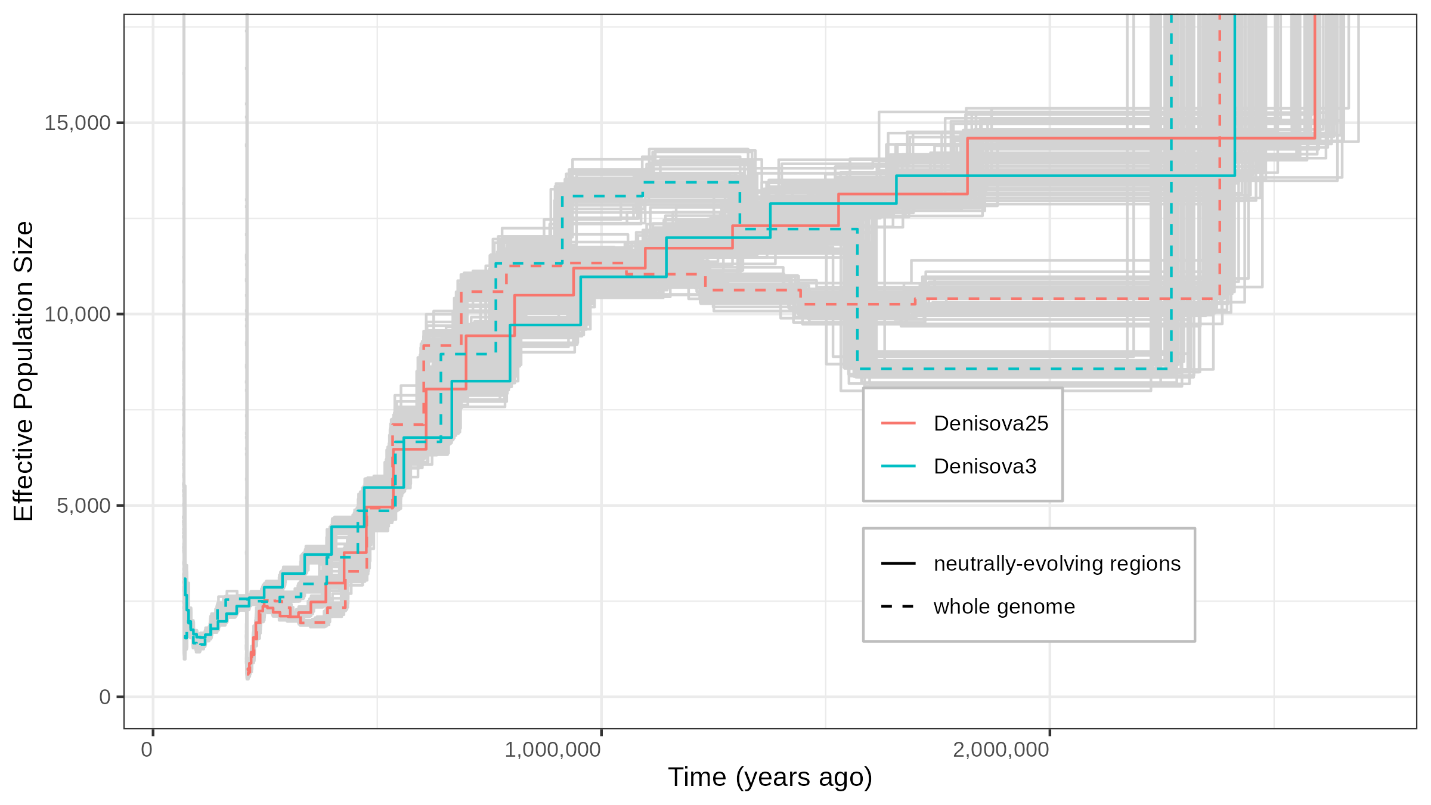

**Figure S36: Effect of background selection on the Denisovan** **demographic histories estimated with PSMC**. Regions with more than 10% of their genetic diversity lost presumably due to background selection (B-score lower than 0.9^104^) were filtered out to retain “neutrally-evolving regions” in the Denisovan genomes. Time is calibrated using a mutation rate of 0.5x10^-9^ per base pair per year (corresponding to 1.45x10^-8^ per base pair per generation, for a generation time of 29 years). Grey curves represent uncertainties in the estimates and were obtained with 50 rounds of bootstrapping for each condition.

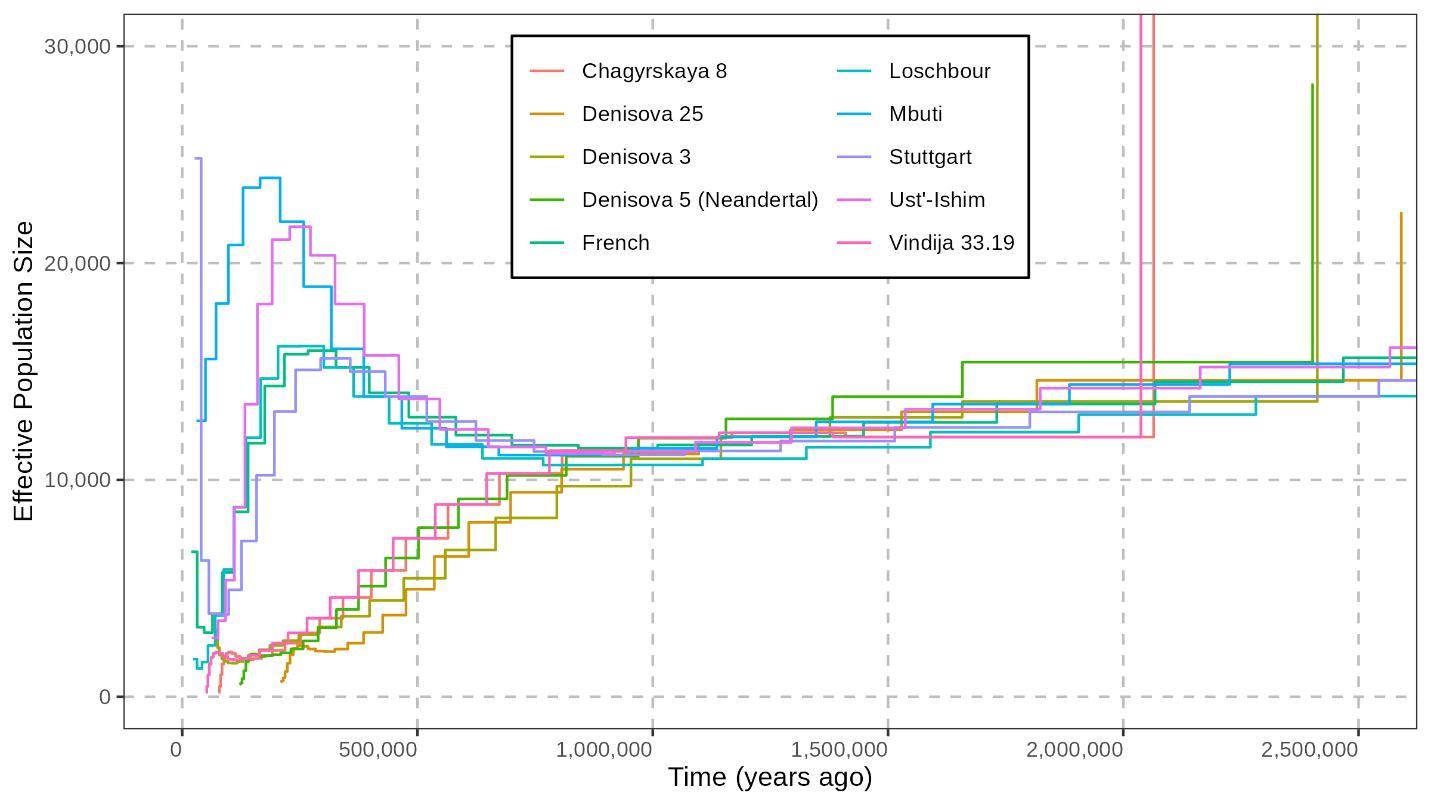

**Figure S37: Corrected demographic histories estimated with PSMC in regions of the genome that presumably evolve neutrally.** See **Figure S36** for more details.

**Table S42**: scrm codes to simulate a diploid genome following the corrected demographic histories inferred from regions in the genome that presumably evolve neutrally (**Figure S36**).

| **Genome** | **Simulation code** |
| --- | --- |
| *Denisova 25* | scrm 2 1 -SC abs -t 273146.285073835 -r 61051.8763405278 2881033286 -en 0.0227 1 0.4724 -en 0.0379 1 0.4905 -en 0.0564 1 0.5858 -en 0.0788 1 0.7782 -en 0.1061 1 1.0328 -en 0.1392 1 1.2909 -en 0.1795 1 1.4943 -en 0.2285 1 1.5778 -en 0.2881 1 1.5481 -en 0.3606 1 1.4722 -en 0.4487 1 1.4066 -en 0.5558 1 1.3939 -en 0.6861 1 1.4677 -en 0.8444 1 1.6530 -en 1.0370 1 1.9821 -en 1.2711 1 2.5136 -en 1.5557 1 3.3069 -en 1.9018 1 4.3130 -en 2.3225 1 5.3622 -en 2.8341 1 6.2904 -en 3.4562 1 6.9973 -en 4.2124 1 7.4687 -en 5.1320 1 7.8163 -en 6.2500 1 8.2078 -en 7.6094 1 8.7605 -en 9.2622 1 9.7322 -en 13.7151 1 14.9112 |
| *Denisova 3* | scrm 2 1 -SC abs -t 1160133.15532523 -r 239253.606519953 2881033286 -en 0.0056 1 0.4845 -en 0.0093 1 0.4172 -en 0.0138 1 0.3558 -en 0.0192 1 0.3023 -en 0.0258 1 0.2751 -en 0.0338 1 0.2570 -en 0.0435 1 0.2450 -en 0.0553 1 0.2430 -en 0.0696 1 0.2550 -en 0.0869 1 0.2791 -en 0.1080 1 0.3095 -en 0.1335 1 0.3410 -en 0.1644 1 0.3721 -en 0.2019 1 0.4064 -en 0.2474 1 0.4494 -en 0.3026 1 0.5058 -en 0.3696 1 0.5838 -en 0.4508 1 0.6976 -en 0.5493 1 0.8586 -en 0.6688 1 1.0635 -en 0.8137 1 1.2950 -en 0.9895 1 1.5253 -en 1.2028 1 1.7227 -en 1.4614 1 1.8835 -en 1.7751 1 2.0232 -en 2.1556 1 2.1377 -en 3.1770 1 6.3084 |
| *Denisova 5* (Neandertal) | scrm 2 1 -SC abs -t 108666.337066188 -r 24510.2345545199 2881033286 -en 0.0537 1 0.9797 -en 0.0896 1 1.0637 -en 0.1332 1 1.4029 -en 0.1863 1 2.0212 -en 0.2509 1 2.7190 -en 0.3296 1 3.2090 -en 0.4253 1 3.3016 -en 0.5417 1 3.1922 -en 0.6834 1 3.1350 -en 0.8558 1 3.1779 -en 1.0657 1 3.2603 -en 1.3210 1 3.3976 -en 1.6317 1 3.7055 -en 2.0098 1 4.3220 -en 2.4699 1 5.3324 -en 3.0298 1 6.7466 -en 3.7111 1 8.5519 -en 4.5401 1 10.7248 -en 5.5490 1 13.0820 -en 6.7766 1 15.2942 -en 8.2705 1 17.1076 -en 10.0884 1 18.5903 -en 12.3005 1 19.9896 -en 14.9924 1 21.4762 -en 18.2681 1 23.1930 -en 22.2543 1 25.8701 -en 33.0074 1 47.4039 |
| *Chagyrskaya 8* | scrm 2 1 -SC abs -t 108926.404911748 -r 23732.7839088342 2881033286 -en 0.0442 1 0.3390 -en 0.0739 1 0.8035 -en 0.1100 1 1.6987 -en 0.1542 1 2.5236 -en 0.2080 1 3.0490 -en 0.2736 1 3.3584 -en 0.3536 1 3.4604 -en 0.4512 1 3.3522 -en 0.5703 1 3.1157 -en 0.7156 1 2.9009 -en 0.8927 1 2.8423 -en 1.1088 1 2.9533 -en 1.3723 1 3.1992 -en 1.6938 1 3.5836 -en 2.0859 1 4.1418 -en 2.5641 1 4.9307 -en 3.1474 1 6.0631 -en 3.8588 1 7.6603 -en 4.7266 1 9.7521 -en 5.7851 1 12.2244 -en 7.0761 1 14.8312 -en 8.6507 1 17.2183 -en 10.5713 1 18.9842 -en 12.9140 1 19.9876 -en 15.7713 1 20.3569 -en 19.2564 1 20.0251 -en 28.6922 1 56.1435 |
| *Vindija 33.19* | scrm 2 1 -SC abs -t 1258650.8628694 -r 283756.693032278 2881033286 -en 0.0041 1 0.0807 -en 0.0068 1 0.0826 -en 0.0102 1 0.1276 -en 0.0142 1 0.1821 -en 0.0191 1 0.2297 -en 0.0251 1 0.2589 -en 0.0324 1 0.2663 -en 0.0412 1 0.2611 -en 0.0520 1 0.2517 -en 0.0650 1 0.2423 -en 0.0809 1 0.2353 -en 0.1003 1 0.2353 -en 0.1238 1 0.2490 -en 0.1524 1 0.2810 -en 0.1871 1 0.3312 -en 0.2294 1 0.4012 -en 0.2808 1 0.4981 -en 0.3433 1 0.6316 -en 0.4194 1 0.8088 -en 0.5118 1 1.0267 -en 0.6243 1 1.2588 -en 0.7610 1 1.4597 -en 0.9272 1 1.5950 -en 1.1294 1 1.6584 -en 1.3753 1 1.6819 -en 1.6743 1 1.7356 -en 2.4800 1 4.3852 |
| *Ust’-Ishim* | scrm 2 1 -SC abs -t 380965.456259404 -r 68533.7947164062 2881033286 -en 0.0899 1 1.1275 -en 0.1477 1 2.0471 -en 0.2160 1 3.0735 -en 0.2970 1 4.7213 -en 0.3929 1 6.8175 -en 0.5065 1 8.7362 -en 0.6411 1 9.8682 -en 0.8004 1 9.9561 -en 0.9891 1 9.2917 -en 1.2125 1 8.2545 -en 1.4772 1 7.2054 -en 1.7907 1 6.3553 -en 2.1619 1 5.7817 -en 2.6015 1 5.4858 -en 3.1222 1 5.4155 -en 3.7389 1 5.5002 -en 4.4692 1 5.6994 -en 5.3341 1 6.0066 -en 6.3585 1 6.4059 -en 7.5717 1 6.8588 -en 9.0084 1 7.3255 -en 10.7101 1 7.7870 -en 12.7253 1 8.2387 -en 15.1120 1 8.6682 -en 17.9387 1 9.0557 -en 21.2863 1 9.5028 -en 29.9465 1 11.6002 |
| *Loschbour* | scrm 2 1 -SC abs -t 1014710.1224482 -r 201572.258762274 2881033286 -en 0.0219 1 0.2755 -en 0.0361 1 0.2288 -en 0.0530 1 0.3007 -en 0.0733 1 0.4359 -en 0.0975 1 0.6785 -en 0.1264 1 1.0687 -en 0.1608 1 1.5926 -en 0.2020 1 2.1679 -en 0.2511 1 2.6431 -en 0.3098 1 2.9010 -en 0.3798 1 2.9161 -en 0.4635 1 2.7593 -en 0.5633 1 2.5375 -en 0.6826 1 2.3195 -en 0.8249 1 2.1367 -en 0.9948 1 2.0034 -en 1.1978 1 1.9290 -en 1.4400 1 1.9204 -en 1.7293 1 1.9754 -en 2.0746 1 2.0830 -en 2.4869 1 2.2302 -en 2.9792 1 2.4055 -en 3.5669 1 2.5939 -en 4.2686 1 2.7781 -en 5.1065 1 2.9518 -en 6.1068 1 3.1671 -en 8.7270 1 3.7166 |
| *Stuttgart* | scrm 2 1 -SC abs -t 609676420.859231 -r 116949187.179559 2881033286 -en 0.0001 1 0.0074 -en 0.0001 1 0.0019 -en 0.0001 1 0.0011 -en 0.0002 1 0.0011 -en 0.0002 1 0.0015 -en 0.0003 1 0.0021 -en 0.0004 1 0.0031 -en 0.0005 1 0.0039 -en 0.0006 1 0.0045 -en 0.0007 1 0.0047 -en 0.0009 1 0.0045 -en 0.0011 1 0.0041 -en 0.0013 1 0.0038 -en 0.0016 1 0.0035 -en 0.0019 1 0.0034 -en 0.0023 1 0.0033 -en 0.0027 1 0.0034 -en 0.0033 1 0.0035 -en 0.0039 1 0.0037 -en 0.0046 1 0.0039 -en 0.0055 1 0.0041 -en 0.0065 1 0.0044 -en 0.0078 1 0.0046 -en 0.0092 1 0.0048 -en 0.0109 1 0.0051 -en 0.0130 1 0.0059 -en 0.0182 1 0.0117 |
| Mbuti HGDP00982 | scrm 2 1 -SC abs -t 2552708.48332439 -r 519201.481524362 2881033286 -en 0.0186 1 0.9079 -en 0.0304 1 1.1113 -en 0.0442 1 1.2942 -en 0.0605 1 1.4863 -en 0.0795 1 1.6751 -en 0.1019 1 1.7072 -en 0.1281 1 1.5632 -en 0.1589 1 1.3494 -en 0.1950 1 1.1450 -en 0.2374 1 0.9877 -en 0.2871 1 0.8835 -en 0.3454 1 0.8227 -en 0.4139 1 0.7953 -en 0.4942 1 0.7952 -en 0.5884 1 0.8169 -en 0.6990 1 0.8548 -en 0.8288 1 0.9045 -en 0.9811 1 0.9629 -en 1.1597 1 1.0278 -en 1.3694 1 1.0958 -en 1.6154 1 1.1615 -en 1.9040 1 1.2200 -en 2.2427 1 1.2712 -en 2.6401 1 1.3213 -en 3.1065 1 1.3828 -en 3.6536 1 1.5219 -en 5.0491 1 2.0011 |
| French HGDP00533 | scrm 2 1 -SC abs -t 23081604.1608257 -r 5150311.81367475 2881033286 -en 0.0013 1 0.0528 -en 0.0022 1 0.0253 -en 0.0032 1 0.0234 -en 0.0044 1 0.0303 -en 0.0058 1 0.0453 -en 0.0075 1 0.0673 -en 0.0095 1 0.0923 -en 0.0119 1 0.1131 -en 0.0148 1 0.1246 -en 0.0182 1 0.1260 -en 0.0222 1 0.1198 -en 0.0270 1 0.1106 -en 0.0328 1 0.1017 -en 0.0396 1 0.0952 -en 0.0477 1 0.0915 -en 0.0573 1 0.0905 -en 0.0687 1 0.0917 -en 0.0823 1 0.0948 -en 0.0985 1 0.0998 -en 0.1178 1 0.1066 -en 0.1406 1 0.1146 -en 0.1679 1 0.1234 -en 0.2002 1 0.1322 -en 0.2387 1 0.1416 -en 0.2845 1 0.1536 -en 0.3390 1 0.1817 -en 0.4808 1 0.3262 |

### Supplementary Note 8: Relationship to other hominins

#### *D*- and *f4*-statistics

To explore the relationships of *Denisova 25* to archaic and modern humans, we computed *D*- and *f4*-statistics comparing *Denisova 25* to the published high-coverage archaic genomes (*Denisova 3*, *Denisova 5*, *Chagyrskaya 8*, *Vindija 33.19*), to the low-coverage Denisovan genomes (*Denisova 2*^61^, *Denisova 4*^8^, *Denisova 8*^8^, *Denisova 11*^96^) and to 276 modern human genomes from the Simons Genome Diversity Panel (SGDP)^106^. We used the genotype calls as described in **Supplementary Note 3** for the high-coverage archaic genomes, randomly sampled one allele at each covered position in the low-coverage Denisovan genomes (reprocessed as described in the next section), and downloaded the SGDP genomes on March 9^th^ 2023 from the European Nucleotide Archive in the Variant Call Format (VCF) that includes invariable sites. Each genome was filtered independently before merging into a VCF file including the high-coverage archaic and modern human genomes and another VCF file including all archaic genomes. We used the standard set of filters as described in **Supplementary Note 3** for *Denisova 25*, or in Prüfer *et al*. (2017)^45^ and Mafessoni *et al.* (2020)^91^ for the other high-coverage archaic genomes and applied similar filters for the SGDP genomes: the Map35_100% mappability track from Prüfer *et al*. (2014)^48^ (for consistency with the archaic genomes), GC-corrected coverage cutoffs (computed for each genome) removing the top 2.5% and bottom 2.5% of the coverage distribution per GC category (as described in **Supplementary Note 3**), a minimum coverage of 10x, and a mask for simple repeats as identified by TandemRepeatFinder^55^. We only applied the mappability filter for the low-coverage Denisovan genomes. All genomes were then merged with bcftools merge (version 1.4^107^), setting unknown genotypes as missing, and removing both invariable sites and those with more than two alleles. Next, we determined the ancestral allele at these positions as that shared by at least three great ape reference genomes (chimpanzee panTro4, bonobo panpan1.1, gorilla gorGor3, and orangutan ponAbe2), allowing missing data or another allele in one genome as long as this allele was not present in archaic and modern humans (the ancestral allele was otherwise set as missing). Finally, we converted the VCF files into the bed/bim/fam format of Plink (with plink --vcf VCF --make-bed --double-id --set-missing-var-ids @_# --keep-allele-order --vcf-half-call missing) to convert it into the EIGENSTRAT format with *convertf* (version from EIG6.0.1).

*D*- and *f4*-statistics were computed with ADMIXTOOLS^108^ as implemented in *admixr* (version 0.9.1^109^). Positive values indicate an excess of allele sharing between the first and the third population, whereas negative values indicate an excess of allele sharing between the second and the third population. Standard errors were calculated using a weighted block jackknife with a block size of 5Mb (default value).

##### Relationship with other Denisovans

To assess the relationship between *Denisova 25* and other published Denisovan genomes, we reprocessed the raw sequencing data for *Denisova 2*, *Denisova 4*, and *Denisova 8*, ensuring consistency in data processing, in particular by applying a consistent minimum mapping quality cutoff of 25. Demultiplexing for *Denisova 4* and *Denisova 8* was performed allowing one mismatch in the expected library indexes, except for a subset of sequences from two HiSeq lanes, for which perfect index matching of P7 was required, while P5 was disregarded due to sequencing issues. *Denisova 2* sequences were used as demultiplexed in the original study^61^. All sequences were then mapped to the human reference genome (version hg19 with additional decoy sequences, as described in Prüfer *et al*. (2017)^45^ using *BWA*^46^ with parameters adapted to ancient DNA^47^. For each library, PCR duplicates, identified as sequences with identical start and end coordinates, were collapsed using bam-rmdup (<https://github.com/mpieva/biohazard-tools>). Sequences shorter than 35bp and those with mapping quality below 25 were removed, as well as those mapping outside the Map35_100% mappability track^48^. We generated VCF files for *Denisova 2*, *Denisova 4*, *Denisova 8*, and *Denisova 11* (using published sequences for *Denisova 11*, as their processing matched our current filtering^96^) by randomly sampling one allele per covered position using bam-caller (<https://github.com/bodkan/bam-caller>), and then merged these VCF files with those of the high-coverage archaic human genomes as described earlier. Given the high levels of present-day human DNA contamination in some of the low-coverage genomes (60% in *Denisova 2*, 66% in *Denisova 4*, 15% in *Denisova 8*, and <1.7% in *Denisova 11*), we generated an additional set of VCF files by sampling only sequences showing at least one C-to-T substitution within the last three positions at either end for all four low-coverage Denisovan genomes.

We then computed *D*-statistics of the form *D*(*Denisova 3*, *Denisova 25*, other Denisovan, ancestral) to compare the relationship of *Denisova 3* and *Denisova 25* with other Denisovan individuals. We found that *Denisova 4* and *Denisova 11* are more closely related to *Denisova* 3 than to *Denisova 25*, while *Denisova 8* is more closely related to *Denisova 25* than to *Denisova 3*. These results are consistent with the mitochondrial relationships and the age of the individuals, suggesting at least two temporally and genetically distinct Denisovan groups. *Denisova 2* appears equally related to both *Denisova 3* and *Denisova 25*, possibly reflecting an earlier Denisovan population, though the substantial present-day human DNA contamination (60%) in *Denisova 2*, even among sequences exhibiting signs of ancient DNA damage, may obscure the relationship to *Denisova 3* and *Denisova 25*.

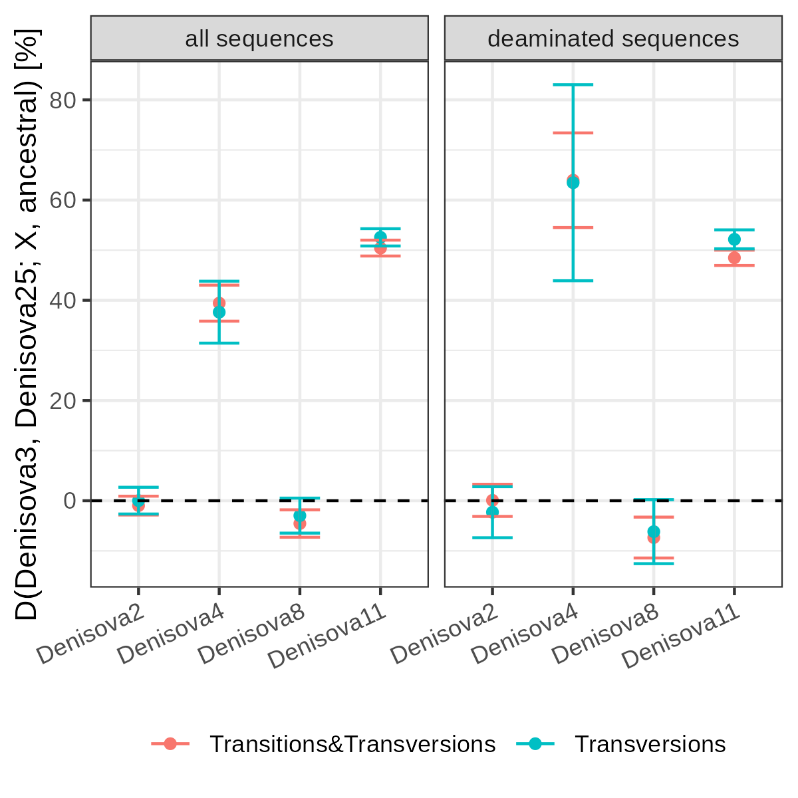

**Figure S38:** **D-statistics comparing derived allele sharing among Denisovans.** D-statistics were calculated using ADMIXTOOLS^108^, as implemented in admixr ^109^, using a random allele at each position of the low coverage Denisovan genomes (X-axis) from either all sequences or those with at least one C-to-T substitution within the last three positions of either end (deaminated sequences). The genomes of panTro4, panPan2, gorGor3, and ponAbe2 were used to determine the ancestral allele. Whiskers on each side of the plotted D values correspond to two standard errors calculated using a Weighted Block Jackknife and a block size of 5Mb across all autosomes. Colours indicate whether the D-statistics were calculated using all polymorphisms (Transitions & Transversions) or by restricting to transversion polymorphisms (Transversions).

##### Relationship with Neandertals

We first computed *D*(*Denisova 3*, *Denisova 25*, Neandertal, ancestral) to compare how both Denisovans are related to Neandertals. We found that *Denisova 25* shares significantly more derived alleles with Neandertals than *Denisova 3* does (**Figure S39**, ***Table S43***), although this is not significant for the *Denisova 5* Neandertal. This suggests that *Denisova 25* carries more Neandertal ancestry than *Denisova 3* (which is consistent with results from **Supplementary Note 9**). Gene flow from Neandertals related to *Denisova 5* into the ancestors of *Denisova 3*^45,48^ may explain why *D*(*Denisova 3*, *Denisova 25*, *Denisova 5*, ancestral) is not significantly different from 0. We next computed statistics of the form *D*(Neandertal #1, Neandertal #2, *Denisova 3* or *Denisova 25*, ancestral) to compare how each Denisovan relates to different Neandertals. *Denisova 3* and *Denisova 25* are equally related to the *Chagyrskaya 8* and *Vindija 33.19* Neandertals (**Figure S39**, **Table S43**), but both Denisovans are significantly more closely related to the *Denisova 5* Neandertal than to other Neandertals (although this is not significant for *Denisova 25* when restricting to transversion polymorphisms). While this is consistent with gene flow from Neandertals related to *Denisova 5* into the ancestors of *Denisova 3*^45,48^, *Denisova 25* predates the split of these Neandertal populations so that the most parsimonious explanation for *Denisova 25* also sharing significantly more derived alleles with *Denisova 5* than other Neandertals is the presence of more Denisovan ancestry in the *Denisova 5* Neandertal genome than in the other two Neandertal genomes.

**Table S43: D-statistics comparing derived allele sharing between Neandertals and Denisovans.** D-statistics were calculated using ADMIXTOOLS^108^ as implemented in admixr^109^. The genomes of panTro4, panPan2, gorGor3, and ponAbe2 were used to determine the ancestral allele. These results are depicted in **Figure S39**.

| **D-statistic** | **polymorphisms** | ***D* [%]** | **Z** | **BABA** | **ABBA** |
| --- | --- | --- | --- | --- | --- |
| D(*Denisova 3*, *Denisova 25*, *Vindija 33.19*, ancestral) | all | -4.8 | -2.7 | 36,264 | 39,930 |
|  | transversions | -4.5 | -2.4 | 11,920 | 13,028 |
| D(*Denisova 3*, *Denisova 25*, *Chagyrskaya 8*, ancestral) | all | -4.4 | -2.5 | 35,944 | 39,206 |
|  | transversions | -3.9 | -2.1 | 11,830 | 12,780 |
| D(*Denisova 3*, *Denisova 25*, *Denisova 5* (Neandertal), ancestral) | all | -3.3 | -1.8 | 37,075 | 39,598 |
|  | transversions | -2.6 | -1.3 | 12,196 | 12,850 |
| D(*Chagyrskaya 8*, *Vindija 33.19*, *Denisova 3*, ancestral) | all | 0.4 | 0.3 | 21,602 | 21,447 |
|  | transversions | 1.0 | 0.6 | 7,024 | 6,890 |
| D(*Chagyrskaya 8*, *Vindija 33.19*, *Denisova 25*, ancestral) | all | -0.4 | -0.3 | 20,420 | 20,566 |
|  | transversions | 0.2 | 0.2 | 6,663 | 6,631 |
| D(*Denisova 5* (Neandertal), *Vindija 33.19*, *Denisova 3*, ancestral) | all | 5.1 | 3.8 | 31,278 | 28,268 |
|  | transversions | 5.0 | 3.5 | 10,113 | 9,147 |
| D(*Denisova 5* (Neandertal), *Vindija 33.19*, *Denisova 25*, ancestral) | all | 3.1 | 2.4 | 29,220 | 27,441 |
|  | transversions | 2.5 | 1.7 | 9,413 | 8,959 |
| D(*Denisova 5* (Neandertal), *Chagyrskaya 8*, *Denisova 3*, ancestral) | all | 4.8 | 3.4 | 30,888 | 28,094 |
|  | transversions | 4.3 | 2.8 | 10,012 | 9,191 |
| D(*Denisova 5* (Neandertal), *Chagyrskaya 8*, *Denisova 25*, ancestral) | all | 3.4 | 2.6 | 28,789 | 26,889 |
|  | transversions | 2.4 | 1.7 | 9,327 | 8,883 |

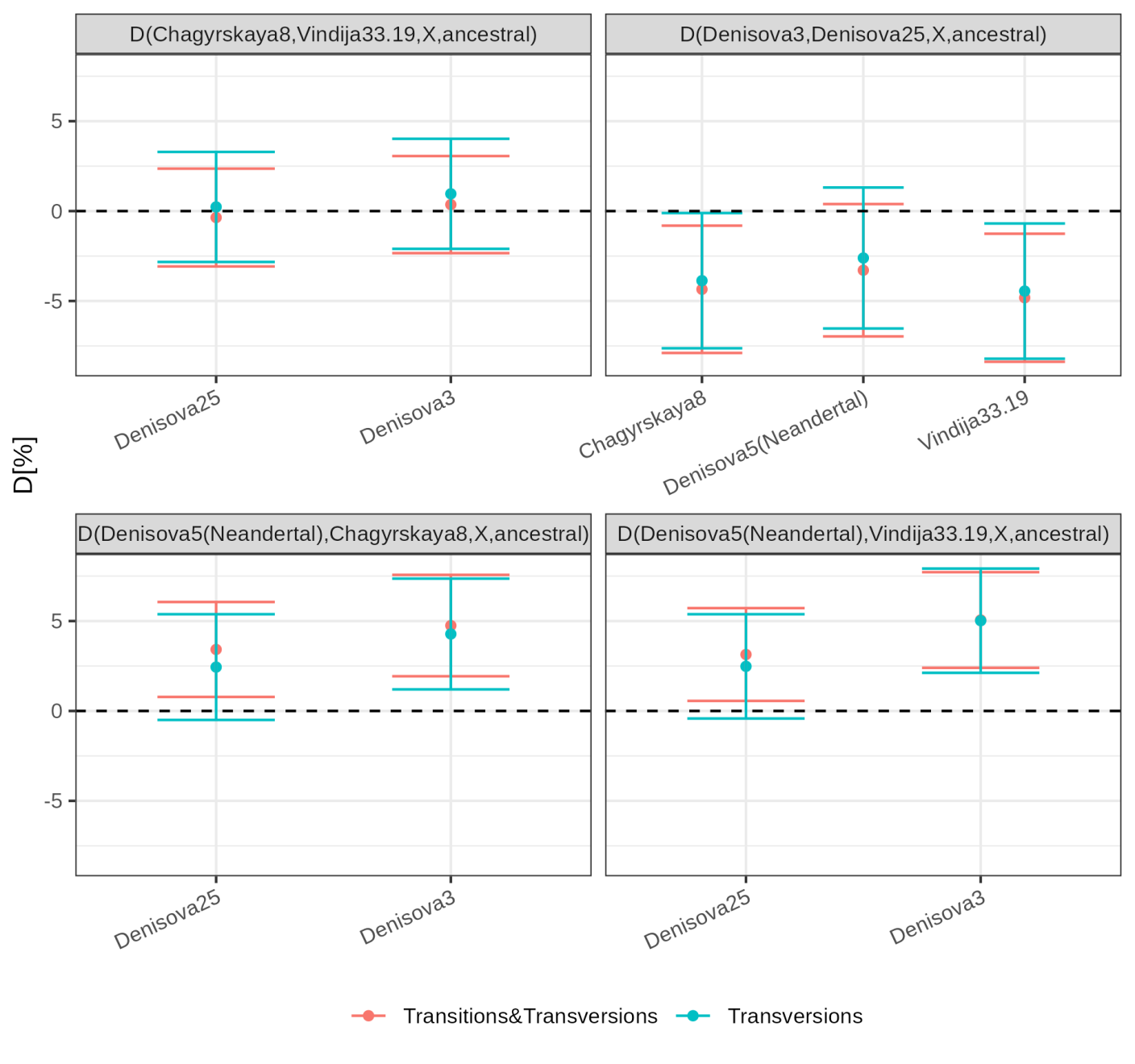

**Figure S39:** **D-statistics comparing derived allele sharing between Neandertals and Denisovans.** D-statistics were calculated using ADMIXTOOLS^108^ as implemented in admixr^109^. The genomes of panTro4, panPan2, gorGor3, and ponAbe2 were used to determine the ancestral allele. Whiskers on each side of the plotted D values correspond to two standard errors calculated using a Weighted Block Jackknife and a block size of 5Mb across all autosomes. Colours indicate whether the computation of the D-statistics were restricted to transversion polymorphisms (Transversions) or all polymorphisms were used (Transitions & Transversions).

##### Relationship with modern humans

We compared how *Denisova 3* and *Denisova 25* are related to present-day human populations using *f4*(*Denisova 3*, *Denisova 25*; present-day human population, ancestral). *Denisova 3* and *Denisova 25* appear equally related to all present-day humans as none of the comparisons are significantly different from 0. However, these genome-wide comparisons do not have the power to detect how the Denisovan ancestry carried by some of these populations is related to *Denisova 3* and *Denisova 25*. We explore and describe this further in **Supplementary Note 11**.

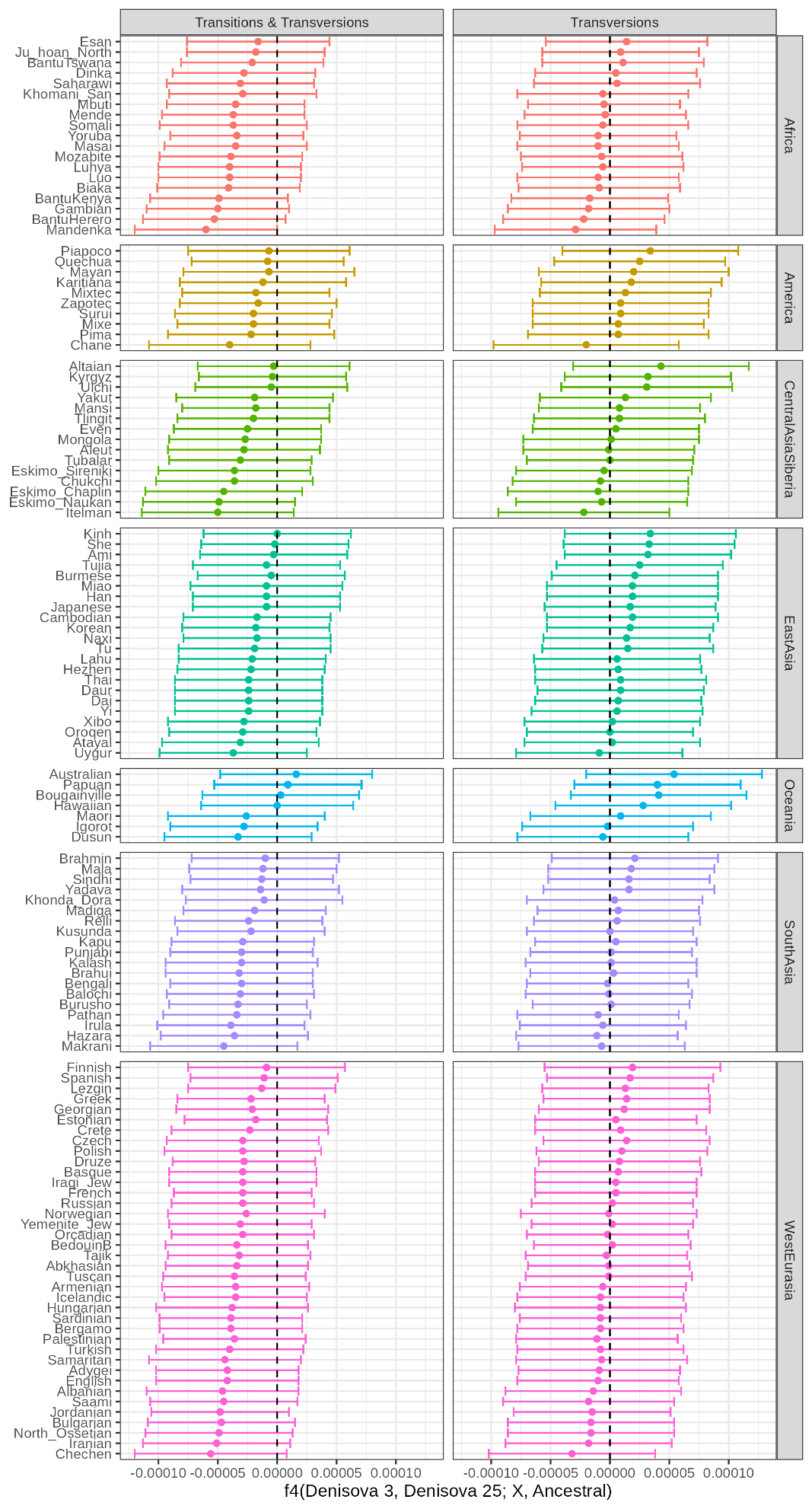

**Figure S40: f4-statistics comparing derived allele sharing between modern humans and two Denisovans.** f4-statistics were calculated using ADMIXTOOLS^108^ as implemented in admixr^109^. Whiskers on each side of the plotted values correspond to two standard errors calculated using a Weighted Block Jackknife and a block size of 5Mb across all autosomes. Colours correspond to different geographical regions.

#### Population split times

To estimate population split times among archaic humans and between archaic and modern humans, previous studies have used the *F(A|B)* statistic^45,48,91^. This statistic corresponds to the proportion of derived alleles carried by a pseudo-haploid genome A at sites where a diploid genome B is heterozygous. When A and B derive from different populations, *F(A|B)* decreases with increasing time between the sampling of B and the split between the two populations. This is because mutations private to B accumulate and are not shared with A. Using the population size history of B inferred with PSMC, simulations can be used to estimate the expected value of *F(A|B)* as a function of split time, enabling calibration of the observed value to infer the population split time.

Here, we assess how gene flow and selection influence the estimates obtained with this approach. We also compare *F(A|B)* to an alternative statistic, divergence, defined here as the number of derived alleles private to a pseudo-haploid version of B divided by the number of positions compared between A and B. Finally, we apply both statistics to estimate split times among archaic humans and between archaic and modern humans.

##### Effect of gene flow on *F(A|B)* and divergence in simulations

Previous studies of archaic genomes have not evaluated the impact of gene flow on split time estimates derived from the *F(A|B)* statistic. However, the multiple instances of gene flow described in this and earlier work^45,48,110,111^, despite generally involving only small fractions of the genome, may bias such estimates. To investigate this, we performed coalescent simulations using *scrm*^95^, based on corrected demographic histories inferred with PSMC (**Supplementary Note 7**). We simulated scenarios with and without gene flow to compare the effects on *F(A|B)* and divergence.

In each simulation, we sampled a diploid genome B at time 0 and a haploid genome A from a distinct population that split from the population of B at a specified time T in the past. We varied T from 0 to up to 900,000 years in steps of 20,000 years, scaling time using a mutation rate of 1.45x10^-8^ per base pair per generation and a generation time of 29 years^83^. Each genome was 2.88Gb in length, with constant recombination rates matching those estimated from PSMC (**Table S42**, **Supplementary Note 7**). Population B followed the corrected demographic history inferred with PSMC for *Denisova 3*, *Denisova 25*, *Denisova 5* (Neandertal), or Mbuti, depending on the population modeled as B (**Table S42**, **Supplementary Note 7**). Population A was modeled with constant size, set to N_0_, the value used to scale theta, since this should have no effect on *F(A|B)* or divergence, which are both computed at sites ascertained in population B.

In simulations including gene flow from Neandertals into Denisovans, we introduced Neandertal ancestry levels similar to those estimated for *Denisova 3* and *Denisova 25* (**Supplementary Note 9**), i.e. 3% and 5%, respectively, with two pulses of admixture per genome (3.55% and 1.5% at 49,000 and 10,000 years before *Denisova 25*; 2.2% and 0.9% at 50,000 and 14,000 years before *Denisova 3*). When neither A nor B was sampled from a Neandertal, we assumed that the Neandertal population contributing gene flow split from Denisovans 550ka, adjusting for the branch shortening of genome B (65ka for *Denisova 3*, and 205ka for *Denisova 25*).

Additional scenarios included 5% gene flow from modern humans into Neandertals at 200ka (adjusted for branch shortening). In these cases, when neither A nor B was sampled from modern humans, we assumed that modern humans split from archaic humans 800ka (more ancient than most published estimates to maximize the impact of such gene flow). In some simulations, we also included gene flow from a “Super-Archaic” hominin who split 1Ma, contributing 5% to Denisovans at 350ka (adjusted for branch shortening).

For each simulation, we computed *F(A|B)* and recorded the average over ten replicates per split time T. Similarly, we computed divergence (as defined above) for each replicate, using a randomly sampled allele at positions heterozygous in the simulated genome B, and recorded the average over the ten replicates for each split time T.

We find that the *F(A|B)* statistic is highly sensitive to even low levels of gene flow (**Figure S41**), whereas divergence is only minimally affected. This difference likely arises because *F(A|B)* is calculated only at heterozygous sites in B, which are enriched for introgressed variants, whereas divergence is computed across all sites. We detail the impact of this below.

###### Denisovan-Denisovan split

For splits between Denisovan populations, Neandertal gene flow into Denisovans causes *F(A|B)*-based estimates to deviate increasingly with deeper split times (**Figure S42**). This leads to larger biases when estimating splits from *Denisova 3*, while estimates from *Denisova 25* remain largely unaffected for splits more recent than ~400ka. In contrast, divergence increases only for very recent splits between Denisovans (**Figure S43**). Gene flow from Neandertals should elevate the divergence between *Denisova 25* and *Denisova 3*, particularly when both individuals carry Neandertal ancestry (see the shoulder in **Figure S43B** when gene flow predates the split), but the effect should be minimal for the divergence on the *Denisova 3* lineage.

###### Neandertal-Denisovan split

For the split between Denisovans and Neandertals who contributed gene flow, the deviation from the *F(A|B)* values expected in the absence of gene flow increases with the split time (**Figure S41**). Therefore, one can expect again a greater effect on the *F(A|B)*-based estimates for *Denisova 3* than for *Denisova 25* because of their age difference. In contrast, the effect on divergence-based estimates is negligible for these levels of Neandertal gene flow into Denisovans.

Modern human ancestry in Neandertals and “Super-Archaic” ancestry in Denisovans are also expected to inflate divergence between Neandertals and Denisovans. Assuming 5% modern human ancestry in Neandertals from a population that split 800ka, and 5% “Super-Archaic” ancestry in Denisovans from a population that split 1Ma, their combined effect leads to a bias in the split time estimate of less than 50,000 years. This deviation decreases with increasing split time (**Figure S44**).

###### Modern-archaic human split

Given gene flow is expected to lead to large biases for *F(A|B)*-based estimates on deep split times (**Figure S42**), the split between modern and archaic humans should be investigated using divergence. Modern human ancestry in Neandertals reduces divergence between modern humans and Neandertals. This effect is expected to be approximately twice as large when divergence is measured on the Neandertal branch compared to the modern human branch (**Figure S45**). Divergence between modern humans and Denisovans could be used as an alternative, but contributions from deeply diverging lineages suggested in both modern humans^112,113^ and Denisovans^48^ (**Supplementary Note 10**) could inflate these estimates. Consequently, divergence-based split time estimates using Denisovans should represent an upper bound for the split time between modern and archaic humans. Estimates using Neandertals provide a lower bound, though both may be affected by gene flow from a divergent population into modern humans.

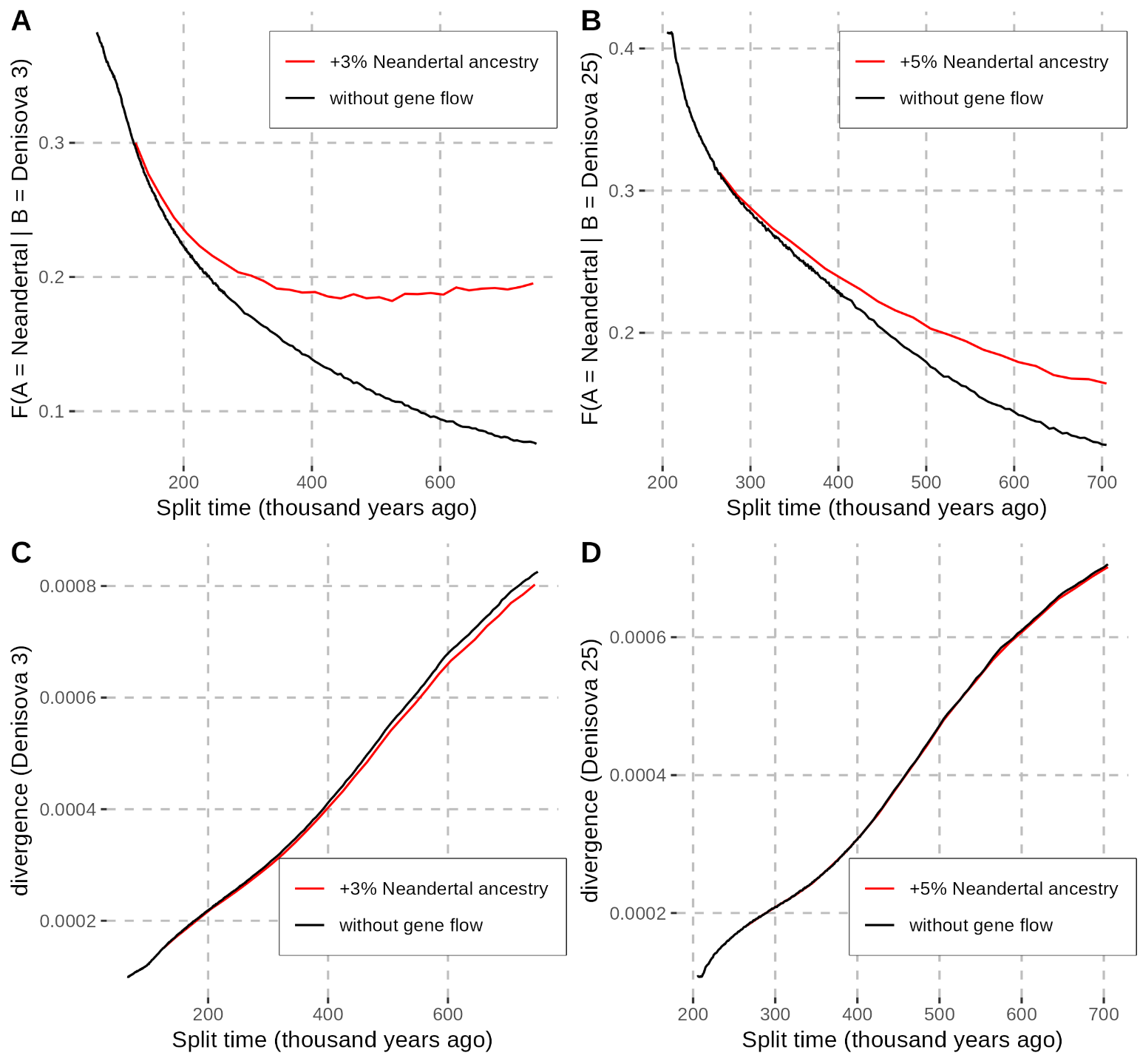

**Figure S41: Effect of Neandertal gene flow on F(A|B) and divergence to estimate the Neandertal-Denisovan split time.** The F(A|B) statistic was computed with coalescent simulations (using scrm) following the corrected demographic history estimated with PSMC for Denisova 3 (**A** and **C**) and Denisova 25 (**B** and **D**). The reported values correspond to the average of the F(A|B) statistic over ten replicates. The red and black curves correspond to simulations with and without additional Neandertal ancestry (in population B) from a population that split as indicated on the x-axis (after correcting for the ages of Denisova 3 and Denisova 25). This ancestry was introduced in two pulses, a first pulse of 2.2% followed by an additional 0.9% (3% in total) inherited 50,000 and 14,000 years before Denisova 3 in panels A and C, and 3.55% followed by 1.5% (5% in total) inherited 50,000 and 10,000 years before Denisova 25 in panels B and D.

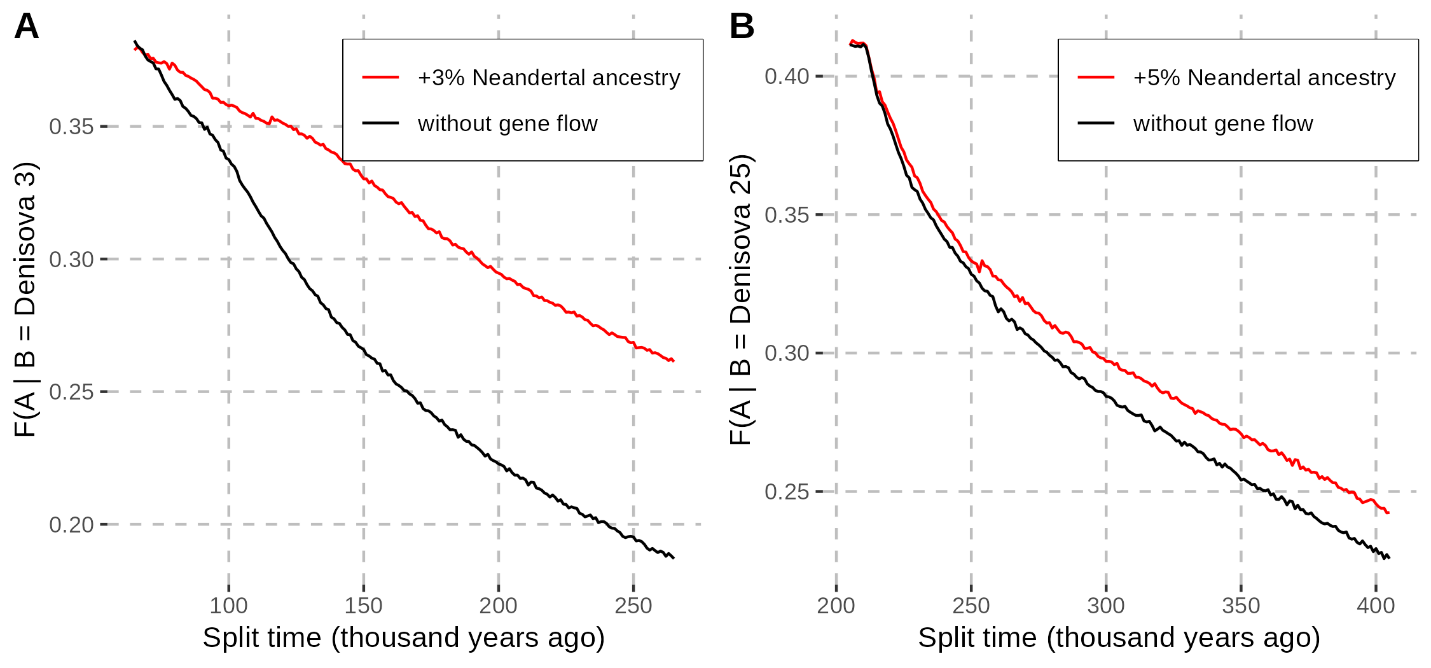

**Figure S42: Effect of Neandertal gene flow on F(A|B) between a Denisovan and an individual other than a Neandertal.** The F(A|B) statistic was computed with coalescent simulations (using scrm) following the corrected demographic history estimated with PSMC for Denisova 3 (**A**) and Denisova 25 (**B**). The reported values correspond to the average of the F(A|B) statistic over ten replicates. The red and black curves correspond to simulations with and without additional Neandertal ancestry (in population B) from a population that split 450,000 years ago (385,000 and 250,000 years before Denisova 3 and Denisova 25, respectively). This ancestry was introduced in two pulses as described in the text above.

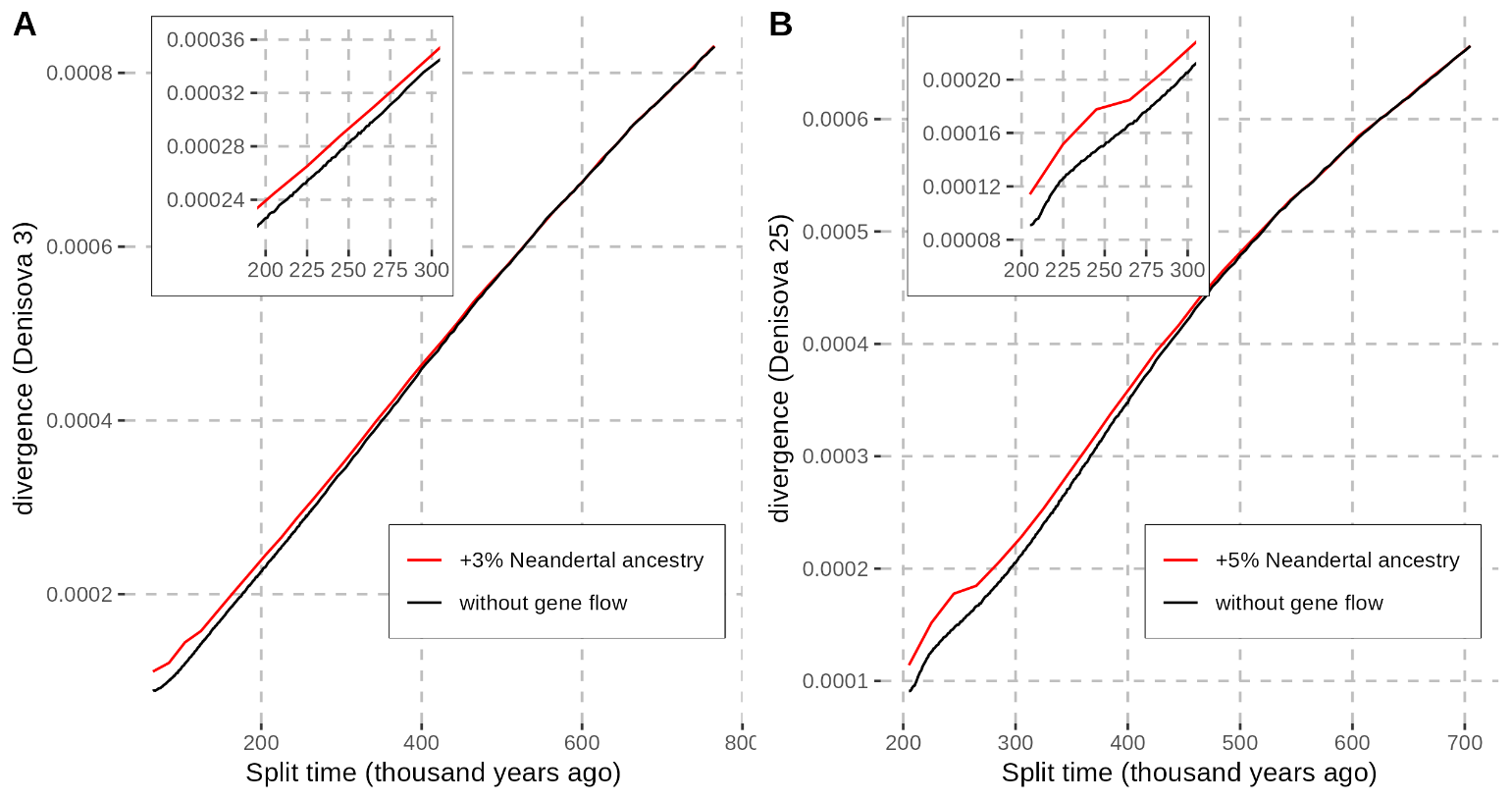

**Figure S43: Effect of Neandertal gene flow on divergence between a Denisovan and an individual other than a Neandertal.** The inset provides more detail for the effect expected for split times between 200ka and 300ka, more relevant for the split between Denisovans. See caption of **Figure S42** for details about the Neandertal gene flow and the layout of the figure.

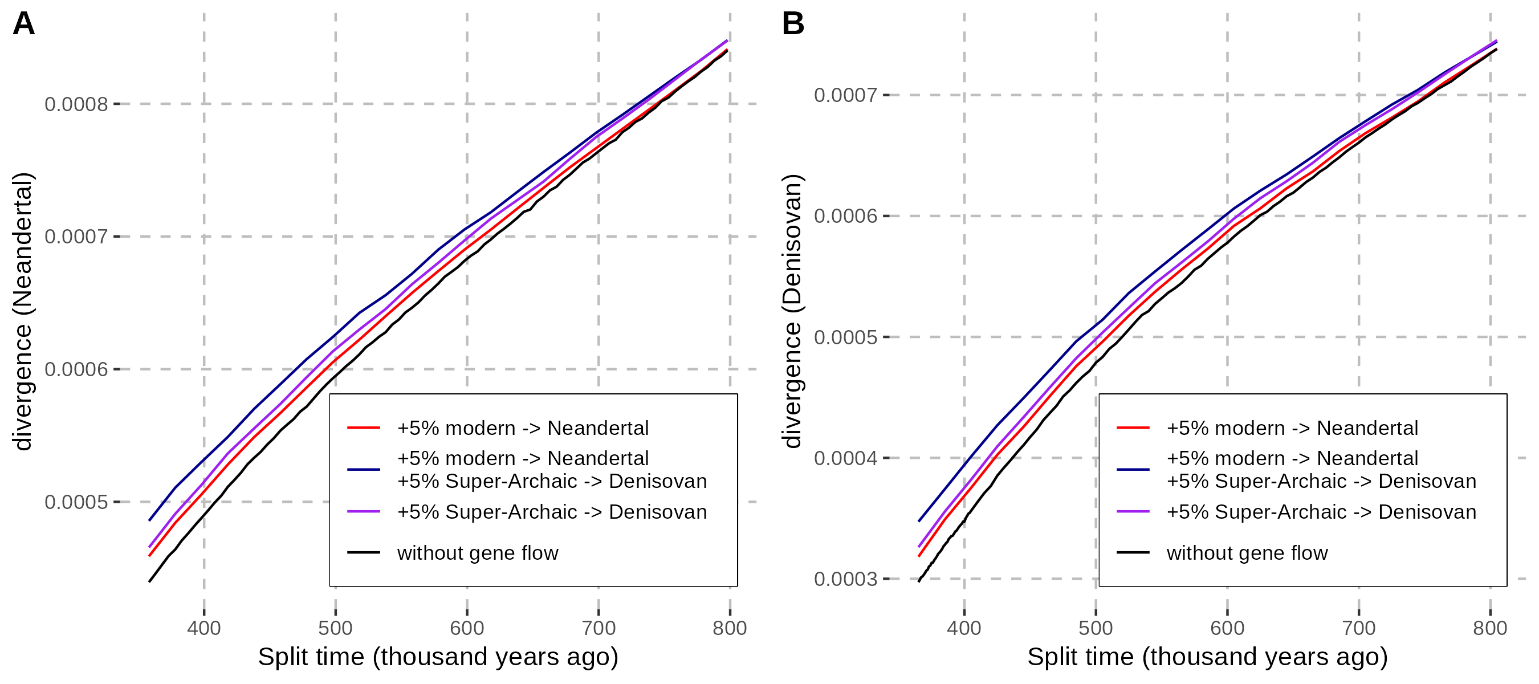

**Figure S44: Effect of modern human and “Super-Archaic” gene flows on divergence between Neandertals and Denisovans.** Divergence was computed with coalescent simulations (using scrm) following the corrected demographic history estimated with PSMC for the Denisova 5 Neandertal (**A**) and Denisova 25 (**B**). The reported values correspond to the average divergence over ten replicates. The colours correspond to simulations with and without gene flow. Modern human and “Super-Archaic” gene flows are described in the text above.

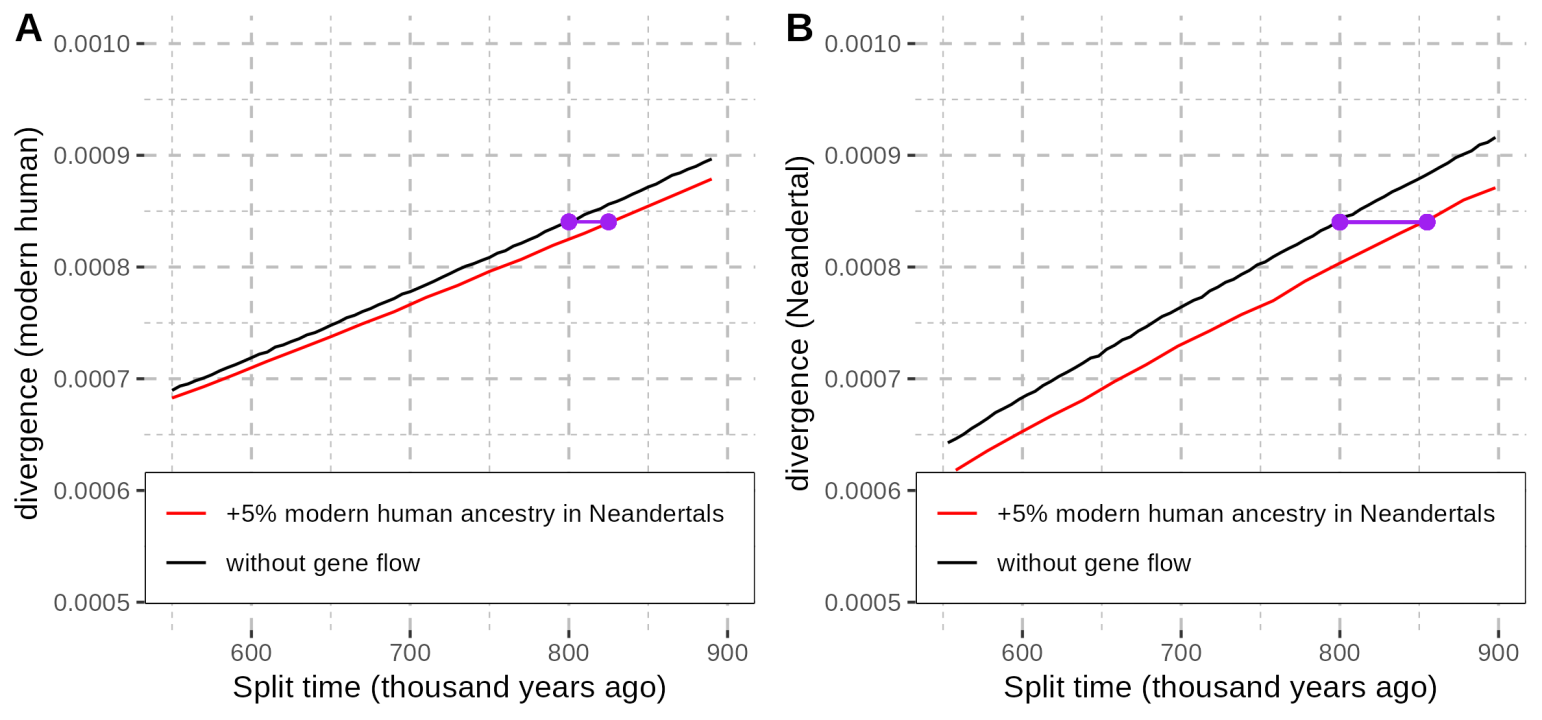

**Figure S45: Effect of modern human gene flow on divergence between modern humans and Neandertals.** Divergence was computed with coalescent simulations (using scrm) following the corrected demographic history estimated with PSMC for Mbuti (**A**) and the Denisova 5 Neandertal (**B**). The reported values correspond to the average divergence over ten replicates. The red and black curves correspond to simulations with and without gene flow, respectively, with 5% modern human ancestry in Neandertals inherited 200ka (after adjustment for the age of the individual, i.e. 0 for the Mbuti and 118ka for Denisova 5). Purple segments show the difference in split time for a divergence that would correspond to 800ka in the absence of gene flow.

##### Split time estimates using *F(A|B)* and divergence computed on the real data

Based on the simulations presented above we expect *F(A|B)* to give an underestimate of the split times in the presence of gene flow (except for between *Denisova 3* and *Denisova 25* when *Denisova 25* is population B). We therefore compared the estimates obtained from the *F(A|B)* and divergence statistics on the real data. We computed *F(A|B)* using only transversions among the genotype calls passing standard filters (**Supplementary Note 3**) to avoid recurrent mutations. Divergence was calculated as the proportion of derived alleles in genome B (after randomly sampling one allele at heterozygous sites) that are private to B relative to A (also randomly sampling one allele at heterozygous sites), using all genotype calls passing the standard filters of A and B. For *F(A|B)* the ancestral alleles were defined as those shared by four great ape reference genomes (panTro4, panpan1.1, gorGor3, ponAbe2), while for the divergence calculation the ancestral alleles were determined using panTro4. We computed standard errors (SE) of both statistics with a block jackknife using a window size of 5Mb.

To convert the observed values of the statistics into split time estimates, we again used coalescent simulations generated with *scrm*^95^ and based on the corrected demographic histories estimated with *PSMC* for population B (**Figure S32**; **Table S41**). As described above, we sampled a diploid genome B at time 0 and a haploid genome A at varying times T in the past to compute a calibration curve for each statistic. We varied T from 1,000 to 200,000 years in steps of 1,000 years, and from 205,000 to 900,000 years in steps of 5,000 years. We computed *F(A|B)* and divergence for each simulation, and calculated an expected value for the statistics with ten replicates for each time T. We then fitted the observed values against the calibration curves to obtain population split time estimates.

###### Uncorrected Denisovan-Denisovan split time estimates

Using *F(A|B)*, we estimated that the populations of *Denisova 3* and *Denisova 25* split 139,000 years and 23,000 years before the two individuals lived, respectively, which correspond to 204ka (2 SE: 196ka-212ka) and 228ka (2 SE: 226ka-232ka), assuming that *Denisova 3* and *Denisova 25* lived 65ka and 205ka (**Table S44**). As observed in simulations including 3 and 5% Neandertal ancestry in these genomes, *F(A|B)* is expected to be inflated for *Denisova 3*, leading to a more recent split time estimate, whereas these levels of Neandertal ancestry should have a minimal effect on the estimates from *Denisova 25*. The opposite should be true for estimates based on divergence, yet we obtained similar estimates (**Table S45**), 183ka (2 SE: 178-188) and 231ka (2 SE: 228-235). We show in the next section that this is likely due to the influence of selection.

###### Uncorrected Neandertal-Denisovan split time estimates

*F(A|B)*-based split time estimates between Neandertals and Denisovans varied between 415ka and 483ka (**Table S46**), closely matching divergence-based estimates (**Table S47**), which varied between 419ka and 505ka, with differences depending on whether divergence was computed along the Neandertal or Denisovan lineage. This is again inconsistent with the effect of gene flow in our simulations, which we show to be likely due to the influence of selection in the next section.

###### Uncorrected modern-archaic human split time estimates

Finally, *F(A|B)*-based population split time estimates between archaic and modern humans varied between 574ka and 670ka (**Table S48**), with most of the variation stemming from differences in how Neandertals and Denisovans relate to modern humans, presumably due to modern human ancestry in the Neandertal genomes^103,110,111,114^, and “Super-Archaic” ancestry in the Denisovan genomes^48^ (**Supplementary Note 10**). This is consistent with previous estimates and similar between *Denisova 3* and *Denisova 25*. Surprisingly, despite the large bias expected for these *F(A|B)*-based estimates in our simulations above, divergence-based estimates are again similar to those *F(A|B)*-based estimates, between 538ka and 640ka. This is likely due to the effect of selection increasing *F(A|B)*-based estimates but decreasing divergence-based estimates (see next section).

**Table S44: Split time estimates between Denisova 3 and Denisova 25 obtained with the F(A|B) statistic.** Standard errors (SE) were computed with a block jackknife using a window size of 5Mb.

| **Genome A** | **Genome B** | ***F(A\|B)* (%) [SE]** | **Estimate (ka)**  **[2 SE]** | **Branch shortening of genome B used for calibration (ka)** |
| --- | --- | --- | --- | --- |
| *Denisova 3* | *Denisova 25* | 35.96 [±0.24] | 228 [226-232] | 205 |
| *Denisova 25* | *Denisova 3* | 22.12 [±0.26] | 204 [196-212] | 65 |

**Table S45: Split time estimates between Denisova 3 and Denisova 25 based on divergence.** Divergence was computed as the proportion of bases with a derived allele in genome B that is not shared with genome A. Standard errors (SE) were computed with a block jackknife using a window size of 5Mb.

| **Genome A** | **Genome B** | **Divergence (B)**  **(×10^-4^) [SE]** | **Estimate (ka)**  **[2 SE]** | **Branch shortening of genome B used for calibration (ka)** |
| --- | --- | --- | --- | --- |
| *Denisova 3* | *Denisova 25* | 1.47 [±0.02] | 231 [228-235] | 205 |
| *Denisova 25* | *Denisova 3* | 2.04 [±0.02] | 183 [178-188] | 65 |

**Table S46: Split time estimates between Denisovans and Neandertals obtained with the F(A|B) statistic.** Standard errors (SE) were computed with a block jackknife using a window size of 5Mb.

| **Genome A** | **Genome B** | ***F(A\|B)* (%)**  **[SE]** | **Estimate (ka) [2 SE]** | **Branch shortening of genome B used for calibration (ka)** |
| --- | --- | --- | --- | --- |
| *Denisova 5* (Neandertal) | *Denisova 3* | 13.42 [±0.34] | 415 [395-440] | 65 |
| *Chagyrskaya 8* |  | 13.04 [±0.33] | 430 [405-455] |  |
| *Vindija 33.19* |  | 13.04 [±0.32] | 430 [405-445] |  |
| *Denisova 5* (Neandertal) | *Denisova 25* | 20.23 [±0.35] | 450 [440-465] | 205 |
| *Chagyrskaya 8* |  | 20.39 [±0.36] | 445 [435-460] |  |
| *Vindija 33.19* |  | 20.31 [±0.36] | 450 [435-465] |  |
| *Denisova 3* | *Denisova 5* (Neandertal) | 16.57 [±0.29] | 478 [458-493] | 118 |
|  | *Chagyrskaya 8* | 13.74 [±0.27] | 464 [444-489] | 74 |
|  | *Vindija 33.19* | 12.41 [±0.27] | 467 [437-487] | 47 |
| *Denisova 25* | *Denisova 5* (Neandertal) | 16.36 [±0.28] | 483 [468-498] | 118 |
|  | *Chagyrskaya 8* | 13.59 [±0.26] | 469 [449-494] | 74 |
|  | *Vindija 33.19* | 12.28 [±0.26] | 472 [442-492] | 47 |

**Table S47: Split time estimates between Denisovans and Neandertals based on divergence.** Divergence was computed as the proportion of bases with a derived allele in genome B that is not shared with genome A. Standard errors (SE) were computed with a block jackknife using a window size of 5Mb.

| **Genome A** | **Genome B** | **Divergence (B)**  **(×10^-4^) [SE]** | **Estimate (ka) [2 SE]** | **Branch shortening of genome B used for calibration (ka)** |
| --- | --- | --- | --- | --- |
| *Denisova 5* (Neandertal) | *Denisova 3* | 5.51 [±0.04] | 505 [495-510] | 65 |
| *Chagyrskaya 8* |  | 5.54 [±0.04] | 505 [500-510] |  |
| *Vindija 33.19* |  | 5.54 [±0.04] | 505 [500-510] |  |
| *Denisova 5* (Neandertal) | *Denisova 25* | 4.82 [±0.04] | 505 [500-510] | 205 |
| *Chagyrskaya 8* |  | 4.84 [±0.04] | 505 [500-510] |  |
| *Vindija 33.19* |  | 4.85 [±0.04] | 505 [505-515] |  |
| *Denisova 3* | *Denisova 5* (Neandertal) | 5.33 [±0.04] | 428 [423-433] | 118 |
|  | *Chagyrskaya 8* | 5.53 [±0.04] | 429 [419-434] | 74 |
|  | *Vindija 33.19* | 5.64 [±0.04] | 422 [417-427] | 47 |
| *Denisova 25* | *Denisova 5* (Neandertal) | 5.23 [±0.04] | 423 [418-428] | 118 |
|  | *Chagyrskaya 8* | 5.43 [±0.26] | 419 [414-424] | 74 |
|  | *Vindija 33.19* | 5.54 [±0.04] | 422 [417-427] | 47 |

**Table S48: Split time estimates between archaic and modern humans obtained with the F(A|B) statistic.** Standard errors (SE) were computed with a block jackknife using a window size of 5Mb.

| **Genome A** | **Genome B** | ***F(A\|B)* (%) [SE]** | **Estimate (ka)**  **[2 SE]** | **Branch shortening of genome B used for calibration (ka)** |
| --- | --- | --- | --- | --- |
| *Denisova 5* (Neandertal) | Mbuti (HGDP00982) | 17.55 [±0.12] | 615 [600-625] | 0 |
| *Chagyrskaya 8* |  | 17.64 [±0.12] | 610 [595-620] | 0 |
| *Vindija 33.19* |  | 17.48 [±0.12] | 620 [605-630] | 0 |
| *Denisova 3* |  | 17.20 [±0.12] | 635 [620-645] | 0 |
| *Denisova 25* |  | 16.98 [±0.12] | 645 [630-660] | 0 |
| Mbuti (HGDP00982) | *Denisova 5* (Neandertal) | 13.29 [±0.24] | 613 [583-643] | 118 |
|  | *Chagyrskaya 8* | 11.54 [±0.25] | 574 [544-594] | 74 |
|  | *Vindija 33.19* | 10.39 [±0.25] | 567 [542-597] | 47 |
|  | *Denisova 3* | 8.51 [±0.22] | 670 [630-705] | 65 |
|  | *Denisova 25* | 13.32 [±0.23] | 645 [620-655] | 205 |

**Table S49: Split time estimates between archaic and modern humans based on divergence.** Divergence was computed as the proportion of bases with a derived allele in genome B that is not shared with genome A. Standard errors (SE) were computed with a block jackknife using a window size of 5Mb.

| **Genome A** | **Genome B** | **Divergence (B)**  **(×10^-4^) [SE]** | **Estimate (ka)**  **[2 SE]** | **Branch shortening of genome B used for calibration (ka)** |
| --- | --- | --- | --- | --- |
| *Denisova 5* (Neandertal) | Mbuti (HGDP00982) | 7.30 [±0.05] | 610 [585-635] | 0 |
| *Chagyrskaya 8* |  | 7.29 [±0.05] | 610 [580-635] | 0 |
| *Vindija 33.19* |  | 7.32 [±0.05] | 615 [585-635] | 0 |
| *Denisova 3* |  | 7.44 [±0.06] | 640 [615-665] | 0 |
| *Denisova 25* |  | 7.34 [±0.05] | 620 [590-640] | 0 |
| Mbuti (HGDP00982) | *Denisova 5* (Neandertal) | 6.71 [±0.05] | 538 [528-548] | 118 |
|  | *Chagyrskaya 8* | 6.88 [±0.05] | 539 [529-544] | 74 |
|  | *Vindija 33.19* | 7.01 [±0.05] | 547 [532-557] | 47 |
|  | *Denisova 3* | 7.04 [±0.05] | 625 [615-635] | 65 |
|  | *Denisova 25* | 6.34 [±0.05] | 625 [615-630] | 205 |

##### Effect of background selection on population split time estimates

All estimates above ignore the effect of selection. Background selection, the removal of genetic diversity linked to deleterious mutations, is expected to increase genetic drift, as it mimics a reduction in effective population size^115^. This effect is not modeled in our simulations to generate the calibration curves and likely inflates the split time estimates based on the *F(A|B)* statistic. In contrast, selection is expected to reduce coalescent times in the ancestral population, thus biasing divergence-based estimates downward.

To correct for the effect of background selection, we restricted our analysis to regions that presumably evolve neutrally, defined as those with a B-score above 0.9^104^, which include about 955Mb. We recalculated *F(A|B)* and divergence within these regions and converted the values to split time estimates using calibration curves based on the demographic histories inferred from the same regions (**Supplementary Note 7**). For generating these calibration curves, we used a mutation rate of 1.58x10^-8^ per base pair per generation (9% higher than genome-wide, as estimated for these regions that presumably evolve neutrally, **Supplementary Note 7**), and a generation time of 29 years.

*F(A|B)*-based estimates for the split time between the ancestors of modern and archaic humans in these regions are about 150,000 years more recent than the estimates obtained with the whole genome (**Table S50**), falling between 417ka and 530ka. Similarly, the estimates for the Neandertal-Denisovan split time decreased by about 70,000 years, ranging between 322ka and 399ka. In contrast, the estimates for the split time between the two Denisovan populations remained similar, 173ka and 220ka. Therefore, while background selection was indeed inflating the uncorrected, genome-wide *F(A|B)*-based estimates, the values corrected for background selection are now more recent and downward biased due to gene flow, as shown by our simulations.

Divergence-based estimates, by contrast, increase when restricted to neutrally-evolving regions (**Table S51**), consistent with the expectation that selection reduces coalescent times in the common ancestral population. Estimated split times between modern and archaic humans range from 717ka to 825ka, those between Neandertals and Denisovans from 504ka to 585ka, and those for the split between the two Denisovan groups are 208ka (2 SE: 202-215ka) and 272ka (2 SE: 264-279ka), for the *Denisova 3* and *Denisova 25* lineages, respectively. The latter likely represent an overestimate, as our simulations suggest that Neandertal gene flow into Denisovans inflates divergence on the *Denisova 25* lineage more than for that of *Denisova 3* (**Figure S43**). Notably, the *F(A|B)*-based estimate for *Denisova 25* (220ka; 2 SE: 218-222ka), which is minimally affected by Neandertal ancestry in our simulations (**Figure S42**), is more consistent with the divergence-based estimate along the *Denisova 3* lineage.

##### Summary

Our simulations show that *F(A|B)* is sensitive to gene flow, particularly for the estimates of the Neandertal-Denisovan and archaic-modern human split times. In contrast, sequence divergence is less affected, given the generally low levels of gene flow. In the real data, the impact of gene flow is masked by the opposing effect of selection. Using the divergence statistic and restricting analyses to putatively neutral regions, we estimate the split between modern and archaic humans at 825-717ka, and between Neandertals and Denisovans at 585-504ka. For the split between Denisovans, the *F(A|B)*-based estimate using *Denisova 25* as B, 15,000 years (±2,000) before *Denisova 25* lived, and the divergence-based estimate using *Denisova 3* as B, 143,000 (2 std errors: 137,000-150,000) years before *Denisova 3* lived, are likely to be the most reliable. Finally, we caution that these estimates should be regarded as approximate, as they rely on the accuracy of the demographic histories estimated with *PSMC* and may be inflated by even low levels of gene flow from deeply diverged populations in modern humans and Denisovans, potentially by up to 50,000 years under the parameters explored here (***Figure S44***).

**Table S50: Split time estimates obtained with the F(A|B) statistic in genomic regions that presumably evolve neutrally.** F(A|B) statistics were computed in regions of the genome with at most 10% of their genetic diversity lost (in modern humans) presumably due to background selection (minimum B-score of 0.9^104^). The calibration curves used for the conversion into split times were computed with simulations using the demographic history estimated with PSMC in the same genomic regions (**Figure S37**). Standard errors (SE) were computed with a block jackknife using a window size of 5Mb.

| **Genome A** | **Genome B** | ***F(A\|B)* (%) [SE]** | **Estimate (ka)**  **[2 SE]** | **Branch shortening of genome B used for calibration** |
| --- | --- | --- | --- | --- |
| *Denisova 5* (Neandertal) | Mbuti (HGDP00982) | 19.16 [±0.13] | 445 [430-455] | 0 |
| *Chagyrskaya 8* |  | 19.20 [±0.13] | 440 [430-450] | 0 |
| *Vindija 33.19* |  | 19.06 [±0.13] | 450 [435-460] | 0 |
| *Denisova 3* |  | 18.71 [±0.13] | 465 [450-475] | 0 |
| *Denisova 25* |  | 18.53 [±0.13] | 470 [460-485] | 0 |
| Mbuti (HGDP00982) | *Denisova 5* (Neandertal) | 15.15 [±0.31] | 478 [453-498] | 118 |
|  | *Chagyrskaya 8* | 12.69 [±0.31] | 484 [449-509] | 74 |
|  | *Vindija 33.19* | 11.82 [±0.28] | 417 [392-442] | 47 |
|  | *Denisova 3* | 9.87 [±0.27] | 505 [475-545] | 65 |
|  | *Denisova 25* | 14.69 [±0.27] | 530 [515-550] | 205 |
| *Denisova 5* (Neandertal) | *Denisova 3* | 14.42 [±0.36] | 325 [310-350] | 65 |
| *Chagyrskaya 8* |  | 14.26 [±0.37] | 330 [315-355] |  |
| *Vindija 33.19* |  | 14.19 [±0.35] | 335 [315-355] |  |
| *Denisova 5* (Neandertal) | *Denisova 25* | 21.58 [±0.39] | 377 [364-390] | 205 |
| *Chagyrskaya 8* |  | 21.77 [±0.40] | 370 [361-385] |  |
| *Vindija 33.19* |  | 21.66 [±0.40] | 375 [363-387] |  |
| *Denisova 3* | *Denisova 5* (Neandertal) | 18.39 [±0.35] | 373 [353-393] | 118 |
|  | *Chagyrskaya 8* | 15.01 [±0.35] | 389 [364-414] | 74 |
|  | *Vindija 33.19* | 14.48 [±0.33] | 322 [302-337] | 47 |
| *Denisova 25* | *Denisova 5* (Neandertal) | 18.13 [±0.34] | 378 [363-398] | 118 |
|  | *Chagyrskaya 8* | 14.91 [±0.34] | 399 [369-419] | 74 |
|  | *Vindija 33.19* | 14.35 [±0.33] | 322 [342-307] | 47 |
| *Denisova 3* | *Denisova 25* | 37.30 [±0.32] | 220 [218-222] | 205 |
| *Denisova 25* | *Denisova 3* | 23.15 [±0.32] | 173 [164-180] | 65 |

**Table S51: Split time estimates based on divergence in genomic regions that presumably evolve neutrally.** Divergence was computed as the proportion of bases with a derived allele in genome B that is not shared with genome A in regions of the genome with at most 10% of their genetic diversity lost (in modern humans) presumably due to background selection (minimum B-score of 0.9^104^). The calibration curves used for the conversion into split times were computed with simulations using the demographic history estimated with PSMC in the same genomic regions (**Figure S37**). Standard errors (SE) were computed with a block jackknife using a window size of 5Mb.

| **Genome A** | **Genome B** | **Divergence (B)**  **(×10^-4^) [SE]** | **Estimate (ka)**  **[2 SE]** | **Branch shortening of genome B used for calibration** |
| --- | --- | --- | --- | --- |
| *Denisova 5* (Neandertal) | Mbuti (HGDP00982) | 8.42 [±0.06] | 805 [780-825] | 0 |
| *Chagyrskaya 8* |  | 8.42 [±0.06] | 805 [780-825] | 0 |
| *Vindija 33.19* |  | 8.42 [±0.06] | 805 [780-825] | 0 |
| *Denisova 3* |  | 8.57 [±0.06] | 825 [805-845] | 0 |
| *Denisova 25* |  | 8.46 [±0.06] | 810 [790-825] | 0 |
| Mbuti (HGDP00982) | *Denisova 5* (Neandertal) | 7.76 [±0.06] | 718 [698-728] | 118 |
|  | *Chagyrskaya 8* | 7.95 [±0.06] | 694 [679-704] | 74 |
|  | *Vindija 33.19* | 8.07 [±0.06] | 717 [702-732] | 47 |
|  | *Denisova 3* | 8.11 [±0.06] | 745 [730-760] | 65 |
|  | *Denisova 25* | 7.35 [±0.06] | 800 [785-820] | 205 |
| *Denisova 5* (Neandertal) | *Denisova 3* | 6.37 [±0.05] | 560 [550-570] | 65 |
| *Chagyrskaya 8* |  | 6.40 [±0.05] | 565 [555-575] |  |
| *Vindija 33.19* |  | 6.39 [±0.05] | 565 [555-575] |  |
| *Denisova 5* (Neandertal) | *Denisova 25* | 5.63 [±0.05] | 585 [570-595] | 205 |
| *Chagyrskaya 8* |  | 5.65 [±0.05] | 585 [575-595] |  |
| *Vindija 33.19* |  | 5.65 [±0.05] | 585 [575-595] |  |
| *Denisova 3* | *Denisova 5* (Neandertal) | 6.18 [±0.05] | 523 [513-538] | 118 |
|  | *Chagyrskaya 8* | 6.40 [±0.05] | 514 [504-529] | 74 |
|  | *Vindija 33.19* | 6.52 [±0.05] | 527 [512-537] | 47 |
| *Denisova 25* | *Denisova 5* (Neandertal) | 6.07 [±0.05] | 513 [503-523] | 118 |
|  | *Chagyrskaya 8* | 6.28 [±0.05] | 504 [489-514] | 74 |
|  | *Vindija 33.19* | 6.40 [±0.05] | 512 [502-522] | 47 |
| *Denisova 3* | *Denisova 25* | 1.73 [±0.04] | 272 [264-279] | 205 |
| *Denisova 25* | *Denisova 3* | 2.36 [±0.04] | 208 [202-215] | 65 |

### Supplementary Note 9: Neandertal ancestry

Genome-wide statistics from **Supplementary Note 8** indicate that *Denisova 3* and *Denisova 25* share significantly different proportions of alleles with Neandertals, suggesting varying levels of Neandertal ancestry among Denisovans. Here, we further characterise Neandertal ancestry in Denisovan genomes and reconstruct the admixture history between these groups.

#### Detection of Neandertal-like segments

We used *admixfrog* (version 0.7.2^97^) to detect Neandertal-like segments in the two high-coverage Denisovan genomes. *Admixfrog* models the genome of an individual (target) as a mosaic of DNA segments from different sources. We used three high-coverage Neandertal genomes as reference for the Neandertal source (including the *Denisova 5 Neandertal*, *Chagyrskaya 8* and *Vindija 33.19* genomes), as well as either *Denisova 3* or *Denisova 25* as reference for the Denisovan source, depending on whether the target is *Denisova 25* or *Denisova 3*, respectively. Standard filters were applied to the VCF file of each archaic genome independently (**Supplementary Note 3**), so that positions with missing or filtered out genotypes in some but not all individuals were included in the reference panel. No further filters were applied, we included all segregating sites in the reference for *admixfrog*. We inferred the ancestral state from the allele present in the chimpanzee reference genome (*panTro4*).

We ran *admixfrog* on the X chromosome separately from the autosomes, and, therefore, fitted the model parameters independently, to account for differences in the length of Neandertal-like segments due to the different modes of transmission of the X chromosome and the autosomes. As no fine-scale recombination map is available for Denisovans, we set the window size either to 0.005 centimorgans (cM) using the African-American recombination map (with linear interpolations when necessary), or to 5,000 base pairs (bp), in which case we used the African-American recombination map^99^ to convert the physical length of identified segments into genetic length. For all downstream analyses (except if stated otherwise), we only considered Neandertal-like segments of 0.05cM or longer, as these can be confidently inferred with *admixfrog*^97,116^. Neandertal-like segments can be erroneously fragmented due to the uneven distribution of the available data along the genome; *admixfrog* includes a penalty that decreases the probability to switch between ancestries when the posterior probability of a given ancestry is high. A high penalty results in fewer short ancestry segments, but may also merge segments that would appear longer than they are. Previous studies have applied penalty values of 0.1, 0.25 or 0.4 for detecting Neandertal and Denisovan ancestry in ancient modern human genomes^98,116-120^, or 0.2 in archaic human genomes^97^. Therefore, we used multiple values (0.01, 0.1, 0.25 and 0.4) to evaluate the results.

We detected between 56Mb and 166Mb of Neandertal ancestry in the *Denisova 25* genome (cumulative length of segments of at least 0.05cM), depending on the parameters used for calling segments (**Table S52**, **Figure S46**). In contrast, we detected between 34Mb and 87Mb of Neandertal ancestry in the *Denisova 3* genome (**Table S52**, **Figure S47**), about half that of *Denisova 25*.

#### Evaluation of identified Neandertal-like segments

We identified more Neandertal-like segments when using a window size of 5,000bp compared to 0.005cM (**Table S52**). Over 80% of the segments detected using a window-size of 0.005cM overlap with those identified with a window size of 5,000bp (***Table S52***), indicating overall consistency in the results. The choice of parameters also affects the proportion of homozygous segments, with higher penalties and window sizes of 5,000bp (instead of 0.005cM) leading to an increase in homozygosity. Depending on the parameters used, the proportion of homozygous Neandertal-like segments varies between 5-35% for *Denisova 25* and 9-41% for *Denisova 3* (**Table S53**).

As the genetic relationship to Neandertal and Denisovan genomes may be informative about the authenticity of the identified segments, we next examined the proportion of alleles shared between these segments and the Neandertal and Denisovan genomes (“branch support”, **Figure S48** and **Figure S49**). We counted sites in the Neandertal-like segments of *Denisova 3* or *Denisova 25* where they carry a derived allele that is seen in the *Denisova 5* Neandertal genome (or in the other Denisovan genome) while a Mbuti genome carries the ancestral version. First, stratifying segments by genetic length (including segments shorter than 0.05cM for this analysis) revealed that short segments share a lower proportion of alleles with the Neandertal genome, suggesting that a larger proportion of short segments are misidentified as Neandertal. However, an alternative explanation is that short segments were inherited from a more distantly related Neandertal population than longer ones. Interestingly, when stratifying segments by physical length (also including segments shorter than 0.05cM), we observe no difference in allele sharing with the Neandertal genome, but whether this supports the possibility that short segments, as determined by genetic length, are differently related to the Neandertal genomes than longer ones is unclear. Second, we found that higher penalties yield segments with reduced allele sharing with the Neandertal genome, implying that high penalties lead to more false positives (misclassification of Denisovan ancestry as Neandertal) and/or low penalties may introduce a reference bias favouring Neandertal-like segments that are closer to the reference Neandertal genomes (missing other Neandertal-like segments). As we could not resolve this issue, we performed subsequent analyses using all tested parameter combinations.

Another intriguing observation is that, despite Neandertal ancestry proportion being below 6% in both *Denisova 25* and *Denisova 3* (as detailed in the next section), up to 17% of Neandertal-like segments in *Denisova 25* overlap with those of *Denisova 3* (**Table S54**; **Figure S50** and **Figure S51**). This unexpectedly high overlap could indicate that some Neandertal ancestry was inherited from common admixture events. However, we used one Denisovan genome as a reference to identify the ancestry in the other Denisovan genome, which may introduce biases. Specifically, Neandertal segments in the reference Denisovan genome should be incorrectly considered as Denisovan ancestry, obscuring similar segments in the target Denisovan genome. For Neandertal ancestry to be detected in the same genomic region in both Denisovans, the two genomes should be sufficiently distinct, making it unlikely that these segments originate from a common admixture event with Neandertals, unless these segments can be detected in the heterozygous state. Indeed, we found that heterozygosity in *Denisova 3* is three times higher in genomic regions where a heterozygous Neandertal-like segment in *Denisova 25* overlaps with a Neandertal-like segment in *Denisova 3* (5.3x10^-4^ compared to 1.7x10^-4^ in segments where *Denisova 3* does not also carry a Neandertal-like segment), consistent with the presence of Neandertal ancestry in both genomes (see examples in **Figure S51**). In contrast, when both genomes carry Neandertal-like segments in the homozygous state, an inspection of their relationship to Neandertals suggests that these overlapping segments arise when one Denisovan is closer to a Neandertal in this genomic region while the other Denisovan is closer to a different Neandertal, with little allele sharing between the two Denisovans (**Figure S50**; **Table S56**). If these segments had originated from the same admixture event, they would exhibit the same relationship to Neandertals, which they do not. While it remains possible that some segments were inherited from a common event, the availability of only two Denisovan genomes limits our ability to test this hypothesis through the identification of shared segments.

Another potential confounding factor is the presence of Denisovan segments in the Neandertal reference genomes^48,97^, which could complicate the identification of Neandertal segments in the Denisovan genomes. Notably, we found that overlapping segments are more often homozygous than other Neandertal-like segments: up to 39% and 57% in *Denisova 3* for window sizes of 0.005cM and 5,000bp, respectively, and up to 26% and 48% in *Denisova 25* for the same window sizes. Since this subset of Neandertal-like segments is more often homozygous than other Neandertal-like segments in both genomes, it is possible that some of these segments actually represent Denisovan ancestry in one or a few Neandertal genomes. In addition, the observed correlation in states between *Denisova 3* and *Denisova 25*, with more overlapping segments being homozygous or heterozygous in both genomes than expected by chance (Pearson's Chi-squared test or Fisher’s exact test depending on the number of observations, see **Table S55**), further supports the possibility that Denisovan ancestry in Neandertal genomes contributes to the shared segments. However, the proportion of alleles shared between these overlapping segments and Neandertal/Denisovan genomes does not differ from that of all Neandertal-like segments (**Table S56**), the majority of which do not overlap. If a substantial fraction of these overlapping segments originated from Denisovan ancestry in Neandertal genomes, we would expect instead an increase in Denisovan-derived alleles, as there is no reason Denisovans would not share alleles in these regions.

To exclude potential confounding effects of Denisovan ancestry in the Neandertal reference genomes, we re-called Neandertal ancestry in *Denisova 3* and *Denisova 25* using a reference that is comprised of only the subset of sites that are monomorphic across all three Neandertal reference genomes (but differ from the other Denisovan genome). Excluding variable positions should effectively mask any Denisovan ancestry in Neandertal genomes, as such segments are unlikely to be shared by all three Neandertals, given the absence of evidence for Denisovan ancestry in *Vindija 33.19* (Mafesonni *et al*., 2020). Between 90% and 94% of segments identified with a window size of 0.005cM overlap with those identified using the reference including all sites (**Table S57**), indicating that most segments were recovered and that Denisovan ancestry in Neandertal genomes does not substantially contribute to the identified segments. In contrast, only 36% to 62% of segments identified with a window size of 5,000bp overlap with those identified with the reference including all sites (**Table S57**), suggesting that these segments are less reliable. As a result, no overlapping homozygous segments were identified using a window size of 0.005cM (**Table S58**), and among the few overlapping homozygous segments identified using a window size of 5kb (8 and 17 with a penalty of 0.25 and 0.4, respectively) only about half were identified with the reference including all sites, suggesting a large proportion of miscalls. Moreover, under these conditions, no correlation in states was observed between the overlapping segments of the two Denisovans (Pearson's Chi-squared test or Fisher’s exact test in **Table S58**), except for a single combination of parameters (5,000bp windows with a penalty of 0.1), where an anti-correlation was detected instead.

Overall, the majority of Neandertal-like segments identified in both Denisovan genomes appear to be genuine. However, our analyses clearly indicate that the identification of these segments is sensitive to reference biases and parameter choices. Therefore, we continue to assess their impact in subsequent analyses. Note that, unless stated otherwise, we analysed the segments called using the reference that contains all sites for subsequent analyses.

**Table S52: Number and breadth of Neandertal-like segments in the Denisova 3 and Denisova 25 genomes.** The counts in the “overlap” rows indicate the number of Neandertal-like segments (or positions) identified with a window size of 0.005cM that overlap with those identified with a window size of 5,000bp.

| **window size** | **penalty** | **# of Neandertal-like segments** | | **# of Neandertal-like segments≥0.05cM** | | **Cumulative length in bp (segments≥0.05cM)** | |
| --- | --- | --- | --- | --- | --- | --- | --- |
|  |  | ***Denisova3*** | ***Denisova25*** | ***Denisova3*** | ***Denisova25*** | ***Denisova3*** | ***Denisova25*** |
| 0.005cM | 0.01 | 2,504 | 3,886 | 279 | 464 | 33,610,825 | 56,489,738 |
|  | 0.1 | 3,227 | 4,692 | 573 | 921 | 55,754,588 | 102,187,710 |
|  | 0.25 | 3,748 | 5,347 | 754 | 1,120 | 68,779,479 | 129,170,116 |
|  | 0.4 | 4,189 | 6,064 | 909 | 1,317 | 80,887,098 | 149,830,617 |
| 5,000bp | 0.01 | 3,684 | 5,608 | 633 | 1,007 | 48,575,000 | 77,885,000 |
|  | 0.1 | 4,814 | 7,084 | 838 | 1,307 | 65,300,000 | 116,440,000 |
|  | 0.25 | 5,534 | 8,153 | 974 | 1,458 | 75,690,000 | 140,665,000 |
|  | 0.4 | 6,122 | 8,818 | 1,092 | 1,572 | 86,510,000 | 165,750,000 |
| overlap | 0.01 | 2,185 | 3,425 | 259 | 412 | 26,668,528 | 39,922,644 |
|  | 0.1 | 2,696 | 4,036 | 498 | 820 | 46,060,981 | 83,389,179 |
|  | 0.25 | 3,054 | 4,416 | 637 | 957 | 56,676,570 | 107,679,090 |
|  | 0.4 | 3,300 | 4,759 | 737 | 1,068 | 65,481,743 | 129,577,122 |

**Table S53: Proportion of Neandertal-like segments in the homozygous state in the Denisova 3 and Denisova 25 genomes.** Segments on the X chromosome were excluded because Denisova 25 only has one X chromosome, and therefore hemizygous segments on this chromosome.

| **window size** | **penalty** | **proportion of homozygous segments** | | | |
| --- | --- | --- | --- | --- | --- |
|  |  | **all** | | **≥0.05cM** | |
|  |  | ***Denisova3*** | ***Denisova25*** | ***Denisova3*** | ***Denisova25*** |
| 0.005cM | 0.01 | 0.39 | 0.30 | 0.09 | 0.05 |
|  | 0.1 | 0.44 | 0.38 | 0.22 | 0.12 |
|  | 0.25 | 0.46 | 0.42 | 0.28 | 0.17 |
|  | 0.4 | 0.48 | 0.47 | 0.32 | 0.22 |
| 5,000bp | 0.01 | 0.44 | 0.36 | 0.32 | 0.22 |
|  | 0.1 | 0.47 | 0.43 | 0.36 | 0.27 |
|  | 0.25 | 0.50 | 0.47 | 0.39 | 0.32 |
|  | 0.4 | 0.51 | 0.50 | 0.41 | 0.35 |

**Table S54: Overlapping Neandertal-like segments between the Denisova 3 and Denisova 25 genomes.**

| **window size** | **penalty** | **# of overlapping Neandertal-like segments (≥0.05cM)** | | **% of overlapping Neandertal-like segments (≥0.05cM)** | | **Cumulative length in bp of overlapping Neandertal ancestry (segments≥0.05cM)** |
| --- | --- | --- | --- | --- | --- | --- |
|  |  | ***Denisova3*** | ***Denisova25*** | ***Denisova3*** | ***Denisova25*** |  |
| 0.005cM | 0.01 | 17 | 17 | 6.1 | 3.7 | 1,565,741 |
|  | 0.1 | 71 | 69 | 12.4 | 7.5 | 5,845,856 |
|  | 0.25 | 122 | 118 | 16.2 | 10.5 | 9,234,173 |
|  | 0.4 | 184 | 175 | 20.2 | 13.3 | 13,194,430 |
| 5,000bp | 0.01 | 92 | 88 | 14.5 | 8.7 | 3,630,000 |
|  | 0.1 | 169 | 164 | 20.2 | 12.5 | 6,800,000 |
|  | 0.25 | 229 | 221 | 23.5 | 15.2 | 9,800,000 |
|  | 0.4 | 288 | 269 | 26.4 | 17.1 | 13,190,000 |

**Table S55: State of Neandertal-like segments overlapping between the Denisova 3 and Denisova 25 genomes.** Only segments that are 0.05cM or longer are included. Segments on the X chromosome were excluded here because Denisova 25 only has one X chromosome, and therefore hemizygous segments on this chromosome. We used a Pearson's Chi-squared test when the expected number of segments was above 5, and a Fisher’s exact test otherwise, to test whether the observed and expected counts significantly differ.

|  |  |  | **state in *Denisova 25*** | | | |  |
| --- | --- | --- | --- | --- | --- | --- | --- |
|  |  |  | **Observed** | | **Expected** | |  |
| **window size** | **penalty** | **state in *Denisova 3*** | **heterozygous** | **homozygous** | **heterozygous** | **homozygous** | **p-value** |
| **0.005cM** | **0.01** | **heterozygous** | 16 | 0 | NA | NA | NA |
|  |  | **homozygous** | 1 | 0 | NA | NA |  |
|  | **0.1** | **heterozygous** | 48 | 4 | 44 | 8 | 0.006 (Fisher) |
|  |  | **homozygous** | 12 | 7 | 16 | 3 |  |
|  | **0.25** | **heterozygous** | 66 | 11 | 61 | 16 | 0.024 (**χ^2^**) |
|  |  | **homozygous** | 30 | 15 | 35 | 10 |  |
|  | **0.4** | **heterozygous** | 97 | 14 | 84 | 27 | <10^-5^ (**χ^2^**) |
|  |  | **homozygous** | 40 | 31 | 53 | 18 |  |
| **5,000bp** | **0.01** | **heterozygous** | 47 | 12 | 44 | 15 | 0.152 (**χ^2^**) |
|  |  | **homozygous** | 21 | 12 | 24 | 9 |  |
|  | **0.1** | **heterozygous** | 76 | 21 | 64 | 33 | <10^-3^ (**χ^2^**) |
|  |  | **homozygous** | 34 | 35 | 46 | 23 |  |
|  | **0.25** | **heterozygous** | 83 | 28 | 64 | 46 | <10^-6^ (**χ^2^**) |
|  |  | **homozygous** | 44 | 74 | 63 | 45 |  |
|  | **0.4** | **heterozygous** | 90 | 32 | 68 | 54 | <10^-7^ (**χ^2^**) |
|  |  | **homozygous** | 61 | 89 | 83 | 67 |  |

**Table S56: Neandertal and Denisovan branch support in Neandertal-like segments inferred by admixfrog in the Denisova 25 genome.** Counts and proportions of derived and ancestral alleles within these segments are reported over all segments. Only segments that are 0.05cM or longer are included. Branch support values were computed separately for homozygous and heterozygous segments, as described in the text above, and for the subsets of segments overlapping Neandertal-like segments of Denisova 3. Segments that overlap those of Denisova 3 may do so partially (see examples in **Figure S50**). For results stratified by the segment length, see **Figure S48** and **Figure S49**.

| **state** | **set** | **penalty** | **Neandertal branch support** | | | **Denisovan branch support** | | |
| --- | --- | --- | --- | --- | --- | --- | --- | --- |
|  |  |  | **# derived** | **# ancestral** | **Proportion derived** | **# derived** | **# ancestral** | **Proportion derived** |
| **homozygous in *Denisova 25*** | **All segments** | **0.01** | 353 | 284 | 0.554 | 24 | 813 | 0.029 |
|  |  | **0.1** | 1,285 | 816 | 0.612 | 80 | 2,617 | 0.030 |
|  |  | **0.25** | 1,820 | 1,372 | 0.570 | 115 | 4,094 | 0.027 |
|  |  | **0.4** | 2,342 | 1,910 | 0.551 | 177 | 5,291 | 0.032 |
|  | **Overlapping Neandertal-like segments of *Denisova 3*** | **0.01** | NA | NA | NA | NA | NA | NA |
|  |  | **0.1** | 149 | 88 | 0.629 | 26 | 319 | 0.075 |
|  |  | **0.25** | 293 | 220 | 0.571 | 31 | 600 | 0.049 |
|  |  | **0.4** | 536 | 445 | 0.546 | 47 | 1,157 | 0.039 |
|  | **Overlapping homozygous Neandertal-like segments of *Denisova 3*** | **0.01** | NA | NA | NA | NA | NA | NA |
|  |  | **0.1** | 81 | 51 | 0.614 | 14 | 155 | 0.083 |
|  |  | **0.25** | 197 | 164 | 0.546 | 16 | 362 | 0.042 |
|  |  | **0.4** | 357 | 331 | 0.519 | 21 | 733 | 0.028 |
| **heterozygous in *Denisova 25*** | **All segments** | **0.01** | 3,857 | 7,387 | 0.343 | 4,471 | 8,022 | 0.358 |
|  |  | **0.1** | 5,967 | 12,486 | 0.323 | 7,214 | 13,112 | 0.355 |
|  |  | **0.25** | 6,712 | 15,002 | 0.309 | 8,565 | 15,687 | 0.353 |
|  |  | **0.4** | 7,311 | 17,294 | 0.297 | 9,755 | 17,707 | 0.355 |
|  | **Overlapping Neandertal-like segments of *Denisova 3*** | **0.01** | 130 | 336 | 0.279 | 202 | 358 | 0.361 |
|  |  | **0.1** | 450 | 1,082 | 0.294 | 550 | 1,190 | 0.316 |
|  |  | **0.25** | 1,308 | 2,872 | 0.313 | 1,607 | 3,228 | 0.332 |
|  |  | **0.4** | 2,970 | 5,854 | 0.337 | 3,345 | 7,008 | 0.323 |

**Table S57: Number of Neandertal-like segments in the Denisova 3 and Denisova 25 genomes when the admixfrog reference does not include variable sites between Neandertals.** For comparison, we also report the number of Neandertal-like segments that overlap those identified with the admixfrog reference that includes all sites.

| **Window size** | **penalty** | **Number of Neandertal-like segments≥0.05cM** | | | |
| --- | --- | --- | --- | --- | --- |
|  |  | ***Denisova 3*** | | ***Denisova 25*** | |
|  |  | **All** | **Overlapping those identified will all sites** | **All** | **Overlapping those identified will all sites** |
| 0.005cM | 0.01 | 236 | 223 | 340 | 319 |
|  | 0.1 | 496 | 448 | 784 | 724 |
|  | 0.25 | 641 | 576 | 1,007 | 913 |
|  | 0.4 | 797 | 700 | 1,154 | 1,040 |
| 5,000bp | 0.01 | 530 | 221 | 856 | 309 |
|  | 0.1 | 767 | 433 | 1,195 | 689 |
|  | 0.25 | 920 | 534 | 1,389 | 856 |
|  | 0.4 | 1,063 | 625 | 1,601 | 934 |

**Table S58: State of Neandertal-like segments overlapping between the *Denisova 3* and *Denisova 25* genomes when the *admixfrog* reference does not include variable sites between Neandertals.** Only segments that are 0.05cM or longer are included. Segments on the X chromosome were excluded here because *Denisova 25* only has one X chromosome, and therefore hemizygous segments on this chromosome. We used a Pearson's Chi-squared test when the expected number of segments was above 5, and a Fisher’s exact test otherwise, to test whether the observed and expected counts significantly differ.

|  |  |  | **state in *Denisova 25*** | | | |  |
| --- | --- | --- | --- | --- | --- | --- | --- |
|  |  |  | **Observed** | | **Expected** | |  |
| **window size** | **penalty** | **state in *Denisova 3*** | **heterozygous** | **homozygous** | **heterozygous** | **homozygous** | **p-value** |
| **0.005cM** | **0.01** | **heterozygous** | 10 | 0 | NA | NA | NA |
|  |  | **homozygous** | 0 | 0 | NA | NA |  |
|  | **0.1** | **heterozygous** | 39 | 5 | 39 | 5 | 1  (Fisher) |
|  |  | **homozygous** | 2 | 0 | 2 | 0 |  |
|  | **0.25** | **heterozygous** | 61 | 18 | 62 | 17 | 0.337  (Fisher) |
|  |  | **homozygous** | 7 | 0 | 6 | 1 |  |
|  | **0.4** | **heterozygous** | 81 | 30 | 83 | 28 | 0.292  (Fisher) |
|  |  | **homozygous** | 11 | 1 | 9 | 3 |  |
| **5,000bp** | **0.01** | **heterozygous** | 35 | 16 | 37 | 14 | 0.096  (Fisher) |
|  |  | **homozygous** | 9 | 0 | 7 | 2 |  |
|  | **0.1** | **heterozygous** | 62 | 32 | 68 | 26 | 0.004  (**χ^2^**) |
|  |  | **homozygous** | 21 | 0 | 15 | 6 |  |
|  | **0.25** | **heterozygous** | 89 | 48 | 93 | 44 | 0.177  (**χ^2^**) |
|  |  | **homozygous** | 29 | 8 | 25 | 11 |  |
|  | **0.4** | **heterozygous** | 107 | 60 | 110 | 57 | 0.822  (**χ^2^**) |
|  |  | **homozygous** | 43 | 17 | 40 | 20 |  |

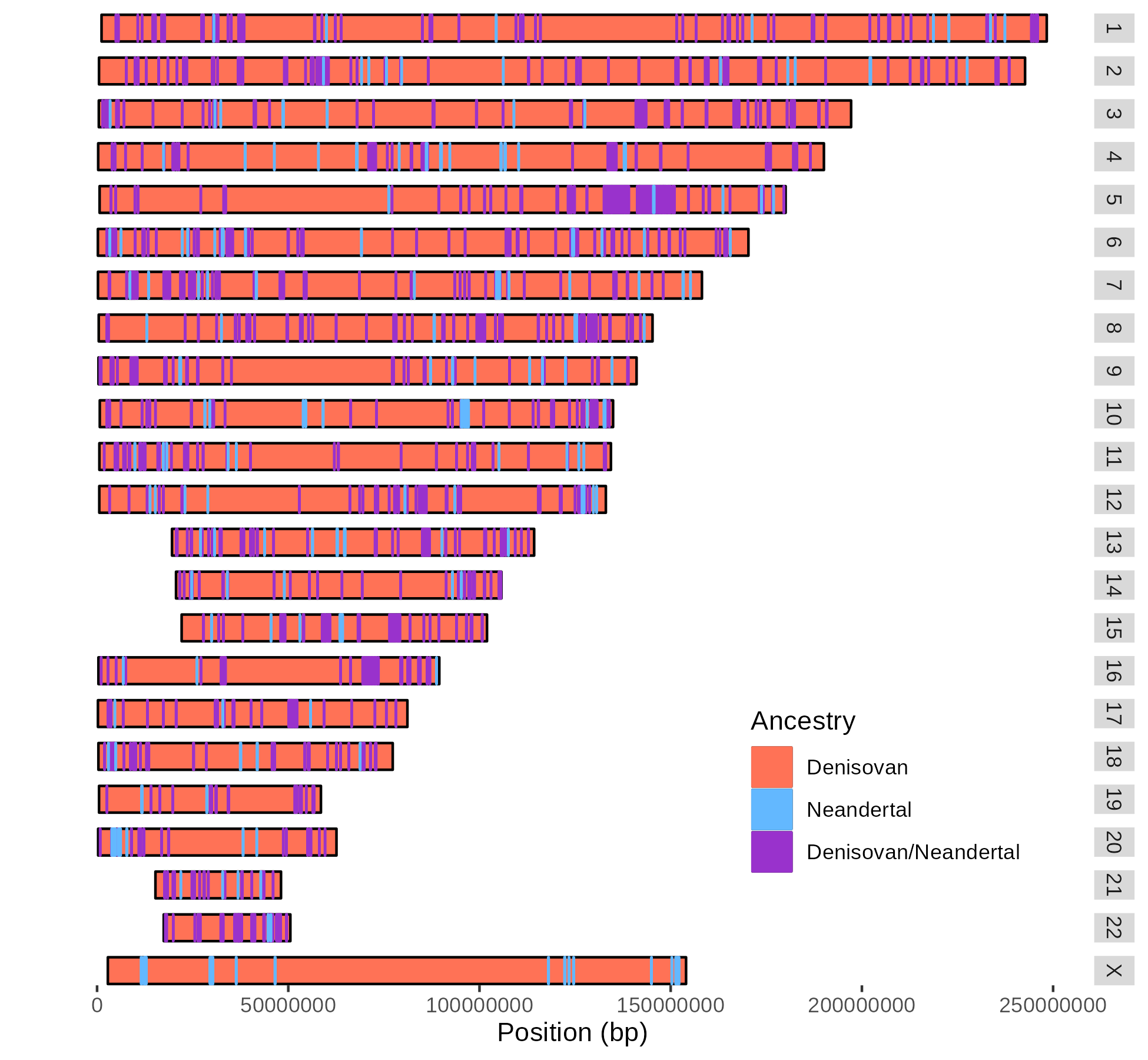

**Figure S46: Map of Neandertal ancestry in the Denisova 25 genome.** Only the Neandertal-like segments of 0.05cM and longer are represented. Window sizes were set to 0.005cM using the African-American recombination map^99^ and a penalty of 0.25 was used to call the segments.

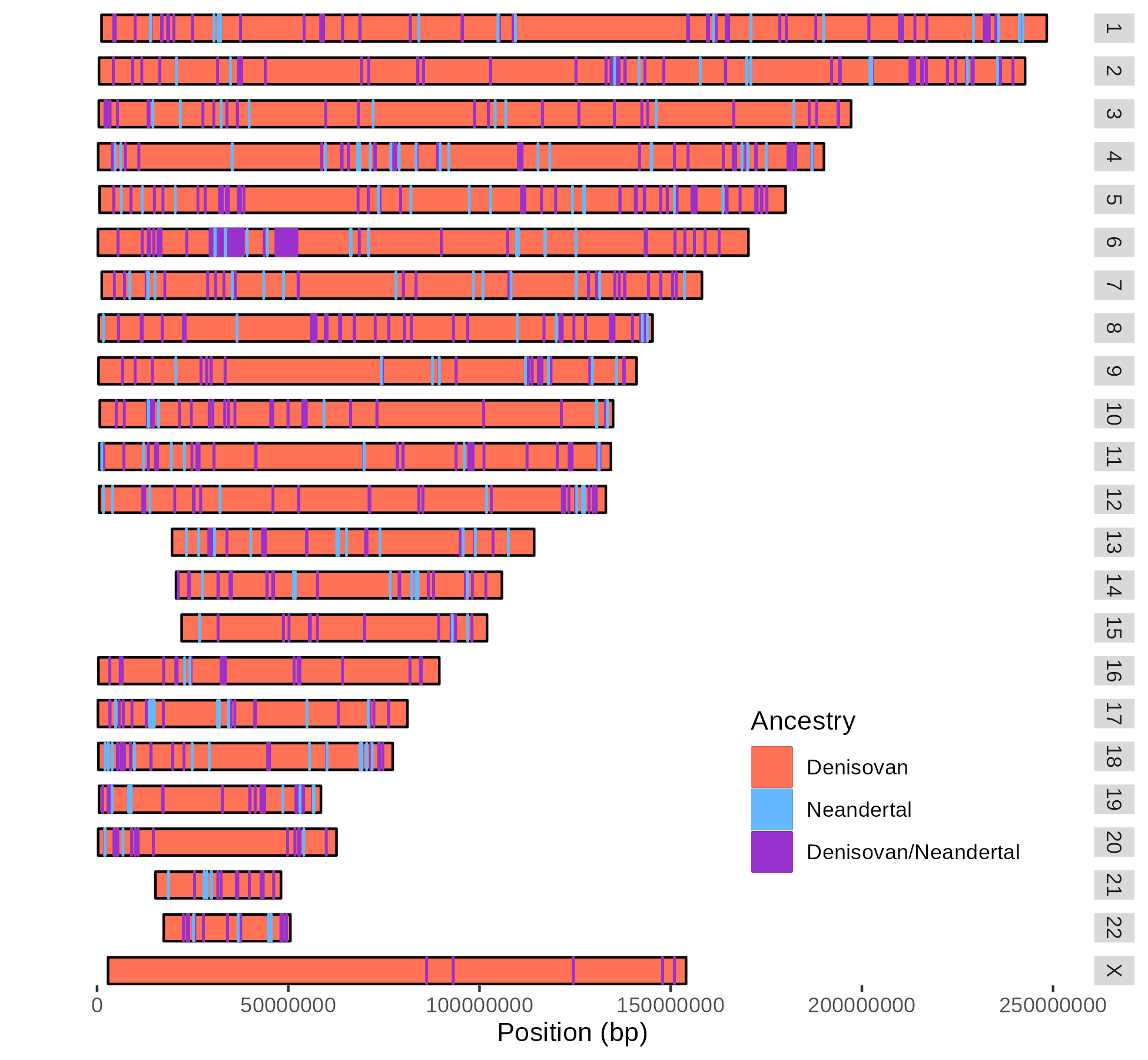

**Figure S47: Map of Neandertal ancestry in the Denisova 3 genome.** Only the Neandertal-like segments of 0.05cM and longer are represented. Window sizes were set to 0.005cM using the African-American recombination map^99^ and a penalty of 0.25 was used to call the segments.

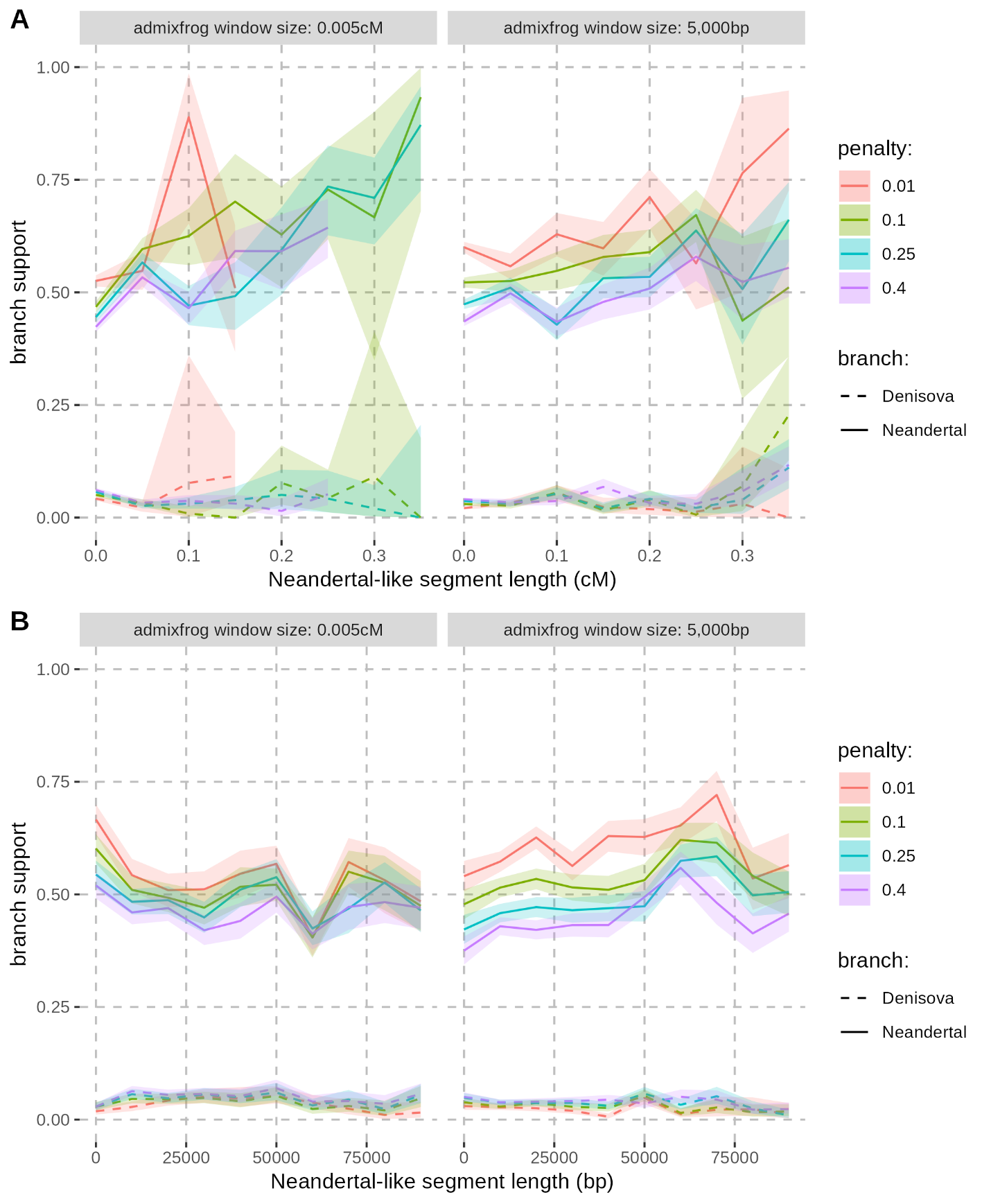

**Figure S48: Neandertal and Denisovan branch support in the homozygous Neandertal-like segments inferred by admixfrog in the Denisova 25 genome.** The branch supports are stratified by the length of segments (x-axis) either in bins of 0.05 centimorgans (**A**) or in bins of 10,000 base pairs (**B**). The shaded areas correspond to 95% binomial confidence intervals. The labels on top correspond to the window size used in admixfrog.

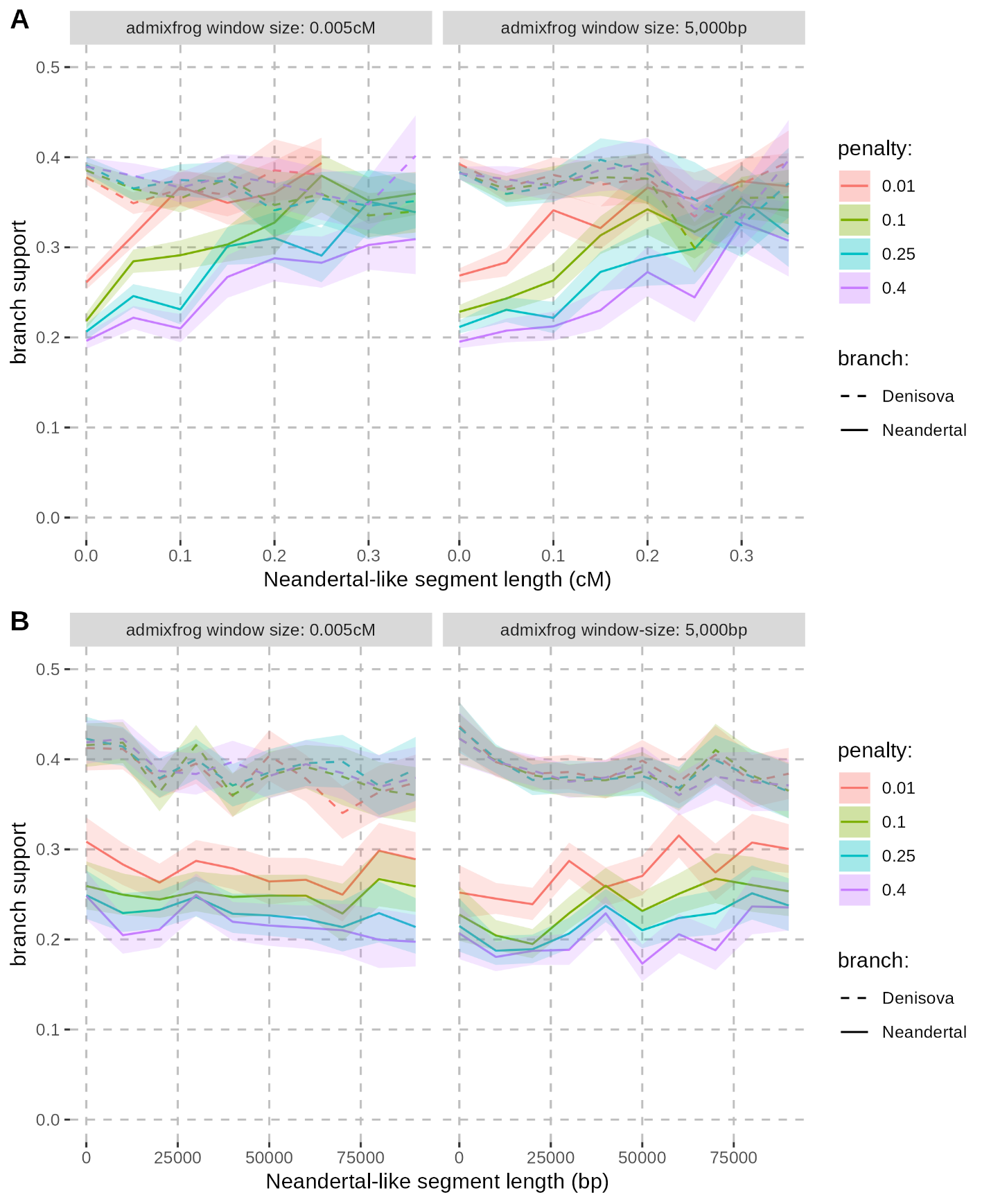

**Figure S49: Neandertal and Denisovan branch support in the heterozygous Neandertal-like segments inferred by admixfrog in the Denisova 25 genome.** The branch supports are stratified by the length of segments (x-axis) either in bins of 0.05 centimorgans (**A**) or in bins of 10,000 base pairs (**B**). The shaded areas correspond to 95% binomial confidence intervals. The labels on top correspond to the window size used in admixfrog.

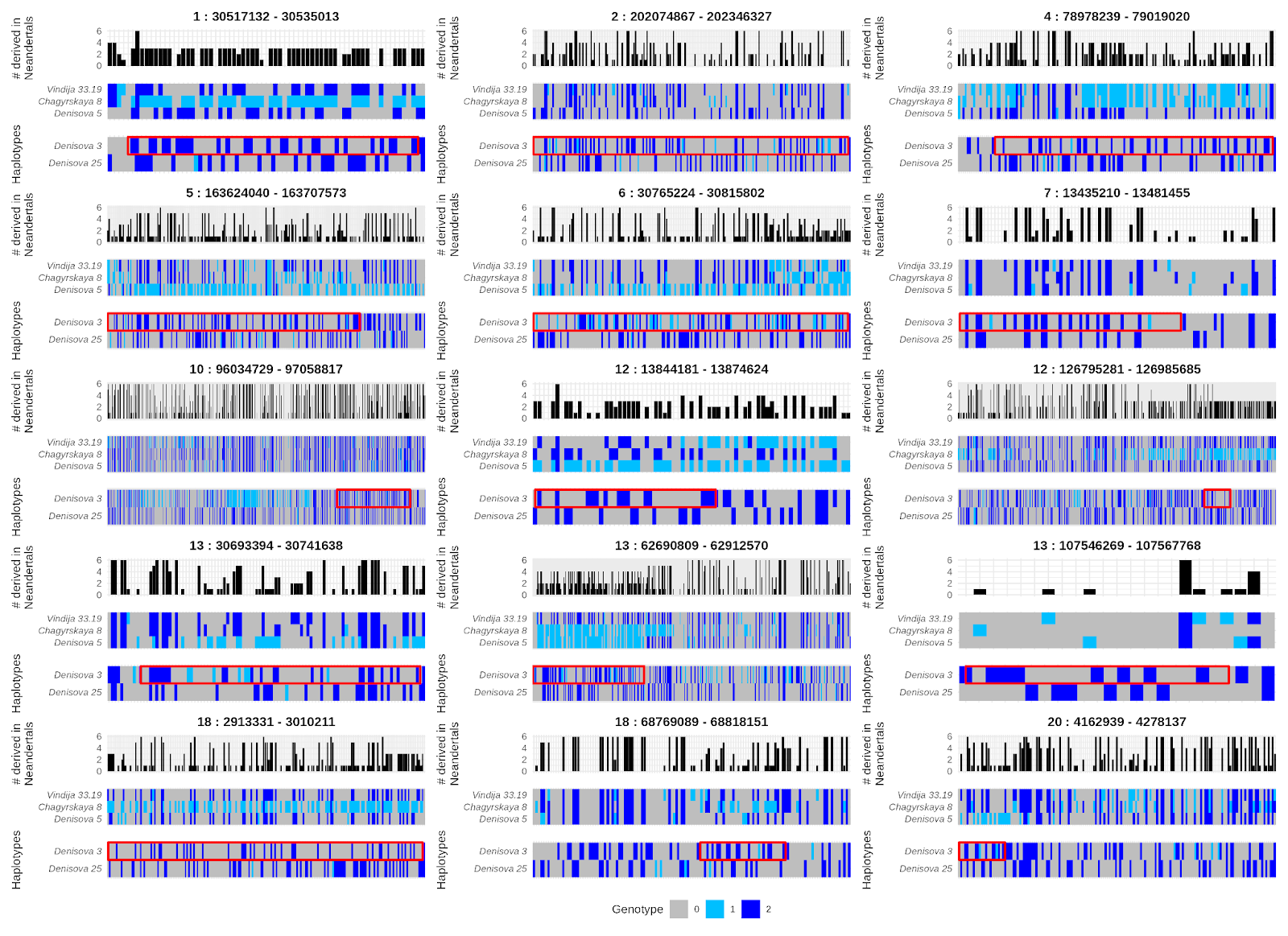

**Figure S50: Haplotype structure of homozygous Neandertal-like segments in Denisova 25 that overlap with those in Denisova 3.** Segment coordinates shown at the top of each panel correspond to the Neandertal-like segment identified in Denisova 25. Neandertal-like segments were identified with admixfrog using a window size of 0.005cM and a penalty of 0.25. The bottom of each panel displays genotypes from the high-coverage archaic genomes, with Denisovans and Neandertals separated, and different colours indicating the number of derived alleles at each position (x-axis). The segment identified in Denisova 3 is marked by a red frame, which is truncated if it extends beyond the boundaries of the corresponding segment in Denisova 25. The top of each panel illustrates the number of derived alleles among Neandertals at each position.

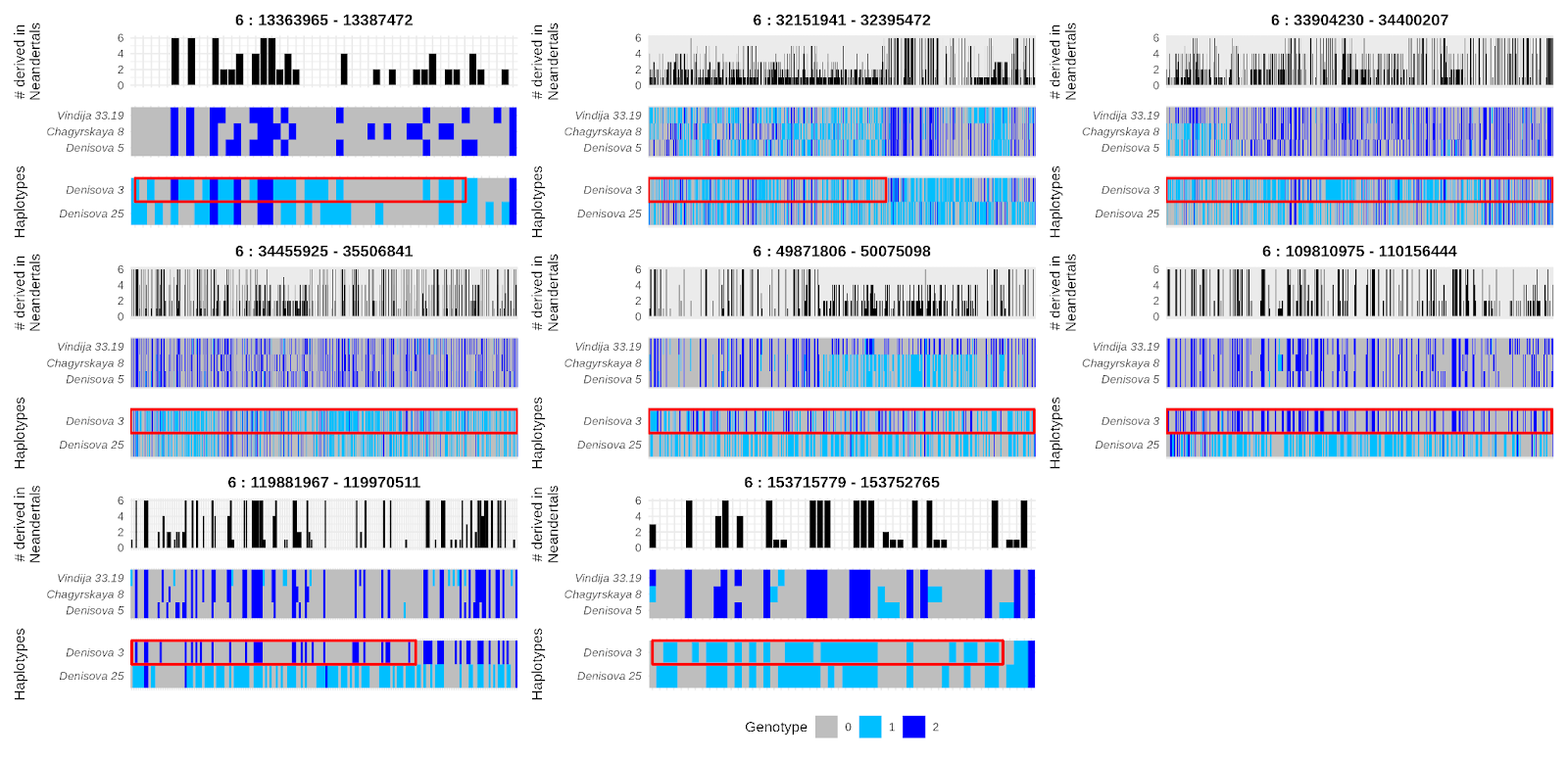

**Figure S51: Haplotype structure of heterozygous Neandertal-like segments in Denisova 25 that overlap with Neandertal-like segments in Denisova 3.** Unlike **Figure S50**, the Neandertal-like segments in Denisova 3 can also be heterozygous. Only segments identified on chromosome 6 are shown for illustration purposes. For details on the figure layout, see **Figure S50**.

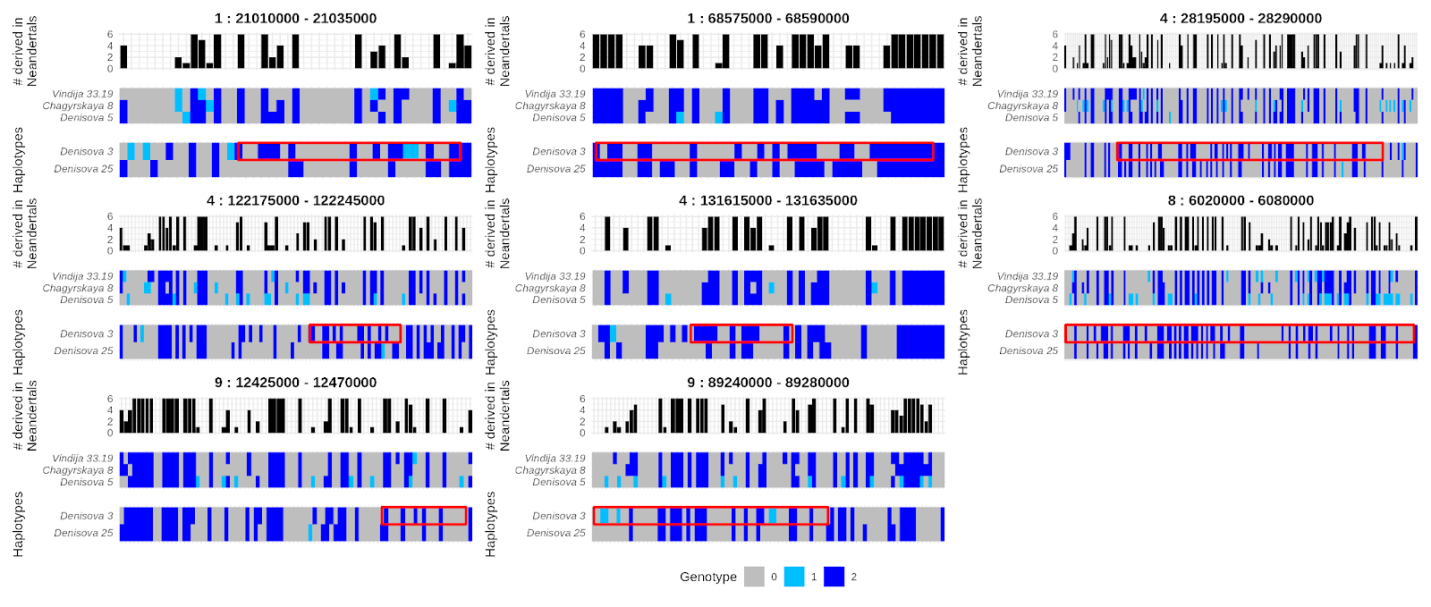

**Figure S52: Haplotype structure of homozygous Neandertal-like segments in Denisova 25 overlapping those of Denisova 3, identified by masking variable positions in Neandertals to prevent confounding with potential Denisovan-like segments in Neandertal genomes.** Unlike **Figure S50**, these Neandertal-like segments were inferred using a reference that did not include variable positions in Neandertal genomes. Furthermore, Neandertal-like segments were identified with admixfrog using a window size of 5,000bp (and a penalty of 0.25 like for **Figure S50**), as no such segments were identified with a window size of 0.05cM. For details on the figure layout, see **Figure S50**.

#### Estimates of Neandertal ancestry proportion

The *f4-ratio* statistic for estimating ancestry proportions requires reference genomes for the source populations. As an unadmixed Denisovan genome and a Neandertal genome closely related to the introgressing Neandertal population(s) are unavailable, we used two alternative approaches to estimate the proportion of Neandertal ancestry in the *Denisova 3* and *Denisova 25* genomes.

We first estimated Neandertal ancestry proportion using an algorithm implemented in admixfrog (as described in Salem *et al*. (2025)^120^). This algorithm performs random samples from the empirical-Bayes posterior probability distribution of hidden states given the observed data and the maximum likelihood estimates of the parameters. It involves drawing the initial state based on stationary probabilities and, iteratively, randomly sampling the state for the next genomic window based on the calculated transition probabilities in the hidden Markov model of *admixfrog*. This was repeated for 200 iterations, each providing different sequences of hidden states that reflect the uncertainty inherent in estimating these states based on the available data. We then counted the number of windows assigned to Neandertal-like segments divided by the total number of windows assigned to any ancestry, providing an estimate of the Neandertal ancestry proportion. We estimated Neandertal ancestry proportion separately for the autosomes and chromosome X, and excluded segments (from any ancestry) that are 0.05cM or shorter.

We estimated that *Denisova 3* and *Denisova 25* carry a little over 3% and 5% Neandertal ancestry, respectively (**Table S59**). However, these estimates are sensitive to the minimum length cutoff applied. For instance, they are substantially lower if we exclude segments that are 0.1cM or shorter, down to 1.6% and 3% for *Denisova 3* and *Denisova 25*, respectively. We therefore compare these estimates to those obtained with another approach described below. Interestingly, irrespective of the minimum length cutoff, we estimated that *Denisova 25* has about the same Neandertal ancestry proportion on the autosomes and X chromosome, whereas *Denisova 3* carries less Neandertal ancestry on the X chromosome than on the autosomes. This comparison may be subject to variation between individuals, so more genomes will be necessary to investigate these potential differences.

The other approach we used to estimate Neandertal ancestry proportion first leverages the ancestry calls from *admixfrog* to infer how the Denisovan and Neandertal components of these genomes are related to the available Neandertal genomes, and then models the genome-wide relationship of *Denisova 3* or *Denisova 25* to Neandertals as a mixture of these two components. Specifically, we measured the proportion of derived alleles one Denisovan shares with a Neandertal, at genomic positions where the other Denisovan genome and an African genome (Mbuti, HGDP00982) carry the ancestral allele. We sampled one allele randomly at heterozygous sites in each genome and defined the ancestral allele as that observed in the Chimpanzee, Bonobo, Gorilla and Orangutan reference genomes (panTro4, panpan1.1, gorGor3, ponAbe2). We first measured this statistic in the homozygous Neandertal-like segments (≥0.05cM) inferred by *admixfrog*, as well as in the regions of the genome with no evidence of Neandertal ancestry by masking all the Neandertal-like segments, including those shorter than 0.05cM. Then, we estimated the Neandertal ancestry proportion *P* by modeling the genome-wide value *S* of the statistic as a linear combination of the values estimated for the Neandertal and Denisovan ancestries (*S_Nea_* and *S_Den_*, respectively) as: *S* = *P* × *S_Nea_* + (1 - *P*) × *S_Den_*.

Because this analysis relies on the authenticity of the Neandertal-like segments called by *admixfrog*, we applied increasing minimum length cutoffs to evaluate the potential effect of including Denisovan ancestry mislabeled as Neandertal-like segments, which should be more frequent among the short segments (**Figure S53** and **Figure S54**). The estimates of the Neandertal ancestry proportion decrease with increasing segment length for both *Denisova 3* and *Denisova 25* but reach a plateau suggesting that uncertainties related to this mislabeling can be mitigated by raising the minimum segment length cutoff. Using segments of at least 0.5cM, we estimated 1.8-2.5% and 3.6-5.2% of Neandertal ancestry in the *Denisova 3* and *Denisova 25* genomes, respectively, depending on the penalty used in *admixfrog* (**Table S60**). However, these could represent underestimates if long and short Neandertal-like segments are not equally related to the Neandertal genome used. This may be true particularly for *Denisova 3* who carries some ancestry closely related to the *Denisova 5* Neandertal (Prüfer *et al*., 2017) and perhaps also Neandertal ancestry from earlier contacts with Neandertals possibly shared with *Denisova 25*.

**Table S59: Estimates of Neandertal ancestry proportion using the empirical-Bayes posterior probability sampling approach implemented in admixfrog.**

| **window size** | **penalty** | **minimum length cutoff** | **Estimated percent of Neandertal ancestry** | | | |
| --- | --- | --- | --- | --- | --- | --- |
|  |  |  | ***Denisova 3*** | | ***Denisova 25*** | |
|  |  |  | **autosomes** | **X** | **autosomes** | **X** |
| 0.005cM | 0.01 | ≥0.05cM | 3.4 [3.3-3.5] | 1.1 [0.8-1.5] | 5.3 [5.1-5.5] | 6.0 [5.0-7.0] |
|  | 0.1 |  | 3.4 [3.3-3.5] | 1.1 [0.8-1.4] | 5.3 [5.1-5.5] | 6.0 [4.9-7.1] |
|  | 0.25 |  | 3.4 [3.3-3.5] | 1.1 [0.8-1.6] | 5.3 [5.1-5.4] | 6.0 [5.1-7.0] |
|  | 0.4 |  | 3.4 [3.3-3.5] | 1.1 [0.8-1.5] | 5.3 [5.1-5.4] | 6.0 [4.9-6.8] |
|  | 0.01 | ≥0.1cM | 1.8 [1.7-1.9] | 0.1 [0.0-0.3] | 3.0 [2.9-3.2] | 3.7 [2.9-4.5] |
|  | 0.1 |  | 1.8 [1.7-1.9] | 0.1 [0.0-0.4] | 3.0 [2.9-3.2] | 3.7 [2.9-4.6] |
|  | 0.25 |  | 1.8 [1.7-1.9] | 0.1 [0.0-0.3] | 3.0 [2.9-3.2] | 3.7 [2.5-4.6] |
|  | 0.4 |  | 1.8 [1.7-1.9] | 0.1 [0.0-0.4] | 3.0 [2.9-3.2] | 3.6 [2.7-4.3] |
| 5,000bp | 0.01 | ≥50kb | 3.0 [3.0-3.1] | 1.6 [1.1-2.0] | 5.1 [5.0-5.2] | 3.3 [3.0-3.8] |
|  | 0.1 |  | 3.0 [3.0-3.1] | 1.6 [1.1-2.1] | 5.1 [5.0-5.2] | 3.3 [2.8-3.9] |
|  | 0.25 |  | 3.0 [2.9-3.1] | 1.6 [1.2-2.0] | 5.1 [5.0-5.2] | 3.3 [2.9-3.8] |
|  | 0.4 |  | 3.0 [2.9-3.1] | 1.6 [1.1-2.0] | 5.1 [5.0-5.2] | 3.3 [2.8-3.7] |
|  | 0.01 | ≥100kb | 1.6 [1.6-1.7] | 0.2 [0.1-0.5] | 3.1 [3.0-3.2] | 2.1 [1.7-2.6] |
|  | 0.1 |  | 1.6 [1.6-1.7] | 0.2 [0.1-0.5] | 3.1 [3.0-3.2] | 2.1 [1.7-2.7] |
|  | 0.25 |  | 1.6 [1.6-1.7] | 0.2 [0.1-0.5] | 3.1 [3.0-3.2] | 2.1 [1.8-2.6] |
|  | 0.4 |  | 1.6 [1.6-1.7] | 0.2 [0.1-0.5] | 3.1 [3.0-3.2] | 2.1 [1.6-2.5] |

**Table S60: Estimates of Neandertal ancestry proportion based on allele sharing with a Neandertal.** Estimates from all segments (≥0.05cM) and from long segments (≥0.5cM) are reported for different window sizes (0.005cM, defined with the African-American recombination map^99^, or 5,000bp) and different penalties. More estimates are provided in **Figure S53** and **Figure S54**.

| **window size** | **penalty** | **Estimated percent of Neandertal ancestry** | | | |
| --- | --- | --- | --- | --- | --- |
|  |  | ***Denisova 3*** | | ***Denisova 25*** | |
|  |  | **≥0.05cM** | **≥0.5cM** | **≥0.05cM** | **≥0.5cM** |
| 0.005cM | 0.01 | 1.8 | NA | 3.6 | NA |
|  | 0.1 | 2.4 | NA | 3.8 | NA |
|  | 0.25 | 2.7 | 1.8 | 4.5 | 3.6 |
|  | 0.4 | 3.1 | 1.9 | 4.9 | 3.6 |
| 5,000bp | 0.01 | 2.8 | 2.0 | 4.3 | 3.6 |
|  | 0.1 | 3.3 | 2.3 | 5.2 | 4.1 |
|  | 0.25 | 3.7 | 2.4 | 5.8 | 4.5 |
|  | 0.4 | 4.0 | 2.5 | 6.3 | 5.2 |

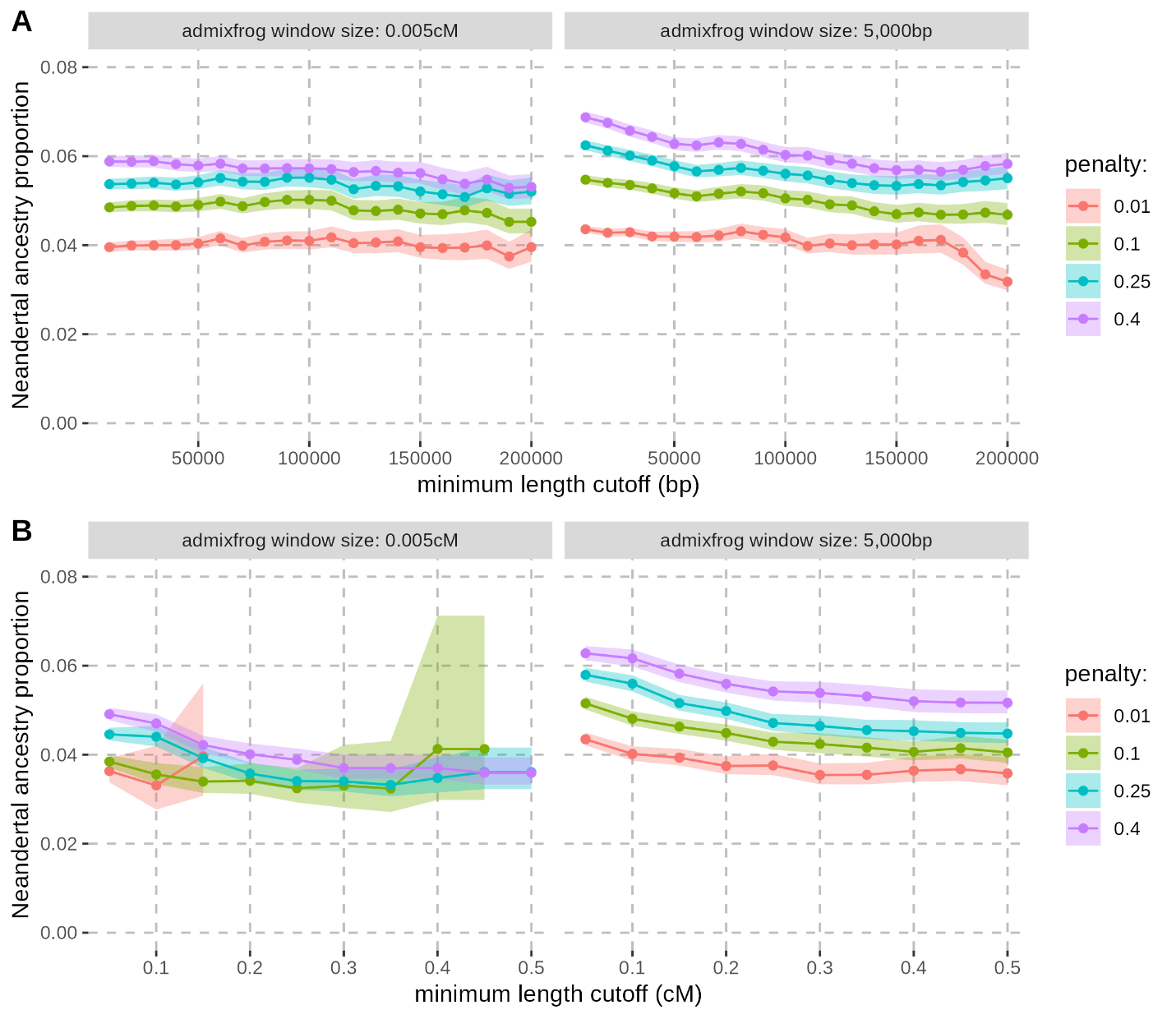

**Figure S53: Effect of the window size, penalty, and minimum segment length on the estimates of Neandertal ancestry proportion in the Denisova 25 genome.** Neandertal-like segments used for the estimates were called in windows of either 5,000bp or 0.005cM, using different penalties (colours), and filtered based on their physical length (**A**) or genetic length (**B**). Shaded areas around the curves represent the binomial 95% confidence intervals computed for each point.

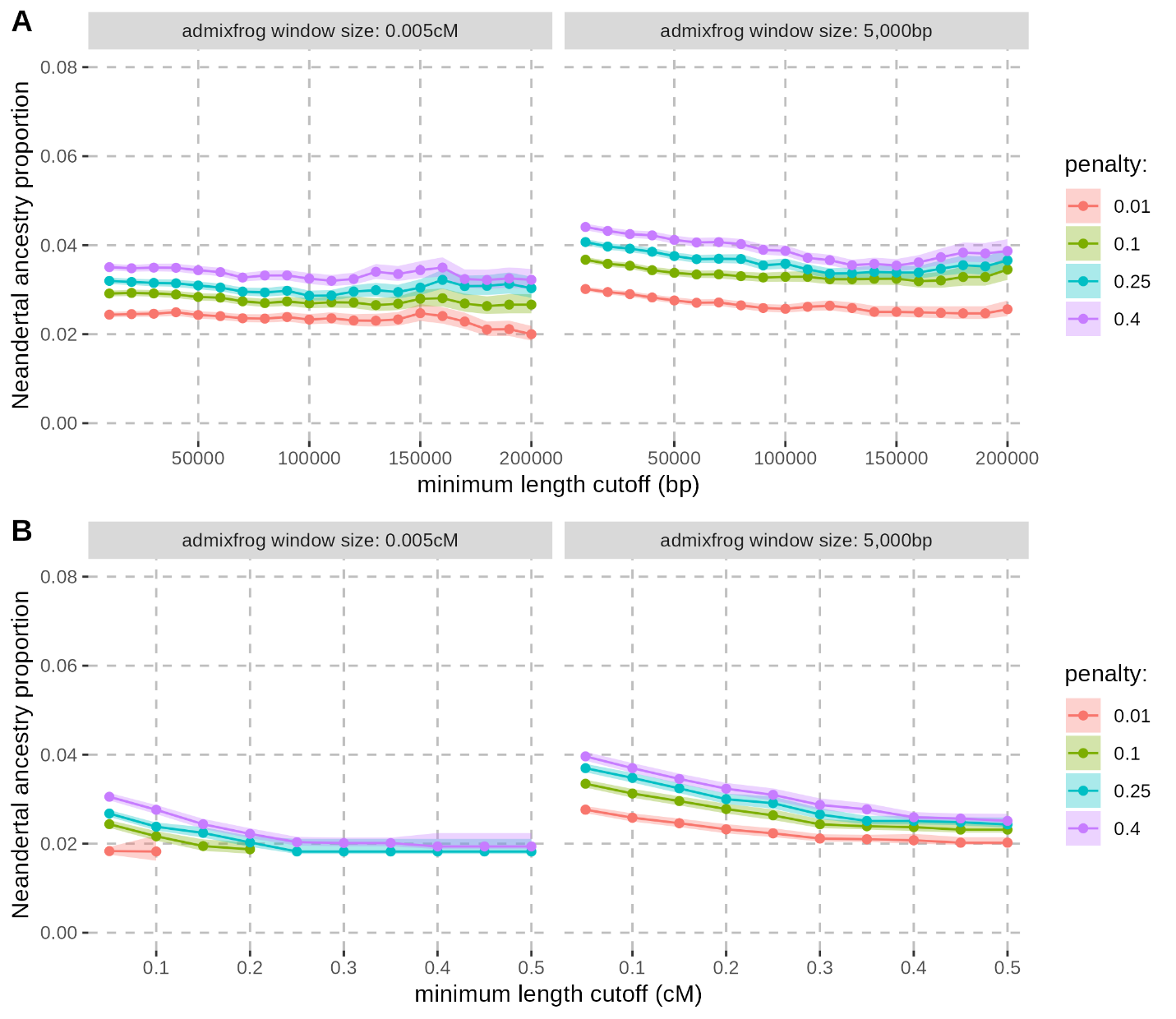

**Figure S54: Effect of the window size, penalty, and minimum segment length on the estimates of Neandertal ancestry proportion in the Denisova 3 genome.** Neandertal-like segments used for the estimates were called in windows of either 5,000bp or 0.005cM, using different penalties (colours), and filtered based on their physical length (**A**) or genetic length (**B**). Shaded areas around the curves represent the binomial 95% confidence intervals computed for each point.

#### Dating of the Neandertal admixture events

The length distribution of Neandertal-like segments detected in the genomes of *Denisova 3* and *Denisova 25* contains information about the history of contacts with Neandertals. After admixture, recombination progressively shortened the Neandertal segments. This shortening results in a length distribution often modeled by an exponential distribution with mean corresponding to the inverse of the number of generations since admixture (minus one, as the chromosomes of the first-generation offspring are not the result of recombinations between the two ancestries^121^). This relationship means that one can infer the age of the introgression from the length of introgressed segments.

We computed the complementary cumulative distribution (CCD) of the lengths of Neandertal-like segments, i.e. the proportion of segments with length equal or higher than a given length X^83^. The logarithm transformation of this cumulative distribution is expected to have a linear relationship with the segment length such as ln(CCD) ~ - (n - 1) x length, with *n* the number of generations since admixture. We thus fitted a line to the logarithm transformation of the CCD (using the lm function in R) to estimate the age of the introgression event from the slope. Yet, for both *Denisova 3* and *Denisova 25*, a line provides a poor fit to the logarithm transformation of the CCD (**Figure S55** and **Figure S56**). This suggests that our assumption of a single admixture event is likely to be invalid and that multiple events occurred. Separating long and short introgressed segments based on an arbitrary length cutoff of 0.2cM and fitting a line for each category provides a better fit to the data (**Figure S57**, **Figure S58**, **Figure S59**, and **Figure S60**). In this section, we describe how we formally tested the number of contacts with Neandertals and estimated possible ages for these contacts.

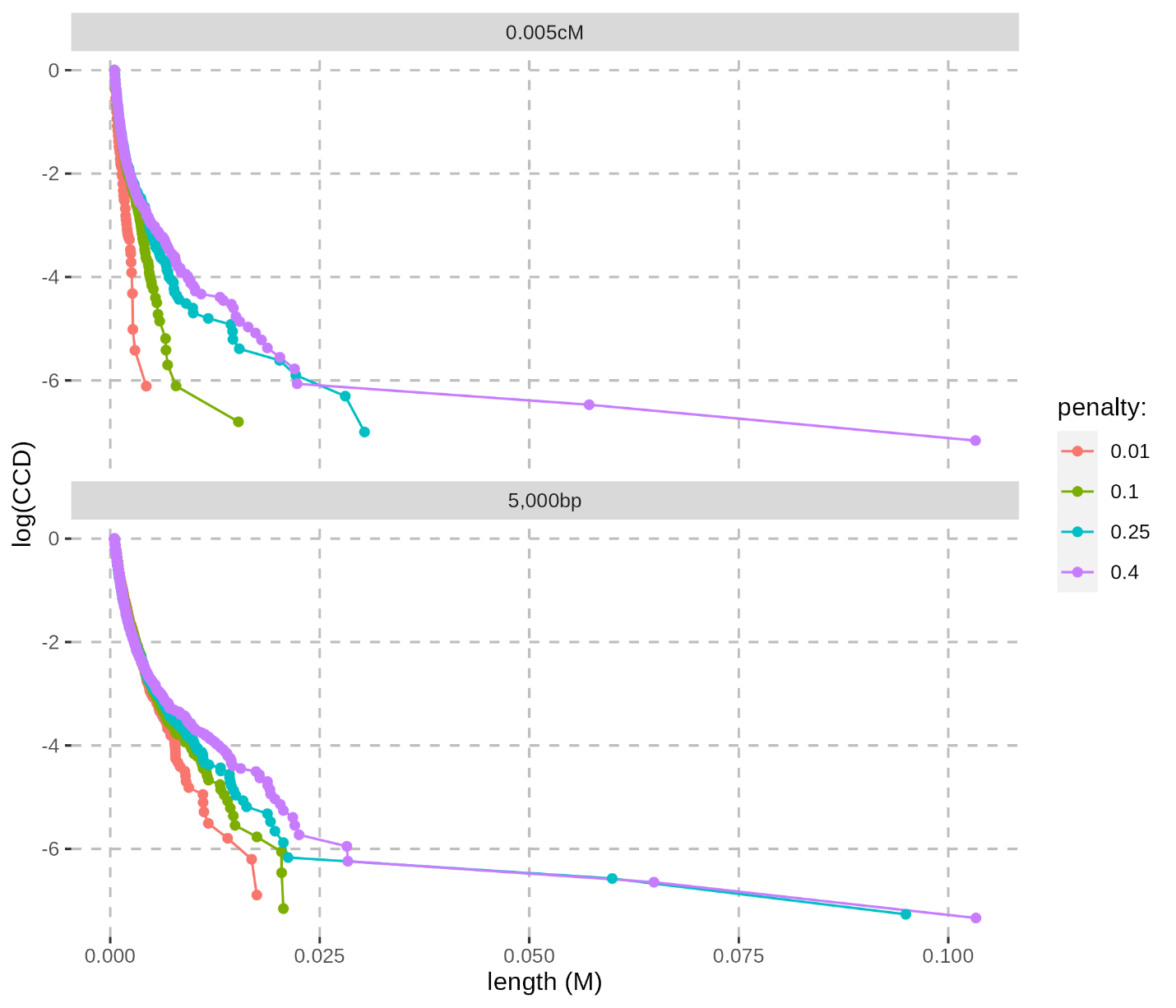

**Figure S55: Logarithm transformation of the Complementary Cumulative Distribution (CCD) of the length of the Neandertal-like segments identified with admixfrog using different window sizes and penalties in the genome of Denisova 25.** Each panel corresponds to the log(CCD) for the segments (≥0.05cM) called with admixfrog using a different window size. The length of homozygous segments was recorded only once. The colours of the distributions correspond to the different penalties. The x-axis corresponds to the genetic length of the Neandertal-like segments in Morgans.

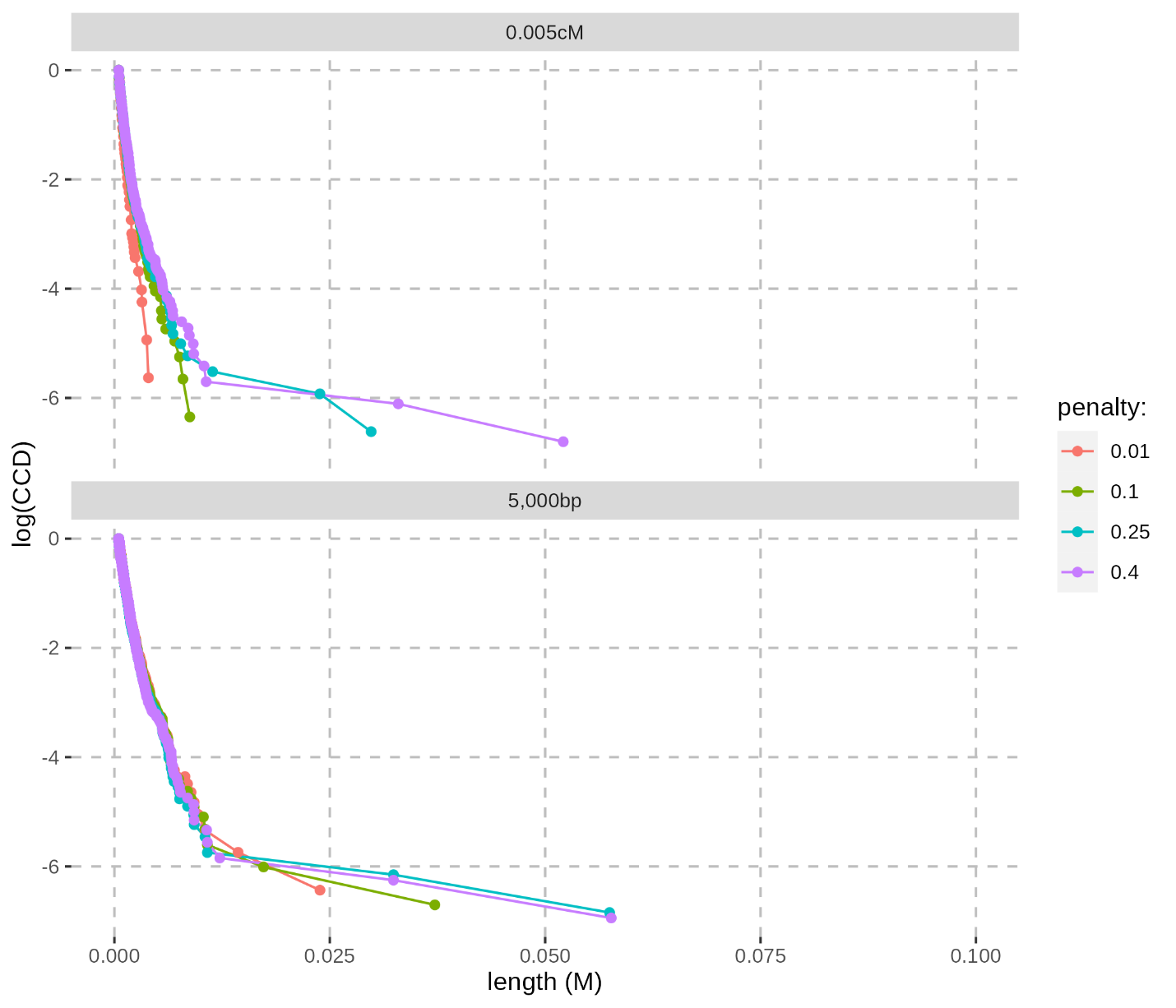

**Figure S56: Logarithm transformation of the Complementary Cumulative Distribution (CCD) of the length of the Neandertal-like segments identified with admixfrog using different window sizes and penalties in the genome of Denisova 3.** Each panel corresponds to the log(CCD) for the segments (≥0.05cM) called with admixfrog using a different window size. The length of homozygous segments was recorded only once. The colours of the distributions correspond to the different penalties. The x-axis corresponds to the genetic length of the Neandertal-like segments in Morgans and the scale is the same as in **Figure S55**, which shows the results for Denisova 25.

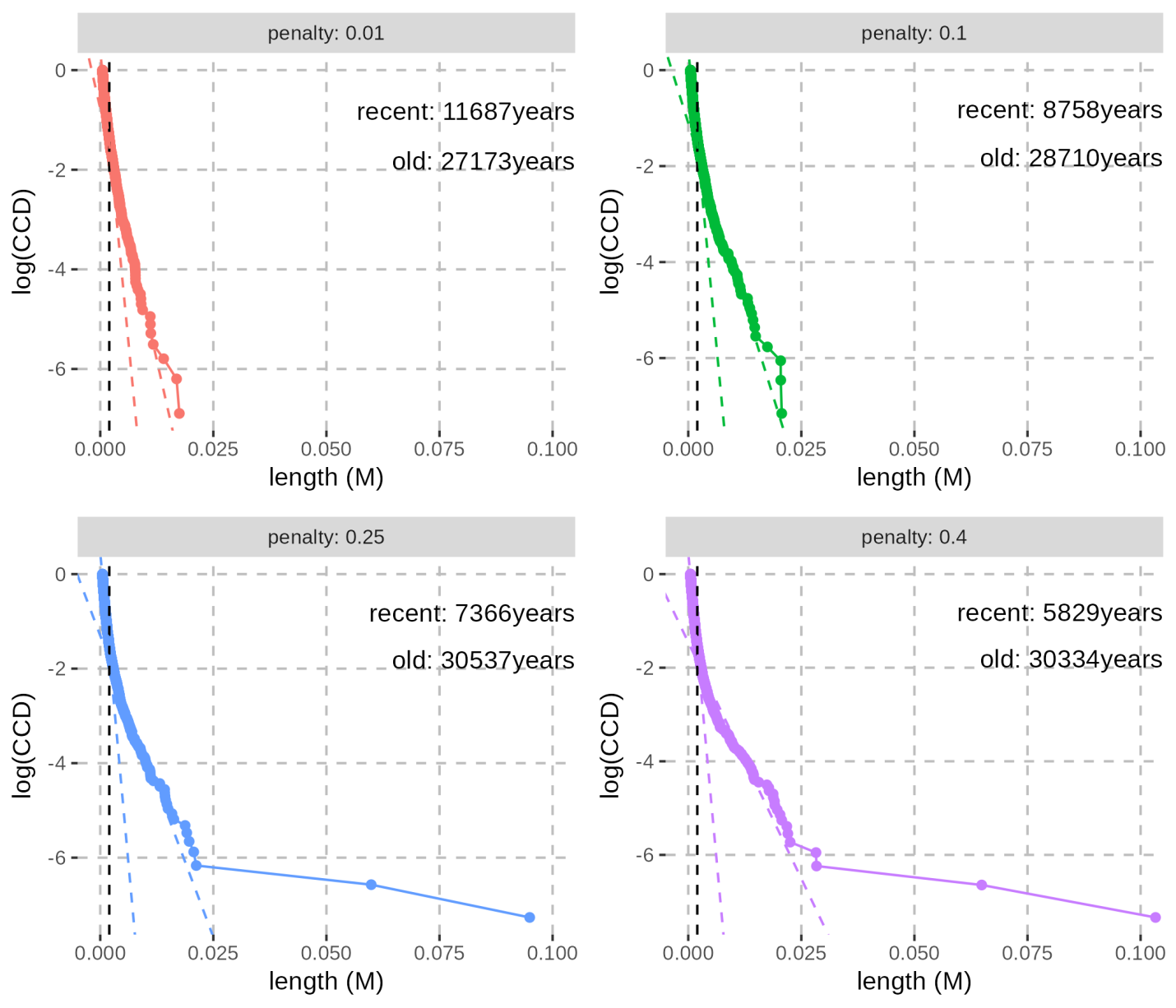

**Figure S57: Linear fits to the logarithm transformation of the Complementary Cumulative Distribution (CCD) of the length of the Neandertal-like segments identified with admixfrog in the genome of Denisova 25 using a window size of 5,000bp.** Each panel corresponds to the log(CCD) for the segments (≥0.05cM) called with admixfrog using a different penalty. The x-axis corresponds to the genetic length of the Neandertal-like segments in Morgans. Neandertal-like segments in the homozygous state were recorded once and those in the heterozygous state were randomly sampled with a probability of 0.5 to get a distribution from a pseudo-haploid genome and avoid potential bias due to the long runs of homozygosity. Slopes corresponding to the estimates of the number of years (assuming a generation time of 29 years) since a recent and an older admixture event with Neandertals, modelled as two single pulses, are shown as dashed coloured lines. The age estimates for both events are reported on the upper right corner. The old and recent events correspond to segments shorter than 0.2cM (vertical dashed line), and at least 0.2cM but no longer than 2.5cM, respectively. The maximum cutoff of 2.5cM was applied to filter out the few longest segments (up to 4) that may represent artefacts of the high penalty used in admixfrog.

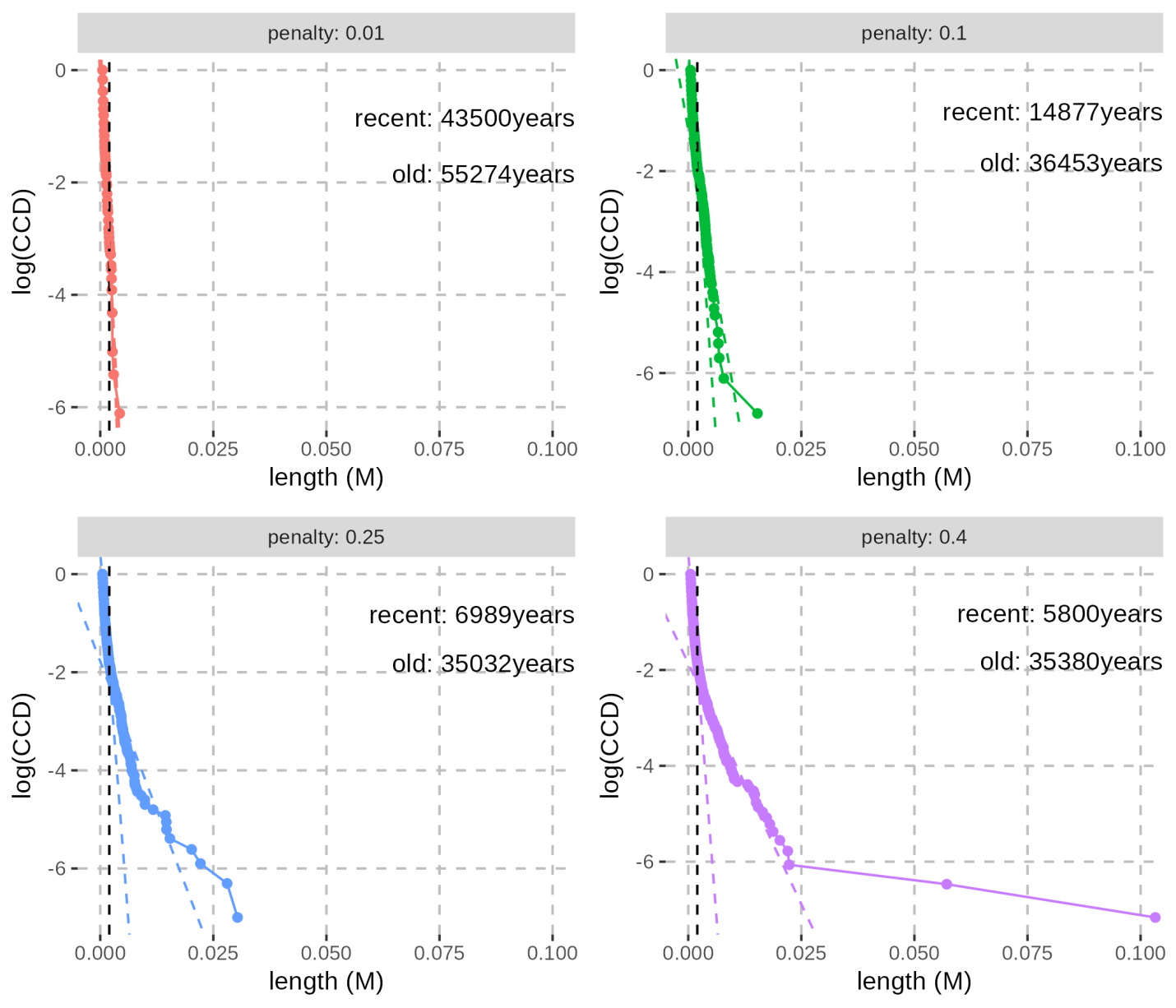

**Figure S58: Linear fits to the logarithm transformation of the Complementary Cumulative Distribution (CCD) of the length of the Neandertal-like segments identified with admixfrog in the genome of Denisova 25 using a window size of 0.005cM.** For more information, see the caption of **Figure S57**, which corresponds to the results obtained with a window site of 5,000bp instead of 0.005cM.

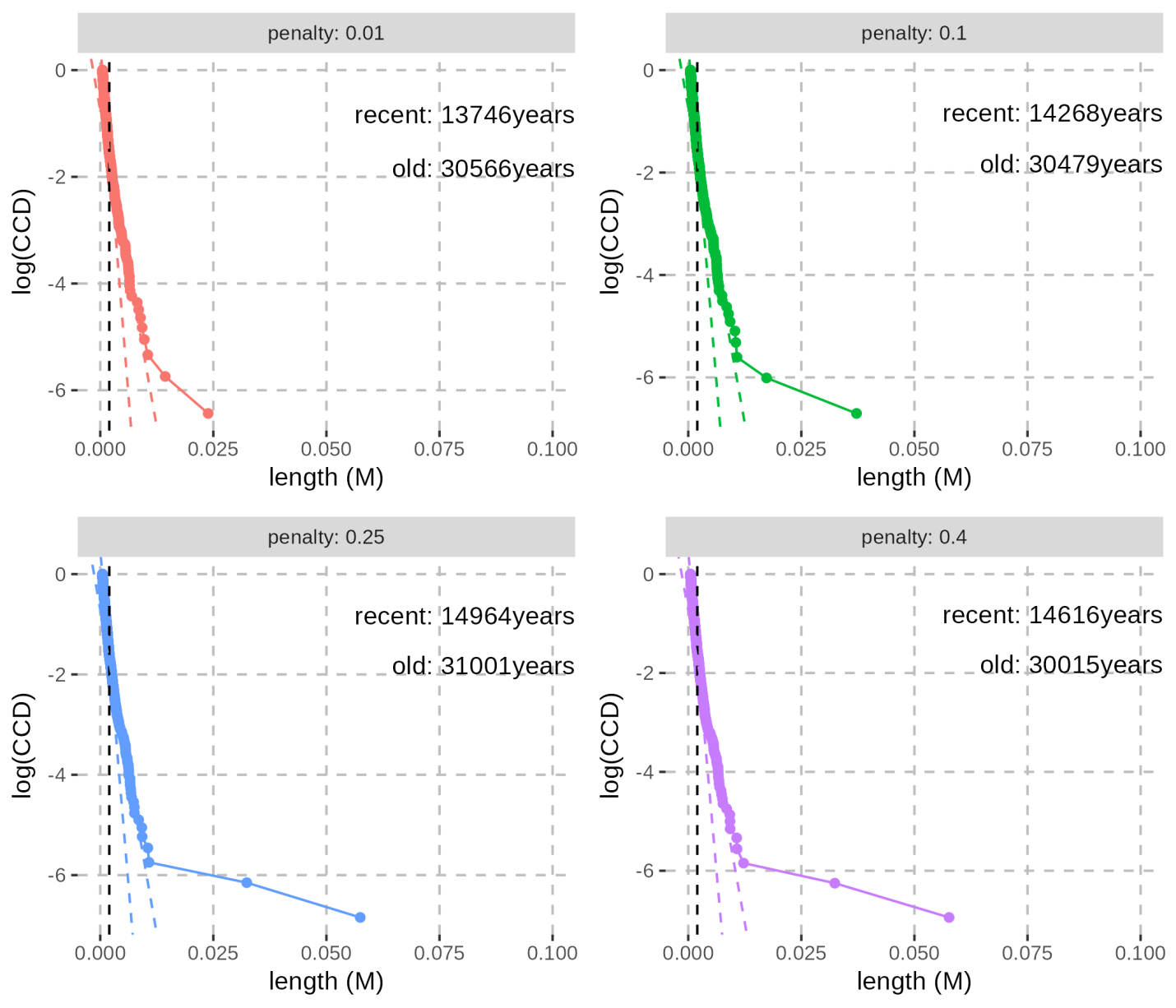

**Figure S59: Linear fits to the logarithm transformation of the Complementary Cumulative Distribution (CCD) of the length of the Neandertal-like segments identified with admixfrog in the genome of Denisova 3 using a window size of 5,000bp.** For more information, see the caption of **Figure S57**.

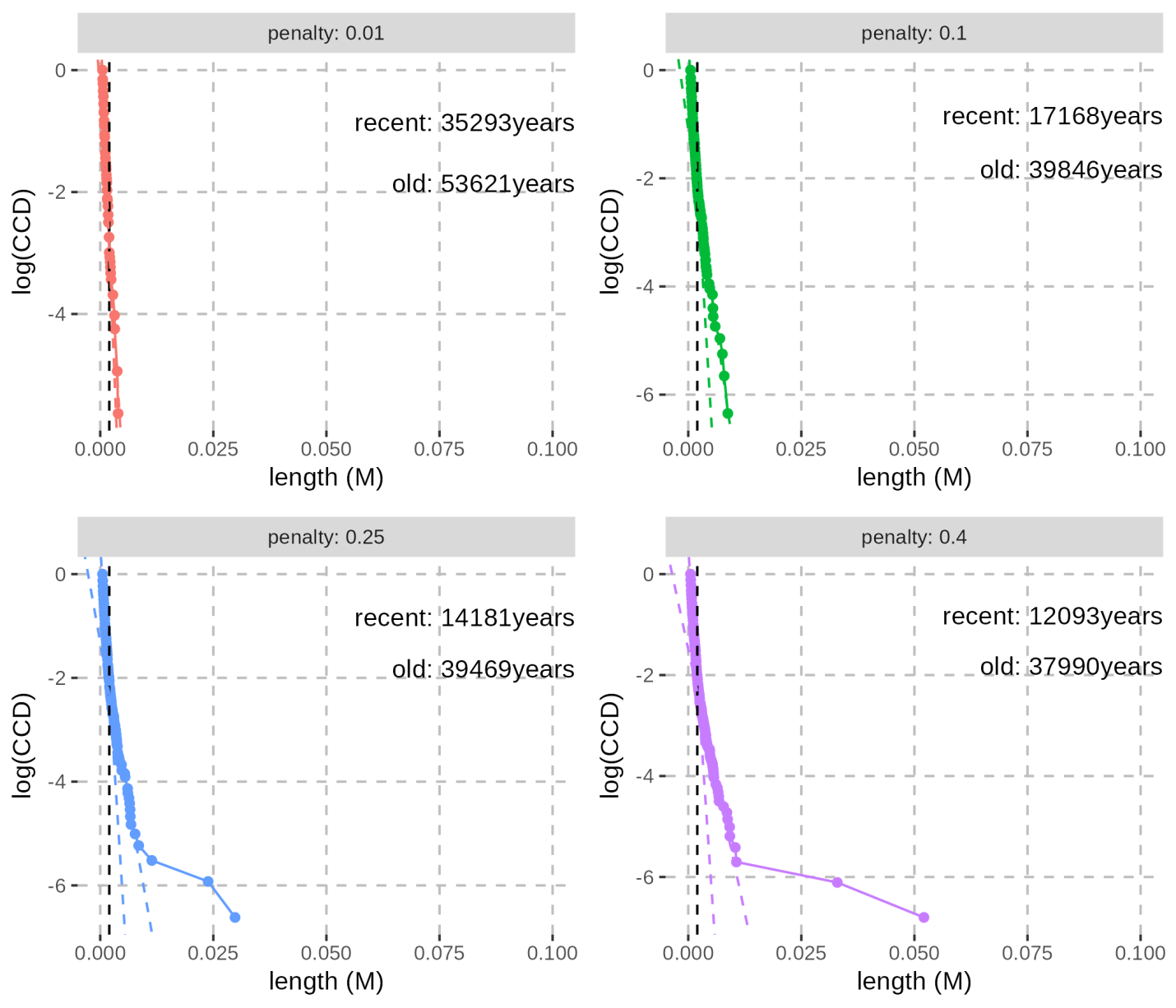

**Figure S60: Linear fits to the logarithm transformation of the Complementary Cumulative Distribution (CCD) of the length of the Neandertal-like segments identified with admixfrog in the genome of Denisova 3 using a window size of 0.005cM.** For more information, see the caption of **Figure S57**.

##### Formal test of the number of contacts with Neandertals

To test whether the ancestors of *Denisova 3* and *Denisova 25* mixed with Neandertals more than once, we computed the likelihood of the observed Neandertal-like segment lengths assuming a single exponential distribution, representing a single event of admixture, or a mixture of two exponential distributions, corresponding to two events of admixture with different ages. To avoid any bias due to the long runs of homozygosity, we computed the length distributions for a pseudo-haploid genome by recording the length of only about half of the Neandertal-like segments in the heterozygous state, sampled randomly, and the length of those in the homozygous state only once. As mentioned above, we only considered segments of at least 0.05cM. Therefore, we used left-truncated distributions to model the decay of the Neandertal-like segment length. We also excluded up to four outlier segments that were longer than 2.5cM and only observed among the calls of *admixfrog* using the highest penalties (**Figure S57** - **Figure S60**), as they might represent multiple segments incorrectly merged.

The maximum likelihood estimate of the rate of the single left-truncated exponential distribution is $\hat{\lambda}=\frac{N}{\sum_{i}^{N} L_{i}-0.0005}$ and the corresponding likelihood is $\prod_{i}^{N} \frac{{\hat{\lambda}e}^{-\hat{\lambda}L_{i}}}{e^{-\hat{\lambda}0.0005}}$, with *N* and *L_i_* correspond to the number of segments longer than 0.05cM and the length of segment *i* (in Morgan), respectively. For the mixture of two left-truncated exponential distributions, we computed the maximum likelihood estimates of the rates, *λ_1_* and *λ_2_*, and the mixture proportion, *p*, using an Expectation-Maximization algorithm. In the expectation step, the probability that segment *i* was inherited from the first or second admixture event is

$p_{1}=\frac{p{\lambda_{1}e}^{-\lambda_{1}(L_{i}-0.0005)}}{{{p\lambda}_{1}e}^{-\lambda_{1}{(L}_{i}-0.0005)}+(1-p){\lambda_{2}e}^{-\lambda_{2}(L_{i}-0.0005)}}$ or $p_{2}=\frac{(1-p){\lambda_{2}e}^{-\lambda_{2}(L_{i}-0.0005)}}{{{p\lambda}_{1}e}^{-\lambda_{1}{(L}_{i}-0.0005)}+(1-p){\lambda_{2}e}^{-\lambda_{2}(L_{i}-0.0005)}}$, respectively. In the maximization step, the parameters were updated as $\hat{\lambda}_{1}=\frac{\sum_{i}^{N} p_{1}}{\sum_{i}^{N} {(L}_{i}-0.0005)p_{1}}$, $\hat{\lambda}_{2}=\frac{\sum_{i}^{N} p_{2}}{\sum_{i}^{N} {(L}_{i}-0.0005)p_{2}}$, and $p=\frac{\sum_{i}^{N} p_{1}}{N}$, and the likelihood is $\prod_{i}^{N} {(p\hat{\lambda}_{1}e}^{-\hat{\lambda}_{1}(L_{i}-0.0005)}+{(1-p)\hat{\lambda}_{2}e}^{-\hat{\lambda}_{2}(L_{i}-0.0005)})$. We considered that the algorithm converged when the change in the log-likelihood was smaller than 10^-8^ or after 1,000 iterations. We then used a likelihood ratio test with two degrees of freedom to test whether a model with two admixture events better fits the data than a model with a single admixture event.

For both *Denisova 3* and *Denisova 25*, we found that the model of two admixture events provides a better fit to the data, irrespectively of the value of the penalty used by *admixfrog* to determine the length of Neandertal-like segments. This is also true for the length distributions computed from diploid genomes, except when we use the lowest penalty of 0.01 and a window size of 0.005cM to call segments. Maximum likelihood estimates of the parameters as well as the corresponding likelihoods are reported in **Table S61** and **Table S62**. We estimated that the ancestors of *Denisova 25* met Neandertals between 2,086 and 1,577 generations, and again between 440 and 242 generations before *Denisova 25* lived (*i.e.*, 60,494-45,733 years and 12,760-7,018 years before, assuming a generation time of 29 years). Likewise, we estimated that the ancestors of *Denisova 3* met Neandertals between 1,967 and 1,531 generations and again between 550 and 412 generations before *Denisova 3* lived (*i.e.*, 57,043-44,399 and 15,950-11,948 years before). Note that these estimates are sensitive to the parameters used to detect Neandertal ancestry, and they likely represent an average over multiple events. The Neandertal ancestry in *Denisova 25* inherited from old events (60-45 thousand years before) could be shared with *Denisova 3* (given the estimated split time between their populations 45-15 thousand years before *Denisova 25* lived, **Supplementary Note 8**). This supports that there may be more events of admixture in the history of the *Denisova 3* population than can be detected in *Denisova 3* from the length distribution. Overall, those results suggest that the populations of both *Denisova 3* and *Denisova 25* met Neandertals multiple times, and perhaps as recently as 12,000 and 7,000 years before they lived, respectively.

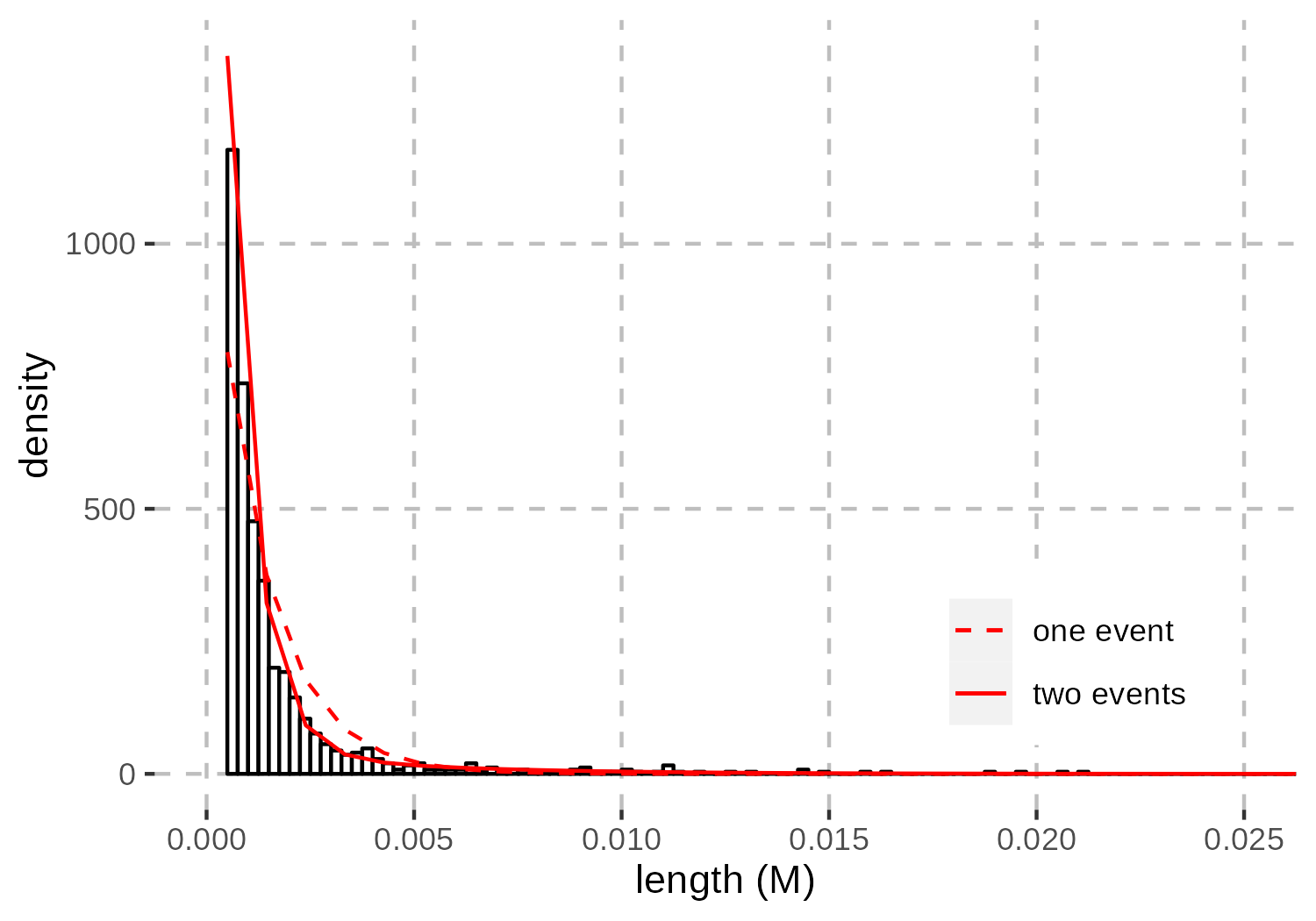

**Figure S61: Example of exponential model fitting to the length distribution of Neandertal-like segments in the genome of Denisova 25.** The histogram corresponds to the length distribution (in Morgans) of Neandertal-like segments called with admixfrog using a window size of 5,000bp and a penalty of 0.25. The red curves correspond to the left-truncated exponential distribution (dashed) or the mixture of two left-truncated exponential distributions (solid) that best fit the data. See **Table S62** for more details about the model fitting for both Denisova 3 and Denisova 25.

**Table S61: Exponential model fitting to the length distribution of Neandertal-like segments in the diploid genomes of Denisova 25 and Denisova 3.** A model with a single exponential distribution (single event) was compared to a model with a mixture of two exponential distributions (two events) with a likelihood ratio test with two degrees of freedom. The exponential distributions were left truncated at 0.0005M as this is the minimum length cutoff applied to the Neandertal-like segments called with admixfrog. The maximum likelihood estimates of the parameter values (λ_1_ and λ_2_, the rates of the exponential distributions corresponding to the recent and old events, respectively, and p, the proportion of the segments from the more recent event) of the mixture of truncated exponential distributions are reported because this model fits best to the data. The results are reported for the different window sizes and penalties used to call the Neandertal-like segments.

| individual | window size | penalty | single event log-likelihood | two events log-likelihood | p-value | *λ_1_* | *λ_2_* | *p* (%) |
| --- | --- | --- | --- | --- | --- | --- | --- | --- |
| *Denisova25* | 5,000bp | 0.04 | 8,779.426 | 9,062.618 | <10^-122^ | 220.64 | 1,534.40 | 18.2 |
|  |  | 0.25 | 8,232.771 | 8,449.207 | <10^-93^ | 285.18 | 1,710.95 | 23.9 |
|  |  | 0.1 | 7,368.985 | 7,509.826 | <10^-61^ | 326.46 | 1,634.71 | 28.2 |
|  |  | 0.01 | 5,722.101 | 5,802.033 | <10^-34^ | 439.22 | 2,013.05 | 42.5 |
|  | 0.005cM | 0.04 | 7,689.948 | 8,017.197 | <10^-142^ | 215.95 | 1,798.12 | 13.8 |
|  |  | 0.25 | 6,643.866 | 6,832.477 | <10^-81^ | 316.62 | 1,963.87 | 19.0 |
|  |  | 0.1 | 5,686.250 | 5,767.668 | <10^-35^ | 512.23 | 2,219.73 | 23.8 |
|  |  | 0.01 | 3,194.618 | 3,196.626 | 0.134 | 616.46 | 2,332.45 | 0.8 |
| *Denisova3* | 5,000bp | 0.04 | 6,409.333 | 6,449.708 | <10^-17^ | 522.08 | 1,665.23 | 32.5 |
|  |  | 0.25 | 5,729.426 | 5,765.920 | <10^-15^ | 537.40 | 1,736.59 | 34.1 |
|  |  | 0.1 | 4,889.948 | 4,930.460 | <10^-17^ | 460.05 | 1,626.62 | 29.1 |
|  |  | 0.01 | 3,672.242 | 3,711.174 | <10^-16^ | 382.04 | 1,504.39 | 22.9 |
|  | 0.005cM | 0.04 | 5,618.186 | 5,693.500 | <10^-32^ | 337.26 | 1,705.75 | 9.5 |
|  |  | 0.25 | 4,674.738 | 4,742.461 | <10^-29^ | 276.70 | 1,622.42 | 6.5 |
|  |  | 0.1 | 3,595.527 | 3,624.006 | <10^-12^ | 354.05 | 1,584.02 | 6.5 |
|  |  | 0.01 | 1,858.642 | 1,858.642 | 0.999 | 770.96 | 1,848.29 | <10^-8^ |

**Table S62: Exponential model fitting to the length distribution of Neandertal-like segments in the pseudo-haploid genomes of Denisova 25 and Denisova 3.** For more information, see the legend of **Table S61**. The only difference is that we computed the length distributions for a pseudo-haploid genome by recording the length of only half of the Neandertal-like segments in the heterozygous state, sampled randomly, and the length of those in the homozygous state only once.

| individual | window size | penalty | single event log-likelihood | two events log-likelihood | p-value | *λ_1_* | *λ_2_* | *p* (%) |
| --- | --- | --- | --- | --- | --- | --- | --- | --- |
| *Denisova25* | 5,000bp | 0.04 | 6,145.799 | 6,319.73 | <10^-75^ | 241.64 | 1,576.83 | 17.7 |
|  |  | 0.25 | 5,560.933 | 5,709.99 | <10^-64^ | 278.24 | 1,729.67 | 20.1 |
|  |  | 0.1 | 4,656.932 | 4,738.892 | <10^-35^ | 344.26 | 1,663.38 | 27.7 |
|  |  | 0.01 | 3,608.453 | 3,664.823 | <10^-24^ | 359.16 | 1,621.81 | 28.8 |
|  | 0.005cM | 0.04 | 3,540.655 | 3,597.297 | <10^-24^ | 400.35 | 1,880.99 | 35.5 |
|  |  | 0.25 | 3,547.222 | 3,601.019 | <10^-23^ | 439.85 | 2,085.57 | 40.4 |
|  |  | 0.1 | 3,555.547 | 3,612.726 | <10^-24^ | 416.79 | 1,977.21 | 38.4 |
|  |  | 0.01 | 3,669.322 | 3,744.53 | <10^-32^ | 381.06 | 2,049.95 | 35.9 |
| *Denisova3* | 5,000bp | 0.04 | 4,588.893 | 4,617.490 | <10^-12^ | 465.85 | 1,530.88 | 21.8 |
|  |  | 0.25 | 3,988.054 | 4,010.651 | <10^-9^ | 550.37 | 1,725.14 | 27.1 |
|  |  | 0.1 | 3,450.871 | 3,477.981 | <10^-11^ | 501.03 | 1,753.22 | 28.6 |
|  |  | 0.01 | 2,468.748 | 2,484.515 | <10^-6^ | 554.31 | 1,817.39 | 31.6 |
|  | 0.005cM | 0.04 | 2,512.043 | 2,526.969 | <10^-6^ | 531.93 | 1,679.64 | 32.4 |
|  |  | 0.25 | 2,417.374 | 2,433.118 | <10^-6^ | 525.76 | 1,708.35 | 31.3 |
|  |  | 0.1 | 2,312.457 | 2,338.218 | <10^-11^ | 411.86 | 1,918.12 | 9.0 |
|  |  | 0.01 | 2,476.259 | 2,497.101 | <10^-9^ | 544.81 | 1,966.65 | 32.7 |

#### Relationship to Neandertals

We assessed the relationship between the Neandertal ancestry in the Denisovan genomes and known Neandertals using *D*-statistics of the form *D*(Neandertal #1, Neandertal #2; *Denisova 3* or *Denisova 25*, ancestor), computed within the Neandertal segments identified with *admixfrog*.

As *Denisova 25* predates known Neandertal populations (given the deepest population split occurred between 145ka and 130ka^45^), his Neandertal ancestry should be equally related to these Neandertals. However, we observed that the Neandertal-like segments of *Denisova 25* are closer to *Vindija 33.19* (and *Chagyrskaya 8* in some instances) than to the *Denisova 5* Neandertal (**Figure S62**). As the method to call Neandertal-like segments relies on these genomes, a reference bias may influence this analysis. To address this, we then computed the same *D*-statistics in the segments identified with the reference comprising only the subset of sites that are monomorphic across all three Neandertal reference genomes (but differ from the other Denisovan genome). We still used all sites that differ among Neandertal genomes to compute these statistics. Although the reference with monomorphic sites in Neandertals reduces the number of Neandertal-like segments identified (compare ***Table S52*** and ***Table S57***), it ensures independence between the variants used in the *D*-statistics and those used to identify the Neandertal-like segments, circumventing reference bias. After these adjustments, we do not detect *D*-statistics significantly different from 0 for *Denisova 25*, whereas *Denisova 3* consistently exhibits a greater allele sharing with the *Denisova 5* Neandertal compared to the *Vindija 33.19* or *Chagyrskaya 8* Neandertals.

We conclude that the Neandertal ancestry in *Denisova 25* is equally related to the available Neandertal genomes and we confirm that at least some of the Neandertal ancestry in *Denisova 3* is more closely related to the *Denisova 5* Neandertal than to *Chagyrskaya 8* or *Vindija 33.19*. *Denisova 3* might also have ancestry from a deeper Neandertal lineage, as indicated by the excess of positions in homozygous Neandertal-like segments where *Denisova 3* is an outgroup to two Neandertals, compared to positions where *Denisova 3* is closer to one of them (**Table S63**). This pattern is unexpected if all of the Neandertal ancestry of *Denisova 3* were more closely related to the *Denisova 5* Neandertal. This is also consistent with the estimates in the previous section that some admixture events between Neandertals and the ancestors of *Denisova 25* predate the split between the ancestors of *Denisova 3* and *Denisova 25*.

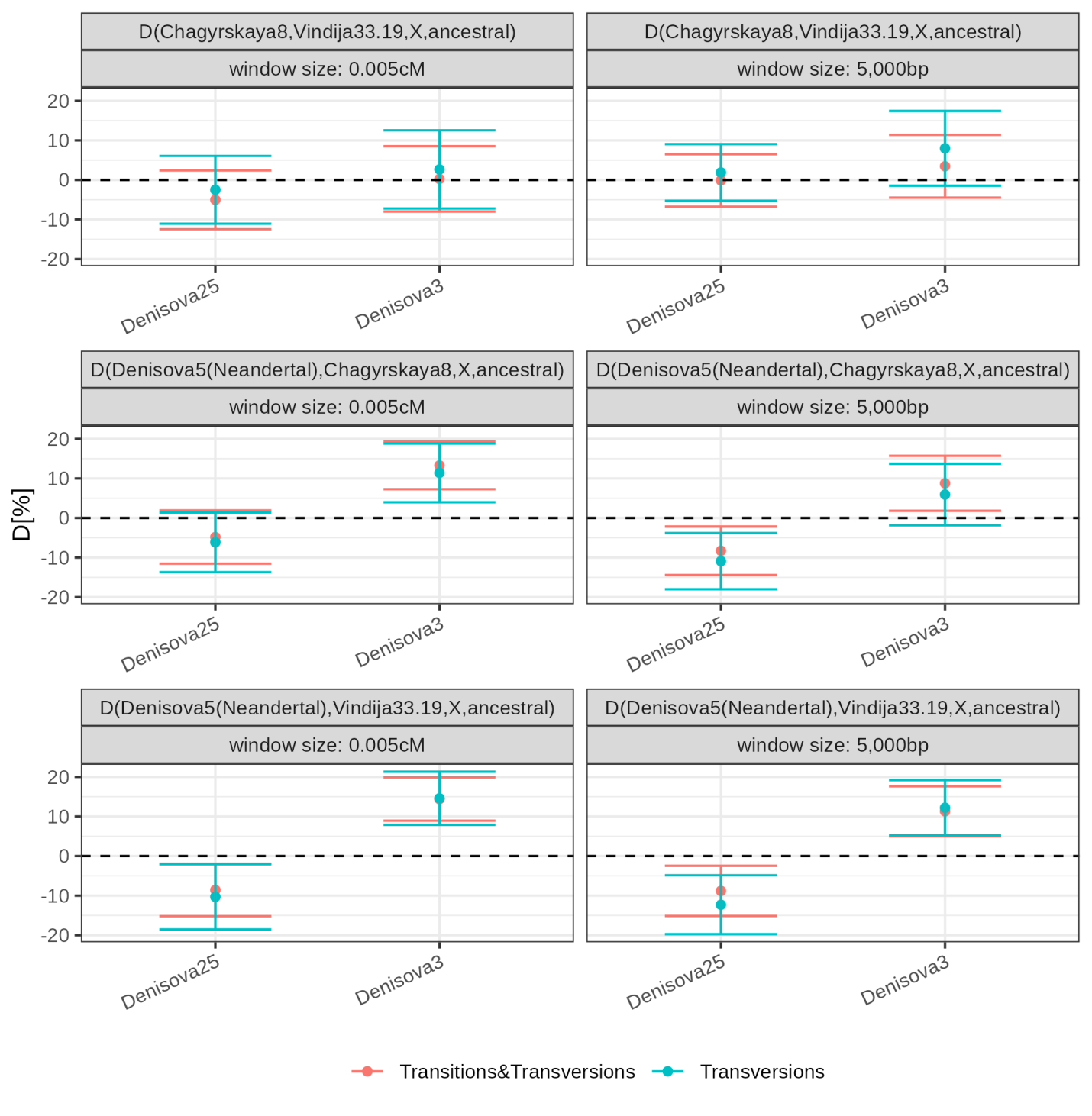

**Figure S62: Biased D-statistics comparing derived allele sharing between Neandertals and the Neandertal ancestry of Denisovans.** D-statistics were calculated using ADMIXTOOLS^108^ as implemented in admixr^109^. The genomes of panTro4, panPan2, gorGor3, and ponAbe2 were used to determine the ancestral allele. Whiskers on each side of the plotted D values correspond to two standard errors calculated using a Weighted Block Jackknife and a block size of 5Mb across all autosomes. Colours indicate whether the analysis was restricted to transversion polymorphisms or based on all polymorphisms. Panels on the left correspond to the D-statistics computed in Neandertal segments identified with admixfrog using windows of 0.005cM, whereas those on the right were computed in Neandertal segments identified using windows of 5,000bp. Only segments of at least 0.05cM were included. Note that these results are affected by reference bias, see **Figure S63** for unbiased D-statistics.

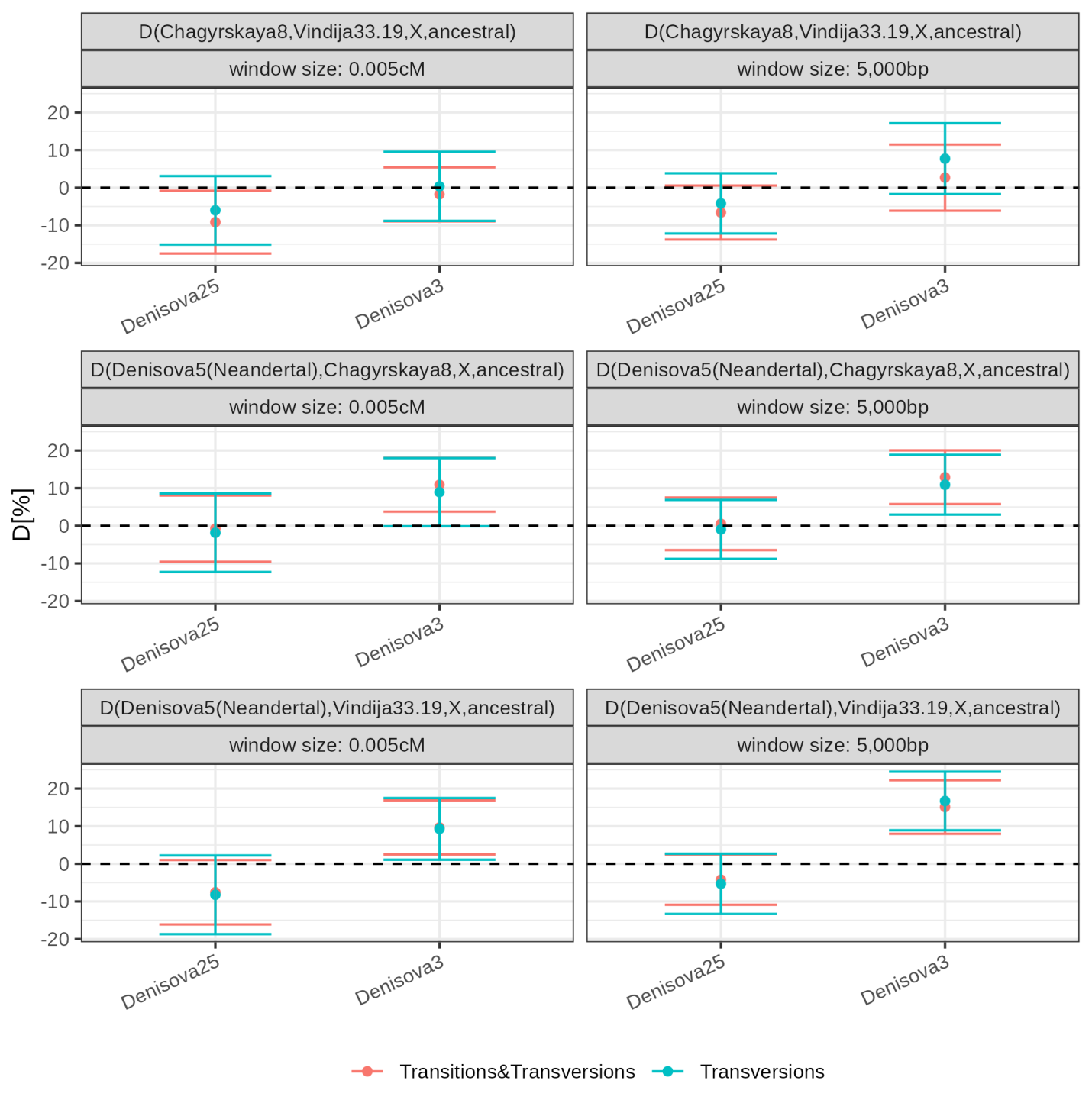

**Figure S63: Unbiased D-statistics comparing derived allele sharing between Neandertals and the Neandertal ancestry of Denisovans.** D-statistics were computed in Neandertal segments identified with admixfrog using a reference that only included monomorphic sites in the Neandertal genomes so that these D-statistics are not subject to reference bias. See the caption of **Figure S62** for more information about the layout of this figure.

**Table S63: Genetic relationship of the introgressing Neandertals to the Vindija 33.19 and Denisova 5 Neandertals.** We report the number of derived and ancestral alleles that Denisova 3 and Denisova 25 carry in genomic regions with homozygous Neandertal-like segments at positions where at least one of the Neandertal genomes (or both, for the “Neandertal” branch) carries a derived allele that is absent from a Mbuti genome and four great ape reference genomes. The numbers reported in bold correspond to the alleles that are informative about the underlying tree topology and support one of the three possible topologies relating the Neandertal ancestry in the Denisovan genomes to the Neandertal genomes. To avoid any reference bias, we used Neandertal-like segments identified with admixfrog only using sites that are monomorphic in the three Neandertal genomes (Denisova 5, Chagyrskaya 8, and Vindija 33.19). We used a window size of 5kb to call segments, and segments of at least 0.05cM.

| **Segments from:** | **penalty** | **Branch** | **derived** | **ancestral** |
| --- | --- | --- | --- | --- |
| *Denisova 3* | 0.4 | Neandertal | 4,209 | **642** |
|  |  | *Denisova 5* (Neandertal) | **357** | 722 |
|  |  | *Vindija 33.19* | **182** | 1,176 |
|  | 0.25 | Neandertal | 3,333 | **482** |
|  |  | *Denisova 5* (Neandertal) | **253** | 531 |
|  |  | *Vindija 33.19* | **135** | 895 |
|  | 0.1 | Neandertal | 2,568 | **338** |
|  |  | *Denisova 5* (Neandertal) | **161** | 385 |
|  |  | *Vindija 33.19* | **115** | 623 |
|  | 0.01 | Neandertal | 1,567 | **172** |
|  |  | *Denisova 5* (Neandertal) | **74** | 175 |
|  |  | *Vindija 33.19* | **59** | 322 |
| *Denisova 25* | 0.4 | Neandertal | 6,454 | **1,289** |
|  |  | *Denisova 5* (Neandertal) | **451** | 1,573 |
|  |  | *Vindija 33.19* | **448** | 1,990 |
|  | 0.25 | Neandertal | 4,947 | **891** |
|  |  | *Denisova 5* (Neandertal) | **360** | 1,181 |
|  |  | *Vindija 33.19* | **372** | 1,450 |
|  | 0.1 | Neandertal | 3,462 | **595** |
|  |  | *Denisova 5* (Neandertal) | **243** | 798 |
|  |  | *Vindija 33.19* | **301** | 926 |
|  | 0.01 | Neandertal | 1,700 | **272** |
|  |  | *Denisova 5* (Neandertal) | **111** | 383 |
|  |  | *Vindija 33.19* | **197** | 428 |

### Supplementary Note 10: Evidence of “Super-Archaic” ancestry

Previous comparisons of *Denisova 3* and Neandertal genomes with present-day African genomes showed that Africans share 7% more derived alleles with Neandertals than with *Denisova 3*. This difference is even larger for derived alleles fixed in Africans, as 13–16% more are shared with Neandertals than with *Denisova 3*^48^. Two non-mutually exclusive hypotheses best explain this observation: 1) Neandertals received gene flow from ancestors or relatives of modern humans, 2) Denisovans received gene flow from a “Super-Archaic” hominin whose ancestors diverged deeply from the lineage leading to Neandertals, Denisovans and present-day humans. While the gene flow from ancestors or relatives of modern human to the ancestors of Neandertals has received increasing support in recent years^103,110,111,114^, evidence for “Super-Archaic” ancestry in Denisovans remains limited by the availability of only one Denisovan genome. The presence of genomic regions where the *Denisova 3* genome is unexpectedly deeply diverged from the genomes of Neandertals and modern humans, is currently best explained by a small contribution from a “Super-Archaic” hominin into Denisovans, however, the methods that have been used give inconsistent estimates of the divergence of the introgressing “Super-Archaic” hominin from other hominins^48,103,110^. The addition of a second, older Denisovan genome could provide further insights into the existence and origin of the “Super-Archaic” ancestry in Denisovans.

#### Stratified *D*-statistics

We first aimed to assess whether “Super-Archaic” ancestry could also be present in the *Denisova 25* genome. Therefore, we repeated the *D*-statistics analysis, stratified by allele frequency in Africans, which previously showed an excess of shared derived alleles between Africans and Neandertals compared to Africans and Denisovans^48^. We computed *D*-statistics of the form *D*(Neandertal, Denisovan, derived allele in Africans, ancestral allele) in 10% frequency bins, using allele frequencies from the African genomes in the SGDP dataset (excluding North African genomes, i.e. Mozabite and Saharawi; from the VCFs described in **Supplementary Note 8**). Additionally, we computed *D*-statistics at positions with fixed derived alleles in Africans, identified by comparing six African genomes (2 Mbuti, HGDP00456 and HGDP00982, 2 Yoruba, HGDP00927 and HGDP00936, and 2 San, HGDP01029 and HGDP01036)^47,48^ to four great ape reference genomes. We ensured that the allele in these African genomes was not present in the four great apes and that at least three great apes carried a single allele that was absent from the SGDP African genomes, allowing for one missing ape allele. Finally, for both polymorphic and fixed positions, we restricted our analysis to transversion substitutions at positions passing the standard filters for both *Denisova 3* and *Denisova 25*. Standard errors were computed using a block jackknife approach in 5Mb windows^93^. We obtained consistent results with both *Denisova 3* and *Denisova 25*, regardless of the Neandertal genome used, suggesting that if “Super-Archaic” ancestry exists, it is present in both Denisovan genomes (**Figure S64**). Stratified *D*-statistics of the form *D*(Denisova 25, *Denisova 3*, derived allele in Africans, ancestral allele) comparing the relationship of *Denisova 3* and *Denisova 25* with Africans show no differences across frequency bins, including at positions with fixed derived alleles in Africans (**Figure S65**). This suggests that any “Super-Archaic” ancestry present in both Denisovans is likely at similar proportions.

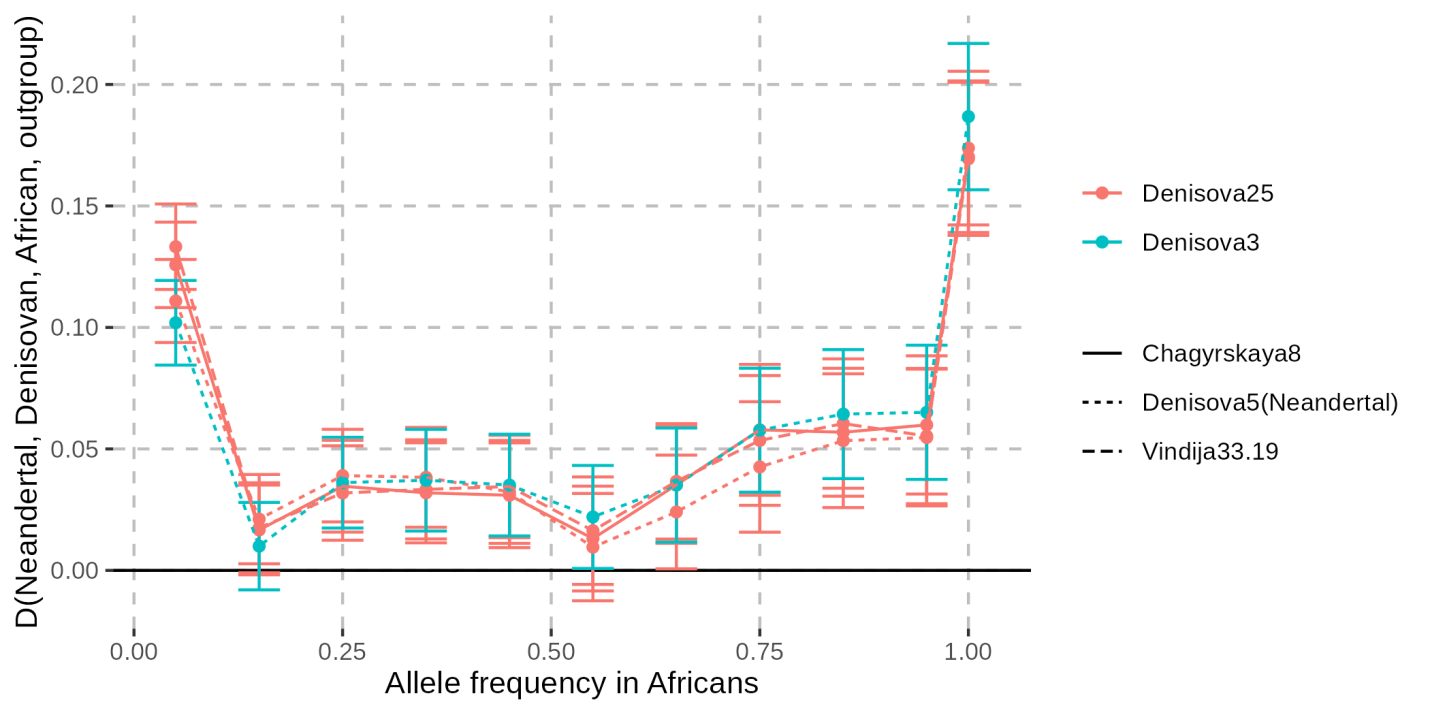

**Figure S64: Stratified *D*-statistics comparing the relationships of Neandertals and Denisovans with Africans.** A positive value indicates that the tested Neandertal is closer to Africans than the tested Denisovan. *D*-statistics were computed in frequency bins of 10%, with results plotted at the midpoint of each bin. Additionally, *D*-statistics were computed at positions where the derived allele is fixed in Africans. Only transversions were used to avoid biases observed in **Supplementary Note 8** likely due to recurrent mutations (**Figure S40**). Note that the results were not weighted by the derived allele frequency, we counted ABBA and BABA site patterns for all sites where at least one African genome carries a derived allele that is either shared with the Neandertal or Denisovan genome. Whiskers represent two standard errors from the estimate, calculated with a block jackknife approach in 5Mb windows. Colours denote different Denisovans, and line types indicate different Neandertals.

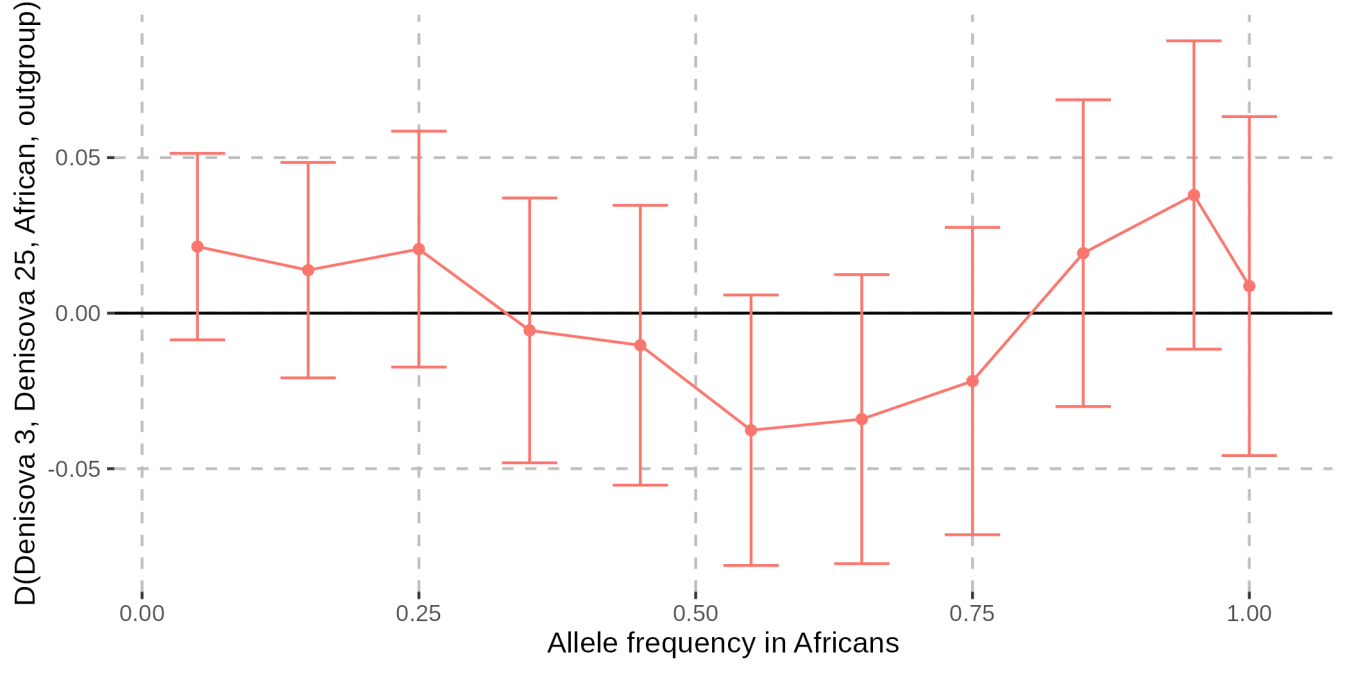

**Figure S65: Stratified *D*-statistics comparing the relationships of *Denisova 3* and *Denisova 25* with Africans.** A positive value indicates *Denisova 3* is closer to Africans than *Denisova 25*, but all estimates are not significantly different from 0. *D*-statistics were computed in 10% frequency bins and plotted at the midpoint of each bin. An additional *D*-statistic was computed at position with fixed derived alleles in Africans. Only transversions were used. Note that the results were not weighted by the derived allele frequency, we counted ABBA and BABA site patterns for all sites where at least one African genome carries a derived allele that is either shared with *Denisova 3* or *Denisova 25*. Whiskers represent two standard errors from the estimate, calculated with a block jackknife approach in 5Mb windows.

#### Evidence of “Super-Archaic” ancestry from temporal patterns in linkage disequilibrium

While stratified *D*-statistics did not reveal differences between *Denisova 3* and *Denisova 25*, any contribution from a “Super-Archaic” hominin should result in longer introgressed segments in the older genome, *Denisova 25*. To test this, without explicitly identifying segments, we compared both genomes for the genotype covariance decay between sites exhibiting alleles potentially inherited from a “Super-Archaic” source. Specifically, we focused on positions where Sub-Saharan Africans from the SGDP dataset are fixed for the derived allele and a Neandertal (*Denisova 5* or *Vindija 33.19*) and a Denisovan (*Denisova 3* or *Denisova 25*) differ, one carrying the African allele and the other the ancestral great ape allele, i.e. sites that drive the putative “Super-Archaic” signal in the stratified *D*-statistics. We measured genotype covariance (i.e. whether the Neandertal or Denisovan carries the African allele) as a function of physical distance between such site pairs. At heterozygous positions in archaic genomes, one allele was randomly sampled. For all pairs within 500kb, we computed the covariance in bins of distance *d* (rounded to the closest lower kb) as:

$cov\left( d \right)=\frac{\sum{(g}_{i}-\bar{g_{i(d)}})\times{(g}_{j}-\bar{g_{j(d)}})}{N(d)-1}$ ,

where $g_{i}$ and $g_{j}$ are the genotypes at positions *i* and *j* (included because the distance between them belongs to bin *d*), with genotypes encoded as 1 (derived) or 0 (ancestral), $\bar{g_{i(d)}}$ and $\bar{g_{j(d)}}$ are the mean genotypes over all possible positions *i* and *j* for distance bin *d*, and $N\left( d \right)$ is the number of pairs in that bin.

Using different combinations of a Denisovan and a Neandertal genome, we observed that the genotype covariance decay differed between *Denisova 3* and *Denisova 25*, but remained consistent across Neandertals (**Figure S66**). This indicates that the signal is specific to Denisovans, consistent with the presence of “Super-Archaic” ancestry in both Denisovans, and with shorter “Super-Archaic” segments in *Denisova 3* than in *Denisova 25*. Supporting this, the covariance in *Denisova 25* is larger at large distances, but lower at shorter distances, compared to *Denisova 3*.

We next explored in simulations whether the observed differences in covariance decay are indeed indicative of “Super-Archaic” ancestry, by comparing the effect of “Super-Archaic” ancestry in Denisovans and modern human ancestry in Neandertals on this covariance statistics. We performed coalescent simulations using scrm under a simplified demographic model (**Figure S67**). We assumed constant but population-specific effective sizes (1,500 for archaic and “Super-Archaic” humans, 15,000 for modern humans, 4,500 for the Neandertal-Denisovan ancestor, and 15,000 for the modern-archaic ancestor), with split times of 400ka (Neandertal-Denisovan), 600ka (modern-archaic), and 6Ma (chimpanzee). We first simulated an admixture event at 250ka from a “Super-Archaic” population (split at 1Ma) into Denisovans, contributing 0%, 1% or 10% of their ancestry. For comparison, we also simulated 10% gene flow from modern humans into Neandertals at 250ka, without “Super-Archaic” ancestry in Denisovans. We sampled individuals at different time points to reflect the ages of the archaic genomes. For simplicity, we assumed the older individuals belonged to the same population as the younger ones, simulating single Neandertal and Denisovan populations. We simulated 1,000 loci of 1Mb each, using a mutation rate of 1.45x10^-8^ per bp per generation, a recombination rate of 1cM/Mb, and a generation time of 25 years. For each individual, we sampled two chromosomes, with one allele randomly sampled at heterozygous sites to mimic empirical data processing. We then computed the genotype covariance as described above. Only the simulations with “Super-Archaic” gene flow produced covariance differences between Denisovans similar to those observed in the real data. In contrast, simulations with modern human gene flow into Neandertals elevated the overall covariance but did not produce covariance differences between the two Denisovan individuals (**Figure S68**). These results suggest that the covariance is more sensitive to “Super-Archaic” ancestry in Denisovans than to modern human ancestry in Neandertals. Although the two Neandertals also differ in age, we do not observe differences in the covariance decay when substituting one for the other, perhaps because modern human introgression into Neandertals introduced relatively few derived alleles not shared with Denisovans, whereas “Super-Archaic” ancestry in Denisovans, due to its greater divergence, introduced a larger number of ancestral alleles that disproportionately affect the covariance signal.

We next investigated how the divergence time of the “Super-Archaic” population and the time of admixture affect the covariance decay. Using simulations, we compared scenarios with divergence times of 1Ma, 1.5Ma, or 2Ma, and admixture events occurring at 250ka, 300ka, or 350ka. As expected, greater divergence led to larger differences in the covariance decay between Denisovans of different ages (**Figure S69**), while older admixture events resulted in smaller differences, consistent with increased recombination breaking down linkage over time (**Figure S70**).

We observed that the decay of covariance in the real data extends over larger genomic distances than in our simulations. Interestingly, covariance is higher in regions with low B-scores (**Figure S71**), a proxy for reduced genetic diversity in modern humans associated with stronger background selection. These low B-score regions also tend to have lower recombination rates, which may explain the slower decay in real data compared to our simulations, where recombination was modeled as uniform. It remains unclear whether the signal is genuinely enriched in low B-score regions or simply more detectable due to reduced recombination limiting the breakdown of linkage.

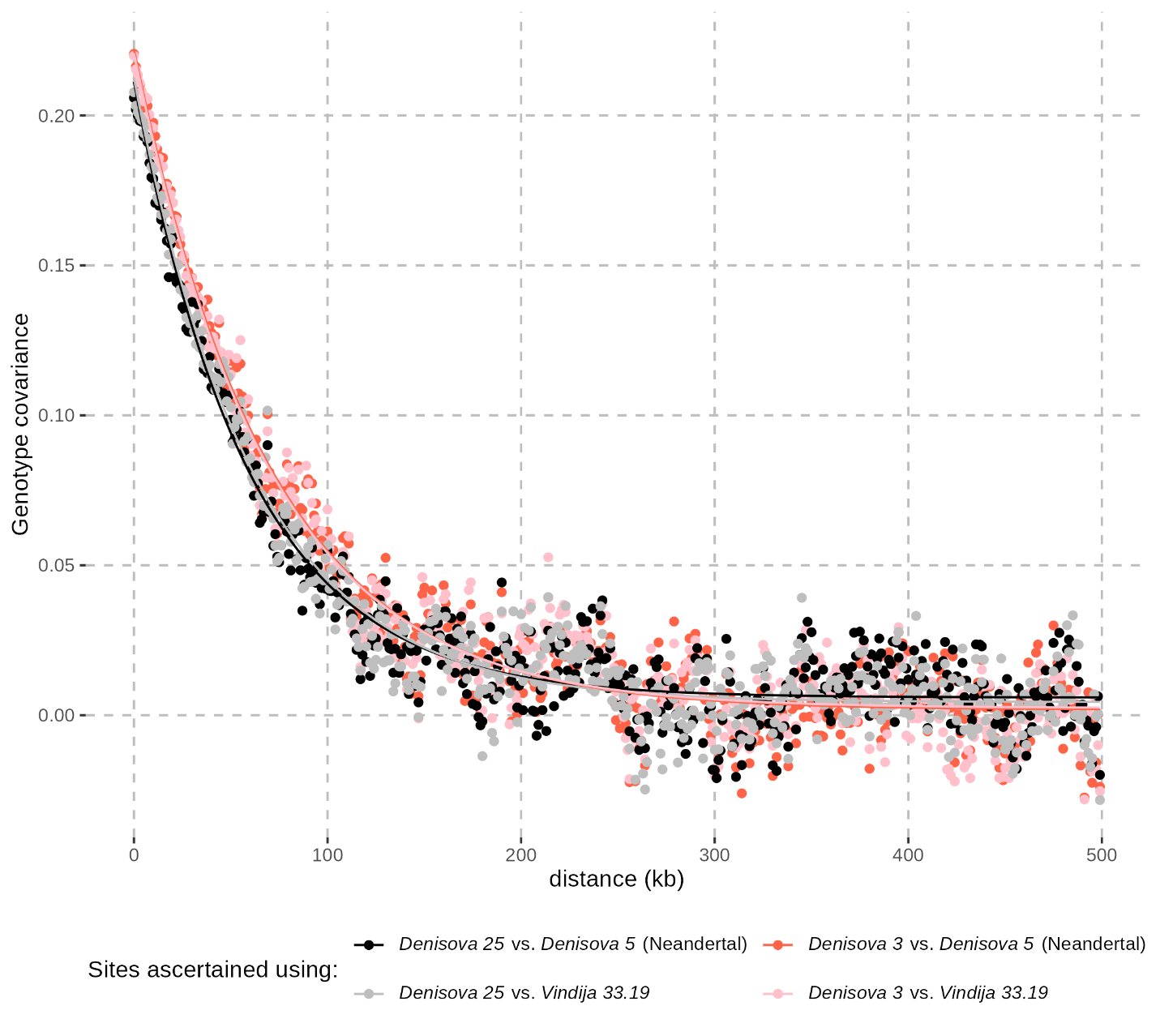

**Figure S66: Decay of genotype covariance as a function of distance between pairs of sites with alleles potentially inherited from a “Super-Archaic” hominin.** Covariance was computed based on whether the Neandertal or Denisovan carries the derived allele fixed in SGDP Sub-Saharan Africans at both sites. At heterozygous sites, one allele was randomly sampled. Each point represents the genome-wide average covariance for pairs of sites binned by distance (rounded down to the nearest kb, x-axis). Curves show exponential fits estimated using the *nls* function in R (model: a × exp(–b × distance) + c). Colours indicate the genome pairs used: grey/black for *Denisova 25* with either the *Denisova 5* or *Vindija 33.19* Neandertal, red/pink for *Denisova 3* with either the *Denisova 5* or *Vindija 33.19* Neandertal.

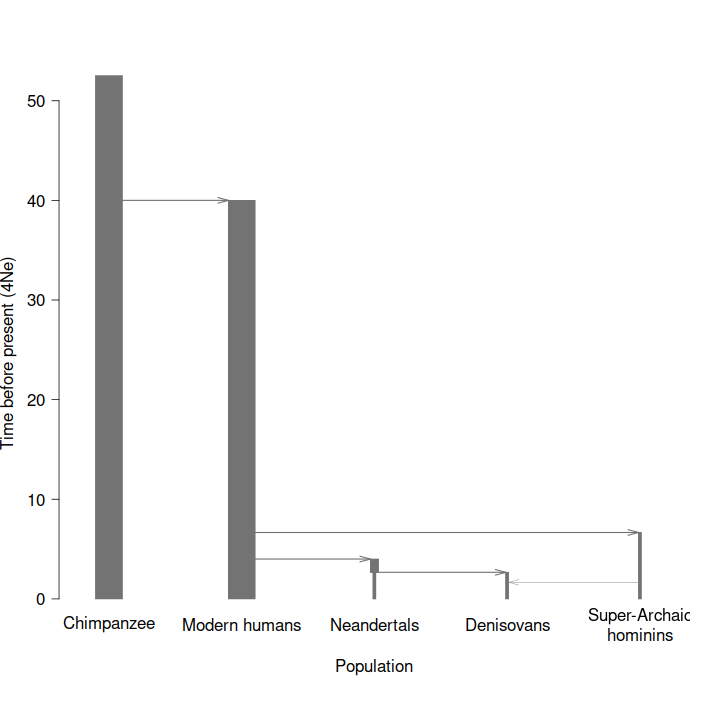

**Figure S67: Tree summarising an example of simulated demographic history.** This scenario includes 1% gene flow from a “Super-Archaic” hominin population into Denisovans, and was reconstructed from a *scrm* command using *POPdemog* (Zhou *et al.*, 2018). Times (generations) are shown in units of 4Nₑ, where Nₑ is the effective population size of Neandertals and Denisovans, assumed to be 1,500 individuals. Across simulations, we varied the divergence time between “Super-Archaic” and other hominins, the timing of admixture, and the proportion of gene flow. In a separate set of simulations, we included gene flow from modern humans into Neandertals instead.

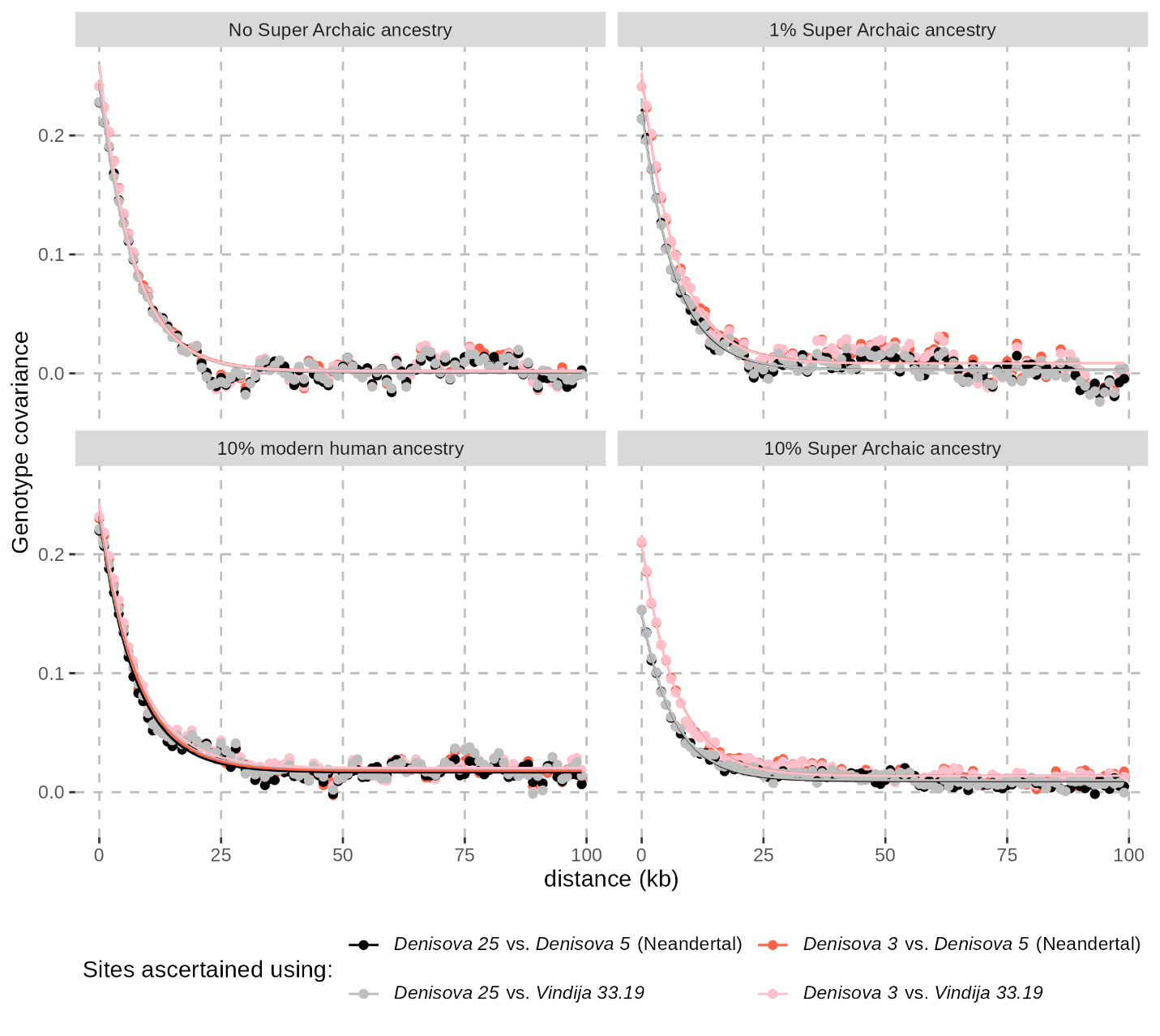

**Figure S68: Decay of genotype covariance in simulations with varying levels of “Super-Archaic” ancestry in Denisovans or modern human ancestry in Neandertal**s. Covariance was computed as for the empirical data (see **Figure S66** for more details), including random sampling of one allele at heterozygous sites. Each point shows the average covariance for pairs of sites binned by distance (rounded down to the nearest kb, x-axis), based on 1,000 simulated loci of 1Mb each. These simulations include gene flow at 250ka, either into Denisovans from a “Super-Archaic” hominin population that diverged 1Ma, or from modern humans into Neandertals. See **Figure S66** for more details about the figure layout, and **Figure S67** and the text above for more details about the simulations.

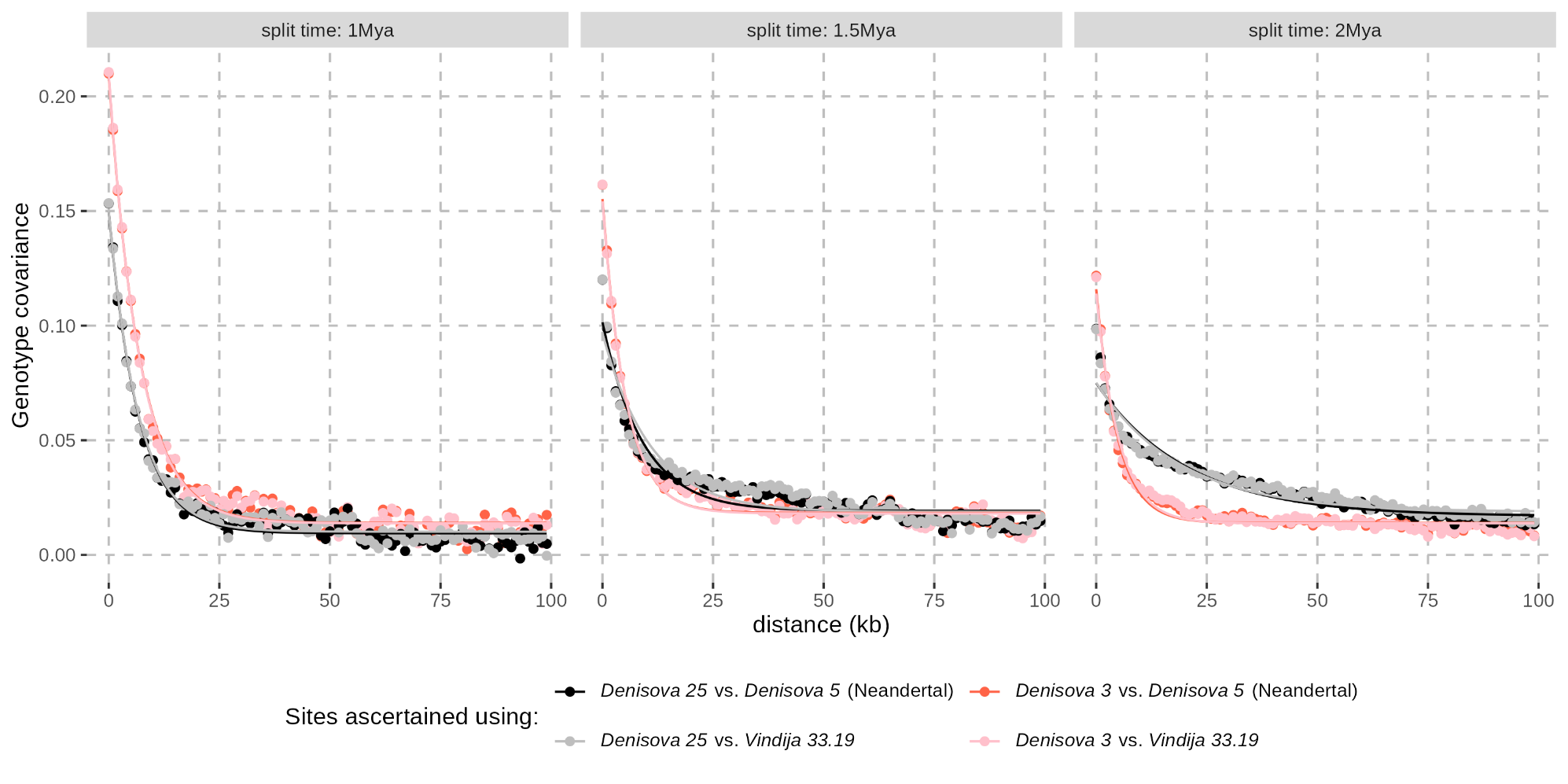

**Figure S69: Decay of genotype covariance in simulations with varying divergence between a “Super-Archaic” hominin and other humans.** Covariance was computed as for the empirical data (see **Figure S66** for more details), including random sampling of one allele at heterozygous sites. Each point shows the average covariance for pairs of sites binned by distance (rounded down to the nearest kb, x-axis), based on 1,000 simulated loci of 1Mb each. These simulations include 10% gene flow from a “Super-Archaic” hominin population into Denisovans at 250ka. See **Figure S66** for more details about the figure layout, and **Figure S67** and the text above for more details about the simulations.

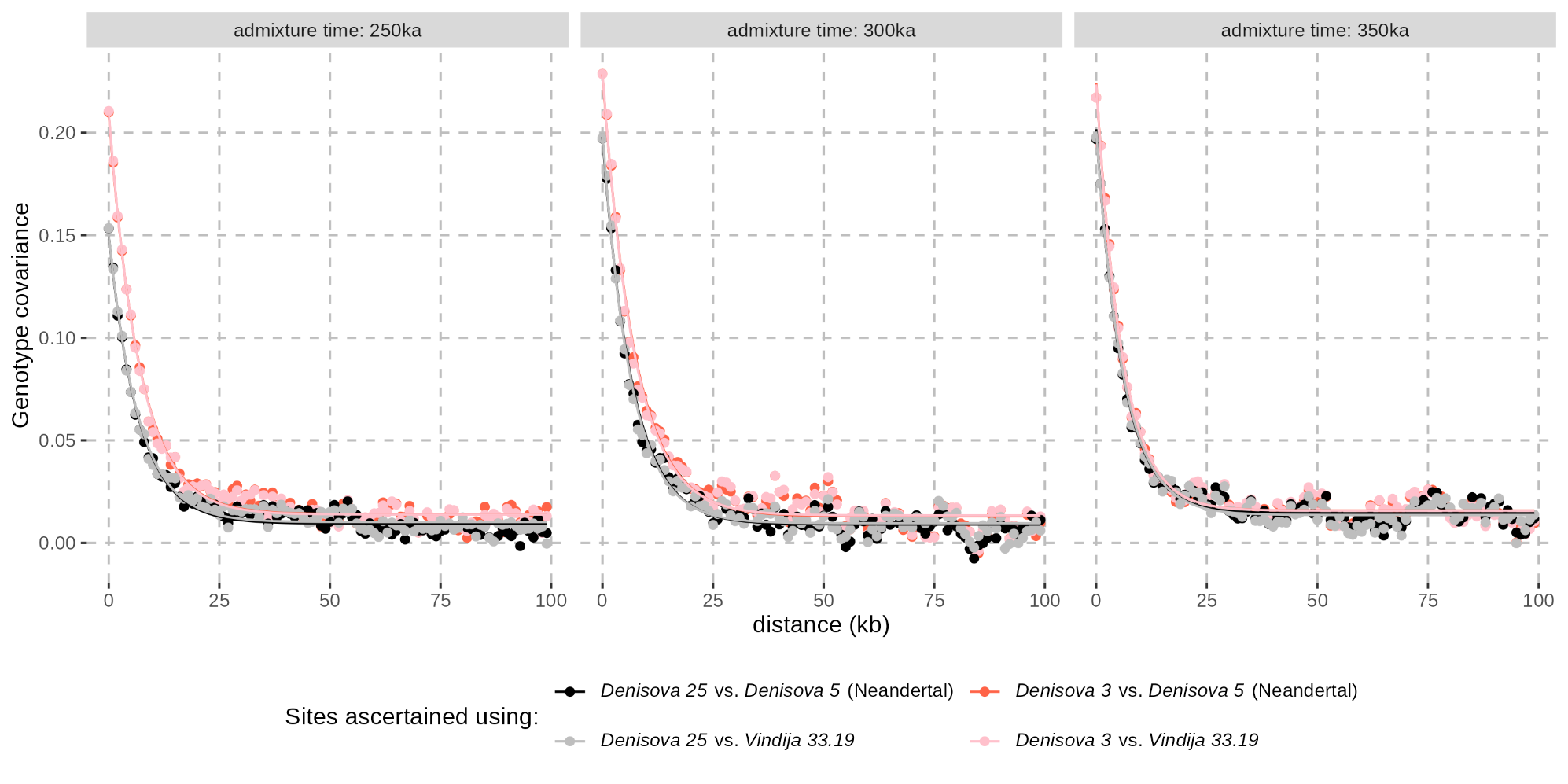

**Figure S70: Decay of genotype covariance in simulations with varying admixture times between Denisovans and “Super-Archaic” hominins.** Covariance was computed as for the empirical data (see **Figure S66** for more details), including random sampling of one allele at heterozygous sites. Each point shows the average covariance for pairs of sites binned by distance (rounded down to the nearest kb, x-axis), based on 1,000 simulated loci of 1Mb each. These simulations include 10% gene flow into Denisovans from a “Super-Archaic” hominin population that diverged 1Ma. See **Figure S66** for more details about the figure layout, and **Figure S67** and the text above for more details about the simulations.

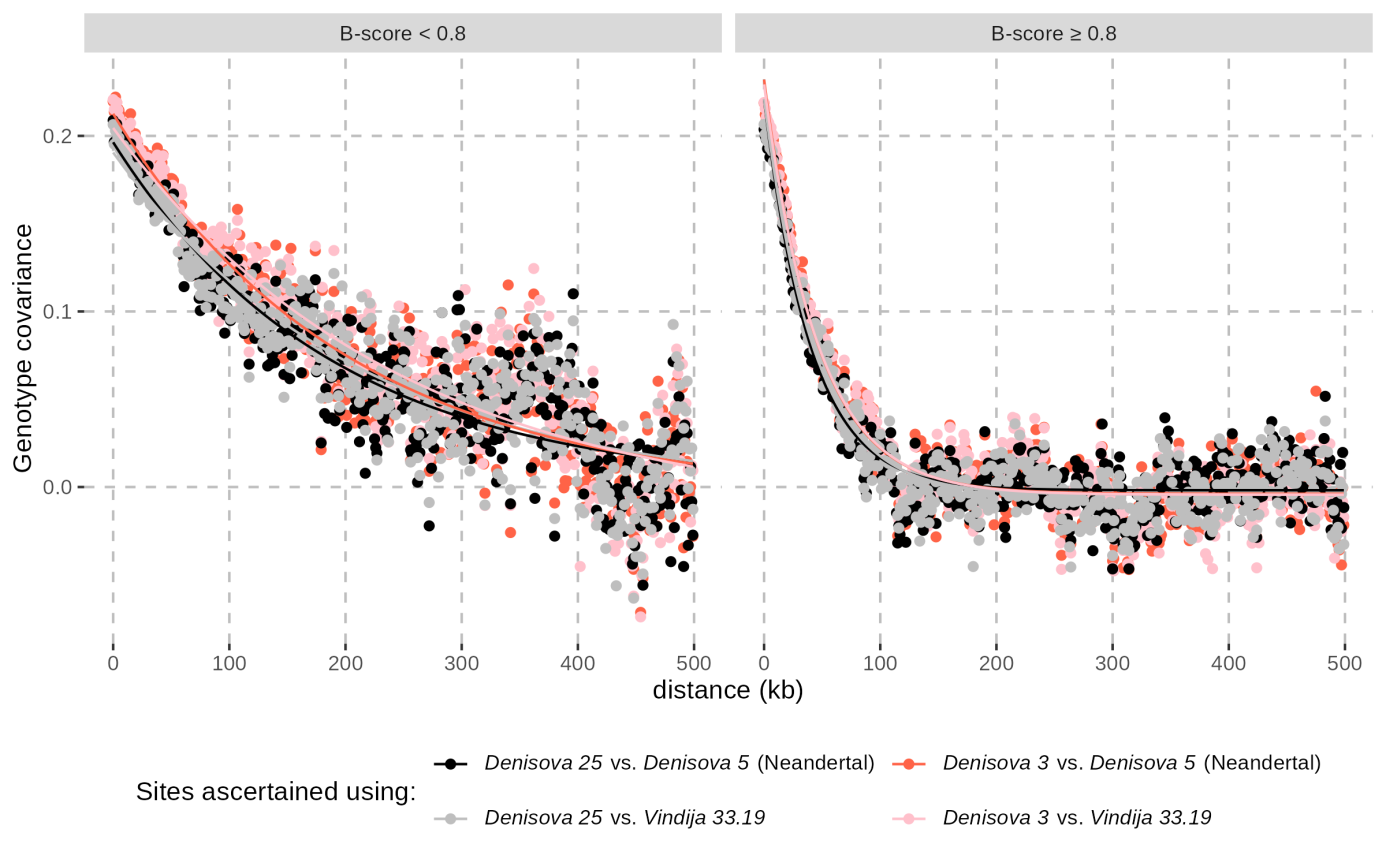

**Figure S71: Decay of genotype covariance in genomic regions with varying B-scores.** See **Figure S66** for more details about the figure layout.

#### Identification of candidate “Super-Archaic” segments using *hmmix*

If the ancestors of the potential “Super-Archaic” hominin diverged deeply from the lineage leading to Neandertals, Denisovans and modern humans, filtering out shared variants among these groups could reveal segments in the Denisovan genomes that still contain a high density of variants from this deeply divergent lineage, assuming that Denisovans do not share these segments with Neandertals or modern humans.

To detect such potential “Super-Archaic” ancestry segments in the Denisovan genomes and, for comparison, the Neandertal genomes, we applied *hmmix* as described in **Supplementary Note 11**. For regions passing the standard filters in the target archaic genome, we assessed mutation rate variation in 100kb windows using *hmmix*, from the number of variants in the outgroup genomes consisting of both Africans from the SGDP dataset (excluding Mozabite and Saharawi; from the VCF described in **Supplementary Note 8**) and either the three Neandertals or the two Denisovans, depending on the target genome. After excluding genetic variation found in the outgroup, we trained the model with default initial parameters in regions that pass the standard filters, providing the relative estimates of mutation rate variation along the genome. Ancestral states were provided using the same reconstruction of the ancestral sequence used in **Supplementary Note 11**. Finally, we applied the posterior decoding algorithm to identify candidate “Super-Archaic” segments using the parameter values that maximize the likelihood. Unlike in **Supplementary Note 11**, we only considered putative “Super-Archaic” segments with a minimum posterior probability of 0.95 for downstream analyses.

We identified 46 and 70 candidate segments for “Super-Archaic” ancestry in the *Denisova 3* and *Denisova 25* genomes, respectively, including 16 segments overlapping between the two. For the Neandertal genomes, we identified 44, 45, and 37 candidate segments in *Denisova 5*, *Chagyrskaya 8*, and *Vindija 33.19*, respectively, with 11 segments overlapping across all three genomes (up to 16 overlapping between *Denisova 5* and *Chagyrskaya 8*). The nearly equivalent number of segments identified in Neandertals and *Denisova 3* suggests either a similar amount of “Super-Archaic” ancestry in Neandertals and Denisovans or a high rate of false positives despite our conservative posterior probability threshold. The higher number of candidate segments in *Denisova 25* may reflect the closer temporal proximity of this individual to a potential admixture event, which should facilitate the detection of any “Super-Archaic” ancestry.

A significant proportion of the candidate segments were also identified in a previous scan for “Super-Archaic” ancestry^103^, supporting their authenticity in *Denisova 3*, as well as in the *Denisova 5* and *Vindija 33.19* Neandertals. Of the 46 segments identified in *Denisova 3*, 15 (~33%) overlap with those reported by Hubisz *et al*.; similarly, of the 44 and 37 segments identified in the *Denisova 5* and *Vindija 33.19* Neandertals, 10 segments in each genome (~23-27%) overlap with those reported by Hubisz *et al.* for the same genomes. Although the previous study predicted low levels of “Super-Archaic” ancestry in Neandertals, the predicted levels only slightly exceeded their estimated false positive rate leading the authors to interpret most signals as likely spurious. It remains possible that our overlapping calls are also false positives for the same reasons. The higher number of candidate segments identified in *Denisova 25* may suggest that at least some of these candidates are genuine.

If the detected “Super-Archaic” ancestry segments are genuine and shared between the two Denisovans from a common admixture event, segments should be longer in *Denisova 25* than in *Denisova 3*, as *Denisova 25* lived closer in time to the admixture event. Consistent with this prediction, the average segment length is 105kb in *Denisova 25* and 89kb in *Denisova 3*. This trend is not observed for segments with lower minimum posterior probabilities: segments with a minimum posterior probability of 0.9 are on average 81kb in *Denisova 3* and 80kb in *Denisova 25*, and those with a minimum posterior probability of 0.8 are on average 60kb in *Denisova 3* and 59kb in *Denisova 25*. In contrast, segments identified in the Neandertal genomes (with a minimum posterior probability of 0.95) are longer and their length increases in more recent genomes (146kb in the ~120,000-year-old *Denisova 5*, 169kb in the ~80,000-year-old *Chagyrskaya 8*, and 184kb in the ~50,000-year-old *Vindija 33.19*), a pattern inconsistent with inheritance of “Super-Archaic” segments from a common admixture event. However, given the dependence of *hmmix* segment length on the minimum posterior probability and the difficulty to accurately determine the length of segments (as described in **Supplementary Note 11** for archaic ancestry in present-day human genomes), these results should be taken with caution.

Another line of evidence that would support the presence of “Super-Archaic” ancestry in the Denisovan genomes is high divergence not only between Neandertals and Denisovans but also between Denisovans in candidate regions. To assess this, we measured divergence as the proportion of alleles the archaic genome with potential “Super-Archaic” ancestry does not share with another archaic genome at sites where the latter differs from great ape genomes. One random allele was sampled at heterozygous positions. In the candidate regions, Denisovan-Neandertal divergence ranged from 10% to 50% of the divergence to Chimpanzee, which is remarkably high (**Figure S72** and **Figure S73**). In contrast, divergence between archaic individuals from the same group (e.g. *Denisova 3* vs. *Denisova 25* at candidate regions for “Super-Archaic” ancestry in *Denisova 25*) was typically below 10%. However, in the Denisovan genomes, about 10 regions per individual also showed >10% divergence to the other Denisovan, a pattern that is extremely rare in Neandertals (just one region), suggesting that some segments in Denisovan genomes are unusually divergent even among Denisovans. Supporting this, several candidate regions in the Denisovan genomes exhibit extremely high heterozygosity, unlike Neandertals, except to a lesser extent in the *Denisova 5* Neandertal genome (**Figure S74**). These observations are consistent with deeply divergent ancestry among Denisovans. Finally, coverage in candidate regions does not deviate from genome-wide levels, arguing against the possibility that these represent cryptic duplications with inflated heterozygosity (**Figure S75**).

Overall, the combination of high divergence, extreme heterozygosity, and overlap with previous scans supports the conclusion that Denisovan genomes retain segments plausibly derived from a “Super-Archaic” lineage. The unexpectedly similar number of candidate segments in *Denisova 3* and the Neandertal genomes, despite a higher count in *Denisova 25*, suggests that either Neandertals carried comparable levels of such ancestry or that many calls are false positives. We found no evidence for “Super-Archaic” ancestry in Neandertals, but the temporal span among Neandertal genomes is much smaller than between the two Denisovans, and any introgressed segments in the ~120ka *Denisova 5* Neandertal may already be short, limiting the differences in genotype covariance decay between Neandertals. The difficulty in detecting segments from a “Super-Archaic” hominin suggests that this hominin was moderately divergent, and admixture may be ancient enough for recombination to erode most tracts.

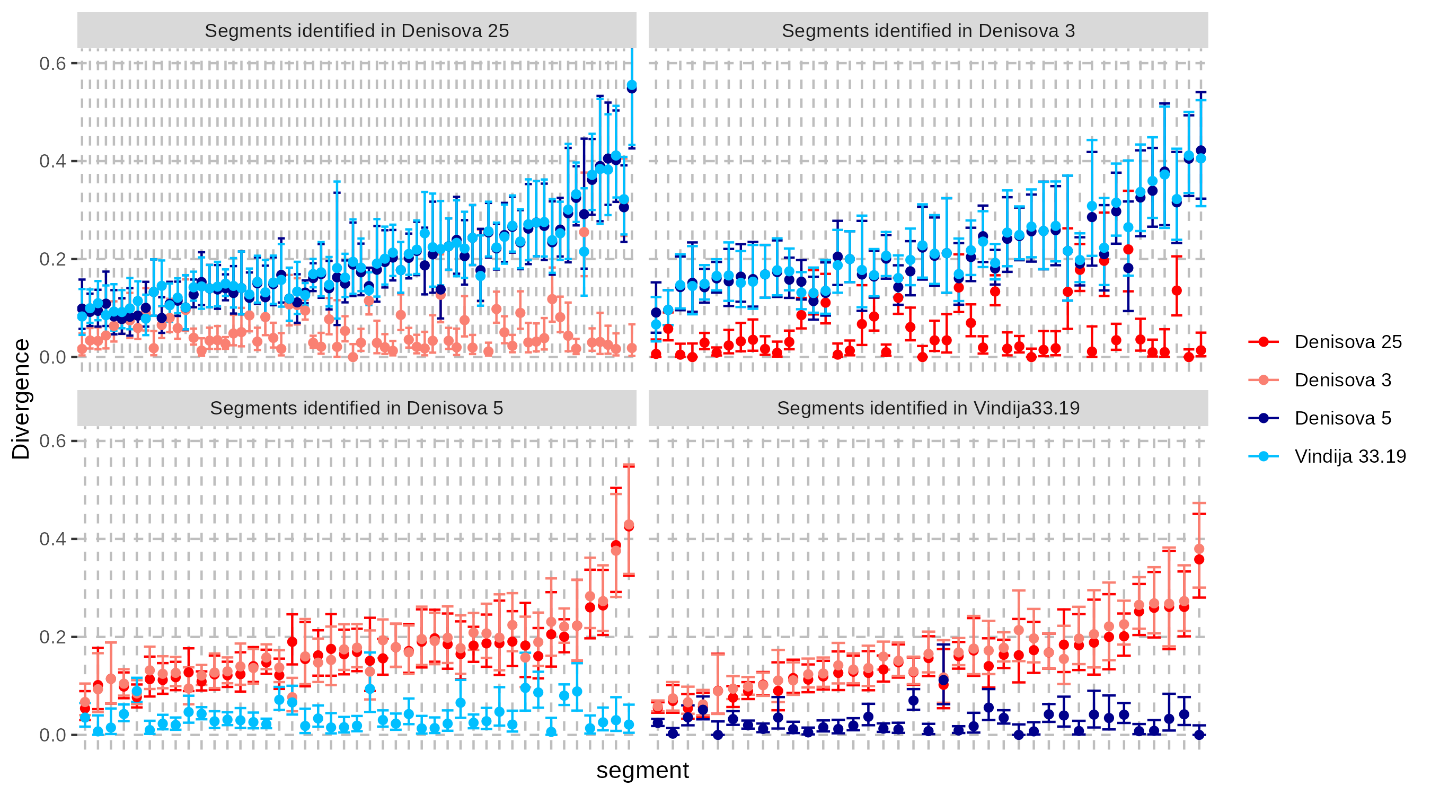

**Figure S72: Pairwise divergence between archaic human genomes in candidate regions for “Super-Archaic” ancestry.** Divergence was calculated as the proportion of derived alleles the archaic genome with potential “Super-Archaic” ancestry (see facet label) does not share with the other archaic genome (indicated by colour) at sites where the latter differs from great ape genomes. One random allele was sampled at heterozygous positions. Whiskers represent the 95% confidence intervals based on a Poisson distribution.

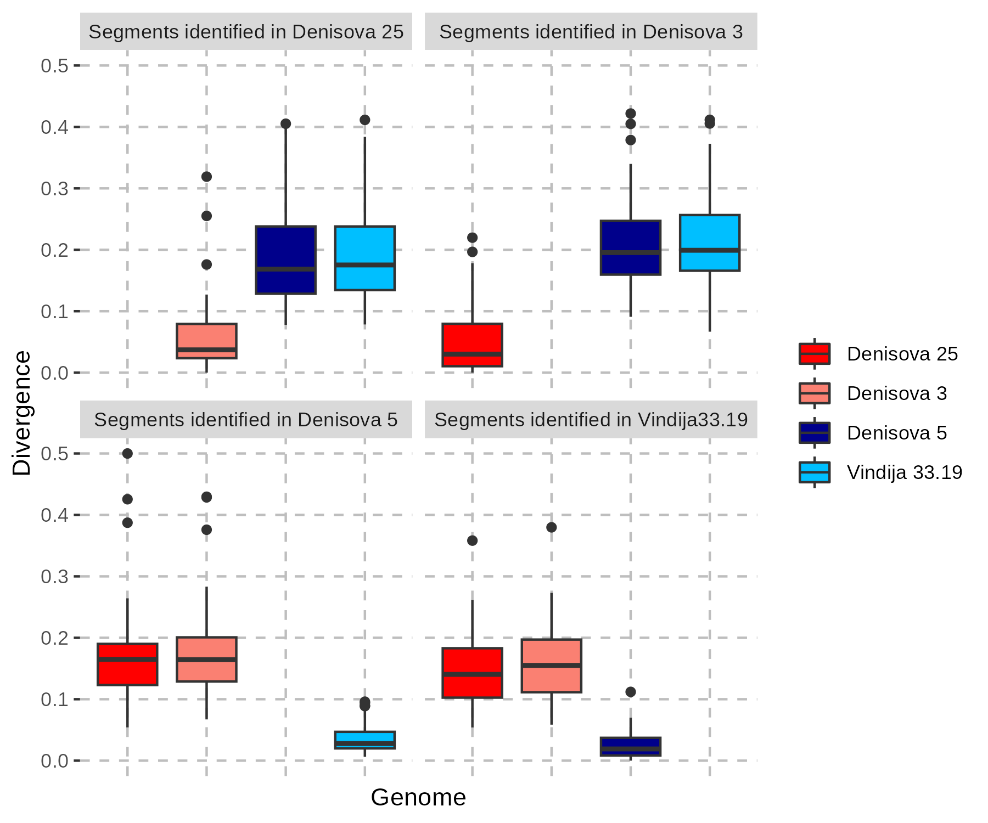

**Figure S73: Summary of pairwise divergence between archaic human genomes in candidate regions for “Super-Archaic” ancestry.** This is a boxplot representation of the data shown in **Figure S72** highlighting more outliers in the divergence between the two Denisovans, when one carries a potential “Super-Archaic” segment, than between Neandertals under similar conditions.

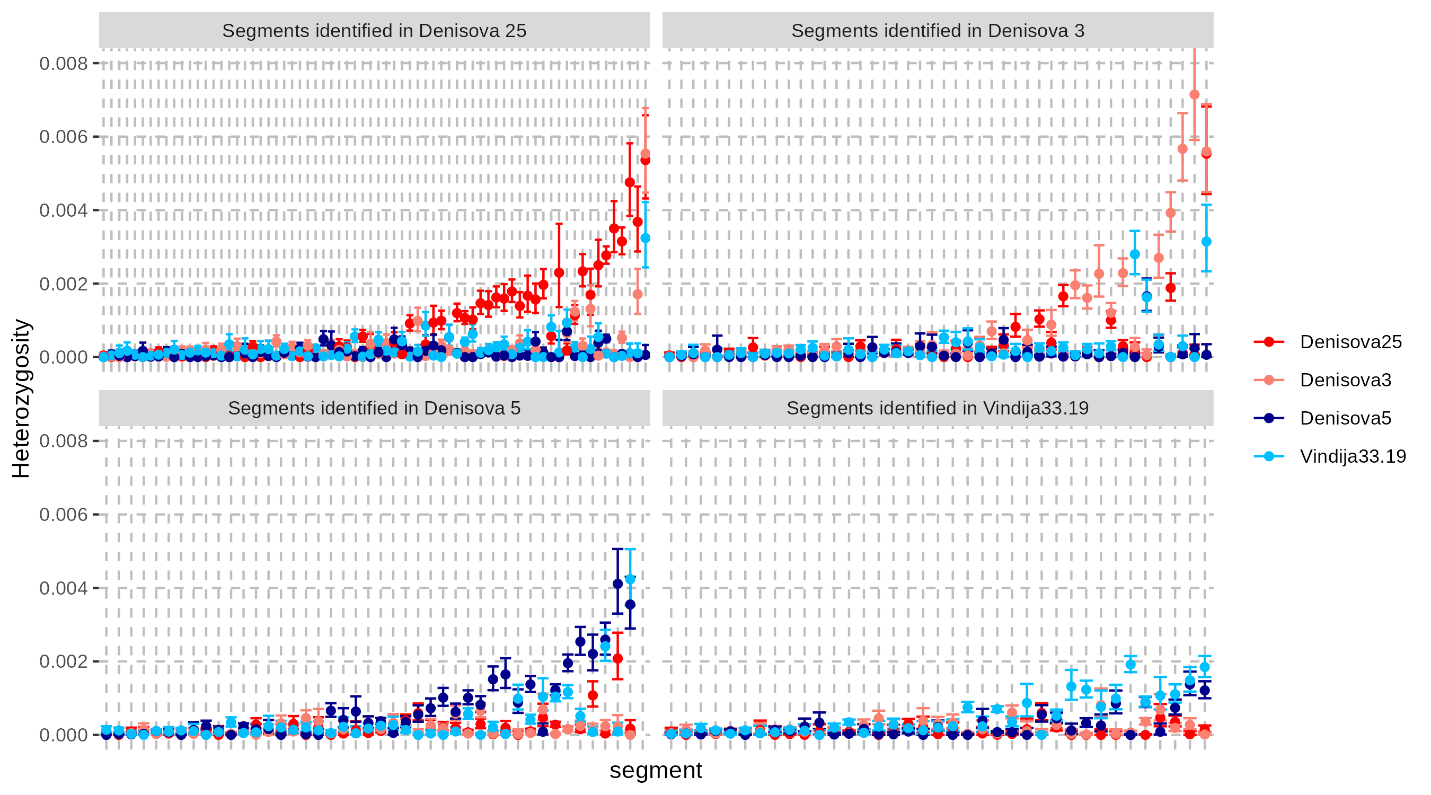

**Figure S74: Heterozygosity of archaic human genomes in candidate regions for “Super-Archaic” ancestry.** Heterozygosity was measured as the proportion of heterozygous sites among those passing the standard filters in all four archaic genomes. Colours indicate the genome for which heterozygosity was calculated in regions where the genome indicated in the facet label carries potential “Super-Archaic” ancestry. Whiskers represent the 95% confidence intervals based on a Poisson distribution.

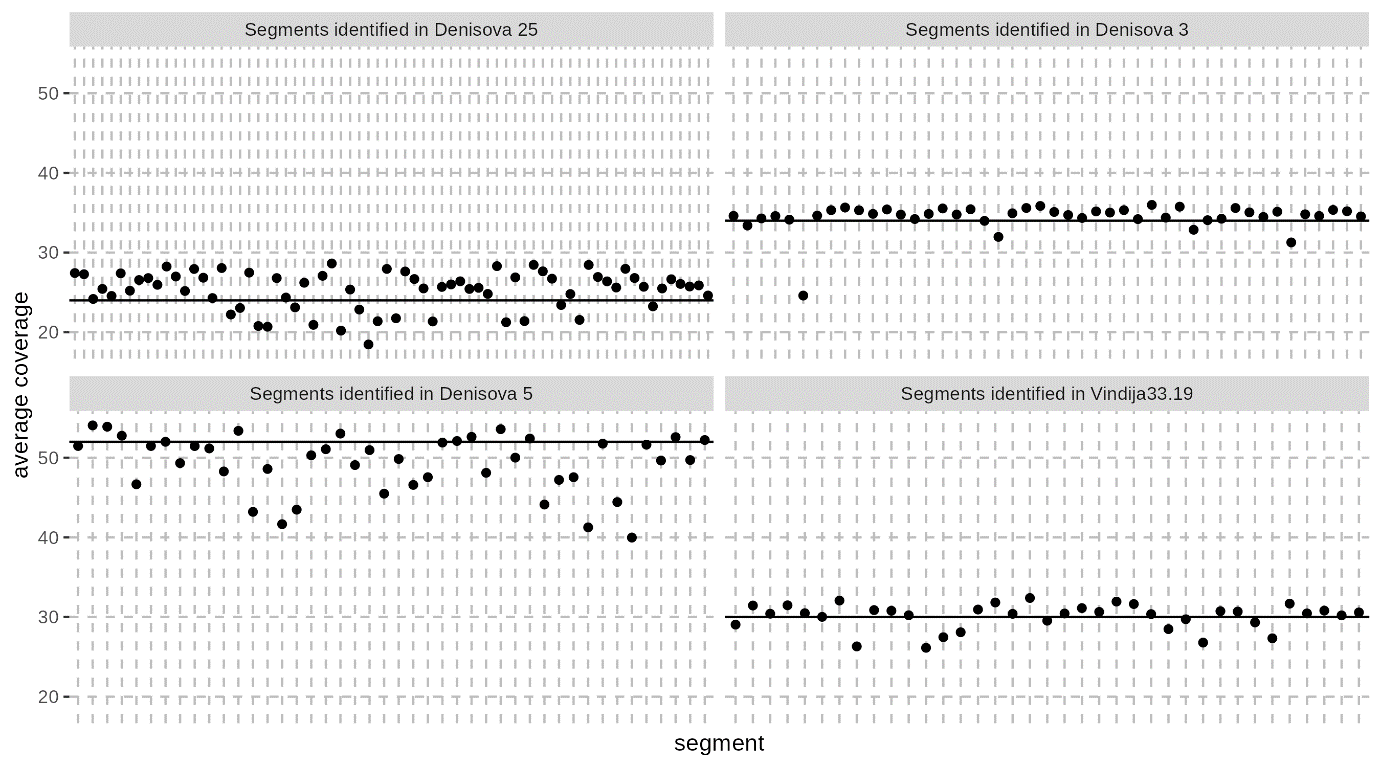

**Figure S75: Average coverage in candidate regions for “Super-Archaic” ancestry.** Horizontal lines correspond to the average genome-wide coverage of the genome of each facet.

### Supplementary Note 11: Relationship to the Denisovan ancestries in present-day humans

Prior research has revealed the existence of multiple Denisovan ancestry components in the genomes of present-day people, each displaying different degrees of divergence from the *Denisova 3* genome. Notably, present-day populations in Oceania and the Philippines exhibit as much as 4% Denisovan ancestry^7,122,123^, from populations that were distantly related to *Denisova 3*^124^. Conversely, present-day Asians and Native Americans carry less than 0.5% of Denisovan ancestry^48,125,126^, some of which was inherited from a population closely related to *Denisova 3*, as shown in East Asians^127^ and Native Americans^124^. Finally, present-day Europeans carry small traces of Denisovan ancestry^80,128^.

With the availability of a second Denisovan genome, we aimed to further characterize how the Denisovan ancestry in present-day people relates to the sequenced Denisovan genomes.

#### Analysis of previously published archaic segments identified with *Sprime*

We downloaded the output from the *Sprime* analysis of non-African genomes of the 1000 Genomes dataset^129^ and from Papuan genomes of the SGDP dataset^106^, as published by Browning *et al.* (2018)^127^. Each file - one for each chromosome and each population - contains a list of Single Nucleotide Polymorphisms (SNPs) with putatively introgressed archaic alleles. These alleles were compared by the authors to a Neandertal (*Denisova 5*) and Denisovan (*Denisova 3*) genomes, categorized as "match" (archaic genome carries the allele), "mismatch" (archaic genome does not carry the allele), or "notcomp" (the position is unavailable in the archaic genome due to missing data or quality issues). Each allele was assigned to a putatively introgressed haplotypes identified by *Sprime*, so that one can compute for each segment the proportion of putative archaic-specific alleles that match a given archaic genome.

We applied standard filters (**Supplementary Note 3**) to the genotype calls of *Denisova 25*, and randomly sampled one allele at heterozygous sites. For each SNP with putatively introgressed alleleson we then determined whether the allele of *Denisova 25* matches or mismatches the allele on the putative archaic segment, or whether the allele of *Densiova 25* was missing or filtered out due to potential quality issues. We then generated two-way density plots of the match rates to the *Denisova 3* and *Denisova 25* genomes, defined as the proportion of alleles a segment matches each genome (**Figure S76**). We only included segments with a match rate ≥10% to at least one Denisovan genome, and with a greater match rate to a Denisovan genome than to the *Denisova 5* Neandertal genome.

In all populations we find that more putatively introgressed segments are more closely related to *Denisova 3* than to *Denisova 25*, though many segments are equidistant (**Figure S76**). A subset of segments in the genomes of present-day East Asians, Native Americans, and some Europeans (e.g. Finnish), match the *Denisova 3* and *Denisova 25* genomes at about 80% and 60% of the positions, respectively, supporting the presence of ancestry inherited from a population closely related to *Denisova 3* in these populations. In contrast, in South Asians and Papuans, the segments tend to only slightly better match *Denisova 3* than *Denisova 25* (as seen in the excess of segments slightly below the diagonal in ***Figure S76***). As the latter signal could be confounded by the Neandertal ancestry of *Denisova 3*, which is closer to the Neandertal population that contributed ancestry to present-day people (**Supplementary Note 9**), we excluded all the segments that overlap regions of the genome where *Denisova 3* or *Denisova 25* carry Neandertal ancestry (at least 0.05cM long, as identified with *admixfrog* with a penalty of 0.4 and a window size of 0.005cM). The results remain unchanged (**Figure S77**), suggesting that a more distantly related Denisovan ancestry component in these populations is slightly closer to *Denisova 3* than *Denisova 25*.

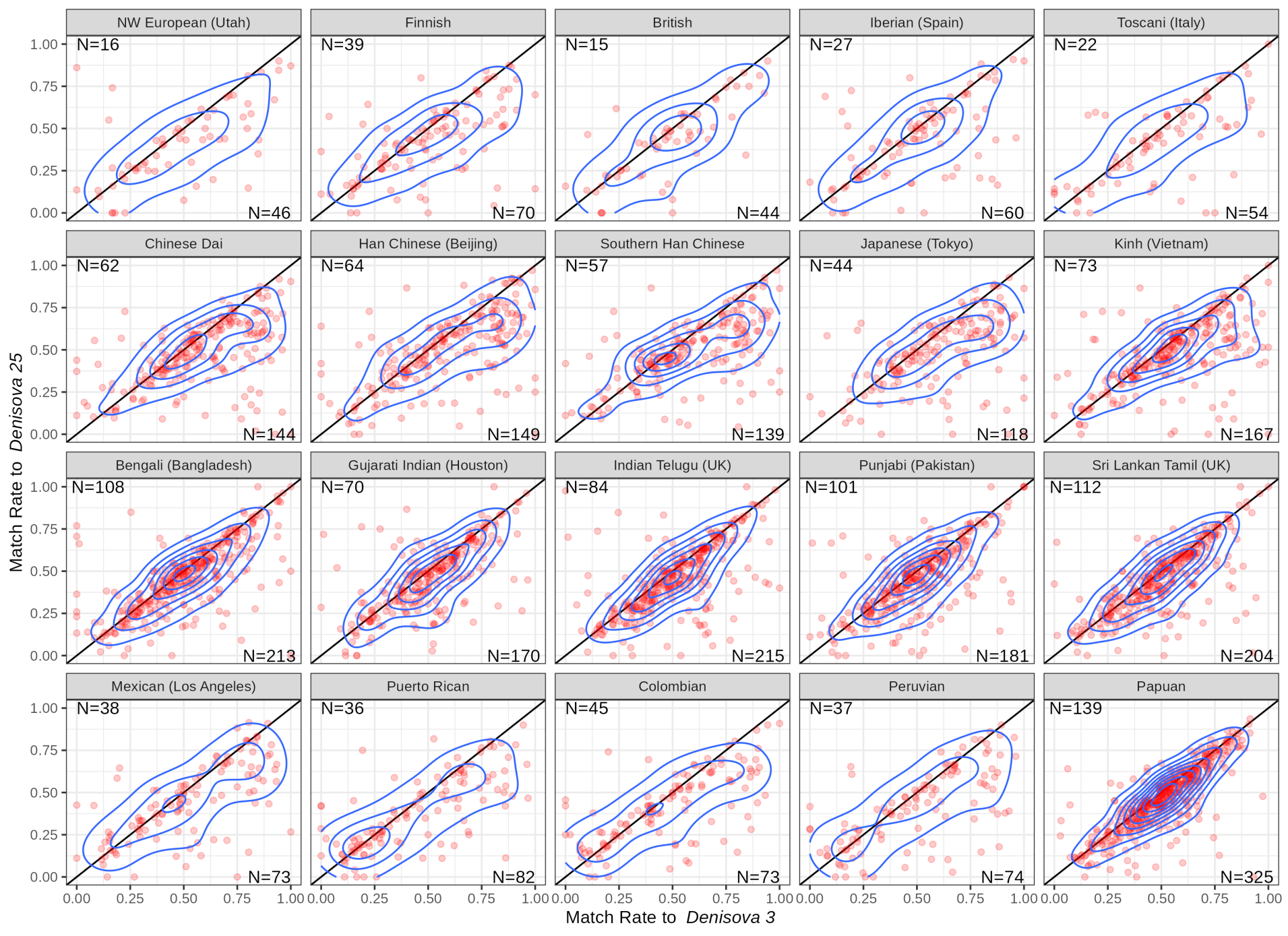

**Figure S76: Match rate to the Denisova 3 and Denisova 25 genomes of putative Denisovan segments identified with Sprime in the genomes of present-day people.** Each point corresponds to a putative introgressed segment called by Browning et al. (2018)^127^, while the contour lines indicate the density distribution of these segments. Only segments that match at least one Denisovan genome in more than ten percent of positions and that match more often a Denisovan genome than the Denisova 5 Neandertal genome are shown. The numbers of segments below and above the diagonal (i.e. closer to Denisova 3 and Denisova 25, respectively) are indicated inside the plots. European populations are given in the first row, East Asian populations in the second row, South Asian populations in the third row, and American and SGDP Papuan populations in the final row.

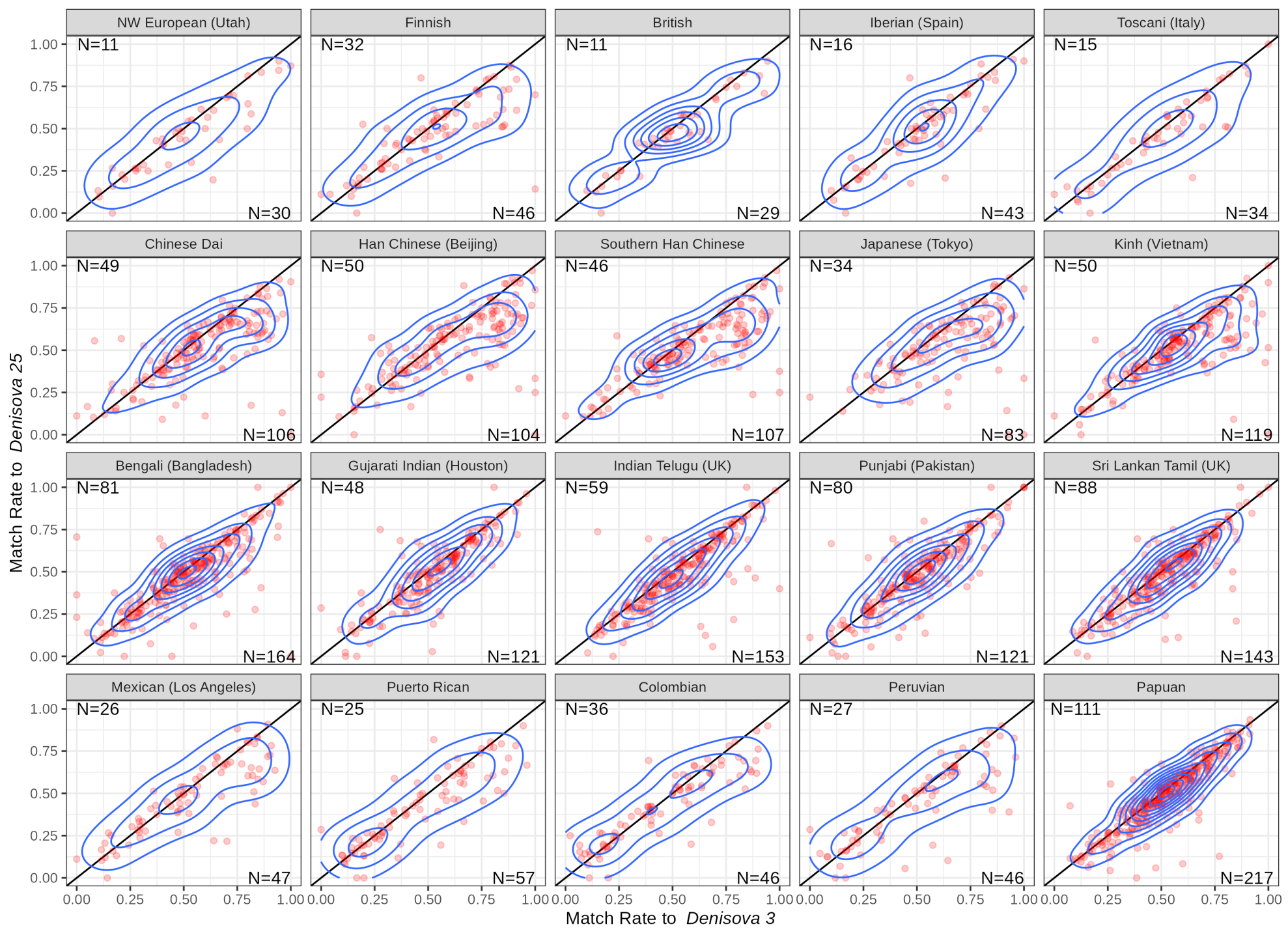

**Figure S77: Match rate to the Denisova 3 and Denisova 25 genomes of putative Denisovan segments identified with Sprime in the genomes of present-day people after excluding genomic regions where the Denisovan genomes carry segments potentially introgressed from Neandertals.** We excluded segments that overlap regions where a Denisovan carries Neandertal segments of at least 0.05cM (as identified with admixfrog with a penalty of 0.4 and a window size of 0.005cM). See **Figure S76** for more information about the layout of these plots.

#### Analysis of archaic segments identified with another reference free method, *hmmix*

Since the method used to call putatively introgressed tracts may influence the results, we used another reference free method, *hmmix*^130^, to identify Denisovan segments in present-day human genomes. Briefly, *hmmix* is a hidden Markov model that scans the genome in sliding non-overlapping windows to identify those with a high density of single nucleotide variants that are absent from an unadmixed outgroup. Regions with a high density of remaining variants are potentially introgressed from a population more divergent than the unadmixed outgroup. Unlike *Sprime*, which provides a tiling of putative archaic segments, *hmmix* operates on a single genome and can identify individual archaic haplotypes.

We used the SGDP dataset as described in **Supplementary Note 8**. Each non-African genome was analyzed separately, as computing the variant density is best achieved by providing callable regions for each genome individually (i.e., regions that pass the filters described in **Supplementary Note 8**, including the minimum 10-fold and GC-corrected coverage cutoffs that are specific to each genome). Focusing on the callable regions of each examined genome, we first assessed the variation of the mutation rate in the same regions in the 40 sub-Saharan African genomes from the SGDP dataset (i.e., all African genomes of the dataset excluding Mozabite and Saharawi genomes) to avoid falsely calling archaic segments in regions with a high mutation rate. The mutation rate was calculated using the density of all variants in 100kb windows divided by the mean variant density of the whole genome (using the mutation_rate command of *hmmix*). After removing variants identified in the 40 sub-Saharan African genomes, the model was then trained with default initial parameters, providing both the relative estimates of mutation rate variation along the genome and the callable regions. Ancestral states were provided to *hmmix* using a reconstruction of the ancestral sequence (available as a Fasta file at: ftp://ftp.ensembl.org/pub/release-74/fasta/ancestral_alleles/homo_sapiens_ancestor_GRCh37_e71.tar.bz2). Finally, we ran the posterior decoding algorithm to identify archaic segments using the parameter values that maximize the likelihood. In that step, we provided all available high-coverage archaic genomes (*Chagyrskaya 8*, *Denisova 3*, *Denisova 5*, *Denisova 25*, and *Vindija 33.19*) to annotate each identified segment with the number of alleles it shares with each archaic genome. We only considered putative archaic segments with a minimum posterior probability of 0.8 for downstream analyses.

Identified segments were then assigned to different groups based on the number of alleles they shared with archaic genomes (as annotated by *hmmix*). If a segment did not share any variant with any archaic genome, that segment was discarded (0.07% of segments). If less than 10% of the variants in a segment were shared with all archaic genomes, the origin of this segment was considered “unknown”. Remaining segments were assigned as “Denisovan” or “Neandertal” if at least one Denisovan or one Neandertal genome shared more alleles with that segment than any Neandertal or any Denisovan, respectively. In the case of a tie, the segments were considered “ambiguous”. To evaluate this assignment, we also analyzed segments following an alternative stricter filtering in which segments were assigned to an archaic group only if a genome from that group shared at least 3 more variants than any genome from the other archaic group.

After removing segments that are shorter than 0.05cM, which are difficult to distinguish from segments originating from incomplete lineage sorting, we detected less than 10Mb of Denisovan ancestry in genomes of non-African individuals outside Oceania (with a minimum of 334kb in a European), and up to 85Mb in Oceanian individuals (**Figure S78**). In contrast, the detected Neandertal ancestry ranges from 35Mb to 79Mb in non-African individuals, with the minimum observed in a Middle Eastern individual and the maximum in an Oceanian. Ambiguous segments generally represent less than 1Mb, with a minimum of 73kb in a Thai individual and a maximum of 2.9Mb in a Papuan, while segments classified as “unknown” span 11Mb to 23Mb, with minimal geographical variation.

We next compared the segments identified with *hmmix* and *Sprime* in the 15 Papuan genomes of the SGDP dataset. We found that 66% of Denisovan segments identified with *Sprime* (i.e. those that match better a Denisovan genome than any Neandertal genome, as described above) are recovered by *hmmix* (**Table S64**). Although this is lower than reported by Skov *et al.* (2018)^130^ (84.9%), including segments that we assigned as unknown or ambiguous raised that recovery rate to 78%, closer to that reported in Skov *et al*. (2018)^130^. We also observed that 23% of segments called by *hmmix* were identified by *Sprime*, showing that *hmmix* identifies more putative Denisovan segments (**Table S65**).

**Table S64: Summary of Denisovan segments called with Sprime and recovered by hmmix.** Segments were identified by Browning et al. (2018)^127^ using Sprime in 15 Papuan genomes from the SGDP dataset and overlapped with those we identified with hmmix in the same genomes. A segment from Sprime is considered recovered by hmmix if any overlap was observed.

|  |  |  | ***Sprime*** |
| --- | --- | --- | --- |
| ***hmmix*** | **All segments excluding Neandertal segments** | **present** | 417 |
|  |  | **absent** | 116 |
|  |  | **overlap proportion** | **78%** |
|  | **All Denisovan segments** | **present** | 366 |
|  |  | **absent** | 167 |
|  |  | **overlap proportion** | **69%** |
|  | **Denisovan segments ≥ 0.05cM** | **present** | 351 |
|  |  | **absent** | 182 |
|  |  | **overlap proportion** | **66%** |

**Table S65: Summary of Denisovan segments called with hmmix and identified by Sprime.** Segments were identified with hmmix in 15 Papuan genomes from the SGDP dataset and overlapped with those identified by Browning et al. (2018)^127^ using Sprime in the same genomes. A segment from hmmix is considered identified by Sprime if any overlap was observed. Note that the numbers of overlapping segments are different from **Table S64** because a segment called with one method could overlap multiple segments from the other method, so it is counted once for the former and multiple times for the latter.

|  | | ***hmmix*** | |
| --- | --- | --- | --- |
|  |  | **All Denisovan segments** | **Denisovan segments ≥0.05cM** |
| ***Sprime*** | **present** | 440 | 407 |
|  | **absent** | 1,783 | 1,357 |
|  | **overlap proportion** | **20%** | **23%** |

We then generated two-way density plots of the match rates of Denisovan segments to the *Denisova 3* and *Denisova 25* genomes (**Figure S79**), similar to those generated with the segments identified with *Sprime*. However, note that the ascertainment by *hmmix* corresponds to variable positions in non-Africans where Africans carry the ancestral allele and where at least one archaic human genome carries the derived allele. In contrast, the ascertainment by *Sprime* corresponds to all putatively introgressed alleles that are on inferred archaic haplotypes. Both ascertainments avoid variants that may be on the modern human background for heterozygous haplotypes and disregard variants that arose on the lineages of the archaic genomes. We pooled segments identified by *hmmix* from all individuals into six geographical groups (as defined by Mallick *et al.* (2016)^106^), and found that all groups carry more segments that are closer to *Denisova 3* than to *Denisova 25*, replicating the results obtained with the segments identified with *Sprime*. In addition, genomes from Central Asia/Siberia carry a set of introgressed Denisovan segments that are more closely related to *Denisova 3*, similar to that seen in East Asia and America. These results are all robust to the different assignment strategies evaluated (**Figure S79**, **Figure S80**).

We computed *D*-statistics for the Denisovan segments identified with *hmmix* in each individual genome to further assess the relationship of the Denisovan ancestry in present-day people to *Denisova 3* and *Denisova 25* (**Figure S81**). Unlike the match rate analysis, which relies on putative archaic alleles identified with *Sprime* or sites ascertained for *hmmix* as invariable in sub-Saharan Africans, these *D*-statistics include all variants that differ between *Denisova 3* and *Denisova 25* in these regions. We observed significant positive *D*-statistics for multiple individuals in all geographical groups, reinforcing that at least some of the Denisovan ancestry in present-day non-Africans is closer to *Denisova 3*. Interestingly, the *D*-values are larger in East Asia, America, Central Asia/Siberia than in South Asia and Oceania, in agreement with the observation of a Denisovan component even more closely related to *Denisova 3* in East Asia, America, and Central Asia/Siberia. By contrast, no individual displays a significant negative *D*-statistic (|Z-score|≥3).

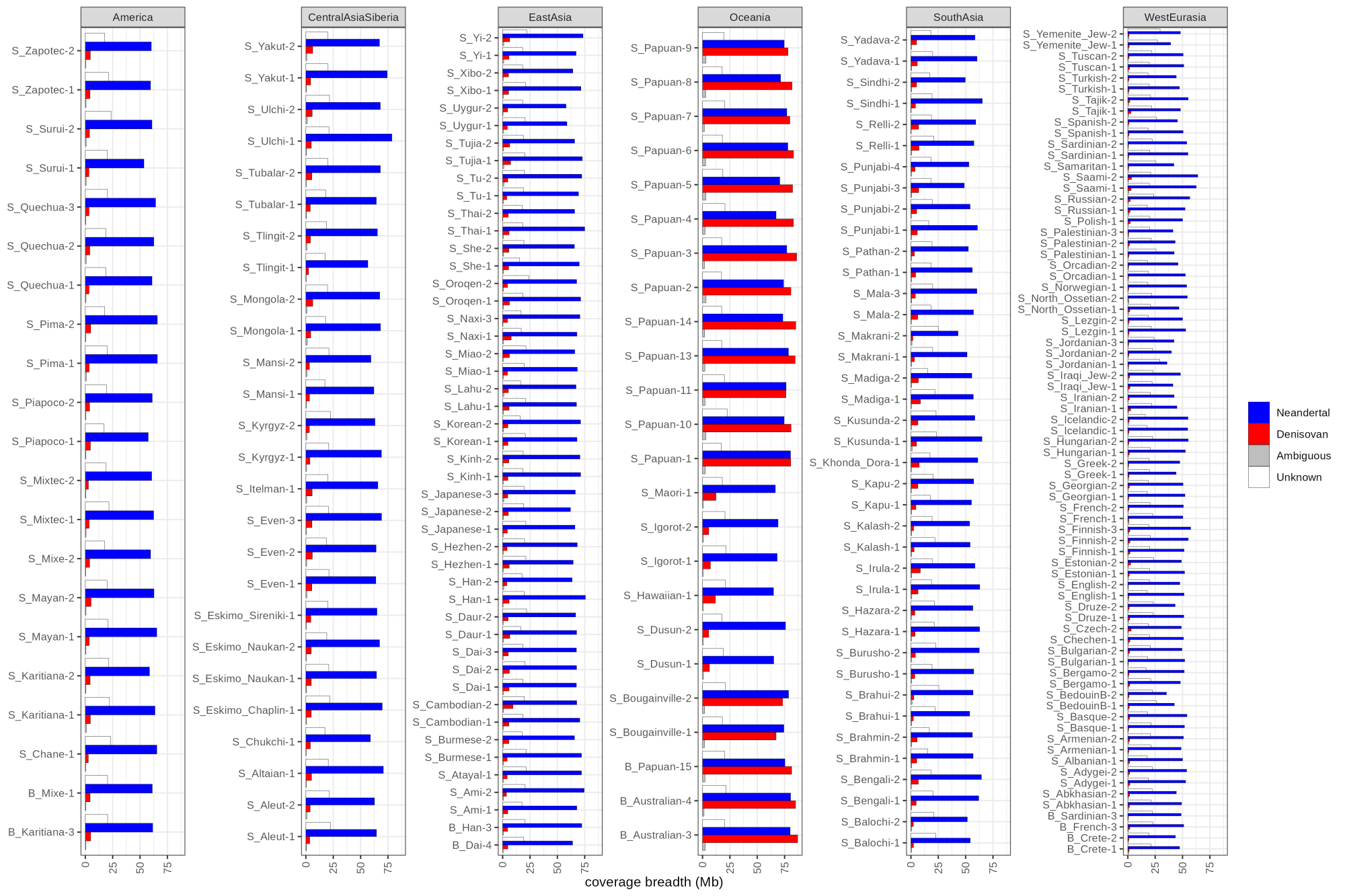

**Figure S78: Amount of the genomes of present-day non-Africans covered by putative archaic segments identified with hmmix.** The coverage breadth is the total number of bases in the genome covered by at least one archaic segment (in Mb). Segments shorter than 0.05cM were excluded. The colours correspond to different segment assignments. Unknown segments are those that do not share enough alleles with the available Neandertal and Denisovan genomes to assign them to a group, while ambiguous segments are those that match equally well the Neandertal and Denisovan genomes.

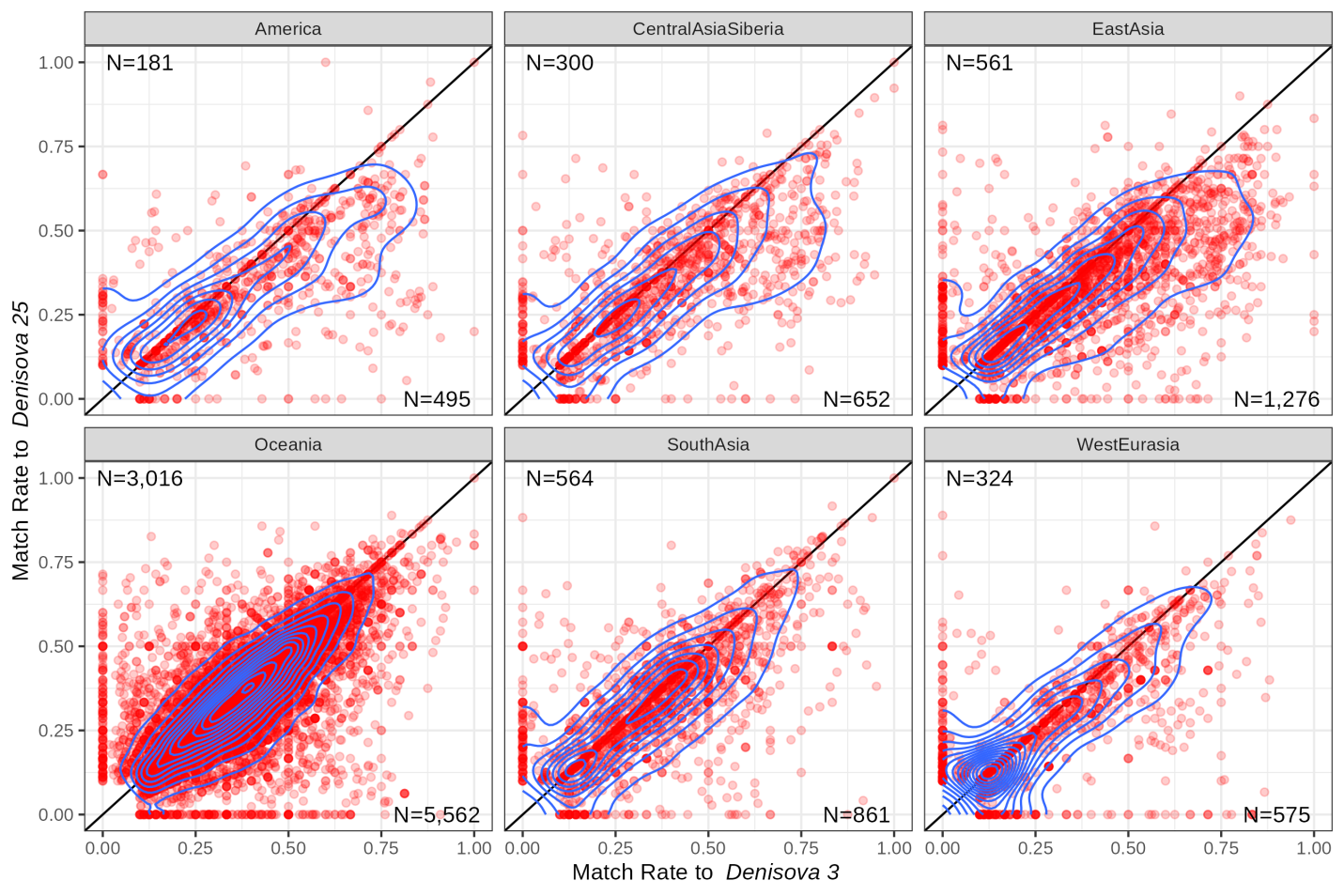

**Figure S79: Match rate of putative Denisovan segments identified with hmmix in the genomes of present-day non-Africans, to the Denisova 3 and Denisova 25 genomes.** Each point corresponds to a putative introgressed segment (≥ 0.05cM). Contour lines indicate the density distribution of these segments. Only segments matching ≥10% of polymorphic positions in at least one Denisovan genome, and more frequently than any high-coverage Neandertal genome, are displayed. The numbers of segments below and above the diagonal (i.e. closer to Denisova 3 and Denisova 25, respectively) are indicated in the plots. Geographical assignments for the individuals are shown atop each panel.

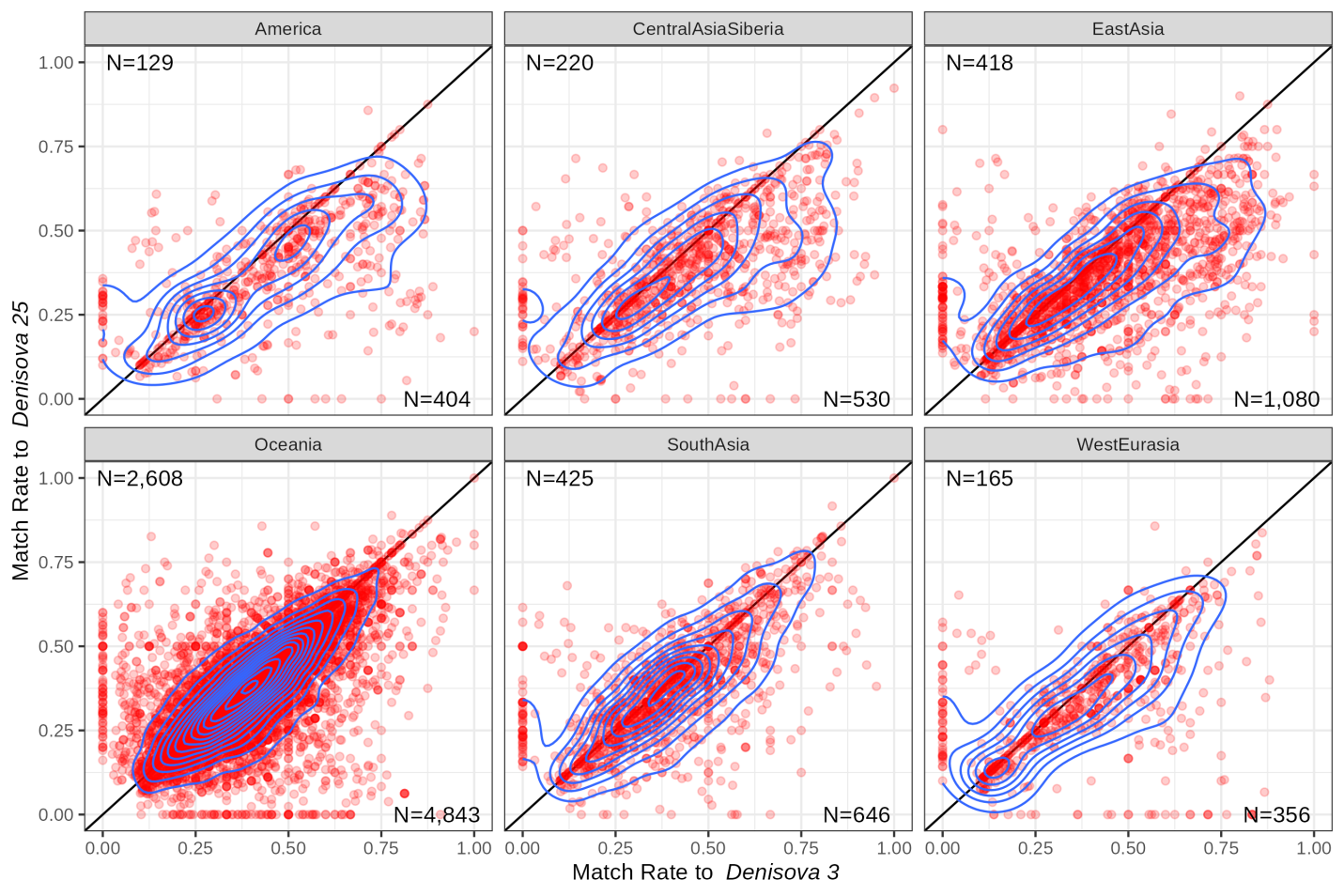

**Figure S80:** **Match rate of more confidently assigned Denisovan segments, identified with hmmix in the genomes of present-day non-Africans, to the Denisova 3 and Denisova 25 genomes.** Displayed segments match at least one Denisovan genome in 10% of polymorphic positions and at least three more positions in a Denisovan genome than all high-coverage Neandertal genomes. Refer to **Figure S79** for additional details.

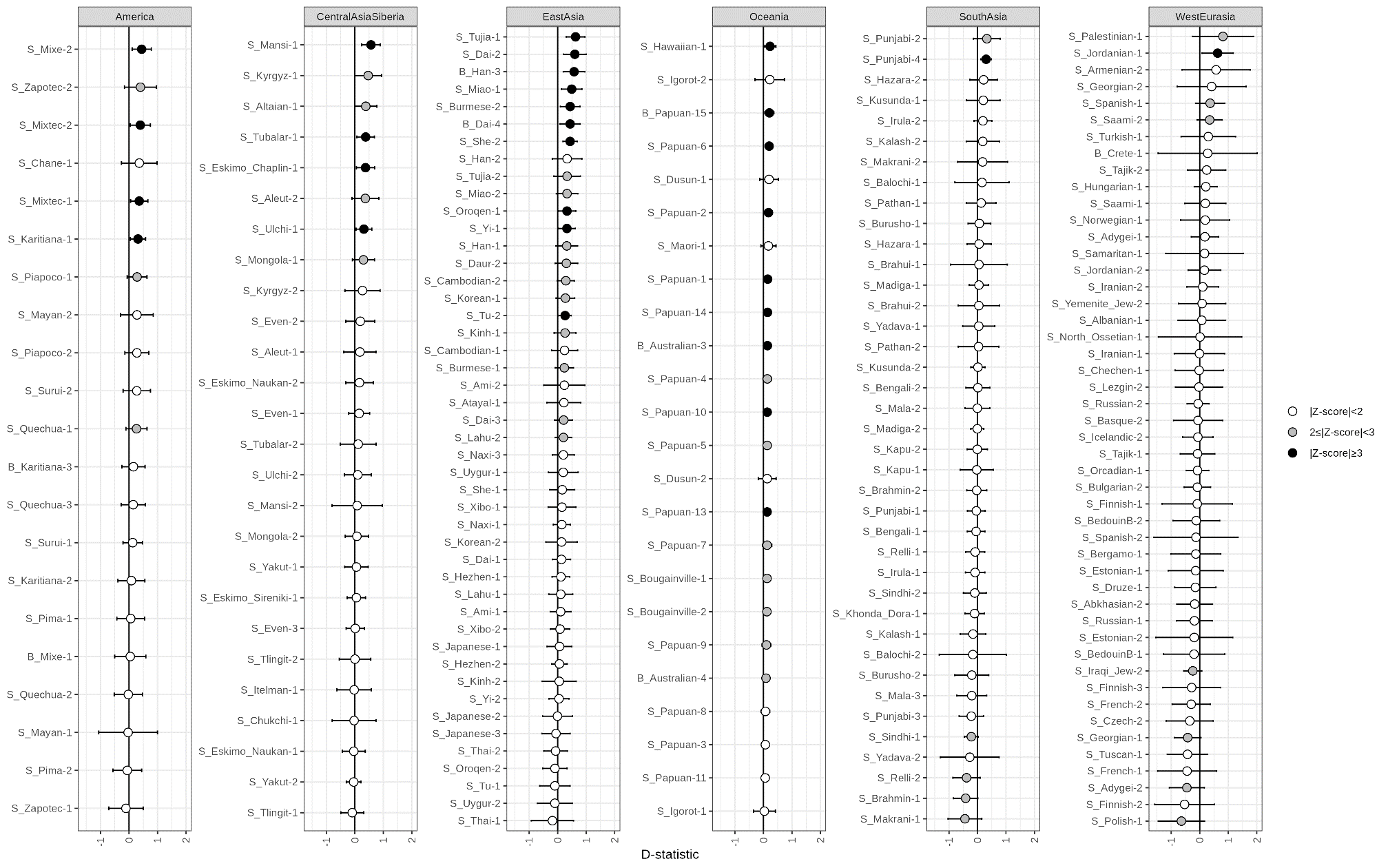

**Figure S81: D-statistics comparing how the Denisovan ancestry of present-day non-Africans relates to Denisova 3 and Denisova 25.** D-statistics were calculated using ADMIXTOOLS^108^ as implemented in admixr ^109^ in regions overlapping Denisovan segments (more confidently assigned set) of each individual, identified with hmmix. Positive values correspond to a closer relationship to Denisova 3. The genomes of panTro4, panPan2, gorGor3, and ponAbe2 were used to determine the ancestral allele. Whiskers on each side of the plotted D values correspond to three standard errors calculated using a Weighted Block Jackknife and a block size of 5Mb across all autosomes. Colours indicate different significance levels.

#### Analysis of archaic segments identified with a reference-based method, *admixfrog*

The availability of a second Denisovan genome may boost the detection power of methods that use reference genomes for detecting archaic ancestry. We tested this by applying *admixfrog* to the SGDP dataset and comparing the Denisovan ancestry detected using *Denisova 3* only and that detected using both *Denisova 3* and *Denisova 25*.

We generated *admixfrog* input files for each non-African genome from the same VCF described earlier (**Supplementary Note 8**) and used a reference set that includes the sub-Saharan Africans from the SGDP dataset. The ancestral state was determined as the allele shared by at least three great ape reference genomes (panTro4, bonobo panpan1.1, gorGor3, ponAbe2), allowing missing data or another allele in one genome as long as this allele was not present in archaic and modern humans (the ancestral allele was otherwise set as missing). We ran *admixfrog* with three states corresponding to Neandertal ancestry, Denisovan ancestry, and modern human ancestry, using the allele frequencies from three Neandertal genomes (*Denisova 5*, *Vindija 33.19* and *Chagyrskaya 8*), one or two Denisovan genome(s) (*Denisova 3* and *Denisova 25*), and the 40 SGDP sub-Saharan African genomes, respectively. The window size was set to 0.005cM using the African-American recombination map^99^ (with linear interpolations when necessary) and the penalty to mitigate erroneous fragmentation of the called segments was set to 0.25. For convenience, we applied *admixfrog* to each chromosome independently. Identified segments shorter than 0.05cM were excluded from further analyses.

We first estimated Denisovan and Neandertal ancestry proportions using an algorithm implemented in *admixfrog*^120^. This algorithm performs random samples from the empirical-Bayes posterior probability distribution of hidden states given the observed data and the maximum likelihood estimates of the parameters. It involves drawing the initial state based on stationary probabilities and, iteratively, randomly sampling the state for the next genomic window based on the calculated transition probabilities in the hidden Markov model of *admixfrog*. This was repeated for 200 iterations, each providing different sequences of hidden states that reflect the uncertainty inherent in estimating these states based on the available data. From these samples, we excluded chromosome X and segments (from any ancestry) that are 0.05cM or shorter. We then counted the number of windows assigned to Denisovan (or Neandertal) segments divided by the total number of windows assigned to any ancestry, providing an estimate of the Denisovan (or Neandertal) ancestry proportion. As this estimate is based on segments longer than 0.05cM, we expect it to be an underestimate of the true proportion.

While estimates of the Neandertal ancestry proportion were unchanged by the number of Denisovan genomes used in the reference set (**Figure S82**), adding a second Denisovan genome increased the Denisovan ancestry proportion estimated by *admixfrog* by an average of 1.8-fold (**Figure S83**). This is reflected in the number of bases covered by the archaic segments detected in each genome (**Figure S84** and **Figure S85**). As an example, Denisovan ancestry in the genomes of Australians, Papuans, and Bougainville islanders increased from an average of 25Mb using *Denisova 3* to 41Mb using both *Denisova 3* and *Denisova 25* (**Table S66**).

We then repeated the match rate analysis, first excluding derived alleles that are seen in Africans to be consistent with the analysis of the segments identified with *Sprime* and *hmmix*, and then counting the number of derived alleles in each segment that are shared with an archaic genome (counting heterozygous sites in the archaic genome as half). Adding a second Denisovan genome in the reference set increased the number of Denisovan segments that match the *Denisova 3* and *Denisova 25* genomes at approximately equal rates (mostly segments with match rates lower than 60%), and, to a lower extent, the number of segments that are slightly closer to *Denisova 25* (**Figure S86**). Few segments were lost by adding *Denisova 25* to the reference compared to only using *Denisova 3*. This analysis also shows that individuals from all geographical groups carry more segments that are closer to *Denisova 3* than to *Denisova 25*, even if we restrict the analysis to transversion polymorphisms (to avoid any residual ancient DNA damage) (**Figure S87**), replicating the results obtained with the segments identified with both *Sprime* and *hmmix*. The segments identified by *admixfrog* also capture the Denisovan component in East Asia, Central Asia/Siberia, and America with an ~80% match rate to *Denisova 3* and ~60% to *Denisova 25*, as detected with the other methods.

*D*-statistics computed in the genomic regions that overlap Denisovan segments were also significantly greater than 0 for multiple individuals in all geographical groups (**Figure S88**), aligning with all the results above that at least some of the Denisovan ancestry in all present-day non-Africans is closer to *Denisova 3*.

**Table S66: Average Denisovan ancestry coverage breadth per SGDP genome by geographic region.**

| **Region** | **Denisovan ancestry coverage breadth (Mb)** | |
| --- | --- | --- |
|  | **using *Denisova 3* only** | **Using both *Denisova 3* and *25*** |
| America | 0.70 | 1.74 |
| Central Asia/Siberia | 1.02 | 1.69 |
| West Eurasia | 0.26 | 0.37 |
| Oceania | 19.25 | 31.38 |
| Oceania (Australia, Papua New Guinea, Bougainville) | 24.87 | 40.65 |
| South Asia | 1.18 | 1.76 |
| East Asia | 1.33 | 1.85 |

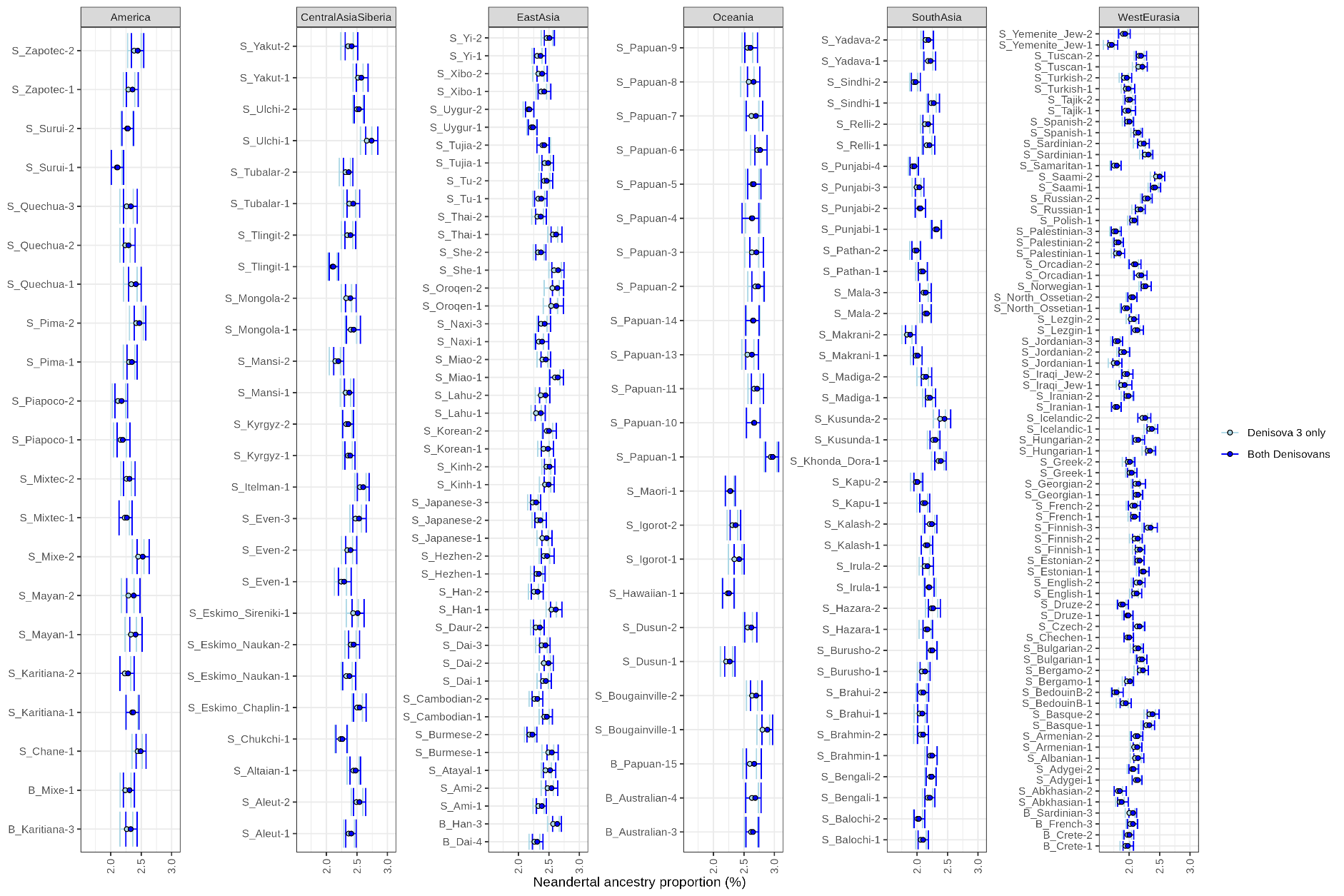

**Figure S82: Neandertal ancestry proportion estimated with admixfrog in the genomes of present-day non-Africans.** Each point corresponds to the average Neandertal ancestry proportion of an individual genome of the SGDP dataset, computed from 200 samples of the empirical-Bayes posterior. The bars correspond to the interval between the minimum and maximum estimates. The different colours correspond to the estimates including one or two Denisovan genomes in the reference set.

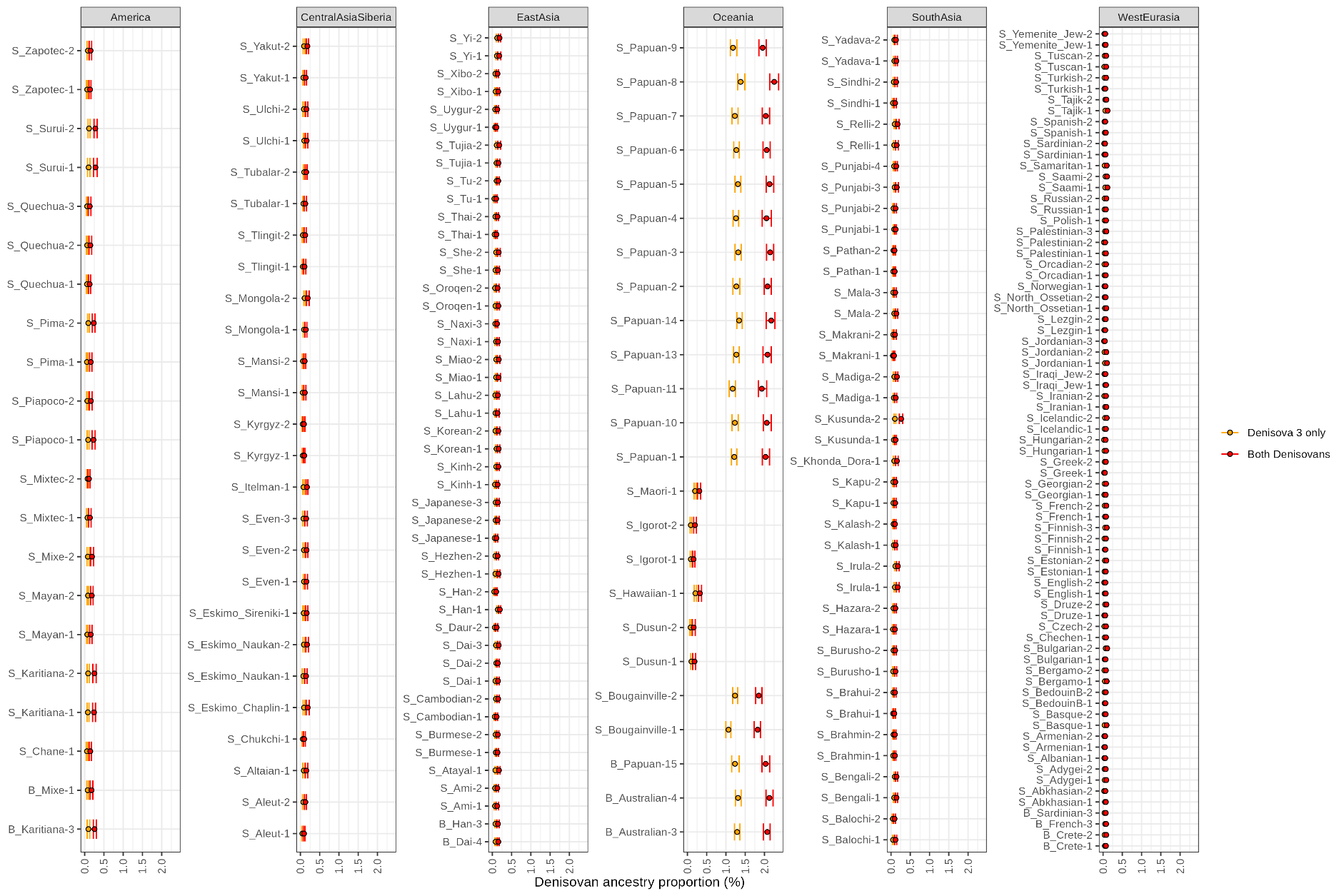

**Figure S83: Denisovan ancestry proportion estimated with admixfrog in the genomes of present-day non-Africans.** Each point corresponds to the average Denisovan ancestry proportion of an individual genome of the SGDP dataset, computed from 200 samples of the empirical-Bayes posterior. The bars correspond to the interval between the minimum and maximum estimates. The different colours correspond to the estimates including one or two Denisovan genomes in the reference set.

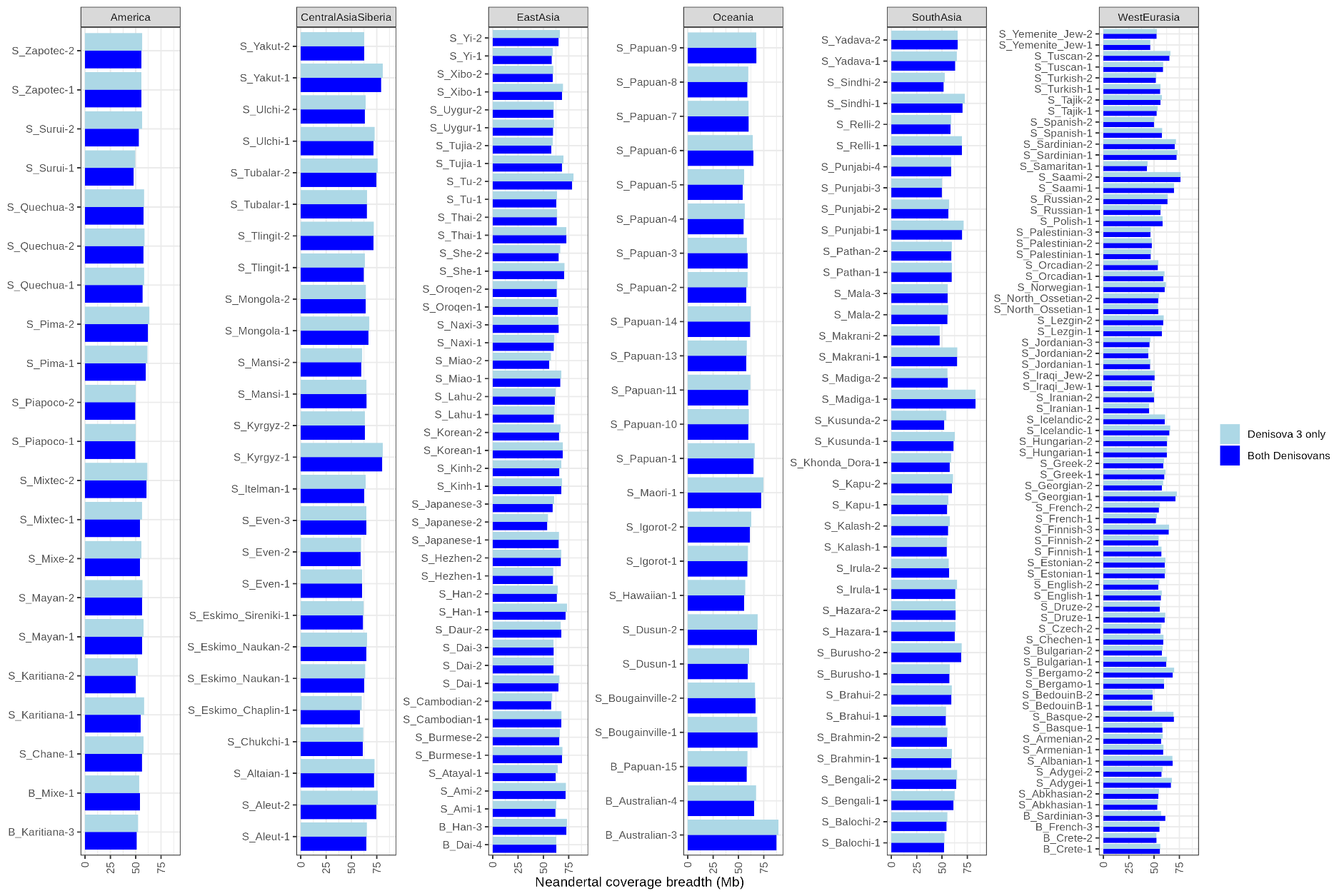

**Figure S84: Coverage breadth of putative Neandertal segments identified with admixfrog in the genomes of present-day non-Africans.** The coverage breadth here is the total number of bases in the genome covered by at least one Neandertal segment (≥0.05cM). The different colours correspond to the Neandertal segments identified when including one or two Denisovan genomes in the reference set.

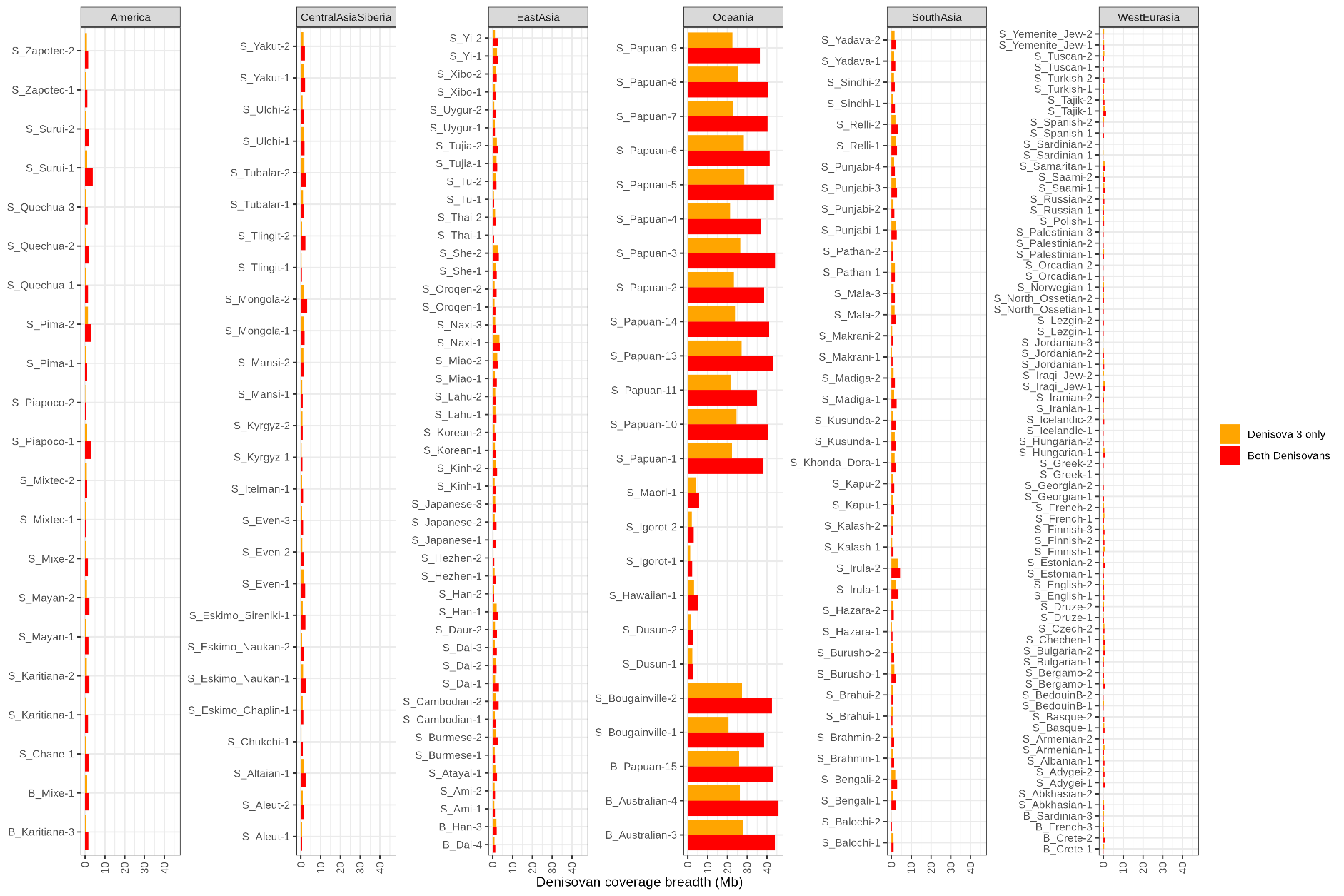

**Figure S85: Coverage breadth of putative Denisovan segments identified with admixfrog in the genomes of present-day non-Africans.** The coverage breadth here is the total number of bases in the genome covered by at least one Denisovan segment (≥0.05cM). The different colours correspond to the Denisovan segments identified when including one or two Denisovan genomes in the reference set.

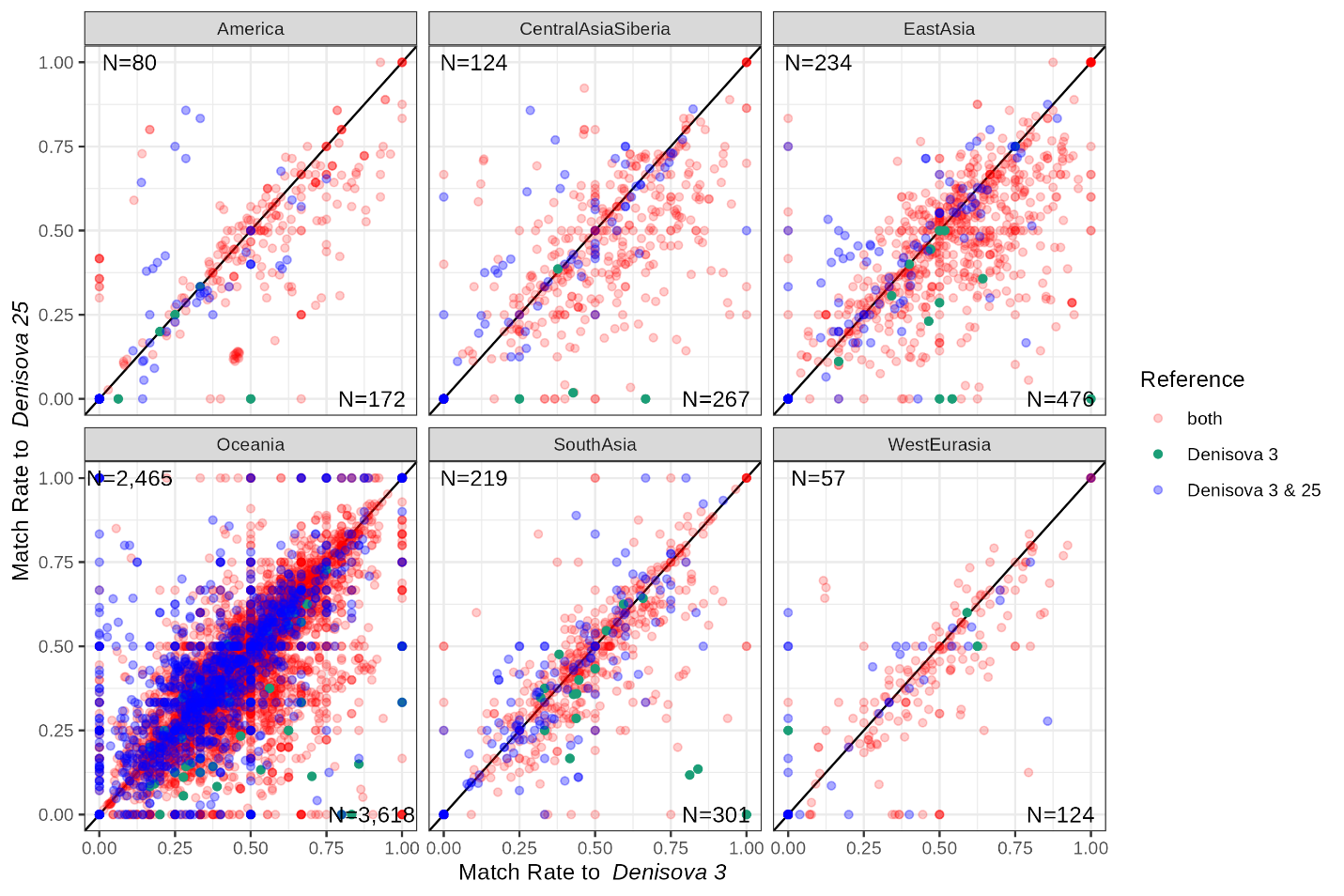

**Figure S86: Match rate of putative Denisovan segments, identified with admixfrog in the genomes of present-day non-Africans, to the Denisova 3 and Denisova 25 genomes.** Each point corresponds to a putative introgressed segment (≥ 0.05cM). The colours indicate whether the segments were identified using both reference sets (i.e. Denisova 3 only, and Denisova 3 & 25, in red) or only one reference set (i.e. Denisova 3 only, in green, or Denisova 3 & 25, in blue). When a segment was identified with both reference sets, we plot the match rates for the segments identified using two Denisovan genomes in the reference set. The numbers of segments below and above the diagonal (i.e. closer to Denisova 3 and Denisova 25, respectively) are indicated in the plots. Geographical assignments for the individuals are shown atop each panel.

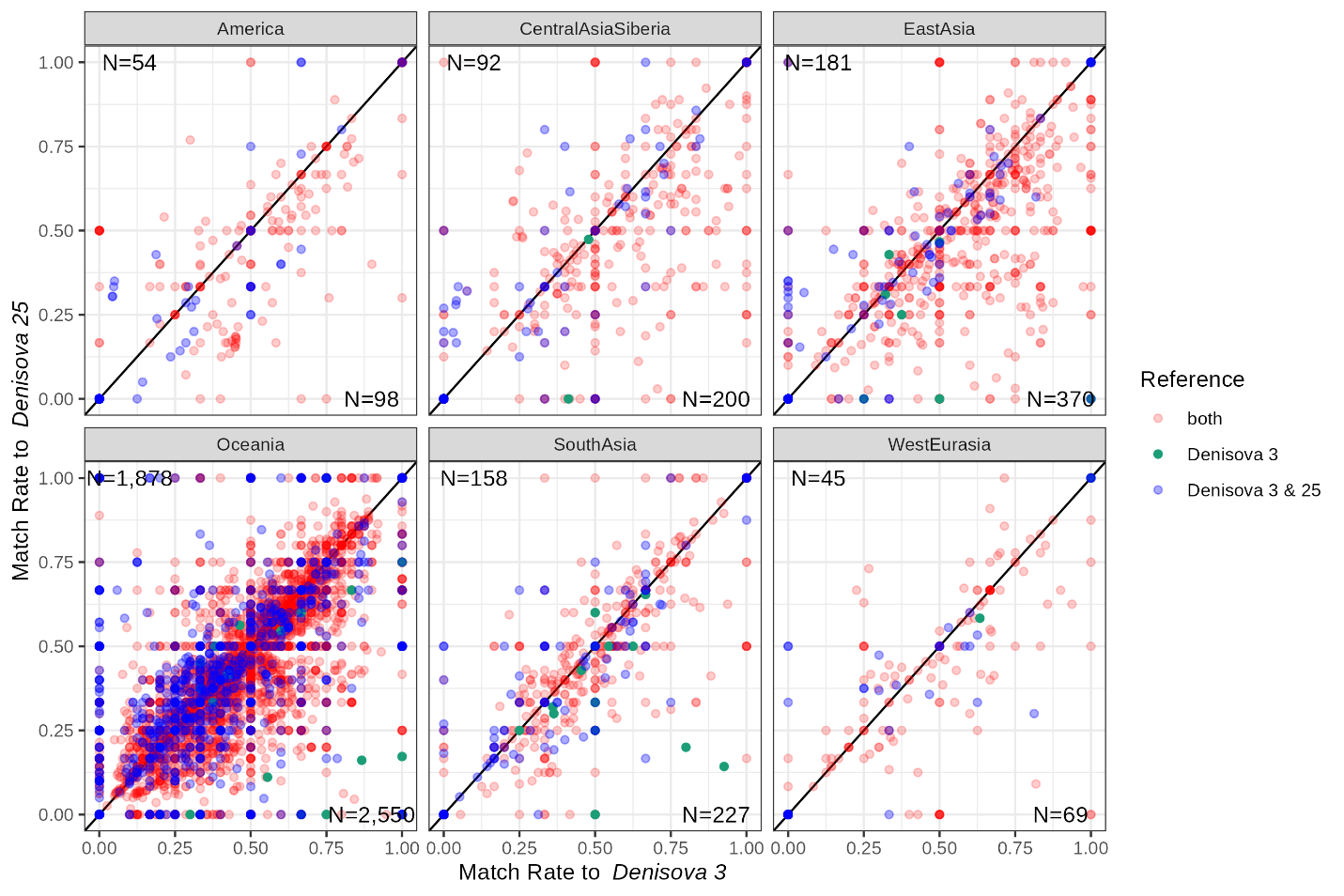

**Figure S87: Match rate of putative Denisovan segments (identified with admixfrog) to the Denisova 3 and Denisova 25 genomes using transversion polymorphisms only.** See **Figure S86** for more details.

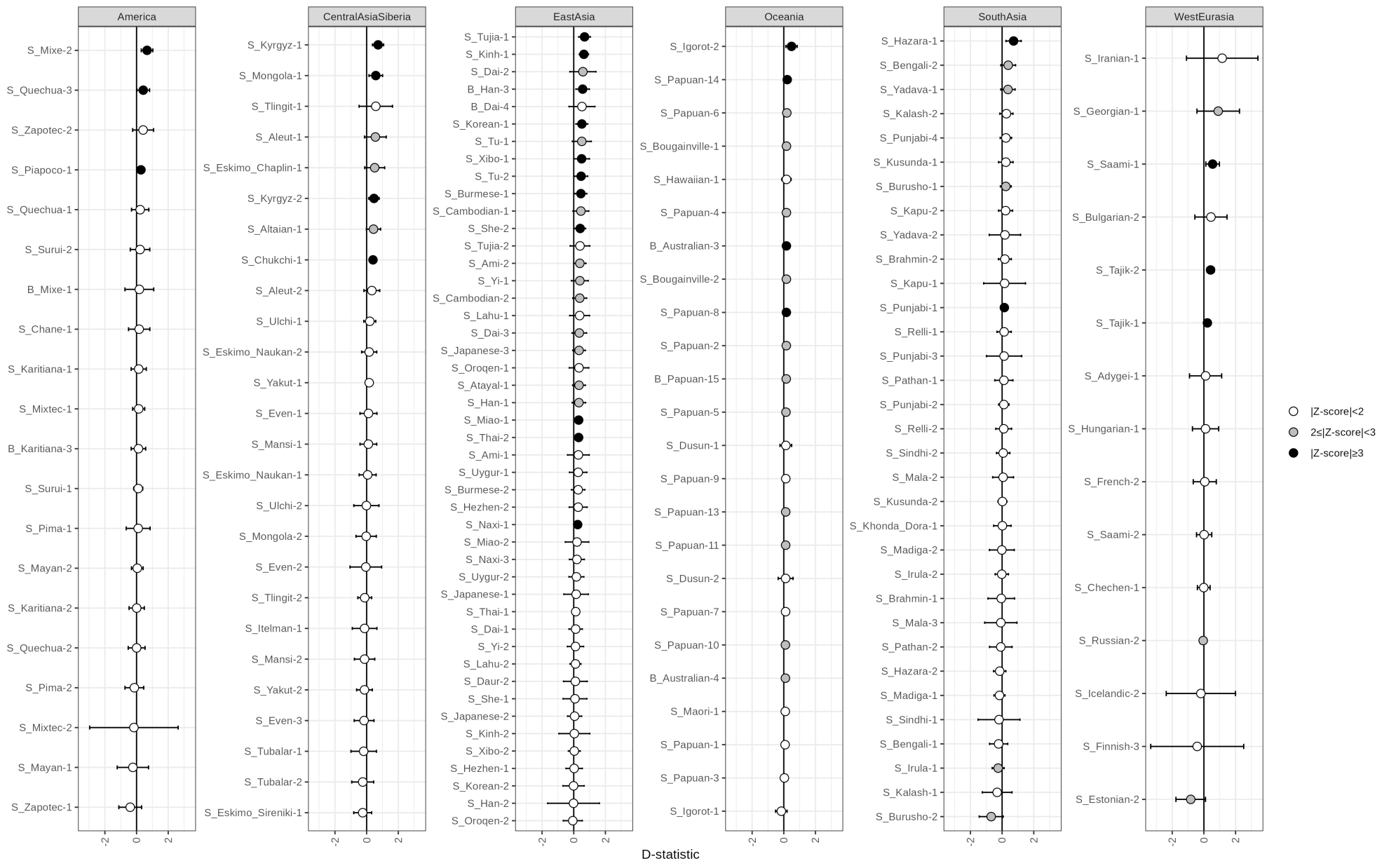

**Figure S88: D-statistics comparing how the Denisovan ancestry of present-day non-Africans relates to Denisova 3 and Denisova 25.** D-statistics were calculated using ADMIXTOOLS^108^ as implemented in admixr^109^ in regions overlapping Denisovan segments of each individual, identified with admixfrog using a reference with two Denisovan genomes. Positive values correspond to a closer relationship to Denisova 3. The genomes of four great apes were used to determine the ancestral allele. Whiskers on each side of the plotted D values correspond to three standard errors calculated using a Weighted Block Jackknife and a block size of 5Mb across all autosomes. Colours indicate different significance levels. For some individuals, particularly in West Eurasia, there were not enough segments to compute D-statistics with admixr.

#### Analysis of archaic segment length distribution and estimates of admixture times

Having identified new sets of Denisovan segments in present-day human genomes, we turned our attention to the length of these segments and ancestry linkage disequilibrium (ALD) curves to reassess the timing and number of gene flows with Denisovans.

##### Length of Denisovan segments

We first compared the average length of Neandertal and Denisovan segments within individual genomes. Surprisingly, in individuals from all populations, Denisovan segments identified with *admixfrog* or *hmmix* are shorter on average than Neandertal segments, both in terms of genetic and physical length (**Figure S89** to **Figure S92**). While this could suggest that at least some Denisovan segments were inherited earlier than Neandertal segments, differences in Neandertal and Denisovan ancestry, including differences in ancestry proportions and divergence from the reference genomes used to detect archaic segments (or assign them to a group), likely introduce biases into this comparison. Notably, there is not much difference in the length of Neandertal and Denisovan segments identified by *hmmix* in Oceanian genomes, as well as in other populations when the assignment of archaic segments to a group is stricter (described in a previous section; **Figure S93** and **Figure S94**), indicating potential biases in the assignment, differences between populations, or uncertainties in the estimates per individual.

Because our estimates per individual are subject to some uncertainty, particularly for Denisovan ancestry as it is rarer than Neandertal ancestry outside Oceania, we compared Denisovan and Neandertal segment lengths after aggregating segments from multiple individuals within the same geographical group. To ensure equitable comparison and avoid counting the same segments multiple times, we identified independent segments within each geographical group by clustering overlapping or directly adjacent segments and randomly selecting one segment from each cluster. This clustering process was performed with *bedtools cluster* after filtering out segments shorter than 0.05cM and down sampling the number of Neandertal segments to match that of Denisovan segments. This down sampling avoids biases that may arise from differences in segment counts, as more Neandertal segments may lead to more overlaps, which may affect long and short segments differently. Despite these precautions, Denisovan segments remained shorter on average than Neandertal segments (**Figure S95A**), except for the segments called with *hmmix* in Oceanians (**Figure S97A**). We also observe large differences in the length of segments called by *hmmix* and *admixfrog*, particularly for genetic lengths, for example, in Oceanians genetic length averages 0.08cM for *admixfrog* compared to 0.19cM for *hmmix* (for Neandertal segments: 0.09cM and 0.18cM, respectively). Moreover, differences in the length of Neandertal segments called with *admixfrog* between some populations likely point to inaccuracies in determining the length of these segments in present-day genomes.

The smaller number of Denisovan segments compared to Neandertal segments in most populations may be associated with a larger relative proportion of false positives, i.e. segments that are closer to the Denisovan or Neandertal genomes due to incomplete lineage sorting. This would not change after down-sampling the number of Neandertal segments. If these segments, which are generally shorter than genuine archaic segments, make up a larger proportion of the introgressed segments identified as Denisovan, this would lower the average length of Denisovan segments. To address this concern, we compared the average length of the 100 longest Denisovan segments to the 100 longest Neandertal segments within each geographical group. Since the number of Neandertal and Denisovan segments may still differ after clustering, we downsampled the larger set (usually the Neandertal set, except for the segments called with *hmmix* in Oceanians) to match the smaller set before comparison. Among these longest segments, the Denisovan segments identified with *admixfrog* remained shorter than the Neandertal segments (**Figure S95B**), while no differences were observed for the segments identified with *hmmix* (**Figure S97B**). We concluded that the observed differences in segment lengths are likely attributed to a higher incidence of false positives among the comparatively fewer Denisovan segments than Neandertal segments.

As the length of segments identified with *admixfrog* is sensitive to the penalty used to call segments, we checked whether the differences between the length of Neandertal and Denisovan segments as well as the inconsistencies in the length of Neandertal segments between populations are resolved using a lower penalty of 0.1. While Denisovan segments remain shorter than Neandertal segments (**Figure S96A**), the confidence intervals for the average length of the 100 longest Denisovan segments and 100 longest Neandertal segments overlap (**Figure S96B**), suggesting that at least some of the length differences are caused by the penalty used and the difficulty in determining the exact length of segments.

##### Admixture linkage disequilibrium curves

The analyses presented here indicate that difficulty with accurately determining the length of archaic segments using both reference-based and reference-free segment detection methods. However, previous work has shown that using admixture linkage disequilibrium decay provides a robust alternative for estimating admixture time from introgressed segment length^131,132^. Linkage disequilibrium decay was previously used to estimate when the ancestors of present-day Oceanians admixed with Denisovans^133^. The authors first identified a set of single nucleotide polymorphisms (SNPs) in the test population where a single randomly chosen allele from the *Denisova 3* genome is derived relative to the human-chimpanzee ancestor, while a single randomly chosen allele from the *Denisova 5* Neandertal genome is ancestral, and all alleles in a panel of sub-Saharan African genomes (44 high-coverage genomes sequenced as part of the Simons Genome Diversity Project, SGDP^106^) are ancestral. Similarly, SNPs with putative Neandertal introgressed alleles were ascertained by identifying sites where the *Denisova 5* Neandertal is derived, *Denisova 3* is ancestral, and all sub-Saharan Africans are ancestral. They then computed the average of Lewontin’s D in the target population for all pairs of ascertained SNPs at a given genetic distance x. By fitting an exponential function to this curve as a function of x, using ordinary least squares within the range of 0.02cM to 1cM, they estimated the time of gene flow from the rate of exponential decay. To estimate standard errors, they used a weighted Block Jackknife approach with 10 Mb blocks, each containing a minimum of 100 SNPs. The dataset included the genomes from 16 Papuans, 2 Native Australians and 2 Bougainville islanders as the target population, with genetic distances measured using the combined Oxford LD-based map^134^.

We repeated this analysis using the African-American recombination map^99^, 40 African genomes from the SGDP dataset instead of 44 (as we excluded North African genomes), and 14 Papuans that are publicly available, instead of the 16 Papuan genomes previously used, and replicated the results obtained by Sankararaman *et al*., i.e. a Denisovan admixture time about 10% more recent than the Neandertal admixture time, at ~1,000 and ~1,100 generations ago, respectively (***Figure S99***). Ascertaining SNPs using *Denisova 25* instead of *Denisova 3* also yielded similar estimates (***Figure S99***). However, as reported previously, these estimates are biased, likely due to errors in the genetic map that can produce systematic underestimates of dates inferred based on linkage disequilibrium. Since population-specific fine-scale recombination maps are now available for the worldwide populations of the 1,000 Genomes dataset^135^, we re-analyzed the admixture linkage disequilibrium decay in the 1,000 Genomes populations (phase 3)^129^, using the same ascertainment strategy described above while measuring genetic distance based on the respective recombination map available for each population^135^. This yielded Neandertal admixture time estimates that are consistent with previous estimates^83,132,136,137^, except for American populations where the estimates are older (***Figure S100***). This discrepancy likely represents uncertainties in inferring recombination rates in these admixed populations, as similar biases were observed when using the recombination map inferred in American populations to measure genetic distance in other populations. A similar trend was observed for the Denisovan admixture time, with estimates ranging between 1,146 and 1,705 generations ago (33ka-49ka) in most populations, and between 2,069 and 2,469 generations ago in American populations (***Figure S100***). Interestingly, Denisovan admixture time estimates in European populations are similar to those obtained in other non-African populations, supporting the presence of Denisovan ancestry in Europeans. In comparison, Denisovan admixture time estimates in African populations from the 1,000 Genomes dataset, which were not used in the ascertainment, differ substantially between populations, ranging from 973 to 2,443 generations ago, usually exceeding the estimates for Neandertal admixture (**Figure S101**).

Neandertal admixture time estimates obtained using SNPs identified using *Denisova 25* and the *Denisova 5* Neandertal genome remain almost the same as those obtained using *Denisova 3*, while Denisovan admixture time estimates increase by a few hundred generations, ranging from 1,398 to 1,974 generations ago in Eurasian populations. If we focus on SNPs for which at least one of the two Denisovan genomes carries an allele that is absent among the three Neandertal genomes (ensuring that a genotype is available for at least one of them), the estimates are more similar to those obtained with *Denisova 3*, suggesting that there may be slightly more sharing of derived alleles due to incomplete lineage sorting when using the older *Denisova 25* genome, leading to estimates that are slightly older.

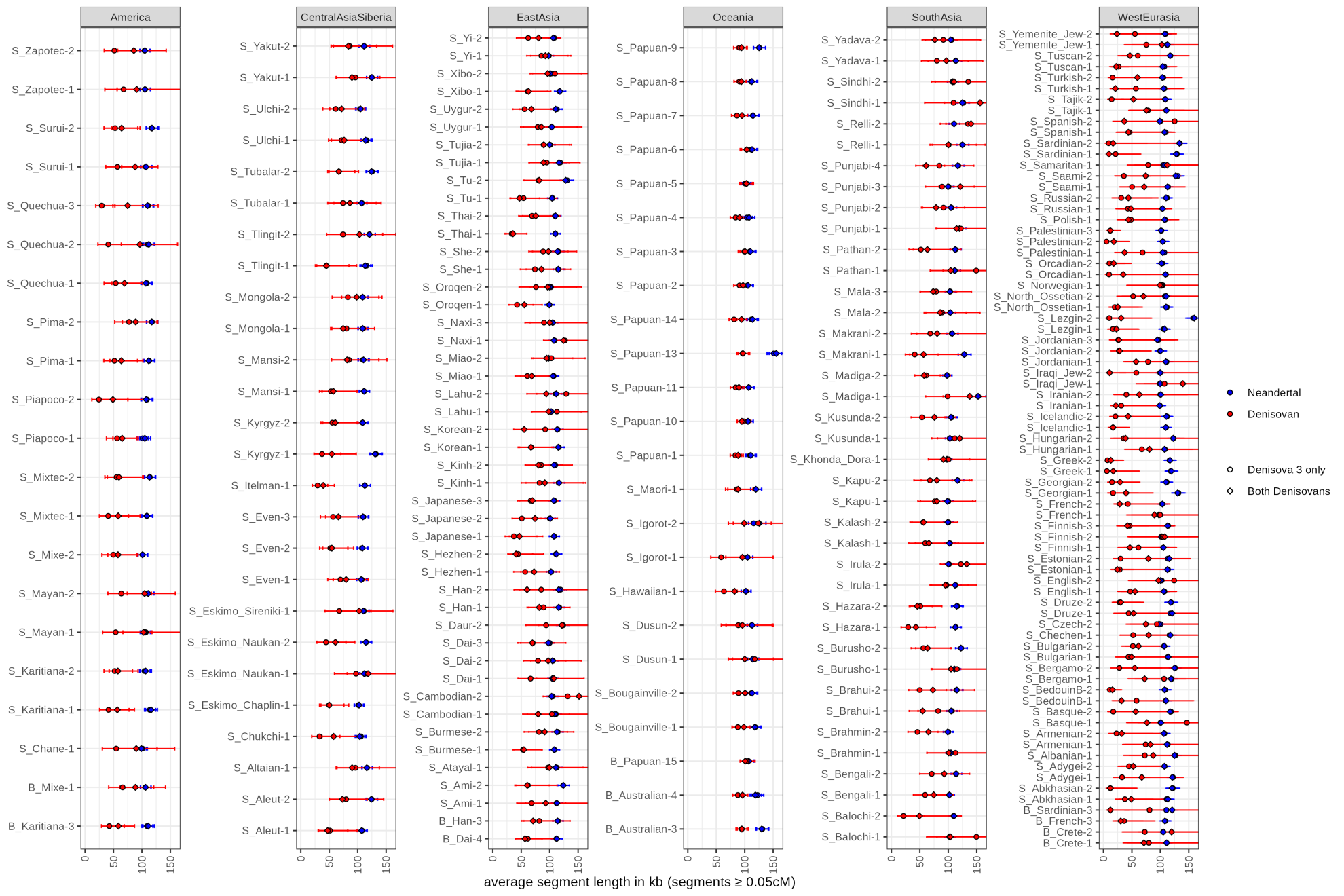

**Figure S89: Per individual average physical length of archaic segments identified with admixfrog.** The shape of the points corresponds to different **r**eferences, using one or two Denisovan genomes. The average length was computed after filtering out segments shorter than 0.05cM. Whiskers correspond to the 95% confidence intervals of the average length computed from the quantiles of a gamma distribution (with shape and rate parameters equal to the number of segments) divided by the average length.

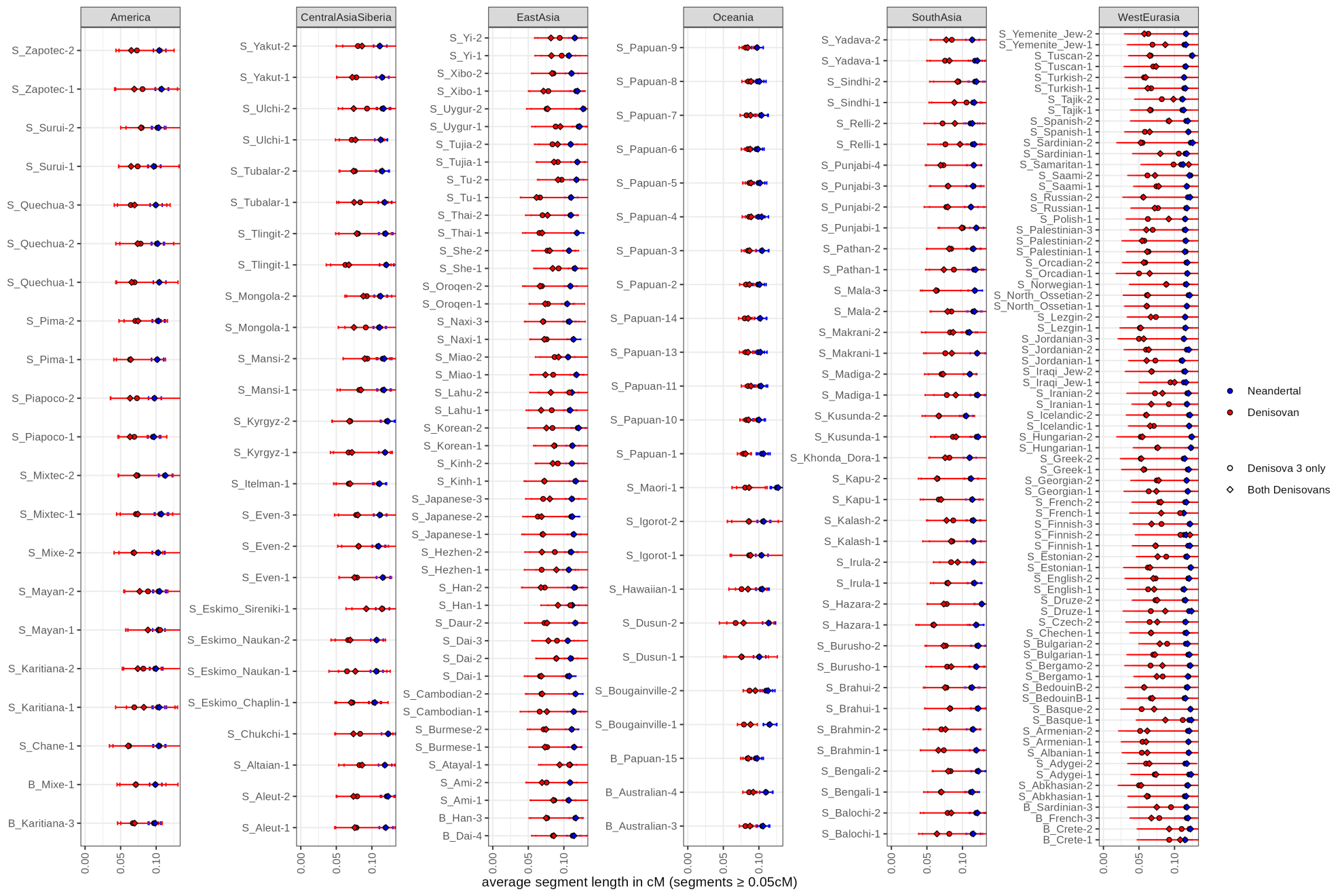

**Figure S90: Per individual average genetic length of archaic segments identified with admixfrog.** See **Figure S89** for additional details.

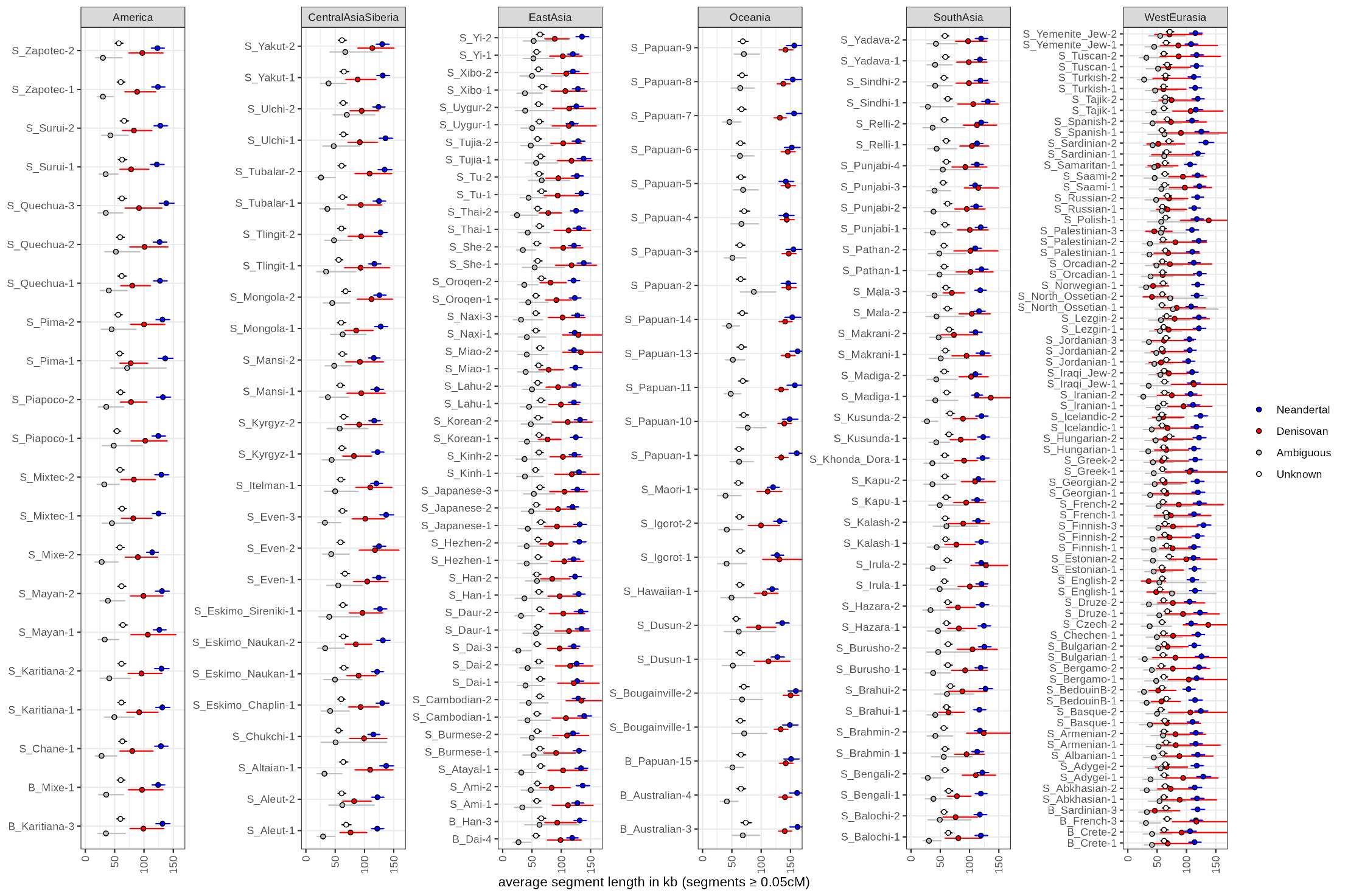

**Figure S91: Per individual average physical length of archaic segments identified with hmmix.** Unknown segments are those that do not share enough alleles with the available Neandertal and Denisovan genomes to assign them to a group, while ambiguous segments are those that match equally well the Neandertal and Denisovan genomes. The average length was computed after filtering out segments shorter than 0.05cM. Whiskers correspond to the 95% confidence intervals of the average length computed from the quantiles of a gamma distribution (with shape and rate parameters equal to the number of segments) divided by the average length.

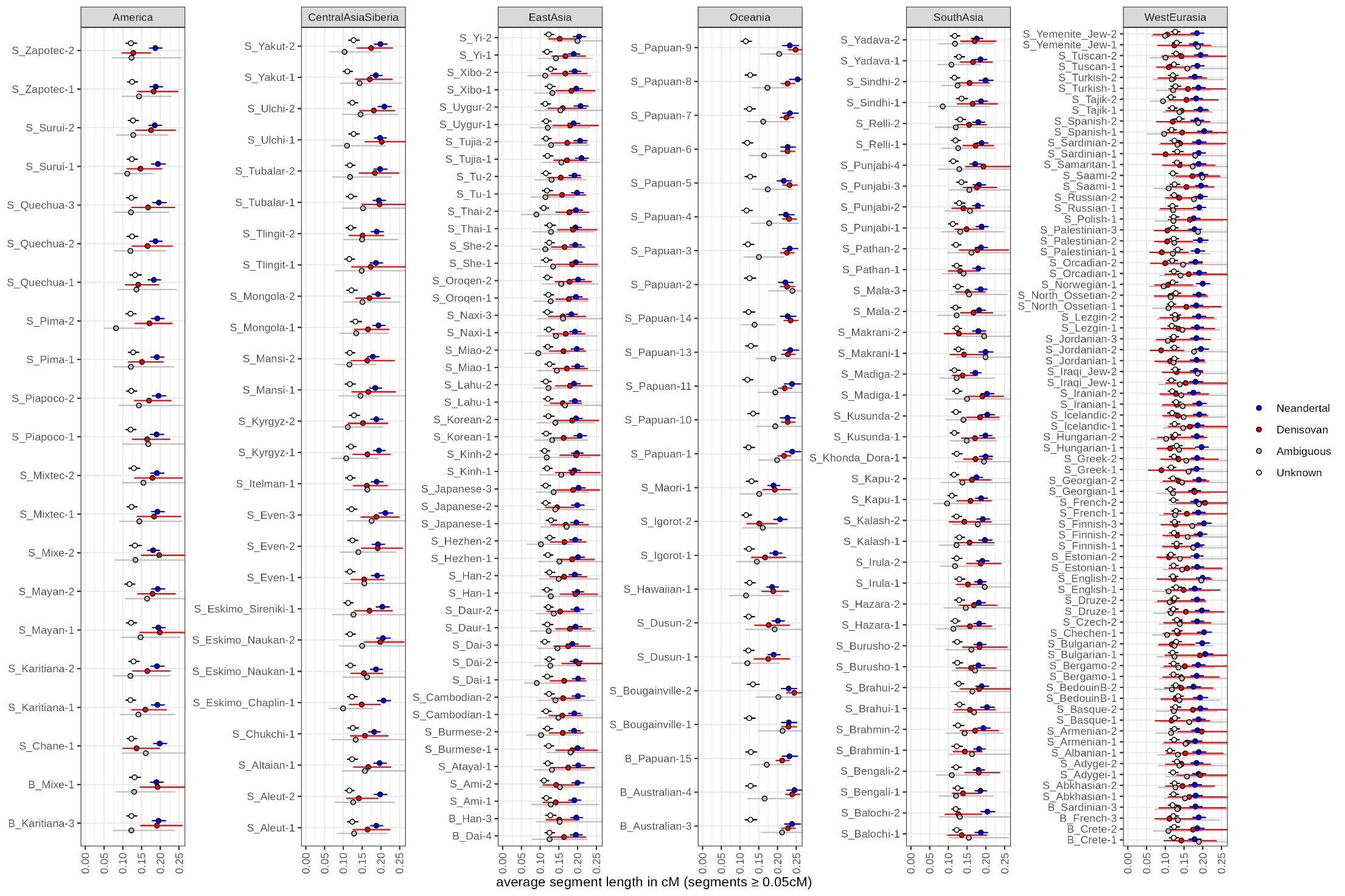

**Figure S92: Per individual average genetic length of archaic segments identified with hmmix.** See **Figure S91** for additional details.

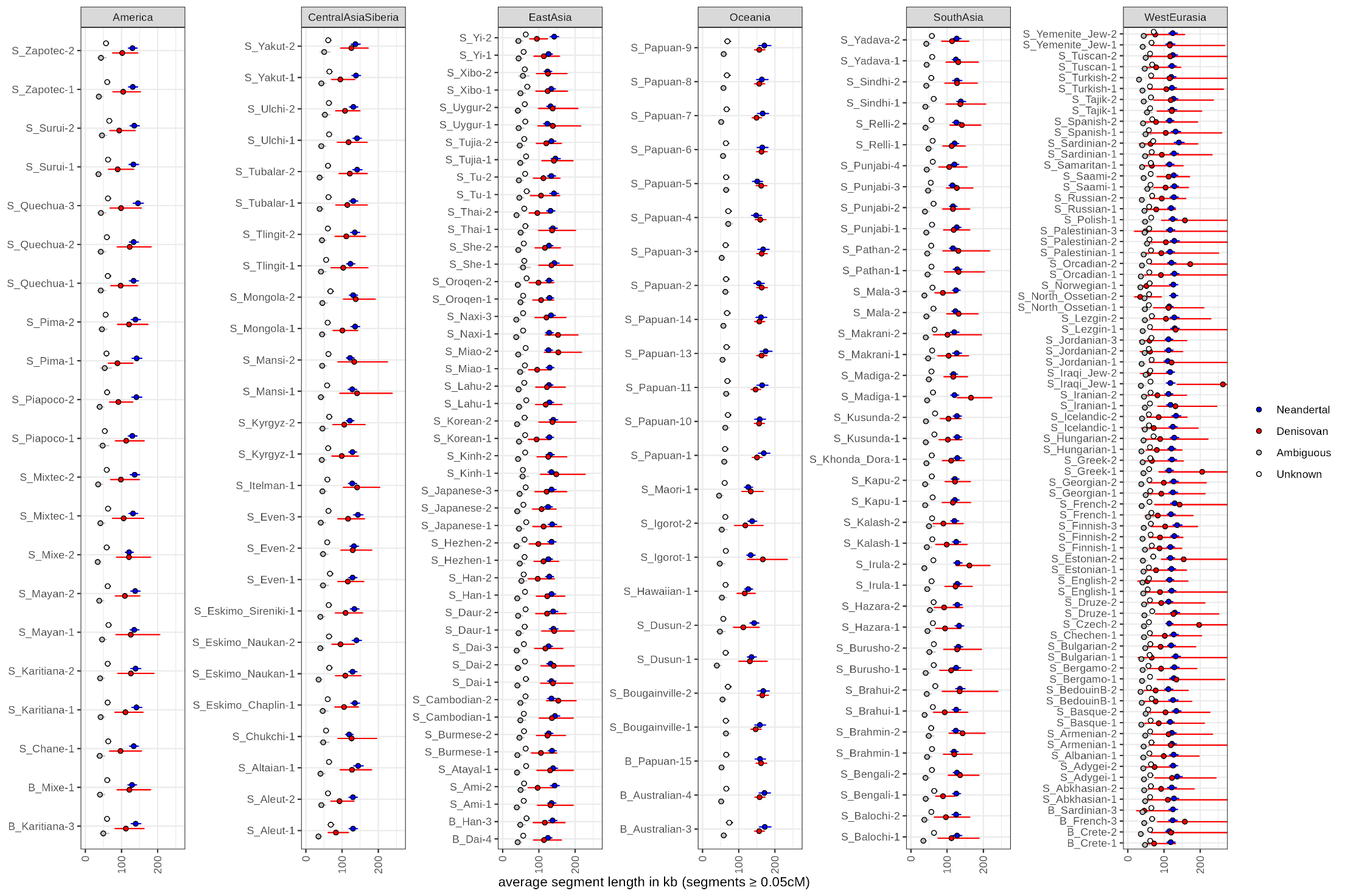

**Figure S93: Per individual average physical length of archaic segments identified with hmmix and more confidently assigned to Neandertals and Denisovans.** See **Figure S91** for additional details.

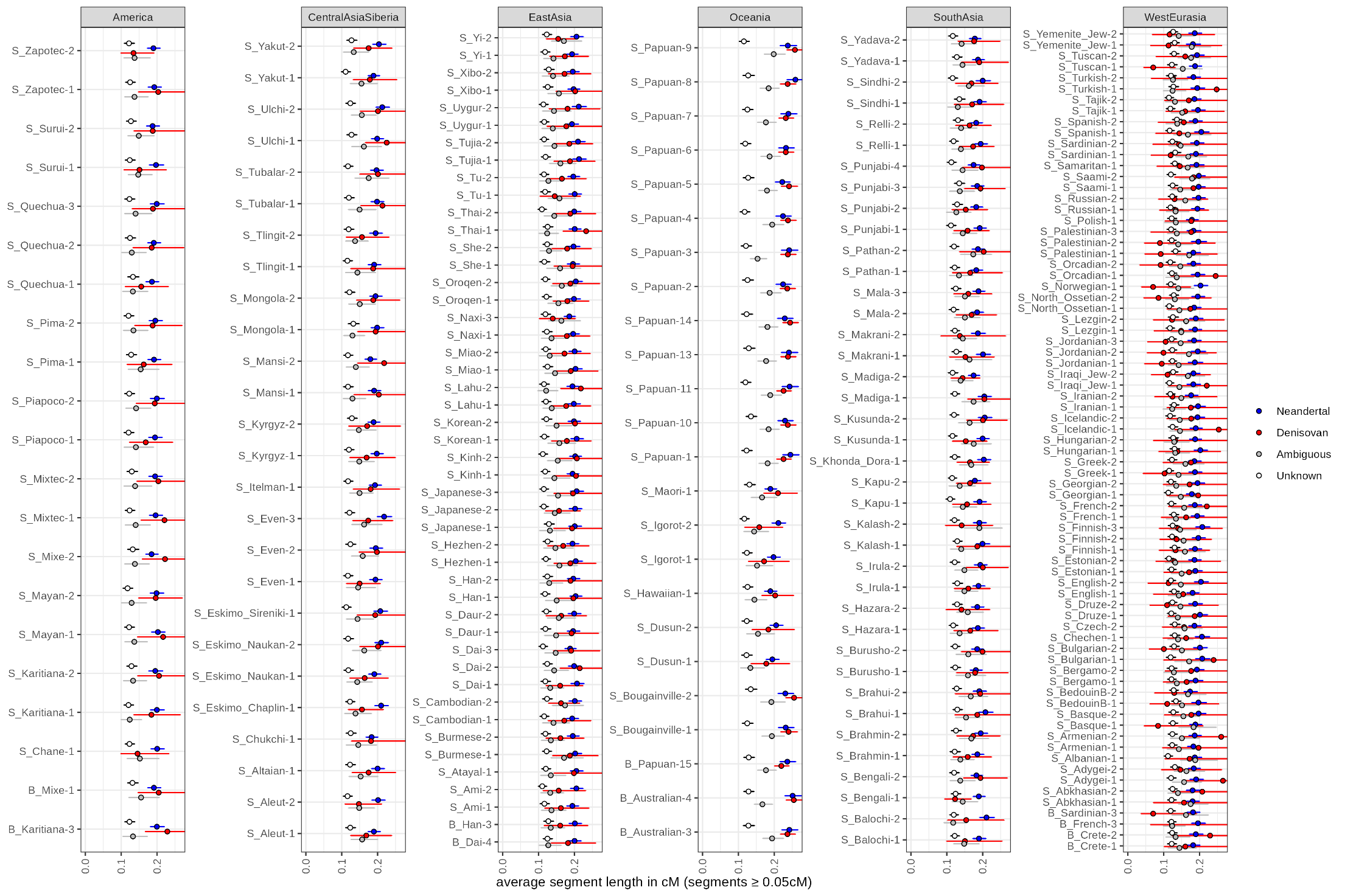

**Figure S94: Per individual average genetic length of archaic segments identified with hmmix and more confidently assigned to Neandertals and Denisovans.** See **Figure S91** for additional details.

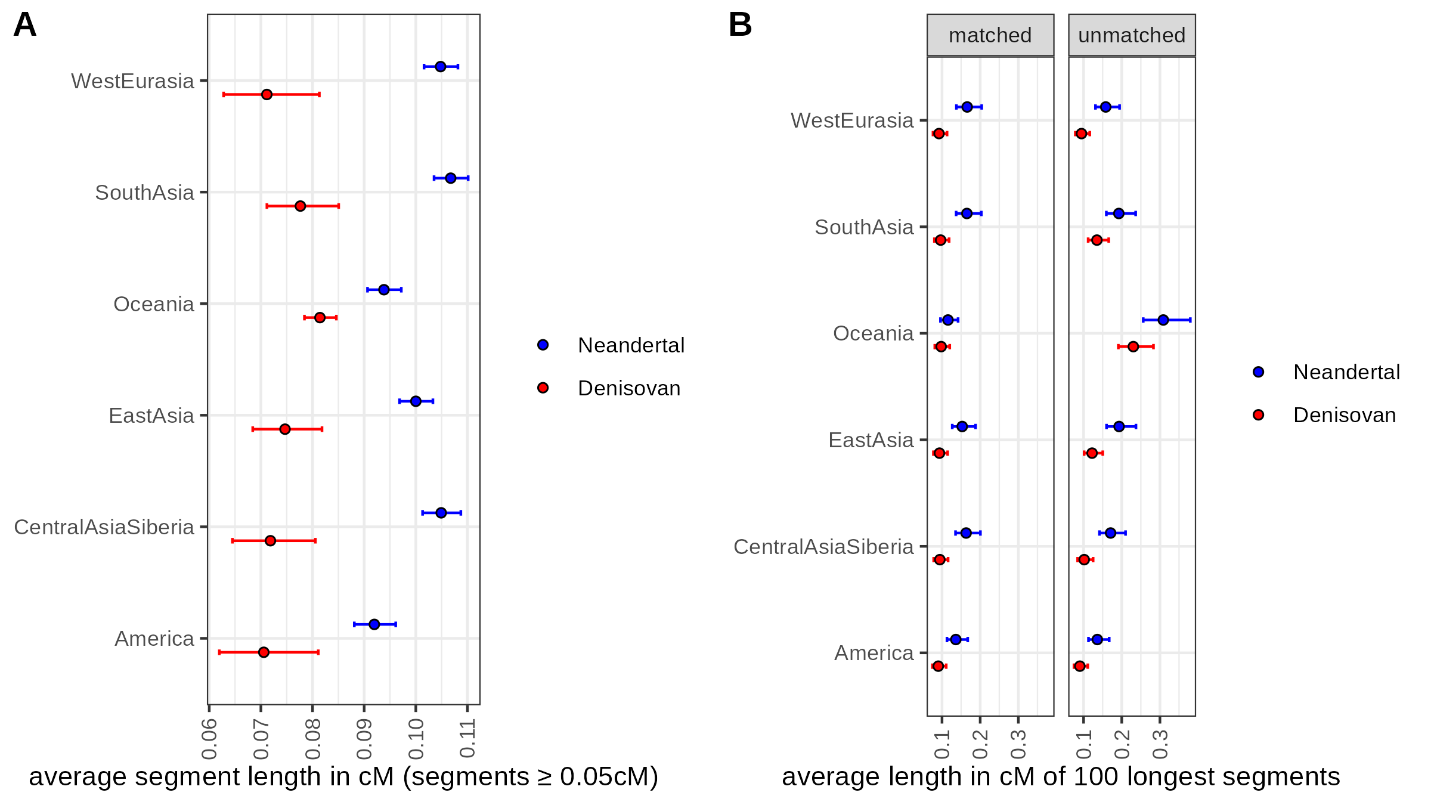

**Figure S95: Per population average genetic length of archaic segments identified with admixfrog.** The average length was computed from an aggregated and independent set of segments (identified using the reference set including two Denisovan genomes) as described above. Whiskers correspond to the 95% confidence intervals of the average length computed from the quantiles of a gamma distribution (with shape and rate parameters equal to the number of segments) divided by the average length. Panel A shows the average for all segments, whereas panel B shows the average for the 100 longest segments, after matching the number of Neandertal and Denisovan segments in all populations (matched) or each population independently (unmatched).

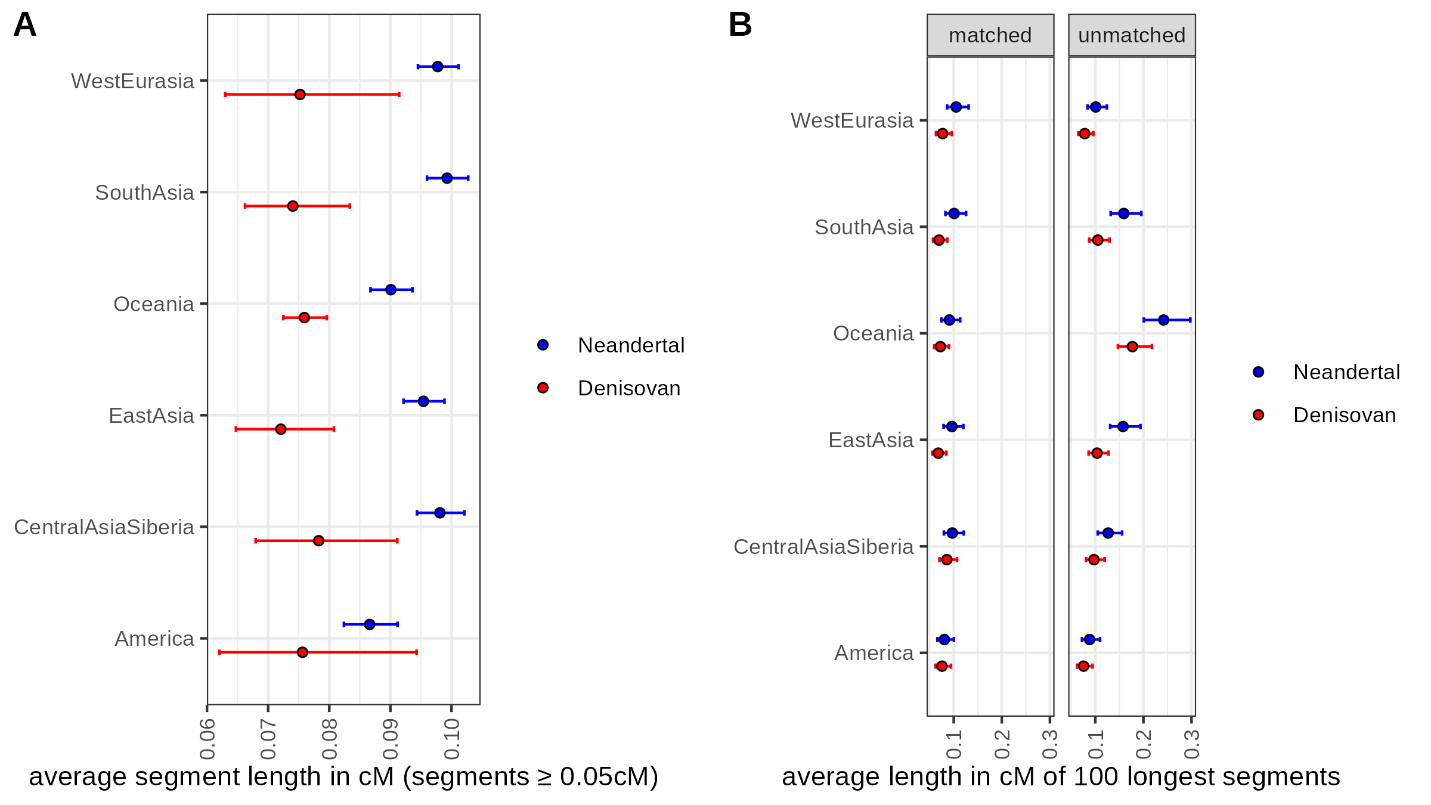

**Figure S96: Per population average genetic length of archaic segments identified with admixfrog using a penalty of 0.1.** See **Figure S95** for additional details.

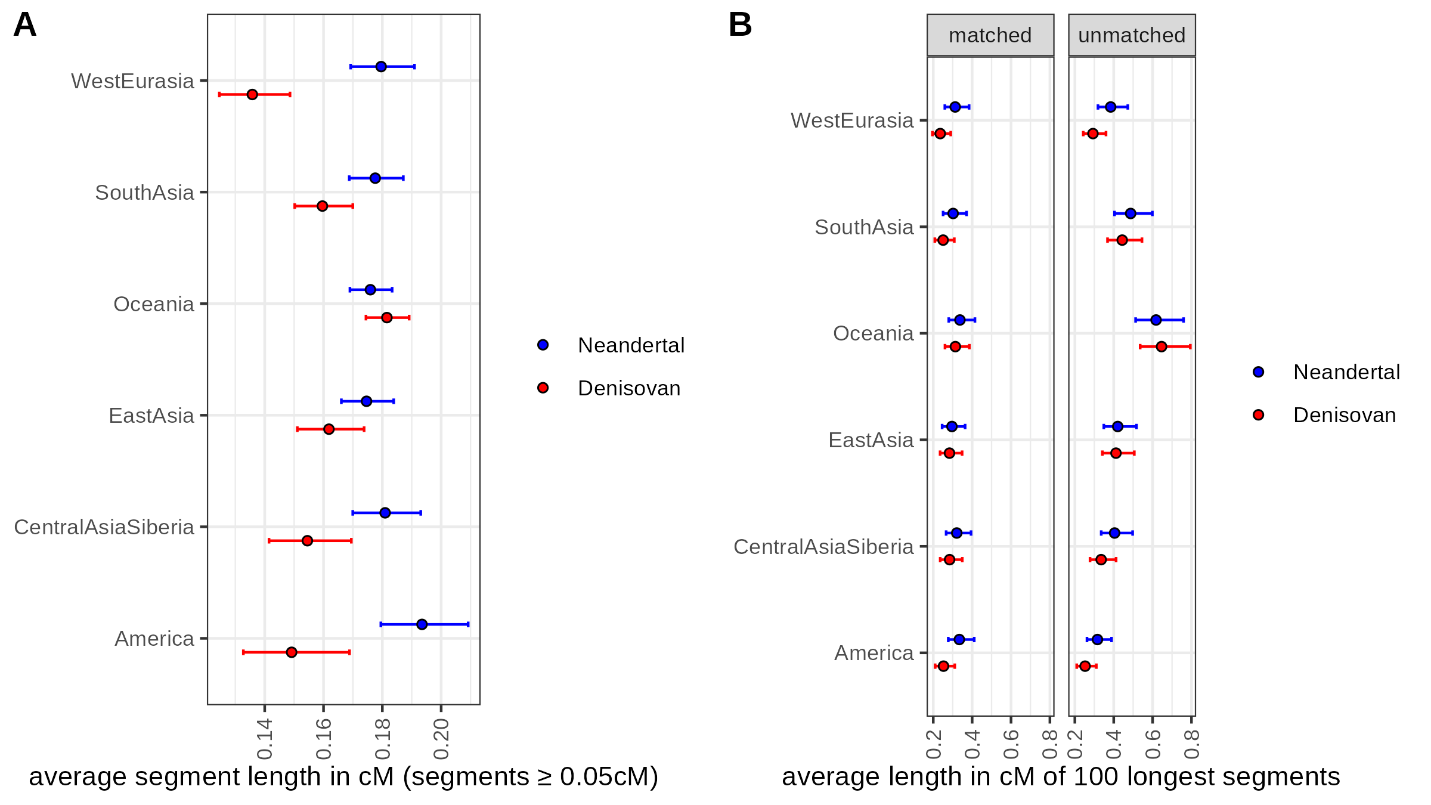

**Figure S97: Per population average genetic length of archaic segments identified with hmmix.** See **Figure S95** for additional details.

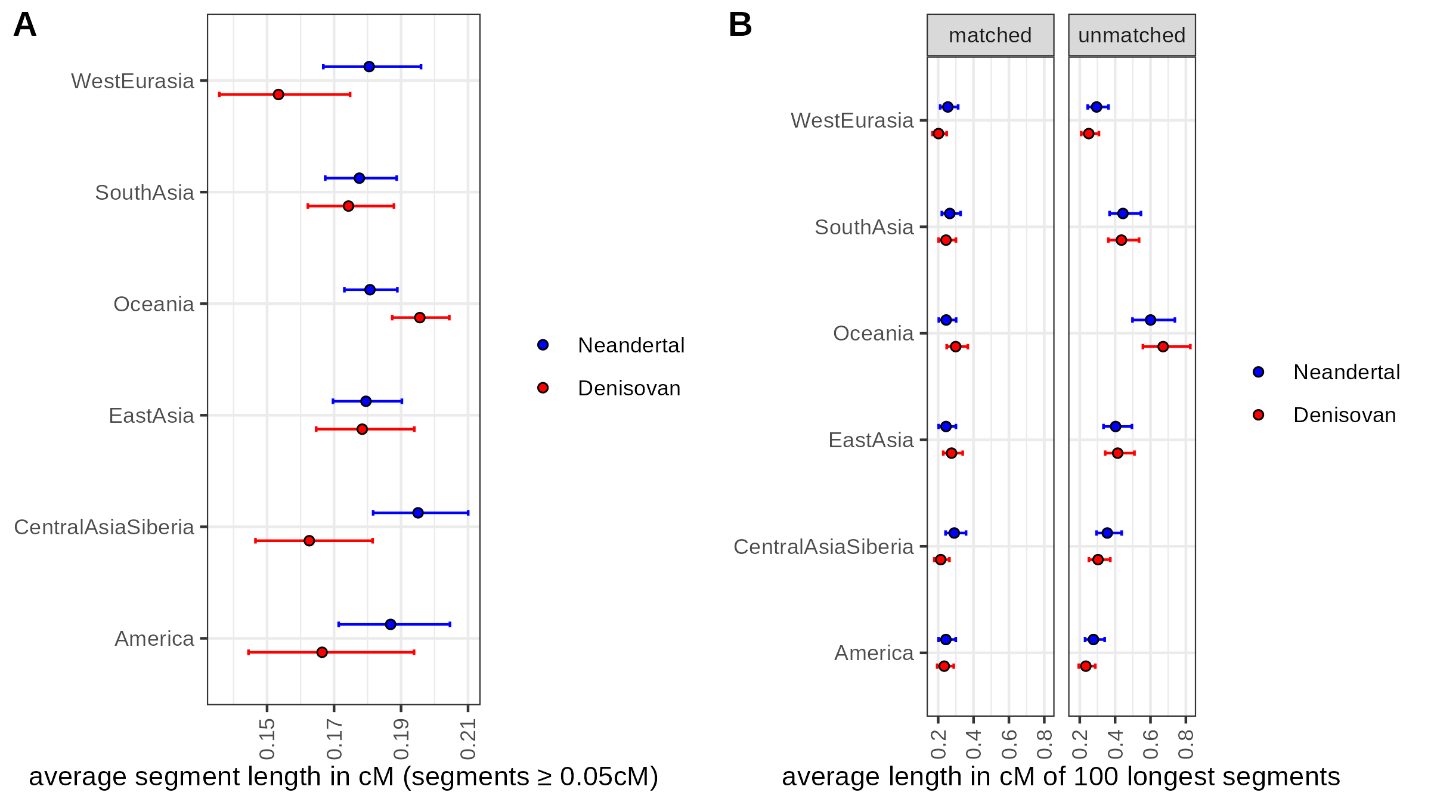

**Figure S98: Per population average genetic length of archaic segments identified with hmmix and more confidently assigned to Neandertals and Denisovans.** See **Figure S95** for additional details.

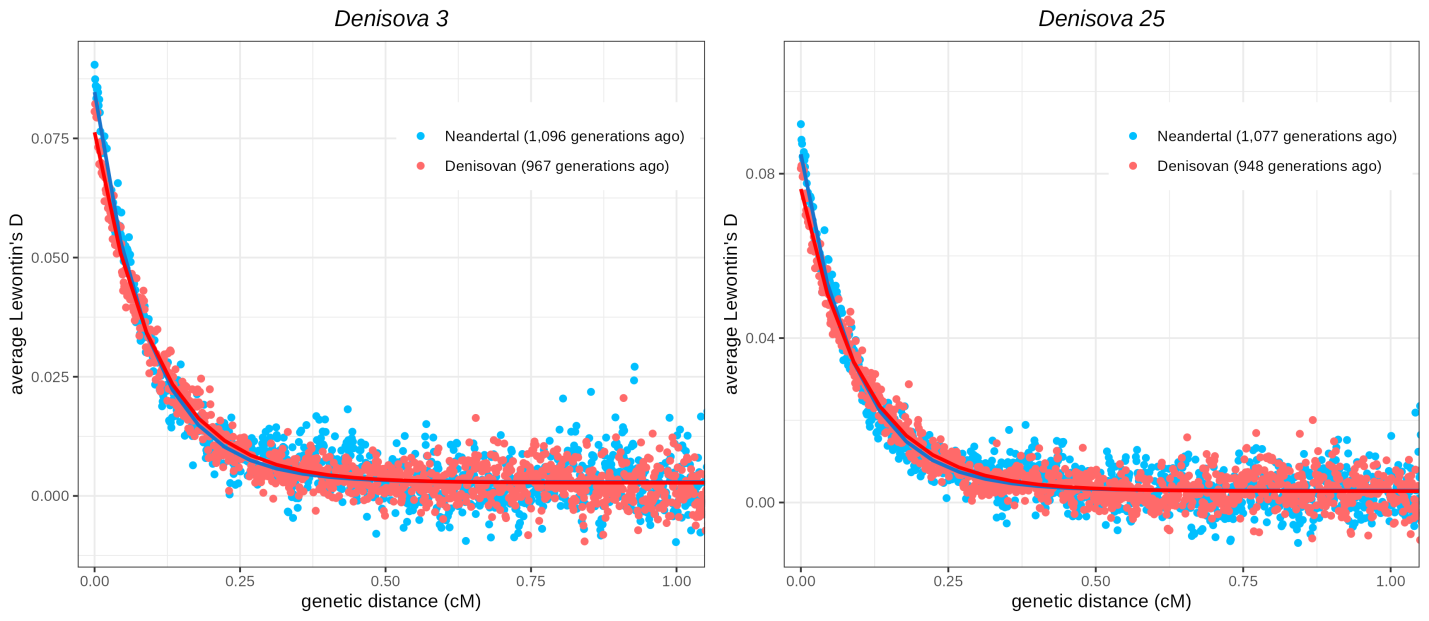

**Figure S99: Linkage disequilibrium decay between putative Neandertal or Denisovan introgressed alleles in present-day Oceanians.** Linkage disequilibrium was measured in a set of Oceanian genomes from the SGDP dataset^106^ (14 Papuans, 2 Australians and 2 Bougainville islanders) using the implementation of Lewontin’s D in PLINK^138^ for pairs of putative archaic variants that are less than 1Mb apart and less than 10 single nucleotide polymorphisms apart. We then computed the average in bins of 0.001cM. Putative Denisovan and Neandertal variants were identified as those that differ between the Denisova 3 (left) or Denisova 25 (right) and Denisova 5 (Neandertal) genomes (after randomly sampling one allele at heterozygous positions), and that are absent in African genomes from the SGDP dataset^106^ and great ape genomes (panTro4, panpan1.1, gorGor3, ponAbe2). Genetic distances were computed using the African-American recombination map^99^. We fit an exponential curve (using the Nonlinear Least Square function in R) after excluding the pairs that are less than 0.02cM apart and report the estimated date of admixture for both Neandertal and Denisovan introgression.

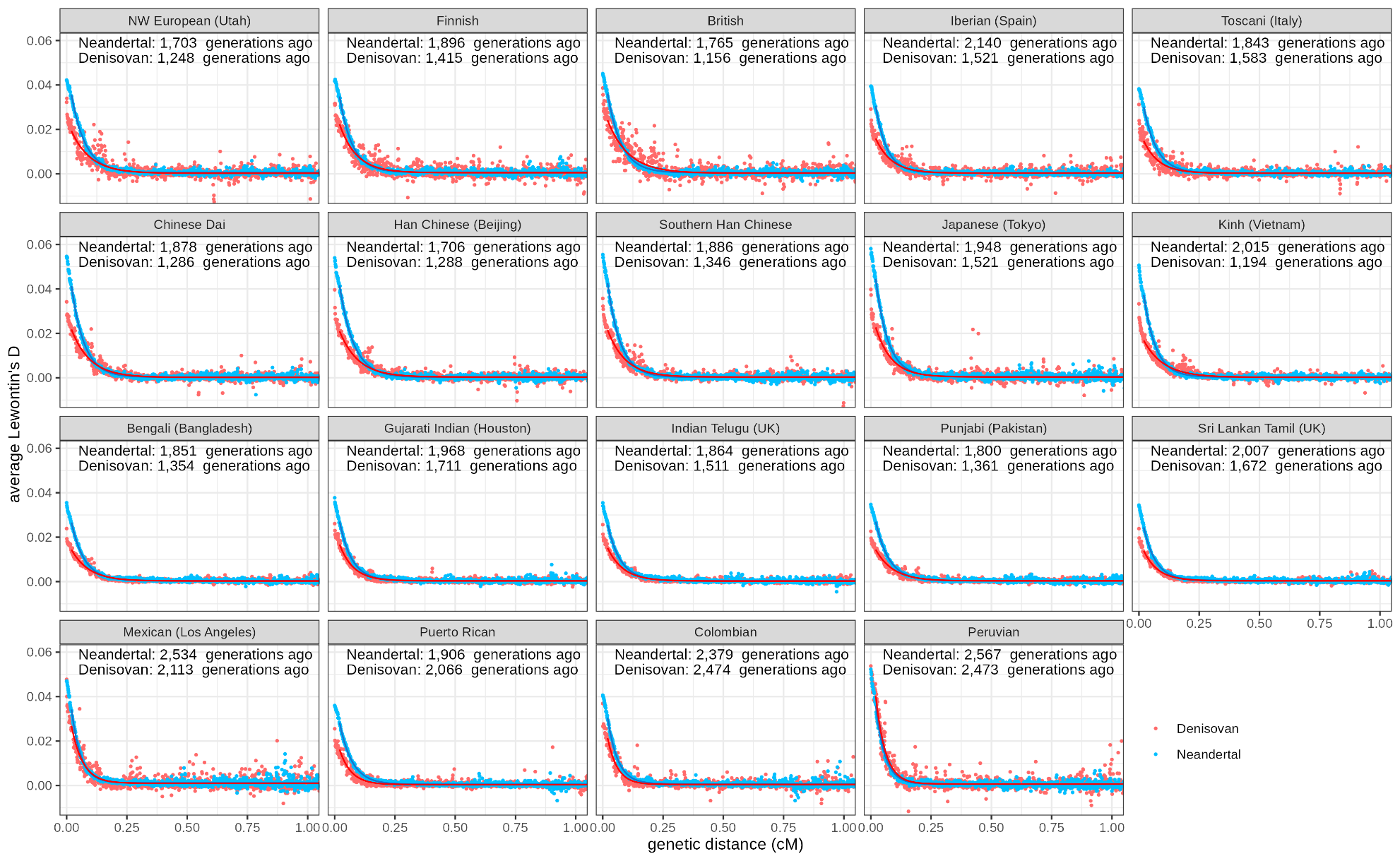

**Figure S100: Linkage disequilibrium decay between putative Neandertal or Denisovan introgressed alleles in present-day human populations.** Linkage disequilibrium was measured in the 1,000 Genomes dataset (phase 3)^129^ using the implementation of Lewontin’s D in PLINK^138^ for pairs of putative archaic variants that are less than 1Mb apart and less than 10 single nucleotide polymorphisms apart. We then computed the average in bins of 0.001cM. Putative Denisovan and Neandertal variants were identified as those that differ between the Denisova 3 (Denisovan) and Denisova 5 (Neandertal) genomes (after randomly sampling one allele at heterozygous positions), and that are absent in African genomes from the SGDP dataset^106^ and great ape genomes (panTro4, panpan1.1, gorGor3, ponAbe2). Genetic distances were computed using the recombination map inferred with pyrho for each population^135^. We fit an exponential curve (using the Nonlinear Least Square function in R) after excluding the pairs that are less than 0.02cM apart and report the estimated date of admixture for both Neandertal and Denisovan introgression in each panel.

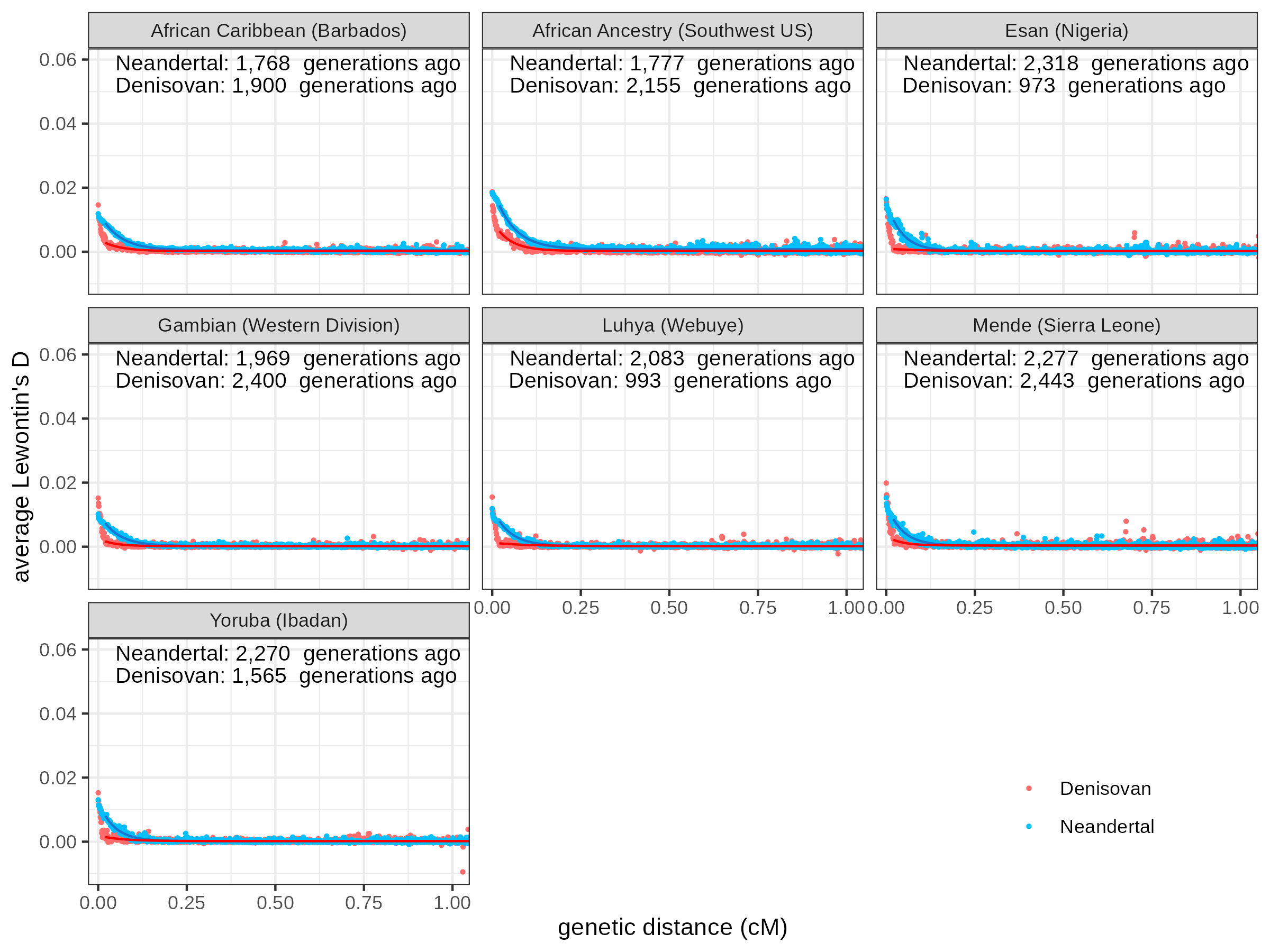

**Figure S101: Linkage disequilibrium decay between putative Neandertal or Denisovan introgressed alleles in present-day African populations.** Linkage disequilibrium was measured in the 1,000 Genomes dataset (phase 3)^129^ for pairs of putative archaic variants. Although these variants were identified by their absence in African genomes from the SGDP dataset^106^ (as described in **Figure S100**), some are present in the genomes of individuals with African ancestry in the 1,000 Genomes dataset and were used in this analysis. See **Figure S100** for more details.

#### Modeling of the match rate distributions and evidence for a third more divergent Denisovan component

Although some Denisovan ancestry in all populations is closer to *Denisova 3* than *Denisova 25*, there could be additional ancestry from more divergent Denisovans. We tested this possibility by modeling the rate at which Denisovan-like segments in present-day human genomes match the Denisovan genomes and evaluated whether including a more divergent Denisovan ancestry improves the fit to observed match rate distributions.

In previous work, match rates were fitted using normal distributions^124,127,139^, but the match rate distributions in real data exhibit a heavy lower tail because of the underlying exponential distribution of coalescent times. Furthermore, short segments lacking sufficient informative positions were filtered out to neglect uncertainties in the match rate of each segment to the Denisovan genome. To address these limitations, we modeled match rate distributions as beta-binomial distributions. The observed match rate for each segment follows a binomial distribution in which the underlying probability of sharing a derived allele with the Denisovan genome is not fixed between segments but randomly drawn from a beta distribution. This approach accommodates uncertainties in the match rate estimates and accounts for variation in coalescent times between segments in present-day human genomes and the Denisovan genome.

We computed match rates in each segment identified with *admixfrog* (using two Denisovan genomes in the reference) or with *hmmix* (most confidently assigned to Denisovans). Our approach differs from previous studies in two more aspects. First, we computed match rates as the proportion of shared derived alleles, rather than a proportion of matches over all bases, ensuring robustness to variations in mutation rates along the genome. Second, we used sets of independent segments (as described in the previous section) to avoid redundant inclusion of match rates for the same segments. Unlike previous sections where we focused on sites derived in present-day human genomes, here, we computed the match rates at positions where *Denisova 3* (or *Denisova 25*) carry a derived allele absent in an Mbuti outgroup (*S_Mbuti-1*; carrying the ancestral allele seen in four great ape genomes). This approach to measure match rates allows us to compare the divergence on the *Denisova 3* (or *Denisova 25*) lineage for the present-day humans and the other Denisovan without considering age differences and excludes modern human alleles from the other chromosome for heterozygous segments. More specifically, we considered a match between the putatively introgressed haplotype in the present-day human genome and the Denisovan genome if at least one allele of the present-day individual was also the derived allele seen in the Denisovan genome. In cases of heterozygous derived alleles in the Denisovan genome, we counted matches and mismatches as half to measure divergence on a single lineage.

We noticed that the average match rate of segments identified with *hmmix* decreases towards their ends (**Figure S102**), suggesting a tendency for *hmmix* to overestimate segment lengths and include the modern human background at segment edges. The average match rates of segments identified with *hmmix* are also lower compared to those identified with *admixfrog*, suggesting a higher incidence of false positives with *hmmix* and/or potential reference bias with *admixfrog*. We therefore focused on segments identified with *admixfrog* for a conservative analysis of the potential existence of a more divergent Denisovan component. We see that Native Americans exhibit lower match rates to *Denisova 3* and *Vindija 33.19* for both Denisovan-like and Neandertal-like segments, respectively, than other populations (except Oceanians, where some Denisovan segments may be misidentified as Neandertal segments; **Figure S102** and**Figure *S103***), suggesting a higher false positive rate in Native Americans (i.e. modern human ancestry misidentified as archaic ancestry). Consistent with this observation, the average match rate in Native Americans (and Oceanians) for segments of at least 0.1cM does not stand out from that of other populations (**Figure S104**). It is likely that the results for Native Americans are confounded by greater genetic drift between the allele frequencies of Native Americans and those of Africans used as reference for distinguishing archaic and modern human ancestry. Therefore, in subsequent analysis, we compared the results for segments of at least 0.05cM and those of at least 0.1cM.

We used an Expectation-Maximization algorithm to estimate the parameters of a mixture of two beta-binomial distributions and evaluated whether this mixture fits the data better than a single beta-binomial distribution using a likelihood ratio test. However, a single beta-binomial distribution fits the data better due to its flexibility in fitting various shapes with large variance, which is unlikely to accurately reflect the expected variance in coalescent time for the small effective population size of Denisovans. We therefore used a different approach leveraging the availability of two Denisovan genomes. We computed the match rate of one Denisovan genome to the other, in the same genomic regions identified as Denisovan-like in the genomes of the present-day human test population. We then tested whether this distribution fits the match rate distribution computed for present-day human genomes or whether the data is better explained by the presence of an additional Denisovan component in present-day humans that cannot simply be explained by the diversity observed between *Denisova 3* and *Denisova 25*

Leveraging the availability of a second Denisovan genome, we therefore fitted the data using the match rate distribution observed between the two Denisovans within the same segments identified as Denisovan ancestry in present-day human genomes. This allowed us to test whether the match rate of the Denisovan ancestry in present-day humans fits to the observed match rate between *Denisova 3* and *Denisova 25*, or whether it supports the presence of a more divergent Denisovan ancestry component.

The match rates of Denisovan-like segments in East Asians and Central Asians/Siberians fit the rate at which *Denisova 25* matches *Denisova 3* indicating that there is no evidence for a more divergent Denisovan ancestry component in these populations. In contrast, the match rate distributions in Oceanians and South Asians significantly deviate toward lower match rates, suggesting the presence of a more divergent Denisovan ancestry component in these populations (**Figure S105**). Although the match rate distributions in Native Americans and West Eurasians include considerably lower match rates than observed in Oceanians and South Asians, these likely represent a larger proportion of false positives (i.e. Neandertal or modern human ancestry) among the few Denisovan-like segments identified in Native Americans and West Eurasians. Restricting the analysis to segments of 0.1cM or longer removes most segments with a low match rate to *Denisova 3* in both populations, which is consistent with an increased false positive rate in these populations, while the results remain consistent for other populations (**Figure S106**). As the difference in match rates for Native Americans and West Eurasians compared to *Denisova 25* and *Denisova 3* are then only marginally significant (p>0.01), it remains unclear whether these two populations also carry a more divergent Denisovan ancestry.

For comparison, we evaluated match rates to *Denisova 25*, using *Denisova 3* to compute the expected match rate distribution for Denisovan ancestry originating on the *Denisova 3* lineage. All populations, including East Asians and Central Asians/Siberians, show lower than expected match rates, suggesting the potential presence of divergent Denisovan components in all populations (**Figure S107**). However, incorrect determination of segment length introducing some modern human ancestry, as observed with *hmmix*, or some segments mistaken as Denisovan ancestry, could bias the observed match rates compared to expectations. Given the short lineage of *Denisova 25*, match rates of *Denisova 3* to *Denisova 25* are much higher, making this comparison more sensitive to errors. After restricting to segments of 0.1cM or longer, the match rates in Central Asians/Siberians and West Eurasian fit the observed distribution of match rates of *Denisova 3* to *Denisova 25* (**Figure S108**).

To further investigate the presence of a more divergent Denisovan component in South Asians and Oceanians, potentially absent in East Asians and Central Asians/Siberians, we examined allele counts supporting a closer relationship to either *Denisova 3*, *Denisova 25*, or their common ancestor. If present-day humans only inherited Denisovan ancestry from a population closer to *Denisova 3* than *Denisova 25*, we expect the greatest support for the *Denisova 3* branch and equal support for the ancestral and *Denisova 25* branches, due to incomplete lineage sorting. We computed shared derived alleles with *Denisova 3* or *Denisova 25* as described above and also counted positions where both Denisovans share a derived allele absent in the Denisovan-like segments in present-day human genomes. These calculations were conducted on sets of independent segments (as described in the previous section), and the results were compared across populations (i.e. dividing by the total number of informative positions in each population, to compare proportions instead of counts of positions). We obtained confidence intervals from 1,000 bootstraps of the segments. In Oceanians and South Asians, the ancestral branch shows the highest support, whereas in East Asians and Central Asians/Siberians, the *Denisova 3* branch shows the highest support (**Figure S109**). This supports the presence of a Denisovan component in Oceanians and South Asians that is divergent to both Denisovan genomes, whereas East Asians and Central Asians/Siberians mostly carry ancestry closer to *Denisova 3*. In Oceanians, the *Denisova 3* branch also shows higher support than the *Denisova 25* branch, confirming the presence of some Denisovan ancestry closer to *Denisova 3* than to *Denisova 25*. In South Asians, the confidence intervals overlap, but the *D*-statistics and match rate analyses in previous sections suggest that South Asians also carry such ancestry. Interestingly, there is no evidence for more support of the ancestral branch over the *Denisova 25* branch in East Asians and Central Asians/Siberians, although the support for the ancestral branch is slightly higher. Focusing on segments with a minimum length of 0.1cM to mitigate potential false positives among shorter segments leads to the same support for the ancestral and *Denisova 25* branches in Central Asians/Siberians and decreased the support for the ancestral branch in East Asians. These results indicate that there is no evidence for a more divergent Denisovan component in East Asians and Central Asians/Siberians, whereas such a component exists in Oceanians and likely in South Asians. Its presence in Native Americans and Western Europeans is unclear, warranting further investigation with additional genomes to identify more Denisovan-like segments and reduce confounding false positives.

Finally, we calculated the proportion of Denisovan-like segments shared across populations to assess whether these segments were inherited from a common admixture event, in particular for the divergent Denisovan component present in South Asians and Oceanians. We first compared the sets of independent segments from each population (i.e. after randomly sampling one segment for those that overlap in the genomes of individuals from a same geographical group) identified with *admixfrog*. We found that between 12% and 66% of Denisovan-like segments overlap between pairs of Eurasian or American populations, whereas less than 8% of the segments in Oceanians overlap with those in other populations (**Figure S110**). Notably, only about 35% of segments in South Asians overlap with those in Oceanians, compared to over 40% of segments in other populations that overlap with those in Oceanians. Therefore, there is no evidence that the divergent Denisovan ancestry detected in both South Asians and Oceanians was inherited from a shared admixture event. These results remained consistent when we aggregated segments in each population by merging overlapping segments instead of randomly sampling one (**Figure S111**).

Next, we examined shared breakpoints at the 5’ and 3’ ends of segments between population pairs, i.e. segments identified with *admixfrog* that start or end in the same genomic window. Since breakpoints indicate past recombination events that are likely inherited from a common ancestor, shared breakpoints serve as evidence for shared Denisovan ancestry from a common admixture event. While there is evidence of shared breakpoints among Eurasians and Native Americans, suggesting that they inherited at least some Denisovan ancestry from a common event — either in their ancestral population or later shared through gene flow — Oceanians show almost no shared breakpoints with other populations (**Figure S112** and **Figure *S113***). This suggests that Oceanians inherited their Denisovan ancestry from separate admixture events, or isolation and genetic drift eroded the signal of shared Denisovan ancestry. As Neandertal segments share breakpoints across all populations (**Figure S114**), supporting a common event of admixture with Neandertals, the lack of shared Denisovan-like segment breakpoints with Oceanians is therefore likely due to separate Denisovan admixture events rather than to isolation or genetic drift. Additionally, there is no evidence that the segments in South Asians share more breakpoints with those in Oceanians than other populations, reinforcing the conclusion that the divergent Denisovan ancestry in South Asians and Oceanians stem from distinct admixture events. To further assess breakpoint sharing, we compared segments from one population against the aggregated set of segments from all other populations. We found that less than 4% of segment breakpoints in Oceanians are shared with other populations, whereas 12% to 46% of breakpoints in other populations are shared with each other (**Table S67**). We note that the small number of segment breakpoints Oceanians share with other groups could be due to Neandertal segments that are misidentified as Denisovan, or result from incomplete lineage sorting, as the number of shared segment breakpoints decreases to less than 2% for longer segments.

**Table S67: Number and proportion of Denisovan-like segment breakpoints individuals from a geographical group (first column) share with individuals from any other group.**

| **Geographical group** | **End type** | **Segments of at least 0.05cM** | | | **Segments of at least 0.1cM** | | |
| --- | --- | --- | --- | --- | --- | --- | --- |
|  |  | **# shared** | **# total** | **proportion** | **# shared** | **# total** | **proportion** |
| America | 5’ | 77 | 216 | 36% | 7 | 37 | 19% |
|  | 3’ | 76 | 216 | 36% | 7 | 37 | 19% |
| Central Asia Siberia | 5’ | 149 | 303 | 49% | 25 | 55 | 46% |
|  | 3’ | 148 | 303 | 49% | 17 | 55 | 31% |
| East Asia | 5’ | 171 | 477 | 36% | 23 | 114 | 20% |
|  | 3’ | 174 | 477 | 36% | 27 | 114 | 24% |
| Oceania | 5’ | 92 | 2,670 | 3% | 9 | 923 | 1% |
|  | 3’ | 97 | 2,670 | 4% | 16 | 923 | 2% |
| South Asia | 5’ | 107 | 493 | 22% | 13 | 107 | 12% |
|  | 3’ | 111 | 493 | 23% | 13 | 107 | 12% |
| Western Eurasia | 5’ | 91 | 233 | 39% | 8 | 38 | 21% |
|  | 3’ | 99 | 233 | 42% | 15 | 38 | 39% |

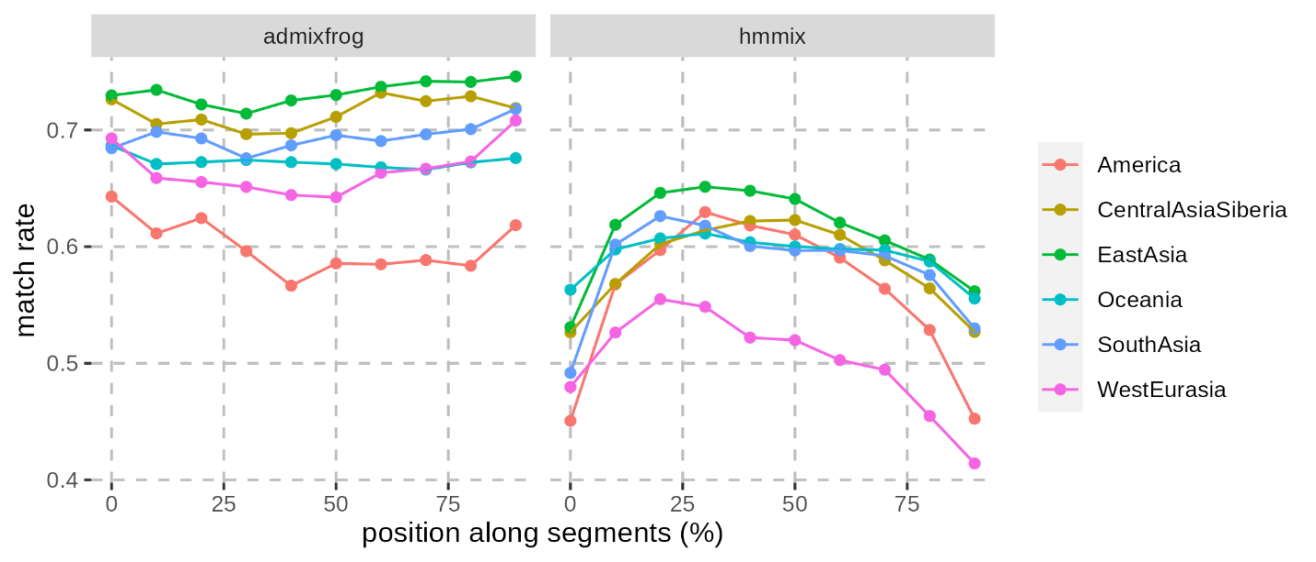

**Figure S102: Average match rate to Denisova 3 along putative Denisovan segments in present-day human genomes.** For this analysis, we computed the match rates at positions where Denisova 3 carries a derived allele absent from a Mbuti outgroup (S_Mbuti-1; carrying the ancestral allele seen in four great ape genomes). The average match rate was computed in 10 non-overlapping sliding windows representing 10% of the length of segments. We used an aggregated and independent set of segments for each geographical group (identified using the reference with two Denisovan genomes for the segments identified with admixfrog) as described above. The facets show the match rate for segments identified with admixfrog and hmmix and the colours correspond to the different geographical groups.

**Figure S103: Average match rate to Vindija33.19 along putative Neandertal segments in present-day human genomes after different minimum length cutoffs.** The facets show the match rate for segments of at least 0.05cM (left) and 0.1cM (right) identified with admixfrog. Those match rates were computed as described for Denisovan segments, using a set of aggregated and independent segments identified with admixfrog in each geographical group. We used the same reference used for detecting Denisovan segments, i.e. including both Denisovan genomes. See **Figure S102** for additional details.

**Figure S104: Average match rate to Denisova 3 along putative Denisovan segments in present-day human genomes after different minimum length cutoffs.** The facets show the match rate for segments of at least 0.05cM (left) and 0.1cM (right) identified with admixfrog. See **Figure S102** for more details.

**Figure S105: Modeling of the distributions of match rate to Denisova 3 of putative Denisovan segments in present-day human genomes.** For this analysis, we computed the match rates at positions where Denisova 3 carries a derived allele absent from a Mbuti outgroup (S_Mbuti-1; carrying the ancestral allele seen in four great ape genomes). Those match rates were computed for the set of aggregated and independent segments for each geographical group identified with admixfrog, using both Denisovan genomes in the reference. We fitted mixtures of two beta-binomial distributions using the match rate of Denisova 25 to Denisova 3 within the same genomic segments as the expected distribution for one component (blue distribution) and tested whether adding a second component (red distribution) provides a better fit to the data. The distributions were fitted using an Expectation-Maximization algorithm and the reported p-values were computed using a likelihood ratio test.

**Figure S106: Modeling of the distributions of match rate to Denisova 3 of putative Denisovan** **segments of at least 0.1cM in present-day human genomes.** See **Figure S105** for additional details.

**Figure S107: Modeling of the distributions of match rate to Denisova 25 of putative Denisovan segments in present-day human genomes.** For this analysis, we computed the match rates at positions where Denisova 25 carries a derived allele absent from a Mbuti outgroup (S_Mbuti-1; carrying the ancestral allele seen in four great ape genomes). We fitted mixtures of two beta-binomial distributions using the match rate of Denisova 3 to Denisova 25 within the same genomic segments as the expected distribution for one component (blue distribution) and tested whether adding a second component (red distribution) provides a better fit to the data. The distributions were fitted using an Expectation-Maximization algorithm and the reported p-values were computed using a likelihood ratio test.

**Figure S108: Modeling of the distributions of match rate to Denisova 25 of putative Denisovan segments of at least 0.1cM in present-day human genomes.** See **Figure S107** for additional details.

**Figure S109: Proportion of positions in putative Denisovan segments in present-day human genomes that support one of the three possible topologies relating these segments to Denisova 3 and Denisova 25.** The support for each topology was evaluated by counting the number of derived alleles in the Denisovan-like segments of present-day human genomes that are shared with Denisova 3 or Denisova 25, or the number of ancestral alleles at positions in these segments where Denisova 3 and Denisova 25 share a derived allele. We computed these counts on sets of independent segments, and obtained the proportion of positions supporting each branch by dividing by the total number of positions in each population. Whiskers correspond to 95% confidence intervals computed with 1,000 bootstraps of the segments. The facets show the results for segments of at least 0.05cM (left) and 0.1cM (right) identified with admixfrog.

**Figure S110: Overlap of putative archaic segments in the genomes of present-day humans after randomly sampling segments that overlap within each geographical group.** The left and right panels show the proportion of Denisovan and Neandertal segments, respectively, from the population indicated in the columns that overlap segments from the populations indicated in the rows. We used segments of at least 0.05cM identified with admixfrog, using both Denisovan genomes in the reference.

**Figure S111: Overlap of putative archaic segments in the genomes of present-day humans after merging segments that overlap within each geographical group.** See **Figure S110** for additional details.

**Figure S112: Proportion of breakpoints of putative Denisovan segments of at least 0.05cM that are shared between the genomes of present-day humans from different geographical groups.** The left and right facets show the proportion of segments from the population indicated in the columns that share a 3’ and 5’ end, respectively, with segments from the populations indicated in the rows. We used segments of at least 0.05cM identified with admixfrog, using both Denisovan genomes in the reference.

**Figure S113: Proportion of breakpoints of putative Denisovan segments of at least 0.1cM that are shared between the genomes of present-day humans from different geographical groups.** See **Figure S112** for additional details.

**Figure S114: Proportion of breakpoints of putative Neandertal segments of at least 0.05cM that are shared between the genomes of present-day humans from different geographical groups.** See **Figure S112** for additional details.

### Supplementary Note 12: Analysis of adaptive introgressed Denisovan DNA

Securely identified Denisovan fossils from the Altai Mountains^8,61,62,64^, the Tibetan Plateau^9,140,141^, Northeast China^142,143^, and the coast of Taiwan^144^ indicate that Denisovans inhabited much of Asia for hundreds of thousands of years and likely adapted to diverse local environments. Several studies have identified Denisovan DNA segments at high-frequency in present-day humans that are candidates for adaptive introgression^124,133,139,145-149^, suggesting that at least some of these segments may have provided benefits to modern humans migrating into these new environments. The increased power to detect Denisovan ancestry segments using a second Denisovan genome (**Supplementary Note 11**), combined with the possibility to identify introgressed DNA from distinct Denisovan lineages depending on their relationship to *Denisova 3* and *Denisova 25*, allows for a systematic screen for further candidates of adaptive introgression and investigation of their likely source population.

Denisovan-like segments were identified with *admixfrog* as described in **Supplementary Note 11**, in a set of diverse present-day individuals^106^. Following Iasi *et al.* (2024)^119^, for each 0.005cM bin along the genome of each individual, we calculated the sum of posterior probabilities for states including Denisovan ancestry (homozygous Denisovan, heterozygous Denisovan/modern human, and heterozygous Denisovan/Neandertal). Bins with a joint posterior probability of at least 0.8 were assigned as most likely Denisovan DNA if they overlap a Denisovan-like segment (as identified in **Supplementary Note 11**) with a minimum length of 0.05cM and containing at least one variant matching either of the Denisovan reference genomes. Analyses were further restricted to bins located within 20kb windows containing at least one polymorphism in the *admixfrog* reference genome panel (callable regions). The posterior probability of Denisovan ancestry for all other bins was set to zero, and bins with no Denisovan DNA in any individual were excluded.

Because Denisovan DNA was introduced into modern humans at different proportions in multiple independent events of admixture with different Denisovan populations, we assess adaptive introgression in each SGDP-defined Superpopulation separately.

For each group, we tested whether the frequency of Denisovan DNA in a bin deviates significantly from the average across all bins with Denisovan DNA in that group. Specifically, we calculated a Z-score for bin *i*:

$Z=\frac{(\mu i-ū)}{\sigma̅}$, with 𝜇*_i_* denoting the mean posterior probability across all individuals in a group for bin *i*, and $ū$ and $\sigma̅$ corresponding to the mean and standard deviation. Because the number of individuals in each Superpopulation varies widely (21-75 individuals), and Denisovan ancestry levels in most populations are very low (<0.2%), even a single segment in one individual may deviate significantly enough from the genome-wide average to be considered an outlier. We therefore computed the average frequency ($ū)$ and its standard deviation ($\sigma̅)$ of Denisovan introgressed segments only in bins with Denisovan ancestry in at least one individual (i.e: across the introgressed genome), instead of the genome-wide frequency of Denisovan ancestry. We defined outlier bins as those with Z ≥ 4.5 and merged consecutive outlier bins within 0.05cM of each other into candidate regions of adaptive introgression.

For comparison, if we called candidates by detecting outliers in the genome-wide levels of Denisovan ancestry this would yield 1,631 outliers (139,219,005bp) across all superpopulations. These numbers are implausibly high; for example, in South Asia, this approach identifies 376 outliers although we only identified 569 Denisovan-like DNA segments in total (373 non-overlapping segments). Therefore, we focus on candidates identified by calling outliers compared to the introgressed genome, which resulted in 38 candidates spanning 3,886,059 bp across all superpopulations (**Supplementary Data 1**).

#### Geographical distribution and frequency of Denisovan-like DNA segments in candidate regions

When comparing the overlap of candidate regions between superpopulations, we observed that most candidate regions are only identified as outliers in a single superpopulation (36 out of 38), with the highest numbers in Oceanians (27 candidates) and South Asians (4 candidates), and none detected in Central Asia-Siberia (**Figure S115A**). Only one candidate region is shared between two groups, East-Asia and West-Eurasia.

Next, we compared how often a candidate region overlaps Denisovan-like DNA segments in any given superpopulation (**Figure S115B**). Most candidates are unique to a superpopulation, in the sense that they do not overlap with Denisovan-like DNA segments in other superpopulations (24 out of 38). There are twenty-two unique candidates identified in Oceania and 2 in South Asia, which do not overlap any Denisovan-like DNA segments identified in other superpopulations.

We further estimated the frequency of Denisovan-like DNA segments for each adaptively introgressed candidate by counting individuals in the superpopulation where it is a candidate with at least one Denisovan-like DNA segment (≥0.05cM and ≥1 match to the Denisovan reference) overlapping the region (**Figure S115D**). Frequencies ranged from 10% (a West Eurasian candidate) to 75 % (an Oceanian candidate). Finally, we quantified the sharing of Denisovan-like DNA segments that overlap candidate regions across superpopulations by correlating the presence of segments in candidate regions between pairs of superpopulations. Oceanian candidates show ananti-correlation with those in other superpopulations (**Figure S115C**) indicating that the Denisovan ancestry in Oceanians was inherited independently from other populations. South Asian candidates also show reduced correlation with other superpopulations, compared to the higher correlations observed between East-Asians, Native Americans and West-Eurasians. This pattern is consistent with that some of the Denisovan ancestry in South Asians may have been inherited independently from other populations.

**Figure S115: Distribution of candidate regions.** **A**) Number of candidate regions per SGDP superpopulation. **B**) Number of superpopulations with Denisovan ancestry overlapping a candidate region. **C**) Correlation of Denisovan ancestry presence/absence across candidate regions between superpopulations. Crosses indicate non-significant correlation. **D**) Frequency of Denisovan ancestry segments overlapping candidate regions per superpopulation, stratified by whether the Denisovan ancestry segments in this region are shared (at least one segment overlapping the region in two superpopulations) or unique to the superpopulation. Red bars indicate if the region is a candidate in the superpopulation, blue indicates the region is a candidate in another superpopulation.

#### Comparison to previously identified candidates for adaptive introgression

We next compared our candidates for Denisovan adaptive introgression with those previously reported in seven studies^124,133,139,145-148^. To assemble this comparative set, we took the union of the candidate regions identified in these studies, resulting in a total of 884 regions. Of the 38 candidates identified, 22 were novel and 16 (42%) matched previously reported regions.

Differences in populations analyzed, methods for calling Denisovan segments, and statistics to identify candidates for adaptive introgression likely contribute to discrepancies between studies. We therefore focused on the candidates identified by Sankararaman *et al.* (2016)^133^, which used the same individuals as in our dataset. Since this study only focused on Oceanians we only compared candidates that we identified in this population. Among the 48 candidate regions identified by Sankaraman *et al.*, 10 (21%) were also recovered in our scan (**Figure S116A**), while 12 of our 27 candidates (44%) were also present in their dataset (four candidates in our scan are merged into two in the Sankaraman *et al*. set). When we stratify the candidates by their population frequency (as inferred in our scan), we observed increased overlap for higher frequency candidates, though some high-frequency candidates were unique to our analysis (**Figure S116B**).

Two regions identified by Sankararaman *et al.* (2016) show no Denisovan segments meeting our criteria. In region 1:63,700,000-63,800,000, we detected 6 Denisovan segments with a genetic length of 0.01 cM (366,821 bp). In region 6:126,700,000-127,100,000, we found 3 Neandertal and 14 Denisovan segments, with Denisovan segments reaching a maximum length of 0.035cM (92,036bp). In both cases, the segments fell below our length cutoff of 0.05cM and were excluded. More generally, we did not identify candidates with frequencies below ~0.2, whereas some candidates from Sankararaman *et al.* (2016) have a frequency as low as 0.04. This likely reflects differences in inferred Denisovan segment frequencies, perhaps because we use an additional Denisovan genome. We explore this in the next section.

**Figure S116: Comparison of our candidates in Oceanians to those identified by Sankararaman et al. (2016)^133^.** **A**) Proportions of regions found by Sankararaman et al. (2016) (left) or in this study (right) that were replicated in both studies, not found in this study or not found by Sankararaman et al. (2016). **B**) Frequency of Denisovan ancestry segments we identified within candidate regions from our scan and within regions reported by Sankararaman et al. (2016).

#### Comparison between using one or two Denisovan reference genomes

To assess whether using an additional Denisovan genome as reference increases the power to detect candidate regions for adaptive introgression, we repeated the analysis with segments called using only *Denisova 3* as the reference (**Supplementary Note 11**). This yielded 23 unique candidates across all superpopulations, fewer than the 38 found using both Denisovans as reference. Comparison of the D3 only to the D3+D25 callsets with the regions identified by Sankararaman *et al.* (2016) shows that 10 candidates unique to our study are identified only when using both Denisovan genomes rather than *Denisova 3* alone (**Figure S117A**). Thus, the differences between the D3 only and D3+D25 callset largely reflects additional candidates detected when including the second Denisovan genome (**Figure S117B**). **Figure S117C** shows the frequency differences between the callsets for each region. The largest increase in candidate regions when using both Denisovan genomes was observed in Oceania, suggesting improved sensitivity to detect Denisovan ancestry in this superpopulation.

**Figure S117: Comparison of Superpopulation-specific candidate regions identified using admixfrog call sets using both Denisovans or only Denisova 3.** A) Proportion of candidate regions that overlap or are unique to the three different callsets, using one or two Denisovans, or identified in Sankararaman et al. (2016). Striped sections in the pie charts indicate overlap with the Sankararaman callset. Facets indicate to which callset the comparison was made to. B) Proportion of candidates identified using one or two Denisovan genomes as reference. Facets indicate to which callset the comparison was made to. C) Population frequency of candidate regions identified using one and/or two Denisovan genomes as reference across multiple superpopulations.

#### Differences in the origin of the candidates of Denisovan adaptive introgression between populations

As we identified multiple Denisovan ancestry components in the genomes of present-day humans, we next investigated the origin of the Denisovan-like DNA segments in the candidate regions for adaptive introgression. We classified the Denisovan segments overlapping each candidate region into three different categories based on their match rate to *Denisova 3*, relative to the match rate of *Denisova 25* to *Denisova 3* in the same genomic region (as described in **Supplementary Note 11**). Segments matching *Denisova 3* more closely than *Denisova 25* does at the same locus were classified as “Denisova 3-like”; those matching *Denisova 3* and *Denisova 25* equally well as “ambiguous”; and those matching *Denisova 3* less well than *Denisova 3* matches *Denisova 25* as “ancestral”. Each region was then assigned to one of three categories: “mostly *Denisova 3*-like” if at least 70 % of the segments are “*Denisova 3*-like”, “mostly ancestral” if at least 70 % of the segments are “ancestral”, or “mixed”. No regions were classified as “mostly ambiguous” as the maximum proportion of ambiguous segments seen in any region is 60%.

**Figure S118: Composition of Denisovan ancestry components among candidate regions for adaptive introgression.** **A**) Number of candidate regions mostly exhibiting Denisova 3-like segments, “ancestral” segments, or segments of mixed origin. **B**) Same as **A**, shown as proportions of the number of candidate regions per superpopulation.

Most candidates in Oceania and South Asia are composed mostly of segments that are classified as “ancestral” (**Figure S118**), whereas no such candidates were found in Native Americans or East Asians. To test whether specific ancestry components are enriched among the candidates for adaptive introgression compared to the rest of the genome, we sampled *n* random genomic regions where we identified Denisovan segments (without replacement), where *n* is the number of candidates identified in the respective superpopulation. Each random genomic region had to overlap at least two Denisovan segments (otherwise a mixed category would not be possible), and we recorded how many were classified as “mostly *Denisova 3*-like”, “mostly ancestral” or “mixed”. We repeated this procedure 1,000 times and computed empirical p-values using the *ecdf* function from the *stats* package in *R,* comparing the observed versus expected number of regions in each ancestry category (**Figure S119**).

We found that candidate regions in Oceania (and in Oceania plus South Asia combined) are significantly enriched for “mostly ancestral” segments, whereas candidate regions in East Asia, the Americas, and West Eurasia are enriched for “mostly *Denisova 3*-like” segments. When pooling the latter individuals with those from South Asia, however, the enrichment for candidates with “mostly *Denisova 3-*like” segments is no longer observed, which is consistent with a different composition of candidates in South Asians compared to these other groups.

**Figure S119: Expected vs observed number of candidate regions with “mostly Denisova 3-like”, “mostly ancestral”, or “mixed” Denisovan ancestry segments (different colours).** Lines indicate the observed number of candidate regions from a particular category. Distributions were obtained from randomly sampling introgressed segments. Solid and dotted lines indicate whether the difference is significant and not significant, respectively, using a p-value cutoff of 0.05. The facets show the results for candidates identified in different superpopulations.

#### Genes overlapping candidate regions

We annotated candidate regions using the GENCODE Havana gene set (release 44, GRCh37), considering only genes. Among all 38 candidates, 31 overlap with 96 genes (including different isoforms), while 7 regions had no overlapping genes (**Figure S120**).

The number of genes per region is positively correlated with region length, with a maximum of 23 genes in a single region. To determine whether the regions identified are enriched for genes, we randomly placed the candidate regions per-superpopulation in the Denisovan introgressed genome (union of all segments passing our filters in that superpopulation) or the whole callable genome (as defined by our callable regions mask) with the same total length as the candidate regions and recorded the number of overlapping genes in each replicate (1,000 replicates in total). We obtained p-values by counting how often the random sample had more or less genes than the candidate regions. We found no enrichment or depletion of genes relative to expectation for either background (**Table S68**). However, when assessing the distribution of Denisovan-like introgressed DNA in callable regions genome-wide we see that Denisovan DNA is depleted in genic regions in Oceanians (p-value: 0.001), replicating a previous result (Sankararaman *et al.* 2016) and in South Asians and Americans (p-values: 0.004/0.005), but not in West-Eurasians or East-Asians.

**Figure S120: Genes in outlier regions.** **A**) Number of outlier regions with a given number of genes. **B**) The length of the outlier region in base pairs versus the number of genes.

**Table S68: Enrichment/Depletion of genes in candidate regions or all Denisovan-like DNA.**

| **Superpopulation** | **Test** | **Background** | **observed genes overlapping** | **mean random genes overlapping** | **sd random genes overlapping** | **p_value_over** | **p_value_under** |
| --- | --- | --- | --- | --- | --- | --- | --- |
| Oceania | candidate regions | Densiovan Segments | 84 | 61.612 | 13.39316366 | 0.064 | 0.942 |
| Oceania | candidate regions | All callable genomic regions | 84 | 65.005 | 15.75243201 | 0.122 | 0.888 |
| Oceania | Densiovan Segments | All callable genomic regions | 5514 | 5929.938 | 145.7929482 | 0.999 | **0.001** |
| SouthAsia | candidate regions | Densiovan Segments | 4 | 8.982 | 5.677491516 | 0.893 | 0.196 |
| SouthAsia | candidate regions | All callable genomic regions | 4 | 10.045 | 6.266757315 | 0.937 | 0.124 |
| SouthAsia | Densiovan Segments | All callable genomic regions | 822 | 968.696 | 57.7872906 | 0.996 | **0.004** |
| EastAsia | candidate regions | Densiovan Segments | 2 | 3.369 | 2.242736915 | 0.823 | 0.405 |
| EastAsia | candidate regions | All callable genomic regions | 2 | 3.397 | 2.232123064 | 0.823 | 0.399 |
| EastAsia | Densiovan Segments | All callable genomic regions | 864 | 888.295 | 53.27152927 | 0.678 | 0.336 |
| America | candidate regions | Densiovan Segments | 4 | 3.496 | 1.90567983 | 0.454 | 0.752 |
| America | candidate regions | All callable genomic regions | 4 | 4.505 | 2.833373079 | 0.609 | 0.579 |
| America | Densiovan Segments | All callable genomic regions | 177 | 235.261 | 26.09950312 | 0.996 | **0.005** |
| WestEurasia | candidate regions | Densiovan Segments | 2 | 4.59 | 5.212395713 | 0.713 | 0.436 |
| WestEurasia | candidate regions | All callable genomic regions | 2 | 4.032 | 3.634306562 | 0.757 | 0.408 |
| WestEurasia | Densiovan Segments | All callable genomic regions | 242 | 256.353 | 25.70254875 | 0.715 | 0.298 |

#### Gene Ontology Enrichment of genes in candidate regions

We performed a Gene Ontology (GO) enrichment analysis for all candidate regions that overlap genes. We used a hypergeometric test to compare genes in these candidate regions to those in regions with Denisovan ancestry segments found in at least one non-African individual in our data, that overlap a gene but do not overlap the candidate regions being tested. To determine our background set, we filtered Deniosvan haplotypes to include only segments longer than 0.05 cM and with at least one variant matching either Denisovan genome. We excluded haplotypes outside of the callable region mask. Overlapping Denisovan haplotypes were merged. We then randomly resampled regions with Denisovan ancestry to account for the spatial clustering of functionally-related genes along the genome, i.e. genes with similar function tend to be in physical proximity in the genome and will therefore be more likely to be on the same Denisovan introgressed haplotypes. Since our background set consists of Denisovan haplotypes, we also account for other features that affect the likelihood of a gene to be on a Denisovan candidate haplotype, such as gene length, recombination rate, and background selection.

To perform the test, we used the *R* package *GOfuncR*^150^. We used a custom annotation set that included all genes overlapping the Denisovan introgressed regions based on the GENCODE Havana gene set (release 44, GRCh37). We corrected for multiple testing performing a Family-Wise Error Rate correction, which shuffles the candidates within the background regions and recomputes p-values, 10,000 times. We circularized the chromosome used for the background to prevent edge effects when placing the random regions.

We identified a significant enrichment (corrected p-values<0.05) in two related GO categories (biological process): *sensory perception of sound* and *sensory perception of mechanical stimulus*, which involve the genes *ATP6V0A4* and *COL4A3* that are both candidates for adaptive introgression in South Asians (**Figure S121**). **Table S69** shows the highest three GO terms per superpopulation per ontology category.

**Figure S121: Denisovan-like DNA segments overlapping the ATP6V0A4 and COL4A3 genes in significant GO categories in South Asia.** Color and shade indicate segment ancestry classification and the certainty of that classification. Box indicates location of candidate region with Z score indicated to its right. Green rectangles at the bottom indicate the positions and lengths of genes overlapping the candidate region. Derived states of functional variants are indicated with filled dots for derived SNPs in the UTR and circles for eQTLs.

**Table S69: Gene Ontology enrichment result.**

| **Ontology category** | **Node ID** | **Node name** | **raw_p_overrep** | **FWER_overrep** | **Region** | **Genes** |
| --- | --- | --- | --- | --- | --- | --- |
| biological_process | GO:0006356 | regulation of transcription by RNA polymerase I | 0.0007 | 0.5472 | Oceania | ERBB2 |
| biological_process | GO:0008631 | intrinsic apoptotic signaling pathway in response to oxidative stress | 0.0019 | 0.7784 | Oceania | MAP3K5, VNN1 |
| biological_process | GO:0036473 | cell death in response to oxidative stress | 0.0019 | 0.7784 | Oceania | MAP3K5, VNN1 |
| cellular_component | GO:0000785 | chromatin | 0.0167 | 0.8355 | Oceania | MED1, NEUROD2, SALL4, TMPO |
| cellular_component | GO:0055037 | recycling endosome | 0.0133 | 0.7770 | Oceania | ATP9A, TF |
| cellular_component | GO:0140534 | endoplasmic reticulum protein-containing complex | 0.0175 | 0.8536 | Oceania | MAP3K5, ORMDL3, SRPRB, UGT3A1 |
| molecular_function | GO:0001042 | RNA polymerase I core binding | 0.0274 | 0.9936 | Oceania | ERBB2 |
| molecular_function | GO:0001594 | trace-amine receptor activity | 0.0043 | 0.7594 | Oceania | TAAR1, TAAR2 |
| molecular_function | GO:0008227 | G protein-coupled amine receptor activity | 0.0185 | 0.9523 | Oceania | TAAR1, TAAR2 |
| biological_process | GO:0001678 | cellular glucose homeostasis | 0.0082 | 0.3936 | EastAsia | PRKCE |
| biological_process | GO:0002221 | pattern recognition receptor signaling pathway | 0.0082 | 0.3936 | EastAsia | PRKCE |
| biological_process | GO:0002224 | toll-like receptor signaling pathway | 0.0082 | 0.3936 | EastAsia | PRKCE |
| cellular_component | GO:0005739 | mitochondrion | 0.0970 | 0.5626 | EastAsia | PRKCE |
| cellular_component | GO:0005794 | Golgi apparatus | 0.0746 | 0.5285 | EastAsia | PRKCE |
| cellular_component | GO:0048471 | perinuclear region of cytoplasm | 0.0373 | 0.4516 | EastAsia | PRKCE |
| molecular_function | GO:0003785 | actin monomer binding | 0.0077 | 0.2841 | EastAsia | PRKCE |
| molecular_function | GO:0004699 | calcium-independent protein kinase C activity | 0.0077 | 0.2841 | EastAsia | PRKCE |
| molecular_function | GO:0030546 | signaling receptor activator activity | 0.0077 | 0.2841 | EastAsia | PRKCE |
| biological_process | GO:0051641 | cellular localization | 0.0021 | 0.1692 | WestEurasia | AGFG1, PRKCE |
| biological_process | GO:0051649 | establishment of localization in cell | 0.0014 | 0.1567 | WestEurasia | AGFG1, PRKCE |
| biological_process | GO:0097435 | supramolecular fiber organization | 0.0008 | 0.1183 | WestEurasia | AGFG1, PRKCE |
| cellular_component | GO:0005635 | nuclear envelope | 0.0882 | 0.8045 | WestEurasia | AGFG1 |
| cellular_component | GO:0005643 | nuclear pore | 0.0226 | 0.4768 | WestEurasia | AGFG1 |
| cellular_component | GO:0005794 | Golgi apparatus | 0.0148 | 0.2130 | WestEurasia | PRKCE |
| molecular_function | GO:0003785 | actin monomer binding | 0.0154 | 0.4989 | WestEurasia | PRKCE |
| molecular_function | GO:0004699 | calcium-independent protein kinase C activity | 0.0154 | 0.4989 | WestEurasia | PRKCE |
| molecular_function | GO:0030546 | signaling receptor activator activity | 0.0154 | 0.4989 | WestEurasia | PRKCE |
| biological_process | GO:0007600 | sensory perception | 0.0037 | 0.0676 | SouthAsia | ATP6V0A4, COL4A3 |
| **biological_process** | **GO:0007605** | **sensory perception of sound** | **0.0004** | **0.0231** | **SouthAsia** | **ATP6V0A4, COL4A3** |
| **biological_process** | **GO:0050954** | **sensory perception of mechanical stimulus** | **0.0004** | **0.0231** | **SouthAsia** | **ATP6V0A4, COL4A3** |
| cellular_component | GO:0000220 | vacuolar proton-transporting V-type ATPase, V0 domain | 0.0155 | 0.4254 | SouthAsia | ATP6V0A4 |
| cellular_component | GO:0005903 | brush border | 0.0155 | 0.4254 | SouthAsia | ATP6V0A4 |
| cellular_component | GO:0016471 | vacuolar proton-transporting V-type ATPase complex | 0.0155 | 0.4254 | SouthAsia | ATP6V0A4 |
| molecular_function | GO:0004866 | endopeptidase inhibitor activity | 0.0210 | 0.5722 | SouthAsia | COL4A3 |
| molecular_function | GO:0005178 | integrin binding | 0.0105 | 0.4920 | SouthAsia | COL4A3 |
| molecular_function | GO:0008191 | metalloendopeptidase inhibitor activity | 0.0105 | 0.4920 | SouthAsia | COL4A3 |
| biological_process | GO:0021536 | diencephalon development | 0.0070 | 0.5518 | America | CNTNAP2 |
| biological_process | GO:0021756 | striatum development | 0.0070 | 0.5518 | America | CNTNAP2 |
| biological_process | GO:0021761 | limbic system development | 0.0070 | 0.5518 | America | CNTNAP2 |
| cellular_component | GO:0008076 | voltage-gated potassium channel complex | 0.0063 | 0.3211 | America | CNTNAP2 |
| cellular_component | GO:0030673 | axolemma | 0.0063 | 0.3211 | America | CNTNAP2 |
| cellular_component | GO:0032589 | neuron projection membrane | 0.0063 | 0.3211 | America | CNTNAP2 |
| molecular_function | GO:0005488 | binding | 0.9747 | 0.7987 | America | CNTNAP2, FAM184A |
| molecular_function | GO:0005515 | protein binding | 0.8955 | 0.7987 | America | CNTNAP2, FAM184A |
| molecular_function | GO:0019899 | enzyme binding | 0.1605 | 0.7077 | America | CNTNAP2 |

#### Functional variants in candidate regions

##### Identification of variants in candidate genes

For each candidate region overlapping a gene, we computed the allele frequencies of variants that overlap candidate regions. We grouped the individuals into carriers (with a Denisovan segment overlapping the candidate region), and non-carriers (Africans and non-Africans, excluding carriers). We filtered the alleles overlapping candidate regions as follows: (i) alternative allele frequency <5% in non-carriers, (ii) carriers and Africans should not have the same allele frequency, and (iii) alternative allele frequency >0% in carriers. Allele frequencies are in **Supplementary Data 1**.

##### Annotation of variants

We annotated the variants in introgressed regions to identify those that may affect how cells function (**Table S70**). As variants at higher frequencies are more likely to have been retained by positive selection than by chance, we used a minimum allele frequency of 25% among carriers of the introgressed region, yielding 1228 variants for analysis. To further evaluate their potential biological impact, we examined where these variants fall relative to genes using a database of gene annotations (GENCODE, version 43^151^). The genomic context of variants provides clues about their functional roles: variants in exons can directly alter protein sequence, while intronic or intergenic variants may affect gene regulation and expression. Of the variants identified, 3.7% are in exonic regions, 49.9% in intronic regions, and 46.4% in intergenic regions of protein-coding genes. Within the coding region of exons, six variants are missense changes (**Figure S122**). Variants in non-coding RNAs also contribute to differences in gene regulation. 31.2% of variants overlap long non-coding or long intergenic non-coding RNAs, while none overlap microRNAs.

To identify variants that may affect gene expression, we overlapped them with regulatory regions. Variants located in regulatory regions can alter when, where, and the level at which genes are expressed. Such changes could affect cellular function and contribute to adaptation. 42.7% of variants are in an open chromatin region identified with DNase I hypersensitive site sequencing in 733 samples from 438 human tissues or cells (ENCODE phase 4)^152^ or from 53 human tissues or cells^153^. A smaller proportion are in promoter-proximal regions (within 1,000 base pairs upstream of transcription start sites; GENCODE^151^, version 43, MANE-select canonical transcripts), representing 1.5% of variants that may directly influence transcription initiation. In addition, 38.8% of variants are in regions that may act as enhancers, identified through combinations of 12 histone marks^153^, are transcribed enhancers^154^, or are human DNA sequences that enable transcriptional activity in developing mice^155^.

To further pinpoint variants that could alter gene expression, we examined their association with expression quantitative trait loci (eQTL) or splicing quantitative trait loci (sQTL) (GTEx, version 10)^156^. 4.5% of variants are significantly correlated with differences in gene expression, and about half of these are correlated with differences in alternative splicing of transcripts.

We also used measures of genetic conservation to prioritize variants that are more likely to have functional effects, as sequences that have been preserved within or across organisms are often biologically important and subject to evolutionary constraint. 0.9% of variants are in the most constrained percentile of the human genome, identified using constraint measurements in 1 kb windows in more than 75,000 human genomes in the genome aggregation database (gnomAD) (Gnocchi score ≥ 4)^157^. A different measure of constraint that takes mammalian genomes into account identifies 2.6% of variants in the top 5% constrained regions of the human genome (B-score < 50)^104^. Genetic conservation quantified at the nucleotide-level revealed that 4% of variants occur at positions significantly conserved in vertebrates and 4.7% in mammals (phyloP score ≥ 1.3, provided by the UCSC Genome Browser and calculated from the 100-way vertebrate genome alignment or the 447-way mammalian genome alignment that includes 223 primates)^158^.

To better understand how these variants may affect phenotypes, we overlapped them with genome-wide association studies (NHGRI-EBI GWAS Catalog, retrieved on 12 October 2024)^159^, which identified one variant associated with a trait. We also used gene ontology terms (retrieved on 23 October 2024 from geneontology.org)^160,161^ and human phenotype ontology terms (retrieved on 12 December 2024 from hpo.jax.org)^162^ to identify the cellular functions or disease associations of genes with variants. Gene expression profiles in tissues and cell types were obtained from The Human Protein Atlas (www.proteinatlas.org)^163,164^.

**Table S70: Annotation of single-nucleotide variants in candidate regions of adaptive introgression.**

| **Annotation of single-nucleotide variants (SNV)** | **SNV** | **% of all SNV** | **Genes** |
| --- | --- | --- | --- |
| Total SNV (Allele frequency ≥ 25% in carriers) | 1228 | 100.0 | . |
| Location relative to genes | . | . | . |
| Exonic | 45 | 3.7 | 22 |
| Coding sequence | 13 | 1.1 | 10 |
| Missense changes | 6 | 0.5 | 4 |
| 5 prime UTR | 6 | 0.5 | 6 |
| 3 prime UTR | 27 | 2.2 | 13 |
| Intronic | 613 | 49.9 | 48 |
| Intergenic | 570 | 46.4 | . |
| Long non-coding RNA | 383 | 31.2 | 25 |
| microRNA | 0 | 0.0 | 0 |
| Transcription-related | . | . | . |
| Open chromatin region | 524 | 42.7 | . |
| ENCODE phase 4 | 406 | 33.1 | . |
| Roadmap Epigenomics | 343 | 27.9 | . |
| Promoter-proximal | 18 | 1.5 | 12 |
| Enhancer | 477 | 38.8 | . |
| Histone marks HMM (Roadmap Epigenomics) | 477 | 38.8 | . |
| In vivo activity in developing mice (VISTA enhancers) | 1 | 0.08 | . |
| Transcribed enhancers (FANTOM5) | 6 | 0.5 | . |
| Expression quantitative loci (eQTL) | 55 | 4.5 | . |
| Splicing quantitative loci (sQTL) | 29 | 2.4 | . |
| Evolutionary constraint | . | . | . |
| Top 1% constrained regions of genome (gnomAD) | 11 | 0.9 | . |
| Top 5% constrained regions of genome (B-score) | 32 | 2.6 | . |
| Position conserved in vertebrates (phyloP) | 49 | 4.0 | . |
| Position conserved in mammals (phyloP) | 58 | 4.7 | . |
| Trait and disease associations | . | . | . |
| GWAS | 1 | 0.1 | . |

**Figure S122: Denisovan-like DNA segments overlapping genes in candidate regions where the introgressed variant causes an amino acid change.** Haplotypes are classified as either Denisova 3-like (blue) or ancestral (red), i.e. falling outside the variation of both Denisova 3 and Denisova 25. Green rectangles at the bottom indicate the positions and lengths of the gene overlapping the candidate region. Only the genes with missense changes are shown. Nucleotide triplet is indicated on the segments, in black if it is derived and gray if it is ancestral. Ancestral nucleotide triplets are also indicated in the genes.

##### Description of introgressed variants with functional annotations

###### Introgressed region in Americans

Region 6:119392450-119433612

A variant (6:119399619) in the 5' UTR of *FAM184A* occurs in a region that is among the top 1% constrained regions of the genome (Gnocchi score 5.5). The introgressed allele has a frequency of 40% in Native Americans (HGDP via gnomAD, version 4.1). The ancestral allele is significantly conserved in primates (p-value = 0.016): none of the 223 genomes carry the introgressed allele, while everyone carries the ancestral allele, aside from thirteen that have no base call. FAM184A is located in processing bodies of the cytoplasm that control messenger RNA degradation and translational repression (GO:0000932), as well as centriolar satellites involved in cell division and cilium assembly (GO:0034451).

###### Introgressed region in East Asians

Region 2:46343745-46409161

An introgressed allele (2:46349271) is associated with lower red blood cell count (0.063 to 0.092 unit decrease, p-value = 1e-24, beta = 0.077). The variant overlaps an enhancer region identified from histone marks in the heart ventricles, lung, mucosa of the stomach and colon, fetal and adult intestines, skeletal muscle, and the HeLa-S3 cervical cancer cell line. While the variant is in the intron of *PRKCE*, the nearest transcription start site is *EPAS1*, located 115,000 base pairs downstream of the introgressed region.

###### Introgressed regions in South Asians

Region 7:138479314-138501786

There is a variant (7:138482795) located in the 5' UTR of both *TMEM213* and *ATP6V0A4*, which are transcribed in opposite directions on different strands. The allele frequency of the introgressed allele is 58% in carriers.

Another variant (7:138482405) is in the promoter-proximal region of *TMEM213*, located 292 base pairs upstream the transcription start site. The introgressed allele would disrupt the binding of transcription factors ZIC2 and ZIC3, which are predicted to bind to the nine nucleotides encompassing the variant (p-value = 1e-4.5, JASPAR Core 2024 predicted transcription factor binding sites provided by the UCSC Genome Browser). *TMEM213* encodes transmembrane protein 213 and is one of only eight genes enriched for specific expression in the kidney and salivary gland (The Human Protein Atlas).

In addition, two variants located in *ATP6V0A4* overlap regions with histone marks indicative of enhancers in the blood, brain, embryo, endoderm, esophagus, mesendoderm, pancreas, thymus, and trophoblast. In several of these tissues, *ATP6V0A4* has many phenotype associations in humans, including abnormal: blood ion concentration (HP:0003111), metabolism/homeostasis (HP:0001939), digestive system physiology (HP:0025032), renal morphology (HP:0012210), distal renal tubular acidosis (HP:0008341), upper urinary tract (HP:0010935), urine calcium concentration (HP:0011280), fluid regulation (HP:0011032), skeletal morphology (HP:0011842), reduced bone mineral density (HP:0004349), decreased body weight (HP:0004325), growth delay (HP:0001510), inner ear (HP:0011389), and bilateral sensorineural hearing impairment (HP:0008619). The gene encodes ATPase H+ transporting V0 subunit a4, which has proton-transporting ATPase activity (GO:0046961). It is involved in the regulation of pH (GO:0006885), renal tubular secretions (GO:0097254), ossification (GO:0001503), the sensory perception of sound (GO:0007605), and synaptic vesicle lumen acidification (GO:0097401).

Region 2:228062789-228095040

Seven introgressed variants are eQTL where the introgressed alleles are correlated with lower expression of *MFF* in the cerebral cortex, esophagus muscularis, skeletal muscle, and fibroblasts. The variant at 2:228072045 has the lowest p-value (4.4e-6) and largest normalized effect size (-0.33) occurring in the cerebral cortex. The introgressed allele has a frequency of 31% in carriers. The gene codes for mitochondrial fission factor which localizes to the mitochondrial outer membrane (GO:0005741) and controls the division of mitochondria. It positively regulates the fission of mitochondria (GO:0090141) and peroxisomes (GO:0016559). It is also a component of synaptic vesicles (GO:0008021). *MFF* has many human phenotype associations with the nervous system, including abnormality of central nervous system electrophysiology (HP:0030178), microcephaly (HP:0000252), neurodevelopmental delay (HP:0012758), upper motor neuron dysfunction (HP:0002493), motor delay (HP:0001270), muscle weakness (HP:0001324), weakness of the muscles that move the eye (HP:0000602) and trunk (HP:0009062), inability to walk (HP:0002540), epileptic spasm (HP:0011097), delayed speech and language development (HP:0000750), sensory behavioral abnormality (HP:5200046), visual impairment (HP:0000505), auditory sensitivity (HP:0025112). The gene also affects head circumference (HP:0040195) and gastrointestinal tract function (HP:0012719).

There are an additional nine introgressed alleles that are eQTL correlated with higher expression of the long non-coding RNA MFF-DT in fibroblasts (p-value = 7.4e-6, normalized effect size = 0.24). One of these variants (2:228079834) overlaps a putative enhancer region identified in both bone marrow and chondrocyte cells derived from mesenchymal stem cells.

###### Introgressed regions in West Eurasians

Region 2:228243909-228457183

The gene *TM4SF20* has an introgressed variant (2:228244107) in the promoter-proximal region located 86 base pairs upstream the transcription start site. This position is significantly conserved in 447 mammalian genomes (p-value = 1.3e-5) and every primate genome carries the ancestral allele. Relative to the ancestral allele, the introgressed allele is associated with lower expression of *TM4SF20* in fibroblasts (p-value = 3.2e-13, normalized effect size = -0.45). It is found at an allele frequency of 55% in carriers. *TM4SF20* encodes transmembrane 4 L six family member 20, which localizes to the plasma membrane (GO:0005886), endoplasmic reticulum membrane (GO:0005789), and focal adhesions (GO:0005925), where it down-regulates proteolysis (GO:0045861). The gene is expressed almost exclusively in the intestines, particularly in proximal enterocytes (The Human Protein Atlas). However, its phenotype associations in humans are related to nervous system physiology (HP:0012638) and morphology (HP:0012639), hyperintensity of cerebral white matter on MRI (HP:0030890), delayed speech and language development (HP:0000750), abnormal speech pattern (HP:0002167), and autistic behavior (HP:0000729).

The promoter-proximal introgressed allele is also correlated with lower expression of nearby genes: *MFF* in fibroblasts (p-value = 9.3e-12, normalized effect size = -0.33) and *AGFG1* in fibroblasts, tibial artery, testis, and the cerebellum (p-value = 1.6e-9, normalized effect size = -0.43). However, it is correlated with higher expression of *MFF-DT* in fibroblasts, tibial nerves, sun exposed skin, subcutaneous adipose, stomach, pancreas, and the sigmoid colon (p-value = 5.1e-7, normalized effect size = 0.48). Lower expression of *MFF* and higher expression of *MFF-DT* is also associated with introgressed alleles on a different haplotype in South Asians (described above).

There are fourteen other introgressed variants that are eQTLs with similar metrics for these four genes. Nearly all overlap open chromatin regions and putative enhancers, suggesting the variants may alter how the genes are regulated. Ten of the eQTL are also correlated with alternative splicing of *AGFG1* in the testis, with position 2:228275698 being the most significant (p-value = 6.5e-9, normalized effect size = 0.76). The gene, which encodes ArfGAP with FG repeats 1, is involved in spermatid nucleus differentiation (GO:0007289) and assembly of the acrosome in sperm cells, which contains enzymes for penetrating egg cells during fertilization (GO:0001675). AGFG1 is also part of the nuclear pore (GO:0005643), where it is involved in mRNA export from the nucleus (GO:0006406). The protein serves as a co-factor in human immunodeficiency virus (HIV-1) replication by facilitating the movement of viral RNA from the nucleus to the cytoplasm for replication (The Human Protein Atlas).

###### Introgressed regions in Oceanians

Region 3:13325993-13519969

Introgressed variants overlap several parts of the gene *NUP210* that encodes nucleoporin 210, which is involved in transporting mRNA (GO:0051028) and proteins (GO:0015031) as part of the nuclear pore (GO:0005643). One variant (3:13363815) is a derived missense change where glycine replaces glutamic acid at amino acid position 1598. The ancestral allele at this position is significantly conserved in vertebrates (p-value = 0.0036). The introgressed allele is at a frequency of 50% in carriers. There are also two variants (3:13357955; 3:13358771) in the 3' UTR. Another variant (3:13462283) overlaps the promoter-proximal region in one of the highest constrained regions of the genome (top 1%, Gnocchi score 4.8). There is also a variant (3:13455743) in a transcribed enhancer active in T cells, dendritic cells, monocytes, and natural killer cells.

Region 3:133332322-133475438

Two introgressed variants introduce missense changes in *TOPBP1*: the first (3:133336155) results in a cysteine to tyrosine substitution at amino acid position 981, the second (3:133342288) results in a glycine to alanine substitution at amino acid position 1203. The allele frequency in carriers is 72%.

*TOPBP1* also has three variants in the promoter-proximal region. At one position (3:133381647), the introgressed allele would likely disrupt binding of transcription factors PHOX2B, POU4F1, POU4F2, POU4F3, CDX4, HOXD9, HOXA10, and HOXB13 (JASPAR Core 2024 predicted transcription factor binding sites provided by the UCSC Genome Browser). Introgression reintroduces the ancestral state of the allele, whereas the derived allele is found at a frequency of >99.9% globally among present-day people (gnomAD 4.1) and shared with high-coverage Neandertal genomes (Prüfer *et al*., 2014; Prüfer *et al*., 2017; Mafessoni *et al*., 2020).

Two other introgressed variants (3:133356300, 3:133366166) in the intron of *TOPBP1* also re-introduce the ancestral allele at positions where derived alleles arose on the modern human lineage and are found at frequencies >99.9% in present-day people. *TOPBP1* encodes DNA topoisomerase II binding protein 1. As part of the spindle pole (GO:0000922), it is involved in checkpoint signaling during DNA replication (GO:0000076). It is also active at sites of DNA damage (GO:0090734), participating in double-strand break repair via homologous recombination (GO:0000724).

The gene *TF* also has two promoter-proximal variants (3:133464468, 3:133464851), one of which would likely disrupt binding of the transcription factors ZNF75A and ZNF677 that are predicted to bind to the locus (JASPAR Core 2024 predicted transcription factor binding sites provided by the UCSC Genome Browser). The alleles have a frequency around 40% in carriers. *TF* encodes transferrin, which transports iron (GO:0006826) and regulates intracellular iron homeostasis by binding to transferrin receptors on the cell surface (GO:1990459). Mutations in this gene can result in anemia (HP:0001903), indicating it influences red blood cell and hemoglobin concentrations. TF is also involved in the antibacterial humoral response (GO:0019731), osteoclast differentiation (GO:0030316), positive regulation of bone resorption (GO:0045780), and arthritis (HP:0001369).

Additionally, three introgressed alleles are eQTL correlated with higher expression of *TF* in breast mammary tissues (p-value = 2.7e-05, normalized effect size = 0.65), but lower expression of the nearby gene *SRPRB* in skeletal muscle (p-value = 8.4e-07, normalized effect size = -0.30) (3: 133434021). The introgressed alleles are found at an allele frequency of 32% in carriers. *SRPRB* encodes signal recognition particle receptor subunit beta, which is involved in protein targeting to the endoplasmic reticulum (GO:0045047) where it is embedded in the membrane (GO:0005789).

Region 3:133479642-133575618

The gene *SRPRB* (described above) has two introgressed variants in the promoter-proximal region (3:133524282, 3:133524581). Furthermore, lower expression of *SRPRB* is associated with three introgressed alleles (3:133561587, 3:133564857, 3:133566350) in skeletal muscle, subcutaneous adipose, heart atrial appendage, and the esophagus (gastroesophageal junction and muscularis). The most significant eQTL is in the gastroesophageal junction of the esophagus (p-value = 7.62e-6, normalized effect size = -0.42). The three eQTL variants are located downstream the transcription start site, whereas the eQTL of *SRPRB* at the locus described above are located upstream the transcription start site. At two of the positions, the ancestral alleles are conserved in vertebrates (phyloP = 1.3). The introgressed derived alleles are found at frequencies ranging from 70-75% in carriers.

In addition, there are numerous introgressed variants in 3' UTR of the following genes: two in *SRPRB*, two in *TF* (described above), and three in *RAB6B*. The gene *RAB6B* is located in the Golgi membrane (GO:0000139), where it is involved in retrograde transport from the endosome to the Golgi (GO:0042147) and retrograde vesicle-mediated transport to the endoplasmic reticulum (GO:0006890). It is also located in the presynapse (GO:0098793) and involved in neuron projection development (GO:0031175). It has GTPase activity (GO:0003924) and enables myosin V binding (GO:0031489).

Region 5:35904806-36034020

Two missense variants occur in the gene *UGT3A1* resulting in a glycine to valine substitution at amino acid positions 103 (5:35968124) and an arginine to tryptophan substitution at amino acid position 411 (5:35955811). The introgressed allele frequency in carriers at the first variant is 59%, whereas it is 73% for the second variant.

*UGT3A1* also has two variants in the 3' UTR (5:35952902, 5:35953351) with allele frequencies around 73% in carriers. This gene encodes UDP glycosyltransferase family 3 member A1, which has glucuronosyltransferase activity (GO:0015020) as part of the UDP-N-acetylglucosamine transferase complex (GO:0043541). It is most highly expressed in the liver and kidneys (The Human Protein Atlas).

Region 6:132943935-133036518

There are two variants in the 3' UTR of *VNN1* (6:133003135, 6:133004018). The introgressed allele frequency in carriers of the first position is 29%, but nearly twice as high at 61% for the second position. *VNN1* encodes vanin 1 that plays a role in the metabolism of coenzyme A (GO:0015938) by cleaving pantetheine to form vitamin B5. It is also involved in the innate immune response (GO:0045087) and the inflammatory response- both acute (GO:0002526) and chronic (GO:0002544). Furthermore, it positively regulates T cell differentiation in the thymus (GO:0033089). The gene is most highly expressed in the liver, gallbladder, and duodenum; in blood cells, it is most highly expressed in monocytes (The Human Protein Atlas).

Region 6:137935444-138101913

Three variants (6:138020511, 6:138037108, 6:138039111) overlap transcribed enhancers active in T cells, dendritic cells, mast cells, mesenchymal cells, monocytes, or natural killer cells (FANTOM5). The first two positions overlap the long intergenic non-coding RNA *LINC03004*, where the second position is in a 20-bp region significantly conserved in vertebrates (phastCons, p-value < 0.01). The second and third position have an allele frequency of 82% in Oceanians (HGDP via gnomAD, version 4.1).

Region 6:154688569-154726767

The 3' UTR of *CNKSR3* contains four variants where the introgressed derived alleles are found at frequencies of 35-40% in carriers. At one position (6:154713000), the ancestral allele is conserved in vertebrates (p-value = 0.03). *CNKSR3* encodes the connector enhancer of kinase suppressor of ras 3. It is located in the apical plasma membrane (GO:0016324) where it positively regulates sodium ion transport (GO:0010765), but negatively regulates the ERK1 and ERK2 cascade (GO:0070373) as well as peptidyl-serine phosphorylation (GO:0033137).

Region 10:79529618-79644112

*DLG5*, encoding discs large MAGUK scaffold protein 5, has an introgressed variant (10:79551987) in the 3' UTR that occurs at an allele frequency of 67% in carriers. Another variant (10:79576772) is an eQTL where the introgressed allele is correlated with increased *DLG5* expression in the thyroid (p-value = 1.17e-06, normalized effect size = 0.56) and whole blood (p-value = 8.01e-06, normalized effect size = 0.28). The introgressed allele is also correlated with alternative splicing of *DLG5* in the thyroid (p-value = 1.3e-9, normalized effect size = -0.97), as well as in esophagus muscularis, subcutaneous adipose, and sun exposed skin of the lower leg.

*DLG5* has several functions. It is involved in intracellular signal transduction (GO:0035556)- particularly hippo signaling (GO:0035331, GO:0035332), smoothened signaling (GO:0045880), and beta-catenin binding (GO:0008013)- regulating cell proliferation (GO:0008285), migration (GO:0030336), and apoptosis (GO:0042981). Located at cell junctions (GO:0030054), the gene is involved in establishing and maintaining epithelial cell apical basal polarity (GO:0045197) and the epithelial to mesenchymal transition (GO:0001837). In the brain, it is located in glutamatergic synapses (GO:0098978) and the postsynaptic density (GO:0014069) where it contributes to postsynapse organization (GO:0099173) and dendritic spine development (GO:0060999). It also has a role in midbrain development (GO:0030901) and neuroepithelial cell differentiation (GO:0060563). Aside from the brain, the gene is involved in processes of other organs, including epithelial tube branching involved in lung morphogenesis (GO:0060441), metanephric collecting duct development (GO:0072205), and negative regulation of T cell proliferation (GO:0042130).

Given its role in multiple systems, mutations in *DLG5* are associated with numerous phenotypes in humans: coarse facial features (HP:0000280), cleft palate (HP:0000175), upper-limb joint contracture (HP:0100360), joint contracture of the hand (HP:0009473), split hand (HP:0001171), broad palm (HP:0001169), dystonia (HP:0001332), nervous system morphology (HP:0002011) and physiology (HP:0012638), hydrocephalus (HP:0000238), septal defects of the heart atria and ventricles (HP:0001631, HP:0001629), multicystic kidney dysplasia (HP:0000003), and dilatation of the renal pelvis (HP:0010946).

Region 12:98785180-99013277

There is a variant (12:98943974) in the 3' UTR of *TMPO* where the introgressed derived allele is found at 61% frequency in carriers. The ancestral allele is significantly conserved in vertebrates (p-value = 0.0095). This gene encodes thymopoietin that is involved in binding of lamin (GO:0005521), cadherin (GO:0045296), and DNA (GO:0003677) as part of the nuclear inner membrane (GO:0005637). It has associations with numerous heart-related phenotypes including cardiovascular system morphology (HP:0030680), left ventricular systolic dysfunction (HP:0025169), dilated cardiomyopathy (HP:0001644), arrhythmia (HP:0011675), congestive heart failure (HP:0001635), and thromboembolic stroke (HP:0001727). It is also involved in metabolism/homeostasis (HP:0001939), loss of fat tissue (HP:0100578), fatigue (HP:0012378), breathing difficulty that occurs with exercise or exertion and improves with rest (HP:0002875), the sensation of breathlessness when lying down that is relieved by sitting or standing (HP:0012764), muscle physiology and morphology (HP:0011804, HP:0011805), functional abnormality of the inner ear (HP:0011389), and sensorineural hearing impairment (HP:0000407).

Another introgressed allele (12:98785374) in an intergenic region is correlated with increased expression of *TMPO* (p-value = 4.05e-6, normalized effect size = 0.42) and *SLC9A7P1* (p-value = 2.61e-6, normalized effect size = 0.52) in the pancreas. The allele is also correlated with increased expression of long intergenic non-coding RNA *LINC02453* in the tibial nerve and skeletal muscle (p-value = 7.7e-8, normalized effect size = 0.47). This gene has been shown to inhibit HIV-1 replication by regulating the viral productive cycle (Chen *et al*., 2024).

Additionally, there are two variants (12:98997300, 12:98997628) in the 3' UTR of *SLC25A3* where the introgressed allele is found at a frequency of 75% in carriers. This gene encodes solute carrier family 25 member 3, which is located in the plasma membrane (GO:0005886), extracellular exosome (GO:0070062), and mitochondrial inner membrane (GO:0005743) where it has phosphate transmembrane transporter activity (GO:0005315). In humans it is associated with mitochondrial shape (HP:0012087), metabolism/homeostasis (HP:0001939), circulating lactate concentration (HP:0002151), lactic acidosis (HP:0003128), cardiovascular system physiology (HP:0011025), myocardium morphology (HP:0001637), congestive heart failure (HP:0001635), pattern of respiration (HP:0002793), breathing difficulty (HP:0002094), skin morphology (HP:0011121), cyanosis (HP:0000961), muscle physiology (HP:0011804) and morphology (HP:0011805), muscle tone (HP:0003808), and body weight (HP:0004325).

Region 17:37387412-38082831

*ERBB2*, encoding Erb-B2 receptor tyrosine kinase 2, carries a missense change (17:37873628) with the introgressed allele resulting in an alanine to aspartic acid substitution. The allele is found at a frequency of 58% in carriers. The ancestral allele is significantly conserved in mammals and primates (p-value = 8.55e-9). The gene is part of the ERBB2-ERBB3 (GO:0038133) and ERBB2-EGFR signaling pathways (GO:0038134) where it serves as a cell surface receptor protein (GO:0007169) and positively regulates protein phosphorylation (GO:0001934). It is involved in the development of the heart (O:0007507), neuromuscular junctions (GO:0007528), neurons (GO:0030182), and Schwann cells (GO:0014044). The gene is also involved in motor neuron axon guidance (GO:0008045), neurotransmitter receptor localization to the postsynaptic membrane (GO:0099645), immature T cell proliferation in the thymus (GO:0033080) and wound healing (GO:0038134). There are numerous phenotype associations in humans, including morphology of the central and peripheral nervous system (HP:0002011, HP:0000759), gastrointestinal tract (HP:0012718), foot (HP:0001760), limbs (HP:0040064), upper eyelid (HP:0000508), lung (HP:0002088), trachea (HP:0005607), respiratory system (HP:0012252), large intestine (HP:0002250), stomach (HP:0002577), enteric ganglion (HP:0004362), and glial cells (HP:0100705). The gene influences body weight (HP:0004325), height (HP:0000002), urine metabolite levels (HP:0033354), peripheral axonal degeneration (HP:0000764), inflammatory response (HP:0012649), gastrointestinal inflammation (HP:0004386), abdominal pain (HP:0002027), abnormality of the eye (HP:0000478), functional abnormality of the inner ear (HP:0011389), and sensorineural hearing impairment (HP:0000407).

Another variant (17:37827612) overlaps the 3' UTR of *PGAP3*. The introgressed allele is found at a frequency of 46% in carriers. An intronic variant (17:37832950) that is significantly conserved in mammals (p-value = 0.009) overlaps an enhancer active in the developing heart. *PGAP3* encodes post-GPI attachment to proteins phospholipase 3. It is located in the membrane of the Golgi (GO:0000139) and endoplasmic reticulum (GO:0005789), where it is involved in the formation of a glycosylphosphatidylinositol (GPI) anchor that attaches some membrane proteins to the lipid bilayer of the cell membrane (GO:0006506) and has hydrolase activity acting on ester bonds (GO:0016788). This gene is associated with 286 phenotypes in humans, many of which are related to craniofacial or skeletal morphology: coarse facial features (HP:0000280), thin upper lip vermilion (HP:0000219), tented upper lip vermilion (HP:0010804), downturned corners of mouth (HP:0002714), cleft palate (HP:0000175), upward slanting of the opening between the eyelids (HP:0000582), eye folds (HP:0000286), highly arched eyebrow (HP:0002553), bulbous nose (HP:0000414), wide nasal bridge (HP:0000431), broad nasal tip (HP:0000455), cupped ear (HP:0000378), large earlobe (HP:0009748), high palate (HP:0000218), bifid uvula (HP:0000193), small lower jaw (HP:0000347), chin prominence (HP:0000303), head circumference (HP:0040194, HP:0040195), skull morphology (HP:0000929), round face (HP:0000311), short phalanx of finger (HP:0009803), shortening of all distal phalanges of the fingers (HP:0006118), small nail (HP:0001792), and funnel chest (HP:0000767). It is also associated with autistic behavior (HP:0000729) and language impairment (HP:0002463).

The gene *IKZF3* also contains two variants in the 3' UTR (17:37915360, 17:37921258). Another variant (17:37926671) in an intron of *IKZF3* overlaps a transcribed enhancer active in T cells and natural killer cell; the introgressed allele would likely disrupt binding of transcription factor RUNX1 that is predicted to bind to this locus (JASPAR Core 2024 predicted transcription factor binding sites provided by the UCSC Genome Browser). The three introgressed alleles are found at frequencies of 54-58% in carriers. The gene encodes Ikaros family zinc finger protein 3, a transcription factor (GO:0001228) involved in the differentiation of B cells (GO:0030183) and T cells (GO:0030217) in response to bacteria (GO:0009617).

In addition, there are 16 introgressed alleles that are eQTL, many which are in highly constrained regions of the genome and have an allele frequency of 26-29% in carriers. They are correlated with expression differences in 32 different tissues of the following genes (reporting the lowest p-value and the normalized effect size for the introgressed allele): *STARD3* (4.85e-15, -0.55), *STAC2* (1.50e-13, 0.45), *GSDMB* (4.93e-13, -0.66), *FBXL20* (7.89e-11, -0.35), *GSDMA* (1.26e-08, 0.62), PGAP3 (3.18e-08, -0.24), *ENSG00000214546* (3.13e-07, -0.39), *ENSG00000264968* (1.03e-06, 0.42), *MIEN1* (1.20e-06, -0.28), *ORMDL3* (1.24e-06, -0.22), *PLXDC1* (1.90e-06, 0.51), *MED24* (4.56e-06, 0.21), *PNMT* (9.46e-06, 0.23), *GRB7* (1.72e-05, -0.24), *LINC00672* (7.79e-05, 0.86), *ZPBP2* (1.35e-04, 0.18), *MED1* (1.47e-04, 0.08), *ENSG00000266469* (2.06e-04, -0.42), *CASC3* (2.83e-04, 0.16). All 16 are also correlated with alternative splicing of the following genes: *GSDMB* (2.02E-34, 1.69), *STARD3* (2.92E-17, -2.15), *GRB7* (7.30E-11, 0.72), *ERBB2* (1.60E-09, 0.78), *PGAP3* (1.36E-07, -0.50), *ZPBP2* (6.93E-06, -0.64).

Region 19:51967869-52036108

The promoter-proximal region of *SIGLEC6* has three variants (19:52035229, 19:52035462, 19:52035925) and that of *SIGLEC12* has one variant (19:52005696). The introgressed allele near *SIGLEC12* has an allele frequency of 81% in carriers, while those near *SIGLEC6* have frequencies of 53-61%. In Oceanians, the *SIGLEC6* alleles have allele frequencies of 89% (HGDP via gnomAD, version 4.1). These genes encode Sialic Acid Binding Ig Like Lectin 6 and 12, which bind sialic acid (GO:0033691) and are involved in cell adhesion (GO:0007155). *SIGLEC6* is mostly expressed in syncytiotrophoblasts and cytotrophoblasts of the placenta, where it is up-regulated in preeclampsia, a condition in pregnancy characterized by high blood pressure (The Human Protein Atlas). *SIGLEC12* is mostly expressed in lymphoid tissues, particularly the spleen, as well as the intestines and lung. Both genes are expressed in blood granulocytes, while *SIGLEC12* is also expressed in macrophages, Kupffer cells, and Hofbauer cells.

Region 21:39632604-39687139

*KCNJ15*, encoding potassium inwardly rectifying channel subfamily J member 15, has two variants (21:39675096, 21:39675367) in the 3' UTR where the introgressed allele is found at frequencies of 71% in carriers. It is involved in potassium ion import across the plasma membrane (GO:1990573).

#### Potential candidate genes of recurrent adaptive introgression

The sharing of candidate regions between superpopulation can be a sign of recurrent independent selection of these variants. There are only two candidates which overlap each other, found in East-Asians and West-Eurasians on chromosome 2 overlapping the *PRKCE* gene (**Figure S123**). Interestingly this gene is only ~100 kb downstream from the *EPAS1* gene, thought of being adaptively introgressed from Denisovan and playing a role in high altitude adaptation in Tibetans^145^. However, it is not believed that West-Eurasians obtained their Denisovan-like DNA from an independent event, rather it was most likely contributed by gene flow from East-Asia^165^ (**Supplementary Note 11**). This is also supported by the observation that the segments overlap between the superpopulations, indicating a common origin. One explanation therefore could be that East-Asians, having the Denisovan-like DNA segments already in high frequency due to selection, simply contributed them to West-Eurasians in a high proportion without additional selection in West-Eurasians. The frequency of Denisovan-like DNA segments in East-Asians is 23% and 12% in West-Eurasians; this would require a high admixture proportion of East-Asians into West-Eurasians of around 50%. Given the proportions between the two populations estimated from present-day individuals are between 2-4%^166^, a more likely explanation would be that the segments were contributed by East-Asians in a lower proportion and selected recurrently in West-Eurasian to reach the frequency seen today. However, a contribution from an unsampled ancient East-Asian population with much higher Denisovan frequency at this locus cannot be excluded.

Functional analysis revealed a variant (T -> C) on the introgressed Denisovan-like DNA segments associated with lower red blood cell count. This variant is not seen in the SGDP Sub-Sahrana-Africans nor in the sequenced Neandertals or Denisovan, and might be a variant specific to the introgressing Denisovan population. The variant is shared by all individuals with a Denisovan-like DNA segment at this site, however the variant is outside the candidate region in West-Eurasians.

Another possible sign of recurrent selection is the presence of introgressed DNA from different Denisovan populations at the same genomic location. Therefore, we investigated regions harbouring Denisovan-like DNA segments that are “mixed”, focussing on those with at least 10 introgressed segments (enough segments to confidently call the mixed category). Only three such regions were identified in Oceanians, one on chromosome 6 overlapping *TAAR1*, *TAAR2*, *HLFP1* and *CCNG1P*, another on chromosome 20 overlapping *ATP9A*, *RP5-827A12.2* and *SALL4* and one on chromosome 3 overlapping *TF* and *RAB6B*.

As a segment could be differently classified based only on few differences to *Denisova 3* and *Denisova 25*, we assessed the certainty of the classification by measuring the difference between the match rate of each introgressed segment to *Denisova 3* to the match rate of *Denisova 25* to *Denisova 3* at the same locus. A small difference indicates uncertainty, while a large difference suggests high confidence. These candidates clearly correspond to Denisovan haplotypes of different origins.

When looking for functional variants in the candidate region, we found the derived state of two SNPs and one eQTL shared by a majority of individuals (≥75%) carrying Denisovan-like DNA segment at these loci for the candidate on chromosome 3 and one derived SNP shared by all at the candidate region on chromosome 6 (**Figure S124**). The different Denisovan-like haplotypes share these variants. If these variants would be the cause for the selection they would be selected recurrently irrespective of the haplotype.

**Figure S123: Individuals carrying Denisovan-like DNA segments overlapping candidate regions between West-Eurasians and East-Asians.** Haplotypes are classified as either Denisova 3-like (blue) or ancestral, i.e. falling outside the variation of both Denisova 3 and Denisova 25 (red). Green rectangles at the bottom indicate the positions and lengths of genes overlapping the candidate region. Grey box indicates the location of the candidate region. The derived states of the functional variant are indicated with black dots.

**Figure S124: Denisovan-like DNA segments at two candidate regions for adaptive introgression with distinct Denisovan haplotypes.** Haplotypes are classified as either Denisova 3-like (blue) or ancestral, i.e. falling outside the variation of both Denisova 3 and Denisova 25 (red). The shading intensity of the haplotypes reflects the certainty of the classification, with darker colours indicating higher confidence. Green rectangles at the bottom indicate the positions and lengths of genes overlapping the candidate region. Grey box indicates the location of the candidate region. Derived states of functional variants are indicated with filled dots for derived SNPs in the UTR and circles for eQTLs.

### Supplementary Note 13: A catalogue of Denisovan-derived single nucleotide changes

Little phenotypic information is available for Denisovans, as few remains have been attributed to this group. However, the Denisovan genome has the potential to provide such information. Using the *Denisova 3* genome, a catalogue of 692,818 single nucleotide changes (SNCs) that may be specific to Denisovans has previously been assembled^47^. Since only a single Denisovan genome was available, only a subset of these are shared among all Denisovans: a later analysis of the DNA sequences retrieved from three additional Denisovans showed that they carry the same base as *Denisova 3* at only 53.8% to 76.2% of the positions^61^. The availability of a second high-coverage Denisovan genome allows us to narrow down the list of changes that may have arisen specifically on the Denisovan lineage before the ancestors of *Denisova 3* and *Denisova 25* separated over 200,000 years ago.

We first applied the standard filters described in **Supplementary Note 3** to the high-coverage genomes of *Denisova 3*, *Denisova 5* (Neandertal), *Chagyrskaya 8*, *Vindija 33.19*, and *Denisova 25*, independently, using the snpAD genotype calls from this study, Prüfer *et al*. (2017)^45^, and Mafessoni *et al*. (2020)^91^. Positions not passing these filters were marked as missing (“N”).

We then identified sites where either one of the Denisovans is homozygous and differs from all high-coverage Neandertals, one Mbuti (HGDP00456)^47^ and four great ape reference genomes (panTro4, panpan1.1, gorGor3, ponAbe2). One allele was chosen randomly at heterozygous positions in the Mbuti genome and we required that at least one Neandertal had a genotype call, and that a base was available in at least two great ape reference genomes.

We next excluded positions where the derived allele in *Denisova 3* or *Denisova 25* was shared with African individuals from the 1000 Genomes dataset (Phase 3)^129^, excluding African-Americans (ASW) and African-Caribbeans (ACB). We further removed sites for which we could not reliably identify the ancestral allele, defined as that shared by a majority of great apes (at least three, or two if some were missing data) and present in at least one genome from a Neandertal or the Mbuti. This resulted in 378,986 and 285,303 putative Denisovan-derived SNCs using *Denisova 3* and *Denisova 25*, respectively. The catalogue of SNCs identified with *Denisova 3* is smaller than the published catalogue as we can now use three high-coverage Neandertal genomes and also used only African genomes instead of all the genomes from the 1000 Genomes dataset.

Finally, we compared the SNCs identified with *Denisova 3* or *Denisova 25*, and identified 245,991 SNCs that are shared by both Denisovan genomes, allowing a missing genotype in one. We then used genotypes from a larger set of individuals^167^ to restrict to sites where the Denisovan-specific SNCs are at <0% frequency in any African population (Human Genome Diversity Project: Bantu Kenya, Bantu South Africa, Biaka, Mandenka, Mbuti, San, Yoruba; 1000 Genomes Project: Esan, Gambian, Luhya, Mende, Yoruba). This resulted in a final set of 237,112 SNCs shared by *Denisova 3* and *Denisova 25*.

#### Single nucleotide changes affecting the sequence of genes

We annotated the SNCs to determine their potential functional impact based on their location relative to gene features (GENCODE, version 43, all transcript isoforms)^151^ (**Table S71**). For protein-coding genes, 3.0% of SNCs are in exons, 42.7% in introns, and 54.3% in intergenic regions. Within exons, 2.0% of all SNCs are in 5’ or 3’ untranslated regions (UTR), while 1.1% are in translated regions.

Those in translated regions may result in missense changes that alter the amino acid sequence of proteins, potentially affecting their structure, stability, or function. We find 1,307 Denisovan-derived missense changes in 1,142 genes (dbNSFP)^168^. Ten missense changes were classified as potentially impactful by PrimateAI-3D (version 1.0) (PrimateAI-3D score between 0.8 and 1), a deep learning approach that predicts the impact of missense changes by combining nucleotide conservation in 811 individuals from 236 primate species with protein 3D structure information (**Table S72**)^169^. For five of the changes, the Denisovan-derived allele is not observed in any present-day human genomes from a database of 807,162 individuals (gnomAD, version 4)^170^. The other half are found at very low allele frequencies, with the highest frequency being 0.02%.

Furthermore, four SNCs result in a start codon loss, one SNC results in a stop codon loss, and eleven SNCs result in a stop codon gain (Ensembl Variant Effector Predictor) (**Table S73**). Such changes may potentially produce truncated or extended proteins and disrupt normal cellular function.

Apart from protein-coding genes, 18% of SNCs overlap long non-coding RNA or long intergenic non-coding RNA, and 14 SNCs overlap microRNAs, which play key roles in transcriptional and post-transcriptional regulation.

#### Single nucleotide changes potentially affecting gene expression

Changes in gene expression caused by single nucleotide variants can influence gene regulatory networks, with downstream effects on phenotypes and organismal fitness. To identify SNCs that may influence RNA expression, we overlapped the catalogue with expression quantitative trait loci (eQTL) (GTex Consortium, version 10)^156^, splicing quantitative trait loci (sQTL) (GTex Consortium, version 10), and putative regulatory regions in humans^152-155^ (**Table S74**).

There are 658 positions within eQTL associated with gene expression variability in humans, of which 366 match the Denisovan-derived allele (305 where both Denisovan genomes have the derived allele call). Expression differences of 139 protein-coding genes are correlated with alleles derived in both Denisovan genomes in 19 tissues (***Figure S125***). Many of the eQTL are also correlated with alternative splicing of transcripts. Of the 321 positions within sQTL, 149 match the Denisovan-derived allele (117 where both Denisovan genomes have the derived allele call) and 73% of these are also eQTL.

In open chromatin regions, which correspond to regulatory regions that are accessible for transcription factor binding, there are 95,113 SNCs (40.1% of all SNCs)^152,153^. To further characterize regulatory regions, we overlapped the SNCs with promoter and enhancer regions that can influence when, where, and the level at which genes are expressed. Promoter-proximal regions, classified here as 1000 base-pairs upstream the transcription start site, contain 4,272 SNC that may impact transcription initiation of 3,510 genes. Enhancer regions, which can be identified with histone modifications (hidden Markov model using 12 histone marks from Roadmap Epigenomics Consortium *et al*., 2015^153^), reporter assay activity (VISTA enhancers from Visel *et al*., 2007^155^), or transcribed enhancers (FANTOM5 eRNA from Andersson *et al*., 2014^154^), contain 80,831 SNCs (34.1% of all SNCs).

As transcription factors recognize specific motifs to bind within regulatory regions and exert effects on gene expression, we used a set of 2,060 motifs for 615 transcription factors to identify 19,625 SNCs (8.3% of all SNCs) predicted to disrupt or create a transcription factor binding site using the tool *FunSeq2*^171^. 2,474 of these SNCs are in a region highly conserved in 100 vertebrate genomes (phastCons score > 0.95), of which 917 SNCs are in a highly conserved base (phyloP score > 1.3).

#### Single nucleotide changes in regions with evolutionary relevance

We next identified SNCs that may have been important in human evolutionary history by overlapping the catalogue of Denisovan-derived SNCs with conserved regions and those with signatures of Denisovan introgression or lack thereof (**Table S74**).

Regions with low levels of genetic variation in humans or conserved for millions of years among mammalian and vertebrate genomes are likely functionally important, as evolutionary conservation indicates selective pressure to maintain sequences essential for proper gene function and organismal fitness. In constrained regions of the genome identified using patterns of genetic variation across 76,156 human genomes, there are 3,333 SNCs in the most constrained percentile (Gnocchi score ≥ 4)^157^. 126 are missense changes, 53 are in the 5’ UTR, 138 are in the 3’ UTR, 1,994 are in intronic regions, and 881 are in intergenic regions. In “ultra-conserved” regions of the genome, there are two SNCs in regions where the sequence is identical in at least 200 base-pairs of the human, mouse, and rat genomes^172^, and two SNCs in regions where the sequence is identical in at least 20 base-pairs across 235 mammalian genomes (UCSC Genome Browser via Christmas *et al*., 2023^173^) (**Table S75)**. Nucleotide-level conservation quantified using 447 genomes from primates and mammals identified 328 missense changes and 89 SNCs in promoter-proximal regions occur at positions where the ancestral allele has high levels of conservation (phyloP score > 1.3 where both Denisovan genomes carry the derived allele in canonical transcripts) (UCSC Genome Browser).

Gene flow from Denisovans to modern humans represents an evolutionarily relevant source of variation, as introgressed alleles can alter gene regulation and protein function, thereby affecting cellular processes. We found that 32,691 and 61,131 SNCs are in regions with Denisovan introgression in non-Africans based on segments of at least 0.05cM identified with *admixfrog* and *hmmix*, respectively (**Supplementary Section 11**). Denisovan-derived SNCs could also have had a deleterious effect when introduced in modern humans and therefore purged from the gene pool. There are 3,898 SNCs in “deserts” of archaic introgression, which are unexpectedly long regions devoid of Neandertal and Denisovan ancestry in the genomes of non-Africans^146^.

#### Single nucleotide changes associated with traits

To identify SNC that may impact traits, we used a database of genetic variants associated with traits from genome-wide association studies in present-day humans (EMBL-EBI catalogue, accessed on October 12, 2024).  There are 41 positions where Denisovans have a SNC that are associated with variability of 74 traits (**Table S74**; **Table S76**). At 11 positions, the Denisovan-derived allele is associated with 16 traits.

A query of the Online Mendelian Inheritance in Man (OMIM) database revealed two positions where a single-nucleotide variant is implicated in a disease at the same positions where Denisovans carry a single-nucleotide change. While the Denisovan alleles are not reported to result in the diseases, their occurrence at the same position suggests they could influence the affected traits. The disease-associated alleles at both positions are observed at higher frequencies globally than the Denisovan-derived alleles (gnomAD, version 4), with conflicting reports in ClinVar regarding the pathogenicity of these variants.

        	One position (16:3820723) results in a missense substitution in *CREBBP* that is implicated in Rubinstein-Taybi syndrome 1, characterized by facial features including high-arched eyebrows, elongated face, prominent nose, high-arched palate, short broad thumbs, and broad halluces, as well behavioral traits including short attention span, dyslexia, dyscalculia, and reading difficulties (OMIM). The disease-associated allele results in a threonine to alanine substitution, while the Denisovan-derived allele results in a threonine to serine substitution. The allele is present in the *Denisova 3* genome, but no allele call is available for the Denisova 25 genome.

        	The other position (11:18332490) occurs in the intron of *HPS5* and activates an alternative splicing event, which is implicated in Hermansky-Pudlak Syndrome 5- characterized by oculocutaneous albinism, lack of platelet dense bodies, and prolonged bleeding (OMIM). The disease-associated allele at this position is cytosine, the ancestral state is thymine, and both Denisovan genomes have adenine.

#### Single nucleotide changes with combinations of annotations

We combined the annotations described above to pinpoint SNCs more likely to alter cellular functions. The SNCs have a derived allele in both Denisovan genomes, unless specifically noted that one of the genomes has no confident allele call. Analyses were restricted to the canonical transcript for each gene, as defined by MANE-Select using criteria including expression and coding region conservation (GENCODE, version 43)^151^. Although alternative isoforms provide a full repertoire of possible changes, SNCs in canonical transcripts are most likely to be biologically relevant. To describe the function of genes associated with SNCs, we utilized gene ontology terms (obtained from geneontology.org on October 23, 2024) and human phenotype ontology terms (obtained from hpo.jax.org on December 12, 2024). Gene expression information was obtained from The Human Protein Atlas (proteinatlas.org). In some cases where noted, predicted transcription factor binding sites were identified using the 2024 JASPAR Transcription Factor Binding Site Database and manually inspected in the UCSC Genome Browser to determine if the Denisovan-derived allele occurred at an important position in the motif position-weight matrix and likely to disrupt binding.

**Missense changes**

Of the 1,307 Denisovan-derived missense changes identified, 726 have an allele call in both Denisovan genomes in 646 genes. We conducted a gene ontology (GO) enrichment analysis with *GOfuncR*^150^ using these missense changes weighted by their PrimateAI-3D score to prioritize those more likely to have a functional impact. However, no GO terms were significantly enriched for genes with missense changes after correction for multiple testing using the Wilcoxon rank-sum test and a family-wise error rate derived from 1,000 random permutations.

**Genes with missense changes likely to be impactful based on PrimateAI-3D scores**

*DACH2* (X:85404043)

The Denisovan-derived missense change in *DACH2*, encoding Dachshund family transcription factor 2, results in a glycine to glutamic acid substitution at amino acid number 140 of 599. The change, which occurs in the DNA-binding domain of the protein (UniProt), has a pathogenicity score of 0.907. The ancestral allele is highly conserved in 447 mammalian genomes (p-value = 2.4E-08) and the Denisovan allele is not found among 807,162 human genomes (gnomAD, version 4). The Denisova 3 genome does not have an allele called at this position. DACH2 has RNA polymerase II-specific DNA-binding transcription factor activity (GO:0000981) and is involved in the development of primary female sexual characteristics (GO:0046545). It is expressed in the brain, epididymis, fallopian tube, and retina (The Human Protein Atlas).

*DENND1B* (1:197564430)

The Denisovan allele results in a serine to proline substitution at amino acid number 319 of 775, which has a pathogenicity score of 0.887. The ancestral allele is highly conserved in 447 mammalian genomes (p-value = 4.7E-08) and the Denisovan allele is not found among 807,162 human genomes (gnomAD, version 4). The *Denisova 25* genome shares this change with *Denisova 3*, but the genotype is in a region masked by our filters. The gene encodes DENN domain containing 1B and is located in clathrin-coated vesicles (GO:0030136), where it is involved in endocytosis (GO:0006897) and protein transport (GO:0015031). It enables small GTPase binding (GO:0031267), phosphatidylinositol phosphate binding (GO:1901981), and guanyl-nucleotide exchange factor activity (GO:0005085). In addition, this gene is involved in regulating the immune response (GO:0050776) by participating in the T cell receptor signaling pathway (GO:0050852) and regulating T-helper 2 cell cytokine production (GO:2000553).

*UNC93A* (6:167705033)

The Denisovan allele results in a phenylalanine to serine substitution at amino acid number 19 of 457. The ancestral allele is highly conserved in 447 mammalian genomes (p-value = 2.3E-07). The missense change occurs in the transmembrane domain of the protein and has a pathogenicity score of 0.884. The Denisovan allele is not found among 807,162 human genomes (gnomAD, version 4). The gene encodes Unc-93 homolog A, located in the plasma membrane (GO:0005886). UNC93A is most highly expressed in the skin, liver, intestine, and esophagus (The Human Protein Atlas).

*SLC49A3* (4:676668)

*SLC49A3*, encoding solute carrier family 49 member 3, has a Denisovan-derived missense change resulting in a glycine to glutamic acid substitution at amino acid number 388 of 560. The change, which occurs in the transmembrane domain of the protein, has a pathogenicity score of 0.841. The ancestral allele is highly conserved in 447 mammalian genomes (p-value = 1.47E-08). The Denisovan allele is found at a frequency of 0.0006274% among 807,162 human genomes (gnomAD, version 4). The gene is involved in transmembrane transport (GO:0055085) and is expressed across a variety of tissues.

*PLCE1* (10:96025626)

The missense change in *PLCE1*, encoding Phospholipase C epsilon 1, results in a leucine to phenylalanine substitution at amino acid number 1398 of 2302. The missense change has a pathogenicity score of 0.837 and is in the PI-PLC X-box catalytic domain of the protein, close to the active site which is 9 residues away (UniProt). The Denisovan-derived allele (no call in the *Denisova 3* genome) is not found among 807,162 human genomes (gnomAD, version 4) and the ancestral allele is highly conserved in 447 mammalian genomes (p-value = 4.09E-07). The gene has phosphatidylinositol phospholipase C activity (GO:0004435), guanyl-nucleotide exchange factor activity (GO:0005085), and can release sequestered calcium ions into the cytosol (GO:0051209). It is part of several intracellular signal transduction pathways (GO:0035556), including epidermal growth factor receptor signaling (GO:0007173), phospholipase C-activating G protein-coupled receptor signaling (GO:0007200), phosphatidylinositol-mediated signaling (GO:0048015), calcium-mediated signaling (GO:0019722), and Ras protein signal transduction (GO:0007265). It positively regulates lamellipodium assembly (GO:0010592), glomerulus development (GO:0032835), and is involved in the biosynthesis of diacylglycerol (GO:0006651). Mutations in PLCE1 can lead to the following phenotypes in humans: abnormal physiology of the kidney (HP:0012211), urinary system (HP:0011277), immune system (HP:0010978), respiratory system (HP:0002795), nervous system (HP:0012638), and digestive system (HP:0025032); abnormal morphology of the renal glomerular mesangium (HP:0001966), renal cortex (HP:0011035), renal corpuscle (HP:0031263), lung (HP:0002088), facial soft tissue (HP:0011799), and peritoneum (HP:0002585); fluid regulation (HP:0011032), low blood albumin level (HP:0003073) urine metabolite level (HP:0033354), foamy urine (HP:0031504), proteinuria (HP:0000093), stage 5 chronic kidney disease (HP:0003774), nephritis (HP:0000123), respiratory tract infection (HP:0011947), increased inflammatory response (HP:0012649), fever (HP:0001945), peritonitis (HP:0002586), abdominal pain (HP:0002027), serositis (HP:0045073), temperature regulation (HP:0004370), facial edema (HP:0000282), periorbital edema (HP:0100539), headache (HP:0002315), emotional state (HP:0100851), and irritability (HP:0000737).

*PIGG* (4:494258)

The Denisovan-derived missense change in PIGG, encoding phosphatidylinositol glycan anchor biosynthesis class G, results in a leucine to phenylalanine substitution at amino acid number 76 of 983. The change has a pathogenicity score of 0.835 and the ancestral allele is conserved in 447 mammalian genomes (p-value = 0.03). The Denisovan allele is found at a frequency of 0.001363% among 807,162 human genomes (gnomAD, version 4) and is reported to be implicated in intellectual disability, autosomal recessive 53, although with uncertain significance (ClinVar Variation ID 864305). PIGG is located in the endoplasmic reticulum membrane (GO:0005789), where it is involved in GPI anchor biosynthetic processes (GO:0006506) and enables CP2 mannose-ethanolamine phosphotransferase activity (GO:0051267). There are 440 human phenotype associations, including craniofacial morphology affecting the skull size (HP:0000240), head circumference (HP:0040195), high forehead (HP:0000348), high anterior hairline (HP:0009890), midface (HP:0000309), orbital region (HP:0000315), downward slanting of the opening between the eyelids (HP:0000494), eye folds (HP:0000286), increased distance between eyes (HP:0000316), iris coloboma (HP:0000612), megalocornea (HP:0000485), retina (HP:0000479), highly arched eyebrow (HP:0002553), posteriorly rotated ears (HP:0000358), middle ear (HP:0000370), preauricular pit (HP:0004467), wide nasal bridge (HP:0000431), short philtrum (HP:0000322), thin upper lip vermilion (HP:0000219), downturned corners of mouth (HP:0002714), oral cavity (HP:0000163), palate (HP:0000174), mandible (HP:0000277), decreased projection of lower jaw (HP:0000347), hypodontia (HP:0000668), taurodontia (HP:0000679), molar morphology (HP:0011071), and dental root (HP:0006486). Other skeletal morphology effects include thumb length (HP:0009778), finger phalanx (HP:0005918), long and slender fingers (HP:0001166), hyperextensibility of the finger joints (HP:0001187), limb bone morphology (HP:0002813), thorax (HP:0000765), rib fusion (HP:0000902), hunched back (HP:0002808), pubic bone (HP:0003172), flat foot (HP:0001763), and body height (HP:0000002). Associations with other parts of the body include morphology of the skin adnexa (HP:0011138), hair (HP:0001595), skeletal muscle (HP:0011805), internal female genitalia (HP:0000008), testis (HP:0000035), and penis (HP:0000036). The nervous system has several associations including microcephaly (HP:0000252), motor delay (HP:0001270), autistic behavior (HP:0000729), delayed speech and language development (HP:0000750), delayed ability to walk (HP:0031936), severe intellectual disability (HP:0010864), hearing impairment (HP:0000365), and chronic infections of the middle ear (HP:0000389).

*COPB1* (11:14501220)

*COPB1*, encoding COPI coat complex subunit beta 1, has a Denisovan-derived missense change in Denisova 3 (the Denisova 25 genome shares this change, but the genotype is in a region masked by our coverage filters) resulting in an alanine to glycine substitution at amino acid number 418 of 953 with a pathogenicity score of 0.824. The ancestral allele is highly conserved in 447 mammalian genomes (p-value = 7.5E-09). The Denisovan allele has a frequency of 0.02116% among 807,162 human genomes and is found in the homozygous state in two South Asians (gnomAD, version 4). The gene is involved in vesicle-mediated transport from the endoplasmic reticulum to the Golgi (GO:0006888) and within the Golgi (GO:0006891). It forms part of the COPI vesicle coat (GO:0030126) and can also be found in ficolin-1-rich granule membranes (GO:0101003). This gene has 135 phenotype associations in humans. It impacts skeletal morphology (HP:0011842), including skull size (HP:0000240), head circumference (HP:0040195), orbital region (HP:0000315), anterior eye segment (HP:0004328), upward slanting of the opening between the eyelids (HP:0000582), high, narrow palate (HP:0002705), oral cavity (HP:0000163), vertebral column curvature (HP:0010674), limb bones (HP:0002813), reduction in girth of the finger from proximal to distal (HP:0001182), and flat foot (HP:0001763). The gene also affects the musculature (HP:0003011), muscle physiology (HP:0011804), muscle tone (HP:0003808), body weight (HP:0004323), obesity (HP:0001513), skin elasticity (HP:0010647), epidermal thickening (HP:0011368), skin hyperpigmentation (HP:0000953), cafe-au-lait spots (HP:0000957), hair morphology (HP:0001595), hair quantity (HP:0011362), and hirsutism (HP:0001007). COPB1 also plays a role in the lower urinary tract (HP:0010936), bladder (HP:0000009), urinary incontinence (HP:0000020), forebrain morphology (HP:0100547), microcephaly (HP:0000252), upper motor neurons (HP:0002493), neurodevelopmental delay (HP:0012758), delayed speech and language development (HP:0000750), inability to walk (HP:0002540), and severe intellectual disability (HP:0010864).

*KIF14* (1:200573017)

The Denisovan allele results in an alanine to serine substitution at amino acid number 605 of 1648 of *KIF14*, which encodes kinesin family member 14. The change has a pathogenicity score of 0.82 and is in the kinesin motor domain of the protein (UniProt). The Denisovan allele has a frequency of 0.001301% among 807,162 human genomes (gnomAD, version 4) and is reported to be implicated in several conditions including Microcephaly 20 primary autosomal recessive, lethal fetal cerebrorenogenitourinary agenesis/hypoplasia syndrome, and KIF14-related disorder, although all with uncertain significance (ClinVar Variation ID 1033306). The ancestral allele is highly conserved in 447 mammalian genomes (p-value = 8.4E-09).

The gene regulates cellular processes including proliferation (GO:0008284), maturation (GO:1903429), growth (GO:0001558), adhesion (GO:0030155), migration (GO:0030334), microtubule-based movement (GO:0007018), establishment of protein localization (GO:0045184), Rap protein signal transduction (GO:0032487), and integrin activation (GO:0033624). It also regulates cell division (GO:0051301), particularly mitotic cell cycle G1/S transition (GO:2000045), G2/M transition (GO:0010389) and mitotic metaphase chromosome alignment (GO:0007080). In the brain, it is involved in cell proliferation in forebrain (GO:0021846), hippocampus development (GO:0021766), olfactory bulb development (GO:0021772), cerebellar Purkinje cell layer structural organization (GO:0021693), myelination (GO:0031641), and negative regulation of neuron apoptotic processes (GO:0043524).

*KIF14* has 226 human phenotype associations. The gene affects craniofacial morphology, including skull size (HP:0000240), skull bones (HP:0009116), head circumference (HP:0040195), sloping forehead (HP:0000340), orbital region (HP:0000315), ocular globe size (HP:0100887), eye fundus (HP:0001098), visual impairment (HP:0000505), eye movement (HP:0000549), optic nerve (HP:0000587), upward slanting of the opening between the eyelids (HP:0000582), low-set ears (HP:0000369), ear pinna (HP:0000377), wide nasal bridge (HP:0000431), nostrils (HP:0005288), thin upper lip vermilion (HP:0000219), oral cavity (HP:0000163), palate (HP:0000174), bifid uvula (HP:0000193), and decreased projection of lower jaw (HP:0000347). Other parts of the skeleton are also impacted, including the limbs (HP:0040064), tarsal bones (HP:0001850), elbow antecubital pterygium (HP:0009760), talus (HP:0008365), rocker bottom foot (HP:0001838), stature (HP:0004322), skeletal muscle (HP:0011805), joint mobility (HP:0011729), and tendon reflexes (HP:0001347). In the brain, the gene is associated with morphology of the cerebral cortex (HP:0002538), forebrain (HP:0100547), cerebellum (HP:0001317), corpus callosum (HP:0001273), olfactory bulb (HP:0002139), cerebral white matter morphology (HP:0002500), cortical gyration (HP:0002536), microcephaly (HP:0000252), lissencephaly (HP:0001339), ventriculomegaly (HP:0002119), neuronal migration (HP:0002269), neurodevelopmental delay (HP:0012758), motor delay (HP:0001270), severe intellectual disability (HP:0010864), poor speech (HP:0002465), increased volition (HP:5200263), short attention span (HP:0000736), and attention deficit hyperactivity disorder (HP:0007018). The gene also affects the reproductive system: female genitalia (HP:0010460), vaginal atresia (HP:0000148), uterus (HP:0008684), and amniotic fluid levels (HP:0001562); the genitourinary system: the ureter (HP:0000069), bladder (HP:0000014), and kidney (HP:0000104); and, skin morphology (HP:0011121).

*VWA8* (13:42306256)

The Denisovan-derived missense change in *VWA8* results in a serine to leucine substitution at amino acid number 821 of 1905. The change has a pathogenicity score of 0.819. The Denisovan allele has a frequency of 0.00111% among 807,162 human genomes (gnomAD, version 4) and is found in the homozygous state in one South Asian. The ancestral allele is highly conserved in 447 mammalian genomes (p-value = 7.9E-09). The gene encodes Von Willebrand factor A domain containing 8, which is located in mitochondria (GO:0005739) and peroxisomes (GO:0005777) where it has ATP hydrolysis activity (GO:0016887). Phenotype associations in humans are related to the eye (HP:0000478), including abnormal retinal morphology (HP:0000479), iris morphology (HP:0000525), retinal degeneration (HP:0000546), rod-cone dystrophy (HP:0000510), macular degeneration (HP:0000608), visual impairment (HP:0000505), lazy eye (HP:0000646), reduced visual acuity (HP:0007663), and poor night vision (HP:0000662).

*GPR158* (10:25701374)

The Denisovan allele results in a leucine to proline substitution at amino acid number 436 of 1215 of *GPR158*, which encodes G protein-coupled receptor 158. The change has a pathogenicity score of 0.801 and is in the transmembrane domain of the protein (UniProt). The Denisovan allele is not found in 807,162 human genomes (gnomAD, version 4) and the ancestral allele is highly conserved in 447 mammalian genomes (p-value = 4.7E-08). GPR158 regulates the G protein-coupled receptor signaling pathway (GO:0007186) and protein localization to the plasma membrane (GO:0072659). It is involved in cognition (GO:0050890) and brain development (GO:0007420), where it can be found in the presynaptic membrane (GO:0042734) as well as the postsynaptic density membrane (GO:0098839) regulating synapse organization (GO:0050807) and neurotransmitter secretion (GO:0001956). The gene is mostly expressed in the brain and retina (The Human Protein Atlas).

**Shared gene ontology germs with at least two genes**

Protein binding (GO:0005515): COPB1, DENND1B, KIF14, PLCE1, SLC49A3, UNC93A, VWA8

Membrane (GO:0016020): COPB1, KIF14, PIGG, SLC49A3, UNC93A

Plasma membrane (GO:0005886): COPB1, GPR158, KIF14, PLCE1, UNC93A

Cytosol (GO:0005829): COPB1, DENND1B, KIF14, PLCE1

Nucleus (GO:0005634): DACH2, GPR158, KIF14

ATP binding (GO:0005524): KIF14, VWA8

ATP hydrolysis activity (GO:0016887): KIF14, VWA8

Cytoplasm (GO:0005737): KIF14, VWA8

Endoplasmic reticulum membrane (GO:0005789): COPB1, PIGG

Golgi membrane (GO:0000139): COPB1, PLCE1

G protein-coupled receptor signaling pathway (GO:0007186): GPR158, PLCE1

Guanyl-nucleotide exchange factor activity (GO:0005085): DENND1B, PLCE1

Intracellular membrane-bounded organelle (GO:0043231): COPB1, DENND1B

Small GTPase binding (GO:0031267): DENND1B, PLCE1

**Shared human phenotype ontology terms with at least four genes**

All (HP:0000001): COPB1, KIF14, PIGG, PLCE1, VWA8

Clinical course (HP:0031797): COPB1, KIF14, PIGG, PLCE1, VWA8

Clinical modifier (HP:0012823)	COPB1, KIF14, PIGG, PLCE1, VWA8

Mendelian inheritance (HP:0034345): COPB1, KIF14, PIGG, PLCE1, VWA8

Mode of inheritance (HP:0000005): COPB1, KIF14, PIGG, PLCE1, VWA8

Onset (HP:0003674): COPB1, KIF14, PIGG, PLCE1, VWA8

Pediatric onset (HP:0410280): COPB1, KIF14, PIGG, PLCE1, VWA8

Phenotypic abnormality (HP:0000118): COPB1, KIF14, PIGG, PLCE1, VWA8

Abnormal eye morphology (HP:0012372): COPB1, KIF14, PIGG, VWA8

Abnormality of head or neck (HP:0000152): COPB1, KIF14, PIGG, PLCE1

Abnormality of mental function (HP:0011446): COPB1, KIF14, PIGG, PLCE1

Abnormality of the eye (HP:0000478): COPB1, KIF14, PIGG, VWA8

Abnormality of the face (HP:0000271): COPB1, KIF14, PIGG, PLCE1

Abnormality of the genitourinary system (HP:0000119): COPB1, KIF14, PIGG, PLCE1

Abnormality of the head (HP:0000234): COPB1, KIF14, PIGG, PLCE1

Abnormality of the nervous system (HP:0000707)	COPB1, KIF14, PIGG, PLCE1

Abnormality of the urinary system (HP:0000079)	COPB1, KIF14, PIGG, PLCE1

Abnormal nervous system physiology (HP:0012638): COPB1, KIF14, PIGG, PLCE1

Autosomal recessive inheritance (HP:0000007): COPB1, KIF14, PIGG, PLCE1

**Missense changes in enzymes with catalytic activity (GO:0003824)**

*DBH* (9:136521734)

The Denisovan allele results in an aspartic acid to glutamic acid substitution at amino acid number 508 of 617 of *DBH*. This gene encodes dopamine beta-hydroxylase, an enzyme that converts dopamine into norepinephrine that is mostly expressed in the adrenal gland and liver (The Human Protein Atlas). It is involved in the positive regulation of vasoconstriction (GO:0045907), cold-induced thermogenesis (GO:0120162), homoiothermy (GO:0042309), vascular associated smooth muscle cell proliferation (GO:1904705), leukocyte mediated immunity (GO:0002443), glucose homeostasis (GO:0042593), chemical synaptic transmission (GO:0007268), memory (GO:0007613), locomotory behavior (GO:0007626), visual learning (GO:0008542), fear response (GO:0042596), maternal behavior (GO:0042711), response to amphetamine (GO:0001975), behavioral response to ethanol (GO:0048149), and response to pain (GO:0048265).

*HSD17B1* (17:40706526)

The Denisovan allele results in an arginine to serine substitution at amino acid number 215 of 328 of *HSD17B1*. This gene encodes 17-beta-hydroxysteroid dehydrogenase type 1, an enzyme involved in the production and metabolism of steroid hormones like estrogen and androgens that is mostly expressed in the placenta (The Human Protein Atlas). It is involved in biosynthesis of steroids (GO:0006694), estrogen (GO:0006703), and testosterone (GO:0061370), as well as the development of adipose tissue (GO:0060612), skeletal muscle tissue (GO:0007519), and bone (GO:0060348).

**Genes with start and stop codon changes**

*C1GALT1C1L* (2:43903301)

The Denisovan-derived allele introduces a stop codon at amino acid number 54 of 315. Among 807,162 human genomes, the allele is found in three individuals (gnomAD, version 4). The gene encodes *C1GALT1* specific chaperone 1 like, which enables glycoprotein-N-acetylgalactosamine 3-beta-galactosyltransferase activity (GO:0016263).

*CCDC7* (10:33103363)

The Denisovan-derived allele introduces a stop codon at amino acid number 1055 of 1385. The allele has a frequency of 0.0076% among 807,162 human genomes (gnomAD, version 4). The gene encodes coiled-coil domain containing 7, which enables protein binding (GO:0005515).

*PON3* (7:94992090)

The Denisovan-derived allele introduces a stop codon at amino acid number 253 of 354. The allele is not found among 807,162 human genomes. The ancestral allele is highly conserved among 447 mammalian genomes (p-value = 8.1E-09). The gene encodes Paraoxonase 3, which is located in the endoplasmic reticulum membrane (GO:0005789) and extracellular exosome (GO:0070062), and enables activity of arylesterase (GO:0004064), aryldialkylphosphatase (GO:0004063), and acyl-L-homoserine-lactone lactonohydrolase (GO:0102007). It is involved in carboxylic acid catabolic processes (GO:0046395), lactone catabolic processes (GO:1901335), renal sodium ion transport (GO:0003096), response to toxic substances (GO:0009636), regulation of the cellular response to drugs (GO:2001038), negative regulation of superoxide anion generation (GO:0032929), and establishment of localization in cell (GO:0051649). PON3 has many human phenotype associations, including depression (HP:0000716), anxiety (HP:0000739), frontotemporal dementia (HP:0002145), neurodegeneration (HP:0002180), language impairment (HP:0002463), motor neuron atrophy (HP:0007373), fatigable weakness of bulbar muscles (HP:0030192), fatigable weakness of swallowing muscles (HP:0030195), fatigable weakness of respiratory muscles (HP:0030196), upper limb muscle weakness (HP:0003484), lower limb muscle weakness (HP:0007340), foot dorsiflexor weakness (HP:0009027), steppage gait (HP:0003376), muscle spasm (HP:0003394), vocal chord spasm (HP:0025425), tongue fasciculations (HP:0001308), sensation of breathlessness in the recumbent position (HP:0012764), jaw hyperreflexia (HP:0033683), and pain (HP:0012531).

*PRXL2B* (1:2520477)

The Denisovan-derived allele introduces a stop codon at amino acid number 193 of 198. The allele is in the genome of a South Asian individual from a collection of 807,162 human genomes (gnomAD, version 4). The ancestral allele is conserved among 447 mammalian genomes (p-value = 0.043). This gene encodes Peroxiredoxin like 2B, which is involved in prostaglandin biosynthetic processes (GO:0001516) by enabling prostaglandin F synthase activity (GO:0047017) and oxidoreductase activity (GO:0016616). It is located in the cytosol (GO:0005829), mitochondria (GO:0005739), endoplasmic reticulum (GO:0005783), extracellular exosome (GO:0070062), and myelin sheath (GO:0043209).

*ZAN* (7:100363121 and 7:100369478)

The Denisovan-derived alleles introduce stop codons at amino acid numbers 1472 and 1754 of 2812. The first position occurs in the Trypsin Inhibitor-Like 2 domain of the protein. The alleles are found in the genome of three individuals from a collection of 807,162 human genomes (gnomAD, version 4). The Denisova 25 genome does not have an allele call at either position. This gene encodes zonadhesin, which is involved in the binding of sperm to the zona pellucida (GO:0007339) and cell-to-cell adhesion (GO:0098609). It is active in the extracellular space (GO:0005615), extracellular matrix (GO:0031012), and plasma membrane (GO:0005886).

*CEP70* (3:138291769)

The Denisovan-derived allele results in a start codon loss at amino acid number 1 of 597 in the Disordered domain of the protein. The allele has a frequency of 0.0013% among 807,162 human genomes (gnomAD, version 4). The gene encodes centrosomal protein 70. It is involved in cilium assembly (GO:0060271) and the regulation of microtubule cytoskeleton organization (GO:0070507), where it is active in the microtubule organizing center (GO:0005815). It is located in the cytosol (GO:0005829) and centrosome (GO:0005813).

**Promoter-proximal changes**

We identified 1,126 single nucleotide changes where both Denisovans carry the derived allele in promoter-proximal regions of 1,052 genes. The SNCs were ranked by the nucleotide-level conservation scores from the 447-way mammalian genome comparison to identify those more likely to have a functional effect (**Figure S126**). To determine if genes with SNCs are involved in shared biological processes, molecular functions, or cellular components, we performed a gene ontology enrichment analysis with GOfuncR and the gene ontology database described above^150^. A Wilcoxon rank-sum test was used to incorporate the conservation score and likelihood of a SNC having a functional effect. The highest score was used for genes with multiple SNC as there could only be one entry per gene. One thousand random permutations were performed to obtain the family-wise error rate (FWER) to correct for multiple testing (**Table S77**). Genes with SNCs at highly conserved positions of the genome were over-represented in the following terms: central nervous system development (GO:0007417) (FWER < 0.001), nervous system development (GO:0007399) (FWER < 0.001), brain development (GO:0007420) (FWER = 0.005), head development (GO:0060322) (FWER = 0.005), and neuron projection membrane (GO:0032589) (FWER = 0.053).

**Changes at conserved positions and predicted to alter transcription factor binding**

*FOXP2* (7:114054475)

A SNC is located 395 base pairs upstream the transcription start site of *FOXP2*, where the ancestral allele is highly conserved in mammals (p-value = 8E-9) and the Denisovan-derived allele would likely disrupt binding of ISL1, a transcription factor predicted to bind strongly to the ancestral allele (JASPAR Core 2024 predicted transcription factor binding sites provided by the UCSC Genome Browser). FOXP2 is expressed in nearly every tissue, while *ISL1* is most highly expressed in the retina, seminal vesicle, vagina, with lower expression in the basal ganglia and hypothalamus of the brain, as well as the stomach, intestines, liver, pancreas, and thymus (The Human Protein Atlas). *FOXP2* encodes the transcription factor forkhead box protein P2. It is involved in the development of the caudate nucleus (GO:0021757), putamen (GO:0021758), and cerebral cortex (GO:0021987). The gene is associated with craniofacial phenotypes in humans, including asymmetry of the posterior skull (HP:0001357), macrocephaly (HP:0000256), horizontal eyebrow (HP:0011228), eye folds (HP:0000286), increased distance between the eyes (HP:0000316), low-set ears (HP:0000369), overfolded ears (HP:0000396), small semicircular canal (HP:0011382), enlarged cochlear aqueduct (HP:0011388), prominent nose (HP:0000448), wide nasal ridge (HP:0012811), increased distance between nasal base and upper lip (HP:0000343), wide mouth (HP:0000154), high, narrow palate (HP:0002705), submucous cleft hard palate (HP:0000176), decreased size of the upper jaw (HP:0000327), prominent fingertip pads (HP:0001212), curvature of the index finger (HP:0040022), and scoliosis (HP:0002650). Other associations include skeletal muscle atrophy (HP:0003202), wry neck (HP:0000473), low muscle tone in the trunk (HP:0008936), poor motor coordination (HP:0007010, HP:0007015), oral-motor apraxia (HP:0007301), speech apraxia (HP:0011098), delayed speech and language development (HP:0000750), incomprehensible speech (HP:0002546), autistic behavior (HP:0000729), short attention span (HP:0000736), hyperactivity (HP:0000752), intellectual disability (HP:0001249), severe global developmental delay (HP:0011344), abnormal basal ganglia morphology (HP:0002134), abnormal caudate nucleus morphology (HP:0002339), hypoplasia of the olfactory bulb (HP:0040326), childhood onset sensorineural hearing impairment (HP:0011474), recurrent ear infections (HP:0410018), recurrent respiratory infections (HP:0002205), asthma (HP:0002099), atrial septal defect (HP:0001631),abnormality of von Willebrand factor (HP:0012146), gastroesophageal reflux (HP:0002020), intrauterine growth retardation (HP:0001511), increased urinary galactose level (HP:0012023), and involuntary loss of urine during sleep (HP:0010677).

*CLCF1* (11:67141996)

The promoter-proximal region of *CLCF1* has a SNC located 928 base pairs upstream the transcription start site where the ancestral allele is highly conserved in mammals (2.2E-8). The Denisovan-derived allele would likely disrupt binding of transcription factors AP1, HMGN3, MAF, NFE2, NFE2L1, NFE2L2, NR3C1, SMARC, TCF7L2, and TFAP2, which are predicted to bind to the ancestral allele (FunSeq2[1] ). The gene encodes cardiotrophin like cytokine factor 1, which is involved in the JAK-STAT cell surface receptor signaling pathway (GO:0007259), cytokine-mediated signaling pathway (GO:0019221), positive regulation of B cell proliferation (GO:0030890) and differentiation (GO:0030183), immunoglobulin production (GO:0002639), and isotype switching to IgE isotypes (GO:0048295). In the brain, it is part of the CRLF/CLCF1 complex (GO:0097058) involved in ciliary neurotrophic factor receptor binding (GO:0005127) that positively regulates astrocyte differentiation (GO:0048711) and negatively regulates neuron apoptotic processes (GO:0043524). Phenotypic associations in humans include large face (HP:0100729), full cheeks (HP:0000293), protruding ears (HP:0000411), wide nose (HP:0000445), upturned nasal tip (HP:0000463), increased distance between nasal base and upper lip (HP:0000343), narrow mouth (HP:0000160), high palate (HP:0000218), small lower jaw (HP:0000347), curving of the finger (HP:0030084), flexion of the finger (HP:0100490), webbed 2nd and 3rd toes (HP:0004691), joint mobility limitation (HP:0001376), flexed joint that cannot be straightened (HP:0001371), thoracolumbar scoliosis (HP:0002944), hunched back (HP:0002808), excessive inward curvature of lower spine (HP:0002938), outward turned elbows (HP:0002967), limited elbow extension (HP:0001377), high level of muscle tone (HP:0001276), sensorimotor neuropathy (HP:0007141), weakness of facial musculature (HP:0030319), seizure (HP:0001250), sudden cardiac death (HP:0001645), respiratory insufficiency (HP:0002093), cognitive impairment (HP:0100543), cold-induced sweating (HP:0025278), and malignant hyperthermia (HP:0002047).

*HOXA3* (7:27192291)

A SNC is located 84 base pairs upstream the transcription start site of HOXA3, where the ancestral allele is highly conserved in mammals (p-value = 3.1E-8) and the Denisovan-derived allele would likely disrupt binding of SOX12, a transcription factor predicted to bind only to the ancestral allele (JASPAR Core 2024 predicted transcription factor binding sites provided by the UCSC Genome Browser). The gene encodes homeobox A3, a transcription factor (GO:0000981) active in the nucleus (GO:0005634) that is involved in anterior posterior pattern specification (GO:0009952), embryonic skeletal system morphogenesis (GO:0048704), animal organ formation (GO:0048645), specification of animal organ position (GO:0010159), positive regulation of stem cell proliferation (GO:2000648), thyroid gland development (GO:0030878), parathyroid gland development (GO:0060017), thymus development (GO:0048538), cartilage development (GO:0051216), glossopharyngeal nerve morphogenesis (GO:0021615), angiogenesis (GO:0001525), and blood vessel remodeling (GO:0001974).

*PAX2* (10:102505033)

The promoter-proximal region of *PAX2* has a SNC located 306 base pairs upstream the transcription start site where the ancestral allele is highly conserved in mammals (p-value = 6.2E-8). The Denisovan-derived allele would likely disrupt binding of SIX5, a transcription factor predicted to bind only to the ancestral allele (*FunSeq2*). PAX2 encodes paired box 2, a transcription factor (GO:0000981) mostly expressed in the kidney and epididymis (The Human Protein Atlas) that is involved in: stem cell differentiation (GO:0048863), epithelial cell proliferation (GO:0050679), mesenchymal to epithelial transition (GO:0060231), mesodermal cell fate specification (GO:0007501), ureteric bud morphogenesis (GO:0001658), ureter maturation (GO:0035799), nephric duct formation (GO:0072179), metanephric distal convoluted tubule development (GO:0072221), metanephric glomerulus development (GO:0072300), metanephros morphogenesis (GO:0003337), mesonephros development (GO:0001823), sensory organ development (GO:0007423), optic nerve formation (GO:2000597), optic chiasma development (GO:0061360), optic cup morphogenesis involved in camera-type eye development (GO:0002072), retinal pigment epithelium development (GO:0003406), visual perception (GO:0007601), inner ear morphogenesis (GO:0042472), cochlea morphogenesis (GO:0090103), vestibulocochlear nerve formation (GO:0021650), neural tube closure (GO:0001843), brain morphogenesis (GO:0048854), axonogenesis (GO:0007409), glial cell differentiation (GO:0010001), response to nutrient levels (GO:0031667), cellular response to retinoic acid (GO:0071300), cellular response to glucose stimulus (GO:0071333), cellular response to epidermal growth factor stimulus (GO:0071364), and negative regulation of cysteine-type endopeptidase activity involved in apoptotic process (GO:0043154).

*NR2F1* (5:92918797)

A SNC is located 131 base pairs upstream the transcription start site of *NR2F1*, where the ancestral allele is highly conserved in mammals (p-value = 8.7E-8) and the Denisovan-derived allele would likely create a binding site for the transcription factors MYOG and TFCP2, which are predicted to bind only to the derived allele (*FunSeq2*). While *NR2F1* and *TFCP2* are expressed in nearly every tissue, *MYOG* is expressed in skeletal muscle and the tongue (The Human Protein Atlas). *NR2F1* encodes nuclear receptor subfamily 2 group F member 1, a transcription factor (GO:0001227) involved in intracellular receptor signaling (GO:0030522), cell differentiation (GO:0030154), nervous system development (GO:0007399), negative regulation of neuron projection development (GO:0010977), and enables retinoic acid-responsive element binding (GO:0044323). Human phenotypes associated with this gene include abnormal facial shape (HP:0001999), protruding ear (HP:0000411), uplifted earlobe (HP:0009909), outer ear rim Darwin tubercle (HP:0011261), prominent ear antihelix (HP:0000395), bulging cornea (HP:0000563), prominent eye folds (HP:0000286), upward slanting of the opening between the eyelids (HP:0000582), prominent nasal bridge (HP:0000426), short nasal bridge (HP:0003194), upturned nasal tip (HP:0000463), wide mouth (HP:0000154), high palate (HP:0000218), pointed chin (HP:0000307), receding chin/lower jaw (HP:0000278), reduction in girth of the finger from proximal to distal (HP:0001182), prominent fingertip pads (HP:0001212), short stature (HP:0004322), slow-growing hair (HP:0002217), delayed skeletal maturation (HP:0002750), low muscle tone (HP:0001252), muscle spasticity (HP:0001257), hearing impairment (HP:0000365), underdeveloped optic nerves (HP:0000609), optic-nerve degeneration (HP:0000648), underdeveloped optic disc (HP:0007766), pale optic discs (HP:0000543), visual field defect (HP:0001123), reduced visual acuity (HP:0007663), involuntary rhythmic eye movements (HP:0000639), lazy eye (HP:0000646), underdevelopment of the corpus callosum (HP:0002079), abnormal hippocampus morphology (HP:0025100), delayed myelination (HP:0012448), global developmental delay (HP:0001263), intellectual disability (HP:0001249), seizures (HP:0001250), autism (HP:0000717), absent speech (HP:0001344), repetitive compulsive behavior (HP:0008762), and attention deficit hyperactivity disorder (HP:0007018).

RHOA and *TCTA* (3:49449532)

A SNC is located in the overlapping promoter-proximal regions of *RHOA* and *TCTA*, which are are transcribed in opposite directions on different strands. The SNC is 124 base pairs upstream the transcription start site of RHOA and 324 base pairs upstream that of *TCTA*. The ancestral allele is highly conserved in mammals (p-value = 5.9E-6) and the Denisovan-derived allele is predicted to create a binding site for the transcription factor SOX17 on the strand that *RHOA* is transcribed from (*FunSeq2*). *RHOA* encodes Ras homolog family member A, which is involved in many processes: cell migration (GO:0016477), cell-substrate adhesion (GO:0010812), cell size (GO:0045792), epithelial cell polarity (GO:0045198), regulation of cell size (GO:0045792), negative chemotaxis (GO:0050919), positive regulation of cytokinesis (GO:0032467), mitotic cleavage furrow formation (GO:1903673), mitotic spindle assembly (GO:0090307), actin cytoskeleton organization (GO:0030036), cytoplasmic microtubule organization (GO:0031122), apical junction assembly (GO:0043297), stress fiber assembly (GO:0043149), focal adhesion assembly (GO:0051893), podosome assembly (GO:0071803), oxidative phosphorylation (GO:0090324), positive regulation of protein serine threonine kinase activity (GO:0071902), apolipoprotein A-I-mediated signaling pathway (GO:0038027), canonical NF-kappaB signal transduction (GO:0043123), Wnt signaling pathway, planar cell polarity pathway (GO:0060071), negative regulation of intracellular steroid hormone receptor signaling pathway (GO:0033144), semaphorin-plexin signaling pathway (GO:0071526), positive regulation of lipase activity (GO:0060193), cellular response to chemokine (GO:1990869), cellular response to cytokine stimulus (GO:0071345), cellular response to lipopolysaccharide (GO:0071222), alpha-beta T cell lineage commitment (GO:0002363), positive regulation of leukocyte adhesion to vascular endothelial cell (GO:1904996), positive regulation of T cell migration (GO:2000406), angiotensin-mediated vasoconstriction involved in regulation of systemic arterial blood pressure (GO:0001998), wound healing, spreading of cells (GO:0044319), aortic valve formation (GO:0003189), kidney development (GO:0001822), skeletal muscle tissue development (GO:0007519), skeletal muscle satellite cell migration (GO:1902766), positive regulation of vascular associated smooth muscle contraction (GO:1904695), forebrain radial glial cell differentiation (GO:0021861), cerebral cortex cell migration (GO:0021795), regulation of neuron projection development (GO:0010975), neuron migration (GO:0001764), substantia nigra development (GO:0021762), negative regulation of motor neuron apoptotic process (GO:2000672), regulation of modification of postsynaptic actin cytoskeleton (GO:1905274), regulation of osteoblast proliferation (GO:0033688), ossification involved in bone maturation (GO:0043931), odontogenesis (GO:0042476), and bone trabecula morphogenesis (GO:0061430).

*RHOA* phenotypic associations in humans include facial asymmetry (HP:0000324), hemifacial hypoplasia (HP:0011332), flat cheekbone (HP:0000272), unequal size of opening between the eyelids (HP:0007946), displaced pupil (HP:0009918), wide nasal bridge (HP:0000431), thick nasal alae (HP:0009928), short fingers or toes (HP:0001156), broad hallux (HP:0010055), widened gap first and second toe (HP:0001852), webbed 3rd-4th toes (HP:0009779), curvature of digit (HP:0030084), oligodontia (HP:0000677), persistence of primary teeth (HP:0006335), decreased width of tooth (HP:0000691), conical tooth (HP:0000698), enamel hypoplasia (HP:0006297), bilateral crossbite (HP:0000689), scarring alopecia of scalp (HP:0004552), optic atrophy (HP:0000648), decreased size of eyeball (HP:0000568), reduced visual acuity (HP:0007663), astigmatism (HP:0000483), eye squint (HP:0000486), glaucoma (HP:0000501), cataract (HP:0000518), near sightedness (HP:0000545), involuntary rhythmic eye movements (HP:0000639), global developmental delay (HP:0001263), and intellectual disability (HP:0001249).

**Multiple single nucleotide changes in promoter-proximal regions**

*FSIP2* (2:186602610, 2:186602759, 2:186602940, 2:186603264)

There are four SNC in the promoter-proximal region of FSIP2. This gene also has four missense changes (2:186653641, 2:186657199, 2:186667097, 2:186669975). FSIP2 encodes fibrous sheath interacting protein 2, which is involved in sperm axoneme assembly (GO:0007288), flagellated sperm motility (GO:0030317), protein localization to cilium (GO:0061512). Human phenotype associations include reduced sperm motility (HP:0012207), absent sperm flagella (HP:0032558), short sperm flagella (HP:0032559), coiled sperm flagella (HP:0032560), irregularly shaped sperm tail (HP:0033393), absent sperm axoneme central pair complex (HP:0033525), and male infertility (HP:0003251) with young adult onset (HP:0011462).

There are nine genes with three SNC in the promoter-proximal region: *C10orf90*, *CCDC180*, *CYTL1*, *MINAR2*, *OR2B6*, *RPL26*, *SIPA1L1*, *SMIM42*, *TMEM258*. There are 73 genes with two SNC in the promoter-proximal region.

**Changes associated with traits in present-day people**

There are seven single nucleotide changes where the derived allele is present in both Denisovan genomes and associated with a trait as reported in the NHGRI-EBI catalog of human genome-wide association studies.

rs17658229 (5:172191052)

A Denisovan-derived allele located in an intergenic region is associated with two traits. One is circulating amounts of C-reactive protein, a biomarker of chronic low-grade inflammation (beta coefficient = 0.056 unit increase [95% confidence interval: 0.036-0.076], p-value = 6E-9). The other is the ratio of the optic cup to optic disk size, where a higher ratio relates to optic nerve degeneration and is a key factor in the clinical assessment of glaucoma- the most common cause of irreversible blindness^174^. The allele with the largest absolute effect size is the Denisovan-derived one, which decreases the ratio (beta coefficient = 0.02 unit decrease [95% CI: 0.012-0.028], p-value = 8E-9)^174^.

The Denisovan-derived allele is associated with decreased expression of the long non-coding RNA *ENSG00000253736* in blood (normalized effect size = -0.39, p-value = 2.8E-8), tibial artery (normalized effect size = -0.69, p-value = 1.7E-7), spleen (normalized effect size = -0.77, p-value = 2.6E-7), EBV-transformed lymphocytes (normalized effect size = -0.87, p-value = 4.7E-7) and subcutaneous adipose (normalized effect size = -0.62, p-value = 7E-6).

It is also associated with alternative splicing of *DUSP1* in subcutaneous adipose (normalized effect size = ±1.2, p-value = 6.3E-16), tibial artery (normalized effect size = ±1.2, p-value = 2.8E-14), sun exposed skin (normalized effect size = ±0.92, p-value = 2.4E-7), and tibial nerve (normalized effect size = ±0.77, p-value = 2E-6). *DUSP1* encodes dual specificity phosphatase 1, which is involved in MAP kinase tyrosine serine threonine phosphatase activity (GO:0017017), negative regulation of the p38MAPK cascade (GO:1903753), negative regulation of cell population proliferation (GO:0008285), negative regulation of cell adhesion (GO:0007162), cellular response to chemokines (GO:1990869), negative regulation of monocyte chemotaxis (GO:0090027), regulation of mitotic cell cycle spindle assembly checkpoint (GO:0090266), and negative regulation of meiotic cell cycle (GO:0051447).

The allele is in a region with histone modifications indicative of enhancers in nine tissues: blood, cervix, fat, GI, heart, lung, muscle, spleen, thymus^153^.

rs45603544 (11:113818507)

A Denisovan-derived allele located in the 3' UTR of *HTR3B* is associated with a decrease in the bacterial taxa Rhodospirillales in the gut (beta coefficient = 0.53 unit decrease [95% confidence interval: 0.31-0.75], p-value = 2E-6). It overlaps a region with histone modifications indicative of enhancers in the fetal thymus and blood cells, including B cells, T cells, natural killer cells, monocytes, and neutrophils^153^. The Denisovan allele is also associated with increased expression of HTR3B in the frontal cortex of the brain (normalized effect size = 0.59, p-value = 7E-8).

rs2237881 (11:2788916)

A Denisovan-derived allele is associated with increased height (beta coefficient = 0.0228 unit increase [95% confidence interval: 0.017-0.028], p-value = 6E-15). It falls in the most constrained percentile of the genome (top 1%, Gnocchi score = 4.63) within the intron of KCNQ1 that is also a tissue invariant open chromatin region^152^. The nearest transcription start sites are those of *CDKN1C*, located 118089 base pairs downstream, and *KCNQ1*, located 322678 base pairs upstream.

rs146948889 (3:48477252)

A Denisovan-derived allele is associated with decreased total cholesterol levels (beta coefficient = 0.0442 unit decrease [95% confidence interval: 0.033-0.055], p-value = 3E-11). The derived allele would likely disrupt binding of the transcription factors SP1 and GABPA, which are predicted to bind strongly to the ancestral allele (*FunSeq2^171^*). Histone modifications indicate that the region is an enhancer in 24 different tissues^153^. The SNC is located in the most constrained percentile of the genome (top 1%, Gnocchi score = 4.2) within the intron of *CCDC51*. The nearest transcription start sites are those of *CCDC51*, located 4277 base pairs downstream, and *PLXNB1*, located 5757 base pairs upstream.

rs372272284 (2:46584859)

A Denisovan-derived allele is associated with decreased oxygenated hemoglobin levels (beta coefficient = 0.386 g/dL decrease, p-value = 6E-9)^175^. It occurs in the intron of *EPAS1*, whose transcription start site is nearest to the SNC and located 60,313 base pairs upstream. The region has histone modifications suggesting it is an enhancer in primary T helper naive cells from peripheral blood and several brain regions^153^.

**"Desert" regions devoid of Denisovan and Neandertal introgression**

**Missense changes**

The genes *C1orf162*, *CPED1*, and *PENK* each have one missense change, while *LRIG2* has three missense changes.

*LRIG2* (1:113616077, 1:113642867, 1:113666514)

There are three Denisovan-derived missense changes in *LRIG2* resulting in the following substitutions: arginine to glycine at amino acid number 17 of 1065, located in the signal peptide that targets it to the endoplasmic reticulum; lysine to glutamic acid at amino acid number 403, located in the extracellular domain; and, valine to methionine at amino acid number 997, located in the cytoplasmic domain (UniProt). The ancestral allele of the SNC resulting in a missense change at amino acid number 403 is highly conserved in 447 mammalian genomes (2.65E-5). *LRIG2* encodes Leucine-rich repeats and immunoglobulin-like domains protein 2, which is involved in the positive regulation of protein localization to the cell surface (GO:0034394), signaling receptor binding (GO:0005102), negative regulation of membrane protein ectodomain proteolysis (GO:0051045), regulation of platelet-derived growth factor receptor signaling pathway (GO:0010640), nerve innervation (GO:0060384), regulation of neuron migration (GO:2001222), negative regulation of axon regeneration (GO:0048681), and the sensory perception of sound (GO:0007605). Phenotype associations in humans include extreme thirst (HP:0001959), spastic/hyperactive bladder (HP:0005340), bladder trabeculation (HP:0032465), dilatation of the bladder (HP:0000021), retrograde movement of urine from the bladder into ureters or kidneys (HP:0000076), recurrent urinary tract infections (HP:0000010), obstruction of the flow of urine through the urethra (HP:0000796), overactive bladder (HP:0000012), lack of the ability to control the urinary bladder leading to involuntary urination (HP:0000805), bowel incontinence (HP:0002607), constipation (HP:0002019), severe distention of the kidney (HP:0000126), renal failure (HP:0000083), undescended testis (HP:0000028), high blood pressure (HP:0000822), and facial grimacing (HP:0000273).

*PENK* (8:57358494)

The Denisovan-derived missense change in *PENK*, encoding proenkephalin, results in a leucine to phenylalanine substitution at amino acid number 7 of 267. The change occurs in signal peptide that targets it to the endoplasmic reticulum (UniProt). *PENK* is mostly expressed in the brain, especially the basal ganglia, as well as the adrenal gland and testis (The Human Protein Atlas). It is involved in G protein-coupled opioid receptor signaling pathway (GO:0038003), cellular response to transforming growth factor beta stimulus (GO:0071560), cellular response to cAMP (GO:0071320), cellular response to virus (GO:0098586), response to lipopolysaccharide (GO:0032496), response to hypoxia (GO:0001666), cellular response to oxidative stress (GO:0034599), response to calcium ion (GO:0051592), response to vitamin D (GO:0071305), response to estradiol (GO:0032355), response to epinephrine (GO:0071871), response to ethanol (GO:0045471), response to nicotine (GO:0035094), response to toxic substance (GO:0009636), osteoblast differentiation (GO:0001649), synaptic signaling via neuropeptide (GO:0099538), chemical synaptic transmission (GO:0007268), transmission of nerve impulse (GO:0019226), regulation of long-term synaptic depression (GO:1900452), glial cell proliferation (GO:0014009), sensory perception (GO:0007600), sensory perception of pain (GO:0019233), locomotory exploration behavior (GO:0035641), response to immobilization stress (GO:0035902), startle response (GO:0001964), aggressive behavior (GO:0002118), and positive regulation of behavioral fear response (GO:2000987).

**Untranslated regions (UTR)**

The 5’ UTR of *PHTF1* and *GPR37* each have one SNC. The 3’ UTR of *VAV3*, *SLC25A24*, *CELSR2*, *PSMA5*, *GNAI3*, *WNT2B*, *BCL2L15*, *OLFML3*, *ROBO2*, *CADM2*, *POT1*, *RAB2A*, *CHD7*, *ASPH*, and *GGH* all have one SNC, while the 3’ UTR of *NKAIN3* has two SNC, and the 3’ UTR of *BPNT2* has three SNC.

**Promoter-proximal regions**

The following genes have a SNC in the promoter-proximal region: *RBM15*, *PROK1*, *DENND2D*, *PIFO*, *INKA2*, *RHOC*, *TAFA3*, *ROBO2*, *GBE1*, *FOXP2*, *ST7*, *ING3*, and *SOX17*.

**Enhancer region**

VISTA enhancer hs1995 (3:80598127, 3:80603261, 3:80603857)

There are three SNC in an 8,058 base pair region with enhancer activity in the facial mesenchyme. The human enhancer was discovered using a gene reporter assay performed in developing mice embryos (VISTA enhancer hs1995). The first SNC occurs at a position where the ancestral allele is highly conserved in vertebrate genomes (p-value = 2.57E-3). The Denisovan-derived allele at the third SNC would likely alter binding of the transcription factors MAFF, MAFB, and NRL, which are all predicted to bind only to the ancestral allele (JASPAR Core 2024 predicted transcription factor binding sites provided by the UCSC Genome Browser). The nearest transcription start sites are for *ROBO1*, located 779,554 base pairs upstream the enhancer, and *GBE1*, located 1,206,037 base pairs downstream the enhancer.

**Table S71: Location of Denisovan-derived single nucleotide changes relative to genes.** Gene annotations are based on GENCODE version 43.

| **Gene category** | **Annotation** | **Number of sites** (out of 237,112) | **Number of Genes** (out of 20,170) |
| --- | --- | --- | --- |
| Protein-coding | 5’ Untranslated region | 1,111 (0.5%) | 1,004 (5%) |
|  | 3’ Untranslated region | 3,659 (1.5%) | 2,730 (13.5%) |
|  | Coding sequence | 2,592 (1.1%) | 2,136 (10.6%) |
|  | Missense change | 1,307 (0.6%) | 1,142 (5.7%) |
|  | Likely impactful missense | 10 (0.004%) | 10 (0.05%) |
|  | Start codon loss | 4 (0.002%) | 4 (0.02%) |
|  | Stop codon loss | 1 (0.0004%) | 1 (0.005%) |
|  | Stop codon gain | 11 (0.005%) | 10 (0.05%) |
|  | Intronic | 101,238 (42.7%) | 11,305 (56.1%) |
|  | Top 1% constrained regions | 1,994 (0.8%) | n/a |
|  | Intergenic | 128,693 (54.3%) | n/a |
|  | Top 1% constrained regions | 881 (0.4%) | n/a |
| Non-coding RNA | Long non-coding RNA & long intergenic non-coding RNA | 44,679 (18.8%) | 9,860 (52.3% of 18,852 genes) |
|  | microRNA | 14 | 14 (0.5% of 3049 genes) |

**Table S72: Denisovan-derived missense changes that are likely impactful.** Missense changes with a PrimateAI-3D pathogenicity score between 0.8 and 1. Alleles with no call in one of the Denisovan genomes are denoted with “NA”.

| **Chr.** | **Position** | **Ancestral allele** | ***Denisova 3* allele** | ***Denisova 25* allele** | **Ancestral amino** | **Derived amino** | **PrimateAI-3D**  **score** | **Gene** | **Gene name** |
| --- | --- | --- | --- | --- | --- | --- | --- | --- | --- |
| X | 85404043 | G | NA | A | G | E | 0.907 | *DACH2* | Dachshund family transcription factor 2 |
| 1 | 197564430 | A | G | NA | S | P | 0.887 | *DENND1B* | DENN domain containing 1B |
| 6 | 167705033 | T | C | C | F | S | 0.884 | *UNC93A* | Unc-93 homolog A |
| 4 | 676668 | C | T | T | G | E | 0.841 | *SLC49A3* | Solute carrier family 49 member 3 |
| 10 | 96025626 | C | NA | T | L | F | 0.837 | *PLCE1* | Phospholipase C epsilon 1 |
| 4 | 494258 | G | T | T | L | F | 0.835 | *PIGG* | Phosphatidylinositol glycan anchor biosynthesis class G |
| 11 | 14501220 | G | C | NA | A | G | 0.824 | *COPB1* | COPI coat complex subunit beta 1 |
| 1 | 200573017 | C | A | A | A | S | 0.820 | *KIF14* | Kinesin family member 14 |
| 13 | 42306256 | G | A | A | S | L | 0.819 | *VWA8* | Von Willebrand factor A domain containing 8 |
| 10 | 25701374 | T | C | C | L | P | 0.801 | *GPR158* | G protein-coupled receptor 158 |

**Table S73: Denisovan-derived single nucleotide changes that alter a start or stop codon.** Alleles with no call in one of the Denisovan genomes are denoted with “NA”.

| **Type of change** | **Chr.** | **Pos.** | **Ancestral allele** | **Denisova 3 allele** | **Denisova 25 allele** | **Gene** | **Gene name** | **Corresponding change** | **Canonical transcript** |
| --- | --- | --- | --- | --- | --- | --- | --- | --- | --- |
| stop codon gain | 2 | 43903301 | A | C | C | *C1GALT1C1L* | C1GALT1 specific chaperone 1 like | L54* | Yes |
|  | 10 | 33103363 | C | T | T | *CCDC7* | Coiled-coil domain containing 7 | R1055* | Yes |
|  | 3 | 115382704 | C | T | T | *GAP43* | Growth associated protein 43 | Q27* | No |
|  | 1 | 1691211 | G | A | A | *NADK* | NAD kinase | R132* | No |
|  | 7 | 94992090 | C | T | T | *PON3* | Paraoxonase 3 | W253* | Yes |
|  | 1 | 2520477 | C | T | T | *PRXL2B* | Peroxiredoxin like 2B | Q193* | Yes |
|  | 9 | 130476495 | G | T | NA | *PTRH1* | Peptidyl-tRNA hydrolase 1 homolog | C141* | No |
|  | 21 | 35772152 | G | A | NA | *SMIM11* | Small integral membrane protein 11 | W52* | No |
|  | 3 | 23365050 | G | A | NA | *UBE2E2* | Ubiquitin conjugating enzyme E2 E2 | W91* | No |
|  | 7 | 100363121 | G | T | NA | *ZAN* | Zonadhesin | E1472* | Yes |
|  | 7 | 100369478 | C | T | NA | *ZAN* | Zonadhesin | Q1754* | Yes |
| stop codon loss | 14 | 56147406 | T | A | A | *KTN1* | Kinectin 1 | *1307R | No |
| start codon loss | 3 | 112935060 | T | A | A | *BOC* | BOC cell adhesion associated, oncogene regulated | M1K | No |
|  | 22 | 19481869 | T | C | C | *CDC45* | Cell division cycle 45 | M1T | No |
|  | 3 | 138291769 | T | C | C | *CEP70* | Centrosomal Protein 70 | M1V | Yes |
|  | X | 153046740 | G | A | NA | *SRPK3* | SRSF Protein Kinase 3 | M1I | No |

**Table S74: Functional and evolutionary annotations of the Denisovan-derived single nucleotide changes.**

| **Annotation category** | **Annotation** | **Number of sites**  (out of 237,112) |
| --- | --- | --- |
| Transcription-related | Expression Quantitative Trait Loci (eQTL) | 658 (0.3%) |
|  | Denisovan allele matches eQTL allele | 366 (0.2%) |
|  | Splicing Quantitative Trait Loci (sQTL) | 321 (0.1%) |
|  | Denisovan allele matches sQTL allele | 149 (0.06%) |
|  | Open chromatin regions | 95,113 (40.1%) |
|  | Roadmap Epigenomics Consortium | 65,464 (27.6%) |
|  | ENCODE phase 4 | 70,890 (29.9%) |
|  | Promoter-proximal regions | 4,272 (1.8%)  in 3,510 genes |
|  | Canonical transcripts only (MANE-Select) | 2,125 (0.9%)  in 1,938 genes |
|  | Enhancer region | 80,831 (34.1%) |
|  | Defined by histone modifications | 80,680 (34%) |
|  | Reporter assay activity | 169 (0.07%) |
|  | Transcribed enhancer | 782 (0.3%) |
|  | Predicted to alter a transcription factor binding site | 19,625 (8.3%) |
|  | In conserved element (phastCons ≥ 0.95) | 2,474 (1%) |
|  | Conserved base (phyloP ≥ 1.3) in conserved element | 917 (0.4%) |
| Evolution-related | Deserts of Neandertal and Denisovan introgression | 3,898 (1.6%) |
|  | Potentially introgressed in modern humans | *admixfrog:*  32,691 (13.8%)  *hmmix:*  61,131 (25.8%) |
|  | Ultra-conserved regions | 4 (0.002%) |
| Trait and Disease-related | Genome-wide association study (GWAS) | 41 (74 associations) |
|  | Denisovan allele matches reported risk allele | 11 (16 associations) |

**Table S75: Single nucleotide changes in ultra-conserved regions of the human genome.** Regions of the human genome with identical sequence in the mouse and rat genomes (at least 200 base pairs) or 235 mammalian genomes (at least 20 base pairs). Reported are the chromosome and position of the single nucleotide change, the ancestral and derived allele, the length of the region with identical sequence in base pairs (bp), the name of the gene with the nearest transcription start site (TSS) upstream or downstream the identical region, and the distance to the TSS.

| **Chr.** | **Position** | **Ancestral allele** | **Denisovan allele** | **Identical region length (bp)** | **Genomes with identical sequence** | **Nearest upstream TSS**  **Gene and distance** | **Nearest downstream TSS**  **Gene and distance** |
| --- | --- | --- | --- | --- | --- | --- | --- |
| 2 | 66092122 | A | T | 28 | 235 mammals | SPRED2 (432,389 bp) | MEIS1 (570,135 bp) |
| 3 | 70872181 | G | A | 219 | Mouse, rat | MDFIC2 (510,392 bp) | FOXP1 (760,698 bp) |
| 7 | 114054475 | C | A | 22 | 235 mammals | PPP1R3A (495,411 bp) | FOXP2 (395 bp) |
| 10 | 77139056 | G | A | 252 | Mouse, rat | COMTD1 (143,341 bp) | ZNF503 (22,466 bp) |

**Table S76: List of 74 traits associated with 41 Denisovan-derived single nucleotide change positions.** Rows highlighted in green represent associations where the reported risk allele matches the Denisovan-derived allele. Rows highlighted in grey represent associations with no reported risk allele. Entries with “NR” correspond to values not reported in the GWAS catalogue.

| **Chr** | **Pos.** | **Ancestral allele** | **Derived allele** | **Trait** | **Reported risk allele** | **P-value** | **OR or Beta** | **95% CI** | **Unit** | **rsID** |
| --- | --- | --- | --- | --- | --- | --- | --- | --- | --- | --- |
| 1 | 2798712 | C | A | Cortical thickness | T | 1.00E-17 | NR | . | . | rs4373707 |
| 1 | 2798712 | C | A | T2 brain MRIs Unsupervised Deep learning derived Imaging Phenotypes (dimension 77) | NR | 8.00E-19 | NR | . | . | rs4373707 |
| 1 | 2798712 | C | A | Vertex-wise cortical surface area | T | 2.00E-37 | 12.78 | NR | z_score increase | rs4373707 |
| 1 | 2798712 | C | A | Vertex-wise cortical thickness | T | 2.00E-22 | 9.76 | NR | z_score increase | rs4373707 |
| 1 | 2798712 | C | A | Whole brain free water diffusion (multivariate analysis) | NR | 1.00E-21 | NR | . | . | rs4373707 |
| 1 | 2798712 | C | A | Whole brain restricted isotropic diffusion (multivariate analysis) | NR | 6.00E-29 | NR | . | . | rs4373707 |
| 1 | 11851927 | C | T | Diastolic blood pressure | T | 9.00E-08 | 0.2752 | [0.17-0.38] | unit increase | rs41394446 |
| 1 | 11851927 | C | T | Pulse pressure | T | 1.00E-07 | 0.3266 | [0.21-0.45] | unit increase | rs41394446 |
| 1 | 11851927 | C | T | Systolic blood pressure | T | 1.00E-11 | 0.5979 | [0.43-0.77] | unit increase | rs41394446 |
| 1 | 21946831 | C | T | Serum alkaline phosphatase levels | T | 3.00E-40 | 0.1203 | [0.1-0.14] | unit increase | rs116068751 |
| 1 | 68907527 | C | G | Facial morphology (right endocanthion-subnasale distance) | C | 3.00E-08 | 0.078 | NR | unit decrease | rs77142479 |
| 1 | 68907527 | C | G | Facial morphology (left endocanthion-subnasale distance) | C | 8.00E-09 | 0.081 | NR | unit decrease | rs77142479 |
| 1 | 119584779 | C | T | Waist circumference adjusted for body mass index | NR | 8.00E-09 | NR | NR | . | rs114473983 |
| 1 | 154979973 | C | T | Height | NR | 7.00E-19 | 0.0500366 | [0.039-0.061] | unit increase | rs150107452 |
| 1 | 154988533 | G | A | A body shape index (BMI-adjusted waist circumference) | C | 4.00E-10 | 0.0446148 | [0.031-0.059] | unit decrease | rs77548871 |
| 1 | 154988533 | G | A | Appendicular lean mass | C | 3.00E-09 | 0.143933 | [0.097-0.191] | unit increase | rs77548871 |
| 1 | 154988533 | G | A | Breast cancer, ovarian cancer or prostate cancer (pleiotropy) | C | 5.00E-09 | 0.0991 | [0.066-0.132] | unit decrease | rs77548871 |
| 1 | 168776296 | T | C | Calcium/calmodulin-dependent protein kinase type 1 levels | G | 1.00E-14 | 0.31698 | [0.24-0.4] | unit decrease | rs78240895 |
| 1 | 201598915 | A | C | Optic disc size | A | 2.00E-10 | 0.02 | NR | . | rs11584075 |
| 1 | 235097741 | G | C | Monocyte count | C | 9.00E-12 | 0.02459202 | [0.018-0.032] | unit decrease | rs115441532 |
| 1 | 235097741 | G | C | Monocyte percentage of white cells | C | 2.00E-12 | 0.02536696 | [0.018-0.032] | unit decrease | rs115441532 |
| 10 | 64300331 | A | C | Breast cancer | A | 2.00E-08 | 1.11 | [1.07-1.15] | . | rs78053936 |
| 11 | 2788916 | T | C | Height | C | 6.00E-15 | 0.0228 | [0.017-0.028] | unit increase | rs2237881 |
| 11 | 30997768 | C | T | Bone mineral density mean | NR | 4.00E-16 | NR | . | . | rs75289607 |
| 11 | 31904941 | G | A | Otosclerosis | T | 1.00E-06 | 0.79 | [0.72-0.87] | . | rs71489821 |
| 11 | 113818507 | A | T | Gut microbiota (bacterial taxa, rank normal transformation method) | T | 2.00E-06 | 0.528377 | [0.31-0.75] | unit decrease | rs45603544 |
| 13 | 44643465 | G | T | Response to paliperidone in schizophrenia (PANSS score) | G | 3.00E-08 | 20.6563 | [NR] | unit decrease | rs181534711 |
| 14 | 101192299 | A | G | Height | A | 7.00E-10 | 0.071 | [0.049-0.094] | unit increase | rs3759556 |
| 14 | 101192299 | A | G | Height | G | 9.00E-22 | 0.046 | [0.037-0.055] | unit decrease | rs3759556 |
| 15 | 55425345 | G | A | Cognitive function (immediate memory) | C | 3.00E-06 | 0.727043 | [0.42-1.03] | unit decrease | rs75799599 |
| 16 | 53338253 | T | G | Triamcinolone acetonide-induced ocular hypertension | NR | 7.00E-13 | NR | . | . | rs371849030 |
| 19 | 10216835 | G | T | Monocyte forward scatter | T | 3.00E-09 | 0.0666624 | [0.045-0.089] | SD units increase | rs73009299 |
| 2 | 46429432 | G | T | Protein quantitative trait loci (liver) | T | 1.00E-08 | 0.4322 | NR | unit increase | rs115180365 |
| 2 | 46584859 | A | G | Hemoglobin levels | G | 8.00E-08 | 0.396 | NR | g/dL decrease | rs372272284 |
| 2 | 46584859 | A | G | Oxygenated hemoglobin levels | G | 6.00E-09 | 0.386 | NR | g/dL decrease | rs372272284 |
| 2 | 72703941 | G | A | Cerebrospinal fluid sTREM-2 levels | NR | 5.00E-06 | 10514.731 | NR | unit decrease | rs115600982 |
| 2 | 124991887 | A | G | Educational attainment (MTAG) | A | 2.00E-18 | 0.0126 | [0.0099-0.0153] | unit decrease | rs4848924 |
| 2 | 124991887 | A | G | Educational attainment (years of education) | A | 4.00E-14 | 0.0117 | [0.0088-0.0146] | unit decrease | rs4848924 |
| 2 | 124991887 | A | G | Highest math class taken (MTAG) | A | 3.00E-21 | 0.0172 | [0.014-0.021] | unit decrease | rs4848924 |
| 2 | 124991887 | A | G | Highest math class taken | A | 1.00E-14 | 0.0192 | [0.014-0.024] | unit decrease | rs4848924 |
| 2 | 218296732 | C | A | Breast cancer | T | 8.00E-07 | 1.0633773 | NR | . | rs16857611 |
| 2 | 218296732 | C | A | Thyroid cancer | C | 2.00E-08 | 0.349 | [0.23-0.47] | unit increase | rs16857611 |
| 2 | 240349095 | A | T | Neuroticism conditioned on cognitive performance (multi-trait conditioning and joint analysis) | NR | 2.00E-09 | NR | . | . | rs4241231 |
| 2 | 240349095 | A | T | Neuroticism conditioned on educational attainment (multi-trait conditioning and joint analysis) | NR | 2.00E-08 | NR | . | . | rs4241231 |
| 2 | 240349095 | A | T | Neuroticism conditioned on highest math class (multi-trait conditioning and joint analysis) | NR | 3.00E-09 | NR | . | . | rs4241231 |
| 2 | 240349095 | A | T | Neuroticism conditioned on self-rated math ability (multi-trait conditioning and joint analysis) | NR | 6.00E-09 | NR | . | . | rs4241231 |
| 2 | 240349095 | A | T | Neuroticism conditioned on Townsend deprivation index (multi-trait conditioning and joint analysis) | NR | 2.00E-09 | NR | . | . | rs4241231 |
| 2 | 240349095 | A | T | Neuroticism | A | 2.00E-06 | 0.0165574 | [0.0097-0.0234] | unit increase | rs4241231 |
| 22 | 48923459 | C | G | Bipolar disorder (body mass index interaction) | NR | 5.00E-06 | NR | . | . | rs80088139 |
| 3 | 48477252 | G | A | Macrophage inflammatory protein 1b levels | G | 5.00E-22 | 0.2944 | [0.23-0.35] | SD units decrease | rs146948889 |
| 3 | 48477252 | G | A | Total cholesterol levels | A | 3.00E-11 | 0.044164 | [0.033-0.055] | unit decrease | rs146948889 |
| 3 | 141094769 | A | T | Blond vs. brown/black hair color | A | 5.00E-58 | 1.131 | NR | . | rs4683605 |
| 3 | 141094769 | A | T | Whole body fat free mass (UKB data field 23101) | A | 7.00E-98 | 0.0278264 | [0.025-0.03] | unit increase | rs4683605 |
| 3 | 141094769 | A | T | Whole body fat mass (UKB data field 23100) | A | 5.00E-21 | 0.0223663 | [0.018-0.027] | unit increase | rs4683605 |
| 3 | 184141482 | G | A | Memory decline x sex interaction | NR | 7.00E-06 | 0.011421 | NR | unit decrease | rs6772679 |
| 4 | 106196951 | A | T | Chronic obstructive pulmonary disease | G | 2.00E-10 | 1.07 | [1.05-1.09] | . | rs2454206 |
| 4 | 106196951 | A | T | Height | A | 6.00E-09 | 0.027 | [0.013-0.041] | unit increase | rs2454206 |
| 4 | 106196951 | A | T | Pulse pressure | A | 2.00E-08 | 0.1068706 | [0.069-0.145] | mmHg decrease | rs2454206 |
| 4 | 106196951 | A | T | Pulse pressure | A | 5.00E-08 | 0.08477454 | [0.053-0.116] | mmHg decrease | rs2454206 |
| 5 | 30716047 | G | A | Bone mineral density mean | NR | 1.00E-08 | NR | . | . | rs115206369 |
| 5 | 109058594 | A | C | Intelligence | A | 4.00E-08 | 5.513 | NR | z-score increase | rs1438660 |
| 5 | 131952398 | G | T | Asthma (childhood onset) | NR | 3.00E-27 | NR | . | . | rs6894017 |
| 5 | 172191052 | T | C | C-reactive protein levels | C | 6.00E-09 | 0.056 | [0.036-0.076] | unit increase | rs17658229 |
| 5 | 172191052 | T | C | Vertical cup-disc ratio | C | 8.00E-09 | 0.02 | [0.012-0.028] | unit decrease | rs17658229 |
| 6 | 30001703 | C | T | Frontal fibrosing alopecia | NR | 5.00E-32 | 2.76 | [2.33-3.27] | . | rs62388754 |
| 6 | 43088094 | A | C | Posterior urethral valves | G | 2.00E-08 | 7.2 | [4.08-12.7] | . | rs144171242 |
| 6 | 126835655 | C | G | Height | T | 6.00E-07 | 4.8 | [2.84-6.76] | % s.d. increase | rs1490388 |
| 6 | 146583120 | G | A | Smoking x experiences in childhood (felt loved by family member) interaction | NR | 2.00E-08 | 0.9806 | [0.64-1.32] | unit decrease | rs73009056 |
| 6 | 160525287 | C | T | Apolipoprotein B levels | C | 1.00E-14 | 0.0205 | [0.015-0.026] | unit decrease | rs2297372 |
| 6 | 160525287 | C | T | Creatinine levels | C | 1.00E-09 | 0.016 | [0.011-0.021] | unit increase | rs2297372 |
| 6 | 160525287 | C | T | Estimated glomerular filtration rate | C | 7.00E-10 | 0.0163 | [0.011-0.021] | unit decrease | rs2297372 |
| 6 | 160525287 | C | T | Low density lipoprotein cholesterol levels | C | 3.00E-14 | 0.0201 | [0.015-0.025] | unit decrease | rs2297372 |
| 6 | 160525287 | C | T | Total cholesterol levels | C | 7.00E-12 | 0.0181 | [0.013-0.023] | unit decrease | rs2297372 |
| 7 | 1883486 | T | A | Smoking initiation (MTAG) | NR | 1.00E-09 | 0.00987474 | [0.0067-0.0131] | unit increase | rs10807752 |

**Figure S125: Single nucleotide changes correlated with expression differences of protein-coding genes.** Scatter plot depicts expression quantitative loci (eQTL) associated with the Denisovan-derived allele carried by both genomes. The color and shape of the point denotes the tissue or cell type where the eQTL is observed. The violin plots, obtained from the GTEx eQTL dashboard, depict normalized expression differences among genotypes for the top twelve most significant eQTL. Listed above each plot is the eQTL GTEx ID (hg38 coordinates) with the red allele corresponding to the Denisovan-derived allele, with the gene name and tissue.

**Figure S126: Single nucleotide changes in promoter-proximal regions ranked by conservation scores from 447 mammalian genomes.** The scatter plot depicts the length of promoter-proximal regions in mappable regions passing filters (described in **Supplementary Note 3**) and the nucleotide-level conservation phyloP score calculated from the 447-way mammalian genome alignment. The gene name appears next to SNC with a phyloP score greater than 5 (p-value = 1E-5).

**Table S77: Gene ontology enrichment results for genes with single nucleotide changes in promoter-proximal regions.** Genes were ranked using the conservation score of the position where single nucleotide changes occurred. Conservation scores are derived from a 447-way alignment of mammalian genomes^176^. Higher scores correspond to stronger levels of conservation and higher likelihood the SNC has a functional effect. A Wilcoxon rank-sum test determined enriched ontology terms. The top 20 terms are shown and sorted by the family-wise error rate.

| **Ontology category** | **Ontology ID** | **Term name** | **Raw p-value (high rank)** | **FWER (high rank)** |
| --- | --- | --- | --- | --- |
| Biological process | GO:0007417 | Central nervous system development | 1.859e-06 | < 0.001 |
| Biological process | GO:0007399 | Nervous system development | 1.954e-06 | < 0.001 |
| Biological process | GO:0007420 | Brain development | 2.541e-05 | 0.005 |
| Biological process | GO:0060322 | Head development | 9.674e-05 | 0.036 |
| Cellular component | GO:0032589 | Neuron projection membrane | 0.00049772 | 0.053 |
| Biological process | GO:0048731 | Stem development | 0.00014296 | 0.071 |
| Biological process | GO:0009887 | Animal organ morphogenesis | 0.00015061 | 0.078 |
| Biological process | GO:0007275 | Multicellular organism development | 0.00016473 | 0.085 |
| Biological process | GO:0072089 | System cell proliferation | 0.00026353 | 0.138 |
| Biological process | GO:0009719 | Response to endogenous stimulus | 0.00034061 | 0.188 |
| Biological process | GO:0022008 | Neurogenesis | 0.00055226 | 0.289 |
| Molecular function | GO:0071837 | HMG box domain binding | 0.00191005 | 0.294 |
| Biological process | GO:0035107 | Appendage morphogenesis | 0.00058795 | 0.31 |
| Biological process | GO:0035108 | Limb morphogenesis | 0.00058795 | 0.31 |
| Biological process | GO:2000648 | Positive regulation of stem cell proliferation | 0.00067015 | 0.344 |
| Biological process | GO:0072091 | Regulation of stem cell proliferation | 0.00069834 | 0.354 |
| Biological process | GO:0007389 | Pattern specification process | 0.00073033 | 0.365 |
| Biological process | GO:0048736 | Appendage development | 0.0009695 | 0.466 |
| Biological process | GO:0060173 | Limb development | 0.0009695 | 0.466 |
| Biological process | GO:0035270 | Endocrine system development | 0.0010813 | 0.511 |
